# Novel Serine/Threonine-O-glycosylation with N-Acetylneuraminic acid and 3-Deoxy-D-manno-octulosonic acid by Maf glycosyltransferases

**DOI:** 10.1101/2020.06.03.131540

**Authors:** Aasawari Khairnar, Sonali Sunsunwal, Ponnusamy Babu, T.N.C. Ramya

**Author notes:** Correspondence and requests for materials should be addressed to T.N.C. Ramya, CSIR-Institute of Microbial Technology Sector 39-A, Chandigarh 160036 INDIA. **Supplementary data:** Supplementary figures S1 to S13, supplementary tables S1 to S23, and additional supplementary data containing SEQUEST-annotated MS/MS spectra of O-glycosylated peptides with Modscore values of ≥ 19 for Site 1 Position_A and Site 1 Position_B are provided as supporting information for this manuscript.

## Abstract

Some bacterial flagellins are O-glycosylated on surface-exposed Serine/Threonine residues with nonulosonic acids such as pseudaminic acid, legionaminic acid, and their derivatives by flagellin nonulosonic acid glycosyltransferases, also called Motility associated factors (Maf). We report here two new glycosidic linkages previously unknown in any organism, Serine/Threonine-O-linked N-Acetylneuraminic acid (Ser/Thr-O-Neu5Ac) and Serine/Threonine-O-linked 3-Deoxy-D-manno-octulosonic acid (Ser/Thr-O-KDO), both catalysed by *Geobacillus kaustophilus* Maf (putative flagellin sialyltransferase) and *Clostridium botulinum* Maf (putative flagellin legionaminic acid transferase). We identified these novel glycosidic linkages in recombinant *G. kaustophilus* and *C. botulinum* flagellins that were co-expressed with their cognate recombinant Maf protein in *Escherichia coli* strains producing the appropriate nucleotide sugar glycosyl donor. The glycosylation of *G. kaustophilus* flagellin with KDO, and that of *C. botulinum* flagellin with Neu5Ac and KDO indicates that Maf glycosyltransferases display donor substrate promiscuity. Maf glycosyltransferases have the potential to radically expand the scope of neoglycopeptide synthesis and posttranslational protein engineering.

**Significance Statement:** Glycosylation, the modification of proteins with sugars, is one of the most common post-translational modifications observed in proteins. While glycosylation is versatile, the most common forms of glycosylation are N-glycosylation, where the N atom of Asparagine is modified with a glycan, and O-glycosylation where the O atom of serine or threonine residues is modified with a glycan. Here, we report a novel type of O-glycosylation in the bacterial flagellin proteins of two Gram-positive bacteria, *Geobacillus kaustophilus* and *Clostridium botulinum*. We demonstrate for the first time that the enzyme flagellin Maf glycosyltransferase is capable of transferring the monosaccharides, N-acetylneuraminic acid and 3-Deoxy-D-manno-octulosonic acid, on to serine and threonine residues of these proteins.

## Introduction

Glycosylation is the most common and versatile post translational modification in proteins ^1^. Initially identified in eukaryotes ^2^, glycans are most commonly N-linked to asparagine or O-linked to serine or threonine ^3^ residues. N-linked and O-linked glycosylation have been well documented in prokaryotic proteins, too ^4, 5, 6, 7^. O-glycosylation occurs in bacterial pilins, flagellins, autotransporter adhesion proteins and several other proteins ^7^.

Flagellins from both Gram-negative and Gram-positive bacteria are known to be O-glycosylated ^7, 8^ with a wide variety of monosaccharide or oligosaccharide glycan structures on varying numbers of glycosites, ranging from two to 19 sites ^9, 10, 11, 12, 13, 14, 15, 16, 17^. The glycan structures that are commonly found on flagellar proteins are derivatives of pseudaminic acid (**Figure 1**) (in *C. jejuni*, *Campylobacter coli*, *Helicobacter pylori*, *Aeromonas caviae*) and legionaminic acid (**Figure 1**) (in *C. coli* and *Clostridium botulinum*), which are non-sialic acid nonulosonic acids ^9, 11, 12, 18^. Nonulosonic acids are nine-carbon sugars with a carboxylate group at C-1 and an exocyclic side chain C-7 to C-9; sialic acids are nonulosonic acids with a glycerol-like exocyclic side chain, eg. N-acetylneuraminic acid (Neu5Ac) ^19^ (**Figure 1**). Also found on flagellar proteins are rhamnose or rhamnose-linked oligosaccharides (in *Pseudomonas aeruginosa*), and N-acetylglucosamine (GlcNAc) (in *L. monocytogenes*) ^10, 13, 14^. Glycans on some flagellins are acylated with acyl groups or peptides, as in the case of *C. jejuni* and *Clostridium difficile* ^15, 16^.

**Figure 1:**
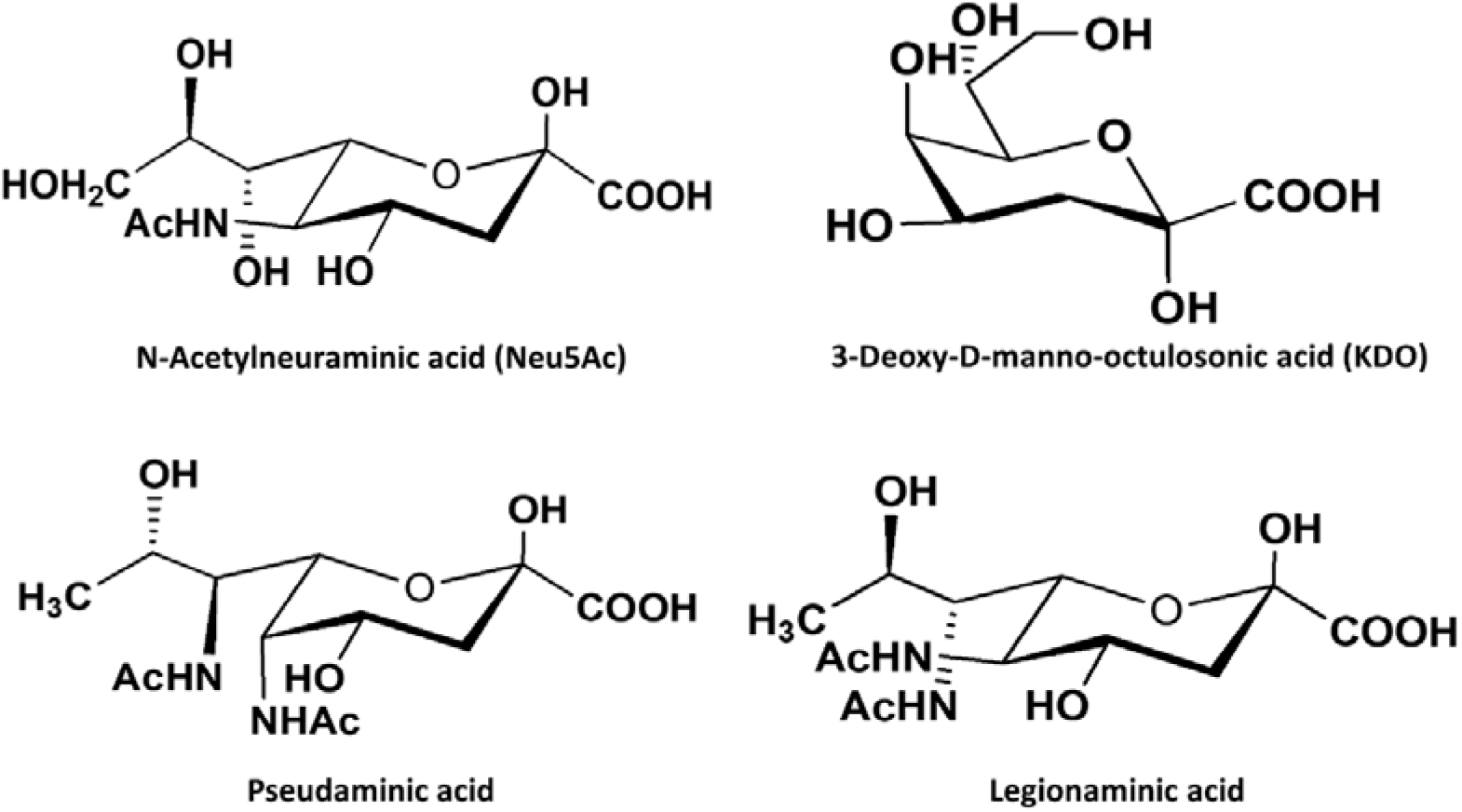
Structures of N-acetylneuraminic acid, 3-Deoxy-D-manno-octulosonic acid, pseudaminic acid and legionaminic acid.

Several lines of evidence indicate that Motility associated factor (Maf) proteins are the flagellin nonulosonic acid glycosyltransferases. The *maf* genes are present in the flagellar locus along with genes responsible for glycan biosynthesis and transport ^20, 21, 22^. Insertional mutagenesis or inactivation of *maf* does not affect biosynthesis of the glycan moiety but results in a non-flagellated, non-motile phenotype in *C. jejuni* ^21, 22, 23^*, H. pylori* ^12^, *Aeromonas hydrophila* ^24, 25, 26^, *A. caviae* ^27^, and *Magnetospirillum magneticum* ^28^. Maf directly interacts with flagellin in vivo in *A. caviae* ^29^. Further, the crystal structure of *M. magneticum* Maf, putative flagellin pseudaminic acid transferase (PDB ID: 5mu5) indicates that the central Maf_flag10 domain (PF01973 or DUF115 in Pfam database) has a modified glycosyltransferase GT-A topology and is structurally related to family GT42 sialyltransferases with striking structural resemblance to *C. jejuni* sialyltransferase Cst-II (PDB ID: 1ro7) ^30^ in the donor substrate binding region ^28^.

In this article, we explicitly demonstrate the flagellin nonulosonic acid glycosyltransferase activity of Maf proteins from Gram-positive bacteria by employing suitable heterologous expression systems. We describe the mass spectrometry analysis of recombinant *Geobacillus kaustophilus* HTA426 flagellins, *Gk*FlaA1 and *Gk*FlaA2, and *C. botulinum* F Str. Langeland flagellin *Cb*Fla heterologously expressed in *E. coli* strains with or without co-expression of the cognate recombinant Maf proteins, *Gk*Maf and *Cb*Maf, respectively. We report two new glycosidic linkages, Serine/Threonine-O-linked N-Acetylneuraminic acid (Neu5Ac) and Serine/Threonine-O-linked 3-Deoxy-D-manno-octulosonic acid (or keto-deoxyoctulosonate; KDO) catalysed by these Maf proteins in their cognate flagellins, and concomitantly demonstrate the donor substrate promiscuity of this class of enzymes.

## Results

### Strategy for heterologous co-expression of Maf and flagellin proteins from *Geobacillus sp*

A recent comparative genomics study of 36 strains of the Gram-positive, aerobic, thermophilic genus *Geobacillus* by Maayer et al demonstrated the occurrence of five distinct types of flagellin glycosylation islands (FGI-I to FGI-V) amongst 18 of these strains ^31^. These islands are present within the flagellar biosynthesis locus and contain glycan biosynthesis and *maf* or glycosyltransferase genes ^31^. Based on the different glycan biosynthesis genes present in the flagellin glycosylation islands (FGI-I to FGI-V), predictions of the glycan moieties on the *Geobacillus* flagellins are possible ^31^. The type-I FGI, present in the type strain, *G. kaustophilus* DSM 7263 (or ATCC 8005 or NBRC 102445, isolated from pasteurized milk in USA ^32, 33, 34^), and strains, *G. kaustophilus* HTA426 (isolated from deep sea sediment in the Mariana trench ^35^) (**Figure 2A**) and *G. thermoglucosidans* C56-YS93 (isolated from a hot spring in Obsidian, USA ^36^), is unique in that it encodes Neu5Ac biosynthetic genes as well as *maf* genes (a single *maf-1* gene in the first and second strains, and two genes, *maf-3* and *maf-4*, in the third strain) ^31^; this suggests that the Maf proteins encoded by them might be flagellin sialyltransferases. In contrast, FGI-II, -III and -IV harbour *maf* genes as well as genes for pseudaminic acid biosynthesis, suggesting that these *maf* genes might encode for flagellin pseudaminic acid transferases, and the type-V FGI has genes for rhamnose biosynthesis but lacks *maf* genes ^31^. Flagellin glycosylation has not been reported in the literature for *Geobacillus kaustophilus* or *G. thermoglucosidans*. The only known glycosylated flagellin from a *Geobacillus* sp. is that of *Geobacillus stearothermophilus* wherein the flagellin was reported to be glycosylated based on positive staining with the Periodic acid-Schiff reagent but without definitive mass spectrometry based characterization ^37^.

**Figure 2:**
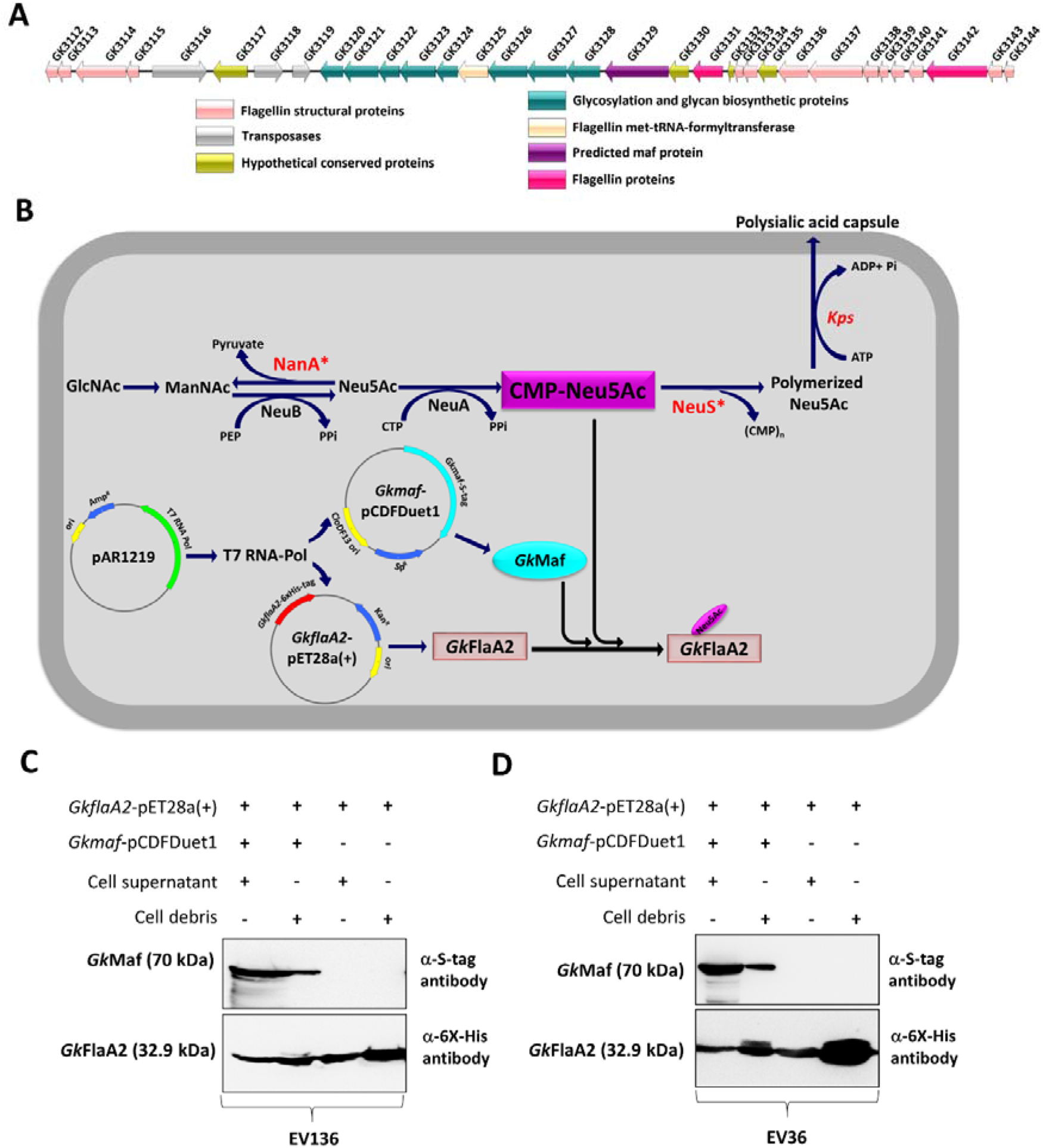
Co-expression of *Gk* FlaA2 with *Gk*Maf. **(A)** Flagellin glycosylation island of *Geobacillus kaustophilus* HTA426. The putative ORFs of Maf and flagellin proteins have been shown in purple and pink colour, respectively. **(B)** Strategy for T7 based co-expression of *GkFlaA2* and *Gk*Maf in recombinant *E. coli* K12:K1 hybrid system overproducing the donor substrate CMP-Neu5Ac and harboring pAR1219 plasmid expressing T7 RNA Polymerase. **(C, D)** Western analysis of cell-free supernatant and cell debris fractions of cell lysates of EV136 **(C)** and EV36 **(D)** cells transformed with *GkflaA2*-pET28a(+) vector and *Gkmaf*-pCDF-Duet-1. Recombinant *Gk*FlaA2 and *Gk*Maf were detected by western analysis with mouse anti-6XHis antibody and mouse anti-S tag antibody, respectively.

Albeit de novo synthesis and/or metabolism of sialic acids (such as Neu5Ac) occur in pathogenic bacteria ^19, 38, 39, 40^ such as *Leptospira* ^41^, *C. jejuni* ^42^, and *E. coli* ^43^, there is no report of any sialic acid being incorporated in flagellins. There is also no evidence, to our knowledge, of protein O-sialylation in any organism to date. Considering the novelty of protein O-linked sialic acids, we decided to study a putative flagellin sialyltransferase protein encoded by *Geobacillus* FGI type-I. We selected the putative flagellin sialyltransferase protein encoded by *maf1* (NCBI accession no. BAD77414.1; protein referred to here as *Gk*Maf) of *G. kaustophilus* HTA426 (which is 100% identical in amino acid sequence to maf1 of *G. kaustophilus* DSM 7263, NCBI accession no. WP_011232599.1), together with its putative acceptor substrates - the two flagellins, encoded by genes *flaA1* (NCBI accession no. BAD77427.1; protein referred to here as *Gk*FlaA1, which does not find a homologue in *G. kaustophilus* DSM 7263), and *flaA2* (NCBI accession no. BAD77416.1, protein referred to here as *Gk*FlaA2, which is 100% identical in amino acid sequence to flagellin of G. kaustophilus DSM 7263, NCBI accession no. WP_011232601.1), for this study considering the presence of a single *maf* gene in this organism as opposed to two *maf* genes in *G. thermoglucosidans* C56-YS93 (*maf-3* and *maf-4*, which are similar to the *maf* genes observed in FGI-II and FGI-III clusters containing pseudaminic acid biosynthesis genes). *Gk*Maf is a member of the Pfam 1973 family ^44^ with the conserved Maf_flag10 domain characteristic of Maf proteins. The two flagellins, *Gk*flaA1 and *Gk*FlaA2 are ~71% identical but vary in length. *Gk*FlaA1 (604 aa long) resembles *Salmonella typhimurium* flagellin in domain organization ^45^ and it has D2 and D3 domains. *Gk*FlaA2 (297 aa long) resembles *Bacillus subtilis* flagellin ^46^ and lacks D2 and D3 domains.

Our strategy for studying the function of the *G. kaustophilus* Maf with putative sialyltransferase function involved co-expressing *Gk*Maf along with *Gk*FlaA1 or *Gk*FlaA2 in a heterologous host producing appropriate nucleotide sugar donor and analysing the recombinant flagellins for the presence of any glycosylation by mass spectrometry (**Figure 2B**). CMP-sialic acid is the donor substrate of sialyltransferases ^47^. In order to ensure sufficient donor substrate availability for the putative flagellin sialyltransferase in our study, we employed as our model heterologous expression system, *E. coli* K12-K1 hybrid strains that were engineered by Vimr and coworkers ^48, 49^ to accumulate abundant CMP-sialic acid in the cytosol. Strains EV136 (*neuS*-) and EV240, (*nanA- neuS-*) accumulate abundant CMP-sialic acid in the cytosol due to their lacking a functional capsule polysialyltransferase (*neuS*) whereas EV36 is the wild type K12-K1 hybrid strain with an insignificant pool of CMP-sialic acids ^49, 50^. We confirmed by borohydride reduction ^51^ and periodate-resorcinol assays ^52^ that chemically competent cells of EV136 and EV240 cells accumulated significant intracellular CMP-sialic acid and total sialic acid (**data not shown**).

We confirmed co-expression of recombinant *Gk*Maf and *Gk*FlaA2 in EV136, EV36 and EV240 cells by western analysis with anti-S-tag and anti-6XHis-tag antibodies, respectively (**Figures 2C and 2D, Supplementary figure S1**) and enriched recombinant *Gk*FlaA2 from these cells by immobilized metal affinity chromatography (**Supplementary figures S2A-C**).

We also singly expressed and purified recombinant *Gk*FlaA2 (**Figures 2C and 2D, Supplementary figures S1 and S2A-C**).

### *Gk*Maf is a flagellin sialyltransferase

We subjected *Gk*FlaA2 to on-blot periodate oxidation and aniline-catalyzed oxime ligation (PAL) with aminooxy-biotin ^53^ in order to detect sialylation, if any. We found that *Gk*FlaA2 co-expressed with *Gk*Maf in EV136, EV36 and EV240 strains displayed less electrophoretic mobility than singly expressed *Gk*FlaA2 and was detected by HRP-conjugated streptavidin following on-blot PAL (**Figure 3A**), suggesting the presence of a vicinal 1,2-diol moiety (such as in sialic acid) that was biotinylated by PAL. The biotinylation was more intense in EV136 and EV240 cells than in EV36 cells. Singly expressed *Gk*FlaA2 from EV136, EV36 and EV240 strains was not biotinylated (**Figure 3A**). This suggested that *Gk*Maf catalysed the sialylation of *Gk*FlaA2 in EV136, EV36 and EV240 cells.

**Figure 3:**
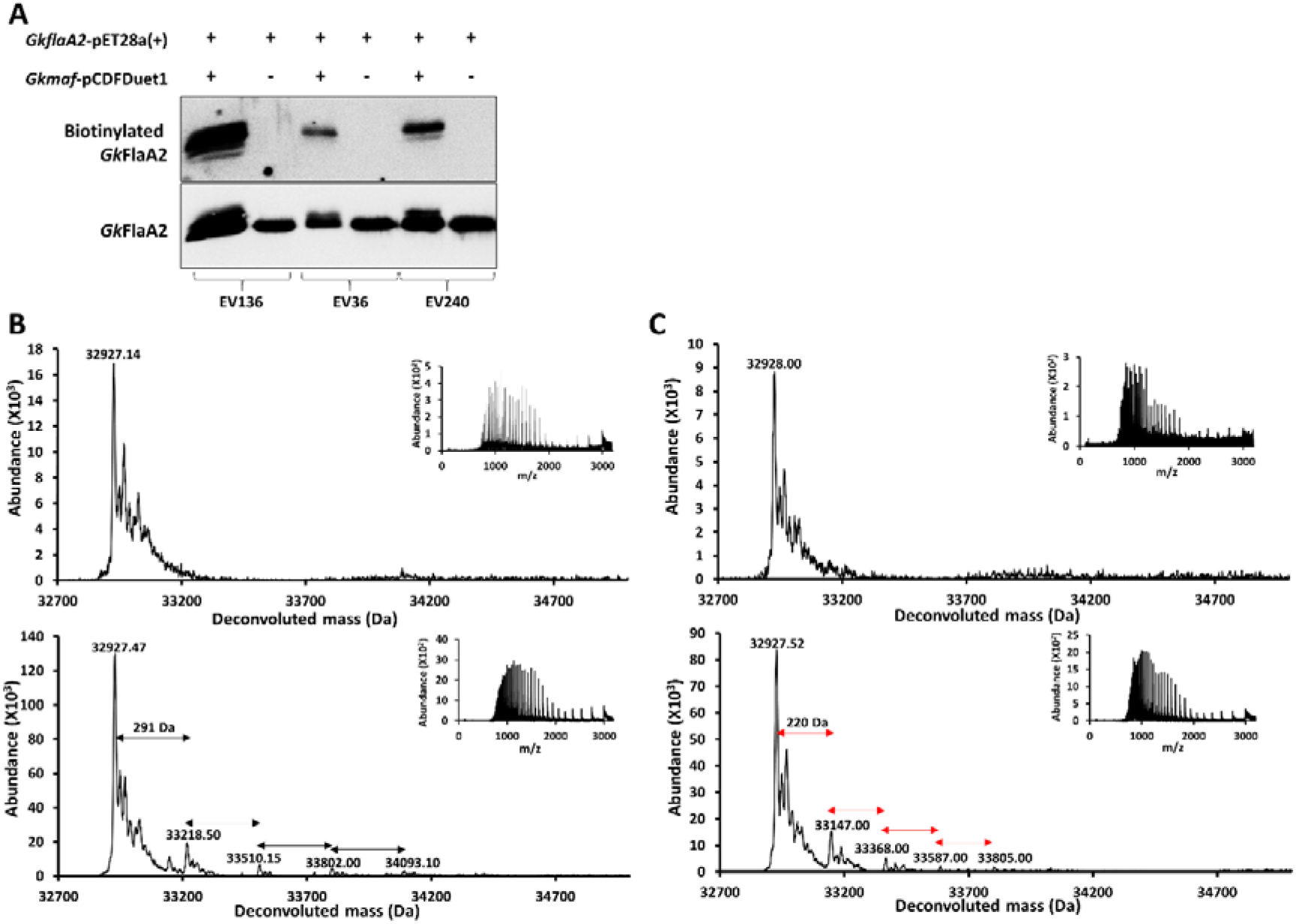
*Gk*Maf glycosylates *Gk*FlaA2. **(A)** On-blot periodate oxidation and biotinylation of *Gk*FlaA2 singly expressed or co-expressed with *Gk*Maf in EV136, EV36 and EV240 cells. Biotinylation was detected with HRP-streptavidin. Total *Gk*FlaA2 protein was detected with mouse anti-6XHis antibody **(B, C)** Intact mass measurement of *Gk*FlaA2 singly expressed (upper panel) or co-expressed with *Gk*Maf and purified from EV136 cells **(B)** or EV36 cells **(C**). The proteins were acetone precipitated and subjected to LC-MS in positive mode ionization. Inset shows the ionization spectrum. The satellite peaks correspond to sodiated adducts.

We also subjected purified *Gk*FlaA2, singly expressed or co-expressed in the presence of *Gk*Maf, from EV36, EV136 and EV240 cells to mass spectrometry for intact mass analysis (**Figure 3B and 3C, and Supplementary Figure S3**). The ionisation spectrum of *Gk*FlaA2 was a multiply charged bell shaped envelope characteristic of intact proteins (**Figures 3B and 3C and Supplementary Figure S3**). Singly expressed *Gk*FlaA2 purified from EV136 cells had a molecular mass of 32.927 kDa (**Figure 3B, upper panel**). The theoretically predicted average molecular mass of recombinant *Gk*FlaA2 is 33.057 kDa, suggesting that the protein undergoes N-ter Met cleavage. In contrast, *Gk*FlaA2 co-expressed with *Gk*Maf in EV136 cells displayed four prominent additional peaks of molecular masses of 33.219 kDa, 33.510 kDa, 33.802 kDa, and 34.093 kDa, each differing by 291 Da from the previous peak, and thus indicative of the presence of up to at least four moieties of 291 Da on *Gk*FlaA2 (**Figure 3B, lower panel**). Similar results were obtained with *Gk*FlaA2 coexpressed with *Gk*Maf in EV240 cells (**Supplementary Figure S3**). The sialic acid, N-acetylneuraminic acid (Neu5Ac), has a molecular mass of 309 Da, and a sialic acid moiety transferred to protein with loss of water (18 Da) would result in the addition of a 291 Da moiety. These results thus indicated that *Gk*Maf is a flagellin sialyltransferase.

### *Gk*Maf displays donor substrate promiscuity

As mentioned above, EV36 cells do not accumulate CMP-sialic acid ^49^ although they actively synthesize the same. *Gk*FlaA2 singly expressed from EV36 cells also had a molecular mass of 32.928 kDa. However, when co-expressed with *Gk*Maf in EV36 cells, *Gk*FlaA2 displayed additional peaks of molecular masses, 33.147 kDa, 33.368 kDa, 33.587 kDa and 33.805 kDa, corresponding to protein modification with up to four moieties of 220 Da. No molecular species corresponding to modification with Neu5Ac (291 Da) was present. The sugar, 3-deoxy-manno-octulosonic acid (or keto-deoxyoctulosonate; KDO) (**Figure 1**) has a molecular mass of 238 Da and a KDO moiety transferred to protein with loss of water (18 Da) would result in the addition of a 220 Da moiety. KDO is known to be present in the lipopolysaccharide lipid-A component of *E. coli* ^54, 55, 56, 57, 58^, and from the apparent Km value (0.29 mM) of CMP-KDO synthetase (from *E. coli* B cells) for KDO ^59^, KDO is likely to be present at sub-millimolar concentration. We conjecture that CMP-3-deoxy-manno-octulosonic acid (CMP-KDO) is present at micromolar concentration in *E. coli* B, *E. coli* K12 and their derivative strains because the Km value of *E. coli* KDO transferase is 88 μM for CMP-KDO ^60^. Considering this, we predicted the 220 Da moiety observed on *Gk*FlaA2 to be KDO. The putative modification of flagellin with KDO, an octulosonic acid by *Gk*Maf is suggestive of the donor substrate promiscuity of *Gk*Maf.

### Glycosylation sites of *Gk*FlaA2

We subjected *Gk*FlaA2, singly expressed or co-expressed along with *Gk*Maf in EV136 and EV36 cells, to protease digestion and liquid-chromatography-tandem mass spectrometry (LC-MS/MS) (performed by Taplin Biomedical Mass Spectrometry Facility, Harvard Medical School) in order to identify sites of O-glycosylation. Parameters used for searching for O-glycosylated peptides are listed (**Supplementary Table S1)**. The FASTA sequence of *Gk*FlaA2 was used as the database for matching spectra to peptides and the reverse sequence was used to ensure specificity of hits. We achieved high sequence coverage (≥ 75%) upon tandem mass spectrometry of tryptic and GluC-digested peptides from *Gk*FlaA2 expressed singly or co-expressed with *Gk*Maf from EV136 as well as EV36 cells (**Figures 4A and 4B, Supplementary Tables S2-S6**). We found ≤5 peptide hits in the searches using the reverse sequence as the database; in contrast, we found >1500 hits to the FASTA sequence of *Gk*FlaA2 in all samples analyzed (**Supplementary Table S6**).

**Figure 4:**
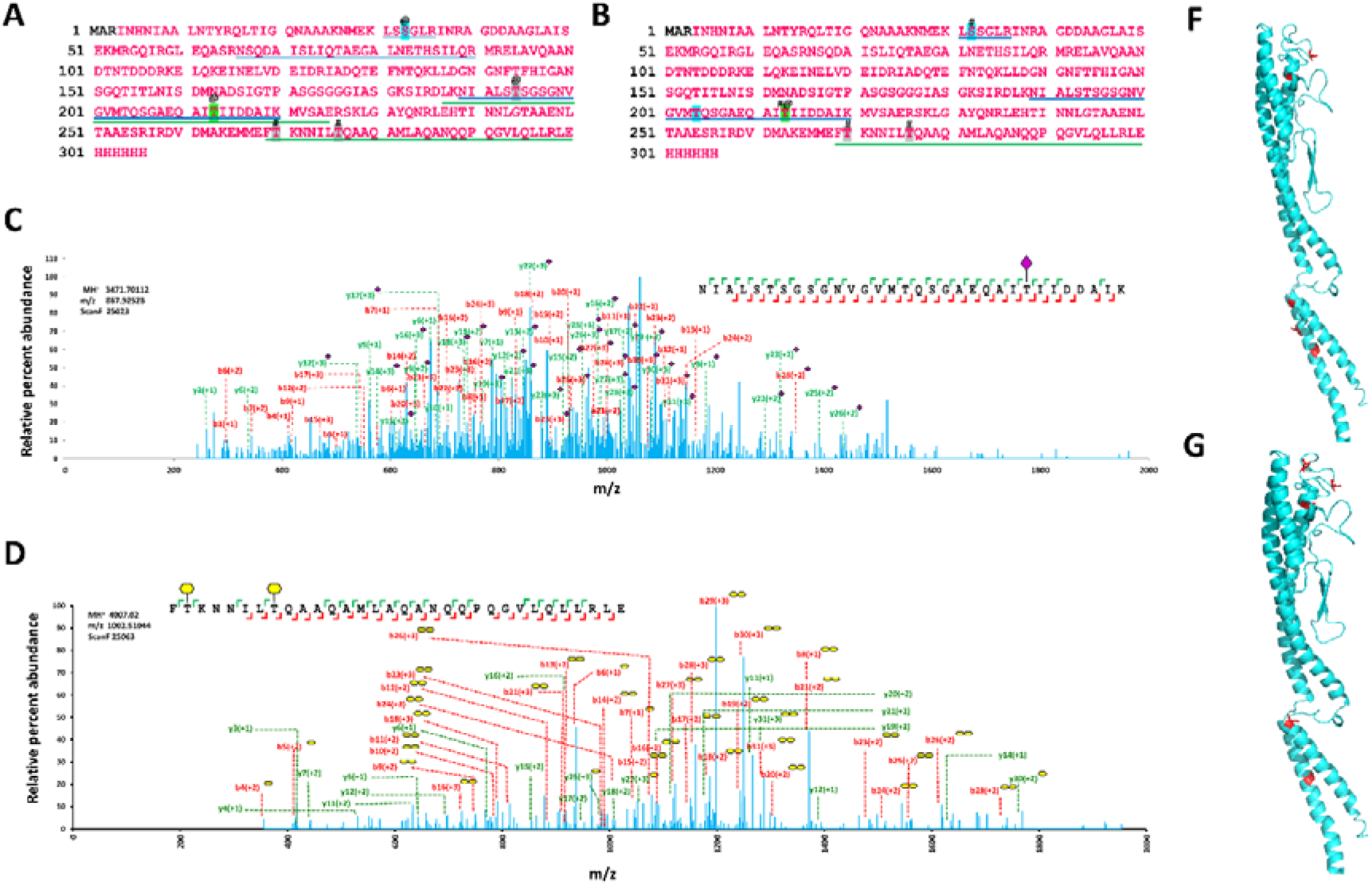
Identification of glycosites in *Gk* FlaA2. **(A,B)** Sequence coverage of tandem mass spectrometry data obtained for *Gk*FlaA2 purified from EV136 **(A)** and EV36 cells **(B)**. The sequence coverage is shown as pink font. The modified sequence coverage is underlined blue (tryptic peptides) or green (Glu-C digested peptides). The sites modified have been highlighted in blue. Neu5Ac is represented by @ symbol and KDO by #. **(C, D)** MS/MS spectra indicating fragment ions of tryptic and Glu-C digested peptides of *Gk*FlaA2 in EV136 **(C**) and EV36 **(D)** cells as per SEQUEST ^97^. y- and b- ions have been shown in green and red colour, respectively. Relative percent abundances (relative to most abundant fragment ion) are plotted on the y-axis. Neu5Ac and KDO are represented by a pink diamond and a yellow hexagon, respectively, as per symbol nomenclature for graphical representations of glycans ^100^. Spectra are of peptides identified and listed in Supplementary tables S7c and S8d. **(F)** and **(G)** show structure model of *Gk*FlaA2, generated by iTASSER ^101^ using 5wjt as template, with Ser/Thr sites modifications (in stick form) as per MS/MS data from EV136 **(F)** and EV36 **(G)** cells.

We employed the Modscore algorithm ^61^ to search for peptides with modifications of 291 Da (Neu5Ac) or 220 Da (KDO) on Ser/Thr residues of *Gk*FlaA2 co-expressed with *Gk*Maf in EV136 and EV36 cells (**Supplementary Tables S7a, S7b, S8a and S8b**). We applied a probabilistic-based approach to assign protein modification sites ^61^, filtering these peptide hits to retain only those with Modscore values of ≥19 (P≤0.01) for Site 1 Position_A and Site 1 Position_B scores in *Gk*FlaA2 co-expressed with *Gk*Maf from EV136 and EV36 cells (**Figures 4A and 4B, Supplementary Tables S7c, S7d, S8c, and S8d, and additional supplementary data**). We expect these glycosylation sites to be confidently assigned. No such peptide hits were obtained in *Gk*FlaA2 singly expressed from EV36 or EV136 cells, confirming that *Gk*Maf was essential for these observed modifications.

Ions indicating confident assignment of the glycosylation sites are evident in the spectra of the modified peptide hits (**Figures 4C and 4D**). For instance, the underlined Thr residue in the peptide, NIALSTSGSGNVGVMTQSGAEQAITIIDDAIK from *Gk*FlaA2 co-expressed with *Gk*Maf in EV136 cells is predicted to be O-sialylated based on the presence of fragment ions retaining a 291 Da moiety (b25 to b28, b30, b31, y8 to y19, y21 to y23, and y25 to y31). Similarly, in the spectrum of the peptide F<u>T</u>KNNIL<u>T</u>QAAQAMLAQANQQPQGVLQLLRLE from *Gk*FlaA2 co-expressed with *Gk*Maf in EV36 with two 220 Da (KDO) modified Thr residues (underlined in the sequence), singly charged ions, b6 and b7, and doubly charged ions, b4 and b5 can be used to assign the KDO moiety on the first (towards N-terminal) Thr residue, b ions, b4 to b7, can be used to assign the KDO moiety on the first Thr residue, y ions, y25, y27 and y30, can be used to assign the KDO moiety on the second Thr residue, and many b ions (b8 to b15, b18 to b21, and b23 to b31) retain the KDO moiety on both residues (**Figure 4C**). (**Figures 4D and 4E**). The spectra of modified peptides with Modscore values 19 ≥ (P ≤ 0.01) for Site 1 Position_A and Site 1 Position_B scores are available as additional **Supplementary data**. The residues identified as O-sialylated are located in and around the loop region of the D1 domain and in the D0 helices (**Figures 4E and 4F**).

### Glycosylation sites of *Gk*FlaA1

As mentioned above, the genome of *G. kaustophilus* HTA426, encodes for two flagellins, *Gk*flaA1 and *Gk*FlaA2. We also studied glycosylation in *Gk*FlaA1, the longer flagellin (theoretically predicted average molecular mass of 65.107 kDa), following a similar strategy of expression in EV136 cells with or without co-expressing *Gk*Maf (**Supplementary Figure S4A**). Following purification (**Supplementary Figures S4B and S4C**), protease digestion and tandem mass spectrometry analysis (**Supplementary Tables S9a, S9b, S9c, S10, and S11)**, the ion fragmentation patterns could be used to assign two glycosylation sites, L<u>S</u>SGLR wherein the underlined Ser residue was modified with a 221 Da (KDO) moiety and LS<u>S</u>GLR wherein the underlined Ser residue was modified with a 291 Da (Neu5Ac) moiety (**Supplementary Figure S5, Supplementary Tables S12a-e, and additional Supplementary data**). These glycosylation sites map to the distal end of the N-terminal D0 domain at the loop adjoining the D1 domain. We did not identify any glycosylation sites in the D2 and D3 domains, or in the conserved Thr residues of the sequence identical D1 loop region corresponding to the O-sialylated T213 and T270 residues in recombinant *Gk*FlaA2, perhaps because *Gk*Maf could not effectively glycosylate *Gk*FlaA1 under these conditions or because these residues are not surface exposed in *Gk*FlaA1 due to the presence of the D2 and D3 domains.

### *C. botulinum* Maf displays promiscuous sialyltransferase activity in *E. coli* strains overproducing CMP-Neu5Ac

Flagellins of different strains of the human pathogen *C. botulinum* have been reported to be modified with various glycan moieties such as a legionaminic acid derivative, 7-acetamido-5-(N-methyl-glutam-4-yl)-amino-3,5,7,9-tetradeoxy-D-glycero-alpha-D-galacto-nonulosonic acid, a di-acetamido-substituted hexuronic acid sugar, a tri-acetamido-substituted hexuronic acid sugar, and a glycan of mass 696 Da ^9^. Flagellins from *C. botulinum* F Str. Langeland and *C. botulinum* strain FE9909ACS Alberta are glycosylated with the legionaminic acid derivative, 7-acetamido-5-(N-methyl-glutam-4-yl)-amino-3,5,7,9-tetradeoxy-D-glycero-alpha-D-galacto-nonulosonic acid, and share 100% amino acid sequence identity, and seven glycosylation sites have been identified in the flagellin from *C. botulinum* strain FE9909ACS Alberta ^9^. The flagellar locus of *C. botulinum* F Str. Langeland comprises genes for flagellum structural components, flagellin genes, genes for flagellin glycan biosynthesis and a single *maf* gene (NCBI accession no. WP_003359189.1; encoding a protein with a Maf_flag10 domain typical of Maf proteins) neighbouring a flagellin gene, *fla* (NCBI accession no. WP_003359201.1) (**Figure 5A**). We selected this Maf-flagellin pair (referred to here as *Cb*Maf and *Cb*Fla, respectively) for heterologous expression in the *E. coli* strains, EV36, EV136 and EV240, with the expectation that *Cb*Maf, being a putative flagellin legionaminic acid transferase, might display donor substrate promiscuity and accept CMP-Neu5Ac instead of CMP-legionaminic acid as donor substrate (**Figure 5B**). We based our assumption on the close stereochemical structural similarity of Neu5Ac and legionaminic acid, and on evidence for the reverse scenario, i.e., that mammalian and engineered bacterial sialyltransferases are able to use CMP-legionaminic acid instead of CMP-Neu5Ac as donor substrate ^62, 63, 64^.

**Figure 5:**
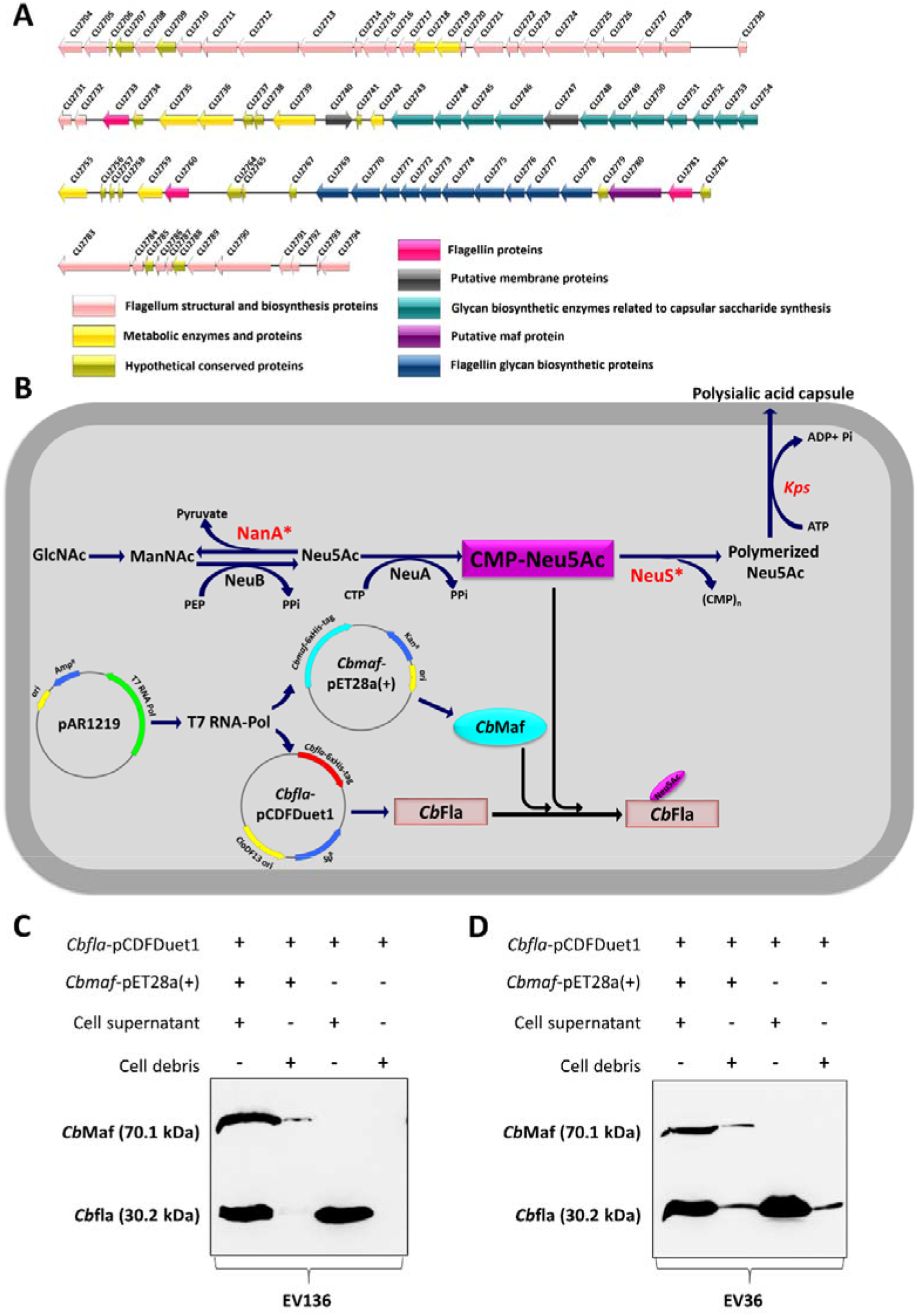
Co-expression of *Cb*Fla with *Cb*Maf. **(A)** Flagellin glycosylation island of *Clostridium botulinum* F strain Langeland. The putative ORFs of Maf and flagellin proteins have been shown in purple and pink colour, respectively. **(B)** Strategy for T7 based co-expression of *Cb*Fla and *Cb*Maf in recombinant *E. coli* K12:K1 hybrid system overproducing the donor substrate CMP-Neu5Ac and harboring pAR1219 plasmid expressing T7 RNA Polymerase. **(C, D)** Western analysis of cell-free supernatant and cell debris fractions of cell lysates of EV36 **(C)** and EV136 **(D)** cells transformed with *Cbfla*-pCDF-Duet-1 vector and *Cbmaf*-pET-28a(+). Recombinant *Cb*Fla and *Cb*Maf proteins were detected by western analysis with mouse anti-6XHis antibody.

We confirmed co-expression of recombinant *Cb*Maf and *Cb*Fla in EV136, EV36 and EV240 cells by western analysis with anti-6XHis-tag antibody (**Figures 4C and 4D, Supplementary figure S6**) and enriched recombinant *Cb*Fla from these cells by immobilized metal affinity chromatography (**Supplementary figure S7**). We also singly expressed and purified recombinant *Cb*Fla (**Figures 4C and 4D, Supplementary figures S6 and S7**).

We performed on-blot periodate oxidation and aniline-catalyzed oxime ligation (PAL) with aminooxy-biotin ^53^ and found that *Cb*Fla was heavily biotinylated in the EV136 and EV240 strains when co-expressed with *Cb*Maf but not when expressed singly (**Figure 6A**). *Cb*Fla, when coexpressed with *Cb*Maf, also displayed less electrophoretic mobility compared to the singly expressed *Cb*Fla in these strains (**Figure 6A**). *Cb*Fla, coexpressed with *Cb*Maf in the EV36 strain, also displayed less electrophoretic mobility than singly expressed *Cb*Fla but displayed relatively little biotinylation (**Figure 6A**). This result suggested that whereas *Cb*Maf displayed robust flagellin sialyltransferase activity in EV136 and EV240 cells, it displayed lower flagellin sialyltransferase activity in EV36 cells, perhaps due to the lower level of CMP-Neu5Ac in this strain.

**Figure 6:**
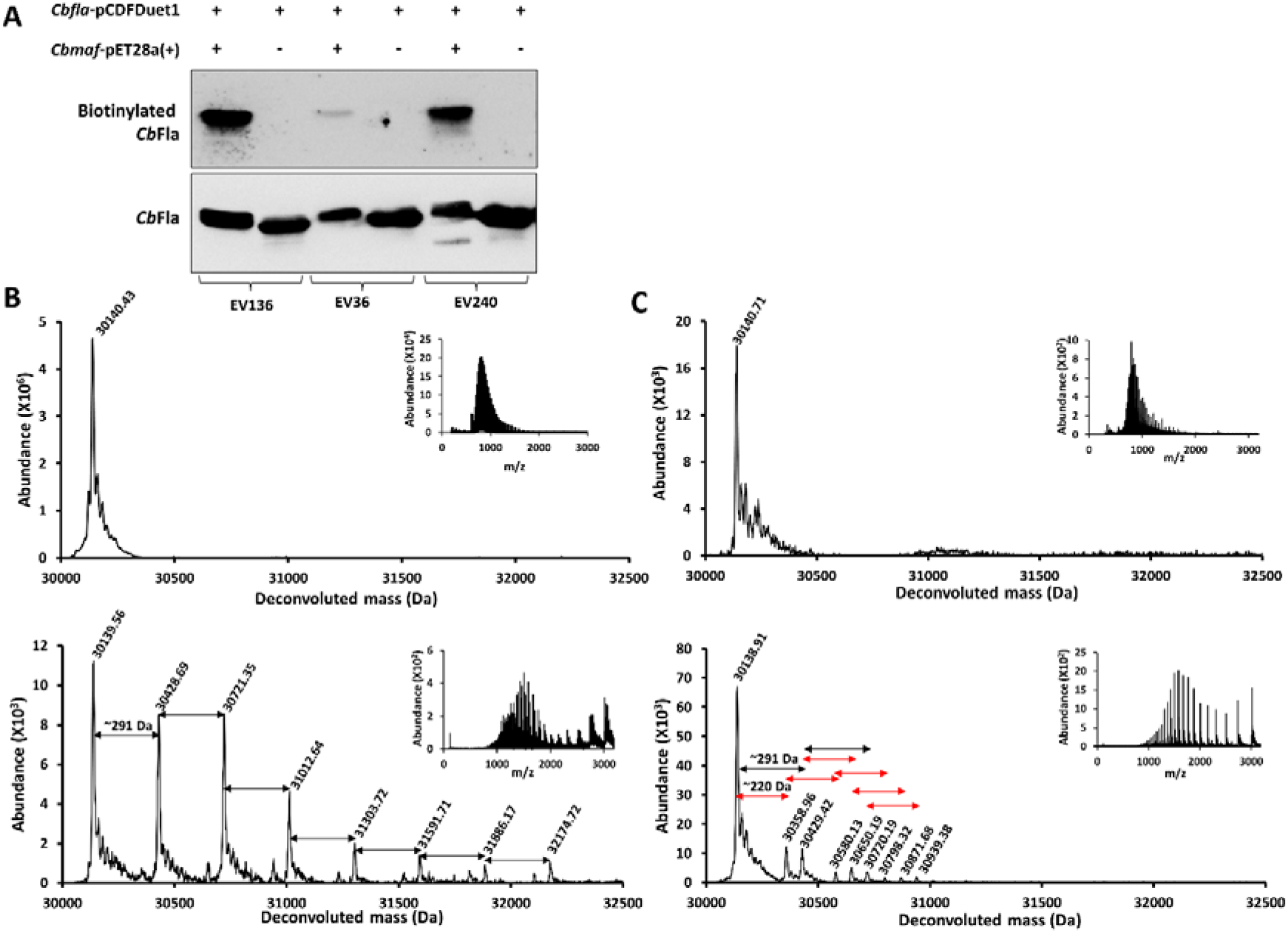
*Cb*Maf glycosylates *Cb*Fla. **(A)** On-blot periodate oxidation and biotinylation of *Cb*Fla singly expressed or co-expressed with *Cb*Maf in EV36, EV136 and EV240 cells. Biotinylation was detected with HRP-streptavidin. Total protein was detected with mouse anti-6XHis antibody **(B, C)** Intact mass measurement of *Cb*Fla singly expressed (upper panel) or co-expressed with *Cb*Maf and purified from EV136 cells **(B)** or EV36 cells **(C**). The proteins were acetone precipitated and subjected to LC-MS in positive mode ionization. Inset shows the ionization spectrum. The satellite peaks correspond to sodiated adducts.

We performed intact mass analysis of *Cb*Fla by liquid chromatography - mass spectrometry. *Cb*Fla singly expressed and purified from EV136 cells had a molecular mass of 30.143 kDa (theoretically predicted average molecular mass: 30.270 kDa) (**Figure 6B upper panel**). This suggests that the recombinant protein undergoes N-ter Met cleavage. However, in addition to a peak of molecular mass 30.139 kDa, the deconvoluted intact mass spectrum of *Cb*Fla purified from EV136 cells co-expressing *Cb*Maf displayed seven additional prominent peaks of molecular masses, 30.431 kDa, 30.722 kDa, 31.016 kDa, 31.305 kDa, 31.595 kDa, 31.886 kDa, and 32.175 kDa, each differing by 291 Da from the previous peak and indicative of the presence of up to at least seven Neu5Ac moieties on *Cb*Fla (**Figure 6B lower panel**).

Singly expressed *Cb*Fla purified from EV36 cells had a molecular mass of 30.141 kDa (**Figure 6C, upper panel**). Intact mass analysis of *Cb*Fla purified from EV36 cells co-expressing *Cb*Maf indicated the presence of two Neu5Ac modified peaks of molecular masses, 30.429 kDa and 30.720 kDa, differing by 291 Da and 582 Da, respectively, from the unmodified peak of molecular mass 30.138 kDa (**Figure 6C lower panel**). Additionally, we observed peaks of molecular masses, 30.358 kDa, 30.580 kDa, and 30.798 kDa, differing by 220 Da, 440 Da, and 660 Da, respectively, from the unmodified peak, and indicative of the presence of up to three KDO moieties (**Figure 6C lower panel**). We also identified peaks of molecular masses 30.650 kDa, corresponding to modification with one Neu5Ac moiety and one KDO moiety, 30.871 kDa, corresponding to modification with one Neu5Ac and two KDO moieties, and 30.939 kDa, corresponding to modification with two Neu5Ac and two KDO moieties (**Figure 6C lower panel**). Similar results were obtained upon intact mass analysis of *Cb*Fla purified from EV240 cells (**Supplementary Figure S8**).

### Glycosylation sites of *Cb*Fla

We obtained ~100 peptide hits with confidently assigned glycosylation sites by tandem mass spectrometry of protease-released peptides from *Cb*Fla purified from EV136 cells or EV36 cells with or without *C*bMaf co-expression followed by searches with Modscore algorithm ^61^ (**Supplementary tables S13a-b, S14, S15, S16, S17, S18a-e and S19a-b, and additional Supplementary data**). A single peptide of *Cb*Fla singly expressed and purified from EV136 cells passed the threshold of Modscore value 19 ≥ (P ≤ 0.01) for Site 1 Position_A and Site 1 Position_B, indicating a low frequency of non-specific hits (**Supplementary table S18c**). The glycosylation sites that we identified in *Cb*Fla include all the seven Ser residues previously identified as glycosylated with the legionaminic acid derivative in *C. botulinum* ^9^. Additionally, we identified five Thr residues in and around the D1 loop region to be glycosylated (**Figure 7A, 7B**). Annotated spectra of a sample peptide from *Cb*Fla expressed in EV136 and EV36 cells, <u>S</u>M*VFQIGANKDQVMELTIAGMGTSALK, is provided, which indicates the presence of fragment ions retaining the glycan moiety, Neu5Ac (291 Da) or KDO (220 Da), respectively (**Figure 7C, 7D**). The structural model of *Cb*Fla depicting the glycosites identified in this study are shown in **Figures 7E and 7F**.

**Figure 7:**
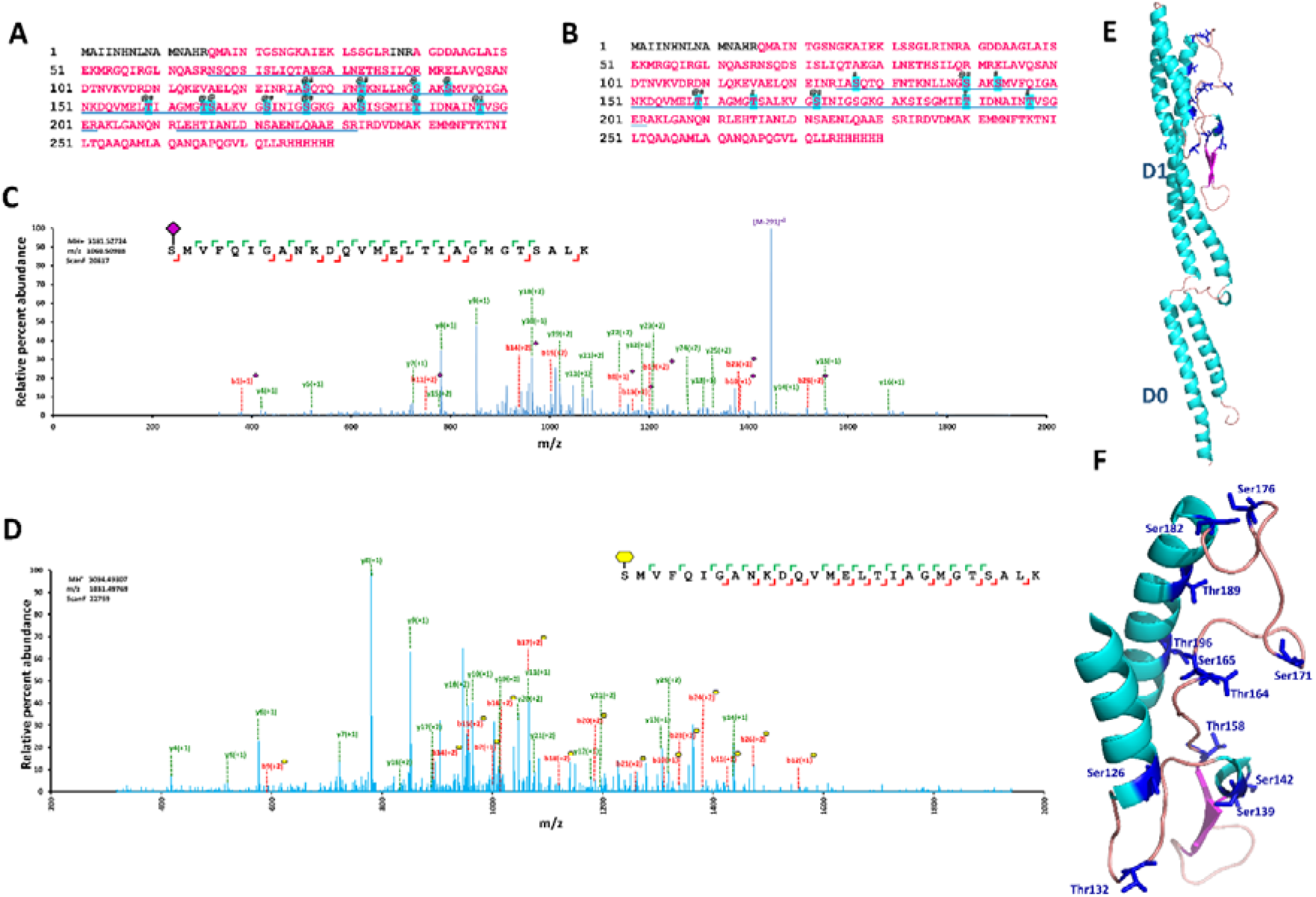
Identification of glycosites in *Cb*Fla. **(A, B)** Sequence coverage of tandem mass spectrometry data obtained for *Cb*Fla purified from EV136 **(A)** and EV36 cells **(B)**. The sequence coverage is shown as pink font. The modified sequence coverage is underlined blue. The sites modified have been highlighted in blue. Neu5Ac is represented by @ symbol and KDO by #. **(C, D)** MS/MS spectra indicating fragment ions of tryptic peptides of *Cb*Fla in EV136 **(C**) and EV36 **(D)** cells as per SEQUEST ^97^. y- and b- ions have been shown in green and red colour, respectively. Relative percent abundances (relative to most abundant fragment ion) are plotted on the y-axis. Neu5Ac and KDO are represented by a pink diamond and a yellow hexagon, respectively, as per symbol nomenclature for graphical representations of glycans ^100^ **(E)** and **(F)** show structure model of *Cb*Fla generated by iTASSER ^101^ using 5wjt as template, with Ser/Thr sites modifications (in stick form) as per MS/MS data.

### *Cb*Maf and *Gk*Maf amino acid residues involved in catalytic activity

Sulzenbacher et al have previously demonstrated that E324 and D415 are critical for the enzyme activity of *M. magneticum* Maf (PDB ID: 5mu5) ^28^. In order to validate the flagellin glycosyltransferase activity observed for *Cb*Maf and *Gk*Maf, we performed site-directed mutagenesis of the corresponding residues in sequence/structure (**Supplementary Figure S9A, S9B, and S9C**) - D272A and D357A in *Cb*Maf, and D285A and N379A in *Gk*Maf. We additionally mutated a Ser residue (S334 and S356 in *Cb*Maf and *Gk*Maf, respectively), conserved in sequence among Maf proteins (**Supplementary Figures S9A, S9B and S9C**), and present in the three-dimensional vicinity of the amino acid residues demonstrated to be important for catalysis, to assess if it was also important for enzymatic activity. We did not observe any flagellin glycosyltransferase activity upon co-expression of any of these Maf mutants with their cognate flagellins – *Cb*Fla/*Gk*FlaA2 (**Figure 8A and Supplementary Figures S10a and S10b**) as evident from lack of sensitivity to periodate oxidation and reaction with aminooxy biotin (**Figure 8B and Supplementary Figures S10c and S10d**), and absence of any modified peaks in intact mass analysis (**Figure 8C**). We conclude that these residues are important for the catalytic activity of Maf.

**Figure 8:**
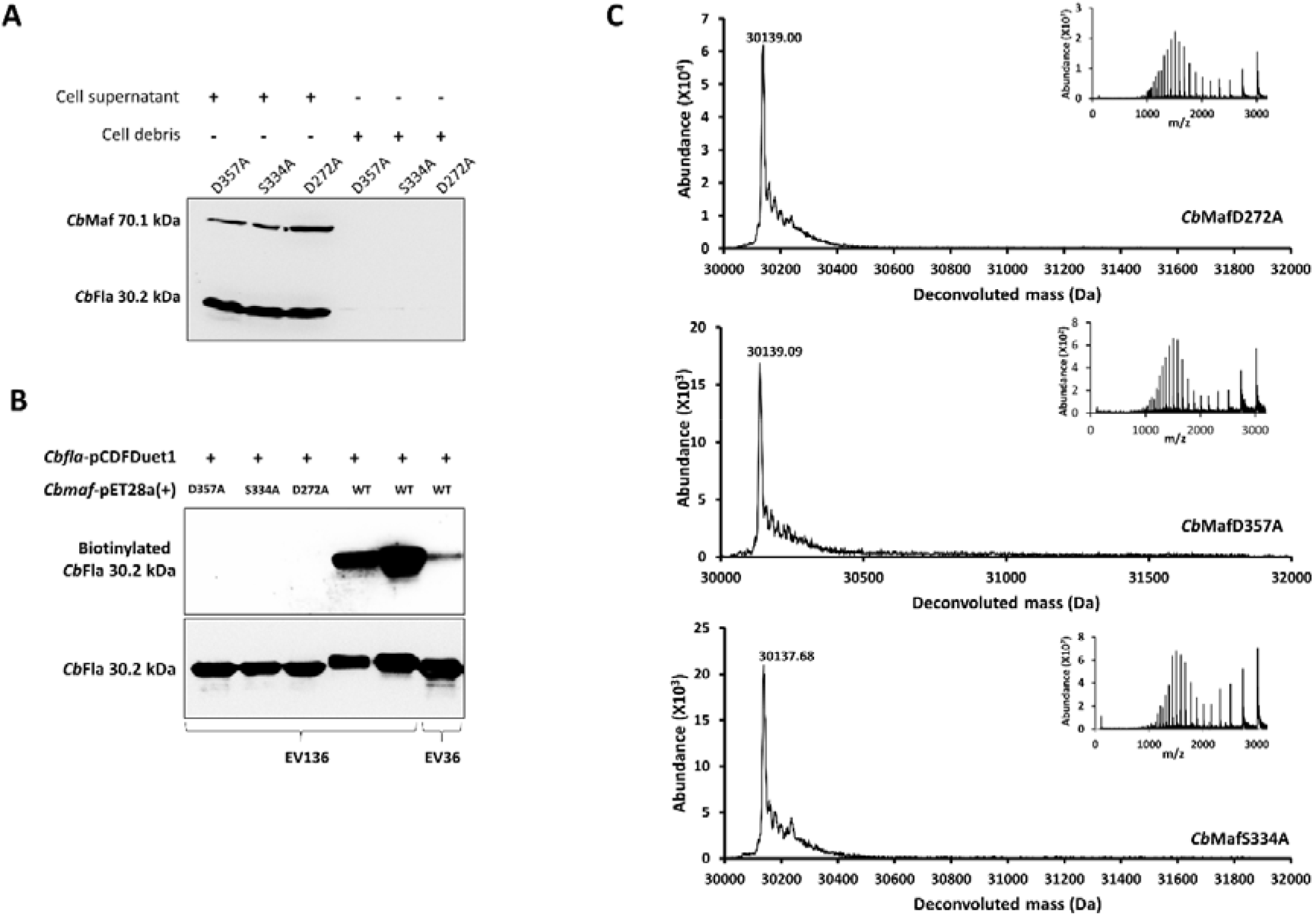
D272, D257 and S334 of *Cb*Maf are required for glycosyltransferase activity. **(A)** Western analysis of *Cb*Fla singly expressed or coexpressed with D272A, D257A and S334A mutants of *Cb*Maf. *Cb*Maf and *Cb*Fla proteins were detected with anti-S tag antibody and mouse anti-6XHis antibody, respectively. **(B)** On-blot periodate oxidation and biotinylation of *Cb*Fla singly expressed or co-expressed with wildtype and mutant *Cb*Maf in EV136 and EV36 cells. Biotinylation was detected with HRP-streptavidin. Total *Cb*Fla protein was detected with mouse anti-6XHis antibody. **(C)** Intact mass measurement of *Cb*Fla singly expressed (upper panel) or co-expressed with *Cb*Maf mutants and purified from EV136 cells. The proteins were acetone precipitated and subjected to LC-MS in positive mode ionization. Inset shows the ionization spectrum. The satellite peaks correspond to sodiated adducts

### *C. botulinum* Maf displays promiscuous KDO transferase activity in *E. coli* BL21

We also assayed *Cb*Maf for flagellin KDO transferase activity in the commonly used *E. coli* lab strain, BL21(DE3), which is known to have KDO2-Lipid(IV)-A in its outer membrane, and should therefore contain CMP-KDO ^57, 65, 66, 67^ (**Figure 9A**). *Cb*Fla, purified from *E. coli* BL21(DE3) cells co-expressing *Cb*Maf (**Figure 9B, Supplementary Figure S11**), was mildly reactive to periodate oxidation and subsequent reaction with aminooxy-biotin (**Figure 9C**) and intact mass analysis confirmed that it was modified by at least six KDO moieties (**Figure 9D**). We subjected *Cb*Fla purified from BL21(DE3) cells to protease digestion, tandem mass spectrometry and Modscore analysis (**Supplementary tables S20a-c, S21a-c, S22, S23a-d**) and assigned glycosylation sites (**Supplementary table S23e and Supplementary Figure S12A-C, and additional Supplementary data**). The glycosylation sites included Ser and Thr residues identified as glycosylated in *Cb*Fla expressed from EV136 cells.

**Figure 9:**
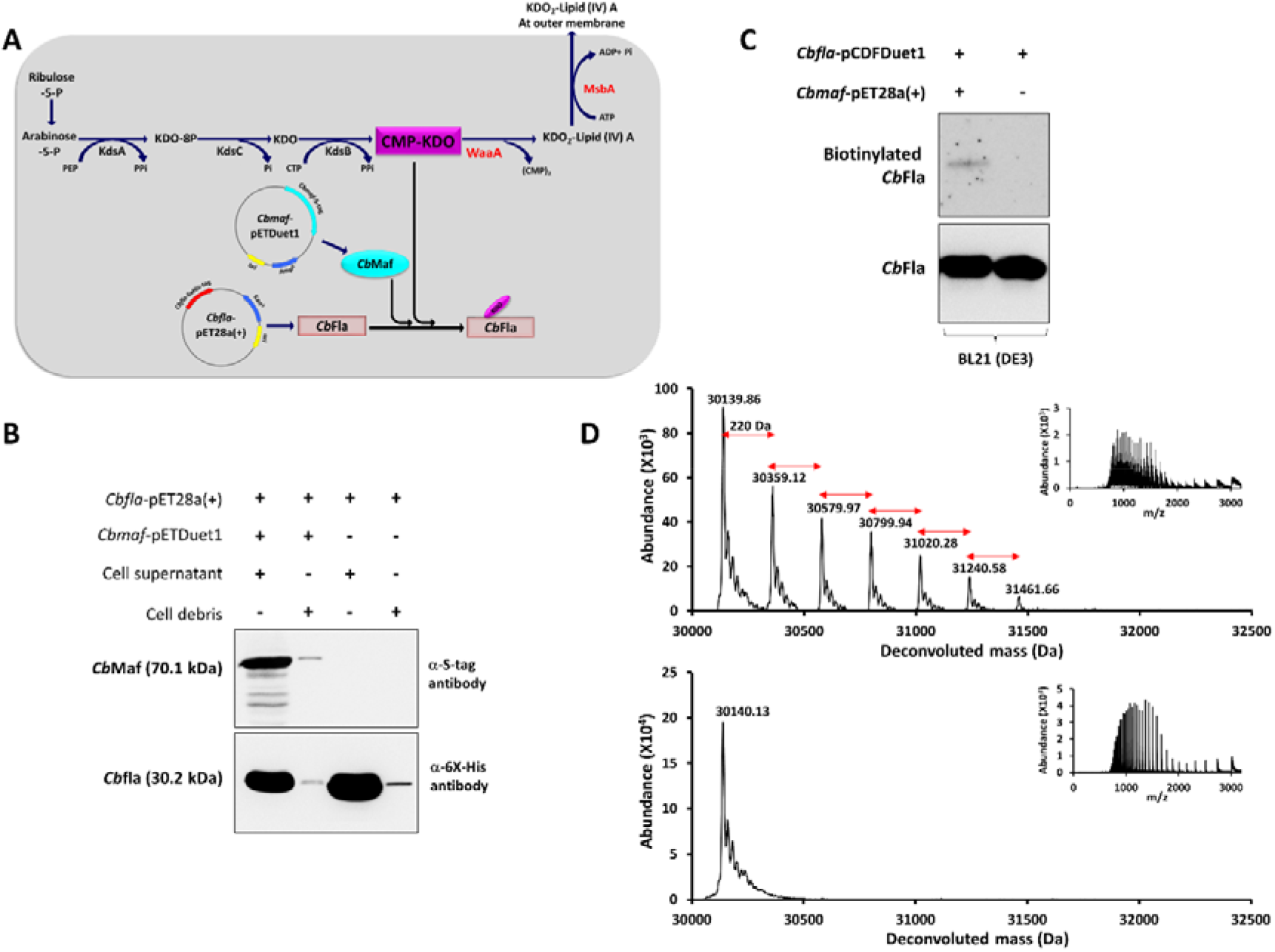
Co-expression of *Cb*Fla and *Cb*Maf in BL21(DE3) cells. **(A)** Strategy for T7 based co-expression of *Cb*Fla and *Cb*Maf in recombinant *E. coli* BL21(DE3) producing CMP-KDO and expressing T7 RNA Polymerase. **(B)** Western analysis of cell-free supernatant and cell debris fractions of cell lysates of BL21(DE3) cells transformed with *Cbfla*-pET-28a(+) vector and *Cbmaf*-pET-Duet-1. Recombinant *Cb*Fla and *Cb*Maf proteins were detected by western analysis with mouse anti-6XHis antibody and mouse anti-S-tag antibody, respectively. **(C)** On-blot periodate oxidation and biotinylation of *Cb*Fla singly expressed or co-expressed with *Cb*Maf in BL21(DE3) cells. Biotinylation was detected with HRP-streptavidin. Total protein was detected with mouse anti-6XHis antibody. **(C)** Intact mass measurement of *Cb*Fla singly expressed (upper panel) or co-expressed with *Cb*Maf and purified from BL21(DE3) cells. The proteins were acetone precipitated and subjected to LC-MS in positive mode ionization. Inset shows the ionization spectrum. The satellite peaks correspond to sodiated adducts.

Our results reiterate the donor substrate promiscuity of *Cb*Maf and indicate that the commonly used *E. coli* BL21(DE3) strain can be employed for further molecular characterization of this class of enzymes.

### Identification of glycan moiety on *Cb*Fla and *Gk*FlaA2

We employed mild acid hydrolysis to release acidic sugar moieties from the glycosylated flagellins, and LC-MS/MS to confirm the identities of the glycan moieties. The tandem mass spectrum of the mild acid hydrolysate of *Cb*Fla purified from EV136 cells matched that of standard Neu5Ac (**Figure 10A**), and the tandem mass spectrum of the mild acid hydrolysate of *Cb*Fla purified from BL21(DE) cells matched that of standard KDO (**Figure 10B**), thereby confirming that the glycans modifying the flagellins in these *E. coli* strains were indeed Neu5Ac and KDO.

**Figure 10:**
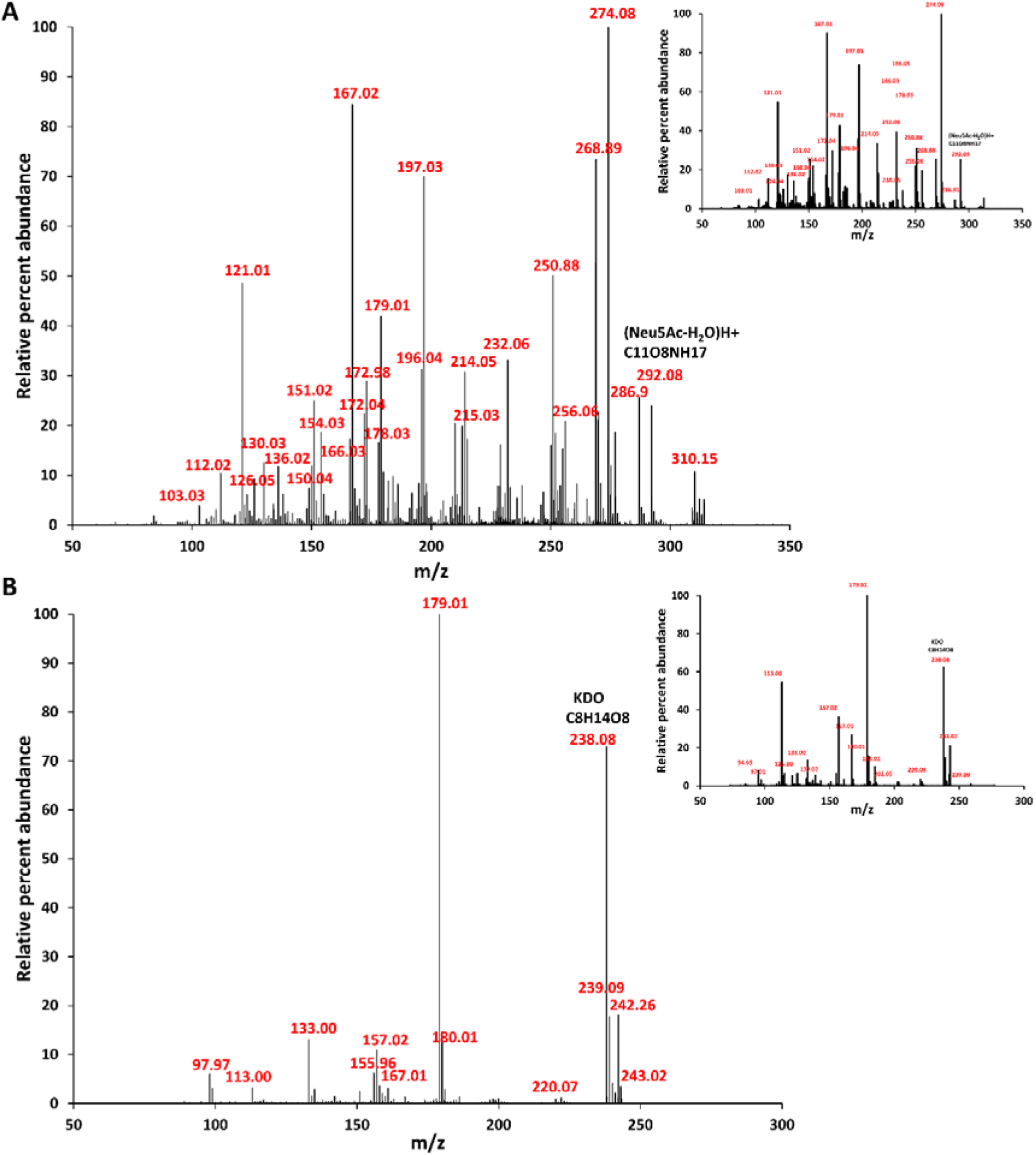
Identification of glycan moiety by tandem mass spectrometry. Tandem mass spectrometry of mild acid extracts of *Cb*Fla co-expressed with *Cb*Maf in EV136 **(A)** and BL21(DE3)cells **(B).** Insets show standard Neu5Ac and KDO fragmentation patterns, respectively.

## Discussion

In this study, we have employed heterologous recombinant co-expression of flagellin and Maf proteins and mass spectrometry to demonstrate that *Gk*Maf, predicted to be a flagellin sialyltransferase based on gene neighbourhood analysis, and *Cb*Maf, a putative flagellin legionaminic acid transferase, can glycosylate Ser/Thr residues of their cognate flagellins with Neu5Ac and KDO. We have also demonstrated here the catalytic involvement of a conserved Ser residue in the Maf_flag10 domain, in addition to residues equivalent to Asp324 and Glu415 residues of *M. magneticum* Maf previously demonstrated to be critical for enzyme activity ^28^.

Sialic acids are the most abundant terminal saccharides in eukaryotic glycoconjugates ^19, 68^. The Carbohydrate-Active enZymes database (CAZy ^69^) family, GT29, comprises eukaryotic sialyltransferases such as ST3Gal, ST6Gal, ST6GalNAc and ST8Sia sialyltransferases that transfer sialic acid (Sia) to glycan residues on glycoconjugates, catalysing the following glycosidic bonds – Siaα2-3Gal, Siaα2-6Gal, Siaα2-6GalNAc and Siaα2-8Sia ^19^. Among prokaryotes, sialic acids are especially prevalent in pathogenic bacteria ^39, 40^. Bacterial sialyltransferases are classified in the CAZy families, GT4, GT38, GT42, GT52, GT38, GT80, GT97 and GT100, and also catalyse transfer of sialic acids to glycan residues to form the glycosidic bonds - Siaα2-3Gal, Siaα2-6Gal, Siaα2-8Sia, and Siaα2-9Sia ^69, 70, 71, 72^. To our knowledge, there is no existing report in the literature of direct Ser/Thr O-sialylation in proteins. Therefore, our study of the putative flagellin sialyltransferase, *Gk*Maf, and the putative legionaminic acid transferase, *Cb*Maf, provides the first evidence for the glycosidic bond, Ser/Thr O-linked Neu5Ac, in any living system. A cursory analysis of flagellar loci in bacterial genomes in the Integrated Microbial Genomes & Microbiomes system v. 5.0 (IMG/M v.5.0) database ^73^ indicate that besides other *Geobacillus kaustophilus* strains, several other bacilli such as *Bacillaceae bacterium* mt8 (Ga0121794_112), *Sporosarcina* sp. P21c (Ga0338388_15), *Cenibacillus caldisaponilyticus* B157 (Ga0302714_1002), *Virgibacillus* sp. SK-1 (Ga0069221_126), *Brevibacillus massiliensis* phR (HGBMDRAFT_CAGW01000083_1.83), *Lysinibacillus sphaericus* LMG 22257 (Ga0175015_11), and *Sporosarcina* sp. P19 (Ga0338387) might also display O-sialylation in their flagellins. Also, *Burkholderia pseudomallei* flagellin is reportedly modified with an unidentified glycan of 291 Da that undergoes fragmentation with neutral losses of water, acetyl group, formic acid, acetamidino group and ammonia; it is possible that this glycan moiety is also Neu5Ac ^74^.

Albeit KDO transferases are involved in the synthesis of lipopolysaccharide and some bacterial and plant polysaccharides ^75, 76, 77, 78, 79^, KDO is not known to modify proteins and there are no reported glycosyltransferases that transfer KDO on to amino acid residues. We could find no entry of any protein modification with KDO in these protein modification databases - FindMod ^80, 81^, UniProt ^82, 83^, DeltaMass (https://abrf.org/delta-mass) ^84, 85^.

Our demonstration of the novel glycosidic linkages, Ser/Thr O-linked Neu5Ac in *Cb*Fla and Ser/Thr O-linked KDO in *Cb*Fla and *Gk*Fla, also highlights the donor substrate promiscuity of *Cb*Maf and *Gk*Maf proteins. Donor substrate promiscuity is quite prevalent in glycosyltransferases (including sialyltransferases ^86^), and is frequently exploited to label glycoconjugates in living cells using metabolic oligosaccharide engineering ^87^ or chemoenzymatic glycan labelling ^88^. In this context, we would like to note that Liu et al have previously demonstrated transfer of azido-labeled pseudaminic acid by *C. jejuni* Maf ^89^.

The non-sialic acid nonulosonic acids, legionaminic acid, pseudaminic acid and their derivatives have been known to decorate Ser/Thr residues in the flagellin of several bacteria such as *C. jejuni, C. coli, H. pylori, A. caviae, A. hydrophila, M. magneticum, Shewanella oneidensis, Treponema denticola, C. botulinum* ^8^. Gene deletion or mutation and complementation studies in the Gram negative bacteria, *A. caviae, A. hydrophila, H. pylori, C. jejuni*, and *M. magneticum*, and structural studies of the *M. magneticum* Maf protein indicate that Maf proteins are flagellin nonulosonic acid glycosyltransferases ^12, 21, 22, 23, 24, 25, 26, 27, 28^. Our study of recombinant *Gk*Maf and *Cb*Maf in *E. coli* strains provides the first direct evidence of the flagellin nonulosonic acid transferase activity of Maf in a heterologous expression system lacking other cognate genes of the flagellar locus. Further, whereas *E. coli* K12 (which is the genetic background of the EV136, EV36 and EV240 strains used here) is known to have a flagellar locus identical to that of *Salmonella* ^90^, *E. coli* B (which is the genetic background of BL21(DE3)) lacks 21 *fli* genes encoding flagellar components and is non-motile ^65^. Therefore, albeit we have not yet successfully demonstrated glycosyltransferase activity in in vitro assays employing recombinant purified *Cb*Maf, *Cb*Fla and CMP-Neu5Ac, the glycosyltransferase activity of *Cb*Maf in BL21(DE3) that we report in this study indicates that Maf might not require any flagellar context to function as a glycosyltransferase. This is in agreement with the notion that native flagellin glycosylation likely occurs in the cytoplasm prior to chaperone binding and protein secretion ^29^. Flagellin glycosylation has been observed in the absence of flagellin chaperone (FlaJ) and FlaJ has been demonstrated to preferentially bind to glycosylated flagellin, thus favoring secretion and assembly of glycosylated flagellin ^29^.

In pathogens such as *Campylobacter jejuni* and *Helicobacter pylori*, flagellin glycosylation is required for flagellum assembly, bacterial motility, and colonization ability, and therefore associated with virulence ^8^. Flagellin glycosylation therefore offers potential as an anti-virulence target, and hence small molecule inhibitors of nonulosonic acid biosynthesis ^91^ or of flagellin glycosyltranferases might inhibit motility and thus virulence. Biosynthetic pathways of flagellin nonulosonic acids, legionaminic acid and pseudaminic acid are well-characterized, and CMP-legionaminic acid and CMP-pseudaminic acid biosynthesis have been reconstituted in vitro using enzymes from *C. jejuni*, and *H. pylori*, respectively ^18, 92, 93, 94^. Albeit these systems are available for studying putative legionaminic acid and pseudaminic acid transferases, our demonstration of donor substrate promiscuity (especially with respect to KDO transferase activity) has the implication that Maf proteins may be easily subjected to high-throughput screening for small molecule inhibitors or to other biochemical characterization using a common lab strain of *E. coli*.

On a broader note, we anticipate that the ability to directly sialylate proteins via Ser/Thr-O-linked sialic acids will dramatically expand the scope of neoglycopeptide synthesis. Further biochemical studies aimed at understanding the sequence/structural prerequisites for glycosylation by Maf will facilitate the development of facile posttranslational protein engineering strategies to install sialic acid moieties and thereby alter the physiochemical properties of recombinant proteins.

## Materials and methods

### Cloning of recombinant Maf and flagellin proteins

The *Clostridium botulinum* F Str. Langeland *maf* (*Cb-maf*, NCBI accession number WP_003359189) and flagellin gene (*Cb-fla,* NCBI accession number WP_003359201), and the *Geobacillus kaustophilus* HTA426 *maf* (*Gk-maf*, NCBI accession number BAD77414.1) and flagellin genes, *flaA1* (*Gk-flaA1*, NCBI accession number BAD77427.1) and *flaA2* (*Gk-flaA2*, NCBI accession number BAD77416.1), were codon optimized for expression in *E. coli* host strains and custom synthesized as clones in pUC57 vector (Genscript, Piscataway, NJ). **Supplementary figure S13** lists details of all expression plasmid constructs used in the study. Cloned inserts in expression plasmids were confirmed by DNA sequencing.

### Expression of recombinant Maf and flagellin proteins in *E. coli* K12-K1 hybrid strains

We obtained *E. coli* K12-K1 hybrid strains EV36, EV136 and EV240 as kind gifts from Prof. Eric R. Vimr. In order to enable overexpression of recombinant proteins under the strong T7 promotor, we used Targe Tron vector pAR1219 (Sigma) that expresses T7 RNA polymerase under the IPTG inducible lac UV5 promotor. For co-expression of Maf and Fla in EV36, EV136 and EV240 cells, we co-transformed *Cb-maf*-pET-28a(+) (pMB1/colE1 *ori*) and *Cb-fla*-pCDFDuet-1 plasmids (CDF *ori*) or the *Gk-maf*-pCDFDuet-1 with *Gk-flaA1*-pET-28a(+) / *Gk-flaA2*-pET-28a(+) plasmids in the cells along with Targe Tron vector pAR1219 (pMB1/colE1 *ori*). We used kanamycin (Goldbio) and ampicillin (Goldbio) as selection markers for pET-28a(+) and pAR1219, respectively (there was no obvious plasmid incompatibility ^95^). We used spectinomycin (Goldbio) as selection marker for pCDFDuet-1 clones as EV36, EV136 and EV240 cells carry selection marker for streptomycin. In the case of EV36 cells, which we found to be inefficient for transformation, we transformed the cells with pAR1219 first and then prepared chemically competent cells of these for further transformation with the Fla and/or Maf expression vectors. We grew secondary cultures of successfully transformed EV36, EV136, or EV240 cells in Luria Bertani liquid medium until the cell density was equivalent to an OD_600nm_ of 0.9, upon which we induced recombinant protein expression by the addition of 1 mM IPTG. Following this, we incubated cells at 37°C with shaking at 200 rpm for 5 hours.

### Expression of recombinant Maf and flagellin proteins in *E. coli* BL21(DE3)

For co-expression in *E. coli* BL21(DE3) (Stratagene), we co-transformed *Cb-maf-*pETDuet-1 (pMB1/colE1 *ori*) and *Cb-fla*-pET-28a(+) plasmids in the cells using ampicillin (Goldbio) and kanamycin (Goldbio) as selection markers, respectively. We grew secondary cultures of successfully transformed *E. coli* BL21(DE3) cells at 37 °C in Luria Bertani liquid medium until the cell density was equivalent to an OD_600nm_ of 0.8, upon which we induced recombinant protein expression by the addition of 1 mM IPTG. Following this, we incubated cells at 37°C with shaking at 200 rpm for 4 hours.

### Western analysis of Maf and flagellin proteins

We confirmed Maf and flagellin co-expression by western blotting the soluble supernatant and insoluble pellet fractions obtained by high-speed centrifugation of cell lysates. We detected co-expressed *Cb*Maf and *Cb*Fla in EV strains by using mouse anti-6X-His tag monoclonal antibody (Invitrogen). We detected the co-expression of *Cb*Maf and *Cb*Fla in *E. coli* BL21(DE3) cells and that of *Gk*Maf and *Gk*FlaA2 in EV strains by using mouse anti-S-tag monoclonal antibody (Novagen) (for the S-tagged Maf proteins) and mouse anti-6X-His tag monoclonal antibody (for the hexahistidine tagged flagellin proteins).

### Purification of flagellin proteins

We purified recombinant hexahistidine tagged flagellin proteins by Ni-NTA metal ion affinity chromatography. Briefly, we lysed cells by sonication in Tris-buffered saline (TBS), i.e., Tris 20 mM, pH 7.5 with 150 mM NaCl, containing N-laurylsarcosine sodium salt at concentration 0.5% for *Cb*Fla and 1% for *Gk*FlaA2. We incubated the cell-free supernatant (obtained following lysis and high-speed centrifugation) with His bind Ni-NTA resin (Pierce), washed the resin with 30 mM imidazole in TBS, and eluted the protein with 300 mM imidazole in TBS. We assessed eluates for purity by SDS-PAGE. We performed acetone precipitation by mixing imidazole eluates with pre-chilled acetone at a ratio of 1:5 and freezing them overnight in a −80 °C freezer. The following day, we washed pellets five times with acetone:water::4:1 and then stored the pellets in a freezer at −20 °C until mass spectrometry analysis.

### Site-directed mutagenesis of Maf

We generated *Cb*Maf mutants - D272A, S334A, and D357A, and *Gk*Maf mutants - D285A, S356A, and N379A, by using appropriate oligonucleotide primers and Quikchange Lightning site directed mutagenesis kit as per manufacturer’s instructions. We co-expressed mutant Maf proteins with the cognate flagellins similar to the wild type Maf proteins.

### Biotinylation of glycosylated flagellin

We performed on-blot biotinylation of sialic acids ^53^. We subjected recombinant, purified flagellins to SDS-PAGE and western blotting on to nitrocellulose membrane. We performed on-blot oxidation with 1 mM sodium metaperiodate in phosphate buffered saline (PBS) (30 mM sodium phosphate, pH 7.4 with 150 mM NaCl) at 4°C for 20 minutes. We then washed the blot with PBS and incubated it in PBS, freshly adjusted to pH 6.7 and containing 250 μM amino-oxybiotinoyl-hydrazine (Invitrogen) and 10 mM aniline, for one hour at room temperature in the dark. We followed this up with washing the blot, overnight blocking in 5% skimmed milk, and probing the blot with HRP-conjugated streptavidin (Jackson ImmunoResearch) followed by detection with Luminata Forte western HRP substrate (Merck-Millipore).

### Mass spectrometry analysis for intact mass measurement

We obtained intact mass data of purified flagellins using Agilent G6550A Quadrupole Time of Flight (Q-TOF) mass spectrometer (CSIR-IMTECH mass spectrometry facility). We dissolved acetone precipitates in 10% acetonitrile, and subjected samples to liquid chromatography in Zorbax 300SB-C8 column (Agilent) at a flow rate of 0.4 ml/minute with acetonitrile as the mobile phase solvent (in concentration gradient). Sample components separated by LC were infused in MS Q-TOF mass spectrometer at a scan rate of 1 scan/minute with the acquisition range 200-3200 m/z. Ionisation source was electrospray ionisation (dual AJS-ESI) in positive ion mode. We analysed the data generated in Agilent Mass-Hunter software in-house. *Cb*Fla typically eluted between 7.4-9 minutes, while *Gk*FlaA2 eluted between 9-9.5 minutes.

*Gk*FlaA2, both expressed singly and co-expressed with *Gk*Maf, was purified by Ni-NTA metal affinity chromatography, acetone precipitated and subjected to intact mass analysis by liquid chromatography coupled electrospray ionisation mass spectrometry. Acetone precipitated purified flagellin is dissolved in 10% acetonitrile to use for mass spectrometry and periodate oxidation and biotinylation.

### Mass spectrometry analysis for identification of glycosylation sites

We obtained results of MS/MS analysis of proteolytic digests of flagellins from Taplin Mass Spectrometry facility, Harvard Medical School, USA employing an LTQ Orbitrap Velos Pro ion-trap mass spectrometer (Thermo Fisher Scientific, Waltham, MA). For this, we subjected the proteins to be analysed to SDS-PAGE and Coomassie staining. Excised gel bands containing the protein (approximately 1 mm^3^ pieces) were subjected to in-gel protease digestion ^96^ at Taplin Mass Spectrometry facility, Harvard Medical School, USA. Gel pieces were washed and dehydrated with acetonitrile for 10 min followed by removal of acetonitrile and drying in a speed-vac, and rehydration in 50 mM bicarbonate. The samples were reduced with 1 mM DTT (in 50 mM ammonium bicarbonate) at 60 °C for 30 minutes, and then cooled to room temperature, Iodoacetamide (in 50 mM ammonium bicarbonate) was added to a final concentration of 5 mM. Following an incubation for 15 minutes at room temperature in the dark, DTT was added to a final concentration of 5 mM to quench the reaction, and sequence grade trypsin was added to a concentration of 5 ng/μl, and the reaction incubated overnight at 37 °C. Peptides were later extracted by removing the ammonium bicarbonate solution, followed by one wash with a solution containing 50% acetonitrile and 1% formic acid. The extracts were desalted by a desalting column made in-house, and then dried in a speed-vac (~1 hr) and stored at 4 ºC until analysis. On the day of analysis the samples were reconstituted in 5 - 10 μl of HPLC solvent A (97.5% water, 2.5% acetonitrile, and 0.1% formic acid). A nano-scale reverse-phase HPLC capillary column was created by packing 2.6 μm C18 spherical silica beads into a fused silica capillary (100 μm inner diameter x ~30 cm length) with a flame-drawn tip (Peng, J and Gygi S.P. Proteomics: The move to mixtures, J. Mass Spec. Oct:36(10):1083-91). After equilibrating the column each sample was loaded via a Famos auto sampler (LC Packings, San Francisco CA) onto the column. A gradient was formed and peptides were eluted with increasing concentrations of solvent B (97.5% acetonitrile, 2.5% water, and 0.1% formic acid). As peptides eluted they were subjected to electrospray ionization and then entered into an LTQ Orbitrap Velos Pro ion-trap mass spectrometer (Thermo Fisher Scientific, Waltham, MA). Peptides were detected, isolated, and fragmented to produce a tandem mass spectrum of specific fragment ions for each peptide. Peptide sequences (and hence protein identity) were determined by matching the protein database with the acquired fragmentation pattern by the software program, SEQUEST (Thermo Fisher Scientific, Waltham, MA) ^97^. All databases included a reverse version of the sequence used and data were filtered to <2% peptide false discovery rate.

The search parameters allowed for a variable modification of 15.994915 Da on methionine. The Modscore algorithm was used to search for modifications of 291 Da (Neu5Ac) or 220 Da (KDO) on Ser/Thr residues. Two scores – Site 1 Position_A and Site 1 Position_B – were used with fragment ion tolerance settings of 0.6 Da and 0.3 Da, respectively. If the Modscore values for Site 1 Position_A and Site 1 Position_B scores were both above 19 for the same site then the location was considered confidently assigned. If the value was 1000 then it was an unequivocal assignment due to there being only one Ser/Thr in the peptide ^61^. Residue numbering and glycosylation site position are according to the amino acid sequence of the recombinant proteins and includes an additional Ala residue following the initial Met residue, and a C-terminal hexahistidine tag.

The mass spectrometry proteomics data reported in the manuscript have been deposited to the ProteomeXchange Consortium ^98^ via the PRIDE ^99^ partner repository with the dataset identifier PXD018167.

### Mass spectrometry analysis for identification of glycan moiety on flagellin

We performed mild acid hydrolysis of flagellin by incubating the protein in 0.1 M HCl at 80 °C for 60 minutes. We subjected these mild acid extracts of purified flagellin to MS/MS analysis at the High Resolution Mass Spectrometry facility at IIT Ropar using a XEVO-G2-XS-QTOF mass spectrometer. Samples were resolved on a C-18 column using a acetonitrile-water gradient on a Waters Acquity UPLC, and subjected to MS/MS in positive ionization mode. For comparison, we subjected standard solutions of Neu5Ac (Sigma) and KDO (Sigma) to MS/MS under the same conditions.

## Acknowledgements

The authors thank Prof. Eric R. Vimr, University of Illinois at Urbana-Champaign for the *E. coli* strains, EV136, EV36, and EV240. The authors acknowledge the Mass Spectrometry facility at CSIR-Institute of Microbial Technology, Chandigarh, the Taplin Biological Mass Spectrometry facility at Harvard Medical School and the High Resolution Mass Spectrometry facility at IIT Ropar, Punjab for their mass spectrometry services. The authors acknowledge CSIR-IMTECH (manuscript communication number 020/2020) for the research facilities and infrastructure.

## Author contributions

RTNC conceived the study. AK and RTNC participated in experimental design. AK performed most of the biochemical experiments and SS performed a few biochemical experiments. RTNC, AK and SS performed data analysis. BP analyzed and interpreted the first intact mass spectrum data of glycosylated flagellin and provided the first critical insight of KDO modification of flagellin in BL21(DE3) cells. AK and RTNC wrote the first draft of the manuscript. All authors participated in editing the manuscript and approved the final version of the manuscript.

