## Supplementary Figures for "Novel Serine/Threonine-O-glycosylation with N-Acetylneuraminic acid and 3-Deoxy-D-manno-octulosonic acid by Maf glycosyltransferases"

T.N.C. Ramya

CSIR-Institute of Microbial Technology

Sector 39-A, Chandigarh 160036 INDIA

**Running Title:**

Ser/Thr O-glycosylation with Neu5Ac and KDO in bacterial flagellins

**Keywords:**

Neu5Ac, sialic acid, KDO, flagellin, Maf, nonulosonic acid, Gram positive, O-glycosylation

**Supplementary data:**

Supplementary figures S1 to S13 are provided in this file. Supplementary tables S1 to S23, and a supplementary data file containing SEQUEST-annotated MS/MS spectra of O-glycosylated peptides with Modscore values of ≥19 for Site 1 Position_A and Site 1 Position_B are provided as separate files of supporting information for this manuscript.

**Supplementary Tables**

***The following supplementary tables are available as worksheets in a separate excel file.***

**Table S1:** Sequest parameters used for searching for Neu5Ac or KDO modified peptides of *Gk*FlaA2

**Table S2a:** Identification of tryptic peptides of *Gk*FlaA2 co-expressed with *Gk*Maf from EV136 cells. Results of dataset 2 (MS/MS results obtained in April 2019).

**Table S2b:** Identification of tryptic peptides of *Gk*FlaA2 co-expressed with *Gk*Maf from EV136 cells. Results of dataset 3 (MS/MS results obtained in July 2019).

**Table S2c:** Identification of Glu-C digested peptides of *Gk*FlaA2 co-expressed with *Gk*Maf from EV136 cells. Results of dataset 3 (MS/MS results obtained in July 2019).

**Table S3a:** Identification of tryptic peptides of *Gk*FlaA2 co-expressed with *Gk*Maf from EV36 cells. Results of dataset 2 (MS/MS results obtained in April 2019).

**Table S3b:** Identification of tryptic peptides of *Gk*FlaA2 co-expressed with *Gk*Maf from EV36 cells. Results of dataset 3 (MS/MS results obtained in July 2019).

**Table S3c:** Identification of Glu-C digested peptides of *Gk*FlaA2 co-expressed with *Gk*Maf from EV36 cells. Results of dataset 3 (MS/MS results obtained in July 2019).

**Table S4:** Identification of tryptic peptides of *Gk*FlaA2 expressed singly from EV136 cells. Results of dataset 2 (MS/MS results obtained in April 2019).

**Table S5:** Identification of tryptic peptides of *Gk*FlaA2 expressed singly from EV36 cells. Results of dataset 2 (MS/MS results obtained in April 2019).

**Table S6:** Details of protein identification of *Gk*FlaA2 expressed singly or co-expressed with *Gk*Maf from EV36 or EV136 cells. Results of datasets 2 and 3 (MS/MS results obtained in April 2019 and July 2019).

**Table S7a:** Results of Modscore algorithm - tryptic peptides of *Gk*FlaA2 co-expressed with *Gk*Maf from EV136 cells. Results of dataset 2 (MS/MS results obtained in April 2019).

**Table S7b:** Results of Modscore algorithm - tryptic and Glu-C digested peptides of *Gk*FlaA2 co-expressed with *Gk*Maf from EV136 cells. Results of dataset 3 (MS/MS results obtained in July 2019).

**Table S7c:** Identification of glycosylated tryptic peptides of *Gk*FlaA2 co-expressed with *Gk*Maf from EV136 cells. Results of dataset 2 (MS/MS results obtained in April 2019).

**Table S7d:** Identification of glycosylated tryptic and Glu-C digested peptides of *Gk*FlaA2 co-expressed with *Gk*Maf from EV136 cells. Results of dataset 3 (MS/MS results obtained in July 2019).

**Table S8a:** Results of Modscore algorithm - tryptic peptides of *Gk*FlaA2 co-expressed with *Gk*Maf from EV36 cells. Results of dataset 2 (MS/MS results obtained in April 2019).

**Table S8b:** Results of Modscore algorithm - tryptic and Glu-C digested peptides of *Gk*FlaA2 co-expressed with *Gk*Maf from EV36 cells. Results of dataset 3 (MS/MS results obtained in July 2019).

**Table S8c:** Identification of glycosylated tryptic peptides of *Gk*FlaA2 co-expressed with *Gk*Maf from EV36 cells. Results of dataset 2 (MS/MS results obtained in April 2019).

**Table S8d:** Identification of glycosylated tryptic and Glu-C digested peptides of *Gk*FlaA2 co-expressed with *Gk*Maf from EV36 cells. Results of dataset 3 (MS/MS results obtained in July 2019).

**Table S9a:** Identification of tryptic peptides of *G*kFlaA1 co-expressed with *Gk*Maf from EV136 cells. Results of dataset 2 (MS/MS results obtained in April 2019).

**Table S9b:** Identification of tryptic peptides of *Gk*FlaA1 co-expressed with *Gk*Maf from EV136 cells. Results of dataset 3 (MS/MS results obtained in July 2019).

**Table S9c:** Identification of AspN digested peptides of *Gk*FlaA1 co-expressed with *Gk*Maf from EV136 cells. Results of dataset 3 (MS/MS results obtained in July 2019).

**Table S10:** Identification of tryptic peptides of *Gk*FlaA1 expressed singly from EV136 cells. Results of dataset 2 (MS/MS results obtained in April 2019).

**Table S11:** Details of protein identification of *Gk*FlaA1 expressed singly or co-expressed with *Gk*Maf from EV136 cells. Results of datasets 2 and 3 (MS/MS results obtained in April 2019 and July 2019).

**Table S12a:** Results of Modscore algorithm - tryptic peptides of *Gk*FlaA1 co-expressed with *Gk*Maf from EV136 cells. Results of dataset 2 (MS/MS results obtained in April 2019).

**Table S12b:** Results of Modscore algorithm - tryptic and AspN digested peptides of *Gk*FlaA1 co-expressed with *Gk*Maf from EV136 cells. Results of dataset 3 (MS/MS results obtained in July 2019).

**Table S12c:** Results of Modscore algorithm - tryptic peptides of *Gk*FlaA1 singly expressed from EV136 cells. Results of dataset 2 (MS/MS results obtained in April 2019).

**Table S12d:** Identification of glycosylated tryptic peptides of *Gk*FlaA1 co-expressed with *Gk*Maf from EV136 cells. Results of dataset 2 (MS/MS results obtained in April 2019).

**Table S12e:** Identification of glycosylated tryptic and AspN digested peptides of *Gk*FlaA1 co-expressed with *Gk*Maf from EV136 cells. Results of dataset 3 (MS/MS results obtained in July 2019).

**Table S13a:** Identification of tryptic peptides of *Cb*Fla co-expressed with *Cb*Maf from EV136 cells. Results of dataset 1 (MS/MS results obtained in January 2019).

**Table S13b:** Identification of tryptic peptides of *Cb*Fla co-expressed with *Cb*Maf from EV136 cells. Results of dataset 2 (MS/MS results obtained in April 2019).

**Table S14:** Identification of tryptic peptides of *Cb*Fla co-expressed with *Cb*Maf from EV36 cells. Results of dataset 2 (MS/MS results obtained in April 2019).

**Table S15:** Identification of tryptic peptides of *Cb*Fla singly expressed from EV136 cells. Results of dataset 2 (MS/MS results obtained in April 2019).

**Table S16:** Identification of tryptic peptides of *Cb*Fla singly expressed from EV36 cells. Results of dataset 2 (MS/MS results obtained in April 2019).

**Table S17:** Details of protein identification of *Cb*Fla expressed singly or co-expressed with *Cb*Maf from EV136 and EV36 cells. Results of datasets 1 and 2 (MS/MS results obtained in January 2019 and April 2019).

**Table S18a:** Results of Modscore algorithm - tryptic peptides of *Cb*Fla co-expressed with *Cb*Maf from EV136 cells. Results of dataset 1 (MS/MS results obtained in January 2019).

**Table S18b:** Results of Modscore algorithm - tryptic peptides of *Cb*Fla co-expressed with *Cb*Maf from EV136 cells. Results of dataset 2 (MS/MS results obtained in April 2019).

**Table S18c:** Results of Modscore algorithm - tryptic peptides of *Cb*Fla singly expressed from EV136 cells. Results of dataset 2 (MS/MS results obtained in April 2019).

**Table S18d:** Identification of glycosylated tryptic peptides of *Cb*Fla co-expressed with *Cb*Maf from EV136 cells. Results of dataset 1 (MS/MS results obtained in January 2019).

**Table S18e:** Identification of glycosylated tryptic peptides of *Cb*Fla co-expressed with *Cb*Maf from EV136 cells. Results of dataset 2 (MS/MS results obtained in April 2019).

**Table S19a:** Results of Modscore algorithm - tryptic peptides of *Cb*Fla co-expressed with *Cb*Maf from EV36 cells. Results of dataset 2 (MS/MS results obtained in April 2019).

**Table S19b:** Identification of glycosylated tryptic peptides of *Cb*Fla co-expressed with *Cb*Maf from EV36 cells. Results of dataset 2 (MS/MS results obtained in April 2019).

**Table S20a:** Identification of tryptic peptides of *Cb*Fla co-expressed with *Cb*Maf from BL21(DE3) cells. Results of dataset 2 (MS/MS results obtained in April 2019).

**Table S20b:** Identification of tryptic peptides of *Cb*Fla co-expressed with *Cb*Maf from BL21(DE3) cells. Results of dataset 3 (MS/MS results obtained in July 2019).

**Table S20c:** Identification of Glu-C digested peptides of *Cb*Fla co-expressed with *Cb*Maf from BL21(DE3) cells. Results of dataset 3 (MS/MS results obtained in July 2019).

**Table S21a:** Identification of tryptic peptides of *Cb*Fla singly expressed from BL21(DE3) cells. Results of dataset 2 (MS/MS results obtained in April 2019).

**Table S21b:** Identification of tryptic peptides of *Cb*Fla singly expressed from BL21(DE3) cells. Results of dataset 3 (MS/MS results obtained in July 2019).

**Table S21c:** Identification of Glu-C digested peptides of *Cb*Fla singly expressed from BL21(DE3) cells. Results of dataset 3 (MS/MS results obtained in July 2019).

**Table S22:** Details of protein identification of *Cb*Fla expressed singly or co-expressed with *Cb*Maf from BL21(DE3) cells. Results of datasets 2 and 3 (MS/MS results obtained in April 2019 and July 2019).

**Table S23a:** Results of Modscore algorithm - tryptic peptides of *Cb*Fla co-expressed with *Cb*Maf from BL21(DE3) cells. Results of dataset 2 (MS/MS results obtained in April 2019).

**Table S23b:** Results of Modscore algorithm - tryptic peptides of *Cb*Fla co-expressed with *Cb*Maf from BL21(DE3) cells. Results of dataset 3 (MS/MS results obtained in July 2019).

**Table S23c:** Results of Modscore algorithm - tryptic peptides of *Cb*Fla singly expressed from BL21(DE3) cells. Results of dataset 2 (MS/MS results obtained in April 2019).

**Table S23d:** Results of Modscore algorithm - tryptic peptides of CbFla singly expressed from BL21(DE3) cells. Results of dataset 3 (MS/MS results obtained in July 2019).

**Table S23e:** Identification of glycosylated tryptic peptides of *Cb*Fla co-expressed with *Cb*Maf from BL21(DE3) cells. Results of dataset 3 (MS/MS results obtained in July 2019).

**Supplementary Figures**

**Supplementary figure 1: *Gk*Maf and *Gk*FlaA2 expression in EV240.** Lysates of EV240 cells transformed with expression vector(s) to singly express *Gk*FlaA2 or co-express *Gk*FlaA2 and *Gk*Maf were subjected to high-speed centrifugation. The cell supernatant and cell debris were subjected to SDS-PAGE and western analysis, and the recombinant proteins *Gk*FlaA2 and *Gk*Maf were detected with anti-6XHis antibody and anti-S tag antibody, respectively.

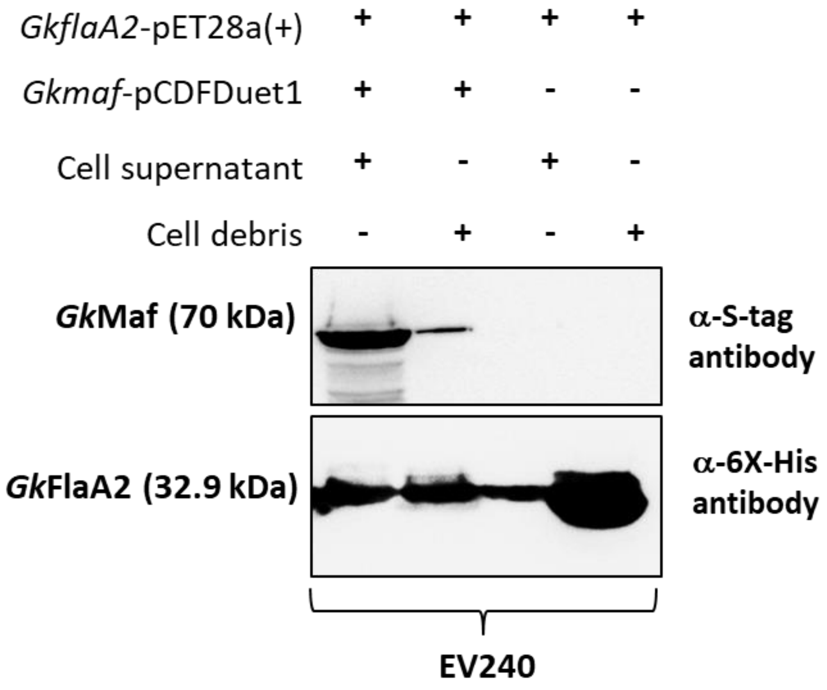

**Supplementary figure 2: *Gk*FlaA2 purification from EV36, EV136 and EV240 cells.**

Coomassie-stained SDS-PAGE gels showing recombinant protein *Gk*FlaA2 singly expressed (i) or co-expressed (ii) with *Gk*Maf in EV36 (**A**), EV136 (**B**) and EV240 (**C**) cells. The recombinant proteins were enriched by Ni-NTA metal ion affinity chromatography. Lanes 1 and 2 show the enriched protein and lane 3 shows the protein molecular weight marker in (**A**) and (**B**). M denotes the protein molecular weight marker in (**C**).

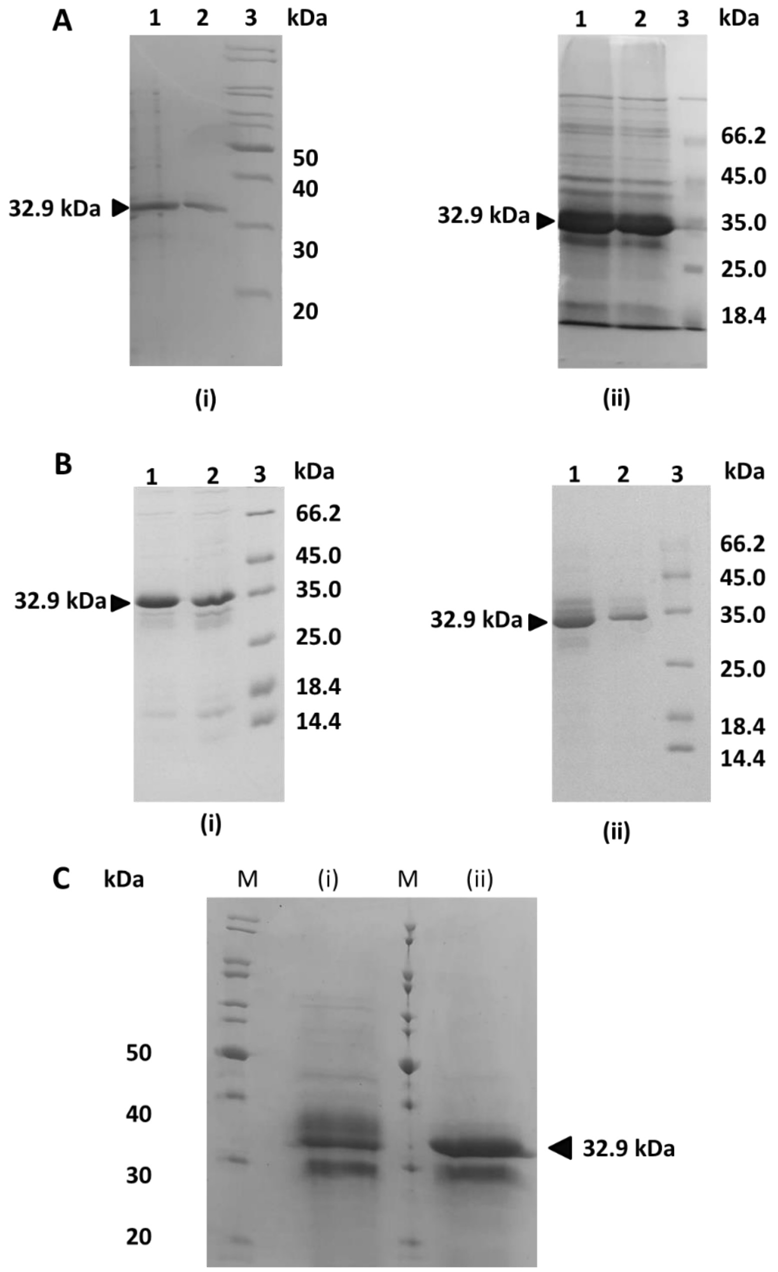

**Supplementary figure 3: Intact mass analysis of *Gk*FlaA2 purified from EV240 cells by ESI-MS analysis.** The *Gk*FlaA2 protein purified from EV240 cells was acetone precipitated and subjected to LC-ESI-MS in positive mode ionization. The deconvoluted mass spectrum is shown indicating intact mass measurement of *Gk*FlaA2. (**A**) Intact mass of *Gk*FlaA2 singly expressed and purified from EV240 cells. (**B**) Intact mass of *Gk*FlaA2 purified from EV240 cells co-expressing *Gk*Maf. Inset shows the ionization spectrum. The satellite peaks correspond to sodiated adducts.

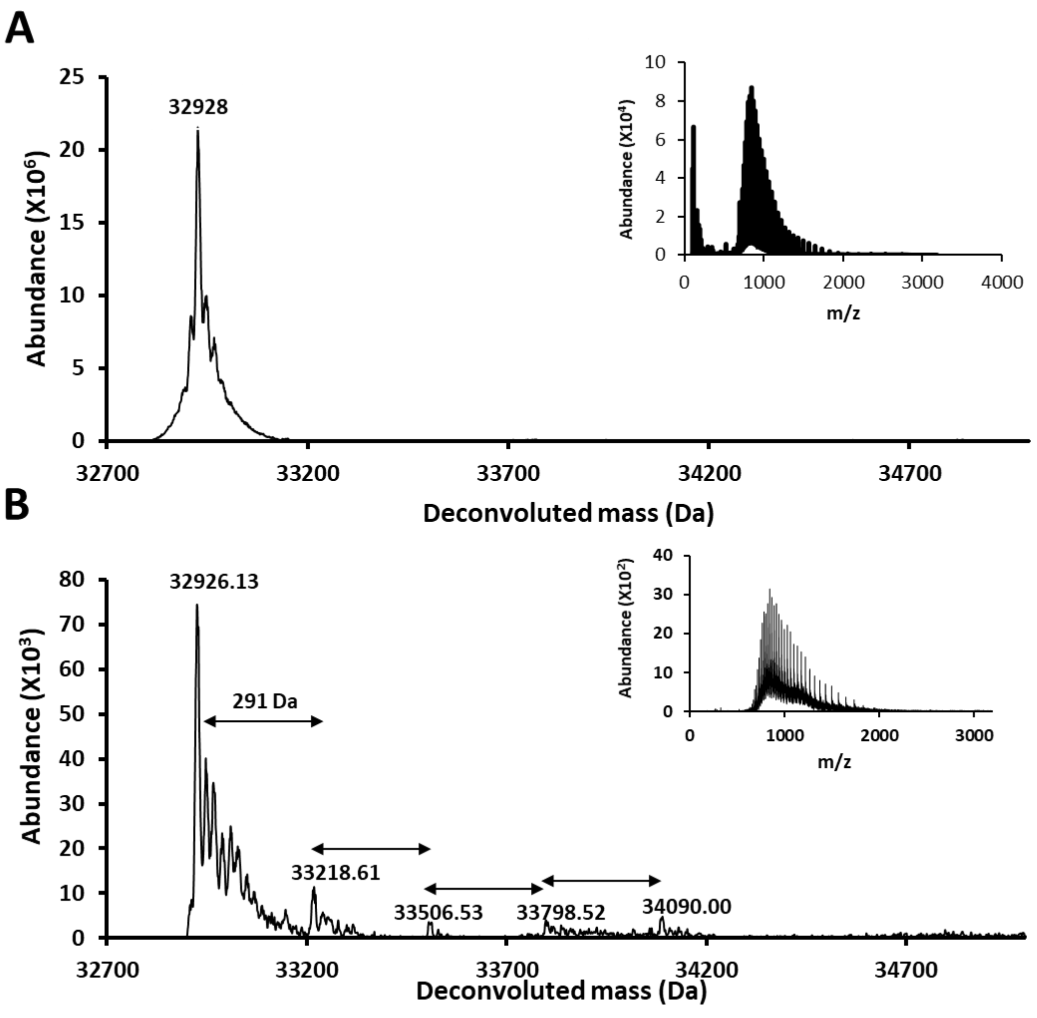

**Supplementary Figure 4: Expression and purification of *Gk*FlaA1 from EV136 cells.** (**A**) Co-expression of *Gk*FlaA1 and *Gk*Maf in EV136 cells transformed with *Gk-maf*-pCDFDuet1 and *Gk-flaA1*-pET-28a(+). (**B**) Purification of *Gk*FlaA1 from EV136 cells transformed with *Gk-flaA1*-pET-28a(+). Lanes 1 and 2: purified *Gk*FlaA1. Lane 3: Molecular marker. (**C**) Purification of *Gk*FlaA1 from EV136 cells transformed with *Gk-maf*-pCDFDuet1 and *Gk-flaA1*-pET-28a(+). Lanes 1 and 2: purified *Gk*FlaA1. Lane 3: Molecular marker.

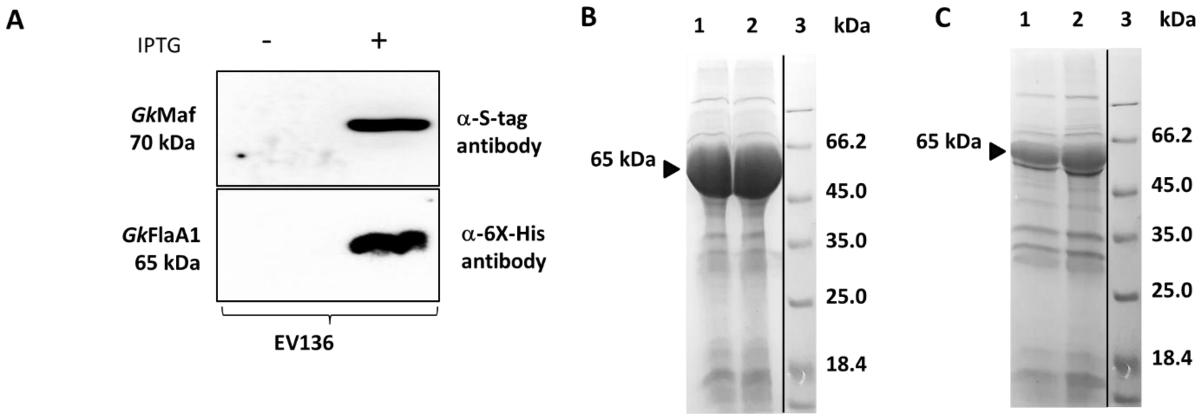

**Supplementary figure 5: Identification of glycosites in *Gk*FlaA1. (A)** Sequence coverage of tandem mass spectrometry data obtained for *Gk*FlaA1 purified from EV136 cells. The sequence coverage is shown as pink font. The modified sequence coverage is underlined blue (for tryptic peptides) and green (for Asp-N digested peptides). The sites modified have been highlighted in blue**.** Neu5Ac is represented by @ symbol and KDO by #. **(B,C)** MS/MS spectra indicating fragment ions of tryptic peptides of *Gk*FlaA1 as per SEQUEST [1]. y- and b-ions have been shown in green and red colour, respectively. Relative percent abundances (relative to most abundant fragment ion) are plotted on the y-axis. Neu5Ac and KDO are represented by a pink diamond and a yellow hexagon, respectively, as per symbol nomenclature for graphical representations of glycans [2]. Peak of mass 274 Da indicates the presence of glycan oxonium ion (Neu5Ac-2$H_2O$). Spectra are of peptides identified and listed in Supplementary tables S12d and S12e. **(D)** shows structure model of *Gk*FlaA1, generated by iTASSER [3] using 5wjt as template, with Ser/Thr glycosites (colored red in stick form) as per MS/MS data.

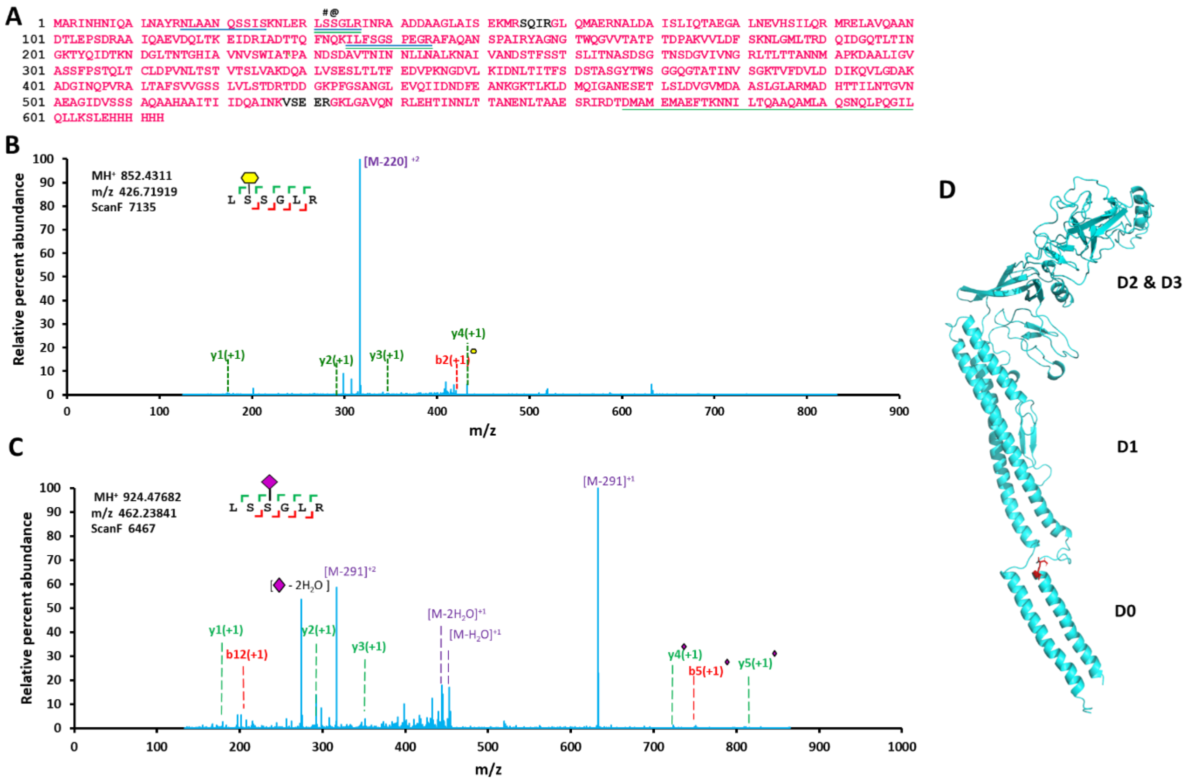

**Supplementary figure 6: *Cb*Fla and *Cb*Maf co-expression in EV 240 cells.** Western analysis of cell-free supernatant and cell debris fractions of cell lysates of EV240 cells transformed with *Cbfla*-pCDF-Duet vector and *Cbmaf*-pET-28a(+). Recombinant *Cb*Fla and *Cb*Maf proteins were detected by western analysis with anti-6XHis antibody.

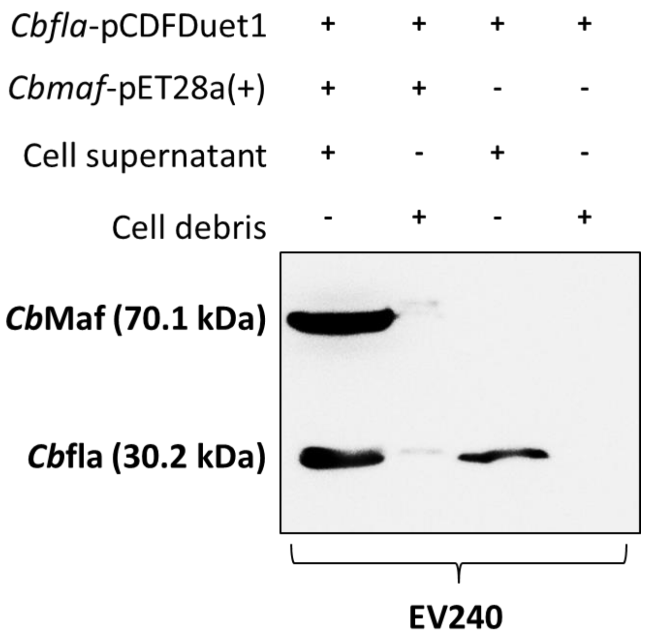

**Supplementary figure 7: *Cb*Fla purification from EV36, EV136 and EV240 cells.**

Coomasie SDS-PAGE gel picture showing recombinant protein *Cb*Fla singly expressed (i) or co-expressed with *Cb*Maf (ii) in EV136 (**A**), EV36 (**B**) and EV240 (**C**) cells. The proteins were purified by Ni-NTA metal ion affinity chromatography. Lanes 1 and 2 show the enriched protein and lane 3 shows the protein molecular weight marker.

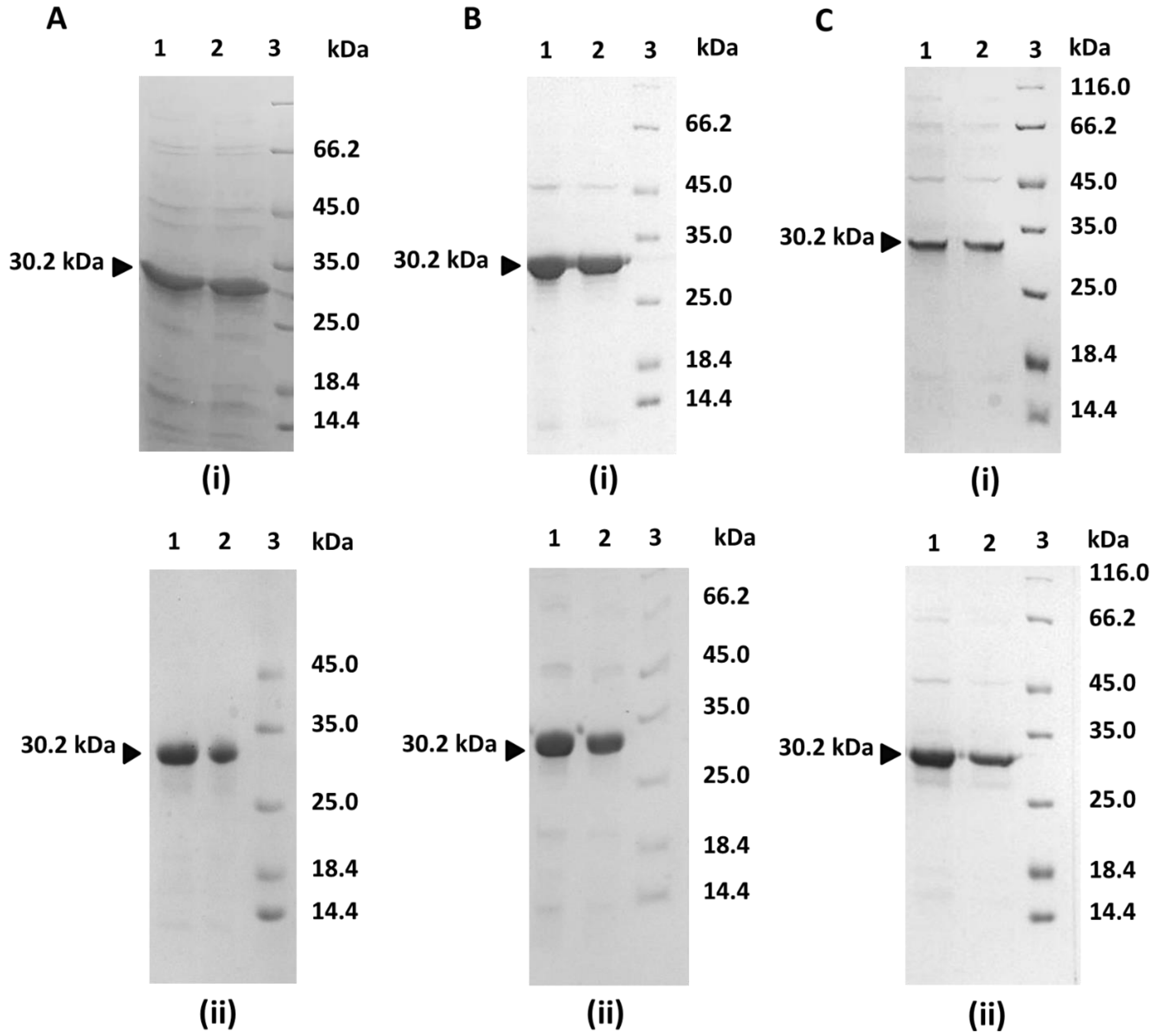

**Supplementary figure 8: Intact mass analysis of *Cb*Fla and *Cb*MFla.** Intact mass measurement of *Cb*Fla singly expressed (**A**) or co-expressed with *Cb*Maf (**B**) and purified from EV240 cells. The proteins were acetone precipitated and subjected to LC-MS in positive mode ionization. Inset shows the ionization spectrum. The satellite peaks correspond to sodiated adducts.

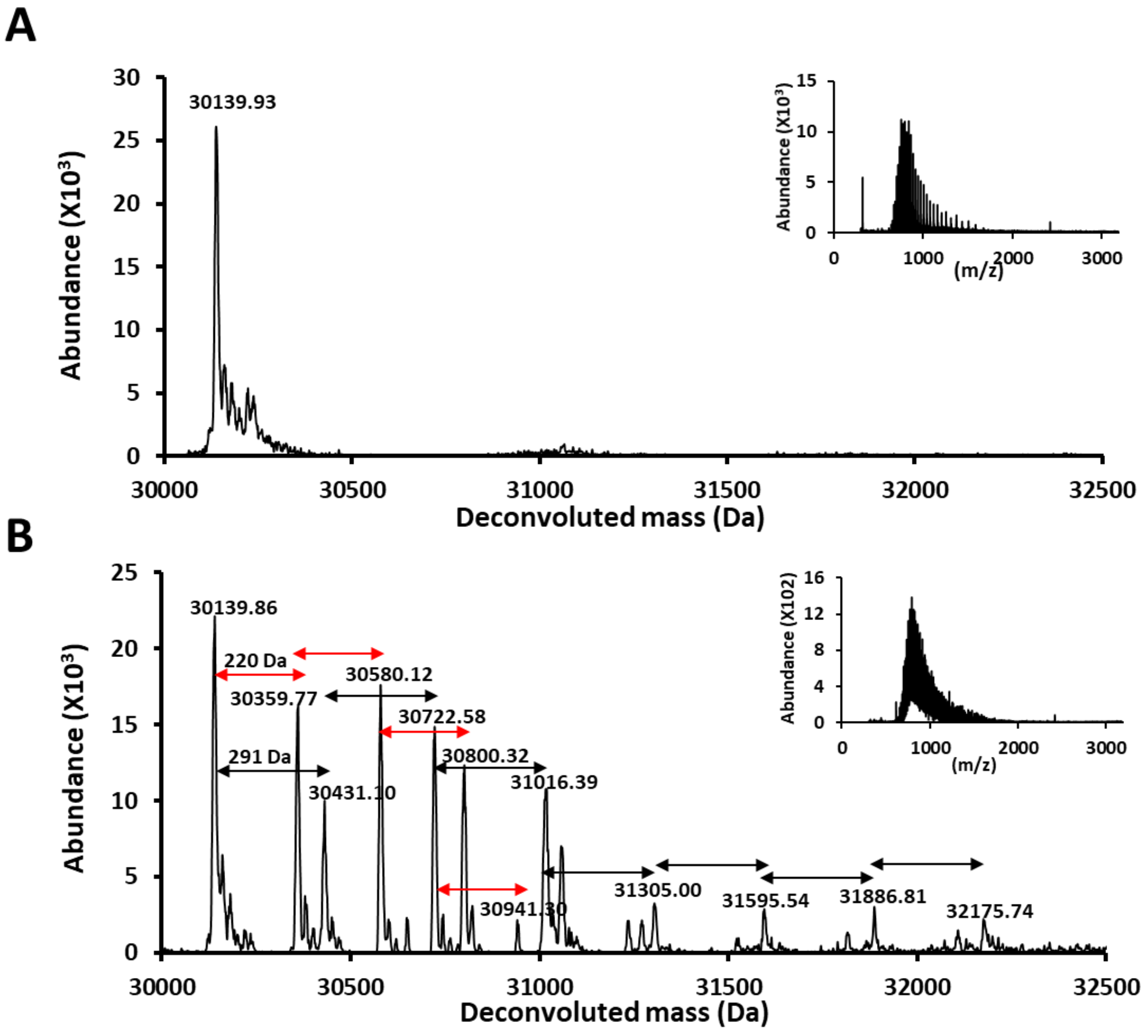

**Supplementary figure 9: Selection of residues for site-directed mutagenesis of *Cb*Maf.** (**A, B**) Sequence alignment of Maf proteins from various bacteria. D272 and D357 of *Cb*Maf are equivalent in position to residues E324 and D415 of *Magnetospirillum magneticum*. The conserved Asp/Glu and Ser residues are marked by red arrows. (**C**) *Cb*Maf domain modelled using *M. magneticum* (PDB id-5mu5) Maf as template (amino acid identity 25.78%). *Cb*Maf and 5mu5 have been depicted in brown and purple colour, respectively.

**A**

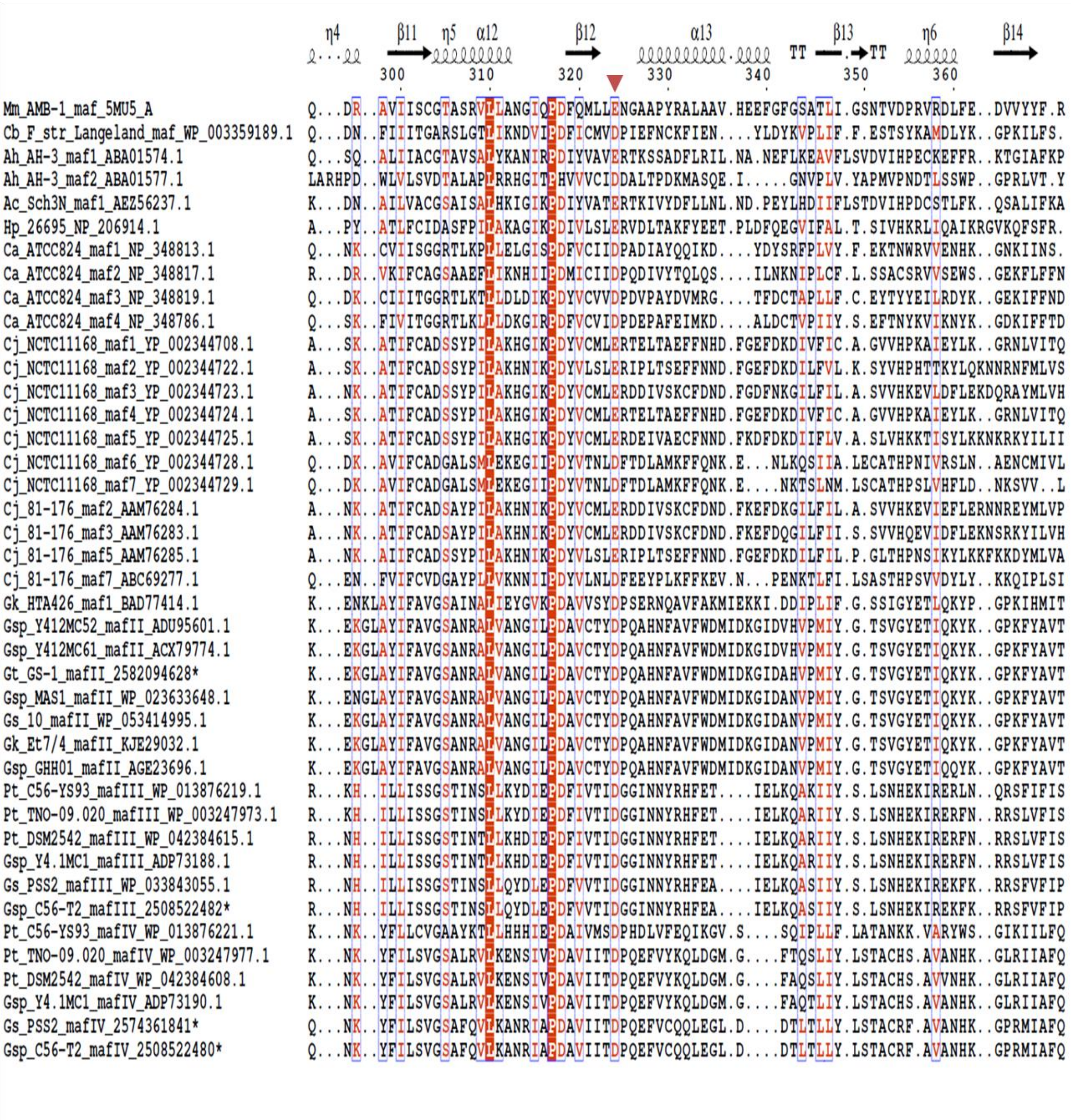

**B**

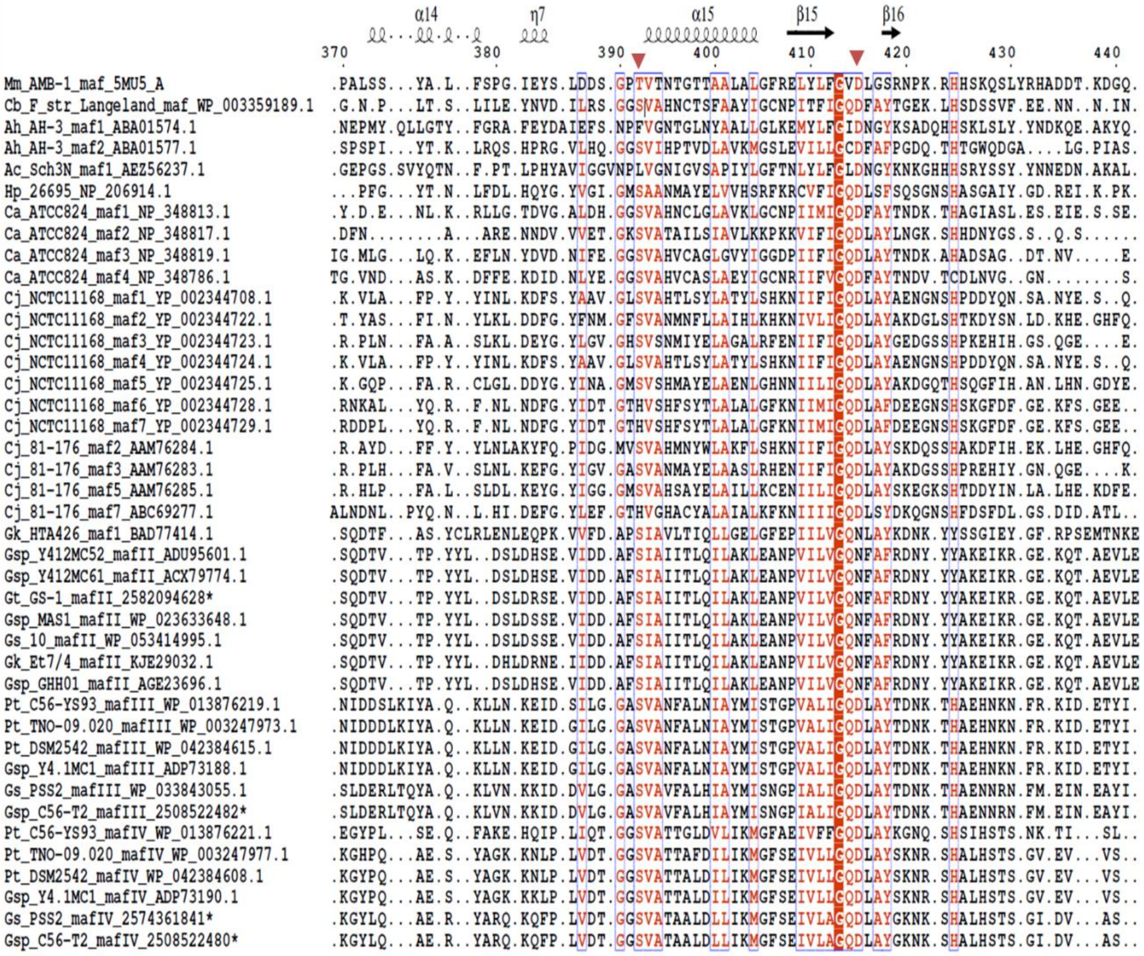
α14
η7
α15
β15
β16
370
380
390
400
410
420
430
440
Mm_AMB-1_maf_5MU5_A
Cb_F_str_Langeland_maf_WP_003359189.1
Ah_AH-3_maf1_ABA01574.1
Ah_AH-3_maf2_ABA01577.1
Ac_Sch3N_maf1_AEZ56237.1
Hp_26695_NP_206914.1
Ca_ATCC824_maf1_NP_348813.1
Ca_ATCC824_maf2_NP_348817.1
Ca_ATCC824_maf3_NP_348819.1
Ca_ATCC824_maf4_NP_348786.1
Cj_NCTC11168_maf1_YP_002344708.1
Cj_NCTC11168_maf2_YP_002344722.1
Cj_NCTC11168_maf3_YP_002344723.1
Cj_NCTC11168_maf4_YP_002344724.1
Cj_NCTC11168_maf5_YP_002344725.1
Cj_NCTC11168_maf6_YP_002344728.1
Cj_NCTC11168_maf7_YP_002344729.1
Cj_81-176_maf2_AAM76284.1
Cj_81-176_maf3_AAM76283.1
Cj_81-176_maf5_AAM76285.1
Cj_81-176_maf7_ABC69277.1
Gk_HTA426_maf1_BAD77414.1
Gsp_Y412MC52_mafII_ADU95601.1
Gsp_Y412MC61_mafII_ACX79774.1
Gt_GS-1_mafII_2582094628*
Gsp_MAS1_mafII_WP_023633648.1
Gs_10_mafII_WP_053414995.1
Gk_Et7/4_mafII_KJE29032.1
Gsp_GHH01_mafII_AGE23696.1
Pt_C56-YS93_mafIII_WP_013876219.1
Pt_TNO-09.020_mafIII_WP_003247973.1
Pt_DSM2542_mafIII_WP_042384615.1
Gsp_Y4.1MC1_mafIII_ADP73188.1
Gs_PSS2_mafIII_WP_033843055.1
Gsp_C56-T2_mafIII_2508522482*
Pt_C56-YS93_mafIV_WP_013876221.1
Pt_TNO-09.020_mafIV_WP_003247977.1
Pt_DSM2542_mafIV_WP_042384608.1
Gsp_Y4.1MC1_mafIV_ADP73190.1
Gs_PSS2_mafIV_2574361841*
Gsp_C56-T2_mafIV_2508522480*

**C**

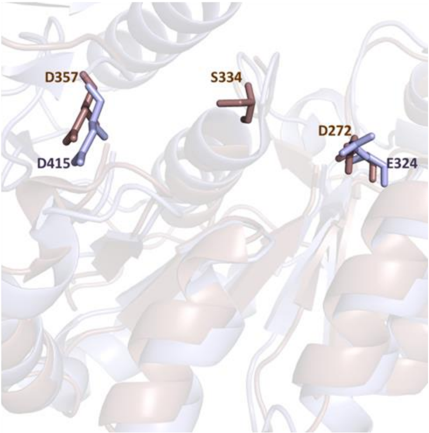
D357
S334
D272
D415
E324

**Supplementary figure 10: D285, S356 and N379 of *Gk*Maf are required for glycosyltransferase activity. (A, B)** Western analysis of *Gk*FlaA2 singly expressed or coexpressed with D285, S356 and N379 mutants of *Gk*Maf. *Gk*Maf and *Gk*FlaA2 proteins were detected with anti-S tag antibody and mouse anti-6XHis antibody, respectively. **(C, D)** On-blot periodate oxidation and biotinylation of *Gk*FlaA2 singly expressed or co-expressed with wildtype and mutant *Gk*Maf in EV136 cells. Biotinylation was detected with HRP-streptavidin. Total protein was detected with mouse anti-6XHis antibody.

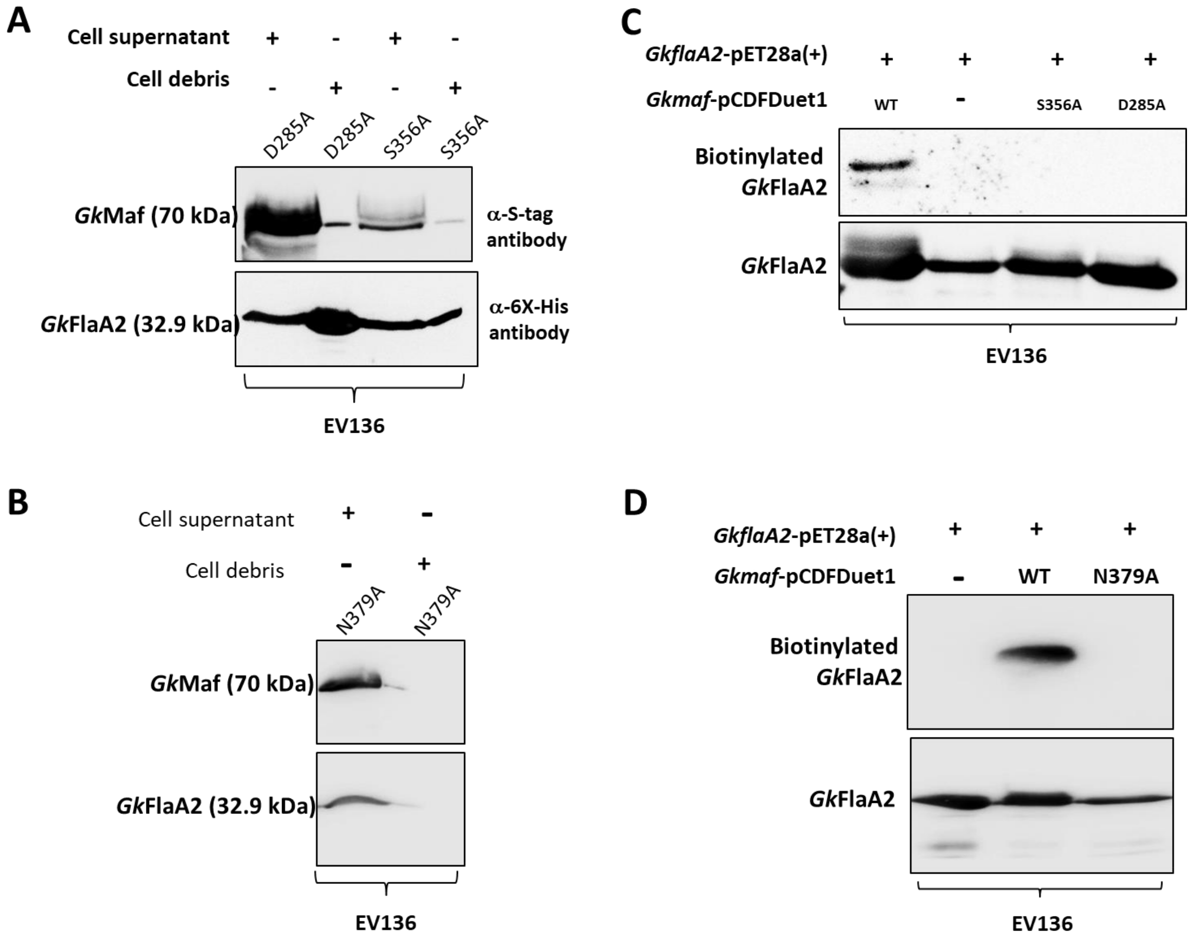

**Supplementary figure 11: *Cb*MafFla purification from BL21(DE3) cells.** Coomasie SDS-PAGE gel picture showing recombinant protein *Cb*Fla singly expressed (**A**) or co-expressed with *Cb*Maf (**B**) in BL21(DE3) cells. The proteins were purified by Ni-NTA metal ion affinity chromatography. Lanes 1 and 2 show the purified protein and lane 3 shows the protein molecular weight marker.

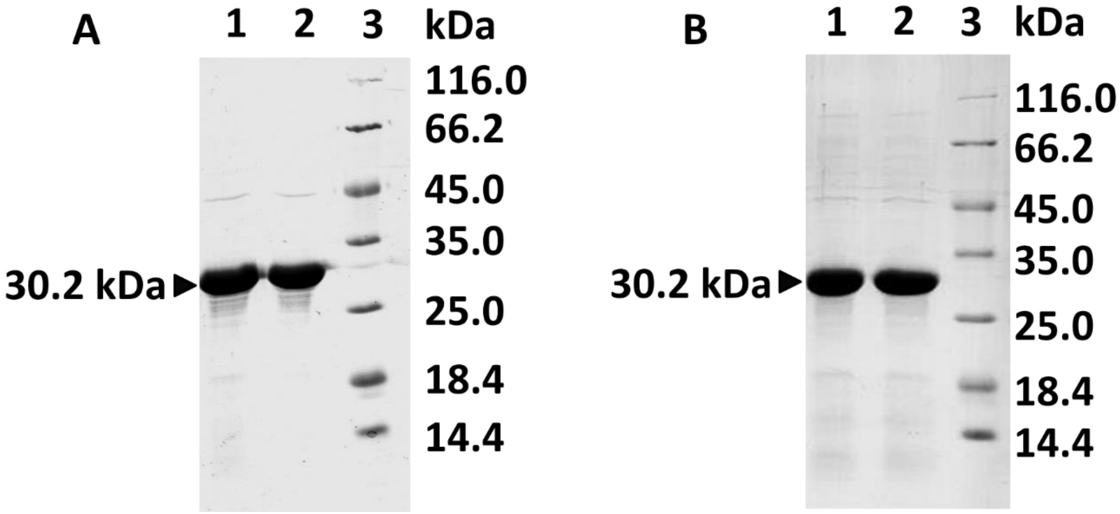

**Supplementary figure 12: Identification of glycosites in *Cb*Fla. (A)** Sequence coverage of tandem mass spectrometry data obtained for *Cb*Fla purified from BL21(DE3) cells. The sequence coverage is shown as pink font. The modified sequence coverage is underlined blue (for tryptic peptides) and green (for Glu-C digested peptides). The sites modified have been highlighted in blue**.** Neu5Ac is represented by @ symbol and KDO by #. **(B, C)** MS/MS spectra indicating fragment ions of tryptic peptides of *Cb*Fla as per SEQUEST [1]. y- and b-ions have been shown in green and red colour, respectively. Relative percent abundances (relative to most abundant fragment ion) are plotted on the y-axis. Neu5Ac and KDO are represented by a pink diamond and a yellow hexagon, respectively, as per symbol nomenclature for graphical representations of glycans [2].

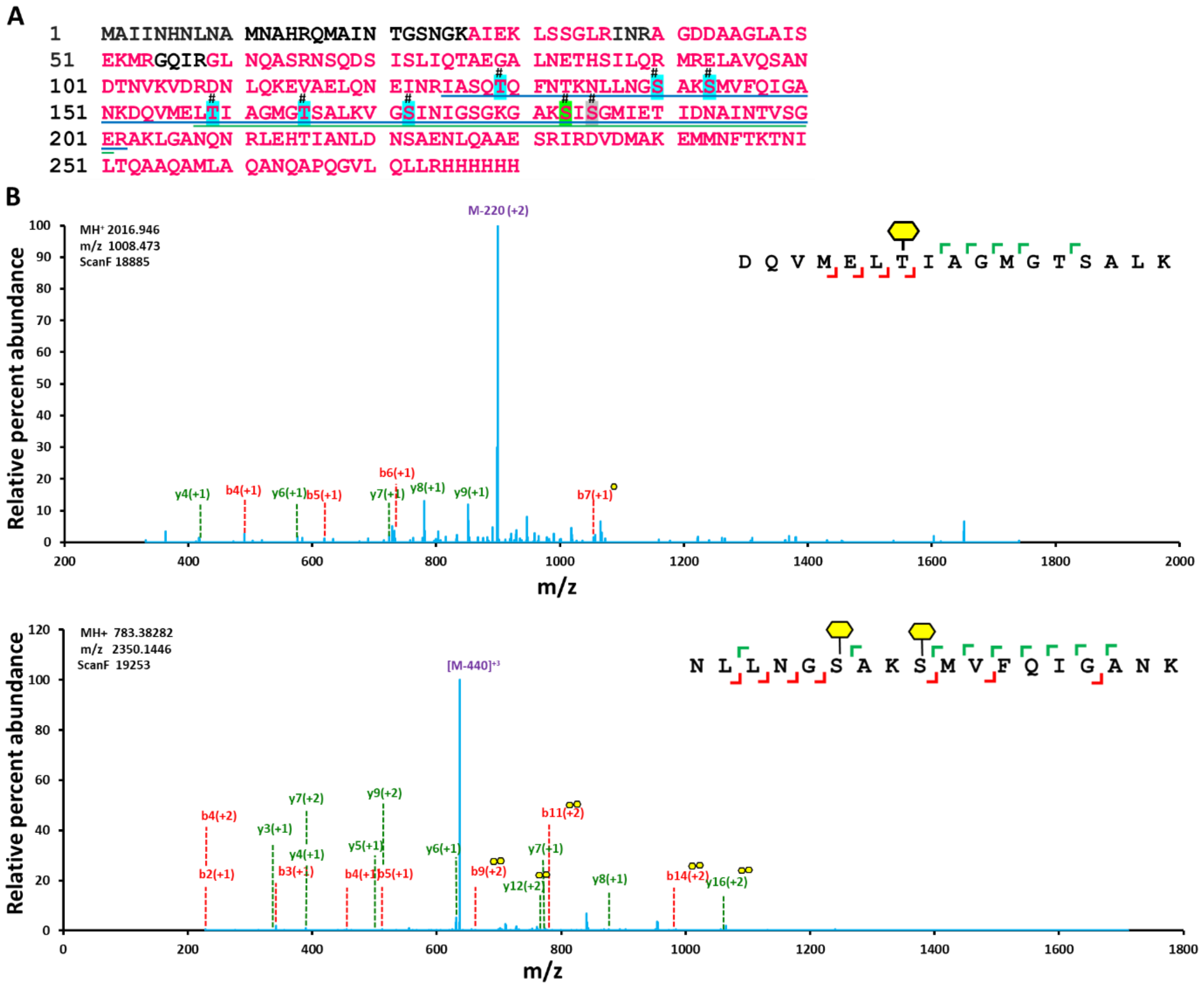

**Supplementary figure 13: Table of all clones and constructs used.**

| Gene locus, symbol, NCBI accession no. | Source | Vector | Restriction sites | Clone | Tag |
|---|---|---|---|---|---|
| ***Cb*****Maf/CLI2780/ WP_003359189.1** | *C. botulinum* F str. Langeland | pETDuet-1 | Nde-I/Xho-I | *Cb*Maf-pETDuet-1 | C-ter S-tag |
| ***Cb*****Maf/CLI2780/ WP_003359189.1** | *C. botulinum* F str. Langeland | pET28a(+) | Nco-I/Xho-I | *Cb*Maf-pET28a(+) | C-ter 6X-His tag |
| ***Cb*****Fla/CLI2781/ WP_003359201.1** | *C. botulinum* F str. Langeland | pET28a(+) | Nco-I/Hind-III | *Cb*Fla-pET28a(+) | C-ter 6X-His tag |
| ***Cb*****Fla/CLI2781/ WP_003359201.1** | *C. botulinum* F str. Langeland | pCDFDuet-1 | Nco-I/Hind-III | *Cb*Fla-pCDFDuet-1 | C-ter 6X-His tag |
| ***Gk*****Maf1/GK3129/ BAD77414.1** | *G. kaustophilus* HTA426 | pCDFDuet-1 | Nco-I/Xho-I | *Gk*Maf1-pCDFDuet-1 | C-ter S-tag |
| ***Gk*****FlaA1/GK3142/ BAD77427.1** | *G. kaustophilus* HTA426 | pET28a(+) | Nco-I/Xho-I | *Gk*FlaA1-pET28a(+) | C-ter 6X-His tag |
| ***Gk*****FlaA2/GK3131/ BAD77416.1** | *G. kaustophilus* HTA426 | pET28a(+) | Nco-I/Xho-I | *Gk*FlaA2-pET28a(+) | C-ter 6X-His tag |
