## Supplementary Data for "Novel Serine/Threonine-O-glycosylation with N-Acetylneuraminic acid and 3-Deoxy-D-manno-octulosonic acid by Maf glycosyltransferases"

**MS/MS data of trypsin digested peptides with site score A and site score B above 19 from *Geobacillus kaustophilus* Flagellin *Gk*F2 protein co-expressed with *Geobacillus kaustophilus* Maf in EV136 cells. Results of dataset 2 (MS/MS results obtained in April 2019).**

| | | | | | |
|---|---|---|---|---|---|
| **Run:** | 77657 | **Scans:** | 6603 - 6603 | **m/z Range:** | 106.12582 to 909.59247 |
| **Search:** | 77657 | **Precursor Mass:** | 462.23936 | | |
| **Mods:** | C=160.03 *+15.99 #+220.06 @+291.1 | **Charge:** | +2 | | |

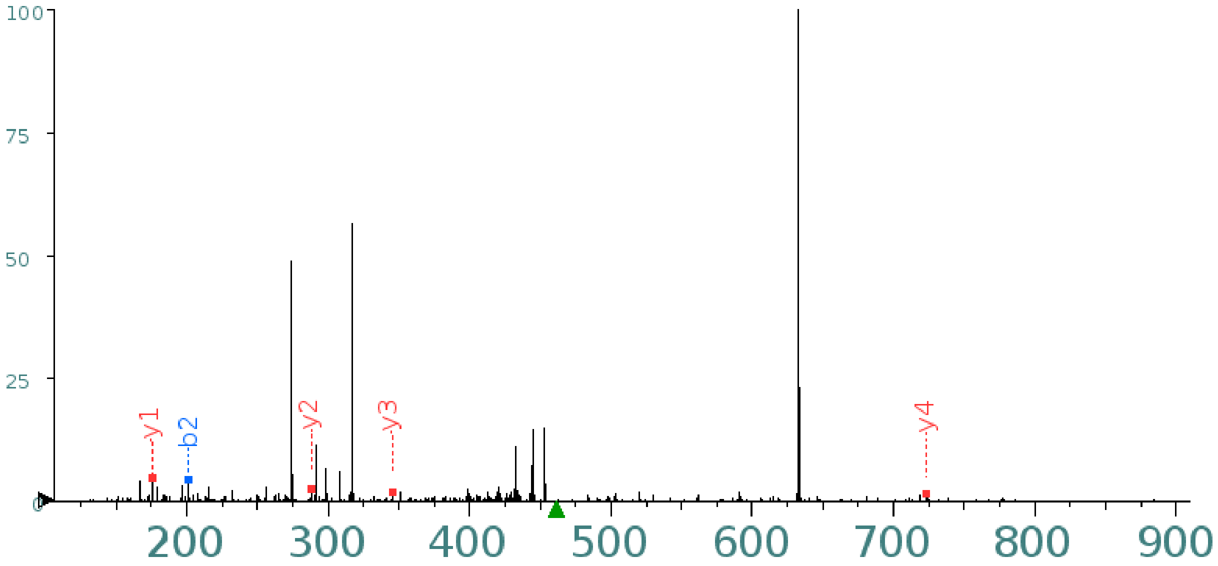

**Sequence**

LSS@GLR

**Predicted Fragmentation Pattern**

| Seq | # | b: Δ Error | b | y | y: Δ Error | +1 |
|---|---|---|---|---|---|---|
| L | 1 | --- | 114.091 | --- | --- | 6 |
| S | 2 | 988.560 | **201.123** | 810.384 | --- | 5 |
| S@ | 3 | --- | 579.251 | **723.352** | 365.562 | 4 |
| G | 4 | --- | 636.272 | **345.224** | 813.956 | 3 |
| L | 5 | --- | 749.356 | **288.203** | 598.821 | 2 |
| R | 6 | --- | --- | **175.119** | 862.432 | 1 |

**Soleil Index**

| | | |
|---|---|---|
| 274.0788 | not explained | 49% |
| 316.7736 | not explained | 57% |
| 632.4745 | not explained | 100% |
| 633.4651 | not explained | 23% |

| | | | | | |
|---|---|---|---|---|---|
| **Run:** | 77657 | **Scans:** | 25346 - 25346 | **m/z Range:** | 303.13318 to 2000 |
| **Search:** | 77657 | **Precursor Mass:** | 1156.89956 | | |
| **Mods:** | C=160.03 *+15.99 #+220.06 @+291.1 | **Charge:** | +3 | | |

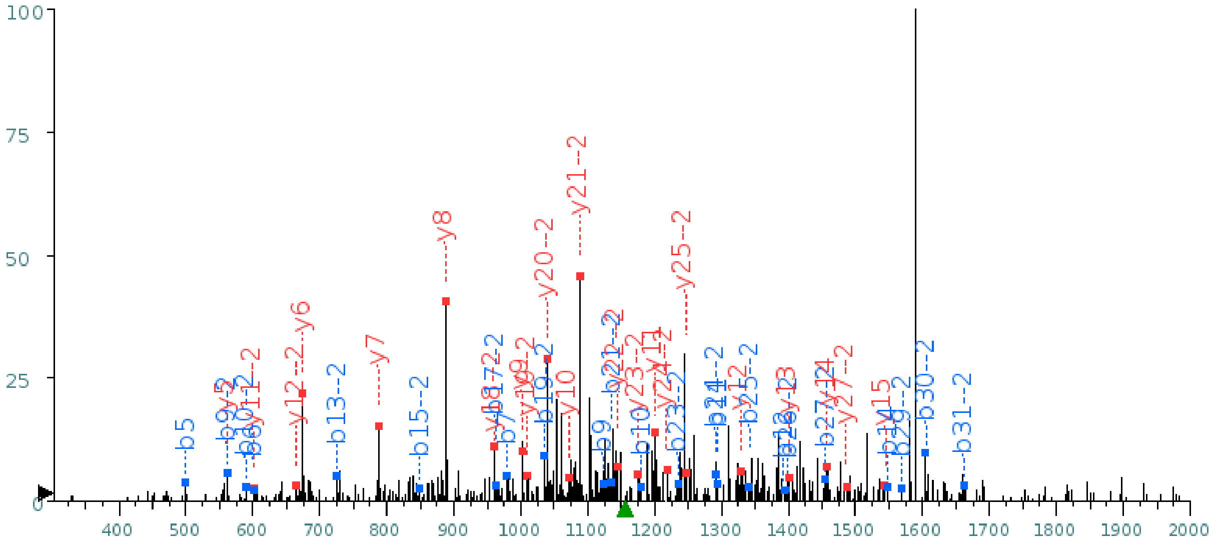

**Sequence**

NIALSTS@GSGNVGVM*TQSGAEQAITIIDDAIK

**Predicted Fragmentation Pattern**

**+1**

| Seq | # | b: Δ Error | b | y | y: Δ Error | +1 |
|---|---|---|---|---|---|---|
| N | 1 | --- | 115.050 | --- | --- | 32 |
| I | 2 | --- | 228.134 | 3354.647 | --- | 31 |
| A | 3 | --- | 299.171 | 3241.563 | --- | 30 |
| L | 4 | --- | 412.255 | 3170.526 | --- | 29 |
| S | 5 | -1651.676 | **499.287** | 3057.442 | --- | 28 |
| T | 6 | 341.374 | **600.335** | 2970.410 | --- | 27 |
| S@ | 7 | -266.006 | **978.463** | 2869.362 | --- | 26 |
| G | 8 | --- | 1035.484 | 2491.234 | --- | 25 |
| S | 9 | 283.333 | **1122.516** | 2434.213 | --- | 24 |
| G | 10 | 675.471 | **1179.538** | 2347.181 | --- | 23 |
| N | 11 | 33.563 | **1293.580** | 2290.160 | --- | 22 |
| V | 12 | 279.644 | **1392.649** | 2176.117 | --- | 21 |
| G | 13 | --- | 1449.670 | 2077.048 | --- | 20 |

| | | | | | | |
|---|---|---|---|---|---|---|
| V | 14 | -7.570 | **1548.739** | 2020.027 | --- | 19 |
| M* | 15 | --- | 1695.774 | 1920.958 | --- | 18 |
| T | 16 | --- | 1796.822 | 1773.923 | --- | 17 |
| Q | 17 | --- | 1924.880 | 1672.875 | --- | 16 |
| S | 18 | --- | 2011.912 | **1544.817** | -512.847 | 15 |
| G | 19 | --- | 2068.934 | **1457.785** | -266.727 | 14 |
| A | 20 | --- | 2139.971 | **1400.763** | 633.790 | 13 |
| E | 21 | --- | 2269.014 | **1329.726** | 10.779 | 12 |
| Q | 22 | --- | 2397.072 | **1200.683** | 22.198 | 11 |
| A | 23 | --- | 2468.109 | **1072.625** | 224.397 | 10 |
| I | 24 | --- | 2581.193 | **1001.588** | -12.158 | 9 |
| T | 25 | --- | 2682.241 | **888.504** | 6.718 | 8 |
| I | 26 | --- | 2795.325 | **787.456** | 136.472 | 7 |
| I | 27 | --- | 2908.409 | **674.372** | -10.475 | 6 |
| D | 28 | --- | 3023.436 | **561.288** | 20.396 | 5 |
| D | 29 | --- | 3138.463 | 446.261 | --- | 4 |
| A | 30 | --- | 3209.500 | 331.234 | --- | 3 |
| I | 31 | --- | 3322.584 | 260.197 | --- | 2 |
| K | 32 | --- | --- | 147.113 | --- | 1 |

+2

| Seq | # | b: Δ Error | b | y | y: Δ Error | +1 |
|---|---|---|---|---|---|---|
| N | 1 | --- | 58.029 | --- | --- | 32 |
| I | 2 | --- | 114.571 | 1677.827 | --- | 31 |
| A | 3 | --- | 150.089 | 1621.285 | --- | 30 |
| L | 4 | --- | 206.631 | 1585.766 | --- | 29 |
| S | 5 | --- | 250.147 | 1529.224 | --- | 28 |
| T | 6 | --- | 300.671 | **1485.708** | 239.984 | 27 |
| S@ | 7 | --- | 489.735 | 1435.185 | --- | 26 |
| G | 8 | --- | 518.246 | **1246.121** | 624.582 | 25 |
| S | 9 | -823.746 | **561.762** | **1217.610** | 574.002 | 24 |
| G | 10 | 815.869 | **590.272** | **1174.094** | 196.668 | 23 |
| N | 11 | --- | 647.294 | **1145.583** | 135.296 | 22 |
| V | 12 | --- | 696.828 | **1088.562** | 674.364 | 21 |
| G | 13 | -1044.447 | **725.339** | **1039.028** | 290.800 | 20 |
| V | 14 | --- | 774.873 | **1010.517** | -33.482 | 19 |
| M* | 15 | 1155.377 | **848.391** | **960.983** | 600.356 | 18 |
| T | 16 | --- | 898.915 | 887.465 | --- | 17 |
| Q | 17 | -370.305 | **962.944** | 836.941 | --- | 16 |
| S | 18 | --- | 1006.460 | 772.912 | --- | 15 |
| G | 19 | -700.279 | **1034.971** | 729.396 | --- | 14 |
| A | 20 | --- | 1070.489 | 700.885 | --- | 13 |
| E | 21 | -507.531 | **1135.010** | **665.367** | 124.997 | 12 |
| Q | 22 | --- | 1199.040 | **600.845** | 976.239 | 11 |
| A | 23 | 206.455 | **1234.558** | 536.816 | --- | 10 |
| I | 24 | -581.229 | **1291.100** | 501.298 | --- | 9 |
| T | 25 | -91.923 | **1341.624** | 444.755 | --- | 8 |
| I | 26 | -545.928 | **1398.166** | 394.232 | --- | 7 |
| I | 27 | 466.860 | **1454.708** | 337.690 | --- | 6 |
| D | 28 | --- | 1512.222 | 281.148 | --- | 5 |
| D | 29 | -419.762 | **1569.735** | 223.634 | --- | 4 |
| A | 30 | -595.369 | **1605.254** | 166.121 | --- | 3 |
| I | 31 | 567.873 | **1661.796** | 130.602 | --- | 2 |
| K | 32 | --- | --- | 74.060 | --- | 1 |

**Soleil Index**

| | | |
|---|---|---|
| 1040.2855 | not explained | 21% |
| 1054.2715 | b2-20 frag loss of ammonia/ y1-10 frag loss of water | 21% |
| 1060.3091 | not explained | 18% |
| 1101.6783 | not explained | 21% |
| 1102.7037 | not explained | 19% |
| 1243.8939 | not explained | 30% |
| 1310.4839 | not explained | 15% |
| 1581.2462 | not explained | 16% |
| 1589.4246 | not explained | 33% |
| 1590.4373 | not explained | 100% |

| | | | | | |
|---|---|---|---|---|---|
| **Run:** | 77657 | **Scans:** | 26388 - 26388 | **m/z Range:** | 231.17789 to 1983.2055 |
| **Search:** | 77657 | **Precursor Mass:** | 863.93006 | | |
| **Mods:** | C=160.03 *+15.99 #+220.06 @+291.1 | **Charge:** | +4 | | |

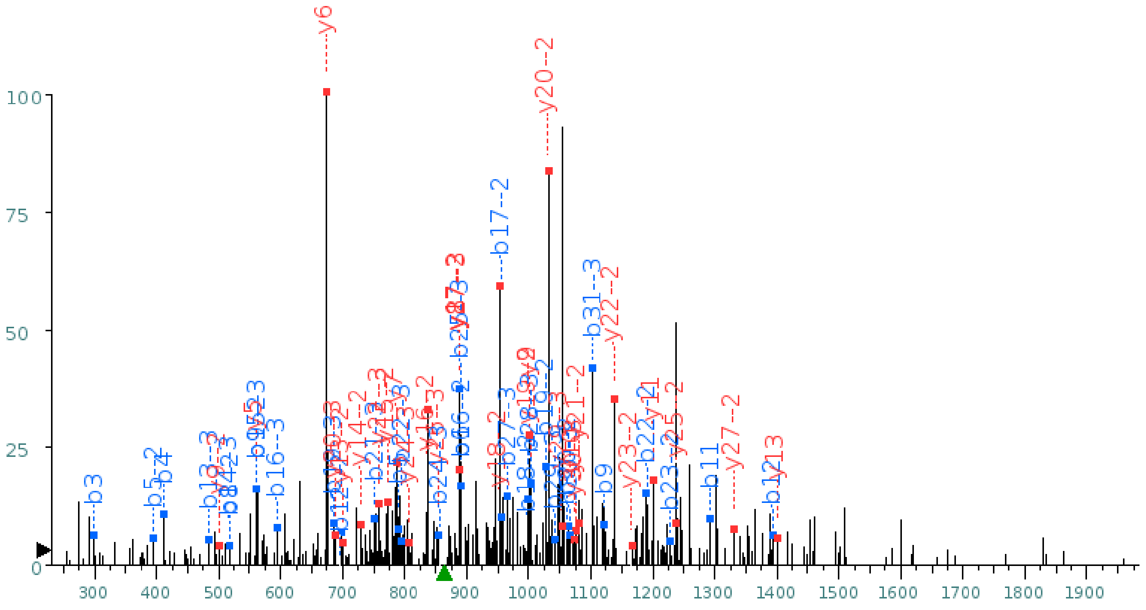

**Sequence**

NIALS@TSGSGNVGVMTQSGAEQAITIIDDAIK

**Predicted Fragmentation Pattern**

| Seq | # | b: Δ Error | b +1 | y +1 | y: Δ Error | +1 |
|---|---|---|---|---|---|---|
| N | 1 | --- | 115.050 | --- | --- | 32 |
| I | 2 | --- | 228.134 | 3338.652 | --- | 31 |
| A | 3 | 1271.244 | **299.171** | 3225.568 | --- | 30 |
| L | 4 | 322.025 | **412.255** | 3154.531 | --- | 29 |
| S@ | 5 | -847.872 | **790.383** | 3041.447 | --- | 28 |
| T | 6 | -363.872 | **891.431** | 2663.319 | --- | 27 |
| S | 7 | --- | 978.463 | 2562.272 | --- | 26 |
| G | 8 | --- | 1035.484 | 2475.240 | --- | 25 |
| S | 9 | -645.812 | **1122.516** | 2418.218 | --- | 24 |
| G | 10 | --- | 1179.538 | 2331.186 | --- | 23 |
| N | 11 | -34.192 | **1293.580** | 2274.165 | --- | 22 |
| V | 12 | 493.528 | **1392.649** | 2160.122 | --- | 21 |
| G | 13 | --- | 1449.670 | 2061.053 | --- | 20 |
| V | 14 | --- | 1548.739 | 2004.032 | --- | 19 |

| | | | | | | |
|---|---|---|---|---|---|---|
| M | 15 | --- | 1679.779 | 1904.963 | --- | 18 |
| T | 16 | --- | 1780.827 | 1773.923 | --- | 17 |
| Q | 17 | --- | 1908.886 | 1672.875 | --- | 16 |
| S | 18 | --- | 1995.918 | 1544.817 | --- | 15 |
| G | 19 | --- | 2052.939 | 1457.785 | --- | 14 |
| A | 20 | --- | 2123.976 | **1400.763** | **-473.740** | 13 |
| E | 21 | --- | 2253.019 | 1329.726 | --- | 12 |
| Q | 22 | --- | 2381.077 | **1200.683** | **336.440** | 11 |
| A | 23 | --- | 2452.114 | **1072.625** | **353.605** | 10 |
| I | 24 | --- | 2565.198 | **1001.588** | **148.515** | 9 |
| T | 25 | --- | 2666.246 | **888.504** | **204.996** | 8 |
| I | 26 | --- | 2779.330 | **787.456** | **-628.068** | 7 |
| I | 27 | --- | 2892.414 | **674.372** | **137.937** | 6 |
| D | 28 | --- | 3007.441 | **561.288** | **139.013** | 5 |
| D | 29 | --- | 3122.468 | 446.261 | --- | 4 |
| A | 30 | --- | 3193.505 | 331.234 | --- | 3 |
| I | 31 | --- | 3306.589 | 260.197 | --- | 2 |
| K | 32 | --- | --- | 147.113 | --- | 1 |

+2

| Seq | # | b: Δ Error | b | y | y: Δ Error | +1 |
|---|---|---|---|---|---|---|
| N | 1 | --- | 58.029 | --- | --- | 32 |
| I | 2 | --- | 114.571 | 1669.830 | --- | 31 |
| A | 3 | --- | 150.089 | 1613.288 | --- | 30 |
| L | 4 | --- | 206.631 | 1577.769 | --- | 29 |
| S@ | 5 | **-650.311** | **395.695** | 1521.227 | --- | 28 |
| T | 6 | --- | 446.219 | **1332.163** | **-35.597** | 27 |
| S | 7 | --- | 489.735 | 1281.639 | --- | 26 |
| G | 8 | **-1858.018** | **518.246** | **1238.123** | **142.153** | 25 |
| S | 9 | **-705.028** | **561.762** | 1209.613 | --- | 24 |
| G | 10 | --- | 590.272 | **1166.097** | **301.618** | 23 |
| N | 11 | --- | 647.294 | **1137.586** | **429.368** | 22 |
| V | 12 | **1257.490** | **696.828** | **1080.564** | **390.593** | 21 |
| G | 13 | --- | 725.339 | **1031.030** | **423.809** | 20 |
| V | 14 | --- | 774.873 | **1002.520** | **-781.661** | 19 |
| M | 15 | --- | 840.393 | **952.985** | **593.387** | 18 |
| T | 16 | **212.338** | **890.917** | **887.465** | **493.017** | 17 |
| Q | 17 | **-169.771** | **954.946** | **836.941** | **889.400** | 16 |
| S | 18 | **343.115** | **998.462** | **772.912** | **163.561** | 15 |
| G | 19 | **38.870** | **1026.973** | **729.396** | **893.224** | 14 |
| A | 20 | **823.203** | **1062.492** | **700.885** | **1045.971** | 13 |
| E | 21 | --- | 1127.013 | 665.367 | --- | 12 |
| Q | 22 | **-739.234** | **1191.042** | 600.845 | --- | 11 |
| A | 23 | **530.928** | **1226.561** | 536.816 | --- | 10 |
| I | 24 | --- | 1283.103 | **501.298** | **-1986.264** | 9 |
| T | 25 | --- | 1333.627 | 444.755 | --- | 8 |
| I | 26 | --- | 1390.169 | 394.232 | --- | 7 |
| I | 27 | --- | 1446.711 | 337.690 | --- | 6 |
| D | 28 | --- | 1504.224 | 281.148 | --- | 5 |
| D | 29 | --- | 1561.738 | 223.634 | --- | 4 |
| A | 30 | --- | 1597.256 | 166.121 | --- | 3 |
| I | 31 | --- | 1653.798 | 130.602 | --- | 2 |
| K | 32 | --- | --- | 74.060 | --- | 1 |

+3

| Seq | # | b: Δ Error | b | y | y: Δ Error | +1 |
|---|---|---|---|---|---|---|
| N | 1 | --- | 39.022 | --- | --- | 32 |
| I | 2 | --- | 76.716 | 1113.555 | --- | 31 |

| | | | | | | |
|---|---|---|---|---|---|---|
| A | 3 | --- | 100.395 | **1075.861** | **-255.102** | **30** |
| L | 4 | --- | 138.090 | **1052.182** | **856.965** | **29** |
| S@ | 5 | --- | 264.132 | 1014.487 | --- | **28** |
| T | 6 | --- | 297.815 | **888.445** | **271.469** | **27** |
| S | 7 | --- | 326.826 | **854.762** | **703.619** | **26** |
| G | 8 | --- | 345.833 | 825.751 | --- | **25** |
| S | 9 | --- | 374.844 | **806.744** | **-378.587** | **24** |
| G | 10 | --- | 393.851 | 777.734 | --- | **23** |
| N | 11 | --- | 431.865 | **758.726** | **-149.972** | **22** |
| V | 12 | --- | 464.888 | 720.712 | --- | **21** |
| G | 13 | **1094.493** | **483.895** | **687.689** | **70.826** | **20** |
| V | 14 | **709.030** | **516.918** | 668.682 | --- | **19** |
| M | 15 | **1368.035** | **560.598** | 635.659 | --- | **18** |
| T | 16 | **-427.603** | **594.280** | 591.979 | --- | **17** |
| Q | 17 | --- | 636.967 | 558.297 | --- | **16** |
| S | 18 | --- | 665.977 | 515.610 | --- | **15** |
| G | 19 | **1038.468** | **684.985** | 486.600 | --- | **14** |
| A | 20 | --- | 708.664 | 467.593 | --- | **13** |
| E | 21 | **-621.096** | **751.678** | 443.914 | --- | **12** |
| Q | 22 | **155.496** | **794.364** | 400.899 | --- | **11** |
| A | 23 | --- | 818.043 | 358.213 | --- | **10** |
| I | 24 | **-436.987** | **855.738** | 334.534 | --- | **9** |
| T | 25 | **-826.368** | **889.420** | 296.839 | --- | **8** |
| I | 26 | --- | 927.115 | 263.157 | --- | **7** |
| I | 27 | **47.656** | **964.810** | 225.462 | --- | **6** |
| D | 28 | **86.261** | **1003.152** | 187.767 | --- | **5** |
| D | 29 | **298.109** | **1041.494** | 149.425 | --- | **4** |
| A | 30 | **609.940** | **1065.173** | 111.083 | --- | **3** |
| I | 31 | **-820.663** | **1102.868** | 87.404 | --- | **2** |
| K | 32 | --- | --- | 49.709 | --- | **1** |

**Soleil Index**

| | | |
|---|---|---|
| **629.7004** | **b2-11 frag loss of water/ b2-11 frag loss of ammonia** | **18%** |
| **675.5515** | **not explained** | **25%** |
| **784.6425** | **not explained** | **17%** |
| **914.2484** | **not explained** | **18%** |
| **945.2249** | **not explained** | **23%** |
| **1047.3364** | **b3-30 frag loss of water/ b3-30 frag loss of ammonia** | **17%** |
| **1048.6769** | **b3-30 frag loss of ammonia** | **23%** |
| **1049.4955** | **not explained** | **20%** |
| **1055.3864** | **y1-10 frag loss of water/ y1-10 frag loss of ammonia** | **93%** |
| **1236.4054** | **not explained** | **52%** |
| **1257.8882** | **not explained** | **21%** |
| **1301.4017** | **not explained** | **19%** |

| | | | | | | |
|---|---|---|---|---|---|---|
| **Run:** | 77657 | **Scans:** | 25023 - 25023 | **m/z Range:** | 219.27527 | to 1988.3891( |
| **Search:** | 77657 | **Precursor Mass:** | 867.92528 | | | |
| **Mods:** | C=160.03 *+15.99 #+220.06 @+291.1 | **Charge:** | +4 | | | |

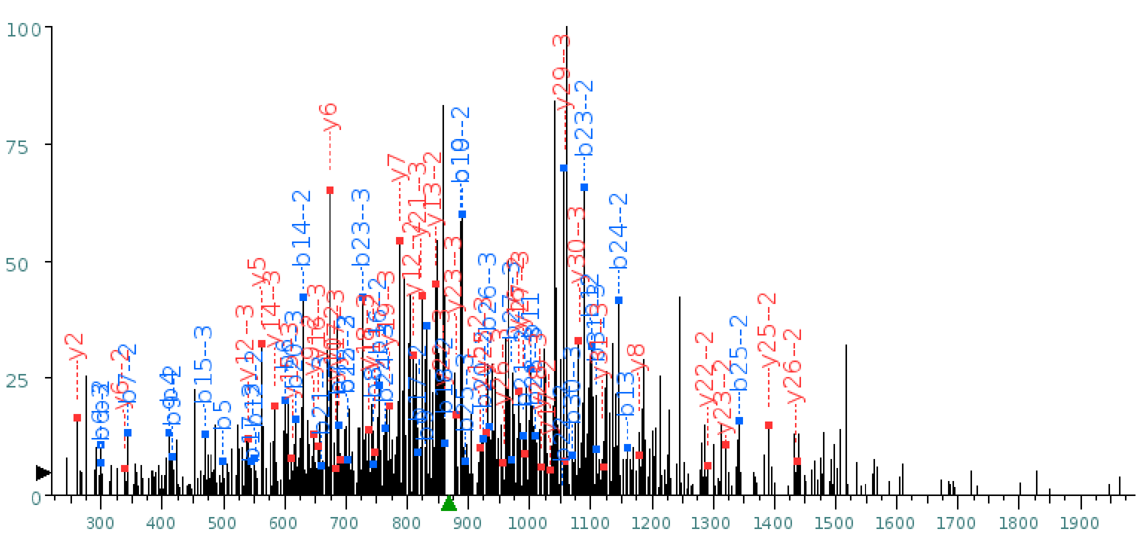

**Sequence**

NIALSTSGSGNVGVM*TQSGAEQAIT@IIDDAIK

**Predicted Fragmentation Pattern**

| | | | +1 | | | |
|---|---|---|---|---|---|---|
| **Seq** | **#** | **b: Δ Error** | **b** | **y** | **y: Δ Error** | **+1** |
| N | 1 | --- | 115.050 | --- | --- | 32 |
| I | 2 | --- | 228.134 | 3354.647 | --- | 31 |
| A | 3 | -73.840 | **299.171** | 3241.563 | --- | 30 |
| L | 4 | -12.246 | **412.255** | 3170.526 | --- | 29 |
| S | 5 | 298.803 | **499.287** | 3057.442 | --- | 28 |
| T | 6 | 356.106 | **600.335** | 2970.410 | --- | 27 |
| S | 7 | 154.846 | **687.367** | 2869.362 | --- | 26 |
| G | 8 | 321.677 | **744.389** | 2782.330 | --- | 25 |
| S | 9 | -147.232 | **831.421** | 2725.308 | --- | 24 |
| G | 10 | 1108.880 | **888.442** | 2638.276 | --- | 23 |
| N | 11 | -738.231 | **1002.485** | 2581.255 | --- | 22 |
| V | 12 | 37.112 | **1101.553** | 2467.212 | --- | 21 |
| G | 13 | 455.494 | **1158.575** | 2368.144 | --- | 20 |
| V | 14 | --- | 1257.643 | 2311.122 | --- | 19 |
| M* | 15 | --- | 1404.679 | 2212.054 | --- | 18 |
| T | 16 | --- | 1505.726 | 2065.018 | --- | 17 |
| Q | 17 | --- | 1633.785 | 1963.971 | --- | 16 |

| | | | | | | |
|---|---|---|---|---|---|---|
| S | 18 | --- | 1720.817 | 1835.912 | --- | 15 |
| G | 19 | --- | 1777.839 | 1748.880 | --- | 14 |
| A | 20 | --- | 1848.876 | 1691.859 | --- | 13 |
| E | 21 | --- | 1977.918 | 1620.821 | --- | 12 |
| Q | 22 | --- | 2105.977 | 1491.779 | --- | 11 |
| A | 23 | --- | 2177.014 | 1363.720 | --- | 10 |
| I | 24 | --- | 2290.098 | 1292.683 | --- | 9 |
| T@ | 25 | --- | 2682.241 | 1179.599 | 202.087 | 8 |
| I | 26 | --- | 2795.325 | 787.456 | 65.643 | 7 |
| I | 27 | --- | 2908.409 | 674.372 | 107.353 | 6 |
| D | 28 | --- | 3023.436 | 561.288 | 155.863 | 5 |
| D | 29 | --- | 3138.463 | 446.261 | --- | 4 |
| A | 30 | --- | 3209.500 | 331.234 | --- | 3 |
| I | 31 | --- | 3322.584 | 260.197 | 30.612 | 2 |
| K | 32 | --- | --- | 147.113 | --- | 1 |

+2

| Seq | # | b: Δ Error | b | y | y: Δ Error | +1 |
|---|---|---|---|---|---|---|
| N | 1 | --- | 58.029 | --- | --- | 32 |
| I | 2 | --- | 114.571 | 1677.827 | --- | 31 |
| A | 3 | --- | 150.089 | 1621.285 | --- | 30 |
| L | 4 | --- | 206.631 | 1585.766 | --- | 29 |
| S | 5 | --- | 250.147 | 1529.224 | --- | 28 |
| T | 6 | -913.438 | 300.671 | 1485.708 | --- | 27 |
| S | 7 | 74.376 | 344.187 | 1435.185 | 695.850 | 26 |
| G | 8 | --- | 372.698 | 1391.669 | -104.742 | 25 |
| S | 9 | 665.502 | 416.214 | 1363.158 | --- | 24 |
| G | 10 | --- | 444.725 | 1319.642 | 723.412 | 23 |
| N | 11 | --- | 501.746 | 1291.131 | 121.479 | 22 |
| V | 12 | -1786.554 | 551.280 | 1234.110 | --- | 21 |
| G | 13 | --- | 579.791 | 1184.575 | --- | 20 |
| V | 14 | 319.951 | 629.325 | 1156.065 | --- | 19 |
| M* | 15 | 892.186 | 702.843 | 1106.530 | --- | 18 |
| T | 16 | 214.037 | 753.367 | 1033.013 | 526.547 | 17 |
| Q | 17 | 127.564 | 817.396 | 982.489 | -456.252 | 16 |
| S | 18 | 88.344 | 860.912 | 918.460 | 703.642 | 15 |
| G | 19 | 6.202 | 889.423 | 874.944 | --- | 14 |
| A | 20 | 199.069 | 924.941 | 846.433 | -37.130 | 13 |
| E | 21 | -526.611 | 989.463 | 810.914 | -153.333 | 12 |
| Q | 22 | 799.952 | 1053.492 | 746.393 | 642.903 | 11 |
| A | 23 | 80.729 | 1089.011 | 682.364 | 1336.951 | 10 |
| I | 24 | 209.446 | 1145.553 | 646.845 | -206.030 | 9 |
| T@ | 25 | -172.923 | 1341.624 | 590.303 | --- | 8 |
| I | 26 | --- | 1398.166 | 394.232 | --- | 7 |
| I | 27 | --- | 1454.708 | 337.690 | -1178.111 | 6 |
| D | 28 | --- | 1512.222 | 281.148 | --- | 5 |
| D | 29 | --- | 1569.735 | 223.634 | --- | 4 |
| A | 30 | --- | 1605.254 | 166.121 | --- | 3 |
| I | 31 | --- | 1661.796 | 130.602 | --- | 2 |
| K | 32 | --- | --- | 74.060 | --- | 1 |

+3

| Seq | # | b: Δ Error | b | y | y: Δ Error | +1 |
|---|---|---|---|---|---|---|
| N | 1 | --- | 39.022 | --- | --- | 32 |
| I | 2 | --- | 76.716 | 1118.887 | 714.047 | 31 |
| A | 3 | --- | 100.395 | 1081.192 | -495.048 | 30 |
| L | 4 | --- | 138.090 | 1057.513 | -707.848 | 29 |
| S | 5 | --- | 167.101 | 1019.819 | -289.929 | 28 |

| | | | | | | |
|---|---|---|---|---|---|---|
| T | 6 | --- | 200.783 | **990.808** | 392.738 | 27 |
| S | 7 | --- | 229.794 | **957.125** | -675.835 | 26 |
| G | 8 | --- | 248.801 | **928.115** | -315.500 | 25 |
| S | 9 | --- | 277.812 | 909.108 | --- | 24 |
| G | 10 | --- | 296.819 | **880.097** | 909.749 | 23 |
| N | 11 | --- | 334.833 | **861.090** | -118.010 | 22 |
| V | 12 | --- | 367.856 | **823.076** | 851.741 | 21 |
| G | 13 | --- | 386.863 | 790.053 | --- | 20 |
| V | 14 | --- | 419.886 | **771.046** | -462.296 | 19 |
| M* | 15 | 978.480 | **468.898** | **738.023** | -985.754 | 18 |
| T | 16 | --- | 502.580 | **689.011** | 870.777 | 17 |
| Q | 17 | -1550.332 | **545.267** | **655.328** | -1054.710 | 16 |
| S | 18 | --- | 574.277 | **612.642** | -1516.894 | 15 |
| G | 19 | --- | 593.284 | **583.632** | -265.270 | 14 |
| A | 20 | 531.440 | **616.963** | 564.624 | --- | 13 |
| E | 21 | -595.818 | **659.978** | **540.945** | -543.949 | 12 |
| Q | 22 | 1146.974 | **702.664** | 497.931 | --- | 11 |
| A | 23 | -116.541 | **726.343** | 455.245 | --- | 10 |
| I | 24 | 1014.349 | **764.038** | 431.566 | --- | 9 |
| T@ | 25 | 143.389 | **894.752** | 393.871 | --- | 8 |
| I | 26 | 218.158 | **932.447** | 263.157 | --- | 7 |
| I | 27 | -518.627 | **970.141** | 225.462 | --- | 6 |
| D | 28 | 430.627 | **1008.484** | 187.767 | --- | 5 |
| D | 29 | --- | 1046.826 | 149.425 | --- | 4 |
| A | 30 | -107.686 | **1070.505** | 111.083 | --- | 3 |
| I | 31 | 763.085 | **1108.200** | 87.404 | --- | 2 |
| K | 32 | --- | --- | 49.709 | --- | 1 |

**Soleil Index**

| | | |
|---|---|---|
| 274.1422 | not explained | 25% |
| 452.3267 | b3-15 frag loss of ammonia | 23% |
| 562.3023 | b2-13 frag loss of water/ b2-13 frag loss of ammonia | 17% |
| 606.3790 | not explained | 20% |
| 626.7971 | not explained | 19% |
| 637.4421 | y3-16 frag loss of water/ y3-16 frag loss of ammonia | 16% |
| 657.4978 | y1-6 frag loss of ammonia | 16% |
| 667.4551 | not explained | 17% |
| 669.7708 | b1-7 frag loss of water/ b1-7 frag loss of ammonia | 31% |
| 675.4809 | not explained | 27% |
| 686.2418 | b2-15 frag loss of ammonia/ b3-22 frag loss of ammonia | 38% |
| 705.0533 | not explained | 19% |
| 741.7830 | not explained | 19% |
| 755.0482 | not explained | 25% |
| 766.2332 | not explained | 20% |
| 793.4726 | y2-12 frag loss of water/ y2-12 frag loss of ammonia/ y2-29 frag loss of water | 22% |
| 795.4584 | not explained | 46% |
| 803.5654 | b2-30 frag doubly charged | 32% |
| 805.8470 | y3-21 frag loss of water/ y3-21 frag loss of ammonia | 43% |

| | | |
|---|---|---|
| 814.7423 | b1-9 frag loss of ammonia | 28% |
| 824.5230 | not explained | 43% |
| 830.3618 | b2-31 frag doubly charged/ y2-13 frag loss of ammonia | 29% |
| 837.6227 | not explained | 27% |
| 841.1799 | not explained | 22% |
| 845.3415 | y1-13 frag doubly charged | 34% |
| 849.8052 | not explained | 54% |
| 850.6237 | not explained | 18% |
| 851.5041 | not explained | 30% |
| 852.4193 | not explained | 26% |
| 854.5467 | not explained | 20% |
| 855.5198 | not explained | 22% |
| 856.2251 | y2-14 frag loss of water | 21% |
| 857.0692 | y2-14 frag loss of water/ y2-14 frag loss of ammonia | 25% |
| 858.6916 | precursor loss of two waters/ y2-14 frag loss of ammonia | 83% |
| 859.6507 | precursor loss of two waters/ b1-18 frag loss of water | 43% |
| 878.3773 | b3-25 frag loss of ammonia | 26% |
| 905.3765 | not explained | 17% |
| 931.0023 | not explained | 28% |
| 937.5279 | not explained | 28% |
| 944.1161 | not explained | 25% |
| 961.6242 | not explained | 34% |
| 962.5018 | not explained | 23% |
| 965.5403 | y2-16 frag loss of ammonia | 27% |
| 966.3724 | y2-16 frag loss of ammonia | 50% |
| 967.4504 | not explained | 21% |
| 971.5252 | b2-21 frag loss of water/ b2-21 frag loss of ammonia | 26% |
| 973.6291 | y3-27 frag loss of water/ y3-27 frag loss of ammonia | 16% |
| 974.8710 | not explained | 17% |
| 983.7057 | b1-11 frag loss of water | 15% |
| 993.8643 | not explained | 23% |
| 994.6129 | not explained | 21% |
| 999.2581 | not explained | 16% |
| 1023.6866 | not explained | 31% |
| 1025.2372 | not explained | 26% |
| 1038.8120 | y3-29 frag loss of water | 19% |
| 1040.1144 | y3-29 frag loss of water/ y3-29 frag loss of ammonia | 84% |
| 1043.5680 | not explained | 44% |
| 1049.2002 | not explained | 29% |
| 1055.5574 | not explained | 23% |
| 1059.9237 | not explained | 29% |

| | | |
|---|---|---|
| 1060.7123 | not explained | 100% |
| 1074.9358 | not explained | 15% |
| 1087.7041 | b1-23 frag doubly charged/ y2-18 frag loss of water | 20% |
| 1090.2723 | b3-31 frag loss of water/ b3-31 frag loss of ammonia/ y2-18 frag loss of ammonia | 41% |
| 1097.7365 | not explained | 23% |
| 1098.6813 | not explained | 30% |
| 1099.8834 | y3-31 frag loss of water | 23% |
| 1103.7490 | not explained | 21% |
| 1109.7006 | not explained | 21% |
| 1122.3098 | not explained | 23% |
| 1124.8276 | not explained | 26% |
| 1131.7750 | not explained | 17% |
| 1136.3926 | not explained | 32% |
| 1140.2216 | b1-13 frag loss of water | 22% |
| 1183.3270 | y1-20 frag doubly charged | 19% |
| 1187.5294 | not explained | 29% |
| 1212.0964 | not explained | 25% |
| 1228.0326 | not explained | 16% |
| 1228.8623 | not explained | 18% |
| 1244.0470 | not explained | 42% |
| 1245.0548 | not explained | 42% |
| 1517.0472 | not explained | 32% |

**MS/MS data of trypsin digested peptides with site score A and site score B above 19 from *Geobacillus kaustophilus* Flagellin *Gk*F2 protein co-expressed with *Geobacillus kaustophilus* Maf in EV136 cells. Results of dataset 3 (MS/MS results obtained in July 2019).**

| Run: | 79491 | Scans: | 29941 - 29941 | m/z Range: | 316.03101 to 2000 |
|---|---|---|---|---|---|
| Search: | 79491 | Precursor Mass: | 1156.90095 | | |
| Mods: | C=160.03 *+15.99 #+220.06 @+291.1 | | | | |

**Sequence**

NIALSTSGSGNVGVM*TQSGAEQAIT@IIDDAIK

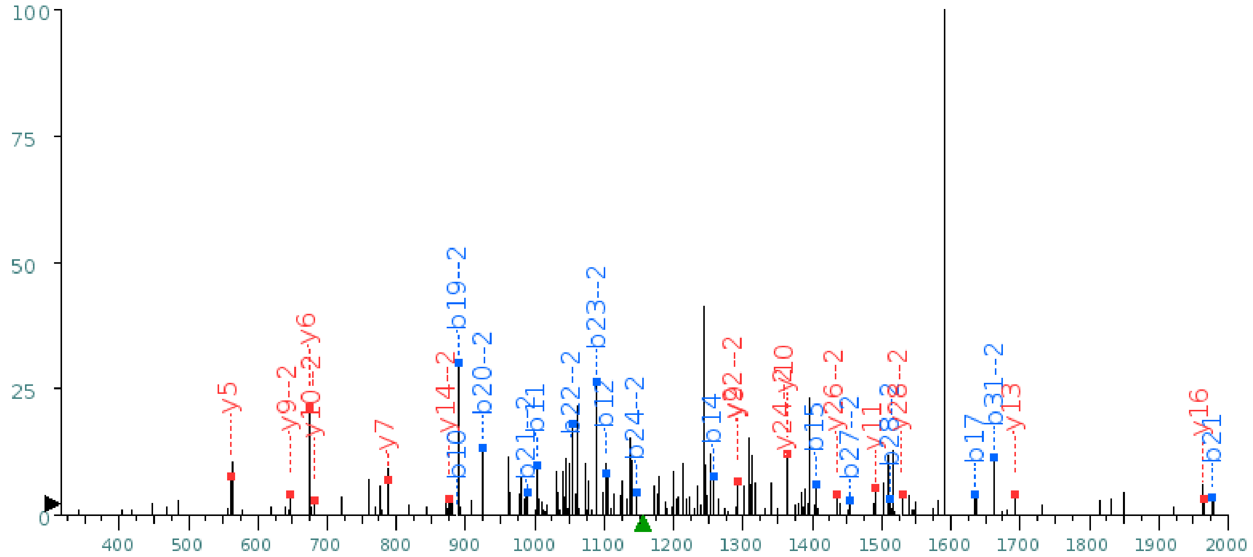

**Predicted Fragmentation Pattern**

+1

| Seq | # | b: Δ Error | b | y | y: Δ Error | +1 |
|---|---|---|---|---|---|---|
| N | 1 | --- | 115.050 | --- | --- | 32 |
| I | 2 | --- | 228.134 | 3354.647 | --- | 31 |
| A | 3 | --- | 299.171 | 3241.563 | --- | 30 |
| L | 4 | --- | 412.255 | 3170.526 | --- | 29 |
| S | 5 | --- | 499.287 | 3057.442 | --- | 28 |
| T | 6 | --- | 600.335 | 2970.410 | --- | 27 |
| S | 7 | --- | 687.367 | 2869.362 | --- | 26 |
| G | 8 | --- | 744.389 | 2782.330 | --- | 25 |
| S | 9 | --- | 831.421 | 2725.308 | --- | 24 |
| G | 10 | 245.473 | **888.442** | 2638.276 | --- | 23 |
| N | 11 | 223.194 | **1002.485** | 2581.255 | --- | 22 |
| V | 12 | 84.204 | **1101.553** | 2467.212 | --- | 21 |
| G | 13 | --- | 1158.575 | 2368.144 | --- | 20 |
| V | 14 | 186.775 | **1257.643** | 2311.122 | --- | 19 |

| | | | | | | |
|---|---|---|---|---|---|---|
| M* | 15 | 30.992 | **1404.679** | 2212.054 | --- | 18 |
| T | 16 | --- | 1505.726 | 2065.018 | --- | 17 |
| Q | 17 | 243.680 | **1633.785** | **1963.971** | 170.128 | 16 |
| S | 18 | --- | 1720.817 | 1835.912 | --- | 15 |
| G | 19 | --- | 1777.839 | 1748.880 | --- | 14 |
| A | 20 | --- | 1848.876 | **1691.859** | 23.066 | 13 |
| E | 21 | 29.625 | **1977.918** | 1620.821 | --- | 12 |
| Q | 22 | --- | 2105.977 | **1491.779** | -585.773 | 11 |
| A | 23 | --- | 2177.014 | **1363.720** | 177.430 | 10 |
| I | 24 | --- | 2290.098 | **1292.683** | -764.767 | 9 |
| T@ | 25 | --- | 2682.241 | 1179.599 | --- | 8 |
| I | 26 | --- | 2795.325 | **787.456** | 189.781 | 7 |
| I | 27 | --- | 2908.409 | **674.372** | 113.325 | 6 |
| D | 28 | --- | 3023.436 | **561.288** | 11.044 | 5 |
| D | 29 | --- | 3138.463 | 446.261 | --- | 4 |
| A | 30 | --- | 3209.500 | 331.234 | --- | 3 |
| I | 31 | --- | 3322.584 | 260.197 | --- | 2 |
| K | 32 | --- | --- | 147.113 | --- | 1 |

+2

| Seq | # | b: Δ Error | b | y | y: Δ Error | +1 |
|---|---|---|---|---|---|---|
| N | 1 | --- | 58.029 | --- | --- | 32 |
| I | 2 | --- | 114.571 | 1677.827 | --- | 31 |
| A | 3 | --- | 150.089 | 1621.285 | --- | 30 |
| L | 4 | --- | 206.631 | 1585.766 | --- | 29 |
| S | 5 | --- | 250.147 | **1529.224** | 480.981 | 28 |
| T | 6 | --- | 300.671 | 1485.708 | --- | 27 |
| S | 7 | --- | 344.187 | **1435.185** | 168.792 | 26 |
| G | 8 | --- | 372.698 | 1391.669 | --- | 25 |
| S | 9 | --- | 416.214 | **1363.158** | 589.778 | 24 |
| G | 10 | --- | 444.725 | 1319.642 | --- | 23 |
| N | 11 | --- | 501.746 | **1291.131** | 436.800 | 22 |
| V | 12 | --- | 551.280 | 1234.110 | --- | 21 |
| G | 13 | --- | 579.791 | 1184.575 | --- | 20 |
| V | 14 | --- | 629.325 | 1156.065 | --- | 19 |
| M* | 15 | --- | 702.843 | 1106.530 | --- | 18 |
| T | 16 | --- | 753.367 | 1033.013 | --- | 17 |
| Q | 17 | --- | 817.396 | 982.489 | --- | 16 |
| S | 18 | --- | 860.912 | 918.460 | --- | 15 |
| G | 19 | -858.158 | **889.423** | **874.944** | -185.784 | 14 |
| A | 20 | 549.863 | **924.941** | 846.433 | --- | 13 |
| E | 21 | 227.839 | **989.463** | 810.914 | --- | 12 |
| Q | 22 | 504.401 | **1053.492** | 746.393 | --- | 11 |
| A | 23 | 17.067 | **1089.011** | **682.364** | 322.166 | 10 |
| I | 24 | 590.520 | **1145.553** | **646.845** | -623.345 | 9 |
| T@ | 25 | --- | 1341.624 | 590.303 | --- | 8 |
| I | 26 | --- | 1398.166 | 394.232 | --- | 7 |
| I | 27 | -532.033 | **1454.708** | 337.690 | --- | 6 |
| D | 28 | 472.148 | **1512.222** | 281.148 | --- | 5 |
| D | 29 | --- | 1569.735 | 223.634 | --- | 4 |
| A | 30 | --- | 1605.254 | 166.121 | --- | 3 |
| I | 31 | 412.003 | **1661.796** | 130.602 | --- | 2 |
| K | 32 | --- | --- | 74.060 | --- | 1 |

**Soleil Index**

| | | |
|---|---|---|
| 1060.3689 | not explained | 22% |
| 1137.5425 | y2-19 frag loss of water | 15% |

| | | |
|---|---|---|
| **1244.0620** | **not explained** | **41%** |
| **1309.4983** | **not explained** | **15%** |
| **1395.5485** | **not explained** | **23%** |
| **1589.9598** | **not explained** | **100%** |

| | | | | | | | |
|---|---|---|---|---|---|---|---|
| **Run:** | 79491 | **Scans:** | 32135 - 32135 | **m/z Range:** | 325.03925 | **to** | 1987.2357[illegible] |
| **Search:** | 79491 | **Precursor Mass:** | 1156.90096 | | | | |
| **Mods:** | C=160.03 *+15.99 #+220.06 @+291.1 | **Charge:** | +3 | | | | |

**Sequence**

NIALST@SGSGNVGVM*TQSGAEQAITIIDDAIK

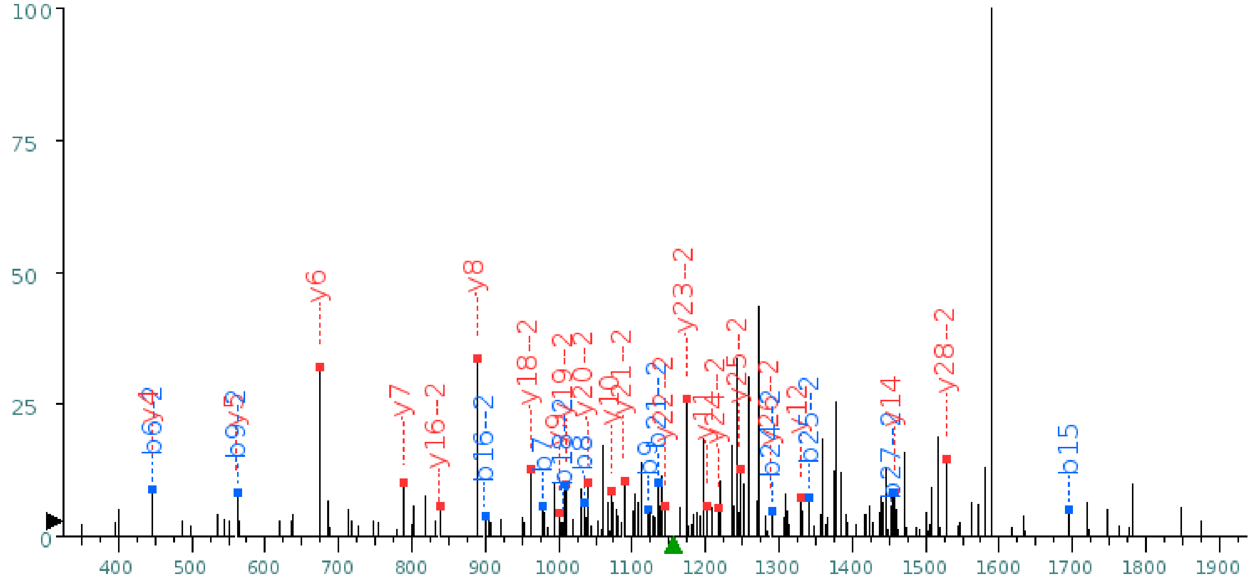

**Predicted Fragmentation Pattern**

**+1**

| Seq | # | b: Δ Error | b | y | y: Δ Error | +1 |
|---|---|---|---|---|---|---|
| N | 1 | --- | 115.050 | --- | --- | 32 |
| I | 2 | --- | 228.134 | 3354.647 | --- | 31 |
| A | 3 | --- | 299.171 | 3241.563 | --- | 30 |
| L | 4 | --- | 412.255 | 3170.526 | --- | 29 |
| S | 5 | --- | 499.287 | 3057.442 | --- | 28 |
| T@ | 6 | --- | 891.431 | 2970.410 | --- | 27 |
| S | 7 | -594.400 | **978.463** | 2578.266 | --- | 26 |
| G | 8 | 665.330 | **1035.484** | 2491.234 | --- | 25 |
| S | 9 | -579.504 | **1122.516** | 2434.213 | --- | 24 |
| G | 10 | --- | 1179.538 | 2347.181 | --- | 23 |

| | | | | | | |
|---|---|---|---|---|---|---|
| N | 11 | --- | 1293.580 | 2290.160 | --- | 22 |
| V | 12 | --- | 1392.649 | 2176.117 | --- | 21 |
| G | 13 | --- | 1449.670 | 2077.048 | --- | 20 |
| V | 14 | --- | 1548.739 | 2020.027 | --- | 19 |
| M* | 15 | **159.347** | **1695.774** | 1920.958 | --- | 18 |
| T | 16 | --- | 1796.822 | 1773.923 | --- | 17 |
| Q | 17 | --- | 1924.880 | 1672.875 | --- | 16 |
| S | 18 | --- | 2011.912 | 1544.817 | --- | 15 |
| G | 19 | --- | 2068.934 | **1457.785** | **151.336** | 14 |
| A | 20 | --- | 2139.971 | 1400.763 | --- | 13 |
| E | 21 | --- | 2269.014 | **1329.726** | **132.490** | 12 |
| Q | 22 | --- | 2397.072 | **1200.683** | **188.897** | 11 |
| A | 23 | --- | 2468.109 | **1072.625** | **-53.126** | 10 |
| I | 24 | --- | 2581.193 | **1001.588** | **210.101** | 9 |
| T | 25 | --- | 2682.241 | **888.504** | **236.376** | 8 |
| I | 26 | --- | 2795.325 | **787.456** | **347.505** | 7 |
| I | 27 | --- | 2908.409 | **674.372** | **355.950** | 6 |
| D | 28 | --- | 3023.436 | **561.288** | **425.168** | 5 |
| D | 29 | --- | 3138.463 | **446.261** | **446.833** | 4 |
| A | 30 | --- | 3209.500 | 331.234 | --- | 3 |
| I | 31 | --- | 3322.584 | 260.197 | --- | 2 |
| K | 32 | --- | --- | 147.113 | --- | 1 |

**+2**

| Seq | # | b: Δ Error | b | y | y: Δ Error | +1 |
|---|---|---|---|---|---|---|
| N | 1 | --- | 58.029 | --- | --- | 32 |
| I | 2 | --- | 114.571 | 1677.827 | --- | 31 |
| A | 3 | --- | 150.089 | 1621.285 | --- | 30 |
| L | 4 | --- | 206.631 | 1585.766 | --- | 29 |
| S | 5 | --- | 250.147 | **1529.224** | **112.169** | 28 |
| T@ | 6 | **540.909** | **446.219** | 1485.708 | --- | 27 |
| S | 7 | --- | 489.735 | **1289.637** | **618.817** | 26 |
| G | 8 | --- | 518.246 | **1246.121** | **557.558** | 25 |
| S | 9 | **-418.632** | **561.762** | **1217.610** | **174.689** | 24 |
| G | 10 | --- | 590.272 | **1174.094** | **-372.563** | 23 |
| N | 11 | --- | 647.294 | **1145.583** | **181.953** | 22 |
| V | 12 | --- | 696.828 | **1088.562** | **557.659** | 21 |
| G | 13 | --- | 725.339 | **1039.028** | **10.095** | 20 |
| V | 14 | --- | 774.873 | **1010.517** | **472.085** | 19 |
| M* | 15 | --- | 848.391 | **960.983** | **411.786** | 18 |
| T | 16 | **720.685** | **898.915** | 887.465 | --- | 17 |
| Q | 17 | --- | 962.944 | **836.941** | **905.852** | 16 |
| S | 18 | **444.654** | **1006.460** | 772.912 | --- | 15 |
| G | 19 | --- | 1034.971 | 729.396 | --- | 14 |
| A | 20 | --- | 1070.489 | 700.885 | --- | 13 |
| E | 21 | **553.401** | **1135.010** | 665.367 | --- | 12 |
| Q | 22 | --- | 1199.040 | 600.845 | --- | 11 |
| A | 23 | --- | 1234.558 | 536.816 | --- | 10 |
| I | 24 | **-515.257** | **1291.100** | 501.298 | --- | 9 |
| T | 25 | **-230.177** | **1341.624** | 444.755 | --- | 8 |
| I | 26 | --- | 1398.166 | 394.232 | --- | 7 |
| I | 27 | **488.489** | **1454.708** | 337.690 | --- | 6 |
| D | 28 | --- | 1512.222 | 281.148 | --- | 5 |
| D | 29 | --- | 1569.735 | 223.634 | --- | 4 |
| A | 30 | --- | 1605.254 | 166.121 | --- | 3 |
| I | 31 | --- | 1661.796 | 130.602 | --- | 2 |
| K | 32 | --- | --- | 74.060 | --- | 1 |

**Soleil Index**

| | | |
|---|---|---|
| 1060.7109 | not explained | 17% |
| 1196.9073 | not explained | 18% |
| 1236.3459 | not explained | 17% |
| 1243.9493 | not explained | 34% |
| 1258.4806 | not explained | 30% |
| 1273.6014 | b2-24 frag loss of water/ b2-24 frag loss of ammonia/ y2-26 frag loss of ammonia | 43% |
| 1360.1007 | not explained | 18% |
| 1377.9930 | not explained | 26% |
| 1470.9103 | not explained | 16% |
| 1517.1450 | not explained | 19% |
| 1589.7202 | not explained | 100% |
| 1590.5444 | not explained | 23% |

**MS/MS data of GluC digested peptides with site score A and site score B above 19 from *Geobacillus kaustophilus* Flagellin *Gk*F2 protein co-expressed with *Geobacillus kaustophilus* Maf in EV136 cells. Results of dataset 3 (MS/MS results obtained in July 2019).**

| | | | | | |
|---|---|---|---|---|---|
| **Run:** | 79492 | **Scans:** | 26030 - 26030 | **m/z Range:** | 317.36453 to 2000 |
| **Search:** | 79492 | **Precursor Mass:** | 998.5103 | | |
| **Mods:** | C=160.03 *+15.99 #+220.06 @+291.1 | **Charge:** | +4 | | |

**Sequence**

FT#KNNILT#QAAQAMLAQANQQPQGVLQLLRLE

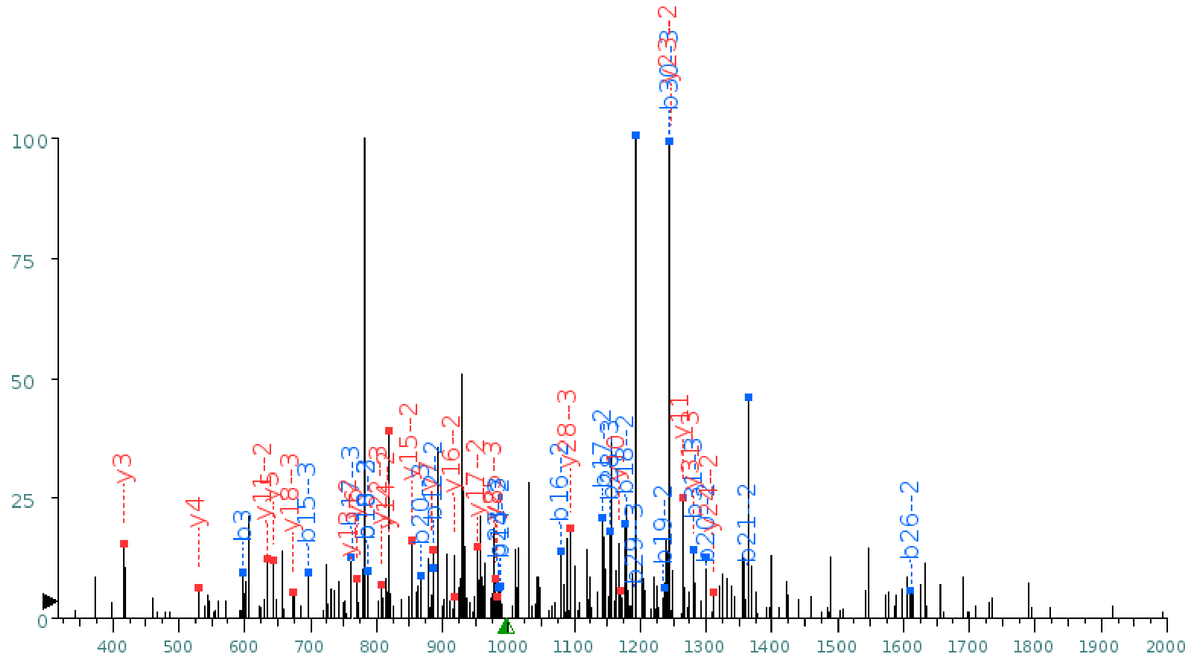

**Predicted Fragmentation Pattern**

| | | | +1 | | | |
|---|---|---|---|---|---|---|
| **Seq** | **#** | **b: Δ Error** | **b** | **y** | **y: Δ Error** | **+1** |
| F | 1 | --- | 148.076 | --- | --- | 32 |
| T# | 2 | --- | 469.182 | 3843.964 | --- | 31 |
| K | 3 | **765.594** | **597.277** | 3522.858 | --- | 30 |
| N | 4 | --- | 711.320 | 3394.763 | --- | 29 |
| N | 5 | --- | 825.362 | 3280.721 | --- | 28 |
| I | 6 | --- | 938.447 | 3166.678 | --- | 27 |
| L | 7 | --- | 1051.531 | 3053.594 | --- | 26 |
| T# | 8 | --- | 1372.637 | 2940.510 | --- | 25 |
| Q | 9 | --- | 1500.695 | 2619.404 | --- | 24 |
| A | 10 | --- | 1571.732 | 2491.345 | --- | 23 |
| A | 11 | --- | 1642.769 | 2420.308 | --- | 22 |
| Q | 12 | --- | 1770.828 | 2349.271 | --- | 21 |
| A | 13 | --- | 1841.865 | 2221.212 | --- | 20 |

| | | | | | | |
|---|---|---|---|---|---|---|
| M | 14 | --- | 1972.906 | 2150.175 | --- | 19 |
| L | 15 | --- | 2085.990 | 2019.135 | --- | 18 |
| A | 16 | --- | 2157.027 | 1906.050 | --- | 17 |
| Q | 17 | --- | 2285.085 | 1835.013 | --- | 16 |
| A | 18 | --- | 2356.122 | 1706.955 | --- | 15 |
| N | 19 | --- | 2470.165 | 1635.918 | --- | 14 |
| Q | 20 | --- | 2598.224 | 1521.875 | --- | 13 |
| Q | 21 | --- | 2726.283 | 1393.816 | --- | 12 |
| P | 22 | --- | 2823.335 | **1265.758** | 232.923 | 11 |
| Q | 23 | --- | 2951.394 | **1168.705** | 606.586 | 10 |
| G | 24 | --- | 3008.415 | 1040.646 | --- | 9 |
| V | 25 | --- | 3107.484 | **983.625** | 544.600 | 8 |
| L | 26 | --- | 3220.568 | **884.556** | 326.425 | 7 |
| Q | 27 | --- | 3348.626 | **771.472** | -45.447 | 6 |
| L | 28 | --- | 3461.710 | **643.414** | -1131.522 | 5 |
| L | 29 | --- | 3574.795 | **530.330** | 74.644 | 4 |
| R | 30 | --- | 3730.896 | **417.246** | 419.495 | 3 |
| L | 31 | --- | 3843.980 | 261.144 | --- | 2 |
| E | 32 | --- | --- | 148.060 | --- | 1 |

+2

| Seq | # | b: Δ Error | b | y | y: Δ Error | +1 |
|---|---|---|---|---|---|---|
| F | 1 | --- | 74.541 | --- | --- | 32 |
| T# | 2 | --- | 235.094 | 1922.486 | --- | 31 |
| K | 3 | --- | 299.142 | 1761.933 | --- | 30 |
| N | 4 | --- | 356.163 | 1697.885 | --- | 29 |
| N | 5 | --- | 413.185 | 1640.864 | --- | 28 |
| I | 6 | --- | 469.727 | 1583.842 | --- | 27 |
| L | 7 | --- | 526.269 | 1527.300 | --- | 26 |
| T# | 8 | --- | 686.822 | 1470.758 | --- | 25 |
| Q | 9 | --- | 750.851 | **1310.205** | 310.187 | 24 |
| A | 10 | 7.414 | **786.370** | **1246.176** | -739.303 | 23 |
| A | 11 | --- | 821.888 | 1210.658 | --- | 22 |
| Q | 12 | -27.723 | **885.918** | 1175.139 | --- | 21 |
| A | 13 | --- | 921.436 | 1111.110 | --- | 20 |
| M | 14 | 695.109 | **986.956** | 1075.591 | --- | 19 |
| L | 15 | --- | 1043.498 | 1010.071 | --- | 18 |
| A | 16 | 220.854 | **1079.017** | **953.529** | -624.689 | 17 |
| Q | 17 | -217.085 | **1143.046** | **918.010** | 732.465 | 16 |
| A | 18 | -240.387 | **1178.565** | **853.981** | 871.527 | 15 |
| N | 19 | 626.449 | **1235.586** | **818.462** | 414.510 | 14 |
| Q | 20 | 725.059 | **1299.616** | **761.441** | 723.972 | 13 |
| Q | 21 | 362.413 | **1363.645** | 697.412 | --- | 12 |
| P | 22 | --- | 1412.171 | **633.382** | 1144.085 | 11 |
| Q | 23 | --- | 1476.201 | 584.856 | --- | 10 |
| G | 24 | --- | 1504.711 | 520.827 | --- | 9 |
| V | 25 | --- | 1554.246 | 492.316 | --- | 8 |
| L | 26 | -353.770 | **1610.788** | 442.782 | --- | 7 |
| Q | 27 | --- | 1674.817 | 386.240 | --- | 6 |
| L | 28 | --- | 1731.359 | 322.211 | --- | 5 |
| L | 29 | --- | 1787.901 | 265.668 | --- | 4 |
| R | 30 | --- | 1865.951 | 209.126 | --- | 3 |
| L | 31 | --- | 1922.493 | 131.076 | --- | 2 |
| E | 32 | --- | --- | 74.534 | --- | 1 |

+3

| Seq | # | b: Δ Error | b | y | y: Δ Error | +1 |
|---|---|---|---|---|---|---|
| F | 1 | --- | 50.030 | --- | --- | 32 |

| | | | | | | |
|---|---|---|---|---|---|---|
| T# | 2 | --- | 157.065 | **1281.993** | 204.240 | 31 |
| K | 3 | --- | 199.764 | 1174.958 | --- | 30 |
| N | 4 | --- | 237.778 | 1132.259 | --- | 29 |
| N | 5 | --- | 275.792 | **1094.245** | -738.309 | 28 |
| I | 6 | --- | 313.487 | 1056.231 | --- | 27 |
| L | 7 | --- | 351.182 | 1018.536 | --- | 26 |
| T# | 8 | --- | 458.217 | **980.841** | 625.197 | 25 |
| Q | 9 | --- | 500.903 | 873.806 | --- | 24 |
| A | 10 | --- | 524.582 | 831.120 | --- | 23 |
| A | 11 | --- | 548.261 | **807.441** | -501.572 | 22 |
| Q | 12 | --- | 590.948 | 783.762 | --- | 21 |
| A | 13 | --- | 614.627 | 741.076 | --- | 20 |
| M | 14 | --- | 658.307 | 717.397 | --- | 19 |
| L | 15 | 456.425 | **696.001** | **673.716** | -16.394 | 18 |
| A | 16 | --- | 719.680 | 636.022 | --- | 17 |
| Q | 17 | -490.754 | **762.367** | 612.343 | --- | 16 |
| A | 18 | 419.577 | **786.046** | 569.656 | --- | 15 |
| N | 19 | --- | 824.060 | 545.977 | --- | 14 |
| Q | 20 | 216.336 | **866.746** | 507.963 | --- | 13 |
| Q | 21 | --- | 909.432 | 465.277 | --- | 12 |
| P | 22 | --- | 941.783 | 422.591 | --- | 11 |
| Q | 23 | 852.128 | **984.469** | 390.240 | --- | 10 |
| G | 24 | --- | 1003.477 | 347.554 | --- | 9 |
| V | 25 | --- | 1036.499 | 328.546 | --- | 8 |
| L | 26 | --- | 1074.194 | 295.524 | --- | 7 |
| Q | 27 | --- | 1116.880 | 257.829 | --- | 6 |
| L | 28 | 58.844 | **1154.575** | 215.143 | --- | 5 |
| L | 29 | 706.745 | **1192.270** | 177.448 | --- | 4 |
| R | 30 | 764.583 | **1244.303** | 139.753 | --- | 3 |
| L | 31 | 200.274 | **1281.998** | 87.720 | --- | 2 |
| E | 32 | --- | --- | 50.025 | --- | 1 |

**Soleil Index**

| | | |
|---|---|---|
| 605.4486 | y2-22 frag doubly charged | 21% |
| 781.0890 | not explained | 100% |
| 892.7509 | b3-21 frag loss of ammonia | 35% |
| 929.7809 | not explained | 51% |
| 931.2443 | not explained | 27% |
| 934.3986 | not explained | 15% |
| 957.6259 | not explained | 21% |
| 979.0923 | not explained | 19% |
| 1032.0631 | not explained | 28% |
| 1089.4017 | not explained | 16% |
| 1145.4504 | not explained | 17% |
| 1155.6515 | not explained | 37% |
| 1166.8704 | not explained | 16% |
| 1179.7834 | not explained | 23% |
| 1239.5653 | not explained | 16% |

| | | | | | |
|---|---|---|---|---|---|
| Run: | 79492 | Scans: | 26110 - 26110 | m/z Range: | 379.26224 to 1980.0737 |
| Search: | 79492 | Precursor | 998.51164 | | |

**Mods:** C=160

**Mass:**

**Sequence**

FT#KNNILT#QAAQAMLAQANQQPQGVLQLLRLE

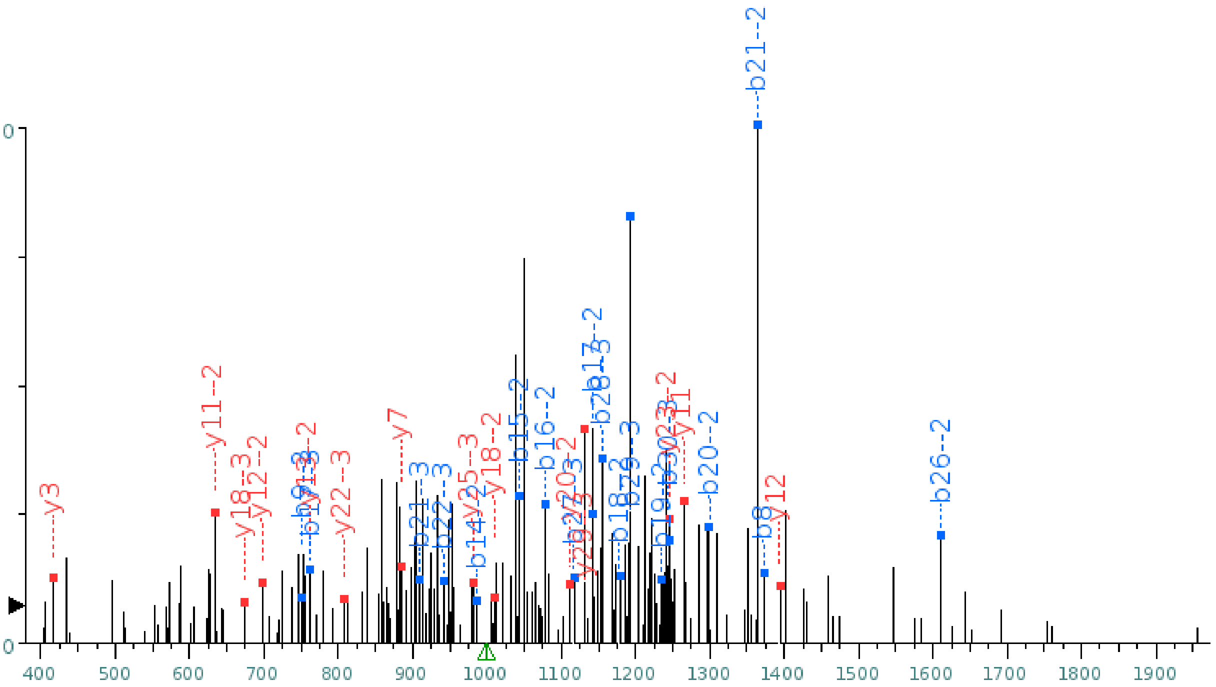

**Predicted Fragmentation Pattern**

| | | | +1 | | | |
|---|---|---|---|---|---|---|
| **Seq** | **#** | **b: Δ Error** | **b** | **y** | **y: Δ Error** | **+1** |
| **F** | **1** | **---** | 148.076 | --- | **---** | **32** |
| **T#** | **2** | **---** | 469.182 | 3843.964 | **---** | **31** |
| **K** | **3** | **---** | 597.277 | 3522.858 | **---** | **30** |
| **N** | **4** | **---** | 711.320 | 3394.763 | **---** | **29** |
| **N** | **5** | **---** | 825.362 | 3280.721 | **---** | **28** |
| **I** | **6** | **---** | 938.447 | 3166.678 | **---** | **27** |
| **L** | **7** | **---** | 1051.531 | 3053.594 | **---** | **26** |
| **T#** | **8** | **-94.807** | **1372.637** | 2940.510 | **---** | **25** |
| **Q** | **9** | **---** | 1500.695 | 2619.404 | **---** | **24** |
| **A** | **10** | **---** | 1571.732 | 2491.345 | **---** | **23** |
| **A** | **11** | **---** | 1642.769 | 2420.308 | **---** | **22** |
| **Q** | **12** | **---** | 1770.828 | 2349.271 | **---** | **21** |
| **A** | **13** | **---** | 1841.865 | 2221.212 | **---** | **20** |
| **M** | **14** | **---** | 1972.906 | 2150.175 | **---** | **19** |
| **L** | **15** | **---** | 2085.990 | 2019.135 | **---** | **18** |
| **A** | **16** | **---** | 2157.027 | 1906.050 | **---** | **17** |

| | | | | | | |
|---|---|---|---|---|---|---|
| Q | 17 | --- | 2285.085 | 1835.013 | --- | 16 |
| A | 18 | --- | 2356.122 | 1706.955 | --- | 15 |
| N | 19 | --- | 2470.165 | 1635.918 | --- | 14 |
| Q | 20 | --- | 2598.224 | 1521.875 | --- | 13 |
| Q | 21 | --- | 2726.283 | **1393.816** | 198.845 | 12 |
| P | 22 | --- | 2823.335 | **1265.758** | 239.767 | 11 |
| Q | 23 | --- | 2951.394 | 1168.705 | --- | 10 |
| G | 24 | --- | 3008.415 | 1040.646 | --- | 9 |
| V | 25 | --- | 3107.484 | 983.625 | --- | 8 |
| L | 26 | --- | 3220.568 | **884.556** | 291.394 | 7 |
| Q | 27 | --- | 3348.626 | 771.472 | --- | 6 |
| L | 28 | --- | 3461.710 | 643.414 | --- | 5 |
| L | 29 | --- | 3574.795 | 530.330 | --- | 4 |
| R | 30 | --- | 3730.896 | **417.246** | 581.193 | 3 |
| L | 31 | --- | 3843.980 | 261.144 | --- | 2 |
| E | 32 | --- | --- | 148.060 | --- | 1 |

+2

| Seq | # | b: Δ Error | b | y | y: Δ Error | +1 |
|---|---|---|---|---|---|---|
| F | 1 | --- | 74.541 | --- | --- | 32 |
| T# | 2 | --- | 235.094 | 1922.486 | --- | 31 |
| K | 3 | --- | 299.142 | 1761.933 | --- | 30 |
| N | 4 | --- | 356.163 | 1697.885 | --- | 29 |
| N | 5 | --- | 413.185 | 1640.864 | --- | 28 |
| I | 6 | --- | 469.727 | 1583.842 | --- | 27 |
| L | 7 | --- | 526.269 | 1527.300 | --- | 26 |
| T# | 8 | --- | 686.822 | 1470.758 | --- | 25 |
| Q | 9 | 172.753 | **750.851** | 1310.205 | --- | 24 |
| A | 10 | --- | 786.370 | **1246.176** | -404.695 | 23 |
| A | 11 | --- | 821.888 | 1210.658 | --- | 22 |
| Q | 12 | --- | 885.918 | 1175.139 | --- | 21 |
| A | 13 | --- | 921.436 | **1111.110** | 172.702 | 20 |
| M | 14 | -291.487 | **986.956** | 1075.591 | --- | 19 |
| L | 15 | -371.246 | **1043.498** | **1010.071** | 0.735 | 18 |
| A | 16 | -904.437 | **1079.017** | 953.529 | --- | 17 |
| Q | 17 | -211.316 | **1143.046** | 918.010 | --- | 16 |
| A | 18 | 168.144 | **1178.565** | 853.981 | --- | 15 |
| N | 19 | -13.254 | **1235.586** | 818.462 | --- | 14 |
| Q | 20 | -140.921 | **1299.616** | **761.441** | 201.028 | 13 |
| Q | 21 | 194.213 | **1363.645** | **697.412** | 637.615 | 12 |
| P | 22 | --- | 1412.171 | **633.382** | 264.560 | 11 |
| Q | 23 | --- | 1476.201 | 584.856 | --- | 10 |
| G | 24 | --- | 1504.711 | 520.827 | --- | 9 |
| V | 25 | --- | 1554.246 | 492.316 | --- | 8 |
| L | 26 | 546.359 | **1610.788** | 442.782 | --- | 7 |
| Q | 27 | --- | 1674.817 | 386.240 | --- | 6 |
| L | 28 | --- | 1731.359 | 322.211 | --- | 5 |
| L | 29 | --- | 1787.901 | 265.668 | --- | 4 |
| R | 30 | --- | 1865.951 | 209.126 | --- | 3 |
| L | 31 | --- | 1922.493 | 131.076 | --- | 2 |
| E | 32 | --- | --- | 74.534 | --- | 1 |

+3

| Seq | # | b: Δ Error | b | y | y: Δ Error | +1 |
|---|---|---|---|---|---|---|
| F | 1 | --- | 50.030 | --- | --- | 32 |
| T# | 2 | --- | 157.065 | 1281.993 | --- | 31 |
| K | 3 | --- | 199.764 | 1174.958 | --- | 30 |
| N | 4 | --- | 237.778 | **1132.259** | -737.280 | 29 |

| | | | | | | |
|---|---|---|---|---|---|---|
| N | 5 | --- | 275.792 | 1094.245 | --- | 28 |
| I | 6 | --- | 313.487 | 1056.231 | --- | 27 |
| L | 7 | --- | 351.182 | 1018.536 | --- | 26 |
| T# | 8 | --- | 458.217 | **980.841** | 999.694 | 25 |
| Q | 9 | --- | 500.903 | 873.806 | --- | 24 |
| A | 10 | --- | 524.582 | 831.120 | --- | 23 |
| A | 11 | --- | 548.261 | **807.441** | 181.837 | 22 |
| Q | 12 | --- | 590.948 | 783.762 | --- | 21 |
| A | 13 | --- | 614.627 | 741.076 | --- | 20 |
| M | 14 | --- | 658.307 | 717.397 | --- | 19 |
| L | 15 | --- | 696.001 | **673.716** | -875.454 | 18 |
| A | 16 | --- | 719.680 | 636.022 | --- | 17 |
| Q | 17 | -1014.334 | **762.367** | 612.343 | --- | 16 |
| A | 18 | --- | 786.046 | 569.656 | --- | 15 |
| N | 19 | --- | 824.060 | 545.977 | --- | 14 |
| Q | 20 | --- | 866.746 | 507.963 | --- | 13 |
| Q | 21 | 4.982 | **909.432** | 465.277 | --- | 12 |
| P | 22 | -428.258 | **941.783** | 422.591 | --- | 11 |
| Q | 23 | --- | 984.469 | 390.240 | --- | 10 |
| G | 24 | --- | 1003.477 | 347.554 | --- | 9 |
| V | 25 | --- | 1036.499 | 328.546 | --- | 8 |
| L | 26 | --- | 1074.194 | 295.524 | --- | 7 |
| Q | 27 | 286.216 | **1116.880** | 257.829 | --- | 6 |
| L | 28 | 440.541 | **1154.575** | 215.143 | --- | 5 |
| L | 29 | 559.191 | **1192.270** | 177.448 | --- | 4 |
| R | 30 | 576.869 | **1244.303** | 139.753 | --- | 3 |
| L | 31 | --- | 1281.998 | 87.720 | --- | 2 |
| E | 32 | --- | --- | 50.025 | --- | 1 |

Soleil Index

| | | |
|---|---|---|
| 433.2753 | b3-20 frag doubly charged | 17% |
| 746.1731 | b3-17 frag loss of ammonia | 17% |
| 752.5732 | b2-24 frag doubly charged/ y1-6 frag loss of water | 17% |
| 837.7693 | b2-27 frag doubly charged/ y2-15 frag loss of ammonia | 18% |
| 857.6837 | y3-24 frag loss of ammonia | 32% |
| 878.1230 | not explained | 31% |
| 882.7185 | not explained | 26% |
| 902.6719 | b2-13 frag loss of water | 20% |
| 905.2378 | b2-13 frag loss of ammonia | 31% |
| 913.7511 | not explained | 28% |
| 925.1642 | b3-22 frag loss of ammonia | 18% |
| 933.9264 | b2-30 frag doubly charged | 29% |
| 949.2609 | not explained | 24% |
| 952.2101 | y1-17 frag doubly charged | 27% |
| 1012.9438 | not explained | 16% |
| 1021.6649 | y1-9 frag loss of water | 16% |
| 1039.2382 | y3-27 frag loss of ammonia | 56% |
| 1048.9424 | not explained | 75% |
| 1141.9745 | b1-17 frag doubly charged | 42% |

| | | |
|---|---|---|
| 1155.7123 | not explained | 23% |
| 1169.7083 | not explained | 21% |
| 1172.4379 | not explained | 15% |
| 1186.5380 | not explained | 19% |
| 1203.2738 | not explained | 19% |
| 1211.8209 | not explained | 32% |
| 1218.6639 | b2-19 frag loss of ammonia | 18% |
| 1219.6948 | not explained | 15% |
| 1221.9297 | not explained | 24% |
| 1240.4908 | not explained | 38% |
| 1284.9235 | not explained | 23% |
| 1295.9159 | not explained | 22% |
| 1308.5107 | not explained | 21% |
| 1350.0975 | not explained | 22% |
| 1364.8862 | not explained | 19% |
| 1401.0641 | not explained | 17% |
| 1402.0265 | not explained | 26% |

| | | | |
|---|---|---|---|
| **Run:** | 79492 | **Scans:** | 26140 - 26140 |
| **Search:** | 79492 | **Precursor Mass:** | 1002.51055 |
| **Mods:** | C=160.03 *+15.99 #+220.06 @+291.1 | **Charge:** | +4 |

**m/z Range:** 259.22281 to 1886.4512

**Sequence**

FT#KNNILT#QAAQAM*LAQANQQPQGVLQLLRLE

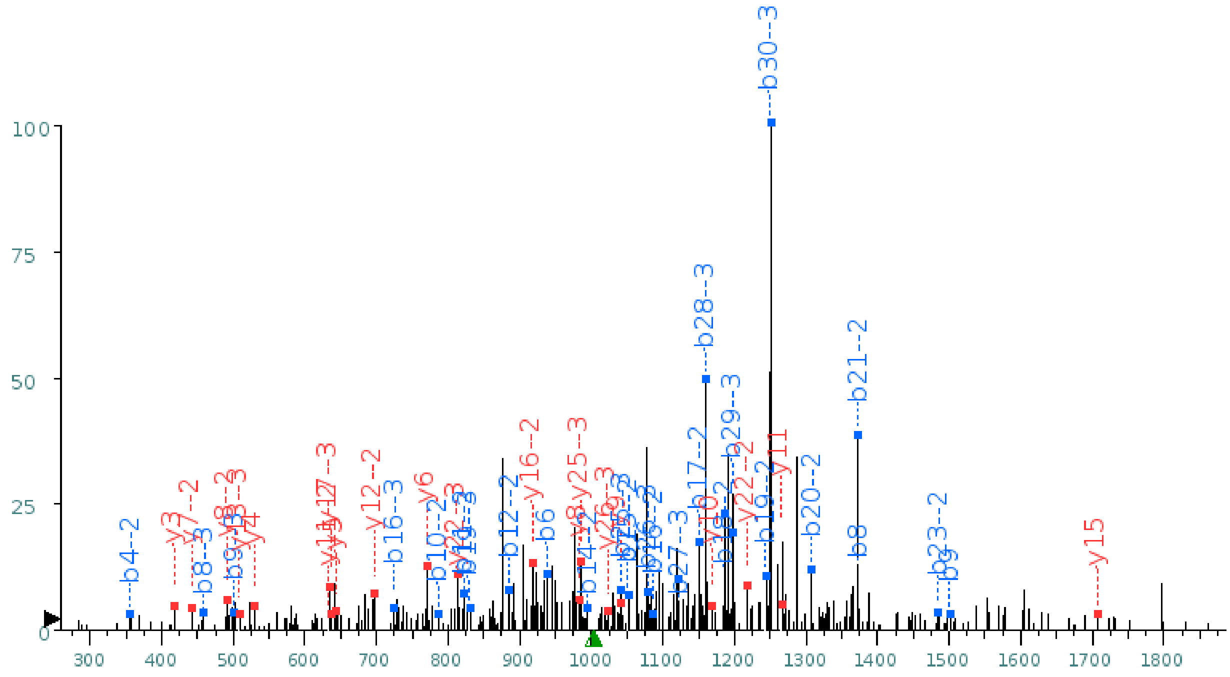

**Predicted Fragmentation Pattern**

| Seq | # | b: Δ Error | b | y | y: Δ Error | +1 |
|---|---|---|---|---|---|---|
| F | 1 | --- | 148.076 | --- | --- | 32 |
| T# | 2 | --- | 469.182 | 3859.959 | --- | 31 |
| K | 3 | --- | 597.277 | 3538.853 | --- | 30 |
| N | 4 | --- | 711.320 | 3410.758 | --- | 29 |
| N | 5 | --- | 825.362 | 3296.715 | --- | 28 |
| I | 6 | -104.871 | **938.447** | 3182.673 | --- | 27 |
| L | 7 | 430.758 | **1051.531** | 3069.588 | --- | 26 |
| T# | 8 | -289.107 | **1372.637** | 2956.504 | --- | 25 |

| Seq | # | b: Δ Error | b | y | y: Δ Error | +1 |
|---|---|---|---|---|---|---|
| Q | 9 | 585.662 | **1500.695** | 2635.398 | --- | 24 |
| A | 10 | --- | 1571.732 | 2507.340 | --- | 23 |
| A | 11 | --- | 1642.769 | 2436.303 | --- | 22 |
| Q | 12 | --- | 1770.828 | 2365.266 | --- | 21 |
| A | 13 | --- | 1841.865 | 2237.207 | --- | 20 |
| M* | 14 | --- | 1988.900 | 2166.170 | --- | 19 |
| L | 15 | --- | 2101.985 | 2019.135 | --- | 18 |
| A | 16 | --- | 2173.022 | 1906.050 | --- | 17 |
| Q | 17 | --- | 2301.080 | 1835.013 | --- | 16 |
| A | 18 | --- | 2372.117 | **1706.955** | 534.587 | 15 |
| N | 19 | --- | 2486.160 | 1635.918 | --- | 14 |
| Q | 20 | --- | 2614.219 | 1521.875 | --- | 13 |
| Q | 21 | --- | 2742.277 | 1393.816 | --- | 12 |
| P | 22 | --- | 2839.330 | **1265.758** | 355.908 | 11 |
| Q | 23 | --- | 2967.389 | **1168.705** | 411.465 | 10 |
| G | 24 | --- | 3024.410 | **1040.646** | 891.625 | 9 |
| V | 25 | --- | 3123.479 | **983.625** | -488.868 | 8 |
| L | 26 | --- | 3236.563 | 884.556 | --- | 7 |
| Q | 27 | --- | 3364.621 | **771.472** | 161.811 | 6 |
| L | 28 | --- | 3477.705 | **643.414** | 365.019 | 5 |
| L | 29 | --- | 3590.789 | **530.330** | 147.250 | 4 |
| R | 30 | --- | 3746.891 | **417.246** | 635.032 | 3 |
| L | 31 | --- | 3859.975 | 261.144 | --- | 2 |
| E | 32 | --- | --- | 148.060 | --- | 1 |

+2

| Seq | # | b: Δ Error | b | y | y: Δ Error | +1 |
|---|---|---|---|---|---|---|
| F | 1 | --- | 74.541 | --- | --- | 32 |
| T# | 2 | --- | 235.094 | 1930.483 | --- | 31 |
| K | 3 | --- | 299.142 | 1769.930 | --- | 30 |
| N | 4 | 458.127 | **356.163** | 1705.883 | --- | 29 |
| N | 5 | --- | 413.185 | 1648.861 | --- | 28 |
| I | 6 | --- | 469.727 | 1591.840 | --- | 27 |
| L | 7 | --- | 526.269 | 1535.298 | --- | 26 |
| T# | 8 | --- | 686.822 | 1478.756 | --- | 25 |
| Q | 9 | --- | 750.851 | 1318.203 | --- | 24 |
| A | 10 | -322.404 | **786.370** | 1254.174 | --- | 23 |
| A | 11 | 966.365 | **821.888** | **1218.655** | 103.074 | 22 |
| Q | 12 | -154.099 | **885.918** | 1183.136 | --- | 21 |
| A | 13 | --- | 921.436 | 1119.107 | --- | 20 |
| M* | 14 | -629.941 | **994.954** | 1083.589 | --- | 19 |
| L | 15 | 463.743 | **1051.496** | 1010.071 | --- | 18 |
| A | 16 | -174.492 | **1087.014** | 953.529 | --- | 17 |
| Q | 17 | 368.661 | **1151.044** | **918.010** | 832.039 | 16 |
| A | 18 | 224.479 | **1186.562** | 853.981 | --- | 15 |
| N | 19 | 246.191 | **1243.584** | 818.462 | --- | 14 |
| Q | 20 | 659.177 | **1307.613** | 761.441 | --- | 13 |
| Q | 21 | 435.427 | **1371.642** | **697.412** | 1059.079 | 12 |
| P | 22 | --- | 1420.169 | **633.382** | 881.544 | 11 |
| Q | 23 | 209.171 | **1484.198** | 584.856 | --- | 10 |
| G | 24 | --- | 1512.709 | 520.827 | --- | 9 |
| V | 25 | --- | 1562.243 | **492.316** | -1861.967 | 8 |
| L | 26 | --- | 1618.785 | **442.782** | 1038.138 | 7 |
| Q | 27 | --- | 1682.814 | 386.240 | --- | 6 |
| L | 28 | --- | 1739.356 | 322.211 | --- | 5 |
| L | 29 | --- | 1795.898 | 265.668 | --- | 4 |
| R | 30 | --- | 1873.949 | 209.126 | --- | 3 |
| L | 31 | --- | 1930.491 | 131.076 | --- | 2 |

| | | | | | | |
|---|---|---|---|---|---|---|
| E | 32 | --- | --- | 74.534 | --- | 1 |

+3

| Seq | # | b: Δ Error | b | y | y: Δ Error | +1 |
|---|---|---|---|---|---|---|
| F | 1 | --- | 50.030 | --- | --- | 32 |
| T# | 2 | --- | 157.065 | 1287.325 | --- | 31 |
| K | 3 | --- | 199.764 | 1180.289 | --- | 30 |
| N | 4 | --- | 237.778 | 1137.591 | --- | 29 |
| N | 5 | --- | 275.792 | 1099.577 | --- | 28 |
| I | 6 | --- | 313.487 | 1061.562 | --- | 27 |
| L | 7 | --- | 351.182 | **1023.868** | 78.559 | 26 |
| T# | 8 | -1309.366 | **458.217** | **986.173** | -412.630 | 25 |
| Q | 9 | -681.570 | **500.903** | 879.138 | --- | 24 |
| A | 10 | --- | 524.582 | 836.451 | --- | 23 |
| A | 11 | --- | 548.261 | **812.772** | -374.082 | 22 |
| Q | 12 | --- | 590.948 | 789.093 | --- | 21 |
| A | 13 | --- | 614.627 | 746.407 | --- | 20 |
| M* | 14 | --- | 663.638 | 722.728 | --- | 19 |
| L | 15 | --- | 701.333 | 673.716 | --- | 18 |
| A | 16 | -631.770 | **725.012** | **636.022** | -113.173 | 17 |
| Q | 17 | --- | 767.698 | 612.343 | --- | 16 |
| A | 18 | --- | 791.377 | 569.656 | --- | 15 |
| N | 19 | 1178.543 | **829.392** | 545.977 | --- | 14 |
| Q | 20 | --- | 872.078 | **507.963** | 1003.867 | 13 |
| Q | 21 | --- | 914.764 | 465.277 | --- | 12 |
| P | 22 | --- | 947.115 | 422.591 | --- | 11 |
| Q | 23 | --- | 989.801 | 390.240 | --- | 10 |
| G | 24 | --- | 1008.808 | 347.554 | --- | 9 |
| V | 25 | 587.008 | **1041.831** | 328.546 | --- | 8 |
| L | 26 | -701.802 | **1079.526** | 295.524 | --- | 7 |
| Q | 27 | -608.137 | **1122.212** | 257.829 | --- | 6 |
| L | 28 | 581.312 | **1159.907** | 215.143 | --- | 5 |
| L | 29 | 400.821 | **1197.601** | 177.448 | --- | 4 |
| R | 30 | 791.970 | **1249.635** | 139.753 | --- | 3 |
| L | 31 | --- | 1287.330 | 87.720 | --- | 2 |
| E | 32 | --- | --- | 50.025 | --- | 1 |

**Soleil Index**

| | | |
|---|---|---|
| 875.5577 | not explained | 34% |
| 905.8472 | not explained | 17% |
| 975.7272 | not explained | 20% |
| 1063.5714 | not explained | 19% |
| 1077.8512 | not explained | 36% |
| 1120.6018 | y3-29 frag loss of ammonia | 16% |
| 1192.2958 | not explained | 35% |
| 1198.8842 | not explained | 27% |
| 1267.1779 | not explained | 17% |
| 1288.4777 | not explained | 34% |

| | | | | | |
|---|---|---|---|---|---|
| **Run:** | 79492 | **Scans:** | 26184 - 26184 | **m/z Range:** | 329.15894 to 1885.2075: |
| **Search:** | 79492 | **Precursor Mass:** | 1002.50979 | | |
| **Mods:** | C=160.03 *+15.99 #+220.06 @+291.1 | **Charge:** | +4 | | |

**Sequence**

FT#KNNILT#QAAQAM*LAQANQQPQGVLQLLRLE

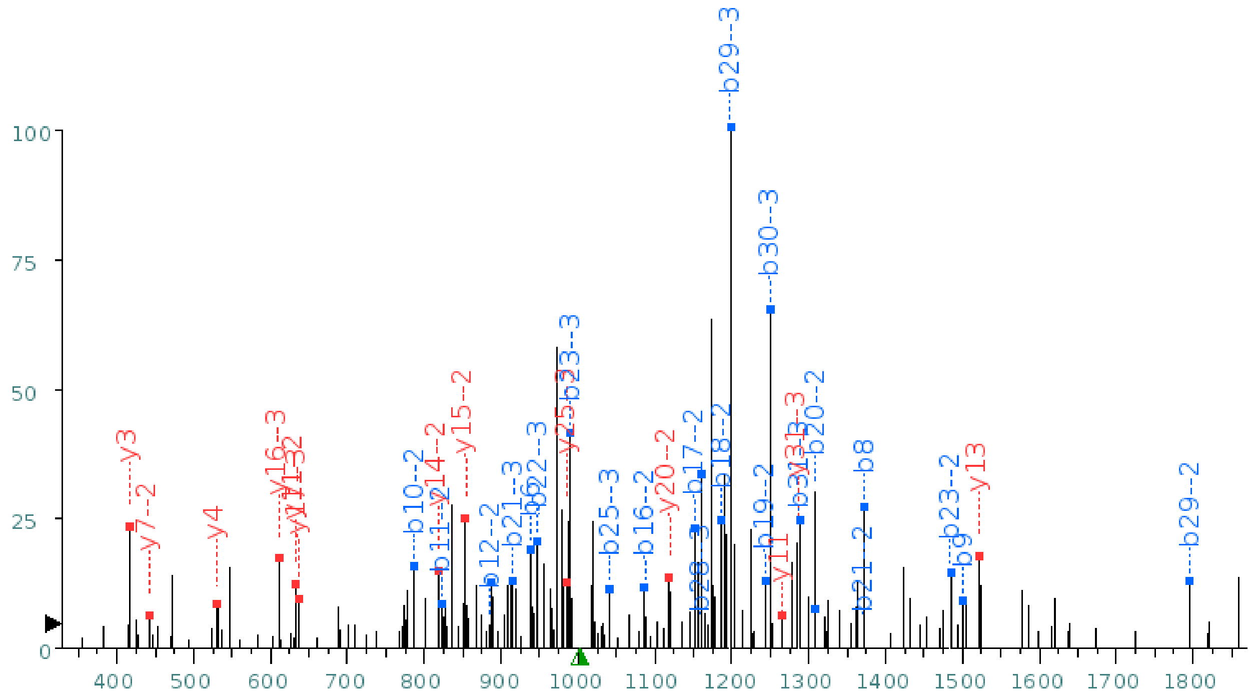

**Predicted Fragmentation Pattern**

+1

| Seq | # | b: Δ Error | b | y | y: Δ Error | +1 |
|---|---|---|---|---|---|---|
| F | 1 | --- | 148.076 | --- | --- | 32 |
| T# | 2 | --- | 469.182 | 3859.959 | --- | 31 |
| K | 3 | --- | 597.277 | 3538.853 | --- | 30 |
| N | 4 | --- | 711.320 | 3410.758 | --- | 29 |
| N | 5 | --- | 825.362 | 3296.715 | --- | 28 |
| I | 6 | -246.184 | **938.447** | 3182.673 | --- | 27 |
| L | 7 | --- | 1051.531 | 3069.588 | --- | 26 |
| T# | 8 | -384.417 | **1372.637** | 2956.504 | --- | 25 |
| Q | 9 | -396.526 | **1500.695** | 2635.398 | --- | 24 |

| | | | | | | |
|---|---|---|---|---|---|---|
| A | 10 | --- | 1571.732 | 2507.340 | --- | 23 |
| A | 11 | --- | 1642.769 | 2436.303 | --- | 22 |
| Q | 12 | --- | 1770.828 | 2365.266 | --- | 21 |
| A | 13 | --- | 1841.865 | 2237.207 | --- | 20 |
| M* | 14 | --- | 1988.900 | 2166.170 | --- | 19 |
| L | 15 | --- | 2101.985 | 2019.135 | --- | 18 |
| A | 16 | --- | 2173.022 | 1906.050 | --- | 17 |
| Q | 17 | --- | 2301.080 | 1835.013 | --- | 16 |
| A | 18 | --- | 2372.117 | 1706.955 | --- | 15 |
| N | 19 | --- | 2486.160 | 1635.918 | --- | 14 |
| Q | 20 | --- | 2614.219 | **1521.875** | **112.364** | 13 |
| Q | 21 | --- | 2742.277 | 1393.816 | --- | 12 |
| P | 22 | --- | 2839.330 | **1265.758** | **256.732** | 11 |
| Q | 23 | --- | 2967.389 | 1168.705 | --- | 10 |
| G | 24 | --- | 3024.410 | 1040.646 | --- | 9 |
| V | 25 | --- | 3123.479 | 983.625 | --- | 8 |
| L | 26 | --- | 3236.563 | 884.556 | --- | 7 |
| Q | 27 | --- | 3364.621 | 771.472 | --- | 6 |
| L | 28 | --- | 3477.705 | 643.414 | --- | 5 |
| L | 29 | --- | 3590.789 | **530.330** | **-1408.758** | 4 |
| R | 30 | --- | 3746.891 | **417.246** | **500.314** | 3 |
| L | 31 | --- | 3859.975 | 261.144 | --- | 2 |
| E | 32 | --- | --- | 148.060 | --- | 1 |

+2

| Seq | # | b: Δ Error | b | y | y: Δ Error | +1 |
|---|---|---|---|---|---|---|
| F | 1 | --- | 74.541 | --- | --- | 32 |
| T# | 2 | --- | 235.094 | 1930.483 | --- | 31 |
| K | 3 | --- | 299.142 | 1769.930 | --- | 30 |
| N | 4 | --- | 356.163 | 1705.883 | --- | 29 |
| N | 5 | --- | 413.185 | 1648.861 | --- | 28 |
| I | 6 | --- | 469.727 | 1591.840 | --- | 27 |
| L | 7 | --- | 526.269 | 1535.298 | --- | 26 |
| T# | 8 | --- | 686.822 | 1478.756 | --- | 25 |
| Q | 9 | --- | 750.851 | 1318.203 | --- | 24 |
| A | 10 | **-519.560** | **786.370** | 1254.174 | --- | 23 |
| A | 11 | **280.446** | **821.888** | 1218.655 | --- | 22 |
| Q | 12 | **783.933** | **885.918** | 1183.136 | --- | 21 |
| A | 13 | --- | 921.436 | **1119.107** | **-747.847** | 20 |
| M* | 14 | --- | 994.954 | 1083.589 | --- | 19 |
| L | 15 | --- | 1051.496 | 1010.071 | --- | 18 |
| A | 16 | **-257.067** | **1087.014** | 953.529 | --- | 17 |
| Q | 17 | **618.273** | **1151.044** | 918.010 | --- | 16 |
| A | 18 | **-383.004** | **1186.562** | **853.981** | **-139.766** | 15 |
| N | 19 | **309.763** | **1243.584** | **818.462** | **1076.029** | 14 |
| Q | 20 | **-130.643** | **1307.613** | 761.441 | --- | 13 |
| Q | 21 | **340.186** | **1371.642** | 697.412 | --- | 12 |
| P | 22 | --- | 1420.169 | **633.382** | **543.404** | 11 |
| Q | 23 | **563.052** | **1484.198** | 584.856 | --- | 10 |
| G | 24 | --- | 1512.709 | 520.827 | --- | 9 |
| V | 25 | --- | 1562.243 | 492.316 | --- | 8 |
| L | 26 | --- | 1618.785 | **442.782** | **-733.522** | 7 |
| Q | 27 | --- | 1682.814 | 386.240 | --- | 6 |
| L | 28 | --- | 1739.356 | 322.211 | --- | 5 |
| L | 29 | **315.948** | **1795.898** | 265.668 | --- | 4 |
| R | 30 | --- | 1873.949 | 209.126 | --- | 3 |
| L | 31 | --- | 1930.491 | 131.076 | --- | 2 |
| E | 32 | --- | --- | 74.534 | --- | 1 |

+3

| Seq | # | b: Δ Error | b | y | y: Δ Error | +1 |
|---|---|---|---|---|---|---|
| F | 1 | --- | 50.030 | --- | --- | 32 |
| T# | 2 | --- | 157.065 | **1287.325** | **574.555** | 31 |
| K | 3 | --- | 199.764 | 1180.289 | --- | 30 |
| N | 4 | --- | 237.778 | 1137.591 | --- | 29 |
| N | 5 | --- | 275.792 | 1099.577 | --- | 28 |
| I | 6 | --- | 313.487 | 1061.562 | --- | 27 |
| L | 7 | --- | 351.182 | 1023.868 | --- | 26 |
| T# | 8 | --- | 458.217 | **986.173** | **-139.169** | 25 |
| Q | 9 | --- | 500.903 | 879.138 | --- | 24 |
| A | 10 | --- | 524.582 | 836.451 | --- | 23 |
| A | 11 | --- | 548.261 | 812.772 | --- | 22 |
| Q | 12 | --- | 590.948 | 789.093 | --- | 21 |
| A | 13 | --- | 614.627 | 746.407 | --- | 20 |
| M* | 14 | --- | 663.638 | 722.728 | --- | 19 |
| L | 15 | --- | 701.333 | 673.716 | --- | 18 |
| A | 16 | --- | 725.012 | **636.022** | **340.058** | 17 |
| Q | 17 | --- | 767.698 | **612.343** | **-1388.785** | 16 |
| A | 18 | --- | 791.377 | 569.656 | --- | 15 |
| N | 19 | --- | 829.392 | 545.977 | --- | 14 |
| Q | 20 | --- | 872.078 | 507.963 | --- | 13 |
| Q | 21 | **781.552** | **914.764** | 465.277 | --- | 12 |
| P | 22 | **405.388** | **947.115** | 422.591 | --- | 11 |
| Q | 23 | **-852.764** | **989.801** | 390.240 | --- | 10 |
| G | 24 | --- | 1008.808 | 347.554 | --- | 9 |
| V | 25 | **92.189** | **1041.831** | 328.546 | --- | 8 |
| L | 26 | --- | 1079.526 | 295.524 | --- | 7 |
| Q | 27 | --- | 1122.212 | 257.829 | --- | 6 |
| L | 28 | **771.647** | **1159.907** | 215.143 | --- | 5 |
| L | 29 | **507.749** | **1197.601** | 177.448 | --- | 4 |
| R | 30 | **503.588** | **1249.635** | 139.753 | --- | 3 |
| L | 31 | **570.607** | **1287.330** | 87.720 | --- | 2 |
| E | 32 | --- | --- | 50.025 | --- | 1 |

**Soleil Index**

| | | |
|---|---|---|
| **547.0785** | **not explained** | **15%** |
| **835.1460** | **y2-15 frag loss of water** | **28%** |
| **955.1144** | **not explained** | **16%** |
| **971.7736** | **b3-23 frag loss of water** | **23%** |
| **972.6251** | **b3-23 frag loss of water/ b3-23 frag loss of ammonia** | **58%** |
| **978.7194** | **b2-14 frag loss of ammonia** | **27%** |
| **988.0920** | **not explained** | **25%** |
| **1018.9606** | **not explained** | **24%** |
| **1156.0173** | **not explained** | **17%** |
| **1173.8466** | **not explained** | **63%** |
| **1191.5934** | **not explained** | **34%** |
| **1192.4285** | **not explained** | **17%** |
| **1193.4783** | **not explained** | **22%** |

| | | |
|---|---|---|
| 1203.5258 | not explained | 20% |
| 1224.7196 | b2-19 frag loss of water | 23% |
| 1277.2906 | not explained | 16% |
| 1284.7776 | not explained | 20% |
| 1308.7411 | not explained | 30% |
| 1423.2242 | not explained | 15% |

| | | | | | |
|---|---|---|---|---|---|
| **Run:** | 79492 | **Scans:** | 26339 - 26339 | **m/z Range:** | 330.38681 to 1814.9265 |
| **Search:** | 79492 | **Precursor Mass:** | 998.51195 | | |
| **Mods:** | C=160.03 *+15.99 #+220.06 @+291.1 | **Charge:** | +4 | | |

**Sequence**

FT#KNNILT#QAAQAMLAQANQQPQGVLQLLRLE

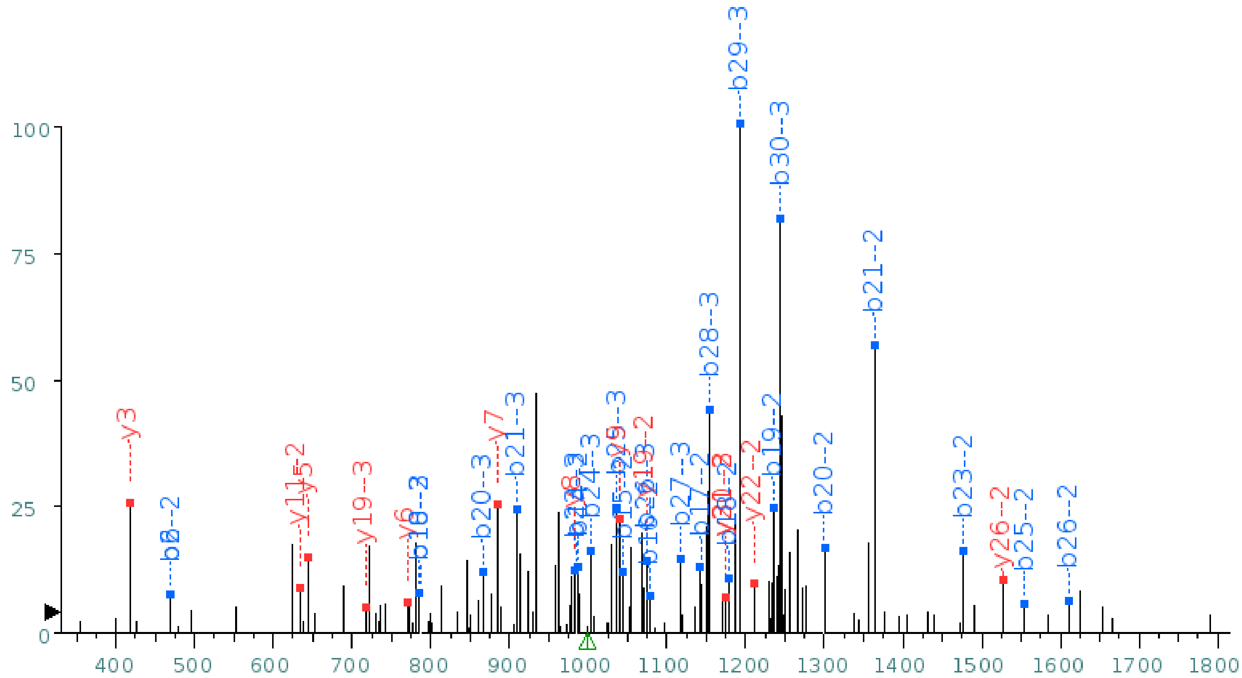

**Predicted Fragmentation Pattern**

**+1**

| Seq | # | b: Δ Error | b | y | y: Δ Error | +1 |
|---|---|---|---|---|---|---|
| F | 1 | --- | 148.076 | --- | --- | 32 |
| T# | 2 | 1506.694 | **469.182** | 3843.964 | --- | 31 |
| K | 3 | --- | 597.277 | 3522.858 | --- | 30 |
| N | 4 | --- | 711.320 | 3394.763 | --- | 29 |
| N | 5 | --- | 825.362 | 3280.721 | --- | 28 |
| I | 6 | --- | 938.447 | 3166.678 | --- | 27 |
| L | 7 | --- | 1051.531 | 3053.594 | --- | 26 |
| T# | 8 | --- | 1372.637 | 2940.510 | --- | 25 |
| Q | 9 | --- | 1500.695 | 2619.404 | --- | 24 |
| A | 10 | --- | 1571.732 | 2491.345 | --- | 23 |
| A | 11 | --- | 1642.769 | 2420.308 | --- | 22 |
| Q | 12 | --- | 1770.828 | 2349.271 | --- | 21 |
| A | 13 | --- | 1841.865 | 2221.212 | --- | 20 |

| | | | | | | |
|---|---|---|---|---|---|---|
| M | 14 | --- | 1972.906 | 2150.175 | --- | 19 |
| L | 15 | --- | 2085.990 | 2019.135 | --- | 18 |
| A | 16 | --- | 2157.027 | 1906.050 | --- | 17 |
| Q | 17 | --- | 2285.085 | 1835.013 | --- | 16 |
| A | 18 | --- | 2356.122 | 1706.955 | --- | 15 |
| N | 19 | --- | 2470.165 | 1635.918 | --- | 14 |
| Q | 20 | --- | 2598.224 | 1521.875 | --- | 13 |
| Q | 21 | --- | 2726.283 | 1393.816 | --- | 12 |
| P | 22 | --- | 2823.335 | 1265.758 | --- | 11 |
| Q | 23 | --- | 2951.394 | 1168.705 | --- | 10 |
| G | 24 | --- | 3008.415 | **1040.646** | 177.314 | 9 |
| V | 25 | --- | 3107.484 | **983.625** | 108.912 | 8 |
| L | 26 | --- | 3220.568 | **884.556** | 291.257 | 7 |
| Q | 27 | --- | 3348.626 | **771.472** | -69.342 | 6 |
| L | 28 | --- | 3461.710 | **643.414** | 13.596 | 5 |
| L | 29 | --- | 3574.795 | 530.330 | --- | 4 |
| R | 30 | --- | 3730.896 | **417.246** | 211.614 | 3 |
| L | 31 | --- | 3843.980 | 261.144 | --- | 2 |
| E | 32 | --- | --- | 148.060 | --- | 1 |

**+2**

| Seq | # | b: Δ Error | b | y | y: Δ Error | +1 |
|---|---|---|---|---|---|---|
| F | 1 | --- | 74.541 | --- | --- | 32 |
| T# | 2 | --- | 235.094 | 1922.486 | --- | 31 |
| K | 3 | --- | 299.142 | 1761.933 | --- | 30 |
| N | 4 | --- | 356.163 | 1697.885 | --- | 29 |
| N | 5 | --- | 413.185 | 1640.864 | --- | 28 |
| I | 6 | 346.327 | **469.727** | 1583.842 | --- | 27 |
| L | 7 | --- | 526.269 | **1527.300** | -305.015 | 26 |
| T# | 8 | --- | 686.822 | 1470.758 | --- | 25 |
| Q | 9 | --- | 750.851 | 1310.205 | --- | 24 |
| A | 10 | -349.277 | **786.370** | 1246.176 | --- | 23 |
| A | 11 | --- | 821.888 | **1210.658** | -139.841 | 22 |
| Q | 12 | --- | 885.918 | **1175.139** | 188.741 | 21 |
| A | 13 | --- | 921.436 | 1111.110 | --- | 20 |
| M | 14 | 510.425 | **986.956** | **1075.591** | -445.282 | 19 |
| L | 15 | 742.475 | **1043.498** | 1010.071 | --- | 18 |
| A | 16 | 332.226 | **1079.017** | 953.529 | --- | 17 |
| Q | 17 | -788.152 | **1143.046** | 918.010 | --- | 16 |
| A | 18 | 77.849 | **1178.565** | 853.981 | --- | 15 |
| N | 19 | -60.186 | **1235.586** | 818.462 | --- | 14 |
| Q | 20 | 482.173 | **1299.616** | 761.441 | --- | 13 |
| Q | 21 | 414.114 | **1363.645** | 697.412 | --- | 12 |
| P | 22 | --- | 1412.171 | **633.382** | 278.236 | 11 |
| Q | 23 | 76.064 | **1476.201** | 584.856 | --- | 10 |
| G | 24 | --- | 1504.711 | 520.827 | --- | 9 |
| V | 25 | 131.755 | **1554.246** | 492.316 | --- | 8 |
| L | 26 | 501.845 | **1610.788** | 442.782 | --- | 7 |
| Q | 27 | --- | 1674.817 | 386.240 | --- | 6 |
| L | 28 | --- | 1731.359 | 322.211 | --- | 5 |
| L | 29 | --- | 1787.901 | 265.668 | --- | 4 |
| R | 30 | --- | 1865.951 | 209.126 | --- | 3 |
| L | 31 | --- | 1922.493 | 131.076 | --- | 2 |
| E | 32 | --- | --- | 74.534 | --- | 1 |

**+3**

| Seq | # | b: Δ Error | b | y | y: Δ Error | +1 |
|---|---|---|---|---|---|---|
| F | 1 | --- | 50.030 | --- | --- | 32 |

| | | | | | | |
|---|---|---|---|---|---|---|
| T# | 2 | --- | 157.065 | 1281.993 | --- | 31 |
| K | 3 | --- | 199.764 | **1174.958** | **343.019** | 30 |
| N | 4 | --- | 237.778 | 1132.259 | --- | 29 |
| N | 5 | --- | 275.792 | 1094.245 | --- | 28 |
| I | 6 | --- | 313.487 | 1056.231 | --- | 27 |
| L | 7 | --- | 351.182 | 1018.536 | --- | 26 |
| T# | 8 | --- | 458.217 | 980.841 | --- | 25 |
| Q | 9 | --- | 500.903 | 873.806 | --- | 24 |
| A | 10 | --- | 524.582 | 831.120 | --- | 23 |
| A | 11 | --- | 548.261 | 807.441 | --- | 22 |
| Q | 12 | --- | 590.948 | 783.762 | --- | 21 |
| A | 13 | --- | 614.627 | 741.076 | --- | 20 |
| M | 14 | --- | 658.307 | **717.397** | **167.770** | 19 |
| L | 15 | --- | 696.001 | 673.716 | --- | 18 |
| A | 16 | --- | 719.680 | 636.022 | --- | 17 |
| Q | 17 | --- | 762.367 | 612.343 | --- | 16 |
| A | 18 | **63.033** | **786.046** | 569.656 | --- | 15 |
| N | 19 | --- | 824.060 | 545.977 | --- | 14 |
| Q | 20 | **-215.330** | **866.746** | 507.963 | --- | 13 |
| Q | 21 | **715.738** | **909.432** | 465.277 | --- | 12 |
| P | 22 | --- | 941.783 | 422.591 | --- | 11 |
| Q | 23 | **-749.736** | **984.469** | 390.240 | --- | 10 |
| G | 24 | **555.677** | **1003.477** | 347.554 | --- | 9 |
| V | 25 | **61.668** | **1036.499** | 328.546 | --- | 8 |
| L | 26 | **854.149** | **1074.194** | 295.524 | --- | 7 |
| Q | 27 | **172.601** | **1116.880** | 257.829 | --- | 6 |
| L | 28 | **73.538** | **1154.575** | 215.143 | --- | 5 |
| L | 29 | **179.521** | **1192.270** | 177.448 | --- | 4 |
| R | 30 | **-115.715** | **1244.303** | 139.753 | --- | 3 |
| L | 31 | --- | 1281.998 | 87.720 | --- | 2 |
| E | 32 | --- | --- | 50.025 | --- | 1 |

**Soleil Index**

| | | |
|---|---|---|
| 624.9415 | y1-5 frag loss of water | 17% |
| 721.4227 | not explained | 17% |
| 780.4904 | not explained | 18% |
| 913.1621 | not explained | 15% |
| 933.9122 | b2-30 frag doubly charged | 47% |
| 962.0579 | b2-31 frag doubly charged/ y2-31 frag loss of water/ y3-25 frag loss of water | 24% |
| 1030.4661 | not explained | 18% |
| 1054.7072 | not explained | 17% |
| 1068.7847 | not explained | 20% |
| 1149.5195 | not explained | 21% |
| 1151.3103 | y1-10 frag loss of water/ y1-10 frag loss of ammonia | 28% |
| 1187.2024 | not explained | 22% |
| 1256.7515 | not explained | 16% |
| 1266.8890 | not explained | 20% |
| 1355.8065 | b1-8 frag loss of ammonia | 18% |

| Run: | 79492 | Scans: | 26833 - 26833 | m/z Range: | 357.2475 to 1951.9144 |
|---|---|---|---|---|---|
| Search: | 79492 | Precursor Mass: | 1002.50997 | | |
| Mods: | C=160.03 *+15.99 #+220.06 @+291.1 | Charge: | +4 | | |

**Sequence**

FT#KNNILT#QAAQAM*LAQANQQPQGVLQLLRLE

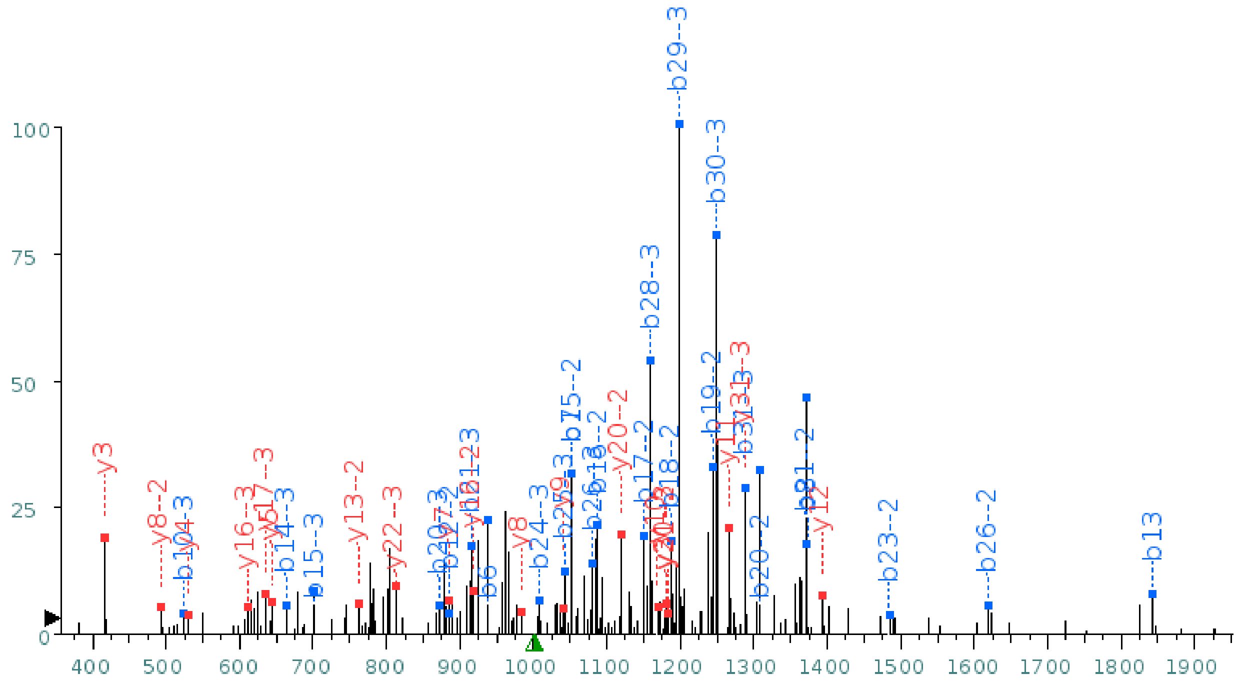

**Predicted Fragmentation Pattern**

+1

| Seq | # | b: Δ Error | b | y | y: Δ Error | +1 |
|---|---|---|---|---|---|---|
| F | 1 | --- | 148.076 | --- | --- | 32 |
| T# | 2 | --- | 469.182 | 3859.959 | --- | 31 |
| K | 3 | --- | 597.277 | 3538.853 | --- | 30 |
| N | 4 | --- | 711.320 | 3410.758 | --- | 29 |
| N | 5 | --- | 825.362 | 3296.715 | --- | 28 |
| I | 6 | -887.801 | **938.447** | 3182.673 | --- | 27 |
| L | 7 | -82.521 | **1051.531** | 3069.588 | --- | 26 |
| T# | 8 | -196.485 | **1372.637** | 2956.504 | --- | 25 |
| Q | 9 | --- | 1500.695 | 2635.398 | --- | 24 |
| A | 10 | --- | 1571.732 | 2507.340 | --- | 23 |
| A | 11 | --- | 1642.769 | 2436.303 | --- | 22 |

| | | | | | | |
|---|---|---|---|---|---|---|
| Q | 12 | --- | 1770.828 | 2365.266 | --- | 21 |
| A | 13 | 154.742 | **1841.865** | 2237.207 | --- | 20 |
| M* | 14 | --- | 1988.900 | 2166.170 | --- | 19 |
| L | 15 | --- | 2101.985 | 2019.135 | --- | 18 |
| A | 16 | --- | 2173.022 | 1906.050 | --- | 17 |
| Q | 17 | --- | 2301.080 | 1835.013 | --- | 16 |
| A | 18 | --- | 2372.117 | 1706.955 | --- | 15 |
| N | 19 | --- | 2486.160 | 1635.918 | --- | 14 |
| Q | 20 | --- | 2614.219 | 1521.875 | --- | 13 |
| Q | 21 | --- | 2742.277 | **1393.816** | 57.002 | 12 |
| P | 22 | --- | 2839.330 | **1265.758** | 128.033 | 11 |
| Q | 23 | --- | 2967.389 | **1168.705** | 538.146 | 10 |
| G | 24 | --- | 3024.410 | **1040.646** | 256.576 | 9 |
| V | 25 | --- | 3123.479 | **983.625** | 890.721 | 8 |
| L | 26 | --- | 3236.563 | **884.556** | 217.877 | 7 |
| Q | 27 | --- | 3364.621 | 771.472 | --- | 6 |
| L | 28 | --- | 3477.705 | **643.414** | 31.998 | 5 |
| L | 29 | --- | 3590.789 | **530.330** | 202.588 | 4 |
| R | 30 | --- | 3746.891 | **417.246** | -20.929 | 3 |
| L | 31 | --- | 3859.975 | 261.144 | --- | 2 |
| E | 32 | --- | --- | 148.060 | --- | 1 |

**+2**

| Seq | # | b: Δ Error | b | y | y: Δ Error | +1 |
|---|---|---|---|---|---|---|
| F | 1 | --- | 74.541 | --- | --- | 32 |
| T# | 2 | --- | 235.094 | 1930.483 | --- | 31 |
| K | 3 | --- | 299.142 | 1769.930 | --- | 30 |
| N | 4 | --- | 356.163 | 1705.883 | --- | 29 |
| N | 5 | --- | 413.185 | 1648.861 | --- | 28 |
| I | 6 | --- | 469.727 | 1591.840 | --- | 27 |
| L | 7 | --- | 526.269 | 1535.298 | --- | 26 |
| T# | 8 | --- | 686.822 | 1478.756 | --- | 25 |
| Q | 9 | --- | 750.851 | 1318.203 | --- | 24 |
| A | 10 | --- | 786.370 | 1254.174 | --- | 23 |
| A | 11 | --- | 821.888 | 1218.655 | --- | 22 |
| Q | 12 | 17.748 | **885.918** | **1183.136** | 479.342 | 21 |
| A | 13 | --- | 921.436 | **1119.107** | -204.556 | 20 |
| M* | 14 | --- | 994.954 | 1083.589 | --- | 19 |
| L | 15 | -49.519 | **1051.496** | 1010.071 | --- | 18 |
| A | 16 | 80.563 | **1087.014** | 953.529 | --- | 17 |
| Q | 17 | -432.930 | **1151.044** | **918.010** | -230.305 | 16 |
| A | 18 | 296.148 | **1186.562** | 853.981 | --- | 15 |
| N | 19 | 167.499 | **1243.584** | 818.462 | --- | 14 |
| Q | 20 | 89.494 | **1307.613** | **761.441** | 882.109 | 13 |
| Q | 21 | -162.283 | **1371.642** | 697.412 | --- | 12 |
| P | 22 | --- | 1420.169 | 633.382 | --- | 11 |
| Q | 23 | 155.813 | **1484.198** | 584.856 | --- | 10 |
| G | 24 | --- | 1512.709 | 520.827 | --- | 9 |
| V | 25 | --- | 1562.243 | **492.316** | 143.059 | 8 |
| L | 26 | -5.333 | **1618.785** | 442.782 | --- | 7 |
| Q | 27 | --- | 1682.814 | 386.240 | --- | 6 |
| L | 28 | --- | 1739.356 | 322.211 | --- | 5 |
| L | 29 | --- | 1795.898 | 265.668 | --- | 4 |
| R | 30 | --- | 1873.949 | 209.126 | --- | 3 |
| L | 31 | --- | 1930.491 | 131.076 | --- | 2 |
| E | 32 | --- | --- | 74.534 | --- | 1 |

**+3**

| Seq | # | b: Δ Error | b | y | y: Δ Error | +1 |
|---|---|---|---|---|---|---|
| F | 1 | --- | 50.030 | --- | --- | 32 |
| T# | 2 | --- | 157.065 | **1287.325** | **293.075** | 31 |
| K | 3 | --- | 199.764 | **1180.289** | **-377.641** | 30 |
| N | 4 | --- | 237.778 | 1137.591 | --- | 29 |
| N | 5 | --- | 275.792 | 1099.577 | --- | 28 |
| I | 6 | --- | 313.487 | 1061.562 | --- | 27 |
| L | 7 | --- | 351.182 | 1023.868 | --- | 26 |
| T# | 8 | --- | 458.217 | 986.173 | --- | 25 |
| Q | 9 | --- | 500.903 | 879.138 | --- | 24 |
| A | 10 | **-299.581** | **524.582** | 836.451 | --- | 23 |
| A | 11 | --- | 548.261 | **812.772** | **884.169** | 22 |
| Q | 12 | --- | 590.948 | 789.093 | --- | 21 |
| A | 13 | --- | 614.627 | 746.407 | --- | 20 |
| M* | 14 | **1183.025** | **663.638** | 722.728 | --- | 19 |
| L | 15 | **686.745** | **701.333** | 673.716 | --- | 18 |
| A | 16 | --- | 725.012 | **636.022** | **609.844** | 17 |
| Q | 17 | --- | 767.698 | **612.343** | **140.937** | 16 |
| A | 18 | --- | 791.377 | 569.656 | --- | 15 |
| N | 19 | --- | 829.392 | 545.977 | --- | 14 |
| Q | 20 | **-491.051** | **872.078** | 507.963 | --- | 13 |
| Q | 21 | **253.792** | **914.764** | 465.277 | --- | 12 |
| P | 22 | --- | 947.115 | 422.591 | --- | 11 |
| Q | 23 | --- | 989.801 | 390.240 | --- | 10 |
| G | 24 | **-185.814** | **1008.808** | 347.554 | --- | 9 |
| V | 25 | **88.323** | **1041.831** | 328.546 | --- | 8 |
| L | 26 | **-451.272** | **1079.526** | 295.524 | --- | 7 |
| Q | 27 | --- | 1122.212 | 257.829 | --- | 6 |
| L | 28 | **52.599** | **1159.907** | 215.143 | --- | 5 |
| L | 29 | **21.398** | **1197.601** | 177.448 | --- | 4 |
| R | 30 | **24.599** | **1249.635** | 139.753 | --- | 3 |
| L | 31 | **289.126** | **1287.330** | 87.720 | --- | 2 |
| E | 32 | --- | --- | 50.025 | --- | 1 |

**Soleil Index**

| | | |
|---|---|---|
| 804.6559 | b2-11 frag loss of water/ b2-11 frag loss of ammonia | 17% |
| 925.1050 | not explained | 19% |
| 961.9150 | not explained | 24% |
| 965.6118 | b2-31 frag doubly charged/ y1-8 frag loss of water/ y1-8 frag loss of ammonia/ y2-31 frag loss of water | 16% |
| 1085.2695 | not explained | 19% |
| 1238.7474 | not explained | 20% |
| 1266.9963 | not explained | 22% |

| | | | | | |
|---|---|---|---|---|---|
| **Run:** | 79492 | **Scans:** | 29491 - 29491 | **m/z Range:** | 268.17917 to 1978.1909 |
| **Search:** | 79492 | **Precursor Mass:** | 998.51341 | | |
| **Mods:** | C=160.03 *+15.99 #+220.06 @+291.1 | **Charge:** | +4 | | |

**Sequence**

FT#KNNILT#QAAQAMLAQANQQPQGVLQLLRLE

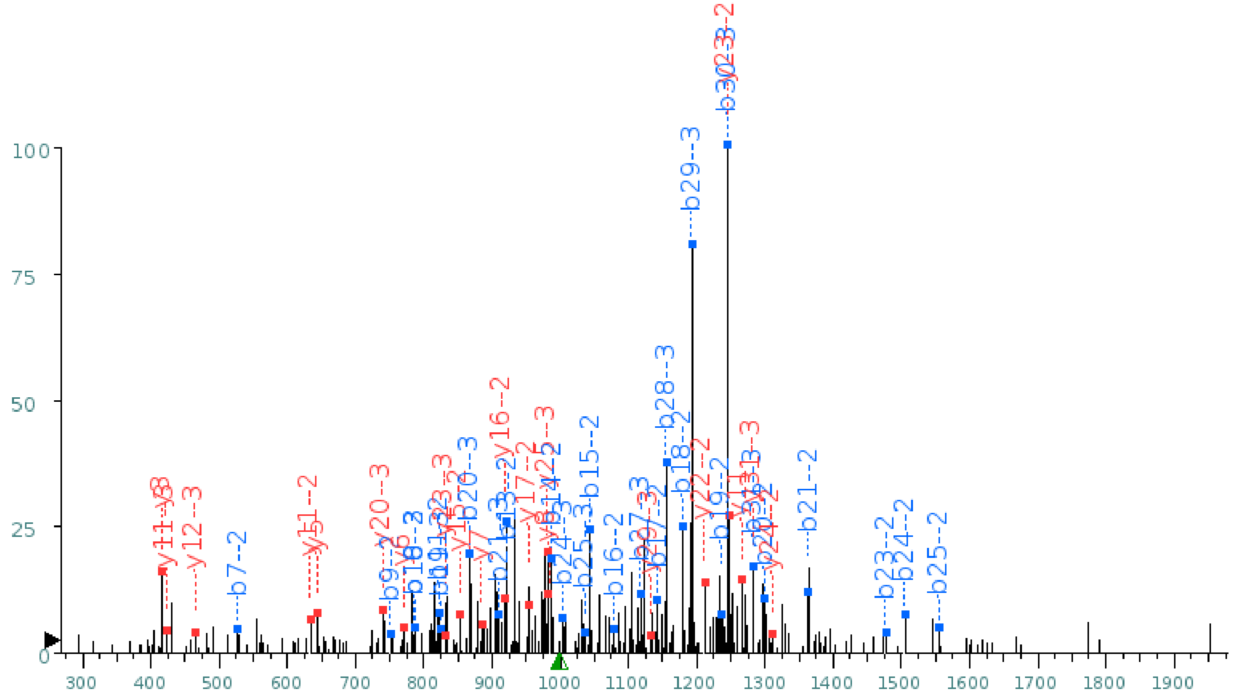

**Predicted Fragmentation Pattern**

| Seq | # | b: Δ Error | b (+1) | y (+1) | y: Δ Error | +1 |
|---|---|---|---|---|---|---|
| F | 1 | --- | 148.076 | --- | --- | 32 |
| T# | 2 | --- | 469.182 | 3843.964 | --- | 31 |
| K | 3 | --- | 597.277 | 3522.858 | --- | 30 |
| N | 4 | --- | 711.320 | 3394.763 | --- | 29 |
| N | 5 | --- | 825.362 | 3280.721 | --- | 28 |
| I | 6 | --- | 938.447 | 3166.678 | --- | 27 |
| L | 7 | --- | 1051.531 | 3053.594 | --- | 26 |
| T# | 8 | --- | 1372.637 | 2940.510 | --- | 25 |
| Q | 9 | --- | 1500.695 | 2619.404 | --- | 24 |
| A | 10 | --- | 1571.732 | 2491.345 | --- | 23 |

| | | | | | | |
|---|---|---|---|---|---|---|
| A | 11 | --- | 1642.769 | 2420.308 | --- | 22 |
| Q | 12 | --- | 1770.828 | 2349.271 | --- | 21 |
| A | 13 | --- | 1841.865 | 2221.212 | --- | 20 |
| M | 14 | --- | 1972.906 | 2150.175 | --- | 19 |
| L | 15 | --- | 2085.990 | 2019.135 | --- | 18 |
| A | 16 | --- | 2157.027 | 1906.050 | --- | 17 |
| Q | 17 | --- | 2285.085 | 1835.013 | --- | 16 |
| A | 18 | --- | 2356.122 | 1706.955 | --- | 15 |
| N | 19 | --- | 2470.165 | 1635.918 | --- | 14 |
| Q | 20 | --- | 2598.224 | 1521.875 | --- | 13 |
| Q | 21 | --- | 2726.283 | 1393.816 | --- | 12 |
| P | 22 | --- | 2823.335 | **1265.758** | **242.466** | 11 |
| Q | 23 | --- | 2951.394 | 1168.705 | --- | 10 |
| G | 24 | --- | 3008.415 | 1040.646 | --- | 9 |
| V | 25 | --- | 3107.484 | **983.625** | **-983.897** | 8 |
| L | 26 | --- | 3220.568 | **884.556** | **283.119** | 7 |
| Q | 27 | --- | 3348.626 | **771.472** | **393.173** | 6 |
| L | 28 | --- | 3461.710 | **643.414** | **-273.719** | 5 |
| L | 29 | --- | 3574.795 | 530.330 | --- | 4 |
| R | 30 | --- | 3730.896 | **417.246** | **24.418** | 3 |
| L | 31 | --- | 3843.980 | 261.144 | --- | 2 |
| E | 32 | --- | --- | 148.060 | --- | 1 |

+2

| Seq | # | b: Δ Error | b | y | y: Δ Error | +1 |
|---|---|---|---|---|---|---|
| F | 1 | --- | 74.541 | --- | --- | 32 |
| T# | 2 | --- | 235.094 | 1922.486 | --- | 31 |
| K | 3 | --- | 299.142 | 1761.933 | --- | 30 |
| N | 4 | --- | 356.163 | 1697.885 | --- | 29 |
| N | 5 | --- | 413.185 | 1640.864 | --- | 28 |
| I | 6 | --- | 469.727 | 1583.842 | --- | 27 |
| L | 7 | **385.313** | **526.269** | 1527.300 | --- | 26 |
| T# | 8 | --- | 686.822 | 1470.758 | --- | 25 |
| Q | 9 | **1203.689** | **750.851** | **1310.205** | **487.439** | 24 |
| A | 10 | **306.454** | **786.370** | **1246.176** | **551.508** | 23 |
| A | 11 | **-597.165** | **821.888** | **1210.658** | **631.228** | 22 |
| Q | 12 | --- | 885.918 | 1175.139 | --- | 21 |
| A | 13 | **-101.251** | **921.436** | 1111.110 | --- | 20 |
| M | 14 | **43.778** | **986.956** | 1075.591 | --- | 19 |
| L | 15 | **211.884** | **1043.498** | 1010.071 | --- | 18 |
| A | 16 | **200.272** | **1079.017** | **953.529** | **49.849** | 17 |
| Q | 17 | **-471.537** | **1143.046** | **918.010** | **-368.946** | 16 |
| A | 18 | **-87.874** | **1178.565** | **853.981** | **-560.025** | 15 |
| N | 19 | **206.622** | **1235.586** | 818.462 | --- | 14 |
| Q | 20 | **114.195** | **1299.616** | 761.441 | --- | 13 |
| Q | 21 | **211.215** | **1363.645** | 697.412 | --- | 12 |
| P | 22 | --- | 1412.171 | **633.382** | **147.237** | 11 |
| Q | 23 | **536.708** | **1476.201** | 584.856 | --- | 10 |
| G | 24 | **600.049** | **1504.711** | 520.827 | --- | 9 |
| V | 25 | **453.894** | **1554.246** | 492.316 | --- | 8 |
| L | 26 | --- | 1610.788 | 442.782 | --- | 7 |
| Q | 27 | --- | 1674.817 | 386.240 | --- | 6 |
| L | 28 | --- | 1731.359 | 322.211 | --- | 5 |
| L | 29 | --- | 1787.901 | 265.668 | --- | 4 |
| R | 30 | --- | 1865.951 | 209.126 | --- | 3 |
| L | 31 | --- | 1922.493 | 131.076 | --- | 2 |
| E | 32 | --- | --- | 74.534 | --- | 1 |

| | | | +3 | | | |
|---|---|---|---|---|---|---|
| Seq | # | b: Δ Error | b | y | y: Δ Error | +1 |
| F | 1 | --- | 50.030 | --- | --- | 32 |
| T# | 2 | --- | 157.065 | **1281.993** | 654.430 | 31 |
| K | 3 | --- | 199.764 | 1174.958 | --- | 30 |
| N | 4 | --- | 237.778 | **1132.259** | 370.025 | 29 |
| N | 5 | --- | 275.792 | 1094.245 | --- | 28 |
| I | 6 | --- | 313.487 | 1056.231 | --- | 27 |
| L | 7 | --- | 351.182 | 1018.536 | --- | 26 |
| T# | 8 | --- | 458.217 | **980.841** | 883.920 | 25 |
| Q | 9 | --- | 500.903 | 873.806 | --- | 24 |
| A | 10 | --- | 524.582 | **831.120** | 518.831 | 23 |
| A | 11 | --- | 548.261 | 807.441 | --- | 22 |
| Q | 12 | --- | 590.948 | 783.762 | --- | 21 |
| A | 13 | --- | 614.627 | **741.076** | -521.784 | 20 |
| M | 14 | --- | 658.307 | 717.397 | --- | 19 |
| L | 15 | --- | 696.001 | 673.716 | --- | 18 |
| A | 16 | --- | 719.680 | 636.022 | --- | 17 |
| Q | 17 | --- | 762.367 | 612.343 | --- | 16 |
| A | 18 | 718.494 | **786.046** | 569.656 | --- | 15 |
| N | 19 | 7.213 | **824.060** | 545.977 | --- | 14 |
| Q | 20 | 595.585 | **866.746** | 507.963 | --- | 13 |
| Q | 21 | -170.348 | **909.432** | **465.277** | -1853.646 | 12 |
| P | 22 | --- | 941.783 | **422.591** | 1582.865 | 11 |
| Q | 23 | --- | 984.469 | 390.240 | --- | 10 |
| G | 24 | -265.692 | **1003.477** | 347.554 | --- | 9 |
| V | 25 | 550.357 | **1036.499** | 328.546 | --- | 8 |
| L | 26 | --- | 1074.194 | 295.524 | --- | 7 |
| Q | 27 | 758.536 | **1116.880** | 257.829 | --- | 6 |
| L | 28 | 489.870 | **1154.575** | 215.143 | --- | 5 |
| L | 29 | 657.258 | **1192.270** | 177.448 | --- | 4 |
| R | 30 | 625.666 | **1244.303** | 139.753 | --- | 3 |
| L | 31 | 650.466 | **1281.998** | 87.720 | --- | 2 |
| E | 32 | --- | --- | 50.025 | --- | 1 |

**Soleil Index**

| | | |
|---|---|---|
| 904.9119 | b2-13 frag loss of ammonia | 16% |
| 933.7770 | b2-30 frag doubly charged | 26% |
| 977.8904 | not explained | 20% |
| 1104.1003 | not explained | 16% |
| 1122.5192 | not explained | 23% |
| 1232.7235 | not explained | 15% |
| 1364.8042 | not explained | 17% |

| | | | | | |
|---|---|---|---|---|---|
| **Run:** | 79492 | **Scans:** | 29606 - 29606 | **m/z Range:** | 303.17358 to 1942.0692 |
| **Search:** | 79492 | **Precursor Mass:** | 998.51166 | | |
| **Mods:** | C=160.03 *+15.99 #+220.06 @+291.1 | **Charge:** | +4 | | |

**Sequence**

FT#KNNILT#QAAQAMLAQANQQPQGVLQLLRLE

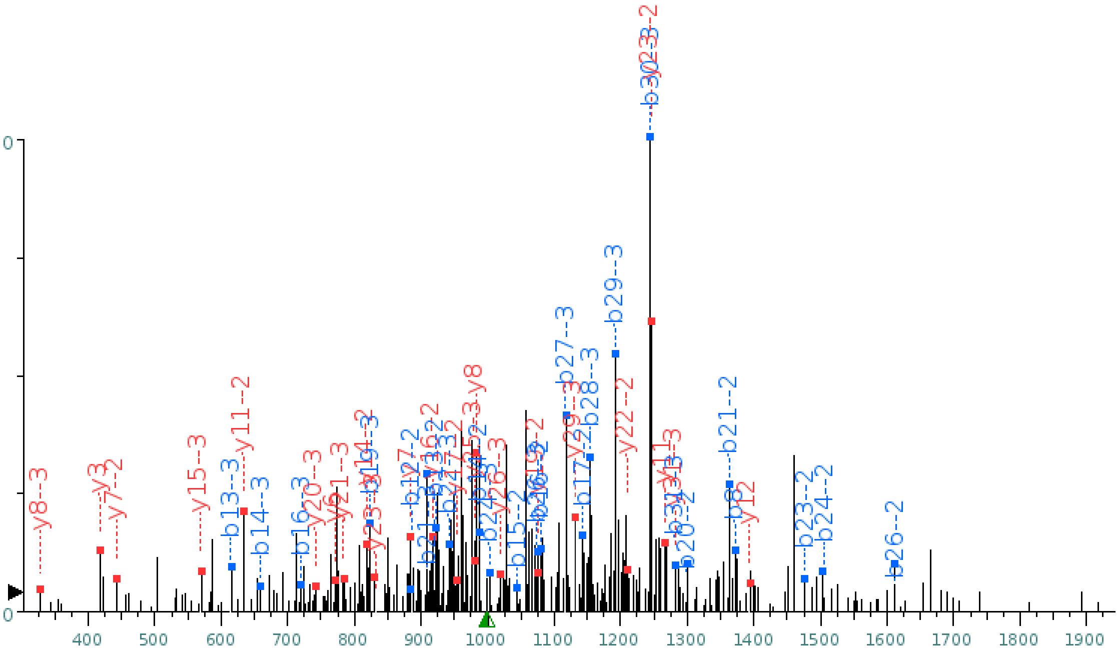

**Predicted Fragmentation Pattern**

| Seq | # | b: Δ Error | b | y | y: Δ Error | +1 |
|---|---|---|---|---|---|---|
| F | 1 | --- | 148.076 | --- | --- | 32 |
| T# | 2 | --- | 469.182 | 3843.964 | --- | 31 |
| K | 3 | --- | 597.277 | 3522.858 | --- | 30 |
| N | 4 | --- | 711.320 | 3394.763 | --- | 29 |
| N | 5 | --- | 825.362 | 3280.721 | --- | 28 |
| I | 6 | --- | 938.447 | 3166.678 | --- | 27 |
| L | 7 | --- | 1051.531 | 3053.594 | --- | 26 |
| T# | 8 | -344.636 | **1372.637** | 2940.510 | --- | 25 |
| Q | 9 | --- | 1500.695 | 2619.404 | --- | 24 |
| A | 10 | --- | 1571.732 | 2491.345 | --- | 23 |
| A | 11 | --- | 1642.769 | 2420.308 | --- | 22 |
| Q | 12 | --- | 1770.828 | 2349.271 | --- | 21 |
| A | 13 | --- | 1841.865 | 2221.212 | --- | 20 |
| M | 14 | --- | 1972.906 | 2150.175 | --- | 19 |

| | | | | | | |
|---|---|---|---|---|---|---|
| L | 15 | --- | 2085.990 | 2019.135 | --- | 18 |
| A | 16 | --- | 2157.027 | 1906.050 | --- | 17 |
| Q | 17 | --- | 2285.085 | 1835.013 | --- | 16 |
| A | 18 | --- | 2356.122 | 1706.955 | --- | 15 |
| N | 19 | --- | 2470.165 | 1635.918 | --- | 14 |
| Q | 20 | --- | 2598.224 | 1521.875 | --- | 13 |
| Q | 21 | --- | 2726.283 | **1393.816** | **290.759** | 12 |
| P | 22 | --- | 2823.335 | **1265.758** | **109.617** | 11 |
| Q | 23 | --- | 2951.394 | 1168.705 | --- | 10 |
| G | 24 | --- | 3008.415 | 1040.646 | --- | 9 |
| V | 25 | --- | 3107.484 | **983.625** | **-914.143** | 8 |
| L | 26 | --- | 3220.568 | **884.556** | **-465.539** | 7 |
| Q | 27 | --- | 3348.626 | **771.472** | **349.534** | 6 |
| L | 28 | --- | 3461.710 | 643.414 | --- | 5 |
| L | 29 | --- | 3574.795 | 530.330 | --- | 4 |
| R | 30 | --- | 3730.896 | **417.246** | **373.160** | 3 |
| L | 31 | --- | 3843.980 | 261.144 | --- | 2 |
| E | 32 | --- | --- | 148.060 | --- | 1 |

**+2**

| Seq | # | b: Δ Error | b | y | y: Δ Error | +1 |
|---|---|---|---|---|---|---|
| F | 1 | --- | 74.541 | --- | --- | 32 |
| T# | 2 | --- | 235.094 | 1922.486 | --- | 31 |
| K | 3 | --- | 299.142 | 1761.933 | --- | 30 |
| N | 4 | --- | 356.163 | 1697.885 | --- | 29 |
| N | 5 | --- | 413.185 | 1640.864 | --- | 28 |
| I | 6 | --- | 469.727 | 1583.842 | --- | 27 |
| L | 7 | --- | 526.269 | 1527.300 | --- | 26 |
| T# | 8 | --- | 686.822 | 1470.758 | --- | 25 |
| Q | 9 | --- | 750.851 | 1310.205 | --- | 24 |
| A | 10 | --- | 786.370 | **1246.176** | **658.835** | 23 |
| A | 11 | --- | 821.888 | **1210.658** | **-379.336** | 22 |
| Q | 12 | **-1061.443** | **885.918** | 1175.139 | --- | 21 |
| A | 13 | **790.111** | **921.436** | 1111.110 | --- | 20 |
| M | 14 | **117.729** | **986.956** | **1075.591** | **191.566** | 19 |
| L | 15 | **803.678** | **1043.498** | 1010.071 | --- | 18 |
| A | 16 | **338.896** | **1079.017** | **953.529** | **574.658** | 17 |
| Q | 17 | **38.945** | **1143.046** | **918.010** | **794.867** | 16 |
| A | 18 | --- | 1178.565 | 853.981 | --- | 15 |
| N | 19 | --- | 1235.586 | **818.462** | **305.414** | 14 |
| Q | 20 | **296.434** | **1299.616** | 761.441 | --- | 13 |
| Q | 21 | **707.045** | **1363.645** | 697.412 | --- | 12 |
| P | 22 | --- | 1412.171 | **633.382** | **833.252** | 11 |
| Q | 23 | **-120.917** | **1476.201** | 584.856 | --- | 10 |
| G | 24 | **-363.086** | **1504.711** | 520.827 | --- | 9 |
| V | 25 | --- | 1554.246 | 492.316 | --- | 8 |
| L | 26 | **225.817** | **1610.788** | **442.782** | **872.559** | 7 |
| Q | 27 | --- | 1674.817 | 386.240 | --- | 6 |
| L | 28 | --- | 1731.359 | 322.211 | --- | 5 |
| L | 29 | --- | 1787.901 | 265.668 | --- | 4 |
| R | 30 | --- | 1865.951 | 209.126 | --- | 3 |
| L | 31 | --- | 1922.493 | 131.076 | --- | 2 |
| E | 32 | --- | --- | 74.534 | --- | 1 |

**+3**

| Seq | # | b: Δ Error | b | y | y: Δ Error | +1 |
|---|---|---|---|---|---|---|
| F | 1 | --- | 50.030 | --- | --- | 32 |
| T# | 2 | --- | 157.065 | **1281.993** | **47.739** | 31 |

| | | | | | | |
|---|---|---|---|---|---|---|
| K | 3 | --- | 199.764 | 1174.958 | --- | 30 |
| N | 4 | --- | 237.778 | 1132.259 | -709.424 | 29 |
| N | 5 | --- | 275.792 | 1094.245 | --- | 28 |
| I | 6 | --- | 313.487 | 1056.231 | --- | 27 |
| L | 7 | --- | 351.182 | 1018.536 | 728.000 | 26 |
| T# | 8 | --- | 458.217 | 980.841 | -259.975 | 25 |
| Q | 9 | --- | 500.903 | 873.806 | --- | 24 |
| A | 10 | --- | 524.582 | 831.120 | -688.976 | 23 |
| A | 11 | --- | 548.261 | 807.441 | --- | 22 |
| Q | 12 | --- | 590.948 | 783.762 | 902.634 | 21 |
| A | 13 | 1310.752 | 614.627 | 741.076 | 712.231 | 20 |
| M | 14 | 359.874 | 658.307 | 717.397 | --- | 19 |
| L | 15 | --- | 696.001 | 673.716 | --- | 18 |
| A | 16 | -280.307 | 719.680 | 636.022 | --- | 17 |
| Q | 17 | --- | 762.367 | 612.343 | --- | 16 |
| A | 18 | --- | 786.046 | 569.656 | 1222.166 | 15 |
| N | 19 | 210.703 | 824.060 | 545.977 | --- | 14 |
| Q | 20 | --- | 866.746 | 507.963 | --- | 13 |
| Q | 21 | 224.795 | 909.432 | 465.277 | --- | 12 |
| P | 22 | -149.811 | 941.783 | 422.591 | --- | 11 |
| Q | 23 | --- | 984.469 | 390.240 | --- | 10 |
| G | 24 | 727.344 | 1003.477 | 347.554 | --- | 9 |
| V | 25 | --- | 1036.499 | 328.546 | -1136.180 | 8 |
| L | 26 | 257.073 | 1074.194 | 295.524 | --- | 7 |
| Q | 27 | 643.827 | 1116.880 | 257.829 | --- | 6 |
| L | 28 | 13.279 | 1154.575 | 215.143 | --- | 5 |
| L | 29 | -220.410 | 1192.270 | 177.448 | --- | 4 |
| R | 30 | 87.266 | 1244.303 | 139.753 | --- | 3 |
| L | 31 | 43.773 | 1281.998 | 87.720 | --- | 2 |
| E | 32 | --- | --- | 50.025 | --- | 1 |

**Soleil Index**

| | | |
|---|---|---|
| 587.8917 | y2-21 frag doubly charged/ y3-30 frag loss of water | 15% |
| 713.5571 | not explained | 16% |
| 774.2305 | not explained | 26% |
| 849.6115 | b3-20 frag loss of water/ b3-20 frag loss of ammonia/ y2-29 frag loss of water | 16% |
| 923.8545 | b3-22 frag loss of water/ b3-22 frag loss of ammonia | 25% |
| 943.9597 | not explained | 20% |
| 948.4988 | not explained | 25% |
| 961.5885 | b2-31 frag doubly charged/ y2-31 frag loss of water | 40% |
| 962.5990 | y3-25 frag loss of water | 20% |
| 975.6805 | not explained | 36% |
| 988.4076 | not explained | 35% |
| 1027.3915 | b2-15 frag loss of ammonia | 35% |
| 1057.4579 | b3-26 frag loss of ammonia/ y2-19 frag loss of water | 43% |
| 1060.9703 | b2-16 frag loss of water | 17% |
| 1066.7499 | not explained | 17% |
| 1081.6608 | not explained | 16% |
| 1107.8544 | not explained | 19% |
| 1184.7782 | not explained | 17% |

| | | |
|---|---|---|
| **1196.6748** | **not explained** | **19%** |
| **1208.1476** | **not explained** | **20%** |
| **1252.7308** | **not explained** | **15%** |
| **1257.8872** | **not explained** | **15%** |
| **1269.7206** | **not explained** | **15%** |
| **1460.5885** | **not explained** | **33%** |

**MS/MS data of trypsin digested peptides with site score A and site score B above 19 from *Geobacillus kaustophilus* *Gk*F2 flagellin protein co-expressed with *Geobacillus kaustophilus* Maf in EV36 cells. Results of dataset 2 (MS/MS results obtained in April 2019).**

| | | | | | |
|---|---|---|---|---|---|
| **Run:** | 77653 | **Scans:** | 7366 - 7366 | **m/z Range:** | 104.09891 to 827.68073 |
| **Search:** | 77653 | **Precursor Mass:** | 426.71938 | **Hit:** | |
| **Mods:** | C=160.03 *+15.99 #+220.06 @+291.1 | **Charge:** | +2 | | |

**Sequence**

LSS#GLR

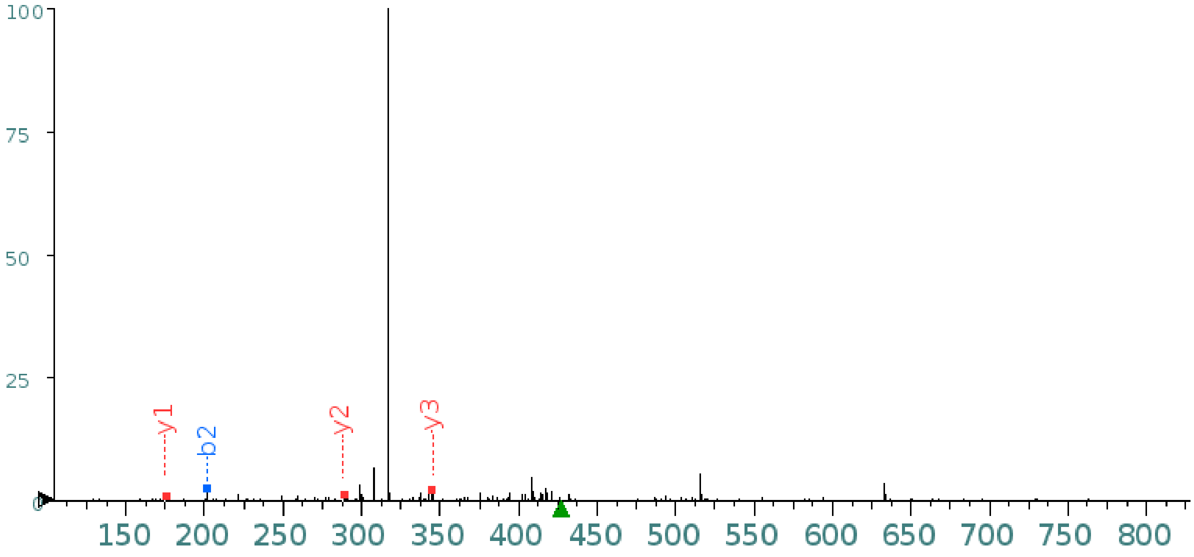

**Predicted Fragmentation Pattern**

| Seq | # | b: Δ Error | b | y | y: Δ Error | +1 |
|---|---|---|---|---|---|---|
| L | 1 | --- | 114.091 | --- | --- | 6 |
| S | 2 | 199.942 | 201.123 | 739.347 | --- | 5 |
| S# | 3 | --- | 508.214 | 652.315 | --- | 4 |
| G | 4 | --- | 565.235 | 345.224 | -2627.105 | 3 |
| L | 5 | --- | 678.319 | 288.203 | 3339.941 | 2 |
| R | 6 | --- | --- | 175.119 | 477.294 | 1 |

**Soleil Index**

| 316.7907 | not explained | 100% |
|---|---|---|

| | | | | | |
|---|---|---|---|---|---|
| **Run:** | 77653 | **Scans:** | 28301 - 28301 | **m/z Range:** | 329.23587 to 2000 |
| **Search:** | 77653 | **Precursor Mass:** | 1133.22205 | | |
| **Mods:** | C=160.03 *+15.99 #+220.06 @+291.1 | **Charge:** | +3 | | |

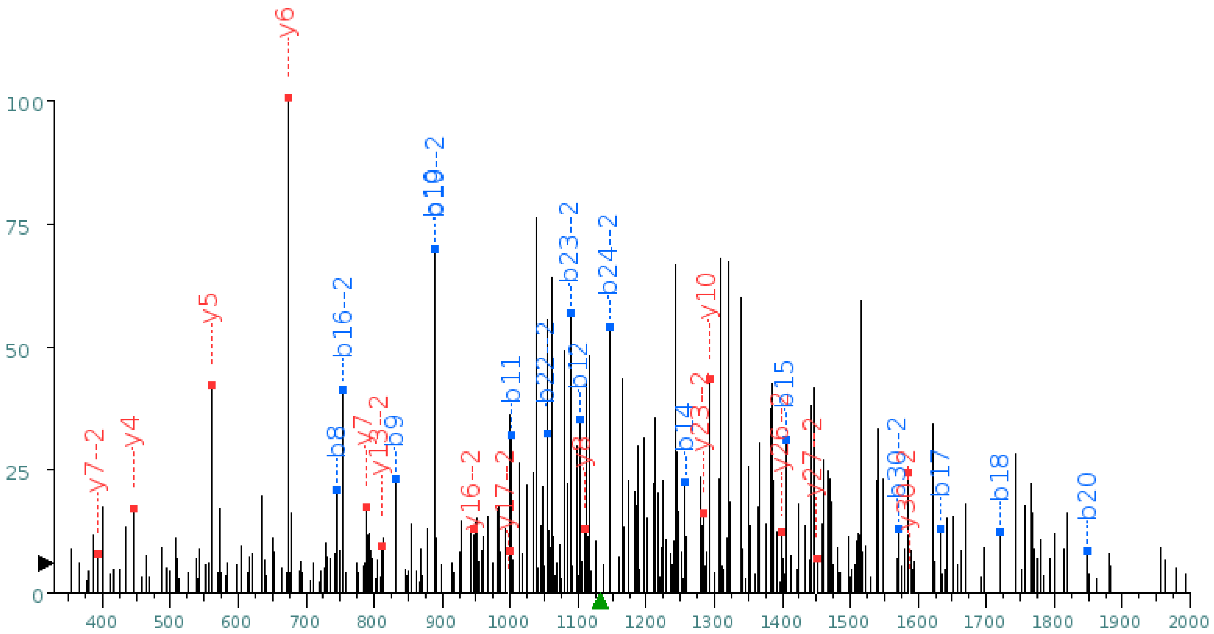

**Sequence**

NIALSTSGSGNVGVM*TQSGAEQAIT#IIDDAIK

**Predicted Fragmentation Pattern**

+1

| Seq | # | b: Δ Error | b | y | y: Δ Error | +1 |
|---|---|---|---|---|---|---|
| N | 1 | --- | 115.050 | --- | --- | 32 |
| I | 2 | --- | 228.134 | 3283.610 | --- | 31 |
| A | 3 | --- | 299.171 | 3170.526 | --- | 30 |
| L | 4 | --- | 412.255 | 3099.489 | --- | 29 |
| S | 5 | --- | 499.287 | 2986.404 | --- | 28 |
| T | 6 | --- | 600.335 | 2899.372 | --- | 27 |
| S | 7 | --- | 687.367 | 2798.325 | --- | 26 |
| G | 8 | -15.050 | **744.389** | 2711.293 | --- | 25 |
| S | 9 | 232.929 | **831.421** | 2654.271 | --- | 24 |
| G | 10 | 171.858 | **888.442** | 2567.239 | --- | 23 |
| N | 11 | -699.026 | **1002.485** | 2510.218 | --- | 22 |
| V | 12 | 256.465 | **1101.553** | 2396.175 | --- | 21 |

| | | | | | | |
|---|---|---|---|---|---|---|
| G | 13 | --- | 1158.575 | 2297.106 | --- | 20 |
| V | 14 | -476.647 | **1257.643** | 2240.085 | --- | 19 |
| M* | 15 | 168.184 | **1404.679** | 2141.017 | --- | 18 |
| T | 16 | --- | 1505.726 | 1993.981 | --- | 17 |
| Q | 17 | 79.133 | **1633.785** | 1892.934 | --- | 16 |
| S | 18 | 174.321 | **1720.817** | 1764.875 | --- | 15 |
| G | 19 | --- | 1777.839 | 1677.843 | --- | 14 |
| A | 20 | 114.997 | **1848.876** | 1620.821 | --- | 13 |
| E | 21 | --- | 1977.918 | 1549.784 | --- | 12 |
| Q | 22 | --- | 2105.977 | 1420.742 | --- | 11 |
| A | 23 | --- | 2177.014 | **1292.683** | -225.307 | 10 |
| I | 24 | --- | 2290.098 | 1221.646 | --- | 9 |
| T# | 25 | --- | 2611.204 | **1108.562** | 116.963 | 8 |
| I | 26 | --- | 2724.288 | **787.456** | 135.387 | 7 |
| I | 27 | --- | 2837.372 | **674.372** | 266.947 | 6 |
| D | 28 | --- | 2952.399 | **561.288** | 285.643 | 5 |
| D | 29 | --- | 3067.426 | **446.261** | 255.831 | 4 |
| A | 30 | --- | 3138.463 | 331.234 | --- | 3 |
| I | 31 | --- | 3251.547 | 260.197 | --- | 2 |
| K | 32 | --- | --- | 147.113 | --- | 1 |

+2

| Seq | # | b: Δ Error | b | y | y: Δ Error | +1 |
|---|---|---|---|---|---|---|
| N | 1 | --- | 58.029 | --- | --- | 32 |
| I | 2 | --- | 114.571 | 1642.309 | --- | 31 |
| A | 3 | --- | 150.089 | **1585.766** | -111.397 | 30 |
| L | 4 | --- | 206.631 | 1550.248 | --- | 29 |
| S | 5 | --- | 250.147 | 1493.706 | --- | 28 |
| T | 6 | --- | 300.671 | **1450.190** | 678.883 | 27 |
| S | 7 | --- | 344.187 | **1399.666** | 130.705 | 26 |
| G | 8 | --- | 372.698 | 1356.150 | --- | 25 |
| S | 9 | --- | 416.214 | 1327.639 | --- | 24 |
| G | 10 | --- | 444.725 | **1284.123** | -200.794 | 23 |
| N | 11 | --- | 501.746 | 1255.613 | --- | 22 |
| V | 12 | --- | 551.280 | 1198.591 | --- | 21 |
| G | 13 | --- | 579.791 | 1149.057 | --- | 20 |
| V | 14 | --- | 629.325 | 1120.546 | --- | 19 |
| M* | 15 | --- | 702.843 | 1071.012 | --- | 18 |
| T | 16 | -533.904 | **753.367** | **997.494** | 805.236 | 17 |
| Q | 17 | --- | 817.396 | **946.970** | 745.753 | 16 |
| S | 18 | --- | 860.912 | 882.941 | --- | 15 |
| G | 19 | -931.854 | **889.423** | 839.425 | --- | 14 |
| A | 20 | --- | 924.941 | **810.914** | 327.995 | 13 |
| E | 21 | --- | 989.463 | 775.396 | --- | 12 |
| Q | 22 | 243.652 | **1053.492** | 710.875 | --- | 11 |
| A | 23 | -70.370 | **1089.011** | 646.845 | --- | 10 |
| I | 24 | 296.995 | **1145.553** | 611.327 | --- | 9 |
| T# | 25 | --- | 1306.106 | 554.785 | --- | 8 |
| I | 26 | --- | 1362.648 | **394.232** | 8.031 | 7 |
| I | 27 | --- | 1419.190 | 337.690 | --- | 6 |
| D | 28 | --- | 1476.703 | 281.148 | --- | 5 |
| D | 29 | --- | 1534.217 | 223.634 | --- | 4 |
| A | 30 | 545.026 | **1569.735** | 166.121 | --- | 3 |
| I | 31 | --- | 1626.277 | 130.602 | --- | 2 |
| K | 32 | --- | --- | 74.060 | --- | 1 |

**Soleil Index**

| | | |
|---|---|---|
| 400.3422 | not explained | 17% |
| 572.7037 | b2-24 frag doubly charged | 17% |
| 634.4526 | not explained | 20% |
| 678.6941 | y2-25 frag doubly charged | 16% |
| 952.2849 | not explained | 15% |
| 966.6547 | not explained | 16% |
| 981.7639 | not explained | 17% |
| 983.5768 | b1-11 frag loss of water | 17% |
| 999.6215 | not explained | 36% |
| 1013.7067 | not explained | 26% |
| 1024.9003 | not explained | 22% |
| 1032.8022 | not explained | 24% |
| 1039.4886 | not explained | 76% |
| 1048.2371 | not explained | 22% |
| 1054.5570 | y2-18 frag loss of ammonia | 56% |
| 1060.5729 | not explained | 64% |
| 1080.5975 | not explained | 49% |

| | | |
|---|---|---|
| 1085.1891 | b1-12 frag loss of ammonia | 22% |
| 1098.8114 | not explained | 30% |
| 1112.2335 | not explained | 43% |
| 1116.7936 | not explained | 48% |
| 1165.3162 | not explained | 44% |
| 1174.3590 | not explained | 23% |
| 1182.6287 | not explained | 21% |
| 1183.5221 | not explained | 16% |
| 1185.8057 | not explained | 18% |
| 1187.1268 | not explained | 30% |
| 1196.8732 | not explained | 31% |
| 1200.7753 | not explained | 15% |
| 1211.0319 | not explained | 22% |
| 1212.0079 | not explained | 36% |
| 1217.7496 | not explained | 20% |
| 1224.7831 | not explained | 23% |
| 1243.6436 | not explained | 67% |
| 1244.4615 | not explained | 29% |
| 1247.2484 | not explained | 16% |
| 1278.2660 | not explained | 24% |
| 1307.5482 | not explained | 23% |
| 1309.5314 | y2-24 frag loss of water | 68% |
| 1320.7664 | not explained | 67% |

| | | |
|---|---|---|
| 1322.0608 | not explained | 19% |
| 1340.0262 | y2-25 frag loss of ammonia | 60% |
| 1350.4004 | not explained | 26% |
| 1365.0099 | not explained | 17% |
| 1367.2200 | not explained | 31% |
| 1383.2891 | y2-26 frag loss of ammonia | 37% |
| 1384.3030 | not explained | 36% |
| 1385.9312 | b1-15 frag loss of water | 43% |
| 1386.8524 | b1-15 frag loss of water/ b1-15 frag loss of ammonia | 23% |
| 1425.0433 | not explained | 18% |
| 1441.3059 | not explained | 38% |
| 1446.6731 | not explained | 41% |
| 1459.9072 | b2-28 frag loss of ammonia | 22% |
| 1460.7430 | not explained | 27% |
| 1468.0458 | not explained | 25% |
| 1470.0258 | not explained | 23% |
| 1472.8055 | not explained | 18% |
| 1516.6953 | b2-29 frag loss of water/ b2-29 frag loss of ammonia | 59% |
| 1538.9541 | not explained | 23% |
| 1540.1377 | not explained | 28% |
| 1540.9956 | not explained | 33% |
| 1546.9185 | not explained | 23% |
| 1621.9996 | not explained | 34% |

| | | |
|---|---|---|
| 1641.1735 | y1-31 frag doubly charged | 15% |
| 1651.0929 | not explained | 16% |
| 1670.1460 | not explained | 18% |
| 1743.1216 | not explained | 28% |
| 1757.3615 | not explained | 18% |
| 1766.0074 | not explained | 22% |
| 1768.2130 | not explained | 16% |
| 1818.8267 | not explained | 16% |

| | | | | | |
|---|---|---|---|---|---|
| **Run:** | 77653 | **Scans:** | 31252 - 31252 | **m/z Range:** | 239.04694 to 1971.2579 |
| **Search:** | 77653 | **Precursor Mass:** | 863.92879 | | |
| **Mods:** | C=160.03 *+15.99 #+220.06 @+291.1 | **Charge:** | +4 | | |

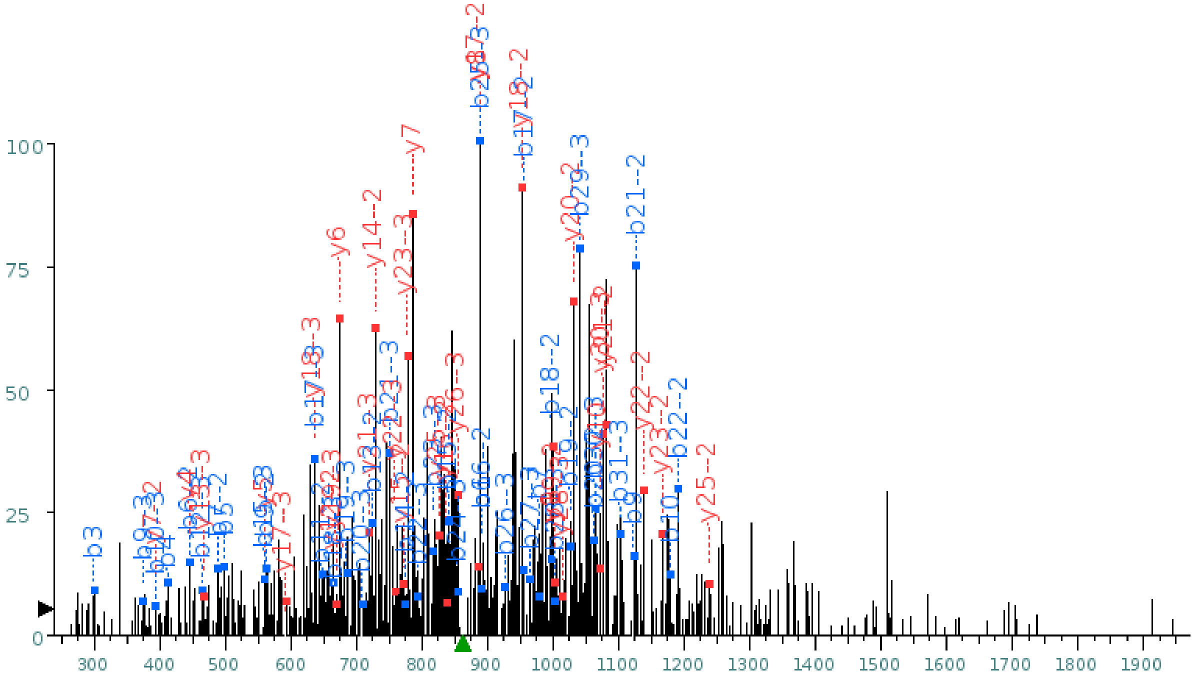

**Sequence**

NIALST@SGSGNVGVMTQSGAEQAITIIDDAIK

**Predicted Fragmentation Pattern**

| Seq | # | b: Δ Error | b | y | y: Δ Error | +1 |
|---|---|---|---|---|---|---|
| N | 1 | --- | 115.050 | --- | --- | 32 |
| I | 2 | --- | 228.134 | 3338.652 | --- | 31 |
| A | 3 | 3203.353 | **299.171** | 3225.568 | --- | 30 |
| L | 4 | 104.778 | **412.255** | 3154.531 | --- | 29 |
| S | 5 | 358.052 | **499.287** | 3041.447 | --- | 28 |
| T@ | 6 | -761.164 | **891.431** | 2954.415 | --- | 27 |
| S | 7 | 95.228 | **978.463** | 2562.272 | --- | 26 |
| G | 8 | --- | 1035.484 | 2475.240 | --- | 25 |
| S | 9 | 686.068 | **1122.516** | 2418.218 | --- | 24 |
| G | 10 | -225.313 | **1179.538** | 2331.186 | --- | 23 |
| N | 11 | --- | 1293.580 | 2274.165 | --- | 22 |
| V | 12 | --- | 1392.649 | 2160.122 | --- | 21 |
| G | 13 | --- | 1449.670 | 2061.053 | --- | 20 |
| V | 14 | --- | 1548.739 | 2004.032 | --- | 19 |

| | | | | | | |
|---|---|---|---|---|---|---|
| M | 15 | --- | 1679.779 | 1904.963 | --- | 18 |
| T | 16 | --- | 1780.827 | 1773.923 | --- | 17 |
| Q | 17 | --- | 1908.886 | 1672.875 | --- | 16 |
| S | 18 | --- | 1995.918 | 1544.817 | --- | 15 |
| G | 19 | --- | 2052.939 | 1457.785 | --- | 14 |
| A | 20 | --- | 2123.976 | 1400.763 | --- | 13 |
| E | 21 | --- | 2253.019 | 1329.726 | --- | 12 |
| Q | 22 | --- | 2381.077 | 1200.683 | --- | 11 |
| A | 23 | --- | 2452.114 | **1072.625** | -267.035 | 10 |
| I | 24 | --- | 2565.198 | **1001.588** | -416.048 | 9 |
| T | 25 | --- | 2666.246 | **888.504** | -4.617 | 8 |
| I | 26 | --- | 2779.330 | **787.456** | -374.358 | 7 |
| I | 27 | --- | 2892.414 | **674.372** | -26.043 | 6 |
| D | 28 | --- | 3007.441 | **561.288** | -133.164 | 5 |
| D | 29 | --- | 3122.468 | **446.261** | -125.502 | 4 |
| A | 30 | --- | 3193.505 | 331.234 | --- | 3 |
| I | 31 | --- | 3306.589 | 260.197 | --- | 2 |
| K | 32 | --- | --- | 147.113 | --- | 1 |

+2

| Seq | # | b: Δ Error | b | y | y: Δ Error | +1 |
|---|---|---|---|---|---|---|
| N | 1 | --- | 58.029 | --- | --- | 32 |
| I | 2 | --- | 114.571 | 1669.830 | --- | 31 |
| A | 3 | --- | 150.089 | 1613.288 | --- | 30 |
| L | 4 | --- | 206.631 | 1577.769 | --- | 29 |
| S | 5 | --- | 250.147 | 1521.227 | --- | 28 |
| T@ | 6 | -31.372 | **446.219** | 1477.711 | --- | 27 |
| S | 7 | -569.034 | **489.735** | 1281.639 | --- | 26 |
| G | 8 | --- | 518.246 | **1238.123** | -606.022 | 25 |
| S | 9 | 1061.092 | **561.762** | 1209.613 | --- | 24 |
| G | 10 | --- | 590.272 | **1166.097** | 441.684 | 23 |
| N | 11 | 1350.152 | **647.294** | **1137.586** | 150.104 | 22 |
| V | 12 | --- | 696.828 | **1080.564** | 122.994 | 21 |
| G | 13 | 157.902 | **725.339** | **1031.030** | 107.385 | 20 |
| V | 14 | 625.605 | **774.873** | **1002.520** | -733.248 | 19 |
| M | 15 | 995.355 | **840.393** | **952.985** | 255.509 | 18 |
| T | 16 | -184.724 | **890.917** | **887.465** | -1002.562 | 17 |
| Q | 17 | -562.814 | **954.946** | **836.941** | 664.334 | 16 |
| S | 18 | -82.967 | **998.462** | **772.912** | 157.245 | 15 |
| G | 19 | -534.935 | **1026.973** | **729.396** | 808.267 | 14 |
| A | 20 | -939.653 | **1062.492** | 700.885 | --- | 13 |
| E | 21 | 461.909 | **1127.013** | **665.367** | -402.239 | 12 |
| Q | 22 | -764.485 | **1191.042** | 600.845 | --- | 11 |
| A | 23 | --- | 1226.561 | 536.816 | --- | 10 |
| I | 24 | --- | 1283.103 | 501.298 | --- | 9 |
| T | 25 | --- | 1333.627 | 444.755 | --- | 8 |
| I | 26 | --- | 1390.169 | **394.232** | -1700.514 | 7 |
| I | 27 | --- | 1446.711 | 337.690 | --- | 6 |
| D | 28 | --- | 1504.224 | 281.148 | --- | 5 |
| D | 29 | --- | 1561.738 | 223.634 | --- | 4 |
| A | 30 | --- | 1597.256 | 166.121 | --- | 3 |
| I | 31 | --- | 1653.798 | 130.602 | --- | 2 |
| K | 32 | --- | --- | 74.060 | --- | 1 |

+3

| Seq | # | b: Δ Error | b | y | y: Δ Error | +1 |
|---|---|---|---|---|---|---|
| N | 1 | --- | 39.022 | --- | --- | 32 |
| I | 2 | --- | 76.716 | 1113.555 | --- | 31 |

| | | | | | | |
|---|---|---|---|---|---|---|
| A | 3 | --- | 100.395 | **1075.861** | -403.723 | 30 |
| L | 4 | --- | 138.090 | 1052.182 | --- | 29 |
| S | 5 | --- | 167.101 | **1014.487** | 269.377 | 28 |
| T@ | 6 | --- | 297.815 | 985.476 | --- | 27 |
| S | 7 | --- | 326.826 | **854.762** | -210.999 | 26 |
| G | 8 | --- | 345.833 | **825.751** | 720.638 | 25 |
| S | 9 | 1726.788 | **374.844** | 806.744 | --- | 24 |
| G | 10 | -732.603 | **393.851** | **777.734** | 594.179 | 23 |
| N | 11 | --- | 431.865 | **758.726** | 1061.138 | 22 |
| V | 12 | 1187.946 | **464.888** | **720.712** | -475.963 | 21 |
| G | 13 | --- | 483.895 | 687.689 | --- | 20 |
| V | 14 | --- | 516.918 | **668.682** | 1114.914 | 19 |
| M | 15 | 1096.192 | **560.598** | **635.659** | 805.802 | 18 |
| T | 16 | --- | 594.280 | **591.979** | 961.204 | 17 |
| Q | 17 | -1249.274 | **636.967** | 558.297 | --- | 16 |
| S | 18 | -1320.471 | **665.977** | 515.610 | --- | 15 |
| G | 19 | 931.487 | **684.985** | 486.600 | --- | 14 |
| A | 20 | 1026.465 | **708.664** | **467.593** | 1403.249 | 13 |
| E | 21 | 5.250 | **751.678** | 443.914 | --- | 12 |
| Q | 22 | -361.635 | **794.364** | 400.899 | --- | 11 |
| A | 23 | 432.164 | **818.043** | 358.213 | --- | 10 |
| I | 24 | 800.046 | **855.738** | 334.534 | --- | 9 |
| T | 25 | -1036.197 | **889.420** | 296.839 | --- | 8 |
| I | 26 | -763.254 | **927.115** | 263.157 | --- | 7 |
| I | 27 | 865.700 | **964.810** | 225.462 | --- | 6 |
| D | 28 | -184.762 | **1003.152** | 187.767 | --- | 5 |
| D | 29 | 85.108 | **1041.494** | 149.425 | --- | 4 |
| A | 30 | -410.726 | **1065.173** | 111.083 | --- | 3 |
| I | 31 | -235.632 | **1102.868** | 87.404 | --- | 2 |
| K | 32 | --- | --- | 49.709 | --- | 1 |

**Soleil Index**

**Index Threshold:** 15%

**Sort By:** m/z

**Show Matched Peaks:** ☐

**Persistent Index Display** ☑

**Matched 42 / 89**

| | | |
|---|---|---|
| 339.2101 | y3-10 frag loss of water | 19% |
| 581.6952 | not explained | 19% |
| 604.4764 | y2-24 frag doubly charged | 16% |
| 620.6768 | b3-17 frag loss of | 25% |

| | | |
|---|---|---|
| | ammonia | |
| 629.7234 | b2-11 frag loss of water/ b2-11 frag loss of ammonia | 34% |
| 642.6124 | not explained | 26% |
| 657.5443 | y1-6 frag loss of ammonia | 16% |
| 664.2943 | y1-12 frag doubly charged | 19% |
| 680.6846 | b2-12 frag loss of ammonia | 15% |
| 686.4811 | not explained | 20% |
| 694.6981 | b2-26 frag doubly charged | 24% |
| 724.0505 | b1-13 frag doubly charged/ b2-27 frag loss of water | 20% |
| 734.4321 | b3-21 frag loss of water/ b3-21 frag loss of ammonia | 19% |
| 738.0872 | y2-27 frag doubly charged | 22% |
| 739.7084 | y2-27 frag doubly charged | 24% |
| 746.2320 | not explained | 39% |
| 754.5956 | y2-15 frag loss of water | 18% |
| 761.7433 | not explained | 23% |
| 769.2066 | y1-7 frag loss of water | 22% |
| 789.5611 | y2-29 frag doubly charged/ y3-24 frag loss of water/ y3-24 | 22% |

| | | |
|---|---|---|
| | frag loss of ammonia | |
| 803.0963 | not explained | 24% |
| 808.2459 | y3-25 frag loss of water/ y3-25 frag loss of ammonia | 39% |
| 809.5438 | y3-25 frag loss of ammonia | 20% |
| 819.4664 | y2-16 frag loss of water/ y2-16 frag loss of ammonia | 23% |
| 821.6245 | b2-15 frag loss of water | 18% |
| 827.7681 | b2-31 frag doubly charged | 16% |
| 828.8359 | not explained | 40% |
| 831.0391 | not explained | 29% |
| 832.0342 | not explained | 18% |
| 834.6539 | y2-31 frag doubly charged | 33% |
| 835.6160 | y1-16 frag doubly charged/ y2-31 frag loss of water | 18% |
| 838.7943 | b3-24 frag loss of ammonia | 20% |
| 842.2485 | not explained | 18% |
| 844.6829 | not explained | 28% |
| 845.7574 | not explained | 36% |
| 846.5587 | not explained | 62% |
| 847.6492 | not explained | 34% |

| | | |
|---|---|---|
| 850.5516 | not explained | 35% |
| 851.8336 | not explained | 28% |
| 853.1442 | not explained | 16% |
| 881.0270 | not explained | 20% |
| 892.9222 | not explained | 19% |
| 900.7787 | not explained | 38% |
| 912.0353 | not explained | 21% |
| 913.8035 | not explained | 25% |
| 915.6275 | not explained | 17% |
| 932.3270 | not explained | 16% |
| 938.4174 | b2-17 frag loss of ammonia | 37% |
| 941.6540 | not explained | 60% |
| 942.4926 | not explained | 37% |
| 947.0243 | b3-27 frag loss of water/ b3-27 frag loss of ammonia | 22% |
| 968.7609 | y3-27 frag loss of ammonia | 19% |
| 971.3627 | not explained | 22% |
| 980.1561 | b2-18 frag loss of water | 21% |
| 980.9012 | b2-18 frag loss of water/ b2-18 frag loss of ammonia | 17% |
| 982.9576 | y1-9 frag loss of water | 29% |
| 988.3249 | not explained | 38% |

| | | |
|---|---|---|
| 995.5878 | y3-28 frag loss of water | 20% |
| 996.9288 | y3-28 frag loss of water/ y3-28 frag loss of ammonia | 49% |
| 1000.2099 | not explained | 32% |
| 1005.4450 | not explained | 15% |
| 1008.8560 | b2-19 frag loss of water | 17% |
| 1018.3328 | b1-8 frag loss of water/ b1-8 frag loss of ammonia | 20% |
| 1020.1891 | not explained | 15% |
| 1025.6105 | b1-19 frag doubly charged | 23% |
| 1032.1215 | not explained | 49% |
| 1033.1647 | not explained | 19% |
| 1049.3724 | not explained | 40% |
| 1053.6354 | y1-10 frag loss of water | 35% |
| 1054.5778 | y1-10 frag loss of water | 67% |
| 1057.3927 | y3-30 frag loss of water | 20% |
| 1071.0062 | not explained | 25% |
| 1081.8273 | not explained | 72% |
| 1083.8282 | not explained | 19% |
| 1084.8444 | b3-31 frag loss of water/ b3-31 frag loss of ammonia | 16% |
| 1098.0072 | not explained | 22% |
| 1101.5873 | not explained | 24% |

| | | |
|---|---|---|
| 1104.4371 | b1-9 frag loss of water | 15% |
| 1149.8982 | y2-23 frag loss of ammonia | 19% |
| 1173.6932 | b2-22 frag loss of water/ b2-22 frag loss of ammonia | 25% |
| 1174.7681 | b2-22 frag loss of ammonia | 21% |
| 1175.5372 | not explained | 23% |
| 1251.6647 | not explained | 18% |
| 1257.8439 | not explained | 23% |
| 1258.7036 | not explained | 19% |
| 1302.3744 | not explained | 23% |
| 1366.8638 | not explained | 19% |
| 1508.6676 | not explained | 29% |
| 1509.7100 | not explained | 22% |

| | | | |
|---|---|---|---|
| **Run:** | 77653 | **Scans:** | 32193 - 32193 |
| **Search:** | 77653 | **Precursor Mass:** | 1133.22144 |
| **Mods:** | C=160.03 *+15.99 #+220.06 @+291.1 | **Charge:** | +3 |

**m/z Range:** 316.32294 **to** 2000

**Sequence**

NIALSTSGSGNVGVM*TQSGAEQAIT#IIDDAIK

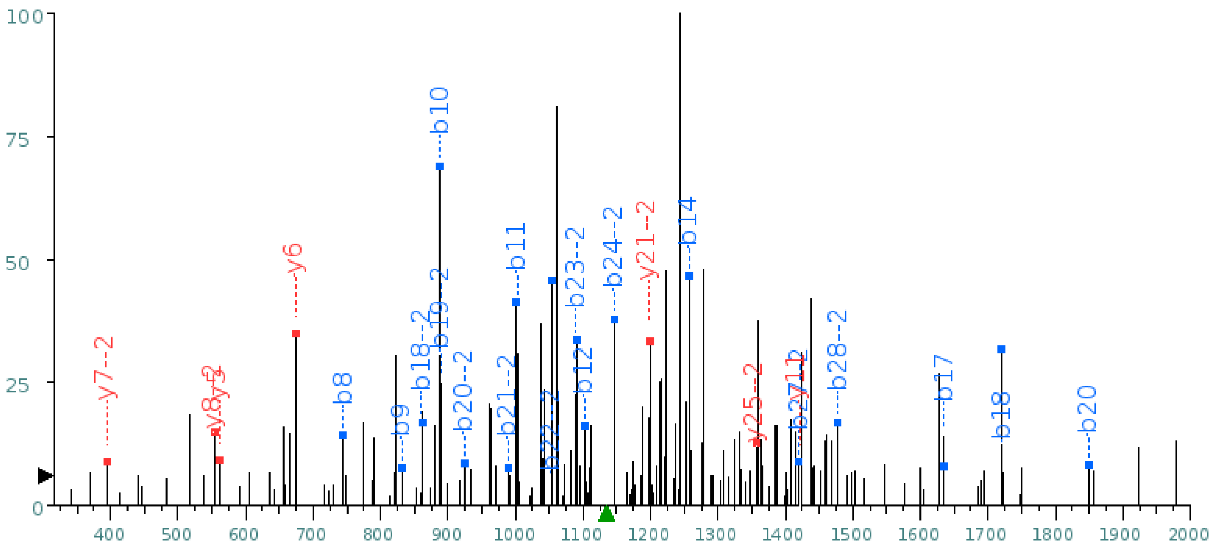

**Predicted Fragmentation Pattern**

+1

| Seq | # | b: Δ Error | b | y | y: Δ Error | +1 |
|---|---|---|---|---|---|---|
| N | 1 | --- | 115.050 | --- | --- | 32 |
| I | 2 | --- | 228.134 | 3283.610 | --- | 31 |
| A | 3 | --- | 299.171 | 3170.526 | --- | 30 |
| L | 4 | --- | 412.255 | 3099.489 | --- | 29 |
| S | 5 | --- | 499.287 | 2986.404 | --- | 28 |
| T | 6 | --- | 600.335 | 2899.372 | --- | 27 |
| S | 7 | --- | 687.367 | 2798.325 | --- | 26 |
| G | 8 | 422.454 | **744.389** | 2711.293 | --- | 25 |
| S | 9 | 54.446 | **831.421** | 2654.271 | --- | 24 |
| G | 10 | 77.352 | **888.442** | 2567.239 | --- | 23 |
| N | 11 | -703.172 | **1002.485** | 2510.218 | --- | 22 |
| V | 12 | 252.920 | **1101.553** | 2396.175 | --- | 21 |
| G | 13 | --- | 1158.575 | 2297.106 | --- | 20 |
| V | 14 | 251.876 | **1257.643** | 2240.085 | --- | 19 |
| M* | 15 | --- | 1404.679 | 2141.017 | --- | 18 |
| T | 16 | --- | 1505.726 | 1993.981 | --- | 17 |
| Q | 17 | 149.798 | **1633.785** | 1892.934 | --- | 16 |
| S | 18 | -558.247 | **1720.817** | 1764.875 | --- | 15 |
| G | 19 | --- | 1777.839 | 1677.843 | --- | 14 |
| A | 20 | 185.556 | **1848.876** | 1620.821 | --- | 13 |
| E | 21 | --- | 1977.918 | 1549.784 | --- | 12 |

| | | | | | | |
|---|---|---|---|---|---|---|
| Q | 22 | --- | 2105.977 | 1420.742 | -501.617 | 11 |
| A | 23 | --- | 2177.014 | 1292.683 | --- | 10 |
| I | 24 | --- | 2290.098 | 1221.646 | --- | 9 |
| T# | 25 | --- | 2611.204 | 1108.562 | --- | 8 |
| I | 26 | --- | 2724.288 | 787.456 | --- | 7 |
| I | 27 | --- | 2837.372 | 674.372 | 242.342 | 6 |
| D | 28 | --- | 2952.399 | 561.288 | 287.817 | 5 |
| D | 29 | --- | 3067.426 | 446.261 | --- | 4 |
| A | 30 | --- | 3138.463 | 331.234 | --- | 3 |
| I | 31 | --- | 3251.547 | 260.197 | --- | 2 |
| K | 32 | --- | --- | 147.113 | --- | 1 |

+2

| Seq | # | b: Δ Error | b | y | y: Δ Error | +1 |
|---|---|---|---|---|---|---|
| N | 1 | --- | 58.029 | --- | --- | 32 |
| I | 2 | --- | 114.571 | 1642.309 | --- | 31 |
| A | 3 | --- | 150.089 | 1585.766 | --- | 30 |
| L | 4 | --- | 206.631 | 1550.248 | --- | 29 |
| S | 5 | --- | 250.147 | 1493.706 | --- | 28 |
| T | 6 | --- | 300.671 | 1450.190 | --- | 27 |
| S | 7 | --- | 344.187 | 1399.666 | --- | 26 |
| G | 8 | --- | 372.698 | 1356.150 | 534.932 | 25 |
| S | 9 | --- | 416.214 | 1327.639 | --- | 24 |
| G | 10 | --- | 444.725 | 1284.123 | --- | 23 |
| N | 11 | --- | 501.746 | 1255.613 | --- | 22 |
| V | 12 | --- | 551.280 | 1198.591 | 99.585 | 21 |
| G | 13 | --- | 579.791 | 1149.057 | --- | 20 |
| V | 14 | --- | 629.325 | 1120.546 | --- | 19 |
| M* | 15 | --- | 702.843 | 1071.012 | --- | 18 |
| T | 16 | --- | 753.367 | 997.494 | --- | 17 |
| Q | 17 | --- | 817.396 | 946.970 | --- | 16 |
| S | 18 | 229.027 | 860.912 | 882.941 | --- | 15 |
| G | 19 | -1026.465 | 889.423 | 839.425 | --- | 14 |
| A | 20 | 804.496 | 924.941 | 810.914 | --- | 13 |
| E | 21 | -898.546 | 989.463 | 775.396 | --- | 12 |
| Q | 22 | 945.118 | 1053.492 | 710.875 | --- | 11 |
| A | 23 | 884.555 | 1089.011 | 646.845 | --- | 10 |
| I | 24 | 74.899 | 1145.553 | 611.327 | --- | 9 |
| T# | 25 | --- | 1306.106 | 554.785 | 1548.021 | 8 |
| I | 26 | --- | 1362.648 | 394.232 | 1692.011 | 7 |
| I | 27 | 591.358 | 1419.190 | 337.690 | --- | 6 |
| D | 28 | -90.208 | 1476.703 | 281.148 | --- | 5 |
| D | 29 | --- | 1534.217 | 223.634 | --- | 4 |
| A | 30 | --- | 1569.735 | 166.121 | --- | 3 |
| I | 31 | --- | 1626.277 | 130.602 | --- | 2 |
| K | 32 | --- | --- | 74.060 | --- | 1 |

Soleil Index

| | | |
|---|---|---|
| 516.7615 | not explained | 18% |
| 656.6475 | y1-6 frag loss of water/ y1-6 frag loss of ammonia | 16% |
| 773.7426 | not explained | 17% |

| | | |
|---|---|---|
| 823.7278 | not explained | 30% |
| 862.5526 | not explained | 19% |
| 880.6052 | not explained | 16% |
| 961.9327 | not explained | 21% |
| 962.8955 | not explained | 20% |
| 1003.7264 | not explained | 16% |
| 1039.0194 | not explained | 37% |
| 1042.7610 | not explained | 24% |
| 1060.3083 | not explained | 81% |
| 1063.0272 | not explained | 22% |
| 1112.6901 | not explained | 16% |
| 1146.5573 | not explained | 18% |
| 1188.3944 | not explained | 20% |
| 1196.7966 | not explained | 18% |
| 1214.5536 | not explained | 25% |
| 1215.3962 | not explained | 26% |
| 1216.4098 | not explained | 16% |
| 1222.7367 | not explained | 48% |
| 1236.0978 | not explained | 17% |
| 1243.9165 | not explained | 100% |
| 1252.5463 | not explained | 21% |
| 1278.6207 | not explained | 48% |
| 1358.2587 | not explained | 37% |

| | | |
|---|---|---|
| 1384.3392 | not explained | 16% |
| 1386.8163 | b1-15 frag loss of water/ b1-15 frag loss of ammonia | 16% |
| 1408.1117 | not explained | 17% |
| 1415.8605 | not explained | 15% |
| 1424.2997 | not explained | 31% |
| 1437.7247 | not explained | 42% |
| 1627.7214 | not explained | 27% |

| Run: | 77653 | Scans: | 33488 - 33488 | m/z Range: | 359.23511 to 2000 |
|---|---|---|---|---|---|
| Search: | 77653 | Precursor Mass: | 1133.22207 | | |
| Mods: | C=160.03 *+15.99 #+220.06 @+291.1 | Charge: | +3 | | |

**Sequence**

NIALST#SGSGNVGVM*TQSGAEQAITIIDDAIK

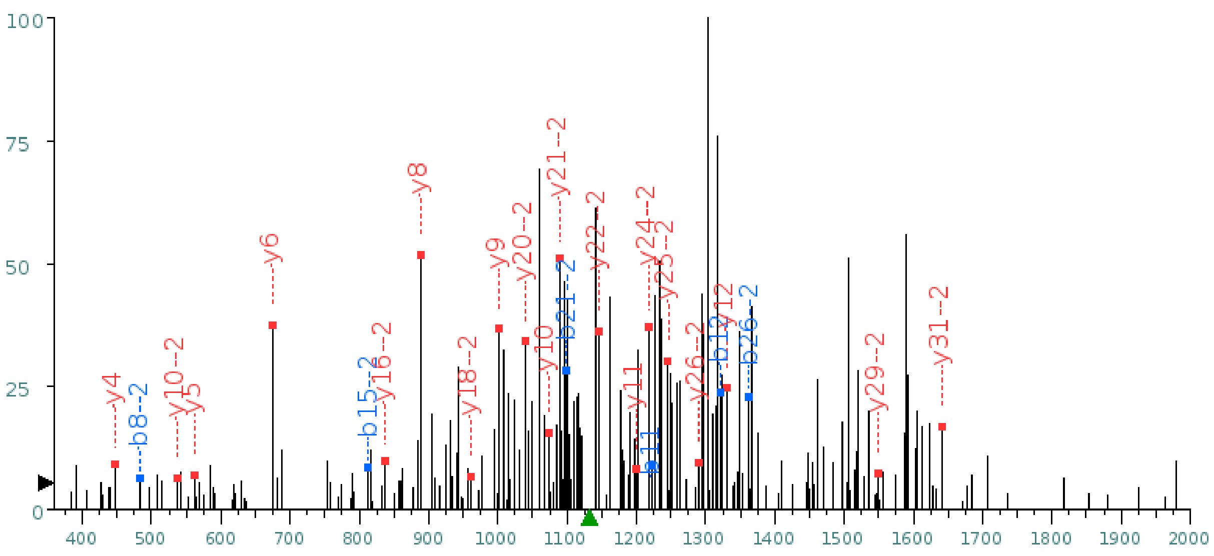

### Predicted Fragmentation Pattern

+1

| Seq | # | b: Δ Error | b | y | y: Δ Error | +1 |
|---|---|---|---|---|---|---|
| N | 1 | --- | 115.050 | --- | --- | 32 |
| I | 2 | --- | 228.134 | 3283.610 | --- | 31 |
| A | 3 | --- | 299.171 | 3170.526 | --- | 30 |
| L | 4 | --- | 412.255 | 3099.489 | --- | 29 |
| S | 5 | --- | 499.287 | 2986.404 | --- | 28 |
| T# | 6 | --- | 820.393 | 2899.372 | --- | 27 |
| S | 7 | --- | 907.425 | 2578.266 | --- | 26 |
| G | 8 | --- | 964.447 | 2491.234 | --- | 25 |
| S | 9 | --- | 1051.479 | 2434.213 | --- | 24 |
| G | 10 | --- | 1108.500 | 2347.181 | --- | 23 |
| N | 11 | -434.857 | 1222.543 | 2290.160 | --- | 22 |
| V | 12 | -374.650 | 1321.612 | 2176.117 | --- | 21 |
| G | 13 | --- | 1378.633 | 2077.048 | --- | 20 |
| V | 14 | --- | 1477.702 | 2020.027 | --- | 19 |
| M* | 15 | --- | 1624.737 | 1920.958 | --- | 18 |
| T | 16 | --- | 1725.785 | 1773.923 | --- | 17 |
| Q | 17 | --- | 1853.843 | 1672.875 | --- | 16 |
| S | 18 | --- | 1940.875 | 1544.817 | --- | 15 |
| G | 19 | --- | 1997.897 | 1457.785 | --- | 14 |
| A | 20 | --- | 2068.934 | 1400.763 | --- | 13 |
| E | 21 | --- | 2197.977 | 1329.726 | -83.416 | 12 |

| Seq | # | b: Δ Error | b | y | y: Δ Error | +1 |
|---|---|---|---|---|---|---|
| Q | 22 | --- | 2326.035 | 1200.683 | 222.840 | 11 |
| A | 23 | --- | 2397.072 | 1072.625 | 591.574 | 10 |
| I | 24 | --- | 2510.156 | 1001.588 | 180.009 | 9 |
| T | 25 | --- | 2611.204 | 888.504 | 76.093 | 8 |
| I | 26 | --- | 2724.288 | 787.456 | --- | 7 |
| I | 27 | --- | 2837.372 | 674.372 | -80.805 | 6 |
| D | 28 | --- | 2952.399 | 561.288 | 399.309 | 5 |
| D | 29 | --- | 3067.426 | 446.261 | 444.784 | 4 |
| A | 30 | --- | 3138.463 | 331.234 | --- | 3 |
| I | 31 | --- | 3251.547 | 260.197 | --- | 2 |
| K | 32 | --- | --- | 147.113 | --- | 1 |

+2

| Seq | # | b: Δ Error | b | y | y: Δ Error | +1 |
|---|---|---|---|---|---|---|
| N | 1 | --- | 58.029 | --- | --- | 32 |
| I | 2 | --- | 114.571 | 1642.309 | -172.418 | 31 |
| A | 3 | --- | 150.089 | 1585.766 | --- | 30 |
| L | 4 | --- | 206.631 | 1550.248 | -175.385 | 29 |
| S | 5 | --- | 250.147 | 1493.706 | --- | 28 |
| T# | 6 | --- | 410.700 | 1450.190 | --- | 27 |
| S | 7 | --- | 454.216 | 1289.637 | 224.534 | 26 |
| G | 8 | -42.355 | 482.727 | 1246.121 | -471.033 | 25 |
| S | 9 | --- | 526.243 | 1217.610 | -46.844 | 24 |
| G | 10 | --- | 554.754 | 1174.094 | --- | 23 |
| N | 11 | --- | 611.775 | 1145.583 | 290.164 | 22 |
| V | 12 | --- | 661.310 | 1088.562 | 815.561 | 21 |
| G | 13 | --- | 689.820 | 1039.028 | 395.994 | 20 |
| V | 14 | --- | 739.354 | 1010.517 | --- | 19 |
| M* | 15 | -341.574 | 812.872 | 960.983 | 768.690 | 18 |
| T | 16 | --- | 863.396 | 887.465 | --- | 17 |
| Q | 17 | --- | 927.425 | 836.941 | 350.850 | 16 |
| S | 18 | --- | 970.941 | 772.912 | --- | 15 |
| G | 19 | --- | 999.452 | 729.396 | --- | 14 |
| A | 20 | --- | 1034.971 | 700.885 | --- | 13 |
| E | 21 | -421.135 | 1099.492 | 665.367 | --- | 12 |
| Q | 22 | --- | 1163.521 | 600.845 | --- | 11 |
| A | 23 | --- | 1199.040 | 536.816 | 1436.742 | 10 |
| I | 24 | --- | 1255.582 | 501.298 | --- | 9 |
| T | 25 | --- | 1306.106 | 444.755 | --- | 8 |
| I | 26 | -296.831 | 1362.648 | 394.232 | --- | 7 |
| I | 27 | --- | 1419.190 | 337.690 | --- | 6 |
| D | 28 | --- | 1476.703 | 281.148 | --- | 5 |
| D | 29 | --- | 1534.217 | 223.634 | --- | 4 |
| A | 30 | --- | 1569.735 | 166.121 | --- | 3 |
| I | 31 | --- | 1626.277 | 130.602 | --- | 2 |
| K | 32 | --- | --- | 74.060 | --- | 1 |

**Soleil Index**

| | | |
|---|---|---|
| 889.6531 | b1-7 frag loss of water/ b1-7 frag loss of ammonia | 20% |
| 904.7061 | not explained | 19% |
| 931.6354 | not explained | 18% |

| | | |
|---|---|---|
| 942.2871 | y2-18 frag loss of water | 29% |
| 994.5796 | not explained | 16% |
| 1009.1902 | y1-19 frag doubly charged | 32% |
| 1015.3002 | not explained | 24% |
| 1025.1014 | not explained | 22% |
| 1044.9302 | not explained | 16% |
| 1048.4615 | not explained | 22% |
| 1060.3735 | not explained | 69% |
| 1068.0153 | not explained | 19% |
| 1085.7759 | not explained | 17% |
| 1091.6099 | b1-10 frag loss of ammonia | 16% |
| 1096.8680 | not explained | 46% |
| 1101.4918 | not explained | 43% |
| 1102.9463 | not explained | 15% |
| 1110.0471 | not explained | 22% |
| 1114.5542 | not explained | 23% |
| 1116.8420 | not explained | 24% |
| 1118.2581 | not explained | 17% |
| 1121.1565 | precursor loss of two waters | 15% |
| 1140.4713 | not explained | 61% |
| 1162.0839 | b1-22 frag doubly charged | 43% |
| 1177.5302 | not explained | 24% |
| 1189.7618 | not explained | 20% |

| | | |
|---|---|---|
| 1202.7990 | not explained | 32% |
| 1227.2379 | y2-25 frag loss of water | 43% |
| 1233.9255 | not explained | 50% |
| 1236.1033 | not explained | 39% |
| 1243.8713 | not explained | 24% |
| 1250.1066 | not explained | 28% |
| 1251.3909 | not explained | 22% |
| 1259.1798 | not explained | 26% |
| 1262.1355 | not explained | 26% |
| 1295.5863 | not explained | 44% |
| 1296.9921 | not explained | 37% |
| 1304.2736 | b1-12 frag loss of water/ b1-12 frag loss of ammonia | 100% |
| 1309.6313 | not explained | 19% |
| 1314.7600 | not explained | 21% |
| 1318.0449 | not explained | 76% |
| 1324.9835 | not explained | 27% |
| 1348.3016 | not explained | 36% |
| 1365.8031 | not explained | 41% |
| 1374.8348 | not explained | 16% |
| 1460.4543 | b1-14 frag loss of water/ b1-14 frag loss of ammonia/ b2-28 frag loss of ammonia | 26% |
| 1498.4941 | not explained | 18% |
| 1505.6744 | not explained | 51% |
| 1521.0509 | not explained | 28% |

| | | |
|---|---|---|
| 1536.2687 | not explained | 20% |
| 1588.1825 | not explained | 16% |
| 1589.4965 | not explained | 56% |
| 1590.9578 | not explained | 27% |
| 1605.8424 | b1-15 frag loss of water | 20% |
| 1611.9448 | not explained | 17% |
| 1622.9609 | not explained | 17% |

**Run:** 77653

**Scans:** 33709 - 33709

**m/z Range:** 337.32477 to 2000

**Search:** 77653

**Precursor Mass:** 1133.22119

**Mods:** C=160.03 *+15.99 #+220.06 @+291.1

**Charge:** +3

**Sequence**

NIALSTSGSGNVGVM*TQSGAEQAIT#IIDDAIK

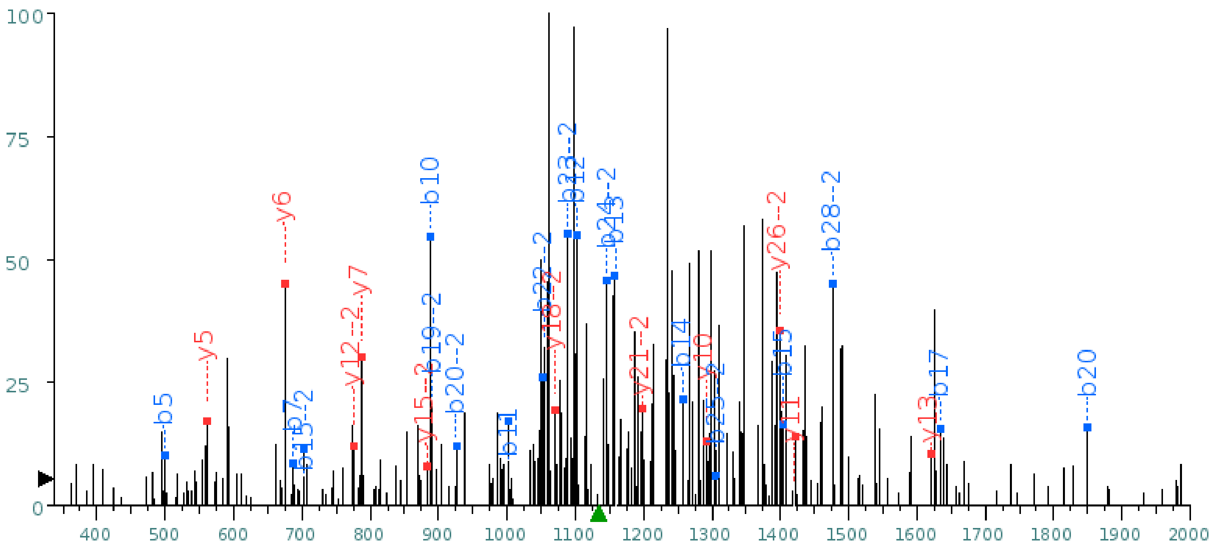

### Predicted Fragmentation Pattern

+1

| Seq | # | b: Δ Error | b | y | y: Δ Error | +1 |
|---|---|---|---|---|---|---|
| N | 1 | --- | 115.050 | --- | --- | 32 |
| I | 2 | --- | 228.134 | 3283.610 | --- | 31 |
| A | 3 | --- | 299.171 | 3170.526 | --- | 30 |
| L | 4 | --- | 412.255 | 3099.489 | --- | 29 |
| S | 5 | 536.919 | **499.287** | 2986.404 | --- | 28 |
| T | 6 | --- | 600.335 | 2899.372 | --- | 27 |
| S | 7 | 15.460 | **687.367** | 2798.325 | --- | 26 |
| G | 8 | --- | 744.389 | 2711.293 | --- | 25 |
| S | 9 | --- | 831.421 | 2654.271 | --- | 24 |
| G | 10 | 57.294 | **888.442** | 2567.239 | --- | 23 |
| N | 11 | -191.168 | **1002.485** | 2510.218 | --- | 22 |
| V | 12 | 317.822 | **1101.553** | 2396.175 | --- | 21 |
| G | 13 | -490.167 | **1158.575** | 2297.106 | --- | 20 |
| V | 14 | 422.106 | **1257.643** | 2240.085 | --- | 19 |
| M* | 15 | 191.639 | **1404.679** | 2141.017 | --- | 18 |
| T | 16 | --- | 1505.726 | 1993.981 | --- | 17 |
| Q | 17 | 167.127 | **1633.785** | 1892.934 | --- | 16 |
| S | 18 | --- | 1720.817 | 1764.875 | --- | 15 |
| G | 19 | --- | 1777.839 | 1677.843 | --- | 14 |
| A | 20 | 21.784 | **1848.876** | **1620.821** | 170.089 | 13 |
| E | 21 | --- | 1977.918 | 1549.784 | --- | 12 |

| | | | | | | |
|---|---|---|---|---|---|---|
| Q | 22 | --- | 2105.977 | 1420.742 | 594.621 | 11 |
| A | 23 | --- | 2177.014 | 1292.683 | -576.308 | 10 |
| I | 24 | --- | 2290.098 | 1221.646 | --- | 9 |
| T# | 25 | --- | 2611.204 | 1108.562 | --- | 8 |
| I | 26 | --- | 2724.288 | 787.456 | -65.193 | 7 |
| I | 27 | --- | 2837.372 | 674.372 | 25.365 | 6 |
| D | 28 | --- | 2952.399 | 561.288 | 272.819 | 5 |
| D | 29 | --- | 3067.426 | 446.261 | --- | 4 |
| A | 30 | --- | 3138.463 | 331.234 | --- | 3 |
| I | 31 | --- | 3251.547 | 260.197 | --- | 2 |
| K | 32 | --- | --- | 147.113 | --- | 1 |

+2

| Seq | # | b: Δ Error | b | y | y: Δ Error | +1 |
|---|---|---|---|---|---|---|
| N | 1 | --- | 58.029 | --- | --- | 32 |
| I | 2 | --- | 114.571 | 1642.309 | --- | 31 |
| A | 3 | --- | 150.089 | 1585.766 | --- | 30 |
| L | 4 | --- | 206.631 | 1550.248 | --- | 29 |
| S | 5 | --- | 250.147 | 1493.706 | --- | 28 |
| T | 6 | --- | 300.671 | 1450.190 | --- | 27 |
| S | 7 | --- | 344.187 | 1399.666 | 575.617 | 26 |
| G | 8 | --- | 372.698 | 1356.150 | --- | 25 |
| S | 9 | --- | 416.214 | 1327.639 | --- | 24 |
| G | 10 | --- | 444.725 | 1284.123 | --- | 23 |
| N | 11 | --- | 501.746 | 1255.613 | --- | 22 |
| V | 12 | --- | 551.280 | 1198.591 | -739.134 | 21 |
| G | 13 | --- | 579.791 | 1149.057 | --- | 20 |
| V | 14 | --- | 629.325 | 1120.546 | --- | 19 |
| M* | 15 | 871.121 | 702.843 | 1071.012 | 437.623 | 18 |
| T | 16 | --- | 753.367 | 997.494 | --- | 17 |
| Q | 17 | --- | 817.396 | 946.970 | --- | 16 |
| S | 18 | --- | 860.912 | 882.941 | -937.621 | 15 |
| G | 19 | -1046.544 | 889.423 | 839.425 | --- | 14 |
| A | 20 | 839.808 | 924.941 | 810.914 | --- | 13 |
| E | 21 | --- | 989.463 | 775.396 | 132.792 | 12 |
| Q | 22 | -901.453 | 1053.492 | 710.875 | --- | 11 |
| A | 23 | 360.950 | 1089.011 | 646.845 | --- | 10 |
| I | 24 | 227.767 | 1145.553 | 611.327 | --- | 9 |
| T# | 25 | -60.581 | 1306.106 | 554.785 | --- | 8 |
| I | 26 | --- | 1362.648 | 394.232 | --- | 7 |
| I | 27 | --- | 1419.190 | 337.690 | --- | 6 |
| D | 28 | -431.624 | 1476.703 | 281.148 | --- | 5 |
| D | 29 | --- | 1534.217 | 223.634 | --- | 4 |
| A | 30 | --- | 1569.735 | 166.121 | --- | 3 |
| I | 31 | --- | 1626.277 | 130.602 | --- | 2 |
| K | 32 | --- | --- | 74.060 | --- | 1 |

Soleil Index

| | | |
|---|---|---|
| 591.9224 | not explained | 30% |
| 593.1652 | y2-9 frag loss of water | 16% |
| 772.8716 | not explained | 16% |

| | | |
|---|---|---|
| 870.2762 | b1-10 frag loss of water | 16% |
| 938.6213 | not explained | 19% |
| 984.8251 | b1-11 frag loss of water/ b1-11 frag loss of ammonia | 19% |
| 1048.6409 | not explained | 15% |
| 1049.8165 | not explained | 50% |
| 1054.5011 | y2-18 frag loss of ammonia | 32% |
| 1059.9304 | not explained | 46% |
| 1060.7821 | not explained | 100% |
| 1076.3118 | not explained | 25% |
| 1078.6810 | not explained | 19% |
| 1079.7351 | not explained | 16% |
| 1097.5793 | not explained | 17% |
| 1098.3300 | not explained | 97% |
| 1102.6414 | y2-19 frag loss of water/ y2-19 frag loss of ammonia | 26% |
| 1115.5992 | not explained | 37% |
| 1142.0931 | b1-13 frag loss of ammonia | 26% |
| 1154.0341 | not explained | 43% |
| 1155.1583 | not explained | 25% |
| 1166.7065 | not explained | 17% |
| 1187.4611 | not explained | 35% |
| 1191.7185 | not explained | 26% |
| 1192.3563 | not explained | 21% |
| 1211.5006 | not explained | 21% |

| | | |
|---|---|---|
| 1215.0104 | not explained | 33% |
| 1232.5011 | not explained | 30% |
| 1235.8878 | not explained | 97% |
| 1237.2759 | y2-22 frag loss of water | 23% |
| 1241.5781 | b1-14 frag loss of ammonia | 48% |
| 1244.0863 | not explained | 26% |
| 1267.5608 | y2-23 frag loss of ammonia | 49% |
| 1270.7163 | not explained | 21% |
| 1279.8716 | not explained | 27% |
| 1281.6049 | not explained | 52% |
| 1287.7682 | b2-25 frag loss of water | 21% |
| 1298.3763 | not explained | 52% |
| 1303.5660 | not explained | 27% |
| 1309.5602 | y2-24 frag loss of water | 36% |
| 1340.1868 | not explained | 21% |
| 1341.9406 | not explained | 15% |
| 1346.0856 | b2-26 frag loss of ammonia | 57% |
| 1366.9333 | not explained | 16% |
| 1373.3726 | not explained | 58% |
| 1388.7097 | not explained | 29% |
| 1392.6414 | not explained | 16% |
| 1395.6946 | not explained | 47% |
| 1402.0475 | b2-27 frag loss of water/ b2-27 frag loss of ammonia/ y1-11 frag loss of water | 19% |

| | | |
|---|---|---|
| 1407.8319 | not explained | 31% |
| 1433.9581 | y2-27 frag loss of ammonia | 15% |
| 1434.8661 | not explained | 32% |
| 1435.8016 | not explained | 18% |
| 1457.9852 | b2-28 frag loss of water | 17% |
| 1460.4762 | b2-28 frag loss of ammonia | 20% |
| 1488.8014 | b1-16 frag loss of ammonia | 32% |
| 1490.1532 | not explained | 32% |
| 1537.5807 | not explained | 23% |
| 1545.0631 | not explained | 16% |
| 1624.3047 | y2-31 frag loss of water/ y2-31 frag loss of ammonia | 40% |

**MS/MS data of trypsin digested peptides with site score A and site score B above 19 from *Geobacillus kaustophilus* *Gk*F2 flagellin protein co-expressed with *Geobacillus kaustophilus* Maf in EV36 cells. Results of dataset 3 (MS/MS results obtained in July 2019).**

| | | | |
|---|---|---|---|
| **Run:** | 79489 | **Scans:** | 26421 - 26421 |
| **Search:** | 79489 | **Precursor Mass:** | 1156.90125 |
| **Mods:** | C=160.03 *+15.99 #+220.06 @+291.1 | **Charge:** | +3 |

**m/z Range:** 337.26114 **to** 1958.2161[illegible]

**Sequence**

NIALST@SGSGNVGVM*TQSGAEQAITIIDDAIK

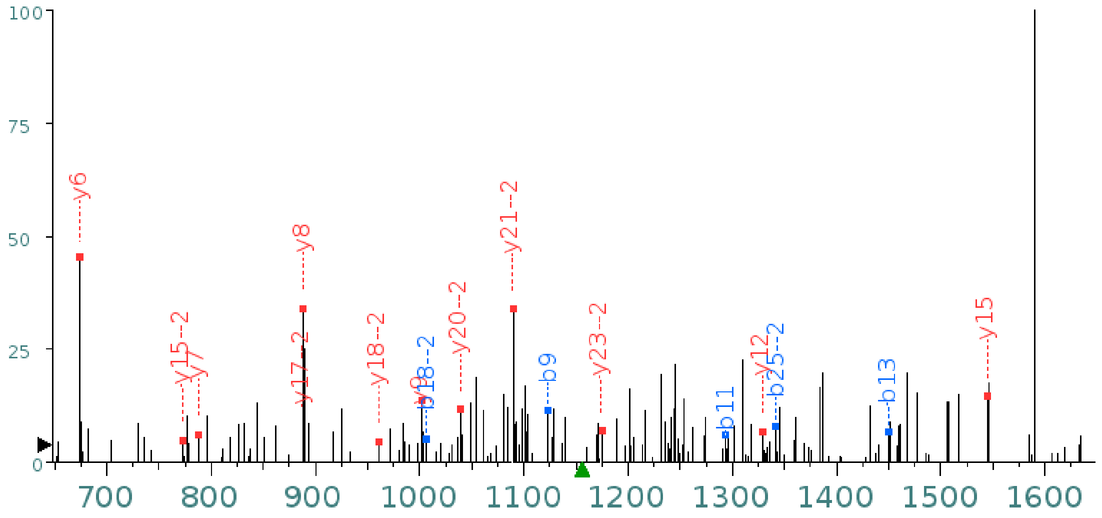

**Predicted Fragmentation Pattern**

+1

| Seq | # | b: Δ Error | b | y | y: Δ Error | +1 |
|---|---|---|---|---|---|---|
| N | 1 | --- | 115.050 | --- | --- | 32 |
| I | 2 | --- | 228.134 | 3354.647 | --- | 31 |
| A | 3 | --- | 299.171 | 3241.563 | --- | 30 |
| L | 4 | --- | 412.255 | 3170.526 | --- | 29 |
| S | 5 | --- | 499.287 | 3057.442 | --- | 28 |
| T@ | 6 | --- | 891.431 | 2970.410 | --- | 27 |

| | | | | | | |
|---|---|---|---|---|---|---|
| S | 7 | --- | 978.463 | 2578.266 | --- | 26 |
| G | 8 | --- | 1035.484 | 2491.234 | --- | 25 |
| S | 9 | 210.617 | 1122.516 | 2434.213 | --- | 24 |
| G | 10 | --- | 1179.538 | 2347.181 | --- | 23 |
| N | 11 | 175.837 | 1293.580 | 2290.160 | --- | 22 |
| V | 12 | --- | 1392.649 | 2176.117 | --- | 21 |
| G | 13 | -39.966 | 1449.670 | 2077.048 | --- | 20 |
| V | 14 | --- | 1548.739 | 2020.027 | --- | 19 |
| M* | 15 | --- | 1695.774 | 1920.958 | 7.926 | 18 |
| T | 16 | --- | 1796.822 | 1773.923 | --- | 17 |
| Q | 17 | --- | 1924.880 | 1672.875 | 540.951 | 16 |
| S | 18 | --- | 2011.912 | 1544.817 | 166.085 | 15 |
| G | 19 | --- | 2068.934 | 1457.785 | --- | 14 |
| A | 20 | --- | 2139.971 | 1400.763 | --- | 13 |
| E | 21 | --- | 2269.014 | 1329.726 | -108.850 | 12 |
| Q | 22 | --- | 2397.072 | 1200.683 | --- | 11 |
| A | 23 | --- | 2468.109 | 1072.625 | --- | 10 |
| I | 24 | --- | 2581.193 | 1001.588 | 286.602 | 9 |
| T | 25 | --- | 2682.241 | 888.504 | -249.987 | 8 |
| I | 26 | --- | 2795.325 | 787.456 | 107.800 | 7 |
| I | 27 | --- | 2908.409 | 674.372 | 27.718 | 6 |
| D | 28 | --- | 3023.436 | 561.288 | 514.469 | 5 |
| D | 29 | --- | 3138.463 | 446.261 | 247.903 | 4 |
| A | 30 | --- | 3209.500 | 331.234 | --- | 3 |
| I | 31 | --- | 3322.584 | 260.197 | --- | 2 |
| K | 32 | --- | --- | 147.113 | --- | 1 |

+2

| Seq | # | b: Δ Error | b | y | y: Δ Error | +1 |
|---|---|---|---|---|---|---|
| N | 1 | --- | 58.029 | --- | --- | 32 |
| I | 2 | --- | 114.571 | 1677.827 | --- | 31 |
| A | 3 | --- | 150.089 | 1621.285 | --- | 30 |
| L | 4 | --- | 206.631 | 1585.766 | --- | 29 |
| S | 5 | --- | 250.147 | 1529.224 | --- | 28 |
| T@ | 6 | 341.998 | 446.219 | 1485.708 | --- | 27 |
| S | 7 | --- | 489.735 | 1289.637 | --- | 26 |
| G | 8 | --- | 518.246 | 1246.121 | --- | 25 |
| S | 9 | -329.255 | 561.762 | 1217.610 | --- | 24 |
| G | 10 | --- | 590.272 | 1174.094 | 496.828 | 23 |
| N | 11 | --- | 647.294 | 1145.583 | --- | 22 |
| V | 12 | --- | 696.828 | 1088.562 | 798.656 | 21 |
| G | 13 | --- | 725.339 | 1039.028 | 214.356 | 20 |
| V | 14 | --- | 774.873 | 1010.517 | --- | 19 |
| M* | 15 | --- | 848.391 | 960.983 | 639.432 | 18 |
| T | 16 | --- | 898.915 | 887.465 | 919.231 | 17 |
| Q | 17 | --- | 962.944 | 836.941 | --- | 16 |
| S | 18 | -158.333 | 1006.460 | 772.912 | 15.522 | 15 |
| G | 19 | --- | 1034.971 | 729.396 | --- | 14 |
| A | 20 | --- | 1070.489 | 700.885 | --- | 13 |
| E | 21 | --- | 1135.010 | 665.367 | --- | 12 |
| Q | 22 | --- | 1199.040 | 600.845 | --- | 11 |
| A | 23 | --- | 1234.558 | 536.816 | --- | 10 |
| I | 24 | --- | 1291.100 | 501.298 | --- | 9 |
| T | 25 | -631.775 | 1341.624 | 444.755 | --- | 8 |
| I | 26 | --- | 1398.166 | 394.232 | --- | 7 |
| I | 27 | --- | 1454.708 | 337.690 | --- | 6 |
| D | 28 | --- | 1512.222 | 281.148 | --- | 5 |
| D | 29 | --- | 1569.735 | 223.634 | --- | 4 |
| A | 30 | --- | 1605.254 | 166.121 | --- | 3 |

| | | | | | |
|---|---|---|---|---|---|
| I 31 | --- | 1661.796 | 130.602 | --- | 2 |
| K 32 | --- | --- | 74.060 | --- | 1 |

Soleil Index

| | | |
|---|---|---|
| 362.2611 | b2-13 frag doubly charged | 2% |
| 381.2674 | not explained | 7% |
| 437.6061 | not explained | 2% |
| 473.4189 | b2-7 frag loss of ammonia | 2% |
| 612.4464 | not explained | 4% |
| 651.2664 | not explained | 1% |
| 652.6708 | not explained | 5% |
| 675.5514 | not explained | 9% |
| 676.6354 | not explained | 2% |
| 682.4309 | y2-13 frag loss of water | 7% |
| 704.4135 | not explained | 5% |
| 730.4091 | not explained | 9% |
| 735.6176 | not explained | 5% |
| 742.4935 | y2-27 frag doubly charged | 3% |
| 776.4021 | not explained | 10% |
| 778.6813 | not explained | 4% |
| 796.4127 | not explained | 10% |
| 809.2631 | not explained | 1% |
| 811.7030 | not explained | 3% |
| 818.4347 | y2-16 frag loss of water | 6% |
| 826.2339 | not explained | 8% |
| 831.6542 | b2-15 frag loss of ammonia/ b2-31 frag loss of water | 9% |
| 835.5948 | y1-16 frag doubly charged | 1% |
| 843.6017 | not explained | 13% |
| 850.5503 | not explained | 5% |
| 861.7155 | not explained | 8% |
| 874.6086 | b1-6 frag loss of ammonia | 2% |
| 889.6003 | not explained | 25% |
| 893.4298 | not explained | 9% |
| 917.3751 | not explained | 7% |
| 925.1288 | not explained | 12% |
| 932.8837 | not explained | 2% |
| 971.7219 | not explained | 7% |
| 980.7905 | not explained | 3% |
| 983.7736 | y1-9 frag loss of water/ y1-9 frag loss of ammonia | 9% |
| 985.7899 | not explained | 4% |
| 990.4353 | not explained | 4% |
| 997.7418 | not explained | 4% |

| | | |
|---|---|---|
| 1003.0237 | not explained | 7% |
| 1016.1038 | b2-19 frag loss of water | 2% |
| 1019.5651 | not explained | 4% |
| 1028.7568 | not explained | 2% |
| 1030.6477 | not explained | 4% |
| 1036.5858 | not explained | 5% |
| 1040.3278 | not explained | 6% |
| 1048.2283 | not explained | 13% |
| 1048.8383 | not explained | 1% |
| 1054.4939 | y1-10 frag loss of water | 19% |
| 1060.5277 | not explained | 11% |
| 1064.7847 | not explained | 1% |
| 1067.7258 | not explained | 2% |
| 1073.9535 | not explained | 4% |
| 1081.0389 | not explained | 15% |
| 1084.2368 | not explained | 12% |
| 1085.0731 | not explained | 3% |
| 1090.2808 | not explained | 9% |
| 1091.6840 | not explained | 8% |
| 1092.9133 | not explained | 9% |
| 1095.3339 | not explained | 4% |
| 1098.9098 | not explained | 12% |
| 1101.6201 | not explained | 17% |
| 1102.9065 | not explained | 7% |
| 1104.2568 | b1-9 frag loss of water | 11% |
| 1107.7684 | not explained | 2% |
| 1126.7894 | y2-22 frag loss of water | 6% |
| 1128.1517 | y2-22 frag loss of water/ y2-22 frag loss of ammonia | 12% |
| 1136.6870 | not explained | 4% |
| 1139.8397 | not explained | 10% |
| 1159.5392 | not explained | 3% |
| 1169.7325 | not explained | 6% |
| 1171.4991 | not explained | 9% |
| 1172.1802 | not explained | 0% |
| 1175.5822 | not explained | 3% |
| 1188.7527 | not explained | 9% |
| 1196.9232 | not explained | 3% |
| 1201.8773 | not explained | 16% |
| 1203.0746 | not explained | 3% |
| 1205.9478 | not explained | 5% |
| 1213.6489 | not explained | 4% |

| | | |
|---|---|---|
| 1216.3073 | b2-23 frag loss of water/ y1-24 frag loss of water | 11% |
| 1222.7423 | not explained | 1% |
| 1231.3944 | not explained | 19% |
| 1235.9255 | not explained | 9% |
| 1239.0704 | not explained | 4% |
| 1239.7430 | not explained | 3% |
| 1240.9822 | not explained | 10% |
| 1243.5929 | not explained | 12% |
| 1244.5833 | not explained | 21% |
| 1247.5217 | not explained | 5% |
| 1249.9590 | not explained | 2% |
| 1252.1797 | not explained | 4% |
| 1253.2672 | not explained | 14% |
| 1253.8715 | not explained | 4% |
| 1257.7852 | not explained | 2% |
| 1262.2903 | not explained | 8% |
| 1272.1545 | b2-24 frag loss of water/ y2-26 frag loss of water/ y2-26 frag loss of ammonia | 6% |
| 1274.2655 | b2-24 frag loss of ammonia | 10% |
| 1295.8431 | not explained | 6% |
| 1301.1962 | not explained | 8% |
| 1302.2606 | not explained | 3% |
| 1309.3099 | not explained | 7% |
| 1310.2396 | not explained | 23% |
| 1312.0499 | y1-12 frag loss of water/ y1-12 frag loss of ammonia | 1% |
| 1315.7064 | not explained | 1% |
| 1318.1952 | not explained | 8% |
| 1330.9386 | not explained | 3% |
| 1332.0753 | not explained | 2% |
| 1333.4170 | not explained | 3% |
| 1336.6030 | not explained | 4% |
| 1345.2572 | not explained | 12% |
| 1348.9916 | not explained | 2% |
| 1358.9738 | not explained | 5% |
| 1361.0831 | not explained | 10% |
| 1369.0433 | not explained | 4% |
| 1372.9308 | not explained | 3% |
| 1375.3583 | b1-12 frag loss of water/ b1-12 frag loss of ammonia | 3% |
| 1383.8986 | y1-13 frag loss of ammonia | 17% |
| 1384.6261 | y1-13 frag loss of ammonia | 10% |
| 1386.7998 | not explained | 20% |
| 1403.1044 | not explained | 1% |

| | | |
|---|---|---|
| 1405.0812 | not explained | 1% |
| 1427.7858 | not explained | 1% |
| 1432.3516 | b1-13 frag loss of water/ b1-13 frag loss of ammonia | 12% |
| 1437.0454 | b2-27 frag loss of water/ b2-27 frag loss of ammonia | 2% |
| 1440.0131 | y1-14 frag loss of water/ y1-14 frag loss of ammonia | 4% |
| 1451.0414 | not explained | 7% |
| 1451.8807 | not explained | 9% |
| 1459.7258 | not explained | 8% |
| 1460.4581 | not explained | 8% |
| 1461.1044 | not explained | 2% |
| 1468.1211 | y2-27 frag loss of water/ y2-27 frag loss of ammonia | 20% |
| 1476.9315 | not explained | 15% |
| 1488.9767 | not explained | 2% |
| 1505.9662 | not explained | 13% |
| 1507.4797 | not explained | 7% |
| 1508.1904 | not explained | 13% |
| 1516.6444 | not explained | 13% |
| 1517.4625 | not explained | 15% |
| 1546.1444 | not explained | 17% |
| 1584.1561 | not explained | 6% |
| 1587.0709 | b2-30 frag loss of water | 2% |
| 1589.4860 | not explained | 100% |
| 1590.6084 | not explained | 54% |
| 1606.4496 | not explained | 2% |
| 1611.9286 | not explained | 2% |
| 1619.3016 | not explained | 3% |
| 1633.1947 | not explained | 4% |
| 1633.8472 | not explained | 6% |
| 1679.3085 | b1-15 frag loss of ammonia | 4% |
| 1703.8990 | not explained | 2% |
| 1721.0128 | not explained | 6% |
| 1722.1476 | not explained | 3% |
| 1759.1554 | not explained | 6% |
| 1765.9222 | not explained | 2% |
| 1848.7783 | not explained | 1% |
| 1884.6929 | not explained | 4% |
| 1926.1436 | not explained | 3% |
| 1933.2162 | not explained | 5% |

| | | | | | |
|---|---|---|---|---|---|
| **Run:** | 79489 | **Scans:** | 27961 - 27961 | **m/z Range:** | 370.21912 to 2000 |
| **Search:** | 79489 | **Precursor Mass:** | 1133.2219 | | |
| **Mods:** | C=160.03 *+15.99 #+220.06 @+291.1 | **Charge:** | +3 | | |

**Sequence**

NIALSTSGSGNVGVM*TQSGAEQAIT#IIDDAIK

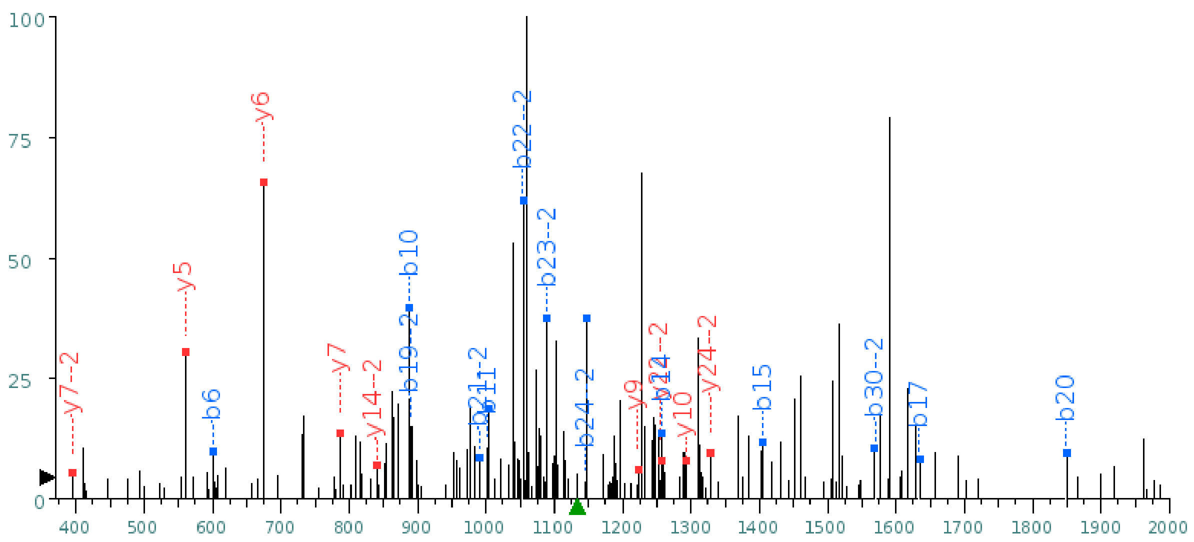

**Predicted Fragmentation Pattern**

| | | | | +1 | | |
|---|---|---|---|---|---|---|
| **Seq** | **#** | **b: Δ Error** | **b** | **y** | **y: Δ Error** | **+1** |
| **N** | **1** | --- | 115.050 | --- | --- | **32** |
| **I** | **2** | --- | 228.134 | 3283.610 | --- | **31** |
| **A** | **3** | --- | 299.171 | 3170.526 | --- | **30** |
| **L** | **4** | --- | 412.255 | 3099.489 | --- | **29** |
| **S** | **5** | --- | 499.287 | 2986.404 | --- | **28** |
| **T** | **6** | **1189.616** | **600.335** | 2899.372 | --- | **27** |

| | | | | | | |
|---|---|---|---|---|---|---|
| S | 7 | --- | 687.367 | 2798.325 | --- | 26 |
| G | 8 | --- | 744.389 | 2711.293 | --- | 25 |
| S | 9 | --- | 831.421 | 2654.271 | --- | 24 |
| G | 10 | 31.878 | **888.442** | 2567.239 | --- | 23 |
| N | 11 | 571.538 | **1002.485** | 2510.218 | --- | 22 |
| V | 12 | --- | 1101.553 | 2396.175 | --- | 21 |
| G | 13 | --- | 1158.575 | 2297.106 | --- | 20 |
| V | 14 | 93.912 | **1257.643** | 2240.085 | --- | 19 |
| M* | 15 | 173.136 | **1404.679** | 2141.017 | --- | 18 |
| T | 16 | --- | 1505.726 | 1993.981 | --- | 17 |
| Q | 17 | 204.172 | **1633.785** | 1892.934 | --- | 16 |
| S | 18 | --- | 1720.817 | 1764.875 | --- | 15 |
| G | 19 | --- | 1777.839 | 1677.843 | --- | 14 |
| A | 20 | 100.211 | **1848.876** | 1620.821 | --- | 13 |
| E | 21 | --- | 1977.918 | 1549.784 | --- | 12 |
| Q | 22 | --- | 2105.977 | 1420.742 | --- | 11 |
| A | 23 | --- | 2177.014 | **1292.683** | -587.747 | 10 |
| I | 24 | --- | 2290.098 | **1221.646** | 780.851 | 9 |
| T# | 25 | --- | 2611.204 | 1108.562 | --- | 8 |
| I | 26 | --- | 2724.288 | **787.456** | 233.787 | 7 |
| I | 27 | --- | 2837.372 | **674.372** | -17.716 | 6 |
| D | 28 | --- | 2952.399 | **561.288** | 326.831 | 5 |
| D | 29 | --- | 3067.426 | 446.261 | --- | 4 |
| A | 30 | --- | 3138.463 | 331.234 | --- | 3 |
| I | 31 | --- | 3251.547 | 260.197 | --- | 2 |
| K | 32 | --- | --- | 147.113 | --- | 1 |

+2

| Seq | # | b: Δ Error | b | y | y: Δ Error | +1 |
|---|---|---|---|---|---|---|
| N | 1 | --- | 58.029 | --- | --- | 32 |
| I | 2 | --- | 114.571 | 1642.309 | --- | 31 |
| A | 3 | --- | 150.089 | 1585.766 | --- | 30 |
| L | 4 | --- | 206.631 | 1550.248 | --- | 29 |
| S | 5 | --- | 250.147 | 1493.706 | --- | 28 |
| T | 6 | --- | 300.671 | 1450.190 | --- | 27 |
| S | 7 | --- | 344.187 | 1399.666 | --- | 26 |
| G | 8 | --- | 372.698 | 1356.150 | --- | 25 |
| S | 9 | --- | 416.214 | **1327.639** | 208.118 | 24 |
| G | 10 | --- | 444.725 | 1284.123 | --- | 23 |
| N | 11 | --- | 501.746 | **1255.613** | 591.045 | 22 |
| V | 12 | --- | 551.280 | 1198.591 | --- | 21 |
| G | 13 | --- | 579.791 | 1149.057 | --- | 20 |
| V | 14 | --- | 629.325 | 1120.546 | --- | 19 |
| M* | 15 | --- | 702.843 | 1071.012 | --- | 18 |
| T | 16 | --- | 753.367 | 997.494 | --- | 17 |
| Q | 17 | --- | 817.396 | 946.970 | --- | 16 |
| S | 18 | --- | 860.912 | 882.941 | --- | 15 |
| G | 19 | -1071.989 | **889.423** | **839.425** | 376.162 | 14 |
| A | 20 | --- | 924.941 | 810.914 | --- | 13 |
| E | 21 | 16.311 | **989.463** | 775.396 | --- | 12 |
| Q | 22 | 899.895 | **1053.492** | 710.875 | --- | 11 |
| A | 23 | 801.299 | **1089.011** | 646.845 | --- | 10 |
| I | 24 | 450.539 | **1145.553** | 611.327 | --- | 9 |
| T# | 25 | --- | 1306.106 | 554.785 | --- | 8 |
| I | 26 | --- | 1362.648 | **394.232** | 2498.563 | 7 |
| I | 27 | --- | 1419.190 | 337.690 | --- | 6 |
| D | 28 | --- | 1476.703 | 281.148 | --- | 5 |
| D | 29 | --- | 1534.217 | 223.634 | --- | 4 |

| | | | | | | |
|---|---|---|---|---|---|---|
| A | 30 | -455.252 | **1569.735** | 166.121 | --- | 3 |
| I | 31 | --- | 1626.277 | 130.602 | --- | 2 |
| K | 32 | --- | --- | 74.060 | --- | 1 |

**Soleil Index**

| | | |
|---|---|---|
| 731.8473 | not explained | 17% |
| 862.7158 | not explained | 22% |
| 865.6359 | y2-15 frag loss of water/ y2-15 frag loss of ammonia | 17% |
| 872.5585 | b2-19 frag loss of ammonia | 20% |
| 975.8459 | not explained | 19% |
| 1039.4210 | not explained | 53% |
| 1040.1381 | not explained | 24% |
| 1060.3192 | not explained | 100% |
| 1072.9030 | b2-23 frag loss of ammonia | 27% |
| 1074.0048 | not explained | 17% |
| 1102.8434 | y2-19 frag loss of water/ y2-19 frag loss of ammonia | 33% |
| 1197.0526 | not explained | 20% |
| 1227.3107 | not explained | 68% |
| 1244.6205 | not explained | 17% |
| 1248.9120 | not explained | 15% |
| 1309.7781 | y2-24 frag loss of water/ y2-24 frag loss of ammonia | 33% |
| 1367.9058 | not explained | 17% |
| 1452.3842 | not explained | 21% |
| 1460.9723 | not explained | 25% |
| 1508.4816 | not explained | 24% |
| 1517.0802 | b2-29 frag loss of water/ b2-29 frag loss of ammonia | 36% |
| 1577.6322 | not explained | 17% |
| 1589.7072 | not explained | 79% |
| 1618.1257 | not explained | 23% |
| 1627.5018 | not explained | 15% |

| | | | | | |
|---|---|---|---|---|---|
| **Run:** | 79489 | **Scans:** | 32487 - 32487 | **m/z Range:** | 331.28336 to 1988.9852[illegible] |
| **Search:** | 79489 | **Precursor Mass:** | 1156.89903 | | |
| **Mods:** | C=160.03 *+15.99 #+220.06 @+291.1 | **Charge:** | +3 | | |

**Sequence**

NIALSTSGSGNVGVM*TQSGAEQAIT@IIDDAIK

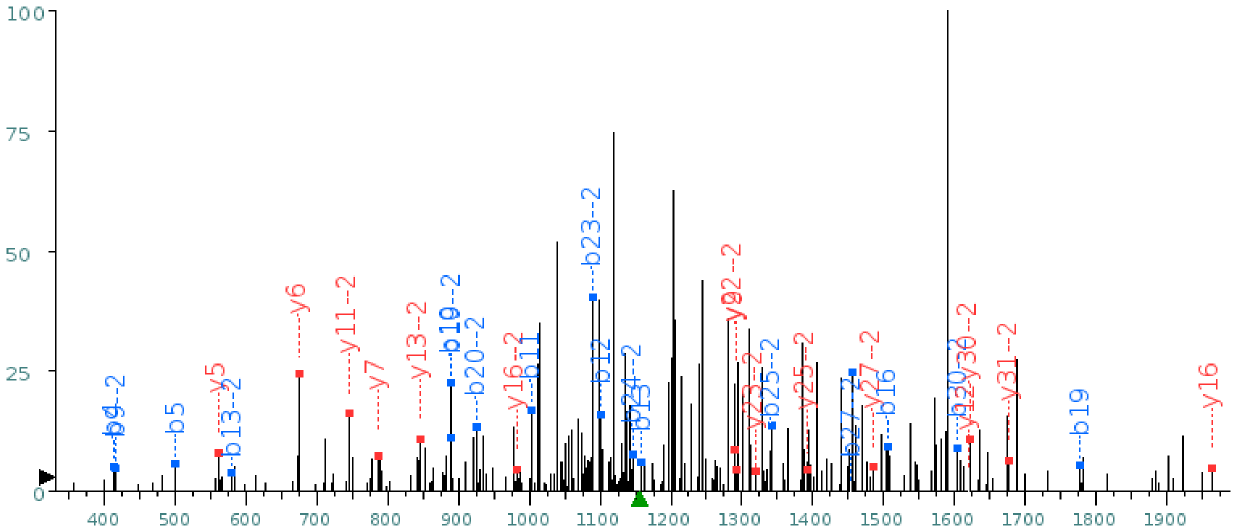

**Predicted Fragmentation Pattern**

| Seq | # | b: Δ Error | b +1 | y +1 | y: Δ Error | +1 |
|---|---|---|---|---|---|---|
| N | 1 | --- | 115.050 | --- | --- | 32 |
| I | 2 | --- | 228.134 | 3354.647 | --- | 31 |
| A | 3 | --- | 299.171 | 3241.563 | --- | 30 |
| L | 4 | **452.062** | **412.255** | 3170.526 | --- | 29 |
| S | 5 | **756.431** | **499.287** | 3057.442 | --- | 28 |
| T | 6 | --- | 600.335 | 2970.410 | --- | 27 |
| S | 7 | --- | 687.367 | 2869.362 | --- | 26 |
| G | 8 | --- | 744.389 | 2782.330 | --- | 25 |

| | | | | | | |
|---|---|---|---|---|---|---|
| S | 9 | --- | 831.421 | 2725.308 | --- | 24 |
| G | 10 | -26.516 | **888.442** | 2638.276 | --- | 23 |
| N | 11 | 255.812 | **1002.485** | 2581.255 | --- | 22 |
| V | 12 | 90.076 | **1101.553** | 2467.212 | --- | 21 |
| G | 13 | 250.287 | **1158.575** | 2368.144 | --- | 20 |
| V | 14 | --- | 1257.643 | 2311.122 | --- | 19 |
| M* | 15 | --- | 1404.679 | 2212.054 | --- | 18 |
| T | 16 | 225.898 | **1505.726** | 2065.018 | --- | 17 |
| Q | 17 | --- | 1633.785 | **1963.971** | 7.435 | 16 |
| S | 18 | --- | 1720.817 | 1835.912 | --- | 15 |
| G | 19 | 194.687 | **1777.839** | 1748.880 | --- | 14 |
| A | 20 | --- | 1848.876 | 1691.859 | --- | 13 |
| E | 21 | --- | 1977.918 | **1620.821** | 417.045 | 12 |
| Q | 22 | --- | 2105.977 | 1491.779 | --- | 11 |
| A | 23 | --- | 2177.014 | 1363.720 | --- | 10 |
| I | 24 | --- | 2290.098 | **1292.683** | 121.855 | 9 |
| T@ | 25 | --- | 2682.241 | 1179.599 | --- | 8 |
| I | 26 | --- | 2795.325 | **787.456** | 301.804 | 7 |
| I | 27 | --- | 2908.409 | **674.372** | 130.698 | 6 |
| D | 28 | --- | 3023.436 | **561.288** | 1324.839 | 5 |
| D | 29 | --- | 3138.463 | 446.261 | --- | 4 |
| A | 30 | --- | 3209.500 | 331.234 | --- | 3 |
| I | 31 | --- | 3322.584 | 260.197 | --- | 2 |
| K | 32 | --- | --- | 147.113 | --- | 1 |

+2

| Seq | # | b: Δ Error | b | y | y: Δ Error | +1 |
|---|---|---|---|---|---|---|
| N | 1 | --- | 58.029 | --- | --- | 32 |
| I | 2 | --- | 114.571 | **1677.827** | -87.556 | 31 |
| A | 3 | --- | 150.089 | **1621.285** | 131.142 | 30 |
| L | 4 | --- | 206.631 | 1585.766 | --- | 29 |
| S | 5 | --- | 250.147 | 1529.224 | --- | 28 |
| T | 6 | --- | 300.671 | **1485.708** | 183.070 | 27 |
| S | 7 | --- | 344.187 | 1435.185 | --- | 26 |
| G | 8 | --- | 372.698 | **1391.669** | 361.695 | 25 |
| S | 9 | 71.959 | **416.214** | 1363.158 | --- | 24 |
| G | 10 | --- | 444.725 | **1319.642** | -594.579 | 23 |
| N | 11 | --- | 501.746 | **1291.131** | -124.623 | 22 |
| V | 12 | --- | 551.280 | 1234.110 | --- | 21 |
| G | 13 | 1248.792 | **579.791** | 1184.575 | --- | 20 |
| V | 14 | --- | 629.325 | 1156.065 | --- | 19 |
| M* | 15 | --- | 702.843 | 1106.530 | --- | 18 |
| T | 16 | --- | 753.367 | 1033.013 | --- | 17 |
| Q | 17 | --- | 817.396 | **982.489** | -353.417 | 16 |
| S | 18 | --- | 860.912 | 918.460 | --- | 15 |
| G | 19 | 433.469 | **889.423** | 874.944 | --- | 14 |
| A | 20 | 240.096 | **924.941** | **846.433** | 289.007 | 13 |
| E | 21 | --- | 989.463 | 810.914 | --- | 12 |
| Q | 22 | --- | 1053.492 | **746.393** | 216.984 | 11 |
| A | 23 | 84.092 | **1089.011** | 682.364 | --- | 10 |
| I | 24 | 667.254 | **1145.553** | 646.845 | --- | 9 |
| T@ | 25 | 671.379 | **1341.624** | 590.303 | --- | 8 |
| I | 26 | --- | 1398.166 | 394.232 | --- | 7 |
| I | 27 | 593.435 | **1454.708** | 337.690 | --- | 6 |
| D | 28 | --- | 1512.222 | 281.148 | --- | 5 |
| D | 29 | --- | 1569.735 | 223.634 | --- | 4 |
| A | 30 | -53.050 | **1605.254** | 166.121 | --- | 3 |
| I | 31 | --- | 1661.796 | 130.602 | --- | 2 |

| K 32 | --- | --- | 74.060 | --- | 1 |
|---|---|---|---|---|---|

**Soleil Index**

| | | |
|---|---|---|
| 1011.5714 | not explained | 26% |
| 1014.2339 | y2-17 frag loss of water | 35% |
| 1039.6550 | not explained | 52% |
| 1090.2452 | y2-18 frag loss of ammonia | 17% |
| 1099.4060 | not explained | 40% |
| 1119.5389 | not explained | 74% |
| 1136.0564 | not explained | 29% |
| 1137.2870 | y2-19 frag loss of water | 16% |
| 1142.0516 | b1-13 frag loss of ammonia | 18% |
| 1195.5309 | not explained | 23% |
| 1200.8722 | not explained | 27% |
| 1203.5034 | not explained | 62% |
| 1206.8008 | not explained | 35% |
| 1214.1454 | not explained | 24% |
| 1227.9694 | not explained | 18% |
| 1239.2469 | b1-14 frag loss of water | 22% |
| 1239.8827 | b1-14 frag loss of water/ b1-14 frag loss of ammonia | 26% |
| 1244.1053 | not explained | 44% |
| 1280.3290 | not explained | 36% |
| 1290.0415 | y1-22 frag doubly charged | 22% |
| 1294.3933 | not explained | 27% |
| 1300.7880 | y2-23 frag loss of water | 20% |
| 1309.6577 | not explained | 34% |
| 1329.0300 | not explained | 26% |
| 1384.7323 | not explained | 31% |
| 1405.9642 | not explained | 27% |
| 1440.9908 | not explained | 23% |
| 1470.0065 | not explained | 18% |
| 1572.4292 | not explained | 19% |
| 1590.0348 | not explained | 100% |
| 1674.2977 | y1-13 frag loss of water/ y1-13 frag loss of ammonia | 16% |
| 1689.3546 | not explained | 27% |

**MS/MS data of GluC digested peptides with site score A and site score B above 19 from *Geobacillus kaustophilus Gk*F2 flagellin protein co-expressed with *Geobacillus kaustophilus* Maf in EV36 cells. Results of dataset 3 (MS/MS results obtained in July 2019).**

| | | | | | |
|---|---|---|---|---|---|
| **Run:** | 79490 | **Scans:** | 24344 - 24344 | **m/z Range:** | 299.13556 to 1977.1405 |
| **Search:** | 79490 | **Precursor Mass:** | 1002.51188 | | |
| **Mods:** | C=160.03 *+15.99 #+220.06 @+291.1 | **Charge:** | +4 | | |

**Sequence**

FT#KNNILT#QAAQAM*LAQANQQPQGVLQLLRLE

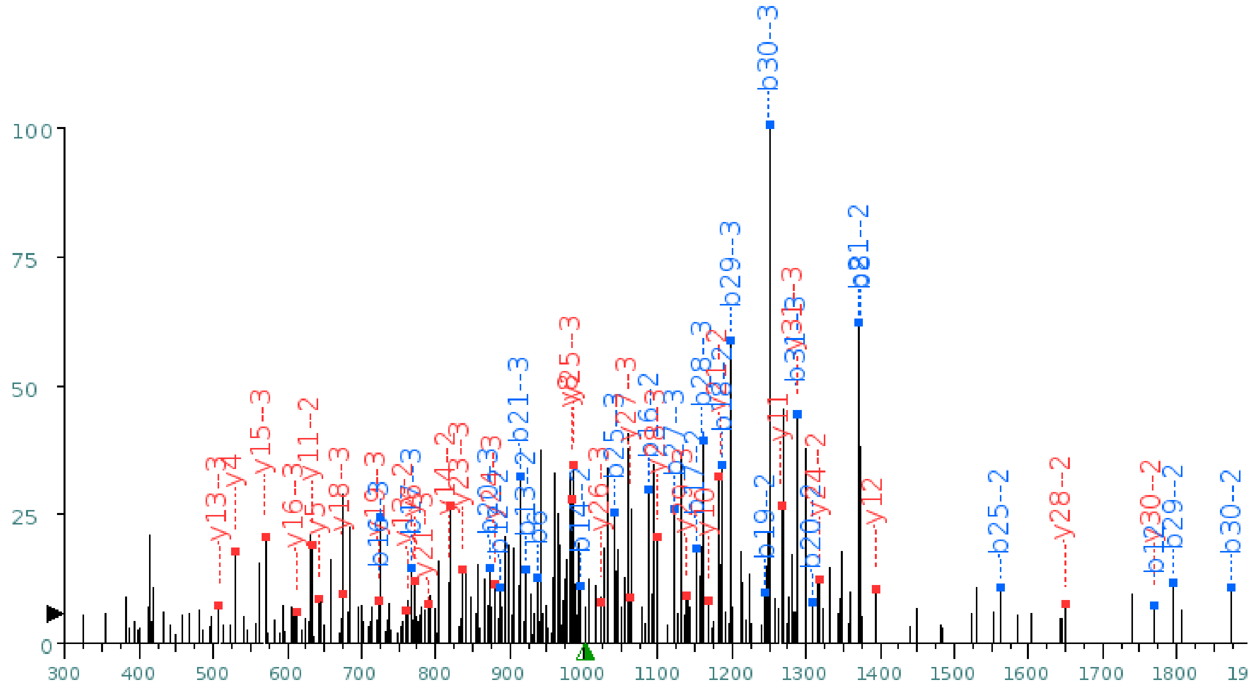

**Predicted Fragmentation Pattern**

| Seq | # | b: Δ Error | b +1 | y +1 | y: Δ Error | +1 |
|---|---|---|---|---|---|---|
| F | 1 | --- | 148.076 | --- | --- | 32 |
| T# | 2 | --- | 469.182 | 3859.959 | --- | 31 |
| K | 3 | --- | 597.277 | 3538.853 | --- | 30 |
| N | 4 | --- | 711.320 | 3410.758 | --- | 29 |
| N | 5 | --- | 825.362 | 3296.715 | --- | 28 |
| I | 6 | -664.503 | **938.447** | 3182.673 | --- | 27 |
| L | 7 | --- | 1051.531 | 3069.588 | --- | 26 |
| T# | 8 | -452.239 | **1372.637** | 2956.504 | --- | 25 |
| Q | 9 | --- | 1500.695 | 2635.398 | --- | 24 |
| A | 10 | --- | 1571.732 | 2507.340 | --- | 23 |

| | | | | | | |
|---|---|---|---|---|---|---|
| A | 11 | --- | 1642.769 | 2436.303 | --- | 22 |
| Q | 12 | -169.798 | 1770.828 | 2365.266 | --- | 21 |
| A | 13 | --- | 1841.865 | 2237.207 | --- | 20 |
| M* | 14 | --- | 1988.900 | 2166.170 | --- | 19 |
| L | 15 | --- | 2101.985 | 2019.135 | --- | 18 |
| A | 16 | --- | 2173.022 | 1906.050 | --- | 17 |
| Q | 17 | --- | 2301.080 | 1835.013 | --- | 16 |
| A | 18 | --- | 2372.117 | 1706.955 | --- | 15 |
| N | 19 | --- | 2486.160 | 1635.918 | --- | 14 |
| Q | 20 | --- | 2614.219 | 1521.875 | --- | 13 |
| Q | 21 | --- | 2742.277 | 1393.816 | 92.203 | 12 |
| P | 22 | --- | 2839.330 | 1265.758 | 483.970 | 11 |
| Q | 23 | --- | 2967.389 | 1168.705 | 394.140 | 10 |
| G | 24 | --- | 3024.410 | 1040.646 | --- | 9 |
| V | 25 | --- | 3123.479 | 983.625 | -389.623 | 8 |
| L | 26 | --- | 3236.563 | 884.556 | --- | 7 |
| Q | 27 | --- | 3364.621 | 771.472 | 428.904 | 6 |
| L | 28 | --- | 3477.705 | 643.414 | 416.773 | 5 |
| L | 29 | --- | 3590.789 | 530.330 | 108.244 | 4 |
| R | 30 | --- | 3746.891 | 417.246 | --- | 3 |
| L | 31 | --- | 3859.975 | 261.144 | --- | 2 |
| E | 32 | --- | --- | 148.060 | --- | 1 |

+2

| Seq | # | b: Δ Error | b | y | y: Δ Error | +1 |
|---|---|---|---|---|---|---|
| F | 1 | --- | 74.541 | --- | --- | 32 |
| T# | 2 | --- | 235.094 | 1930.483 | --- | 31 |
| K | 3 | --- | 299.142 | 1769.930 | 337.200 | 30 |
| N | 4 | --- | 356.163 | 1705.883 | --- | 29 |
| N | 5 | --- | 413.185 | 1648.861 | -104.057 | 28 |
| I | 6 | --- | 469.727 | 1591.840 | --- | 27 |
| L | 7 | --- | 526.269 | 1535.298 | --- | 26 |
| T# | 8 | --- | 686.822 | 1478.756 | --- | 25 |
| Q | 9 | --- | 750.851 | 1318.203 | 311.992 | 24 |
| A | 10 | --- | 786.370 | 1254.174 | --- | 23 |
| A | 11 | --- | 821.888 | 1218.655 | --- | 22 |
| Q | 12 | -23.451 | 885.918 | 1183.136 | -332.217 | 21 |
| A | 13 | 469.123 | 921.436 | 1119.107 | --- | 20 |
| M* | 14 | 16.293 | 994.954 | 1083.589 | --- | 19 |
| L | 15 | --- | 1051.496 | 1010.071 | --- | 18 |
| A | 16 | 427.165 | 1087.014 | 953.529 | --- | 17 |
| Q | 17 | 438.175 | 1151.044 | 918.010 | --- | 16 |
| A | 18 | 738.371 | 1186.562 | 853.981 | --- | 15 |
| N | 19 | 663.384 | 1243.584 | 818.462 | 1094.334 | 14 |
| Q | 20 | 515.115 | 1307.613 | 761.441 | -227.665 | 13 |
| Q | 21 | 272.413 | 1371.642 | 697.412 | --- | 12 |
| P | 22 | --- | 1420.169 | 633.382 | 481.602 | 11 |
| Q | 23 | --- | 1484.198 | 584.856 | --- | 10 |
| G | 24 | --- | 1512.709 | 520.827 | --- | 9 |
| V | 25 | 516.741 | 1562.243 | 492.316 | --- | 8 |
| L | 26 | --- | 1618.785 | 442.782 | --- | 7 |
| Q | 27 | --- | 1682.814 | 386.240 | --- | 6 |
| L | 28 | --- | 1739.356 | 322.211 | --- | 5 |
| L | 29 | -228.934 | 1795.898 | 265.668 | --- | 4 |
| R | 30 | -343.048 | 1873.949 | 209.126 | --- | 3 |
| L | 31 | --- | 1930.491 | 131.076 | --- | 2 |
| E | 32 | --- | --- | 74.534 | --- | 1 |

| | | | +3 | | | |
|---|---|---|---|---|---|---|
| Seq | # | b: Δ Error | b | y | y: Δ Error | +1 |
| F | 1 | --- | 50.030 | --- | --- | 32 |
| T# | 2 | --- | 157.065 | **1287.325** | 158.674 | 31 |
| K | 3 | --- | 199.764 | 1180.289 | --- | 30 |
| N | 4 | --- | 237.778 | **1137.591** | -484.438 | 29 |
| N | 5 | --- | 275.792 | **1099.577** | -447.053 | 28 |
| I | 6 | --- | 313.487 | **1061.562** | 41.175 | 27 |
| L | 7 | --- | 351.182 | **1023.868** | -715.267 | 26 |
| T# | 8 | --- | 458.217 | **986.173** | -403.462 | 25 |
| Q | 9 | --- | 500.903 | **879.138** | 17.806 | 24 |
| A | 10 | --- | 524.582 | **836.451** | 340.144 | 23 |
| A | 11 | --- | 548.261 | 812.772 | --- | 22 |
| Q | 12 | --- | 590.948 | **789.093** | 418.417 | 21 |
| A | 13 | --- | 614.627 | 746.407 | --- | 20 |
| M* | 14 | --- | 663.638 | **722.728** | -419.415 | 19 |
| L | 15 | --- | 701.333 | **673.716** | 1226.509 | 18 |
| A | 16 | 666.486 | **725.012** | 636.022 | --- | 17 |
| Q | 17 | 171.828 | **767.698** | **612.343** | -116.018 | 16 |
| A | 18 | --- | 791.377 | **569.656** | 779.906 | 15 |
| N | 19 | --- | 829.392 | 545.977 | --- | 14 |
| Q | 20 | 624.971 | **872.078** | **507.963** | -1134.960 | 13 |
| Q | 21 | 195.636 | **914.764** | 465.277 | --- | 12 |
| P | 22 | --- | 947.115 | 422.591 | --- | 11 |
| Q | 23 | --- | 989.801 | 390.240 | --- | 10 |
| G | 24 | --- | 1008.808 | 347.554 | --- | 9 |
| V | 25 | 310.035 | **1041.831** | 328.546 | --- | 8 |
| L | 26 | --- | 1079.526 | 295.524 | --- | 7 |
| Q | 27 | 404.126 | **1122.212** | 257.829 | --- | 6 |
| L | 28 | 718.159 | **1159.907** | 215.143 | --- | 5 |
| L | 29 | 465.897 | **1197.601** | 177.448 | --- | 4 |
| R | 30 | 355.430 | **1249.635** | 139.753 | --- | 3 |
| L | 31 | 154.724 | **1287.330** | 87.720 | --- | 2 |
| E | 32 | --- | --- | 50.025 | --- | 1 |

Soleil Index

| | | |
|---|---|---|
| 414.5685 | b3-19 frag doubly charged | 21% |
| 561.4336 | b3-27 frag doubly charged | 16% |
| 630.6921 | not explained | 21% |
| 659.5314 | y2-24 frag doubly charged | 16% |
| 675.6603 | not explained | 29% |
| 683.6199 | b3-15 frag loss of water/ b3-15 frag loss of ammonia | 15% |
| 684.4058 | b3-15 frag loss of ammonia | 23% |
| 803.6722 | b2-11 frag loss of water | 16% |
| 857.6321 | not explained | 15% |
| 894.8301 | not explained | 21% |
| 897.4028 | b2-29 frag doubly charged/ b3-21 frag loss of water/ b3-21 frag loss of ammonia | 19% |
| 904.9591 | b2-13 frag loss of ammonia | 18% |

| | | |
|---|---|---|
| 941.2816 | not explained | 38% |
| 960.2924 | not explained | 33% |
| 965.9703 | b2-31 frag doubly charged/ y1-8 frag loss of water/ y1-8 frag loss of ammonia/ y2-31 frag loss of water | 25% |
| 967.8417 | y3-25 frag loss of water | 19% |
| 975.8750 | not explained | 16% |
| 981.0944 | not explained | 16% |
| 981.9391 | not explained | 33% |
| 984.7358 | not explained | 23% |
| 992.4607 | b3-24 frag loss of ammonia/ y2-18 frag loss of water/ y2-18 frag loss of ammonia | 19% |
| 1027.8898 | not explained | 18% |
| 1032.0641 | not explained | 34% |
| 1046.9413 | not explained | 18% |
| 1059.1705 | not explained | 41% |
| 1064.2806 | not explained | 26% |
| 1094.6733 | not explained | 34% |
| 1130.4529 | not explained | 18% |
| 1131.4674 | not explained | 37% |
| 1184.5316 | not explained | 15% |
| 1198.9347 | not explained | 22% |
| 1212.8677 | not explained | 18% |
| 1250.9712 | not explained | 17% |
| 1270.2784 | b3-31 frag loss of water/ b3-31 frag loss of ammonia/ y3-31 frag loss of water/ y3-31 frag loss of ammonia | 45% |
| 1282.1483 | not explained | 17% |
| 1300.2690 | y2-24 frag loss of water/ y2-24 frag loss of ammonia | 38% |
| 1348.5526 | not explained | 18% |

| | | | | | |
|---|---|---|---|---|---|
| **Run:** | 79490 | **Scans:** | 24436 - 24436 | **m/z Range:** | 269.64487 to 1991.1652[illegible] |
| **Search:** | 79490 | **Precursor Mass:** | 998.5118 | | |
| **Mods:** | C=160.03 *+15.99 #+220.06 @+291.1 | **Charge:** | +4 | | |

**Sequence**

FT#KNNILT#QAAQAMLAQANQQPQGVLQLLRLE

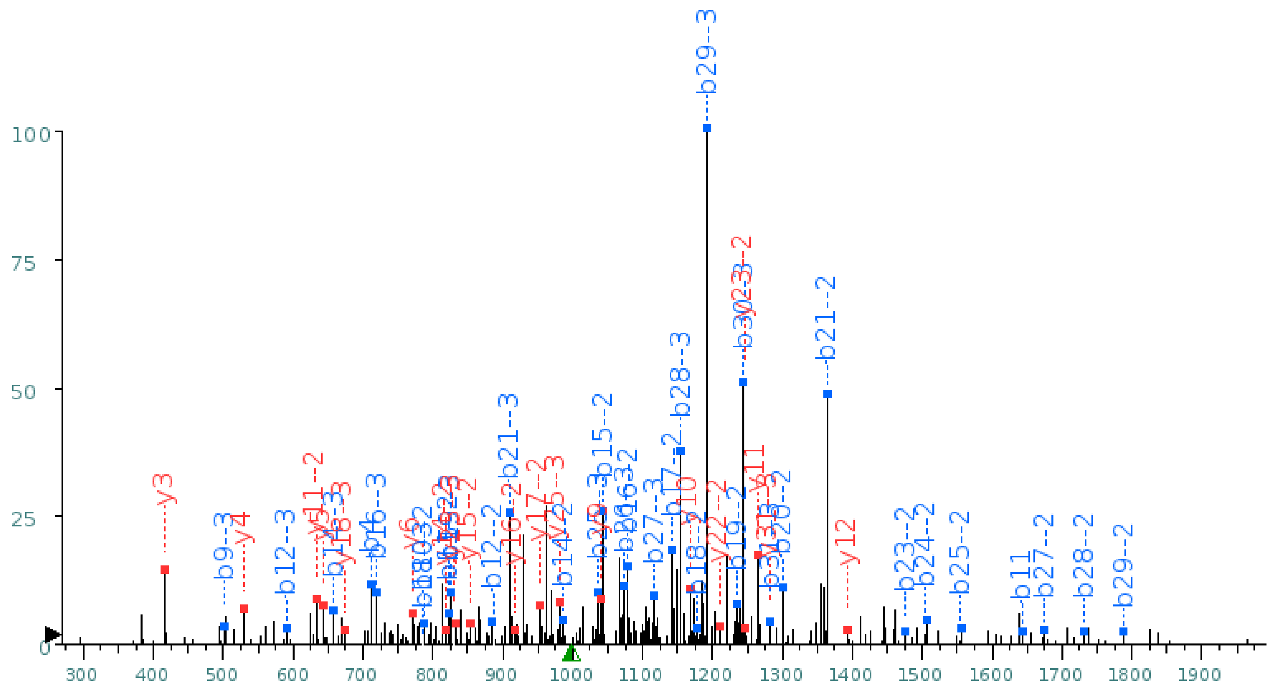

**Predicted Fragmentation Pattern**

| Seq | # | b: Δ Error | b | y | y: Δ Error | +1 |
|---|---|---|---|---|---|---|
| F | 1 | --- | 148.076 | --- | --- | 32 |
| T# | 2 | --- | 469.182 | 3843.964 | --- | 31 |
| K | 3 | --- | 597.277 | 3522.858 | --- | 30 |
| N | 4 | -143.715 | **711.320** | 3394.763 | --- | 29 |
| N | 5 | -1198.305 | **825.362** | 3280.721 | --- | 28 |
| I | 6 | --- | 938.447 | 3166.678 | --- | 27 |
| L | 7 | --- | 1051.531 | 3053.594 | --- | 26 |
| T# | 8 | --- | 1372.637 | 2940.510 | --- | 25 |

| | | | | | | |
|---|---|---|---|---|---|---|
| Q | 9 | --- | 1500.695 | 2619.404 | --- | 24 |
| A | 10 | --- | 1571.732 | 2491.345 | --- | 23 |
| A | 11 | -128.562 | 1642.769 | 2420.308 | --- | 22 |
| Q | 12 | --- | 1770.828 | 2349.271 | --- | 21 |
| A | 13 | --- | 1841.865 | 2221.212 | --- | 20 |
| M | 14 | --- | 1972.906 | 2150.175 | --- | 19 |
| L | 15 | --- | 2085.990 | 2019.135 | --- | 18 |
| A | 16 | --- | 2157.027 | 1906.050 | --- | 17 |
| Q | 17 | --- | 2285.085 | 1835.013 | --- | 16 |
| A | 18 | --- | 2356.122 | 1706.955 | --- | 15 |
| N | 19 | --- | 2470.165 | 1635.918 | --- | 14 |
| Q | 20 | --- | 2598.224 | 1521.875 | --- | 13 |
| Q | 21 | --- | 2726.283 | 1393.816 | 135.195 | 12 |
| P | 22 | --- | 2823.335 | 1265.758 | 126.298 | 11 |
| Q | 23 | --- | 2951.394 | 1168.705 | 310.953 | 10 |
| G | 24 | --- | 3008.415 | 1040.646 | 222.457 | 9 |
| V | 25 | --- | 3107.484 | 983.625 | --- | 8 |
| L | 26 | --- | 3220.568 | 884.556 | --- | 7 |
| Q | 27 | --- | 3348.626 | 771.472 | 215.826 | 6 |
| L | 28 | --- | 3461.710 | 643.414 | 257.418 | 5 |
| L | 29 | --- | 3574.795 | 530.330 | -105.015 | 4 |
| R | 30 | --- | 3730.896 | 417.246 | 335.373 | 3 |
| L | 31 | --- | 3843.980 | 261.144 | --- | 2 |
| E | 32 | --- | --- | 148.060 | --- | 1 |

+2

| Seq | # | b: Δ Error | b | y | y: Δ Error | +1 |
|---|---|---|---|---|---|---|
| F | 1 | --- | 74.541 | --- | --- | 32 |
| T# | 2 | --- | 235.094 | 1922.486 | --- | 31 |
| K | 3 | --- | 299.142 | 1761.933 | --- | 30 |
| N | 4 | --- | 356.163 | 1697.885 | --- | 29 |
| N | 5 | --- | 413.185 | 1640.864 | --- | 28 |
| I | 6 | --- | 469.727 | 1583.842 | --- | 27 |
| L | 7 | --- | 526.269 | 1527.300 | --- | 26 |
| T# | 8 | --- | 686.822 | 1470.758 | --- | 25 |
| Q | 9 | --- | 750.851 | 1310.205 | --- | 24 |
| A | 10 | 490.878 | 786.370 | 1246.176 | 642.107 | 23 |
| A | 11 | 237.693 | 821.888 | 1210.658 | 629.617 | 22 |
| Q | 12 | -328.487 | 885.918 | 1175.139 | --- | 21 |
| A | 13 | --- | 921.436 | 1111.110 | --- | 20 |
| M | 14 | 178.440 | 986.956 | 1075.591 | --- | 19 |
| L | 15 | 550.055 | 1043.498 | 1010.071 | --- | 18 |
| A | 16 | -54.122 | 1079.017 | 953.529 | 649.074 | 17 |
| Q | 17 | -152.238 | 1143.046 | 918.010 | 719.586 | 16 |
| A | 18 | -301.840 | 1178.565 | 853.981 | 750.937 | 15 |
| N | 19 | -129.158 | 1235.586 | 818.462 | 795.862 | 14 |
| Q | 20 | -17.667 | 1299.616 | 761.441 | --- | 13 |
| Q | 21 | -165.817 | 1363.645 | 697.412 | --- | 12 |
| P | 22 | --- | 1412.171 | 633.382 | 262.441 | 11 |
| Q | 23 | 395.106 | 1476.201 | 584.856 | --- | 10 |
| G | 24 | 416.649 | 1504.711 | 520.827 | --- | 9 |
| V | 25 | 488.341 | 1554.246 | 492.316 | --- | 8 |
| L | 26 | --- | 1610.788 | 442.782 | --- | 7 |
| Q | 27 | 67.307 | 1674.817 | 386.240 | --- | 6 |
| L | 28 | -192.856 | 1731.359 | 322.211 | --- | 5 |
| L | 29 | 412.070 | 1787.901 | 265.668 | --- | 4 |
| R | 30 | --- | 1865.951 | 209.126 | --- | 3 |
| L | 31 | --- | 1922.493 | 131.076 | --- | 2 |

| | | | | | | |
|---|---|---|---|---|---|---|
| E | 32 | --- | --- | 74.534 | --- | 1 |

+3

| Seq | # | b: Δ Error | b | y | y: Δ Error | +1 |
|---|---|---|---|---|---|---|
| F | 1 | --- | 50.030 | --- | --- | 32 |
| T# | 2 | --- | 157.065 | **1281.993** | 462.778 | 31 |
| K | 3 | --- | 199.764 | 1174.958 | --- | 30 |
| N | 4 | --- | 237.778 | 1132.259 | --- | 29 |
| N | 5 | --- | 275.792 | 1094.245 | --- | 28 |
| I | 6 | --- | 313.487 | 1056.231 | --- | 27 |
| L | 7 | --- | 351.182 | 1018.536 | --- | 26 |
| T# | 8 | --- | 458.217 | **980.841** | -231.585 | 25 |
| Q | 9 | -1064.970 | **500.903** | 873.806 | --- | 24 |
| A | 10 | --- | 524.582 | **831.120** | 589.546 | 23 |
| A | 11 | --- | 548.261 | 807.441 | --- | 22 |
| Q | 12 | -308.705 | **590.948** | 783.762 | --- | 21 |
| A | 13 | --- | 614.627 | 741.076 | --- | 20 |
| M | 14 | -461.297 | **658.307** | 717.397 | --- | 19 |
| L | 15 | --- | 696.001 | **673.716** | -447.644 | 18 |
| A | 16 | -517.197 | **719.680** | 636.022 | --- | 17 |
| Q | 17 | --- | 762.367 | 612.343 | --- | 16 |
| A | 18 | 902.842 | **786.046** | 569.656 | --- | 15 |
| N | 19 | 381.695 | **824.060** | 545.977 | --- | 14 |
| Q | 20 | --- | 866.746 | 507.963 | --- | 13 |
| Q | 21 | 107.385 | **909.432** | 465.277 | --- | 12 |
| P | 22 | --- | 941.783 | 422.591 | --- | 11 |
| Q | 23 | --- | 984.469 | 390.240 | --- | 10 |
| G | 24 | --- | 1003.477 | 347.554 | --- | 9 |
| V | 25 | -20.416 | **1036.499** | 328.546 | --- | 8 |
| L | 26 | 32.473 | **1074.194** | 295.524 | --- | 7 |
| Q | 27 | 116.111 | **1116.880** | 257.829 | --- | 6 |
| L | 28 | 52.184 | **1154.575** | 215.143 | --- | 5 |
| L | 29 | 20.345 | **1192.270** | 177.448 | --- | 4 |
| R | 30 | -103.940 | **1244.303** | 139.753 | --- | 3 |
| L | 31 | 458.813 | **1281.998** | 87.720 | --- | 2 |
| E | 32 | --- | --- | 50.025 | --- | 1 |

### Soleil Index

| | | |
|---|---|---|
| 929.2618 | not explained | 21% |
| 961.9838 | b2-31 frag doubly charged/ y2-31 frag loss of water/ y3-25 frag loss of water | 27% |
| 1066.9878 | not explained | 17% |
| 1219.5100 | b2-19 frag loss of ammonia | 17% |

| | | | | | |
|---|---|---|---|---|---|
| **Run:** | 79490 | **Scans:** | 24547 - 24547 | **m/z Range:** | 392.2576 to 2000 |
| **Search:** | 79490 | **Precursor Mass:** | 998.51182 | | |
| **Mods:** | C=160.03 *+15.99 #+220.06 @+291.1 | **Charge:** | +4 | | |

**Sequence**

FT#KNNILT#QAAQAMLAQANQQPQGVLQLLRLE

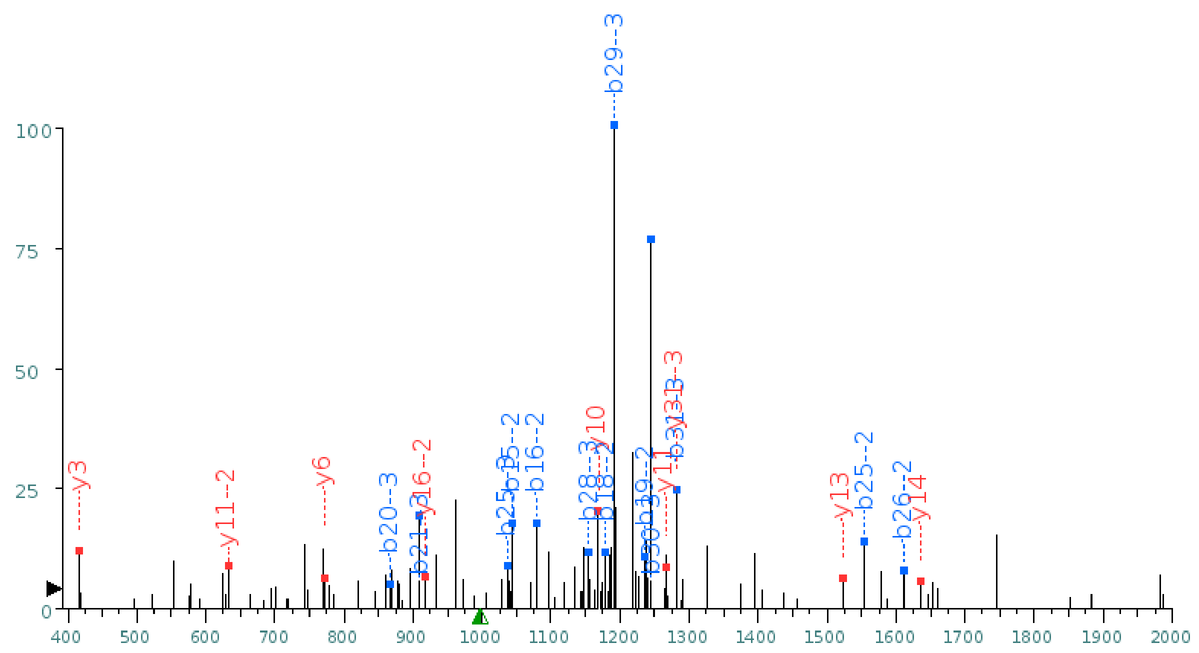

**Predicted Fragmentation Pattern**

| | | | +1 | | | |
|---|---|---|---|---|---|---|
| **Seq** | **#** | **b: Δ Error** | **b** | **y** | **y: Δ Error** | **+1** |
| **F** | **1** | --- | 148.076 | --- | --- | **32** |
| **T#** | **2** | --- | 469.182 | 3843.964 | --- | **31** |
| **K** | **3** | --- | 597.277 | 3522.858 | --- | **30** |
| **N** | **4** | --- | 711.320 | 3394.763 | --- | **29** |
| **N** | **5** | --- | 825.362 | 3280.721 | --- | **28** |
| **I** | **6** | --- | 938.447 | 3166.678 | --- | **27** |
| **L** | **7** | --- | 1051.531 | 3053.594 | --- | **26** |
| **T#** | **8** | --- | 1372.637 | 2940.510 | --- | **25** |
| **Q** | **9** | --- | 1500.695 | 2619.404 | --- | **24** |
| **A** | **10** | --- | 1571.732 | 2491.345 | --- | **23** |
| **A** | **11** | --- | 1642.769 | 2420.308 | --- | **22** |
| **Q** | **12** | --- | 1770.828 | 2349.271 | --- | **21** |
| **A** | **13** | --- | 1841.865 | 2221.212 | --- | **20** |
| **M** | **14** | --- | 1972.906 | 2150.175 | --- | **19** |
| **L** | **15** | --- | 2085.990 | 2019.135 | --- | **18** |
| **A** | **16** | --- | 2157.027 | 1906.050 | --- | **17** |

| Seq | # | b: Δ Error | b | y | y: Δ Error | +1 |
|---|---|---|---|---|---|---|
| Q | 17 | --- | 2285.085 | 1835.013 | --- | 16 |
| A | 18 | --- | 2356.122 | 1706.955 | --- | 15 |
| N | 19 | --- | 2470.165 | **1635.918** | 101.274 | 14 |
| Q | 20 | --- | 2598.224 | **1521.875** | 156.628 | 13 |
| Q | 21 | --- | 2726.283 | 1393.816 | --- | 12 |
| P | 22 | --- | 2823.335 | **1265.758** | 235.525 | 11 |
| Q | 23 | --- | 2951.394 | **1168.705** | -661.287 | 10 |
| G | 24 | --- | 3008.415 | 1040.646 | --- | 9 |
| V | 25 | --- | 3107.484 | 983.625 | --- | 8 |
| L | 26 | --- | 3220.568 | 884.556 | --- | 7 |
| Q | 27 | --- | 3348.626 | **771.472** | 334.038 | 6 |
| L | 28 | --- | 3461.710 | 643.414 | --- | 5 |
| L | 29 | --- | 3574.795 | 530.330 | --- | 4 |
| R | 30 | --- | 3730.896 | **417.246** | 28.734 | 3 |
| L | 31 | --- | 3843.980 | 261.144 | --- | 2 |
| E | 32 | --- | --- | 148.060 | --- | 1 |

+2

| Seq | # | b: Δ Error | b | y | y: Δ Error | +1 |
|---|---|---|---|---|---|---|
| F | 1 | --- | 74.541 | --- | --- | 32 |
| T# | 2 | --- | 235.094 | 1922.486 | --- | 31 |
| K | 3 | --- | 299.142 | 1761.933 | --- | 30 |
| N | 4 | --- | 356.163 | 1697.885 | --- | 29 |
| N | 5 | --- | 413.185 | 1640.864 | --- | 28 |
| I | 6 | --- | 469.727 | 1583.842 | --- | 27 |
| L | 7 | --- | 526.269 | 1527.300 | --- | 26 |
| T# | 8 | --- | 686.822 | 1470.758 | --- | 25 |
| Q | 9 | --- | 750.851 | 1310.205 | --- | 24 |
| A | 10 | --- | 786.370 | 1246.176 | --- | 23 |
| A | 11 | --- | 821.888 | 1210.658 | --- | 22 |
| Q | 12 | --- | 885.918 | 1175.139 | --- | 21 |
| A | 13 | --- | 921.436 | 1111.110 | --- | 20 |
| M | 14 | --- | 986.956 | 1075.591 | --- | 19 |
| L | 15 | 252.458 | **1043.498** | 1010.071 | --- | 18 |
| A | 16 | 250.819 | **1079.017** | 953.529 | --- | 17 |
| Q | 17 | --- | 1143.046 | **918.010** | 305.933 | 16 |
| A | 18 | 151.577 | **1178.565** | 853.981 | --- | 15 |
| N | 19 | 745.137 | **1235.586** | 818.462 | --- | 14 |
| Q | 20 | --- | 1299.616 | 761.441 | --- | 13 |
| Q | 21 | --- | 1363.645 | 697.412 | --- | 12 |
| P | 22 | --- | 1412.171 | **633.382** | 516.451 | 11 |
| Q | 23 | --- | 1476.201 | 584.856 | --- | 10 |
| G | 24 | --- | 1504.711 | 520.827 | --- | 9 |
| V | 25 | 107.178 | **1554.246** | 492.316 | --- | 8 |
| L | 26 | -138.895 | **1610.788** | 442.782 | --- | 7 |
| Q | 27 | --- | 1674.817 | 386.240 | --- | 6 |
| L | 28 | --- | 1731.359 | 322.211 | --- | 5 |
| L | 29 | --- | 1787.901 | 265.668 | --- | 4 |
| R | 30 | --- | 1865.951 | 209.126 | --- | 3 |
| L | 31 | --- | 1922.493 | 131.076 | --- | 2 |
| E | 32 | --- | --- | 74.534 | --- | 1 |

+3

| Seq | # | b: Δ Error | b | y | y: Δ Error | +1 |
|---|---|---|---|---|---|---|
| F | 1 | --- | 50.030 | --- | --- | 32 |
| T# | 2 | --- | 157.065 | **1281.993** | 232.317 | 31 |
| K | 3 | --- | 199.764 | 1174.958 | --- | 30 |
| N | 4 | --- | 237.778 | 1132.259 | --- | 29 |

| | | | | | | |
|---|---|---|---|---|---|---|
| **N** | **5** | --- | 275.792 | 1094.245 | --- | **28** |
| **I** | **6** | --- | 313.487 | 1056.231 | --- | **27** |
| **L** | **7** | --- | 351.182 | 1018.536 | --- | **26** |
| **T#** | **8** | --- | 458.217 | 980.841 | --- | **25** |
| **Q** | **9** | --- | 500.903 | 873.806 | --- | **24** |
| **A** | **10** | --- | 524.582 | 831.120 | --- | **23** |
| **A** | **11** | --- | 548.261 | 807.441 | --- | **22** |
| **Q** | **12** | --- | 590.948 | 783.762 | --- | **21** |
| **A** | **13** | --- | 614.627 | 741.076 | --- | **20** |
| **M** | **14** | --- | 658.307 | 717.397 | --- | **19** |
| **L** | **15** | --- | 696.001 | 673.716 | --- | **18** |
| **A** | **16** | --- | 719.680 | 636.022 | --- | **17** |
| **Q** | **17** | --- | 762.367 | 612.343 | --- | **16** |
| **A** | **18** | --- | 786.046 | 569.656 | --- | **15** |
| **N** | **19** | --- | 824.060 | 545.977 | --- | **14** |
| **Q** | **20** | **242.520** | **866.746** | 507.963 | --- | **13** |
| **Q** | **21** | **664.400** | **909.432** | 465.277 | --- | **12** |
| **P** | **22** | --- | 941.783 | 422.591 | --- | **11** |
| **Q** | **23** | --- | 984.469 | 390.240 | --- | **10** |
| **G** | **24** | --- | 1003.477 | 347.554 | --- | **9** |
| **V** | **25** | **423.522** | **1036.499** | 328.546 | --- | **8** |
| **L** | **26** | --- | 1074.194 | 295.524 | --- | **7** |
| **Q** | **27** | --- | 1116.880 | 257.829 | --- | **6** |
| **L** | **28** | **-8.078** | **1154.575** | 215.143 | --- | **5** |
| **L** | **29** | **94.770** | **1192.270** | 177.448 | --- | **4** |
| **R** | **30** | **50.581** | **1244.303** | 139.753 | --- | **3** |
| **L** | **31** | **228.352** | **1281.998** | 87.720 | --- | **2** |
| **E** | **32** | --- | --- | 50.025 | --- | **1** |

**Soleil Index**

| | | |
|---|---|---|
| **962.0328** | **b2-31 frag doubly charged/ y2-31 frag loss of water/ y3-25 frag loss of water** | **23%** |
| **1219.2465** | **b2-19 frag loss of ammonia** | **33%** |
| **1745.5389** | **y2-30 frag loss of ammonia** | **15%** |

| | | | | | |
|---|---|---|---|---|---|
| **Run:** | 79490 | **Scans:** | 24615 - 24615 | **m/z Range:** | 312.30164 to 1821.3184 |
| **Search:** | 79490 | **Precursor Mass:** | 1002.51124 | | |
| **Mods:** | C=160.03 *+15.99 #+220.06 @+291.1 | **Charge:** | +4 | | |

**Sequence**

FT#KNNILT#QAAQAM*LAQANQQPQGVLQLLRLE

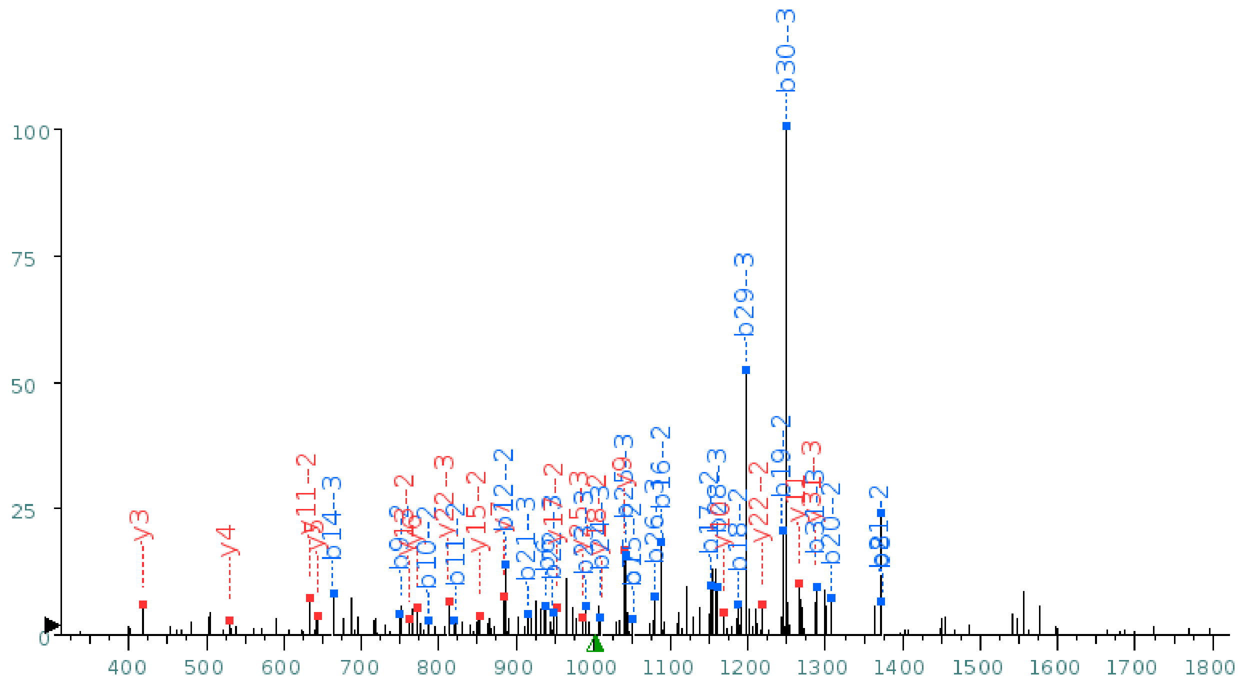

**Predicted Fragmentation Pattern**

| Seq | # | b: Δ Error | +1 b | y | y: Δ Error | +1 |
|---|---|---|---|---|---|---|
| F | 1 | --- | 148.076 | --- | --- | 32 |
| T# | 2 | --- | 469.182 | 3859.959 | --- | 31 |
| K | 3 | --- | 597.277 | 3538.853 | --- | 30 |
| N | 4 | --- | 711.320 | 3410.758 | --- | 29 |
| N | 5 | --- | 825.362 | 3296.715 | --- | 28 |
| I | 6 | -542.603 | **938.447** | 3182.673 | --- | 27 |
| L | 7 | -332.215 | **1051.531** | 3069.588 | --- | 26 |
| T# | 8 | -69.902 | **1372.637** | 2956.504 | --- | 25 |
| Q | 9 | --- | 1500.695 | 2635.398 | --- | 24 |
| A | 10 | --- | 1571.732 | 2507.340 | --- | 23 |
| A | 11 | --- | 1642.769 | 2436.303 | --- | 22 |

| | | | | | | |
|---|---|---|---|---|---|---|
| Q | 12 | --- | 1770.828 | 2365.266 | --- | 21 |
| A | 13 | --- | 1841.865 | 2237.207 | --- | 20 |
| M* | 14 | --- | 1988.900 | 2166.170 | --- | 19 |
| L | 15 | --- | 2101.985 | 2019.135 | --- | 18 |
| A | 16 | --- | 2173.022 | 1906.050 | --- | 17 |
| Q | 17 | --- | 2301.080 | 1835.013 | --- | 16 |
| A | 18 | --- | 2372.117 | 1706.955 | --- | 15 |
| N | 19 | --- | 2486.160 | 1635.918 | --- | 14 |
| Q | 20 | --- | 2614.219 | 1521.875 | --- | 13 |
| Q | 21 | --- | 2742.277 | 1393.816 | --- | 12 |
| P | 22 | --- | 2839.330 | **1265.758** | 133.240 | 11 |
| Q | 23 | --- | 2967.389 | **1168.705** | -126.818 | 10 |
| G | 24 | --- | 3024.410 | **1040.646** | 12.301 | 9 |
| V | 25 | --- | 3123.479 | 983.625 | --- | 8 |
| L | 26 | --- | 3236.563 | **884.556** | 56.045 | 7 |
| Q | 27 | --- | 3364.621 | **771.472** | 156.749 | 6 |
| L | 28 | --- | 3477.705 | **643.414** | 173.692 | 5 |
| L | 29 | --- | 3590.789 | **530.330** | 256.655 | 4 |
| R | 30 | --- | 3746.891 | **417.246** | 173.742 | 3 |
| L | 31 | --- | 3859.975 | 261.144 | --- | 2 |
| E | 32 | --- | --- | 148.060 | --- | 1 |

+2

| Seq | # | b: Δ Error | b | y | y: Δ Error | +1 |
|---|---|---|---|---|---|---|
| F | 1 | --- | 74.541 | --- | --- | 32 |
| T# | 2 | --- | 235.094 | 1930.483 | --- | 31 |
| K | 3 | --- | 299.142 | 1769.930 | --- | 30 |
| N | 4 | --- | 356.163 | 1705.883 | --- | 29 |
| N | 5 | --- | 413.185 | 1648.861 | --- | 28 |
| I | 6 | --- | 469.727 | 1591.840 | --- | 27 |
| L | 7 | --- | 526.269 | 1535.298 | --- | 26 |
| T# | 8 | --- | 686.822 | 1478.756 | --- | 25 |
| Q | 9 | -245.423 | **750.851** | 1318.203 | --- | 24 |
| A | 10 | 242.843 | **786.370** | 1254.174 | --- | 23 |
| A | 11 | -998.076 | **821.888** | **1218.655** | 178.580 | 22 |
| Q | 12 | -184.147 | **885.918** | 1183.136 | --- | 21 |
| A | 13 | --- | 921.436 | 1119.107 | --- | 20 |
| M* | 14 | --- | 994.954 | 1083.589 | --- | 19 |
| L | 15 | -299.204 | **1051.496** | **1010.071** | -308.986 | 18 |
| A | 16 | 306.420 | **1087.014** | **953.529** | -725.064 | 17 |
| Q | 17 | -175.598 | **1151.044** | 918.010 | --- | 16 |
| A | 18 | 493.722 | **1186.562** | **853.981** | 322.140 | 15 |
| N | 19 | 486.997 | **1243.584** | 818.462 | --- | 14 |
| Q | 20 | -129.896 | **1307.613** | **761.441** | 191.813 | 13 |
| Q | 21 | -49.680 | **1371.642** | 697.412 | --- | 12 |
| P | 22 | --- | 1420.169 | **633.382** | 170.550 | 11 |
| Q | 23 | --- | 1484.198 | 584.856 | --- | 10 |
| G | 24 | --- | 1512.709 | 520.827 | --- | 9 |
| V | 25 | --- | 1562.243 | 492.316 | --- | 8 |
| L | 26 | --- | 1618.785 | 442.782 | --- | 7 |
| Q | 27 | --- | 1682.814 | 386.240 | --- | 6 |
| L | 28 | --- | 1739.356 | 322.211 | --- | 5 |
| L | 29 | --- | 1795.898 | 265.668 | --- | 4 |
| R | 30 | --- | 1873.949 | 209.126 | --- | 3 |
| L | 31 | --- | 1930.491 | 131.076 | --- | 2 |
| E | 32 | --- | --- | 74.534 | --- | 1 |

+3

| Seq | # | b: Δ Error | b | y | y: Δ Error | +1 |
|---|---|---|---|---|---|---|
| F | 1 | --- | 50.030 | --- | --- | 32 |
| T# | 2 | --- | 157.065 | **1287.325** | 462.209 | 31 |
| K | 3 | --- | 199.764 | 1180.289 | --- | 30 |
| N | 4 | --- | 237.778 | 1137.591 | --- | 29 |
| N | 5 | --- | 275.792 | 1099.577 | --- | 28 |
| I | 6 | --- | 313.487 | 1061.562 | --- | 27 |
| L | 7 | --- | 351.182 | 1023.868 | --- | 26 |
| T# | 8 | --- | 458.217 | **986.173** | -154.151 | 25 |
| Q | 9 | --- | 500.903 | 879.138 | --- | 24 |
| A | 10 | --- | 524.582 | 836.451 | --- | 23 |
| A | 11 | --- | 548.261 | **812.772** | 245.982 | 22 |
| Q | 12 | --- | 590.948 | 789.093 | --- | 21 |
| A | 13 | --- | 614.627 | 746.407 | --- | 20 |
| M* | 14 | 230.363 | **663.638** | 722.728 | --- | 19 |
| L | 15 | --- | 701.333 | 673.716 | --- | 18 |
| A | 16 | --- | 725.012 | 636.022 | --- | 17 |
| Q | 17 | --- | 767.698 | 612.343 | --- | 16 |
| A | 18 | --- | 791.377 | 569.656 | --- | 15 |
| N | 19 | --- | 829.392 | 545.977 | --- | 14 |
| Q | 20 | --- | 872.078 | 507.963 | --- | 13 |
| Q | 21 | 526.873 | **914.764** | 465.277 | --- | 12 |
| P | 22 | 705.875 | **947.115** | 422.591 | --- | 11 |
| Q | 23 | -154.641 | **989.801** | 390.240 | --- | 10 |
| G | 24 | 941.458 | **1008.808** | 347.554 | --- | 9 |
| V | 25 | 219.043 | **1041.831** | 328.546 | --- | 8 |
| L | 26 | 215.704 | **1079.526** | 295.524 | --- | 7 |
| Q | 27 | --- | 1122.212 | 257.829 | --- | 6 |
| L | 28 | 278.162 | **1159.907** | 215.143 | --- | 5 |
| L | 29 | -48.528 | **1197.601** | 177.448 | --- | 4 |
| R | 30 | 99.319 | **1249.635** | 139.753 | --- | 3 |
| L | 31 | 458.261 | **1287.330** | 87.720 | --- | 2 |
| E | 32 | --- | --- | 50.025 | --- | 1 |

| Run: | 79490 | Scans: | 25063 - 25063 | m/z Range: | 330.5773 to 1979.0393 |
|---|---|---|---|---|---|
| Search: | 79490 | Precursor Mass: | 1002.51044 | | |
| Mods: | C=160.03 *+15.99 #+220.06 @+291.1 | Charge: | +4 | | |

**Sequence**

FT#KNNILT#QAAQAM*LAQANQQPQGVLQLLRLE

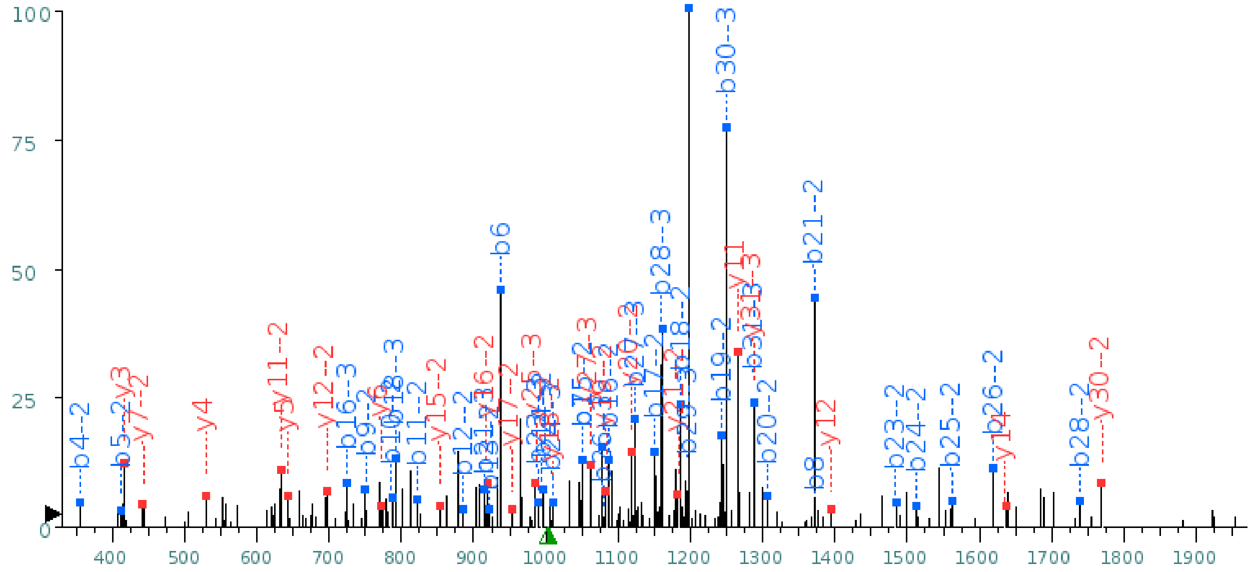

**Predicted Fragmentation Pattern**

| Seq | # | b: Δ Error | b | y | y: Δ Error | +1 |
|---|---|---|---|---|---|---|
| F | 1 | --- | 148.076 | --- | --- | 32 |
| T# | 2 | --- | 469.182 | 3859.959 | --- | 31 |
| K | 3 | --- | 597.277 | 3538.853 | --- | 30 |
| N | 4 | --- | 711.320 | 3410.758 | --- | 29 |
| N | 5 | --- | 825.362 | 3296.715 | --- | 28 |
| I | 6 | -936.929 | **938.447** | 3182.673 | --- | 27 |
| L | 7 | -8.450 | **1051.531** | 3069.588 | --- | 26 |

| | | | | | | |
|---|---|---|---|---|---|---|
| T# | 8 | -393.584 | 1372.637 | 2956.504 | --- | 25 |
| Q | 9 | --- | 1500.695 | 2635.398 | --- | 24 |
| A | 10 | --- | 1571.732 | 2507.340 | --- | 23 |
| A | 11 | --- | 1642.769 | 2436.303 | --- | 22 |
| Q | 12 | --- | 1770.828 | 2365.266 | --- | 21 |
| A | 13 | --- | 1841.865 | 2237.207 | --- | 20 |
| M* | 14 | --- | 1988.900 | 2166.170 | --- | 19 |
| L | 15 | --- | 2101.985 | 2019.135 | --- | 18 |
| A | 16 | --- | 2173.022 | 1906.050 | --- | 17 |
| Q | 17 | --- | 2301.080 | 1835.013 | --- | 16 |
| A | 18 | --- | 2372.117 | 1706.955 | --- | 15 |
| N | 19 | --- | 2486.160 | 1635.918 | 149.466 | 14 |
| Q | 20 | --- | 2614.219 | 1521.875 | --- | 13 |
| Q | 21 | --- | 2742.277 | 1393.816 | 72.414 | 12 |
| P | 22 | --- | 2839.330 | 1265.758 | 146.737 | 11 |
| Q | 23 | --- | 2967.389 | 1168.705 | --- | 10 |
| G | 24 | --- | 3024.410 | 1040.646 | --- | 9 |
| V | 25 | --- | 3123.479 | 983.625 | --- | 8 |
| L | 26 | --- | 3236.563 | 884.556 | --- | 7 |
| Q | 27 | --- | 3364.621 | 771.472 | 158.806 | 6 |
| L | 28 | --- | 3477.705 | 643.414 | 181.278 | 5 |
| L | 29 | --- | 3590.789 | 530.330 | 417.441 | 4 |
| R | 30 | --- | 3746.891 | 417.246 | 207.959 | 3 |
| L | 31 | --- | 3859.975 | 261.144 | --- | 2 |
| E | 32 | --- | --- | 148.060 | --- | 1 |

+2

| Seq | # | b: Δ Error | b | y | y: Δ Error | +1 |
|---|---|---|---|---|---|---|
| F | 1 | --- | 74.541 | --- | --- | 32 |
| T# | 2 | --- | 235.094 | 1930.483 | --- | 31 |
| K | 3 | --- | 299.142 | 1769.930 | -524.937 | 30 |
| N | 4 | -1648.352 | 356.163 | 1705.883 | --- | 29 |
| N | 5 | -2086.750 | 413.185 | 1648.861 | --- | 28 |
| I | 6 | --- | 469.727 | 1591.840 | --- | 27 |
| L | 7 | --- | 526.269 | 1535.298 | --- | 26 |
| T# | 8 | --- | 686.822 | 1478.756 | --- | 25 |
| Q | 9 | -1011.467 | 750.851 | 1318.203 | --- | 24 |
| A | 10 | 814.736 | 786.370 | 1254.174 | --- | 23 |
| A | 11 | 382.414 | 821.888 | 1218.655 | --- | 22 |
| Q | 12 | -194.485 | 885.918 | 1183.136 | -119.168 | 21 |
| A | 13 | -898.109 | 921.436 | 1119.107 | -247.771 | 20 |
| M* | 14 | 702.492 | 994.954 | 1083.589 | -102.854 | 19 |
| L | 15 | 24.550 | 1051.496 | 1010.071 | -551.022 | 18 |
| A | 16 | 6.564 | 1087.014 | 953.529 | -149.369 | 17 |
| Q | 17 | -214.958 | 1151.044 | 918.010 | 623.443 | 16 |
| A | 18 | 29.165 | 1186.562 | 853.981 | 709.116 | 15 |
| N | 19 | 110.189 | 1243.584 | 818.462 | --- | 14 |
| Q | 20 | -88.438 | 1307.613 | 761.441 | --- | 13 |
| Q | 21 | 331.026 | 1371.642 | 697.412 | 282.974 | 12 |
| P | 22 | --- | 1420.169 | 633.382 | 478.136 | 11 |
| Q | 23 | 136.819 | 1484.198 | 584.856 | --- | 10 |
| G | 24 | 402.097 | 1512.709 | 520.827 | --- | 9 |
| V | 25 | 110.522 | 1562.243 | 492.316 | --- | 8 |
| L | 26 | 181.874 | 1618.785 | 442.782 | -1057.071 | 7 |
| Q | 27 | --- | 1682.814 | 386.240 | --- | 6 |
| L | 28 | 114.525 | 1739.356 | 322.211 | --- | 5 |
| L | 29 | --- | 1795.898 | 265.668 | --- | 4 |
| R | 30 | --- | 1873.949 | 209.126 | --- | 3 |

| | | | | | | |
|---|---|---|---|---|---|---|
| L | 31 | --- | 1930.491 | 131.076 | --- | 2 |
| E | 32 | --- | --- | 74.534 | --- | 1 |

+3

| Seq | # | b: Δ Error | b | y | y: Δ Error | +1 |
|---|---|---|---|---|---|---|
| F | 1 | --- | 50.030 | --- | --- | 32 |
| T# | 2 | --- | 157.065 | **1287.325** | **667.747** | 31 |
| K | 3 | --- | 199.764 | 1180.289 | --- | 30 |
| N | 4 | --- | 237.778 | 1137.591 | --- | 29 |
| N | 5 | --- | 275.792 | 1099.577 | --- | 28 |
| I | 6 | --- | 313.487 | **1061.562** | **670.992** | 27 |
| L | 7 | --- | 351.182 | 1023.868 | --- | 26 |
| T# | 8 | --- | 458.217 | **986.173** | **-576.555** | 25 |
| Q | 9 | --- | 500.903 | 879.138 | --- | 24 |
| A | 10 | --- | 524.582 | 836.451 | --- | 23 |
| A | 11 | --- | 548.261 | 812.772 | --- | 22 |
| Q | 12 | --- | 590.948 | 789.093 | --- | 21 |
| A | 13 | --- | 614.627 | 746.407 | --- | 20 |
| M* | 14 | --- | 663.638 | 722.728 | --- | 19 |
| L | 15 | --- | 701.333 | 673.716 | --- | 18 |
| A | 16 | **-470.293** | **725.012** | 636.022 | --- | 17 |
| Q | 17 | --- | 767.698 | 612.343 | --- | 16 |
| A | 18 | **140.831** | **791.377** | 569.656 | --- | 15 |
| N | 19 | --- | 829.392 | 545.977 | --- | 14 |
| Q | 20 | --- | 872.078 | 507.963 | --- | 13 |
| Q | 21 | **-704.906** | **914.764** | 465.277 | --- | 12 |
| P | 22 | --- | 947.115 | 422.591 | --- | 11 |
| Q | 23 | **-53.861** | **989.801** | 390.240 | --- | 10 |
| G | 24 | **699.725** | **1008.808** | 347.554 | --- | 9 |
| V | 25 | --- | 1041.831 | 328.546 | --- | 8 |
| L | 26 | **-899.325** | **1079.526** | 295.524 | --- | 7 |
| Q | 27 | **-79.881** | **1122.212** | 257.829 | --- | 6 |
| L | 28 | **599.392** | **1159.907** | 215.143 | --- | 5 |
| L | 29 | **81.733** | **1197.601** | 177.448 | --- | 4 |
| R | 30 | **118.656** | **1249.635** | 139.753 | --- | 3 |
| L | 31 | **663.799** | **1287.330** | 87.720 | --- | 2 |
| E | 32 | --- | --- | 50.025 | --- | 1 |

**Soleil Index**

| | | |
|---|---|---|
| 965.7003 | b2-31 frag doubly charged/ y1-8 frag loss of water/ y1-8 frag loss of ammonia/ y2-31 frag loss of water | 21% |
| 1198.6040 | not explained | 33% |

| | | | | | |
|---|---|---|---|---|---|
| **Run:** | 79490 | **Scans:** | 25367 - 25367 | **m/z Range:** | 286.00864 to 1922.3885 |
| **Search:** | 79490 | **Precursor Mass:** | 998.51073 | | |
| **Mods:** | C=160.03 *+15.99 #+220.06 @+291.1 | **Charge:** | +4 | | |

**Sequence**

FT#KNNILT#QAAQAMLAQANQQPQGVLQLLRLE

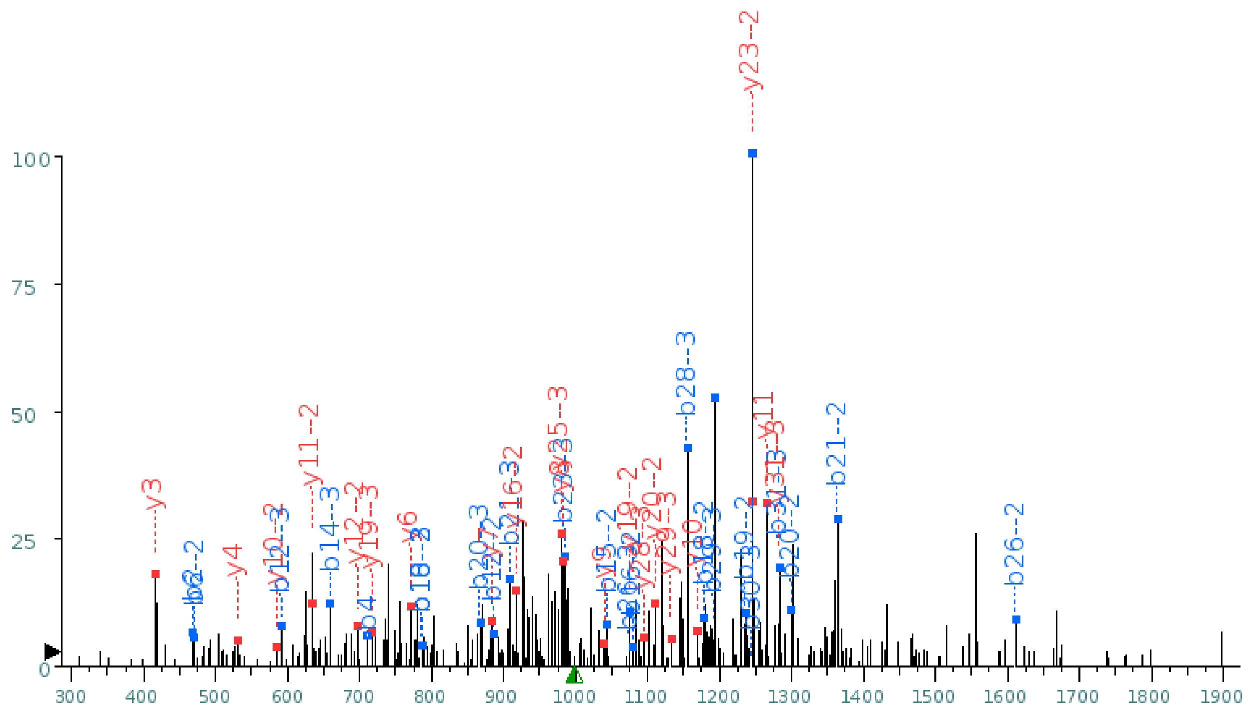

**Predicted Fragmentation Pattern**

**+1**

| Seq | # | b: Δ Error | b | y | y: Δ Error | +1 |
|---|---|---|---|---|---|---|
| F | 1 | --- | 148.076 | --- | --- | 32 |
| T# | 2 | -1588.749 | **469.182** | 3843.964 | --- | 31 |
| K | 3 | --- | 597.277 | 3522.858 | --- | 30 |
| N | 4 | 453.564 | **711.320** | 3394.763 | --- | 29 |
| N | 5 | --- | 825.362 | 3280.721 | --- | 28 |
| I | 6 | --- | 938.447 | 3166.678 | --- | 27 |
| L | 7 | --- | 1051.531 | 3053.594 | --- | 26 |
| T# | 8 | --- | 1372.637 | 2940.510 | --- | 25 |
| Q | 9 | --- | 1500.695 | 2619.404 | --- | 24 |
| A | 10 | --- | 1571.732 | 2491.345 | --- | 23 |
| A | 11 | --- | 1642.769 | 2420.308 | --- | 22 |
| Q | 12 | --- | 1770.828 | 2349.271 | --- | 21 |

| | | | | | | |
|---|---|---|---|---|---|---|
| A | 13 | --- | 1841.865 | 2221.212 | --- | 20 |
| M | 14 | --- | 1972.906 | 2150.175 | --- | 19 |
| L | 15 | --- | 2085.990 | 2019.135 | --- | 18 |
| A | 16 | --- | 2157.027 | 1906.050 | --- | 17 |
| Q | 17 | --- | 2285.085 | 1835.013 | --- | 16 |
| A | 18 | --- | 2356.122 | 1706.955 | --- | 15 |
| N | 19 | --- | 2470.165 | 1635.918 | --- | 14 |
| Q | 20 | --- | 2598.224 | 1521.875 | --- | 13 |
| Q | 21 | --- | 2726.283 | 1393.816 | --- | 12 |
| P | 22 | --- | 2823.335 | **1265.758** | **185.397** | 11 |
| Q | 23 | --- | 2951.394 | **1168.705** | **141.091** | 10 |
| G | 24 | --- | 3008.415 | **1040.646** | **-856.587** | 9 |
| V | 25 | --- | 3107.484 | **983.625** | **-891.391** | 8 |
| L | 26 | --- | 3220.568 | **884.556** | **81.434** | 7 |
| Q | 27 | --- | 3348.626 | **771.472** | **-895.785** | 6 |
| L | 28 | --- | 3461.710 | 643.414 | --- | 5 |
| L | 29 | --- | 3574.795 | **530.330** | **386.967** | 4 |
| R | 30 | --- | 3730.896 | **417.246** | **446.533** | 3 |
| L | 31 | --- | 3843.980 | 261.144 | --- | 2 |
| E | 32 | --- | --- | 148.060 | --- | 1 |

**+2**

| Seq | # | b: Δ Error | b | y | y: Δ Error | +1 |
|---|---|---|---|---|---|---|
| F | 1 | --- | 74.541 | --- | --- | 32 |
| T# | 2 | --- | 235.094 | 1922.486 | --- | 31 |
| K | 3 | --- | 299.142 | 1761.933 | --- | 30 |
| N | 4 | --- | 356.163 | 1697.885 | --- | 29 |
| N | 5 | --- | 413.185 | 1640.864 | --- | 28 |
| I | 6 | **1684.300** | **469.727** | 1583.842 | --- | 27 |
| L | 7 | --- | 526.269 | 1527.300 | --- | 26 |
| T# | 8 | --- | 686.822 | 1470.758 | --- | 25 |
| Q | 9 | --- | 750.851 | 1310.205 | --- | 24 |
| A | 10 | **-259.032** | **786.370** | **1246.176** | **-94.412** | 23 |
| A | 11 | --- | 821.888 | 1210.658 | --- | 22 |
| Q | 12 | **698.218** | **885.918** | 1175.139 | --- | 21 |
| A | 13 | --- | 921.436 | **1111.110** | **-232.059** | 20 |
| M | 14 | --- | 986.956 | **1075.591** | **-604.791** | 19 |
| L | 15 | **413.435** | **1043.498** | 1010.071 | --- | 18 |
| A | 16 | **-103.795** | **1079.017** | 953.529 | --- | 17 |
| Q | 17 | --- | 1143.046 | **918.010** | **-267.556** | 16 |
| A | 18 | **188.231** | **1178.565** | 853.981 | --- | 15 |
| N | 19 | **658.911** | **1235.586** | 818.462 | --- | 14 |
| Q | 20 | **-469.493** | **1299.616** | 761.441 | --- | 13 |
| Q | 21 | **251.032** | **1363.645** | **697.412** | **611.742** | 12 |
| P | 22 | --- | 1412.171 | **633.382** | **375.115** | 11 |
| Q | 23 | --- | 1476.201 | **584.856** | **1327.605** | 10 |
| G | 24 | --- | 1504.711 | 520.827 | --- | 9 |
| V | 25 | --- | 1554.246 | 492.316 | --- | 8 |
| L | 26 | **390.695** | **1610.788** | 442.782 | --- | 7 |
| Q | 27 | --- | 1674.817 | 386.240 | --- | 6 |
| L | 28 | --- | 1731.359 | 322.211 | --- | 5 |
| L | 29 | --- | 1787.901 | 265.668 | --- | 4 |
| R | 30 | --- | 1865.951 | 209.126 | --- | 3 |
| L | 31 | --- | 1922.493 | 131.076 | --- | 2 |
| E | 32 | --- | --- | 74.534 | --- | 1 |

**+3**

| Seq | # | b: Δ Error | b | y | y: Δ Error | +1 |
|---|---|---|---|---|---|---|

| | | | | | | |
|---|---|---|---|---|---|---|
| F | 1 | --- | 50.030 | --- | --- | 32 |
| T# | 2 | --- | 157.065 | **1281.993** | **760.164** | 31 |
| K | 3 | --- | 199.764 | 1174.958 | --- | 30 |
| N | 4 | --- | 237.778 | **1132.259** | **181.567** | 29 |
| N | 5 | --- | 275.792 | **1094.245** | **583.730** | 28 |
| I | 6 | --- | 313.487 | 1056.231 | --- | 27 |
| L | 7 | --- | 351.182 | 1018.536 | --- | 26 |
| T# | 8 | --- | 458.217 | **980.841** | **787.753** | 25 |
| Q | 9 | --- | 500.903 | 873.806 | --- | 24 |
| A | 10 | --- | 524.582 | 831.120 | --- | 23 |
| A | 11 | --- | 548.261 | 807.441 | --- | 22 |
| Q | 12 | **1208.815** | **590.948** | 783.762 | --- | 21 |
| A | 13 | --- | 614.627 | 741.076 | --- | 20 |
| M | 14 | **-32.926** | **658.307** | **717.397** | **169.046** | 19 |
| L | 15 | --- | 696.001 | 673.716 | --- | 18 |
| A | 16 | --- | 719.680 | 636.022 | --- | 17 |
| Q | 17 | --- | 762.367 | 612.343 | --- | 16 |
| A | 18 | **153.241** | **786.046** | 569.656 | --- | 15 |
| N | 19 | --- | 824.060 | 545.977 | --- | 14 |
| Q | 20 | **464.464** | **866.746** | 507.963 | --- | 13 |
| Q | 21 | **340.701** | **909.432** | 465.277 | --- | 12 |
| P | 22 | --- | 941.783 | 422.591 | --- | 11 |
| Q | 23 | **804.344** | **984.469** | 390.240 | --- | 10 |
| G | 24 | --- | 1003.477 | 347.554 | --- | 9 |
| V | 25 | --- | 1036.499 | 328.546 | --- | 8 |
| L | 26 | **694.847** | **1074.194** | 295.524 | --- | 7 |
| Q | 27 | --- | 1116.880 | 257.829 | --- | 6 |
| L | 28 | **750.270** | **1154.575** | 215.143 | --- | 5 |
| L | 29 | **694.885** | **1192.270** | 177.448 | --- | 4 |
| R | 30 | **512.387** | **1244.303** | 139.753 | --- | 3 |
| L | 31 | **756.201** | **1281.998** | 87.720 | --- | 2 |
| E | 32 | --- | --- | 50.025 | --- | 1 |

**Soleil Index**

| | | |
|---|---|---|
| **634.5511** | **not explained** | **22%** |
| **739.8367** | **not explained** | **20%** |
| **927.0315** | **not explained** | **28%** |
| **929.9825** | **not explained** | **18%** |
| **962.2007** | **b2-31 frag doubly charged/ y2-31 frag loss of water/ y3-25 frag loss of water** | **17%** |
| **963.0974** | **y3-25 frag loss of water/ y3-25 frag loss of ammonia** | **18%** |
| **990.0905** | **precursor loss of two waters** | **15%** |
| **1120.1083** | **not explained** | **25%** |
| **1147.2987** | **not explained** | **16%** |
| **1230.7513** | **not explained** | **23%** |
| **1266.9999** | **not explained** | **22%** |
| **1300.8843** | **not explained** | **24%** |
| **1360.3636** | **not explained** | **17%** |
| **1364.7518** | **not explained** | **21%** |
| **1555.5702** | **b1-10 frag loss of ammonia** | **26%** |

| | | | | | |
|---|---|---|---|---|---|
| **Run:** | 79490 | **Scans:** | 25653 - 25653 | **m/z Range:** | 274.14551 to 1799.1095 |
| **Search:** | 79490 | **Precursor Mass:** | 1002.51189 | | |
| **Mods:** | C=160.03 *+15.99 #+220.06 @+291.1 | **Charge:** | +4 | | |

**Sequence**

FT#KNNILT#QAAQAM*LAQANQQPQGVLQLLRLE

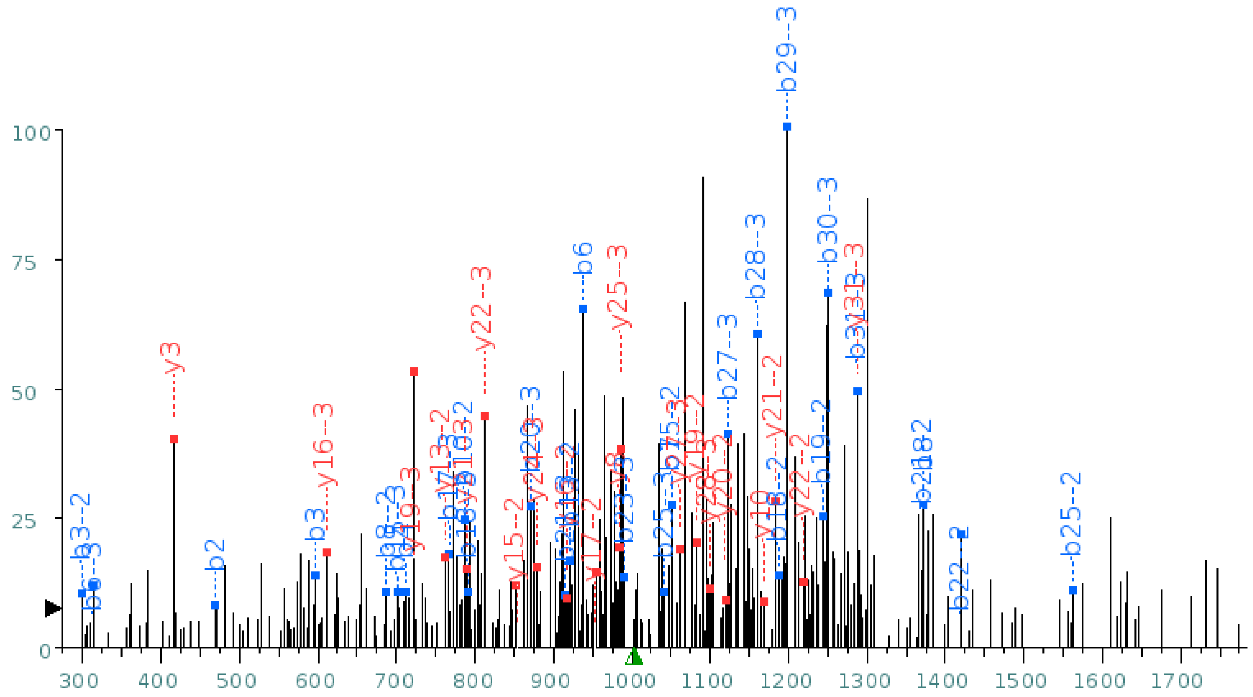

**Predicted Fragmentation Pattern**

| Seq | # | b: Δ Error | b | y | y: Δ Error | +1 |
|---|---|---|---|---|---|---|
| F | 1 | --- | 148.076 | --- | --- | 32 |
| T# | 2 | -1945.538 | **469.182** | 3859.959 | --- | 31 |
| K | 3 | -1539.323 | **597.277** | 3538.853 | --- | 30 |
| N | 4 | 435.903 | **711.320** | 3410.758 | --- | 29 |
| N | 5 | --- | 825.362 | 3296.715 | --- | 28 |
| I | 6 | -696.024 | **938.447** | 3182.673 | --- | 27 |
| L | 7 | -549.491 | **1051.531** | 3069.588 | --- | 26 |
| T# | 8 | 1.426 | **1372.637** | 2956.504 | --- | 25 |
| Q | 9 | --- | 1500.695 | 2635.398 | --- | 24 |
| A | 10 | --- | 1571.732 | 2507.340 | --- | 23 |
| A | 11 | --- | 1642.769 | 2436.303 | --- | 22 |

| | | | | | | |
|---|---|---|---|---|---|---|
| Q | 12 | --- | 1770.828 | 2365.266 | --- | 21 |
| A | 13 | --- | 1841.865 | 2237.207 | --- | 20 |
| M* | 14 | --- | 1988.900 | 2166.170 | --- | 19 |
| L | 15 | --- | 2101.985 | 2019.135 | --- | 18 |
| A | 16 | --- | 2173.022 | 1906.050 | --- | 17 |
| Q | 17 | --- | 2301.080 | 1835.013 | --- | 16 |
| A | 18 | --- | 2372.117 | 1706.955 | --- | 15 |
| N | 19 | --- | 2486.160 | 1635.918 | --- | 14 |
| Q | 20 | --- | 2614.219 | 1521.875 | --- | 13 |
| Q | 21 | --- | 2742.277 | 1393.816 | --- | 12 |
| P | 22 | --- | 2839.330 | 1265.758 | --- | 11 |
| Q | 23 | --- | 2967.389 | **1168.705** | 357.402 | 10 |
| G | 24 | --- | 3024.410 | 1040.646 | --- | 9 |
| V | 25 | --- | 3123.479 | **983.625** | 343.236 | 8 |
| L | 26 | --- | 3236.563 | 884.556 | --- | 7 |
| Q | 27 | --- | 3364.621 | 771.472 | --- | 6 |
| L | 28 | --- | 3477.705 | 643.414 | --- | 5 |
| L | 29 | --- | 3590.789 | 530.330 | --- | 4 |
| R | 30 | --- | 3746.891 | **417.246** | -29.267 | 3 |
| L | 31 | --- | 3859.975 | 261.144 | --- | 2 |
| E | 32 | --- | --- | 148.060 | --- | 1 |

+2

| Seq | # | b: Δ Error | b | y | y: Δ Error | +1 |
|---|---|---|---|---|---|---|
| F | 1 | --- | 74.541 | --- | --- | 32 |
| T# | 2 | --- | 235.094 | 1930.483 | --- | 31 |
| K | 3 | 11.881 | **299.142** | 1769.930 | --- | 30 |
| N | 4 | --- | 356.163 | 1705.883 | --- | 29 |
| N | 5 | --- | 413.185 | 1648.861 | --- | 28 |
| I | 6 | --- | 469.727 | 1591.840 | --- | 27 |
| L | 7 | --- | 526.269 | 1535.298 | --- | 26 |
| T# | 8 | 898.111 | **686.822** | 1478.756 | --- | 25 |
| Q | 9 | --- | 750.851 | 1318.203 | --- | 24 |
| A | 10 | 97.285 | **786.370** | 1254.174 | --- | 23 |
| A | 11 | --- | 821.888 | **1218.655** | -281.840 | 22 |
| Q | 12 | --- | 885.918 | **1183.136** | -56.323 | 21 |
| A | 13 | 644.594 | **921.436** | **1119.107** | 749.250 | 20 |
| M* | 14 | --- | 994.954 | **1083.589** | -3.933 | 19 |
| L | 15 | -516.473 | **1051.496** | 1010.071 | --- | 18 |
| A | 16 | --- | 1087.014 | **953.529** | 796.850 | 17 |
| Q | 17 | --- | 1151.044 | **918.010** | -471.421 | 16 |
| A | 18 | 53.854 | **1186.562** | **853.981** | -1016.302 | 15 |
| N | 19 | 540.145 | **1243.584** | 818.462 | --- | 14 |
| Q | 20 | --- | 1307.613 | **761.441** | 299.893 | 13 |
| Q | 21 | 725.750 | **1371.642** | 697.412 | --- | 12 |
| P | 22 | 573.907 | **1420.169** | 633.382 | --- | 11 |
| Q | 23 | --- | 1484.198 | 584.856 | --- | 10 |
| G | 24 | --- | 1512.709 | 520.827 | --- | 9 |
| V | 25 | 175.983 | **1562.243** | 492.316 | --- | 8 |
| L | 26 | --- | 1618.785 | 442.782 | --- | 7 |
| Q | 27 | --- | 1682.814 | 386.240 | --- | 6 |
| L | 28 | --- | 1739.356 | 322.211 | --- | 5 |
| L | 29 | --- | 1795.898 | 265.668 | --- | 4 |
| R | 30 | --- | 1873.949 | 209.126 | --- | 3 |
| L | 31 | --- | 1930.491 | 131.076 | --- | 2 |
| E | 32 | --- | --- | 74.534 | --- | 1 |

+3

| Seq | # | b: Δ Error | b | y | y: Δ Error | +1 |
|---|---|---|---|---|---|---|
| F | 1 | --- | 50.030 | --- | --- | 32 |
| T# | 2 | --- | 157.065 | 1287.325 | -141.347 | 31 |
| K | 3 | --- | 199.764 | 1180.289 | --- | 30 |
| N | 4 | --- | 237.778 | 1137.591 | --- | 29 |
| N | 5 | --- | 275.792 | 1099.577 | 205.322 | 28 |
| I | 6 | 2350.678 | 313.487 | 1061.562 | -412.059 | 27 |
| L | 7 | --- | 351.182 | 1023.868 | --- | 26 |
| T# | 8 | --- | 458.217 | 986.173 | 700.343 | 25 |
| Q | 9 | --- | 500.903 | 879.138 | -549.151 | 24 |
| A | 10 | --- | 524.582 | 836.451 | --- | 23 |
| A | 11 | --- | 548.261 | 812.772 | 856.282 | 22 |
| Q | 12 | --- | 590.948 | 789.093 | -443.885 | 21 |
| A | 13 | --- | 614.627 | 746.407 | --- | 20 |
| M* | 14 | --- | 663.638 | 722.728 | -649.198 | 19 |
| L | 15 | -957.908 | 701.333 | 673.716 | --- | 18 |
| A | 16 | --- | 725.012 | 636.022 | --- | 17 |
| Q | 17 | -1012.505 | 767.698 | 612.343 | -93.886 | 16 |
| A | 18 | -414.931 | 791.377 | 569.656 | --- | 15 |
| N | 19 | --- | 829.392 | 545.977 | --- | 14 |
| Q | 20 | -59.686 | 872.078 | 507.963 | --- | 13 |
| Q | 21 | 198.837 | 914.764 | 465.277 | --- | 12 |
| P | 22 | --- | 947.115 | 422.591 | --- | 11 |
| Q | 23 | 186.104 | 989.801 | 390.240 | --- | 10 |
| G | 24 | --- | 1008.808 | 347.554 | --- | 9 |
| V | 25 | -139.114 | 1041.831 | 328.546 | --- | 8 |
| L | 26 | --- | 1079.526 | 295.524 | --- | 7 |
| Q | 27 | 82.413 | 1122.212 | 257.829 | --- | 6 |
| L | 28 | 331.907 | 1159.907 | 215.143 | --- | 5 |
| L | 29 | 372.812 | 1197.601 | 177.448 | --- | 4 |
| R | 30 | 747.007 | 1249.635 | 139.753 | --- | 3 |
| L | 31 | -145.297 | 1287.330 | 87.720 | --- | 2 |
| E | 32 | --- | --- | 50.025 | --- | 1 |

**Soleil Index**

| | | |
|---|---|---|
| 383.4539 | b3-17 frag doubly charged | 15% |
| 481.0623 | not explained | 16% |
| 527.3635 | y3-14 frag loss of water | 16% |
| 577.2576 | not explained | 18% |
| 578.2937 | b1-3 frag loss of water | 16% |
| 588.3141 | not explained | 17% |
| 654.4200 | b2-20 frag doubly charged | 22% |
| 766.0508 | not explained | 16% |
| 773.0663 | b3-18 frag loss of water/ y3-21 frag loss of ammonia | 15% |
| 773.7050 | b3-18 frag loss of water/ b3-18 frag loss of ammonia | 34% |
| 777.7345 | not explained | 18% |
| 793.1969 | not explained | 24% |
| 803.4778 | b2-11 frag loss of water | 21% |
| 862.6870 | y3-24 frag loss of ammonia | 17% |

| | | |
|---|---|---|
| 867.3615 | b2-12 frag loss of water/ y1-7 frag loss of water/ y1-7 frag loss of ammonia | 47% |
| 875.6100 | not explained | 29% |
| 896.2570 | b3-21 frag loss of water | 20% |
| 901.6971 | y2-16 frag loss of ammonia | 19% |
| 910.1871 | not explained | 22% |
| 910.8811 | not explained | 20% |
| 912.0953 | not explained | 53% |
| 927.3839 | not explained | 46% |
| 931.6865 | not explained | 38% |
| 959.9536 | not explained | 25% |
| 964.8756 | b2-31 frag doubly charged/ y1-8 frag loss of water/ y2-31 frag loss of water | 49% |
| 967.6576 | y3-25 frag loss of water | 21% |
| 974.0327 | not explained | 34% |
| 976.9843 | b2-14 frag loss of water/ b2-14 frag loss of ammonia | 19% |
| 977.6702 | b2-14 frag loss of water/ b2-14 frag loss of ammonia | 30% |
| 987.6832 | not explained | 48% |
| 1033.6853 | b1-7 frag loss of water/ b1-7 frag loss of ammonia/ b2-15 frag loss of water/ b2-15 frag loss of ammonia | 39% |
| 1038.8363 | not explained | 19% |
| 1047.1327 | not explained | 16% |
| 1068.2797 | b2-16 frag loss of water | 67% |
| 1076.1470 | not explained | 26% |
| 1090.8103 | not explained | 91% |
| 1094.9960 | not explained | 29% |
| 1103.1680 | not explained | 24% |
| 1126.4785 | not explained | 27% |
| 1132.4061 | b2-17 frag loss of water | 15% |
| 1134.8894 | b2-17 frag loss of ammonia | 39% |
| 1143.8998 | not explained | 41% |
| 1147.1300 | not explained | 29% |
| 1148.6627 | not explained | 29% |
| 1149.4210 | not explained | 19% |
| 1154.6931 | not explained | 15% |
| 1193.9574 | not explained | 17% |
| 1195.8467 | not explained | 16% |
| 1198.7008 | not explained | 27% |
| 1209.3702 | not explained | 37% |
| 1212.7089 | not explained | 20% |
| 1221.1680 | not explained | 25% |
| 1229.3539 | not explained | 16% |

| | | |
|---|---|---|
| 1235.5726 | y2-23 frag loss of water | 25% |
| 1245.5648 | not explained | 17% |
| 1251.3167 | not explained | 17% |
| 1256.2367 | not explained | 19% |
| 1257.9237 | not explained | 17% |
| 1271.0295 | b3-31 frag loss of ammonia/ y3-31 frag loss of ammonia | 39% |
| 1275.4844 | not explained | 19% |
| 1290.5729 | b2-20 frag loss of water/ b2-20 frag loss of ammonia | 19% |
| 1301.0880 | y2-24 frag loss of water/ y2-24 frag loss of ammonia | 87% |
| 1309.3370 | not explained | 18% |
| 1366.0870 | not explained | 26% |
| 1369.1094 | not explained | 15% |
| 1378.6725 | not explained | 23% |
| 1383.6879 | not explained | 26% |
| 1611.2692 | not explained | 25% |
| 1732.1880 | not explained | 17% |
| 1746.8448 | not explained | 15% |

| | | | | | | |
|---|---|---|---|---|---|---|
| **Run:** | 79490 | **Scans:** | 27287 - 27287 | **m/z Range:** | 306.32849 | to 1922.0821! |
| **Search:** | 79490 | **Precursor Mass:** | 1002.51102 | | | |
| **Mods:** | C=160.03 *+15.99 #+220.06 @+291.1 | **Charge:** | +4 | | | |

**Sequence**

FT#KNNILT#QAAQAM*LAQANQQPQGVLQLLRLE

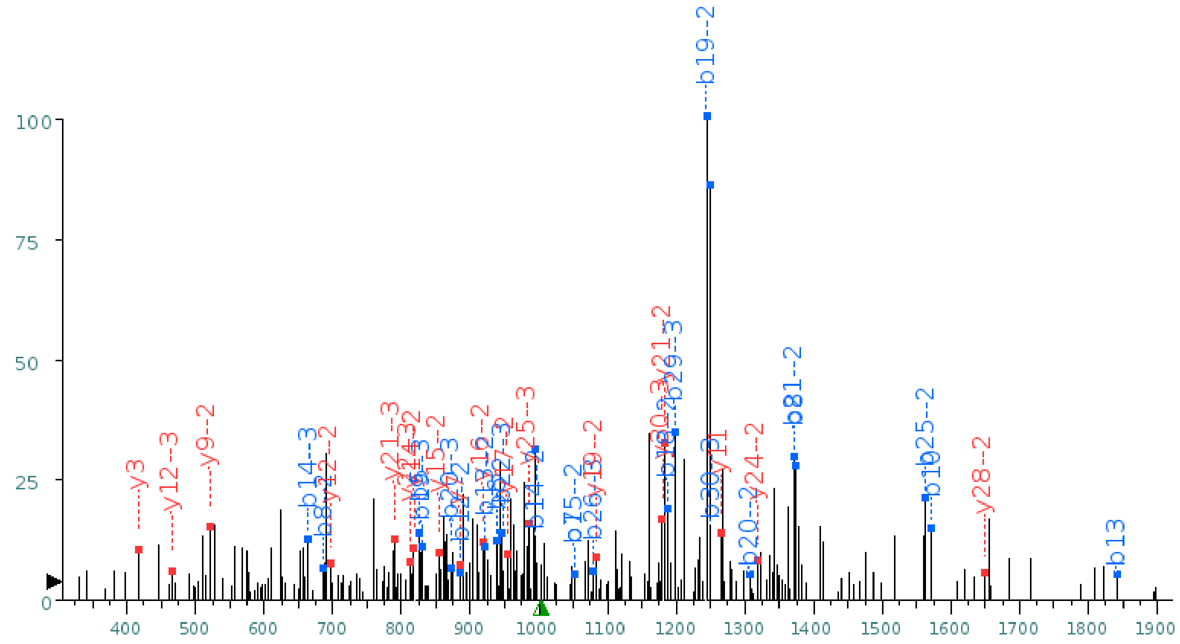

**Predicted Fragmentation Pattern**

| Seq | # | b: Δ Error | b | +1 y | y: Δ Error | +1 |
|---|---|---|---|---|---|---|
| F | 1 | --- | 148.076 | --- | --- | 32 |
| T# | 2 | --- | 469.182 | 3859.959 | --- | 31 |
| K | 3 | --- | 597.277 | 3538.853 | --- | 30 |
| N | 4 | --- | 711.320 | 3410.758 | --- | 29 |
| N | 5 | -799.514 | **825.362** | 3296.715 | --- | 28 |
| I | 6 | -231.349 | **938.447** | 3182.673 | --- | 27 |
| L | 7 | 706.038 | **1051.531** | 3069.588 | --- | 26 |
| T# | 8 | 104.753 | **1372.637** | 2956.504 | --- | 25 |
| Q | 9 | --- | 1500.695 | 2635.398 | --- | 24 |
| A | 10 | 123.484 | **1571.732** | 2507.340 | --- | 23 |
| A | 11 | --- | 1642.769 | 2436.303 | --- | 22 |
| Q | 12 | --- | 1770.828 | 2365.266 | --- | 21 |

| | | | | | | |
|---|---|---|---|---|---|---|
| A | 13 | -361.916 | **1841.865** | 2237.207 | --- | 20 |
| M* | 14 | --- | 1988.900 | 2166.170 | --- | 19 |
| L | 15 | --- | 2101.985 | 2019.135 | --- | 18 |
| A | 16 | --- | 2173.022 | 1906.050 | --- | 17 |
| Q | 17 | --- | 2301.080 | 1835.013 | --- | 16 |
| A | 18 | --- | 2372.117 | 1706.955 | --- | 15 |
| N | 19 | --- | 2486.160 | 1635.918 | --- | 14 |
| Q | 20 | --- | 2614.219 | 1521.875 | --- | 13 |
| Q | 21 | --- | 2742.277 | 1393.816 | --- | 12 |
| P | 22 | --- | 2839.330 | **1265.758** | 166.116 | 11 |
| Q | 23 | --- | 2967.389 | 1168.705 | --- | 10 |
| G | 24 | --- | 3024.410 | 1040.646 | --- | 9 |
| V | 25 | --- | 3123.479 | 983.625 | --- | 8 |
| L | 26 | --- | 3236.563 | **884.556** | 218.291 | 7 |
| Q | 27 | --- | 3364.621 | 771.472 | --- | 6 |
| L | 28 | --- | 3477.705 | 643.414 | --- | 5 |
| L | 29 | --- | 3590.789 | 530.330 | --- | 4 |
| R | 30 | --- | 3746.891 | **417.246** | -64.378 | 3 |
| L | 31 | --- | 3859.975 | 261.144 | --- | 2 |
| E | 32 | --- | --- | 148.060 | --- | 1 |

+2

| Seq | # | b: Δ Error | b | y | y: Δ Error | +1 |
|---|---|---|---|---|---|---|
| F | 1 | --- | 74.541 | --- | --- | 32 |
| T# | 2 | --- | 235.094 | 1930.483 | --- | 31 |
| K | 3 | --- | 299.142 | 1769.930 | --- | 30 |
| N | 4 | --- | 356.163 | 1705.883 | --- | 29 |
| N | 5 | --- | 413.185 | **1648.861** | 225.646 | 28 |
| I | 6 | --- | 469.727 | 1591.840 | --- | 27 |
| L | 7 | --- | 526.269 | 1535.298 | --- | 26 |
| T# | 8 | -1087.175 | **686.822** | 1478.756 | --- | 25 |
| Q | 9 | --- | 750.851 | **1318.203** | -293.902 | 24 |
| A | 10 | --- | 786.370 | 1254.174 | --- | 23 |
| A | 11 | --- | 821.888 | 1218.655 | --- | 22 |
| Q | 12 | -189.661 | **885.918** | **1183.136** | -581.510 | 21 |
| A | 13 | 512.401 | **921.436** | 1119.107 | --- | 20 |
| M* | 14 | -702.424 | **994.954** | **1083.589** | 338.645 | 19 |
| L | 15 | 739.014 | **1051.496** | 1010.071 | --- | 18 |
| A | 16 | --- | 1087.014 | **953.529** | 648.690 | 17 |
| Q | 17 | --- | 1151.044 | **918.010** | 852.880 | 16 |
| A | 18 | 258.413 | **1186.562** | **853.981** | 1135.013 | 15 |
| N | 19 | 643.484 | **1243.584** | **818.462** | -938.354 | 14 |
| Q | 20 | 555.494 | **1307.613** | 761.441 | --- | 13 |
| Q | 21 | -211.427 | **1371.642** | **697.412** | 112.209 | 12 |
| P | 22 | --- | 1420.169 | 633.382 | --- | 11 |
| Q | 23 | --- | 1484.198 | 584.856 | --- | 10 |
| G | 24 | --- | 1512.709 | **520.827** | 1369.030 | 9 |
| V | 25 | 288.293 | **1562.243** | 492.316 | --- | 8 |
| L | 26 | --- | 1618.785 | 442.782 | --- | 7 |
| Q | 27 | --- | 1682.814 | 386.240 | --- | 6 |
| L | 28 | --- | 1739.356 | 322.211 | --- | 5 |
| L | 29 | --- | 1795.898 | 265.668 | --- | 4 |
| R | 30 | --- | 1873.949 | 209.126 | --- | 3 |
| L | 31 | --- | 1930.491 | 131.076 | --- | 2 |
| E | 32 | --- | --- | 74.534 | --- | 1 |

+3

| Seq | # | b: Δ Error | b | y | y: Δ Error | +1 |
|---|---|---|---|---|---|---|

| | | | | | | |
|---|---|---|---|---|---|---|
| F | 1 | --- | 50.030 | --- | --- | 32 |
| T# | 2 | --- | 157.065 | 1287.325 | --- | 31 |
| K | 3 | --- | 199.764 | **1180.289** | -735.783 | 30 |
| N | 4 | --- | 237.778 | 1137.591 | --- | 29 |
| N | 5 | --- | 275.792 | 1099.577 | --- | 28 |
| I | 6 | --- | 313.487 | 1061.562 | --- | 27 |
| L | 7 | --- | 351.182 | 1023.868 | --- | 26 |
| T# | 8 | --- | 458.217 | **986.173** | -237.612 | 25 |
| Q | 9 | --- | 500.903 | 879.138 | --- | 24 |
| A | 10 | --- | 524.582 | 836.451 | --- | 23 |
| A | 11 | --- | 548.261 | **812.772** | 138.488 | 22 |
| Q | 12 | --- | 590.948 | **789.093** | 678.178 | 21 |
| A | 13 | --- | 614.627 | 746.407 | --- | 20 |
| M* | 14 | 50.335 | **663.638** | 722.728 | --- | 19 |
| L | 15 | --- | 701.333 | 673.716 | --- | 18 |
| A | 16 | --- | 725.012 | 636.022 | --- | 17 |
| Q | 17 | --- | 767.698 | 612.343 | --- | 16 |
| A | 18 | --- | 791.377 | 569.656 | --- | 15 |
| N | 19 | 929.598 | **829.392** | 545.977 | --- | 14 |
| Q | 20 | -599.091 | **872.078** | 507.963 | --- | 13 |
| Q | 21 | --- | 914.764 | **465.277** | 567.311 | 12 |
| P | 22 | -946.718 | **947.115** | 422.591 | --- | 11 |
| Q | 23 | --- | 989.801 | 390.240 | --- | 10 |
| G | 24 | --- | 1008.808 | 347.554 | --- | 9 |
| V | 25 | --- | 1041.831 | 328.546 | --- | 8 |
| L | 26 | -393.667 | **1079.526** | 295.524 | --- | 7 |
| Q | 27 | --- | 1122.212 | 257.829 | --- | 6 |
| L | 28 | --- | 1159.907 | 215.143 | --- | 5 |
| L | 29 | 468.239 | **1197.601** | 177.448 | --- | 4 |
| R | 30 | 521.251 | **1249.635** | 139.753 | --- | 3 |
| L | 31 | --- | 1287.330 | 87.720 | --- | 2 |
| E | 32 | --- | --- | 50.025 | --- | 1 |

**Soleil Index**

| | | |
|---|---|---|
| 528.4276 | y3-14 frag loss of water/ y3-14 frag loss of ammonia | 16% |
| 624.7910 | b3-30 frag doubly charged/ y1-5 frag loss of water | 19% |
| 689.2000 | not explained | 30% |
| 758.0835 | not explained | 21% |
| 860.8951 | y3-24 frag loss of water | 17% |
| 890.6830 | not explained | 22% |
| 903.5406 | b2-13 frag loss of water/ b2-13 frag loss of ammonia | 17% |
| 910.7454 | not explained | 16% |
| 943.6384 | not explained | 15% |
| 944.2919 | not explained | 28% |
| 959.7303 | not explained | 21% |
| 962.1755 | not explained | 16% |
| 977.7501 | b2-14 frag loss of water/ b2-14 frag loss of ammonia | 24% |
| 991.8771 | b3-24 frag loss of ammonia/ y2-18 frag loss of water | 15% |

| | | |
|---|---|---|
| 1161.0482 | not explained | 34% |
| 1211.8145 | not explained | 29% |
| 1266.8168 | not explained | 27% |
| 1342.0632 | not explained | 23% |
| 1361.9246 | not explained | 19% |
| 1377.8934 | not explained | 15% |
| 1408.8541 | not explained | 15% |
| 1563.3754 | not explained | 15% |
| 1656.0322 | not explained | 17% |

**MS/MS data of trypsin digested peptides with site score A and site score B above 19 from *Geobacillus kaustophilus Gk*F1 flagellin protein co-expressed with *Geobacillus kaustophilus* Maf in EV136 cells. Results of dataset 2 obtained in April 2019.**

| | | | | | |
|---|---|---|---|---|---|
| **Run:** | 77659 | **Scans:** | 6587 - 6587 | **m/z Range:** | 105.10699 to 904.65009 |
| **Search:** | 77659 | **Precursor Mass:** | 462.23781 | | |
| **Mods:** | C=160.03 *+15.99 #+220.06 @+291.1 | **Charge:** | +2 | | |

**Sequence**

LSS@GLR

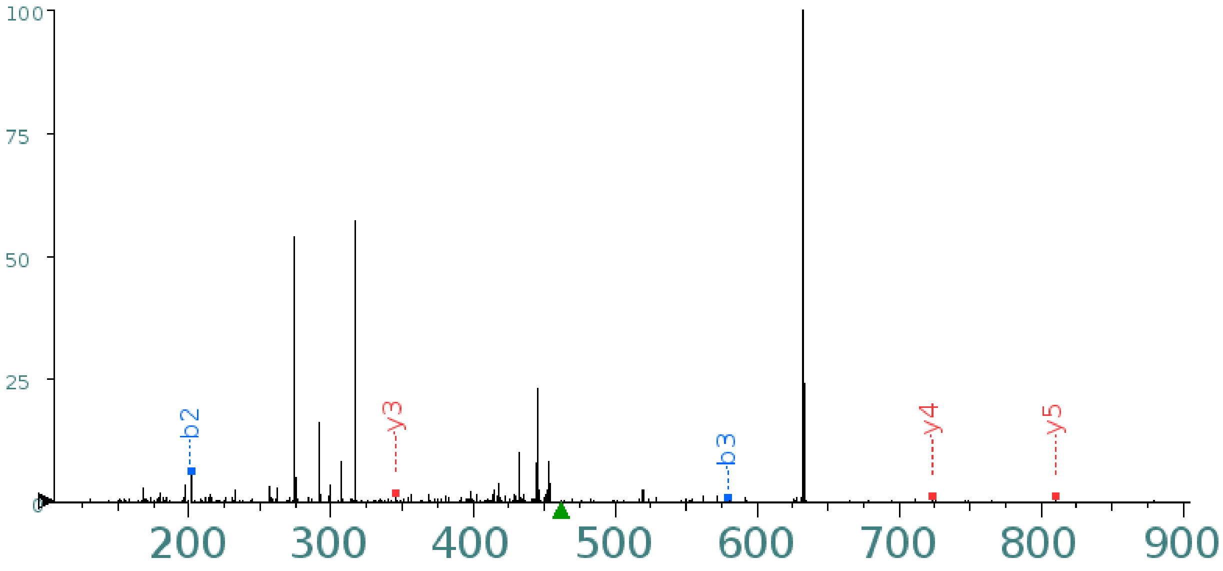

**Predicted Fragmentation Pattern**

| Seq | # | b: Δ Error | b | y | y: Δ Error | +1 |
|---|---|---|---|---|---|---|
| L | 1 | --- | 114.091 | --- | --- | 6 |
| S | 2 | 609.676 | 201.123 | 810.384 | 259.943 | 5 |
| S@ | 3 | -329.955 | 579.251 | 723.352 | 410.500 | 4 |
| G | 4 | --- | 636.272 | 345.224 | 351.196 | 3 |
| L | 5 | --- | 749.356 | 288.203 | --- | 2 |
| R | 6 | --- | --- | 175.119 | --- | 1 |

**Soleil Index**

| | | |
|---|---|---|
| 274.1237 | not explained | 54% |
| 292.1703 | not explained | 16% |
| 316.8489 | not explained | 57% |
| 445.1733 | precursor loss of two waters | 23% |
| 632.4200 | not explained | 100% |
| 633.4184 | not explained | 24% |

| | | | | | |
|---|---|---|---|---|---|
| Run: | 77659 | Scans: | 6672 - 6672 | m/z Range: | 122.29648 to 875.2807 |
| Search: | 77659 | Precursor Mass: | 462.23785 | | |
| Mods: | C=160.03 *+15.99 #+220.06 @+291.1 | Charge: | +2 | | |

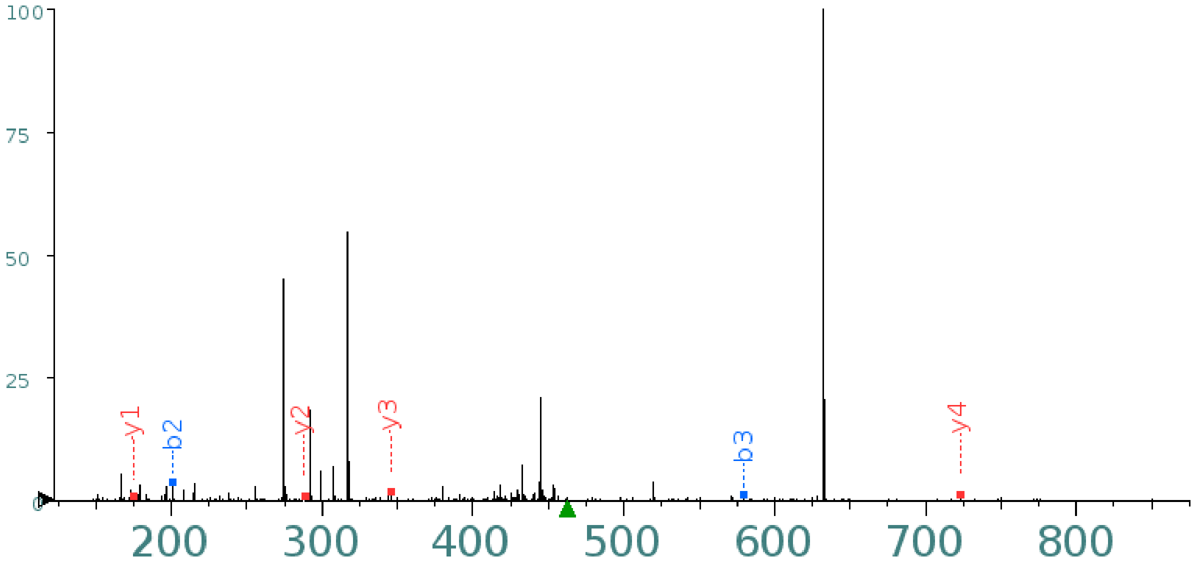

**Sequence**

LSS@GLR

**Predicted Fragmentation Pattern**

| Seq | # | b: Δ Error | b | y | y: Δ Error | +1 |
|---|---|---|---|---|---|---|
| L | 1 | --- | 114.091 | --- | --- | 6 |
| S | 2 | -162.009 | 201.123 | 810.384 | --- | 5 |
| S@ | 3 | 153.248 | 579.251 | 723.352 | 175.223 | 4 |
| G | 4 | --- | 636.272 | 345.224 | 25.567 | 3 |
| L | 5 | --- | 749.356 | 288.203 | 368.841 | 2 |
| R | 6 | --- | --- | 175.119 | 1148.613 | 1 |

**Soleil Index**

| | | |
|---|---|---|
| 274.0883 | not explained | 45% |
| 292.1053 | not explained | 18% |
| 316.6790 | not explained | 54% |
| 445.0871 | precursor loss of two waters | 21% |
| 632.3710 | not explained | 100% |
| 633.3690 | not explained | 21% |

| | | | |
|---|---|---|---|
| **Run:** | 77659 | **Scans:** | 7135 - 7135 |
| **Search:** | 77659 | **Precursor Mass:** | 426.71919 |
| **Mods:** | C=160.03 *+15.99 #+220.06 @+291.1 | **Charge:** | +2 |

**m/z Range:** 100.13631 **to** 853.43837

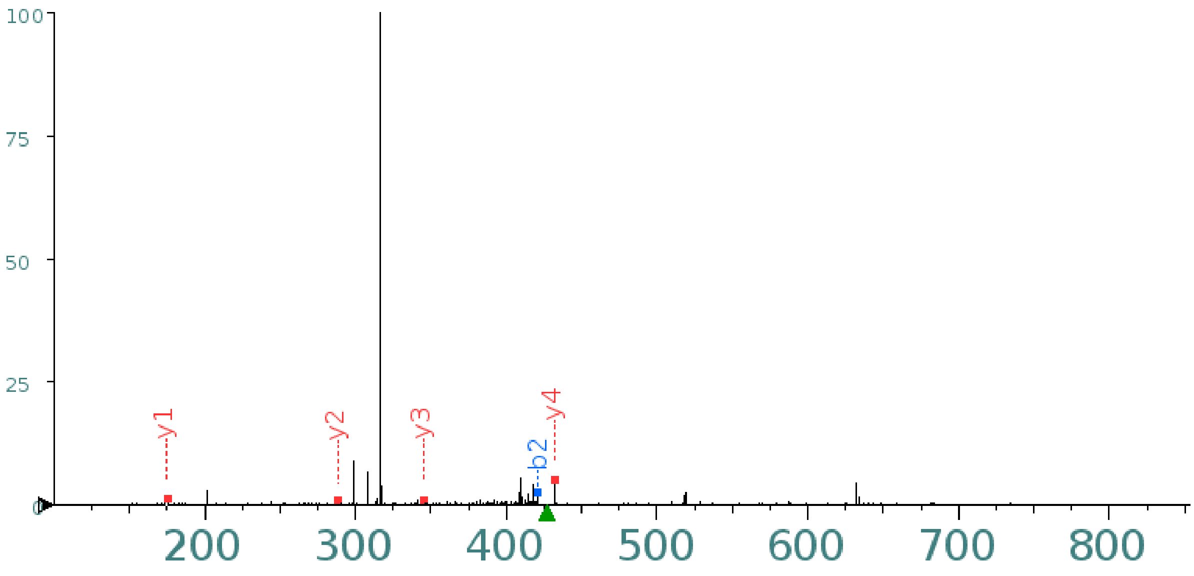

**Sequence**

LS#SGLR

**Predicted Fragmentation Pattern**

| Seq | # | b: Δ Error | b | y | y: Δ Error | +1 |
|---|---|---|---|---|---|---|
| L | 1 | --- | 114.091 | --- | --- | 6 |
| S# | 2 | -493.234 | **421.182** | 739.347 | --- | 5 |
| S | 3 | --- | 508.214 | **432.257** | -2.985 | 4 |
| G | 4 | --- | 565.235 | **345.224** | 1188.460 | 3 |
| L | 5 | --- | 678.319 | **288.203** | 699.179 | 2 |
| R | 6 | --- | --- | **175.119** | 270.332 | 1 |

**Soleil Index**

| | | |
|---|---|---|
| 316.7222 | not explained | 100% |

| | | | | | |
|---|---|---|---|---|---|
| **Run:** | 77659 | **Scans:** | 7454 - 7454 | **m/z Range:** | 96.24421 to 924.47595 |
| **Search:** | 77659 | **Precursor Mass:** | 462.23797 | | |
| **Mods:** | C=160.03 *+15.99 #+220.06 @+291.1 | **Charge:** | +2 | | |

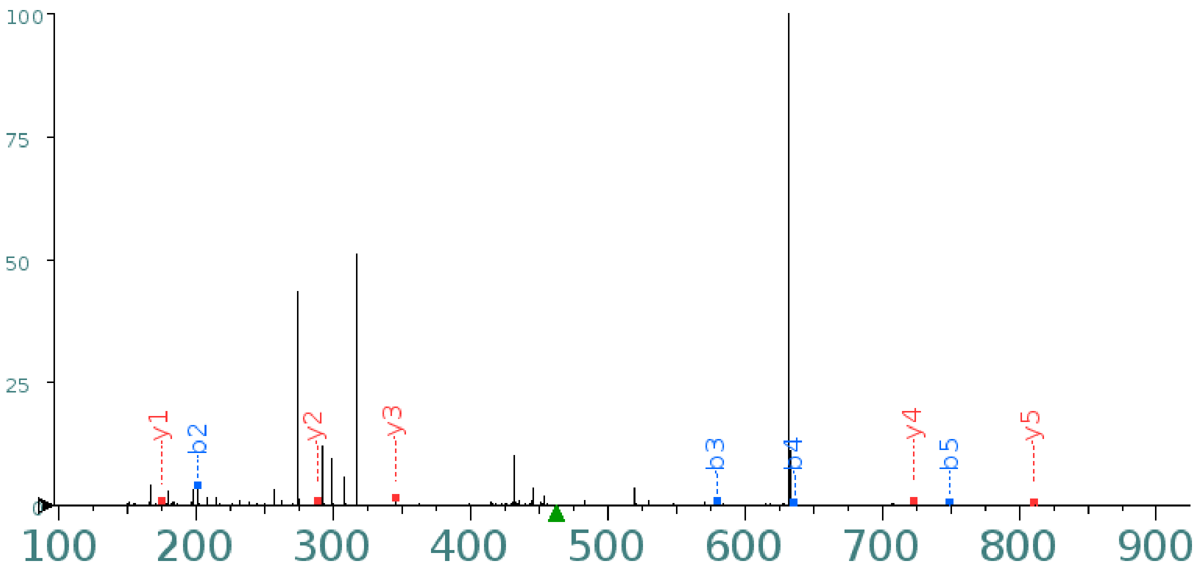

**Sequence**

LSS@GLR

**Predicted Fragmentation Pattern**

| Seq | # | b: Δ Error | b | y | y: Δ Error | +1 |
|---|---|---|---|---|---|---|
| L | 1 | --- | 114.091 | --- | --- | 6 |
| S | 2 | -27.394 | 201.123 | 810.384 | 595.266 | 5 |
| S@ | 3 | 694.071 | 579.251 | 723.352 | 92.133 | 4 |
| G | 4 | -922.129 | 636.272 | 345.224 | 50.494 | 3 |
| L | 5 | 84.917 | 749.356 | 288.203 | 629.703 | 2 |
| R | 6 | --- | --- | 175.119 | 190.728 | 1 |

**Soleil Index**

| | | |
|---|---|---|
| 274.1389 | not explained | 44% |
| 316.7420 | not explained | 51% |
| 632.4542 | not explained | 100% |

| | | | |
|---|---|---|---|
| **Run:** | 77659 | **Scans:** | 7526 - 7526 |
| **Search:** | 77659 | **Precursor Mass:** | 462.2378 |
| **Mods:** | C=160.03 *+15.99 #+220.06 @+291.1 | **Charge:** | +2 |

**m/z Range:** 96.12129 **to** 888.48517

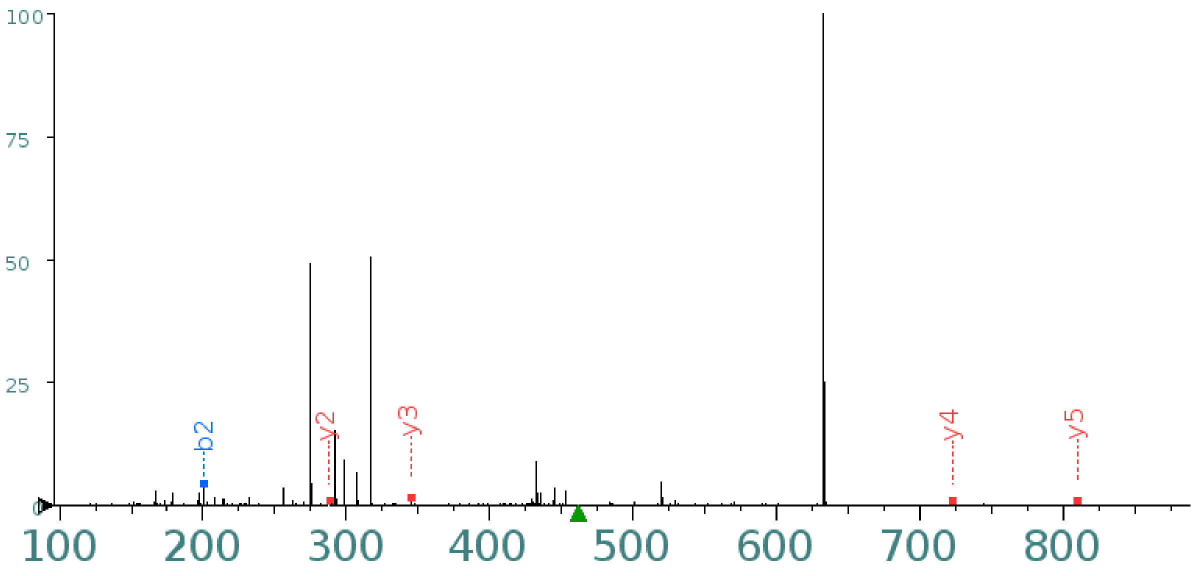

**Sequence**

LSS@GLR

**Predicted Fragmentation Pattern**

| Seq | # | b: Δ Error | b | y | y: Δ Error | +1 |
|---|---|---|---|---|---|---|
| L | 1 | --- | 114.091 | --- | --- | 6 |
| S | 2 | -46.666 | **201.123** | **810.384** | 221.400 | 5 |
| S@ | 3 | --- | 579.251 | **723.352** | 207.274 | 4 |
| G | 4 | --- | 636.272 | **345.224** | 668.402 | 3 |
| L | 5 | --- | 749.356 | **288.203** | 907.129 | 2 |
| R | 6 | --- | --- | 175.119 | --- | 1 |

**Soleil Index**

| | | |
|---|---|---|
| 274.1259 | not explained | 49% |
| 292.0948 | not explained | 15% |
| 316.7641 | not explained | 50% |
| 632.3833 | not explained | 100% |
| 633.4008 | not explained | 25% |

| | | | |
|---|---|---|---|
| Run: | 77659 | Scans: | 7687 - 7687 |
| Search: | 77659 | Precursor Mass: | 462.23778 |
| Mods: | C=160.03 *+15.99 #+220.06 @+291.1 | Charge: | +2 |

m/z Range: 107.19115 to 915.58344

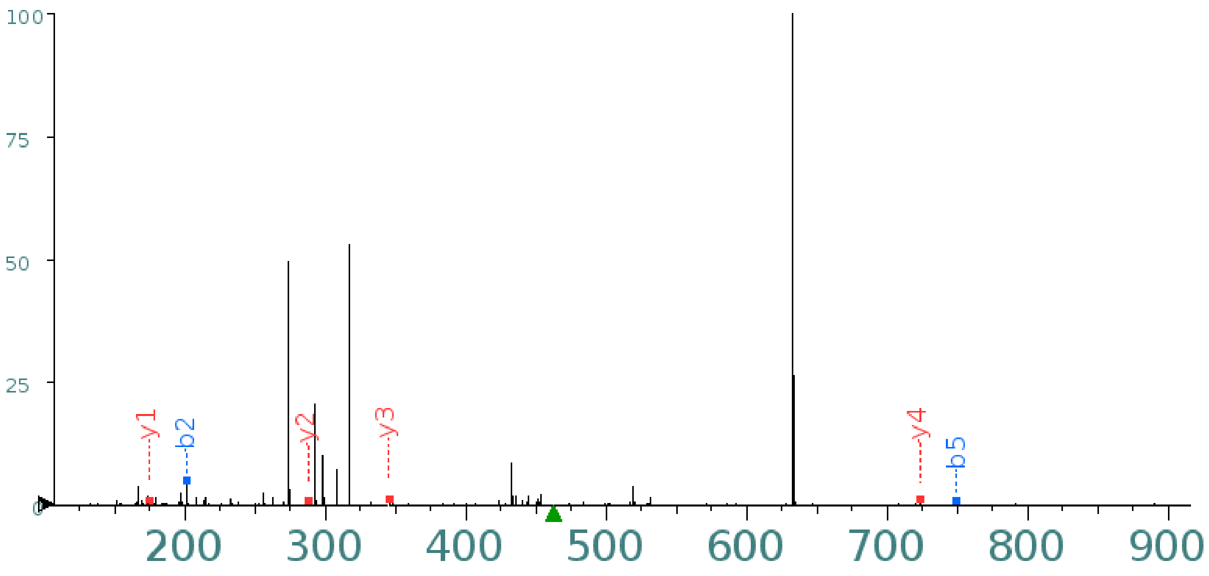

**Sequence**

LSS@GLR

**Predicted Fragmentation Pattern**

| Seq | # | b: Δ Error | b | y | y: Δ Error | +1 |
|---|---|---|---|---|---|---|
| L | 1 | --- | 114.091 | --- | --- | 6 |
| S | 2 | 336.658 | 201.123 | 810.384 | --- | 5 |
| S@ | 3 | --- | 579.251 | 723.352 | 154.557 | 4 |
| G | 4 | --- | 636.272 | 345.224 | 875.731 | 3 |
| L | 5 | 233.597 | 749.356 | 288.203 | 388.944 | 2 |
| R | 6 | --- | --- | 175.119 | 1202.248 | 1 |

**Soleil Index**

| | | |
|---|---|---|
| 274.1238 | not explained | 50% |
| 292.2257 | not explained | 21% |
| 316.6857 | not explained | 53% |

| | | |
|---|---|---|
| **632.3976** | **not explained** | **100%** |
| **633.5460** | **not explained** | **26%** |

| | | | | | |
|---|---|---|---|---|---|
| **Run:** | 77659 | **Scans:** | 7949 - 7949 | **m/z Range:** | 108.1953 to 890.72064 |
| **Search:** | 77659 | **Precursor Mass:** | 462.23774 | | |
| **Mods:** | C=160.03 *+15.99 #+220.06 @+291.1 | **Charge:** | +2 | | |

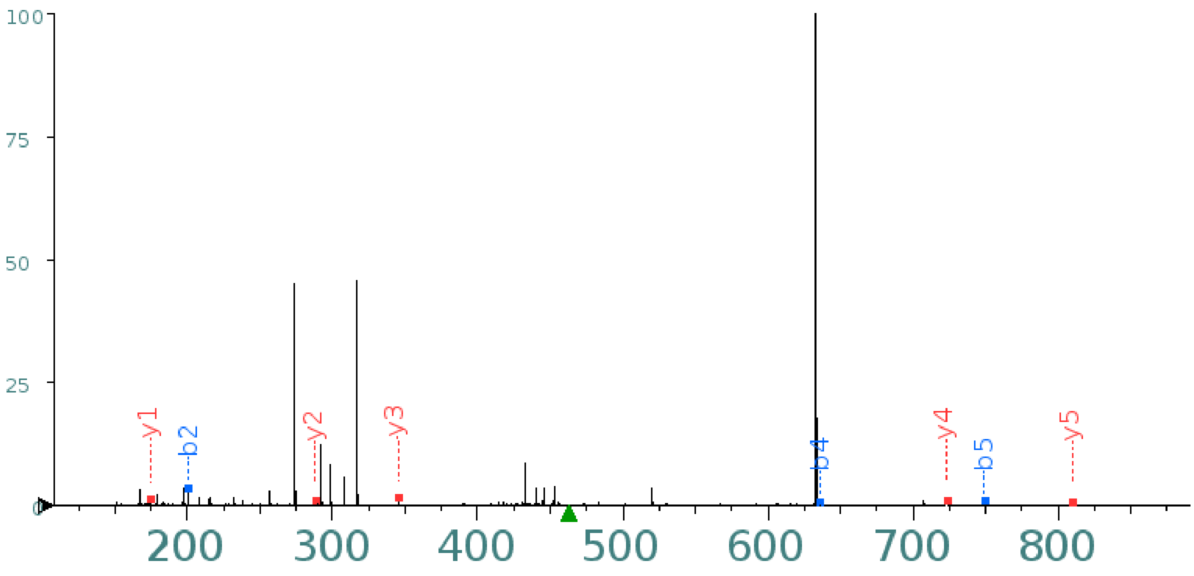

**Sequence**

LSS@GLR

**Predicted Fragmentation Pattern**

| Seq | # | b: Δ Error | b | y | y: Δ Error | +1 |
|---|---|---|---|---|---|---|
| L | 1 | --- | 114.091 | --- | --- | 6 |
| S | 2 | 166.193 | 201.123 | 810.384 | 149.349 | 5 |
| S@ | 3 | --- | 579.251 | 723.352 | 323.908 | 4 |
| G | 4 | 356.139 | 636.272 | 345.224 | 764.972 | 3 |
| L | 5 | 392.488 | 749.356 | 288.203 | 564.553 | 2 |
| R | 6 | --- | --- | 175.119 | 698.789 | 1 |

**Soleil Index**

| | | |
|---|---|---|
| 274.1165 | not explained | 45% |
| 316.7356 | not explained | 46% |
| 632.4685 | not explained | 100% |
| 633.4319 | not explained | 18% |

| | | | |
|---|---|---|---|
| **Run:** | 77659 | **Scans:** | 8038 - 8038 |
| **Search:** | 77659 | **Precursor Mass:** | 462.23772 |
| **Mods:** | C=160.03 *+15.99 #+220.06 @+291.1 | **Charge:** | +2 |

**m/z Range:** 107.3053 **to** 847.60162

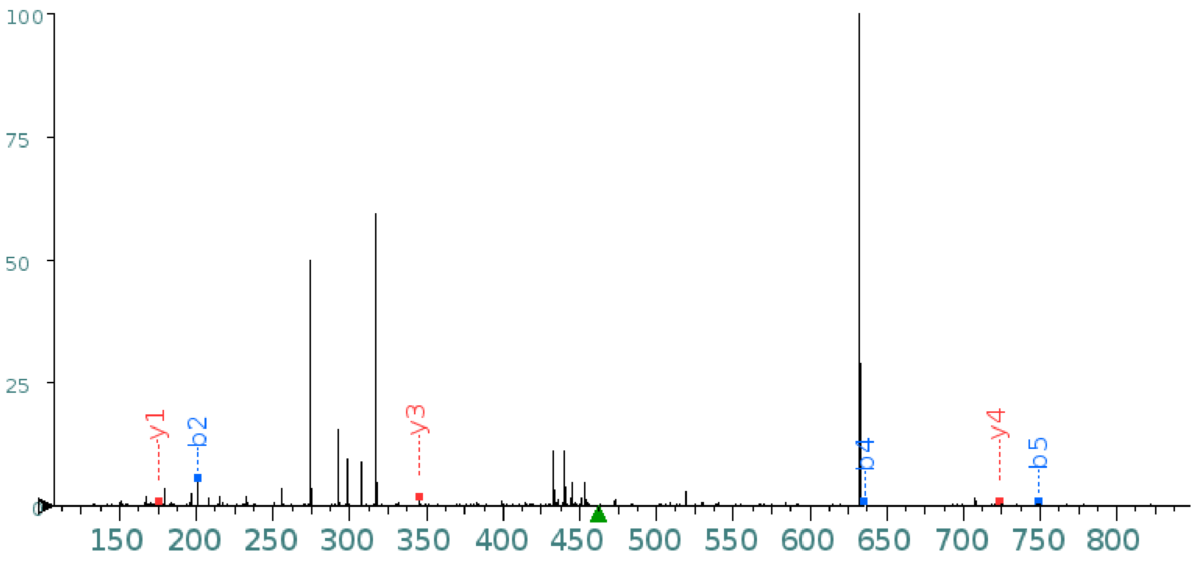

**Sequence**

LSS@GLR

**Predicted Fragmentation Pattern**

| Seq | # | b: Δ Error | b | y | y: Δ Error | +1 |
|---|---|---|---|---|---|---|
| L | 1 | --- | 114.091 | --- | --- | 6 |
| S | 2 | 827.104 | **201.123** | 810.384 | --- | 5 |
| S@ | 3 | --- | 579.251 | **723.352** | 276.768 | 4 |
| G | 4 | -1257.064 | **636.272** | **345.224** | 463.371 | 3 |
| L | 5 | -925.098 | **749.356** | 288.203 | --- | 2 |
| R | 6 | --- | --- | **175.119** | 456.227 | 1 |

**Soleil Index**

| | | |
|---|---|---|
| 274.1456 | not explained | 50% |
| 292.1471 | not explained | 15% |
| 316.7594 | not explained | 59% |
| 632.4393 | not explained | 100% |
| 633.4582 | not explained | 29% |

**MS/MS data of trypsin digested peptides with site score A and site score B above 19 from *Geobacillus kaustophilus Gk*F1 flagellin protein co-expressed with *Geobacillus kaustophilus* Maf in EV136 cells. Results of dataset 3 obtained in July 2019.**

| Run: | 79493 | Scans: | 6093 - 6093 | m/z Range: | 114.965 | to | 890.50812 |
|---|---|---|---|---|---|---|---|
| Search: | 79493 | Precursor Mass: | 462.23802 | | | | |
| Mods: | C=160.03 *+15.99 #+220.06 @+291.1 | Charge: | +2 | | | | |

**Sequence**

LSS@GLR

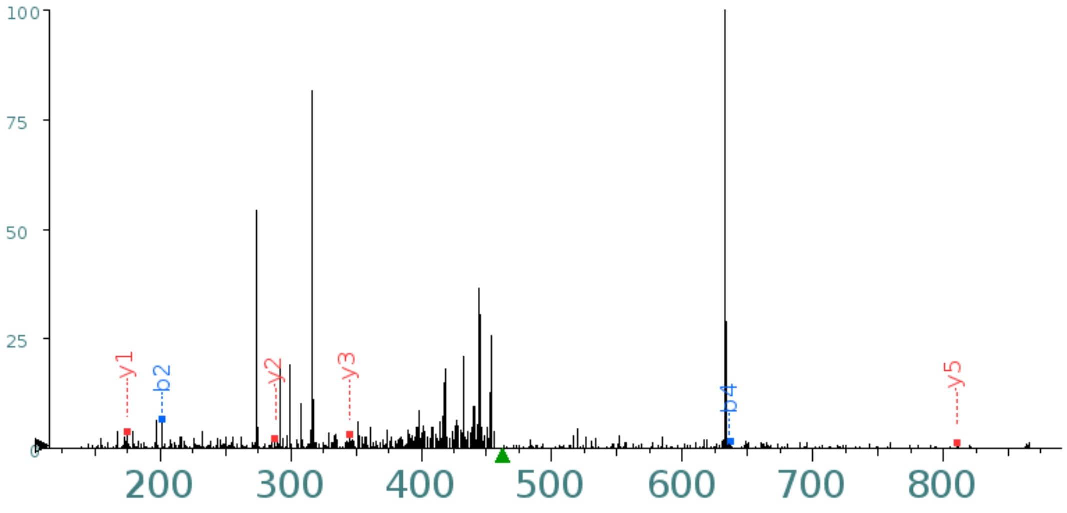

**Predicted Fragmentation Pattern**

| Seq | # | b: Δ Error | b | y | y: Δ Error | +1 |
|---|---|---|---|---|---|---|
| L | 1 | --- | 114.091 | --- | --- | 6 |
| S | 2 | 807.259 | 201.123 | 810.384 | 181.875 | 5 |
| S@ | 3 | --- | 579.251 | 723.352 | --- | 4 |
| G | 4 | 971.167 | 636.272 | 345.224 | 971.026 | 3 |
| L | 5 | --- | 749.356 | 288.203 | -3336.699 | 2 |
| R | 6 | --- | --- | 175.119 | -148.208 | 1 |

**Soleil Index**

| | | |
|---|---|---|
| 274.0312 | not explained | 54% |
| 292.1749 | not explained | 18% |
| 298.8715 | not explained | 19% |

| | | |
|---|---|---|
| 316.6473 | not explained | 82% |
| 417.4123 | not explained | 15% |
| 418.1690 | not explained | 18% |
| 432.2677 | not explained | 21% |
| 444.3224 | precursor loss of two waters | 37% |
| 445.4347 | not explained | 30% |
| 453.3940 | precursor loss of water | 26% |
| 632.3935 | not explained | 100% |
| 633.5339 | not explained | 29% |

| | | | | | |
|---|---|---|---|---|---|
| **Run:** | 79493 | **Scans:** | 6467 - 6467 | **m/z Range:** | 107.935 to 889.43372 |
| **Search:** | 79493 | **Precursor Mass:** | 462.23841 | | |
| **Mods:** | C=160.03 *+15.99 #+220.06 @+291.1 | **Charge:** | +2 | | |

**Sequence**

LSS@GLR

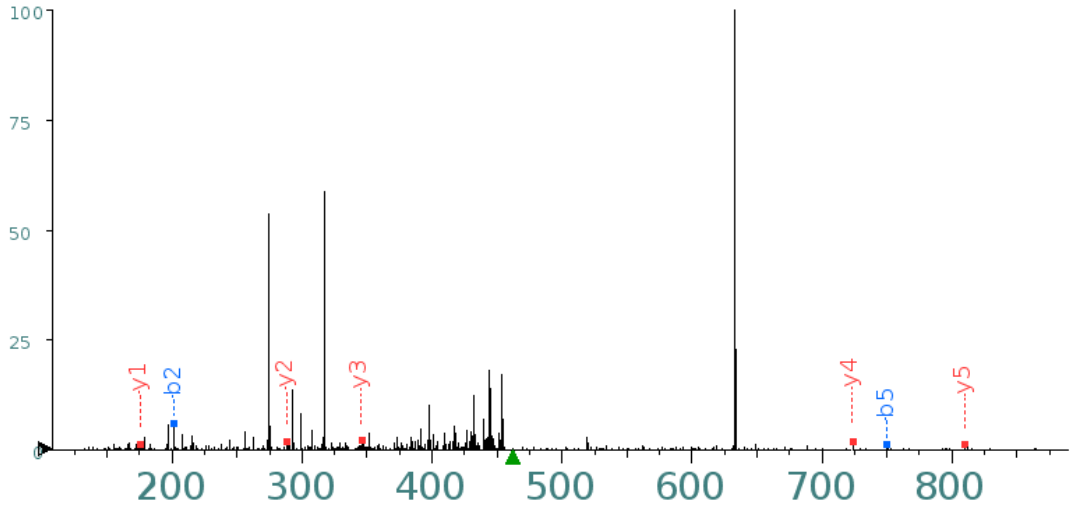

**Predicted Fragmentation Pattern**

| Seq | # | b: Δ Error | b | y | y: Δ Error | +1 |
|---|---|---|---|---|---|---|
| L | 1 | --- | 114.091 | --- | --- | 6 |
| S | 2 | 725.144 | 201.123 | 810.384 | -1059.297 | 5 |
| S@ | 3 | --- | 579.251 | 723.352 | 352.155 | 4 |
| G | 4 | --- | 636.272 | 345.224 | 2769.391 | 3 |
| L | 5 | 334.213 | 749.356 | 288.203 | 268.204 | 2 |
| R | 6 | --- | --- | 175.119 | 968.802 | 1 |

**Soleil Index**

| | | |
|---|---|---|
| 274.0946 | not explained | 54% |
| 316.8179 | not explained | 59% |
| 444.2976 | precursor loss of two waters | 18% |
| 453.0671 | precursor loss of water | 17% |
| 632.5024 | not explained | 100% |
| 633.4593 | not explained | 23% |

**MS/MS data of trypsin digested peptides with site score A and site score B above 19 from *Clostridium botulinum Cb*Fla flagellin protein co-expressed with *Clostridium botulinum* Maf in EV136 cells. Results of dataset 1 (MS/MS results obtained in January 2019).**

| | | | | | |
|---|---|---|---|---|---|
| **Run:** | 76752 | **Scans:** | 2546 - 2546 | **m/z Range:** | 189.19795 to 1178.9063 |
| **Search:** | 76752 | **Precursor Mass:** | 714.85085 | | |
| **Mods:** | C=160.03 *+15.99 #+220.06 @+291.1 | **Charge:** | +2 | | |

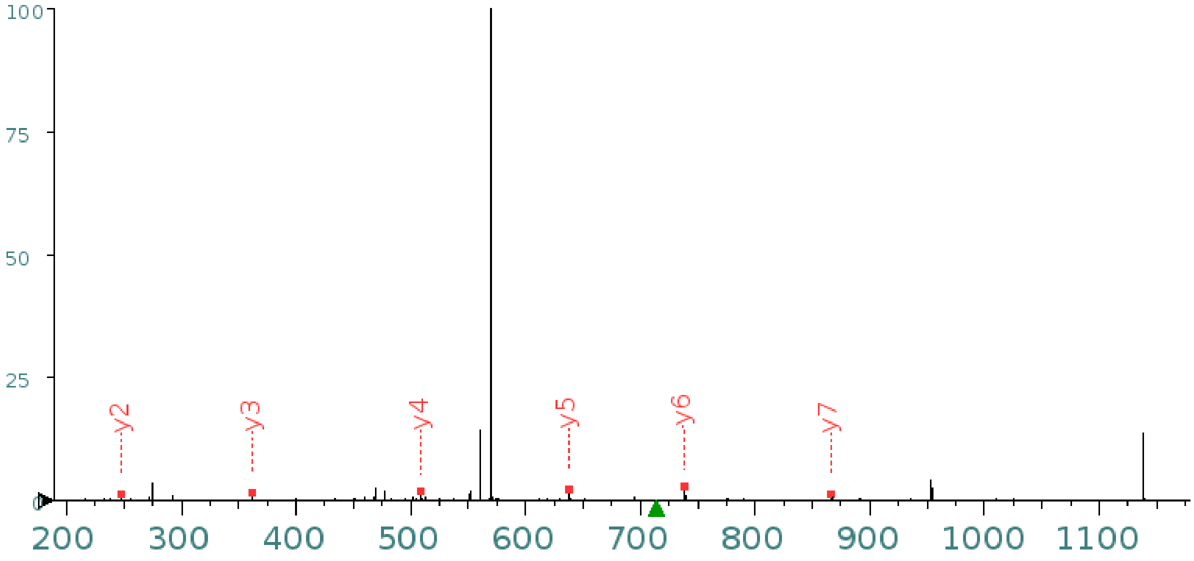

**Sequence**

IAS@QTQFNTK

**Predicted Fragmentation Pattern**

| Seq | # | b: Δ Error | b | y | y: Δ Error | +1 |
|---|---|---|---|---|---|---|
| I | 1 | --- | 114.091 | --- | --- | 10 |
| A | 2 | --- | 185.128 | 1315.601 | --- | 9 |
| S@ | 3 | --- | 563.256 | 1244.564 | --- | 8 |
| Q | 4 | --- | 691.314 | **866.437** | **140.713** | 7 |
| T | 5 | --- | 792.362 | **738.378** | **181.535** | 6 |
| Q | 6 | --- | 920.421 | **637.330** | **67.099** | 5 |
| F | 7 | --- | 1067.489 | **509.272** | **134.742** | 4 |
| N | 8 | --- | 1181.532 | **362.203** | **512.065** | 3 |
| T | 9 | --- | 1282.580 | **248.160** | **654.389** | 2 |
| K | 10 | --- | --- | 147.113 | --- | 1 |

**Soleil Index**

| | | |
|---|---|---|
| **569.5167** | **not explained** | **100%** |

| | | | | | |
|---|---|---|---|---|---|
| **Run:** | 76752 | **Scans:** | 5627 - 5627 | **m/z Range:** | 154.0643 to 1223.6242 |
| **Search:** | 76752 | **Precursor** | 611.81212 | | |

**Mods:** C=160.03 *+15.99 #+220.06 @+291.1

**Mass:**
**Charge:** +2

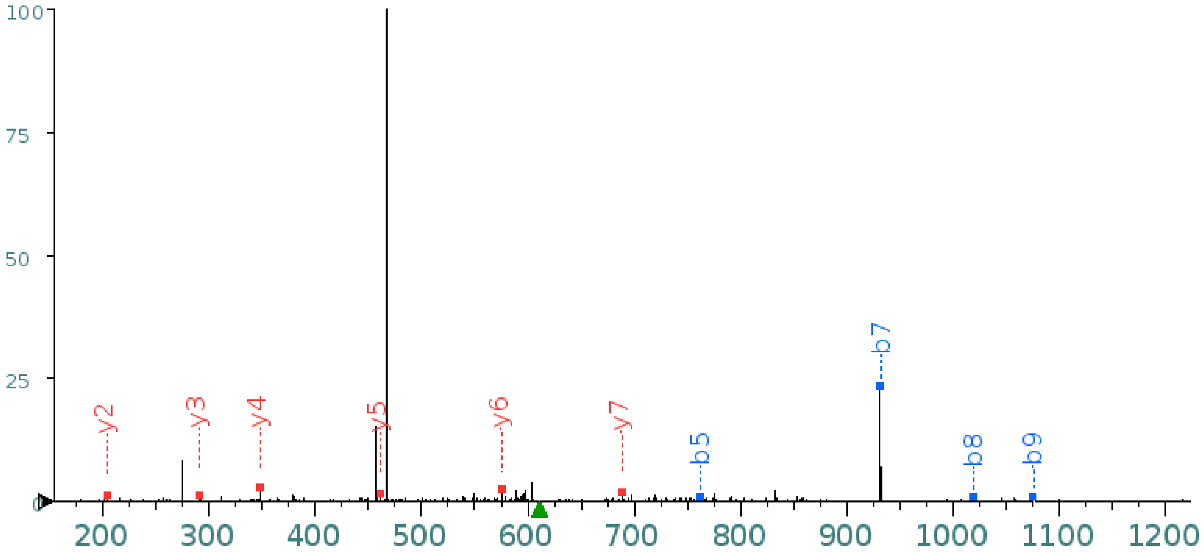

**Sequence**

VGS@INIGSGK

**Predicted Fragmentation Pattern**

| Seq | # | b: Δ Error | b | y | y: Δ Error | +1 |
|---|---|---|---|---|---|---|
| V | 1 | --- | 100.076 | --- | --- | 10 |
| G | 2 | --- | 157.097 | 1123.548 | --- | 9 |
| S@ | 3 | --- | 535.225 | 1066.526 | --- | 8 |
| I | 4 | --- | 648.309 | **688.399** | 219.551 | 7 |
| N | 5 | 139.968 | **762.352** | **575.315** | 12.530 | 6 |
| I | 6 | --- | 875.436 | **461.272** | 569.235 | 5 |
| G | 7 | -890.505 | **932.457** | **348.188** | 668.953 | 4 |
| S | 8 | -767.936 | **1019.489** | **291.166** | 693.663 | 3 |
| G | 9 | -606.000 | **1076.511** | **204.134** | 618.283 | 2 |
| K | 10 | --- | --- | 147.113 | --- | 1 |

**Soleil Index**

| | | |
|---|---|---|
| 457.3243 | not explained | 15% |
| 466.4030 | b1-7 frag doubly charged | 100% |

**Run:** 76752

**Scans:** 6916 - 6916

**m/z Range:** 172.26776 to 1660.8427

| Search: | 76752 | Precursor Mass: | 691.33093 |
|---|---|---|---|
| Mods: | C=160.03 *+15.99 #+220.06 @+291.1 | Charge: | +3 |

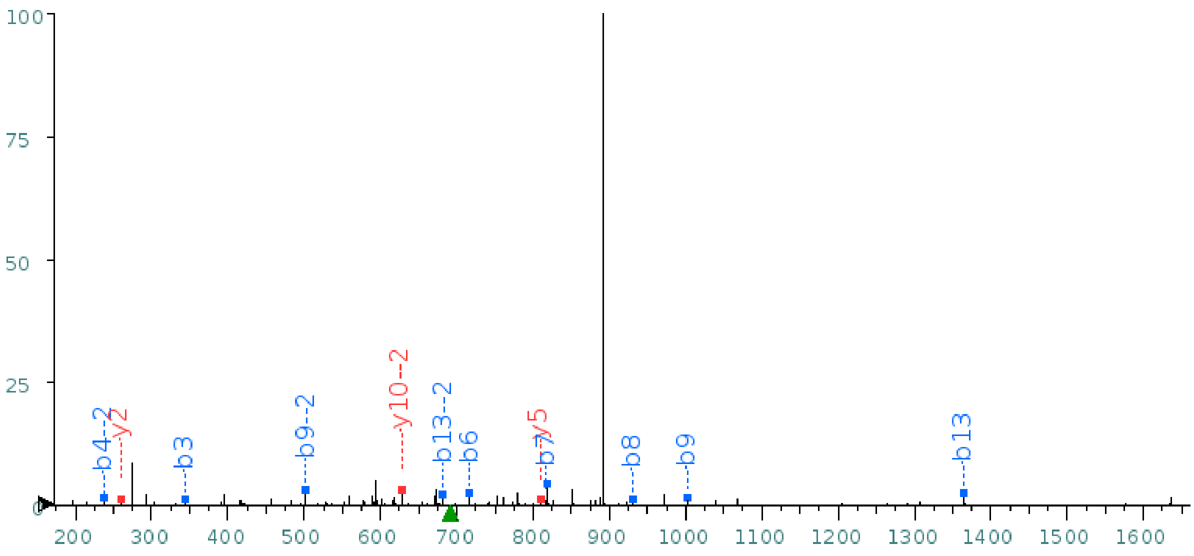

**Sequence**

DQVMELTIAGM*GTS@ALK

**Predicted Fragmentation Pattern**

**+1**

| Seq | # | b: Δ Error | b | y | y: Δ Error | +1 |
|---|---|---|---|---|---|---|
| D | 1 | --- | 116.034 | --- | --- | 17 |
| Q | 2 | --- | 244.093 | 1956.950 | --- | 16 |
| V | 3 | -222.781 | **343.161** | 1828.892 | --- | 15 |
| M | 4 | --- | 474.202 | 1729.823 | --- | 14 |
| E | 5 | --- | 603.244 | 1598.783 | --- | 13 |
| L | 6 | 407.992 | **716.328** | 1469.740 | --- | 12 |
| T | 7 | 1146.626 | **817.376** | 1356.656 | --- | 11 |
| I | 8 | 926.339 | **930.460** | 1255.609 | --- | 10 |
| A | 9 | 21.497 | **1001.497** | 1142.525 | --- | 9 |
| G | 10 | --- | 1058.519 | 1071.487 | --- | 8 |
| M* | 11 | --- | 1205.554 | 1014.466 | --- | 7 |
| G | 12 | --- | 1262.576 | 867.431 | --- | 6 |
| T | 13 | 284.289 | **1363.623** | **810.409** | -949.393 | 5 |
| S@ | 14 | --- | 1741.751 | 709.361 | --- | 4 |
| A | 15 | --- | 1812.788 | 331.234 | --- | 3 |
| L | 16 | --- | 1925.872 | **260.197** | 827.827 | 2 |
| K | 17 | --- | --- | 147.113 | --- | 1 |

**+2**

| Seq | # | b: Δ Error | b | y | y: Δ Error | +1 |
|---|---|---|---|---|---|---|

| | | | | | | |
|---|---|---|---|---|---|---|
| D | 1 | --- | 58.521 | --- | --- | 17 |
| Q | 2 | --- | 122.550 | 978.979 | --- | 16 |
| V | 3 | --- | 172.084 | 914.950 | --- | 15 |
| M | 4 | 2534.041 | **237.604** | 865.415 | --- | 14 |
| E | 5 | --- | 302.126 | 799.895 | --- | 13 |
| L | 6 | --- | 358.668 | 735.374 | --- | 12 |
| T | 7 | --- | 409.192 | 678.832 | --- | 11 |
| I | 8 | --- | 465.734 | **628.308** | -453.798 | 10 |
| A | 9 | 133.591 | **501.252** | 571.766 | --- | 9 |
| G | 10 | --- | 529.763 | 536.247 | --- | 8 |
| M* | 11 | --- | 603.281 | 507.737 | --- | 7 |
| G | 12 | --- | 631.791 | 434.219 | --- | 6 |
| T | 13 | 104.831 | **682.315** | 405.708 | --- | 5 |
| S@ | 14 | --- | 871.379 | 355.184 | --- | 4 |
| A | 15 | --- | 906.898 | 166.121 | --- | 3 |
| L | 16 | --- | 963.440 | 130.602 | --- | 2 |
| K | 17 | --- | --- | 74.060 | --- | 1 |

**Soleil Index**

**891.1617** **not explained** **100%**

| Run: | 76752 | Scans: | 6966 - 6966 | m/z Range: | 210.40036 to 1657.1285 |
|---|---|---|---|---|---|
| Search: | 76752 | Precursor Mass: | 795.63424 | | |
| Mods: | C=160.03 *+15.99 #+220.06 @+291.1 | | | | |

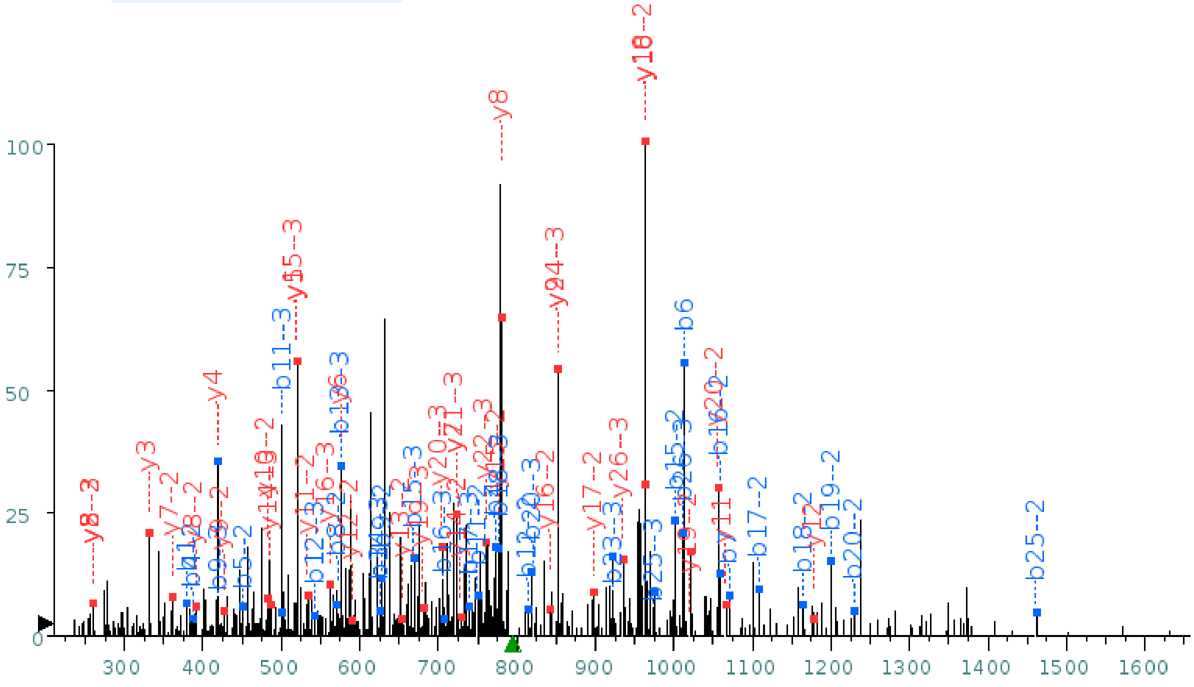

**Sequence**

S@M*VFQIGANKDQVM*ELTIAGM*GTSALK

**Predicted Fragmentation Pattern**

+1

| Seq | # | b: Δ Error | b | y | y: Δ Error | +1 |
|---|---|---|---|---|---|---|
| S@ | 1 | 373.053 | **379.135** | --- | --- | 27 |
| M* | 2 | --- | 526.170 | 2801.388 | --- | 26 |
| V | 3 | 413.421 | **625.239** | 2654.353 | --- | 25 |
| F | 4 | 73.327 | **772.307** | 2555.284 | --- | 24 |
| Q | 5 | --- | 900.366 | 2408.216 | --- | 23 |
| I | 6 | -388.792 | **1013.450** | 2280.157 | --- | 22 |
| G | 7 | 436.002 | **1070.471** | 2167.073 | --- | 21 |
| A | 8 | --- | 1141.508 | 2110.052 | --- | 20 |
| N | 9 | --- | 1255.551 | 2039.015 | --- | 19 |
| K | 10 | --- | 1383.646 | 1924.972 | --- | 18 |
| D | 11 | --- | 1498.673 | 1796.877 | --- | 17 |
| Q | 12 | --- | 1626.732 | 1681.850 | --- | 16 |
| V | 13 | --- | 1725.800 | 1553.791 | --- | 15 |
| M* | 14 | --- | 1872.835 | 1454.723 | --- | 14 |
| E | 15 | --- | 2001.878 | 1307.688 | --- | 13 |
| L | 16 | --- | 2114.962 | **1178.645** | 149.912 | 12 |
| T | 17 | --- | 2216.010 | **1065.561** | 682.577 | 11 |
| I | 18 | --- | 2329.094 | **964.513** | -97.346 | 10 |
| A | 19 | --- | 2400.131 | **851.429** | 149.380 | 9 |
| G | 20 | --- | 2457.152 | **780.392** | 148.361 | 8 |

| | | | | | | |
|---|---|---|---|---|---|---|
| M* | 21 | --- | 2604.188 | **723.371** | -411.260 | 7 |
| G | 22 | --- | 2661.209 | **576.335** | 113.605 | 6 |
| T | 23 | --- | 2762.257 | **519.314** | 320.463 | 5 |
| S | 24 | --- | 2849.289 | **418.266** | 490.164 | 4 |
| A | 25 | --- | 2920.326 | **331.234** | 748.376 | 3 |
| L | 26 | --- | 3033.410 | **260.197** | 725.595 | 2 |
| K | 27 | --- | --- | 147.113 | --- | 1 |

+2

| Seq | # | b: Δ Error | b | y | y: Δ Error | +1 |
|---|---|---|---|---|---|---|
| S@ | 1 | --- | 190.071 | --- | --- | 27 |
| M* | 2 | --- | 263.589 | 1401.198 | --- | 26 |
| V | 3 | --- | 313.123 | 1327.680 | --- | 25 |
| F | 4 | 1773.309 | **386.657** | 1278.146 | --- | 24 |
| Q | 5 | -782.096 | **450.686** | 1204.612 | --- | 23 |
| I | 6 | --- | 507.228 | 1140.582 | --- | 22 |
| G | 7 | --- | 535.739 | 1084.040 | --- | 21 |
| A | 8 | -1427.170 | **571.258** | **1055.530** | 899.022 | 20 |
| N | 9 | -1160.030 | **628.279** | **1020.011** | 559.381 | 19 |
| K | 10 | --- | 692.327 | **962.990** | 737.855 | 18 |
| D | 11 | 1283.467 | **749.840** | **898.942** | -417.390 | 17 |
| Q | 12 | 539.685 | **813.869** | **841.429** | 968.876 | 16 |
| V | 13 | --- | 863.404 | **777.399** | -49.094 | 15 |
| M* | 14 | --- | 936.921 | **727.865** | 971.536 | 14 |
| E | 15 | 873.033 | **1001.443** | **654.347** | 146.251 | 13 |
| L | 16 | 903.835 | **1057.985** | **589.826** | -441.218 | 12 |
| T | 17 | 806.905 | **1108.509** | **533.284** | 262.223 | 11 |
| I | 18 | -73.662 | **1165.051** | **482.760** | 1319.107 | 10 |
| A | 19 | 347.310 | **1200.569** | **426.218** | 322.095 | 9 |
| G | 20 | 266.607 | **1229.080** | **390.700** | -924.584 | 8 |
| M* | 21 | --- | 1302.598 | **362.189** | -2249.985 | 7 |
| G | 22 | --- | 1331.108 | 288.671 | --- | 6 |
| T | 23 | --- | 1381.632 | **260.160** | 865.334 | 5 |
| S | 24 | --- | 1425.148 | 209.637 | --- | 4 |
| A | 25 | 531.958 | **1460.667** | 166.121 | --- | 3 |
| L | 26 | --- | 1517.209 | 130.602 | --- | 2 |
| K | 27 | --- | --- | 74.060 | --- | 1 |

+3

| Seq | # | b: Δ Error | b | y | y: Δ Error | +1 |
|---|---|---|---|---|---|---|
| S@ | 1 | --- | 127.050 | --- | --- | 27 |
| M* | 2 | --- | 176.062 | **934.468** | 621.293 | 26 |
| V | 3 | --- | 209.084 | 885.456 | --- | 25 |
| F | 4 | --- | 258.107 | **852.433** | -1029.461 | 24 |
| Q | 5 | --- | 300.793 | 803.410 | --- | 23 |
| I | 6 | --- | 338.488 | **760.724** | -362.201 | 22 |
| G | 7 | --- | 357.495 | **723.029** | 60.695 | 21 |
| A | 8 | --- | 381.174 | **704.022** | 534.385 | 20 |
| N | 9 | -1714.383 | **419.189** | **680.343** | 277.369 | 19 |
| K | 10 | --- | 461.887 | 642.329 | --- | 18 |
| D | 11 | 269.217 | **500.229** | 599.630 | --- | 17 |
| Q | 12 | -912.742 | **542.915** | **561.288** | -157.827 | 16 |
| V | 13 | 802.311 | **575.938** | **518.602** | 1690.534 | 15 |
| M* | 14 | 874.743 | **624.950** | **485.579** | 1879.011 | 14 |
| E | 15 | 1209.515 | **667.964** | 436.567 | --- | 13 |
| L | 16 | 1383.982 | **705.659** | 393.553 | --- | 12 |
| T | 17 | -684.419 | **739.341** | 355.858 | --- | 11 |
| I | 18 | 418.119 | **777.036** | 322.176 | --- | 10 |

| | | | | | | |
|---|---|---|---|---|---|---|
| A | 19 | --- | 800.715 | 284.481 | --- | 9 |
| G | 20 | -867.916 | **819.722** | **260.802** | -1599.112 | 8 |
| M* | 21 | --- | 868.734 | 241.795 | --- | 7 |
| G | 22 | --- | 887.741 | 192.783 | --- | 6 |
| T | 23 | 1039.622 | **921.424** | 173.776 | --- | 5 |
| S | 24 | --- | 950.434 | 140.094 | --- | 4 |
| A | 25 | 922.015 | **974.114** | 111.083 | --- | 3 |
| L | 26 | -482.812 | **1011.808** | 87.404 | --- | 2 |
| K | 27 | --- | --- | 49.709 | --- | 1 |

**Soleil Index**

| | | |
|---|---|---|
| 344.2639 | y2-7 frag loss of water/ y2-7 frag loss of ammonia | 17% |
| 457.1410 | not explained | 18% |
| 474.4396 | b3-24 frag doubly charged | 22% |
| 484.4609 | not explained | 15% |
| 501.2703 | b2-15 frag doubly charged/ y1-5 frag loss of water/ y3-15 frag loss of water/ y3-15 frag loss of ammonia | 43% |
| 587.4852 | not explained | 29% |
| 614.2974 | b2-20 frag doubly charged | 45% |
| 622.4750 | not explained | 17% |
| 632.4960 | not explained | 64% |
| 637.3392 | y2-13 frag loss of ammonia | 25% |
| 650.5863 | b2-21 frag doubly charged/ b3-15 frag loss of water/ b3-15 frag loss of ammonia | 20% |
| 675.0289 | b2-10 frag loss of water/ b2-10 frag loss of ammonia | 22% |
| 711.2769 | y2-14 frag loss of ammonia | 19% |
| 719.0540 | not explained | 27% |
| 734.4410 | not explained | 22% |
| 752.4711 | not explained | 17% |
| 762.5986 | y1-8 frag loss of water/ y1-8 frag loss of ammonia | 19% |
| 763.6799 | y1-8 frag loss of ammonia | 18% |
| 778.4590 | not explained | 92% |
| 779.3880 | not explained | 34% |
| 781.5604 | not explained | 24% |
| 788.2813 | not explained | 17% |
| 833.8209 | y1-9 frag loss of water/ y1-9 frag loss of ammonia/ y3-24 frag loss of water | 15% |
| 954.4554 | not explained | 23% |
| 955.4347 | b3-25 frag loss of water | 26% |
| 958.0952 | not explained | 23% |
| 969.2045 | not explained | 17% |
| 1237.2545 | b1-9 frag loss of water | 23% |

| Run: | 76752 | Scans: | 7119 - 7119 | m/z Range: | 172.04732 to 1530.7086 |
|---|---|---|---|---|---|
| Search: | 76752 | Precursor Mass: | 691.33083 | | |
| Mods: | C=160.03 *+15.99 #+220.06 @+291.1 | Charge: | +3 | | |

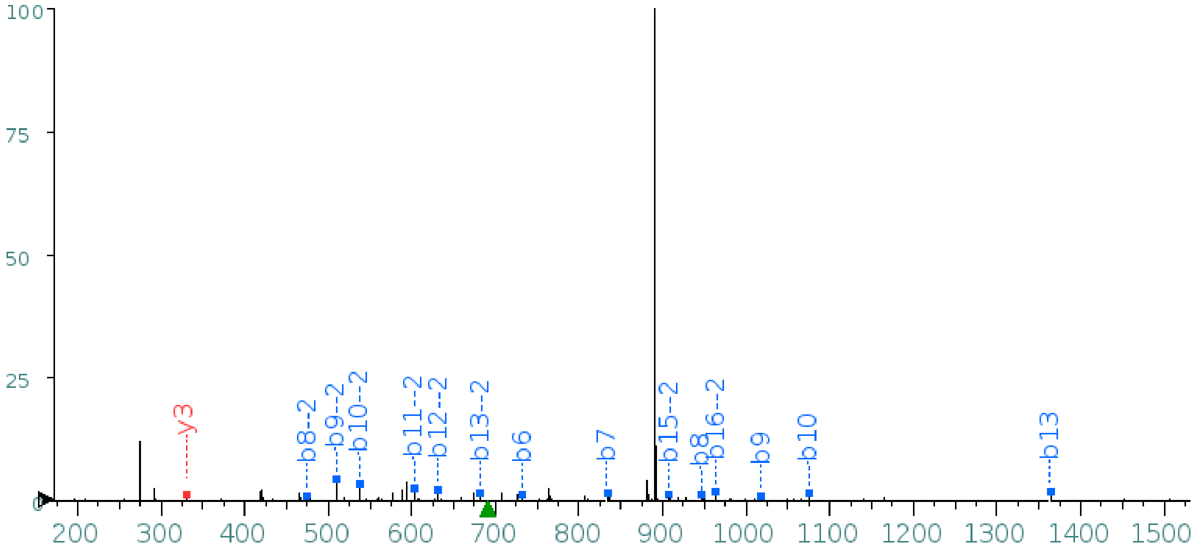

**Sequence**

DQVM*ELTIAGMGTS@ALK

**Predicted Fragmentation Pattern**

+1

| Seq | # | b: Δ Error | b | y | y: Δ Error | +1 |
|---|---|---|---|---|---|---|
| D | 1 | --- | 116.034 | --- | --- | 17 |
| Q | 2 | --- | 244.093 | 1956.950 | --- | 16 |
| V | 3 | --- | 343.161 | 1828.892 | --- | 15 |
| M* | 4 | --- | 490.197 | 1729.823 | --- | 14 |
| E | 5 | --- | 619.239 | 1582.788 | --- | 13 |
| L | 6 | **197.287** | **732.323** | 1453.745 | --- | 12 |
| T | 7 | **342.379** | **833.371** | 1340.661 | --- | 11 |
| I | 8 | **172.033** | **946.455** | 1239.614 | --- | 10 |
| A | 9 | **96.931** | **1017.492** | 1126.530 | --- | 9 |
| G | 10 | **106.743** | **1074.514** | 1055.493 | --- | 8 |
| M | 11 | --- | 1205.554 | 998.471 | --- | 7 |
| G | 12 | --- | 1262.576 | 867.431 | --- | 6 |
| T | 13 | **102.904** | **1363.623** | 810.409 | --- | 5 |
| S@ | 14 | --- | 1741.751 | 709.361 | --- | 4 |
| A | 15 | --- | 1812.788 | **331.234** | **-194.174** | 3 |
| L | 16 | --- | 1925.872 | 260.197 | --- | 2 |
| K | 17 | --- | --- | 147.113 | --- | 1 |

+2

| Seq | # | b: Δ Error | b | y | y: Δ Error | +1 |
|---|---|---|---|---|---|---|

| | | | | | | | |
|---|---|---|---|---|---|---|---|
| D | 1 | --- | 58.521 | --- | | --- | 17 |
| Q | 2 | --- | 122.550 | 978.979 | | --- | 16 |
| V | 3 | --- | 172.084 | 914.950 | | --- | 15 |
| M* | 4 | --- | 245.602 | 865.415 | | --- | 14 |
| E | 5 | --- | 310.123 | 791.898 | | --- | 13 |
| L | 6 | --- | 366.665 | 727.376 | | --- | 12 |
| T | 7 | --- | 417.189 | 670.834 | | --- | 11 |
| I | 8 | **-64.944** | **473.731** | 620.310 | | --- | 10 |
| A | 9 | **339.659** | **509.250** | 563.768 | | --- | 9 |
| G | 10 | **-200.696** | **537.760** | 528.250 | | --- | 8 |
| M | 11 | **1043.758** | **603.281** | 499.739 | | --- | 7 |
| G | 12 | **163.104** | **631.791** | 434.219 | | --- | 6 |
| T | 13 | **-905.813** | **682.315** | 405.708 | | --- | 5 |
| S@ | 14 | --- | 871.379 | 355.184 | | --- | 4 |
| A | 15 | **-216.424** | **906.898** | 166.121 | | --- | 3 |
| L | 16 | **719.235** | **963.440** | 130.602 | | --- | 2 |
| K | 17 | --- | --- | 74.060 | | --- | 1 |

**Soleil Index**

**891.1978** **not explained** **100%**

**Run:** 76752

**Search:** 76752

**Mods:** C=160.03 *+15.99 #+220.06 @+291.1

**Scans:** 7347 - 7347

**Precursor Mass:** 791.63554

**Charge:** +4

**m/z Range:** 203.1037 to 2000

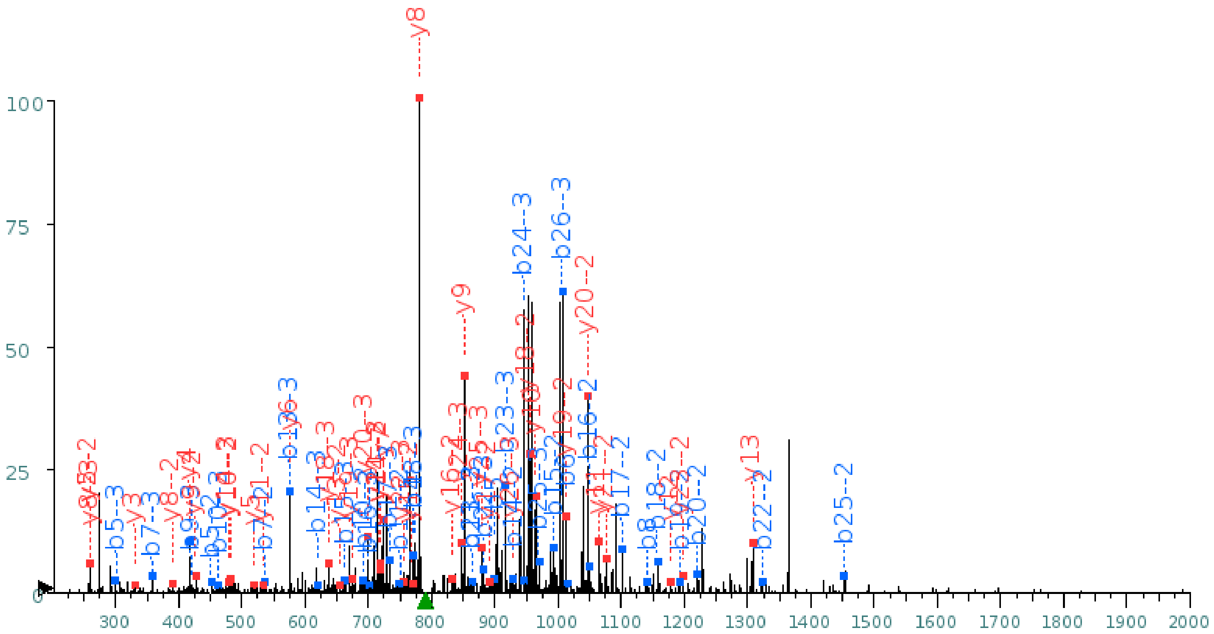

### Sequence

S@M*VFQIGANKDQVMELTIAGM*GTSALK

### Predicted Fragmentation Pattern

+1

| Seq | # | b: Δ Error | b | y | y: Δ Error | +1 |
|---|---|---|---|---|---|---|
| S@ | 1 | --- | 379.135 | --- | --- | 27 |
| M* | 2 | --- | 526.170 | 2785.393 | --- | 26 |
| V | 3 | --- | 625.239 | 2638.358 | --- | 25 |
| F | 4 | -519.977 | **772.307** | 2539.289 | --- | 24 |
| Q | 5 | -782.550 | **900.366** | 2392.221 | --- | 23 |
| I | 6 | 906.943 | **1013.450** | 2264.162 | --- | 22 |
| G | 7 | --- | 1070.471 | 2151.078 | --- | 21 |
| A | 8 | 186.151 | **1141.508** | 2094.057 | --- | 20 |
| N | 9 | --- | 1255.551 | 2023.020 | --- | 19 |
| K | 10 | --- | 1383.646 | 1908.977 | --- | 18 |
| D | 11 | --- | 1498.673 | 1780.882 | --- | 17 |
| Q | 12 | --- | 1626.732 | 1665.855 | --- | 16 |
| V | 13 | --- | 1725.800 | 1537.796 | --- | 15 |
| M | 14 | --- | 1856.840 | 1438.728 | --- | 14 |
| E | 15 | --- | 1985.883 | **1307.688** | 205.022 | 13 |
| L | 16 | --- | 2098.967 | **1178.645** | -533.383 | 12 |
| T | 17 | --- | 2200.015 | **1065.561** | 181.239 | 11 |
| I | 18 | --- | 2313.099 | **964.513** | 64.342 | 10 |
| A | 19 | --- | 2384.136 | **851.429** | -13.610 | 9 |
| G | 20 | --- | 2441.157 | **780.392** | -16.877 | 8 |

| | | | | | | |
|---|---|---|---|---|---|---|
| M* | 21 | --- | 2588.193 | **723.371** | **182.040** | 7 |
| G | 22 | --- | 2645.214 | **576.335** | **260.648** | 6 |
| T | 23 | --- | 2746.262 | **519.314** | **485.695** | 5 |
| S | 24 | --- | 2833.294 | **418.266** | **43.656** | 4 |
| A | 25 | --- | 2904.331 | **331.234** | **484.649** | 3 |
| L | 26 | --- | 3017.415 | **260.197** | **293.952** | 2 |
| K | 27 | --- | --- | 147.113 | --- | 1 |

**+2**

| Seq | # | b: Δ Error | b | y | y: Δ Error | +1 |
|---|---|---|---|---|---|---|
| S@ | 1 | --- | 190.071 | --- | --- | 27 |
| M* | 2 | --- | 263.589 | 1393.200 | --- | 26 |
| V | 3 | --- | 313.123 | 1319.683 | --- | 25 |
| F | 4 | --- | 386.657 | 1270.148 | --- | 24 |
| Q | 5 | **2172.933** | **450.686** | **1196.614** | **543.612** | 23 |
| I | 6 | --- | 507.228 | 1132.585 | --- | 22 |
| G | 7 | **-811.990** | **535.739** | **1076.043** | **445.862** | 21 |
| A | 8 | --- | 571.258 | **1047.532** | **-370.483** | 20 |
| N | 9 | --- | 628.279 | **1012.014** | **-190.444** | 19 |
| K | 10 | **677.475** | **692.327** | **954.992** | **423.589** | 18 |
| D | 11 | **-342.871** | **749.840** | **890.945** | **407.204** | 17 |
| Q | 12 | --- | 813.869 | **833.431** | **313.356** | 16 |
| V | 13 | **238.938** | **863.404** | **769.402** | **326.730** | 15 |
| M | 14 | **778.472** | **928.924** | **719.868** | **-864.935** | 14 |
| E | 15 | **243.742** | **993.445** | **654.347** | **707.295** | 13 |
| L | 16 | **527.630** | **1049.987** | 589.826 | --- | 12 |
| T | 17 | **455.182** | **1100.511** | **533.284** | **845.508** | 11 |
| I | 18 | **552.856** | **1157.053** | **482.760** | **491.861** | 10 |
| A | 19 | **291.450** | **1192.572** | **426.218** | **718.208** | 9 |
| G | 20 | **-793.057** | **1221.082** | **390.700** | **1469.977** | 8 |
| M* | 21 | --- | 1294.600 | 362.189 | --- | 7 |
| G | 22 | **-199.751** | **1323.111** | 288.671 | --- | 6 |
| T | 23 | --- | 1373.635 | **260.160** | **433.751** | 5 |
| S | 24 | --- | 1417.151 | 209.637 | --- | 4 |
| A | 25 | **390.162** | **1452.669** | 166.121 | --- | 3 |
| L | 26 | --- | 1509.211 | 130.602 | --- | 2 |
| K | 27 | --- | --- | 74.060 | --- | 1 |

**+3**

| Seq | # | b: Δ Error | b | y | y: Δ Error | +1 |
|---|---|---|---|---|---|---|
| S@ | 1 | --- | 127.050 | --- | --- | 27 |
| M* | 2 | --- | 176.062 | **929.136** | **550.356** | 26 |
| V | 3 | --- | 209.084 | **880.124** | **-249.647** | 25 |
| F | 4 | --- | 258.107 | **847.101** | **1072.034** | 24 |
| Q | 5 | **-1774.463** | **300.793** | 798.079 | --- | 23 |
| I | 6 | --- | 338.488 | **755.392** | **24.387** | 22 |
| G | 7 | **2251.559** | **357.495** | **717.698** | **292.684** | 21 |
| A | 8 | --- | 381.174 | **698.691** | **327.551** | 20 |
| N | 9 | **-2161.876** | **419.189** | **675.011** | **104.713** | 19 |
| K | 10 | **-1108.652** | **461.887** | **636.997** | **1024.060** | 18 |
| D | 11 | --- | 500.229 | 594.299 | --- | 17 |
| Q | 12 | --- | 542.915 | 555.957 | --- | 16 |
| V | 13 | **949.252** | **575.938** | 513.270 | --- | 15 |
| M | 14 | **-93.878** | **619.618** | **480.248** | **368.759** | 14 |
| E | 15 | **1095.318** | **662.633** | 436.567 | --- | 13 |
| L | 16 | **638.825** | **700.327** | 393.553 | --- | 12 |
| T | 17 | **520.817** | **734.010** | 355.858 | --- | 11 |
| I | 18 | **260.526** | **771.704** | 322.176 | --- | 10 |

| | | | | | | |
|---|---|---|---|---|---|---|
| A | 19 | --- | 795.384 | 284.481 | --- | 9 |
| G | 20 | --- | 814.391 | **260.802** | -2031.759 | 8 |
| M* | 21 | 240.289 | **863.402** | 241.795 | --- | 7 |
| G | 22 | 786.640 | **882.410** | 192.783 | --- | 6 |
| T | 23 | 423.337 | **916.092** | 173.776 | --- | 5 |
| S | 24 | 428.679 | **945.103** | 140.094 | --- | 4 |
| A | 25 | 789.697 | **968.782** | 111.083 | --- | 3 |
| L | 26 | 492.939 | **1006.477** | 87.404 | --- | 2 |
| K | 27 | --- | --- | 49.709 | --- | 1 |

**Soleil Index**

| | | |
|---|---|---|
| 274.0756 | not explained | 20% |
| 714.8589 | not explained | 24% |
| 728.5735 | not explained | 20% |
| 765.9980 | not explained | 25% |
| 904.7445 | not explained | 21% |
| 937.5723 | y2-18 frag loss of water/ y2-18 frag loss of ammonia | 15% |
| 946.2530 | y1-10 frag loss of water | 58% |
| 952.5754 | b3-25 frag loss of ammonia | 60% |
| 958.5984 | not explained | 59% |
| 963.2338 | not explained | 17% |
| 1002.8700 | not explained | 59% |
| 1038.9363 | not explained | 22% |
| 1092.0421 | not explained | 18% |
| 1364.6124 | not explained | 31% |

**Run:** 76752  **Scans:** 7630 - 7630  **m/z Range:** 213.22818 to 1951.1090

**Search:** 76752  **Precursor Mass:** 791.63614

**Mods:** C=160.03 *+15.99 #+220.06 @+291.1  **Charge:** +4

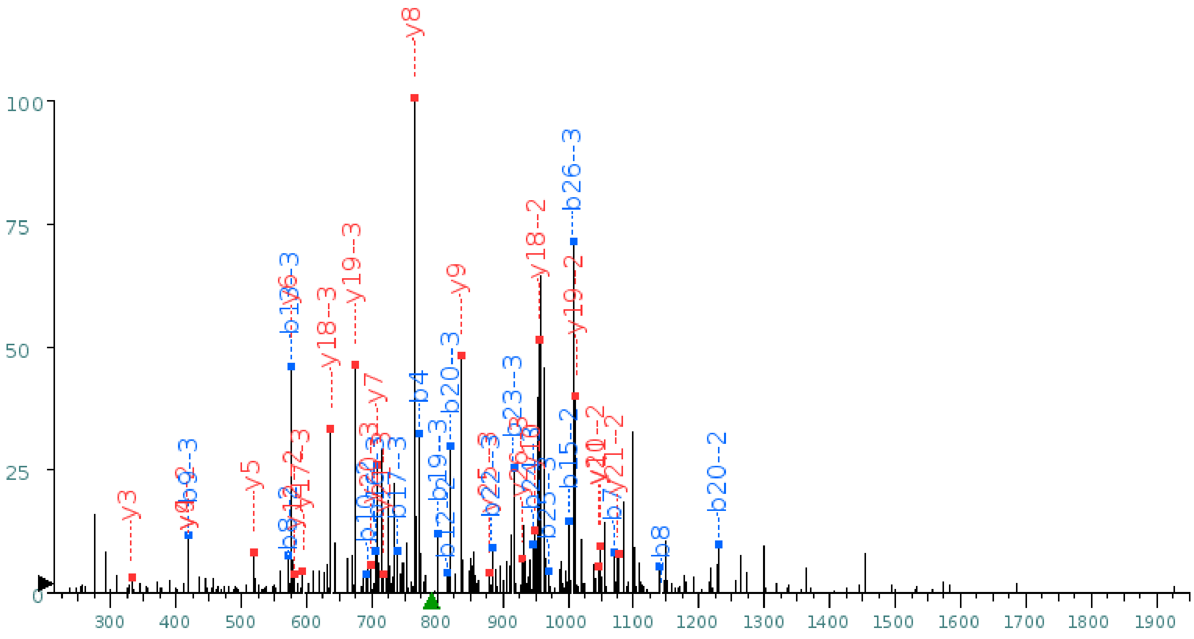

### Sequence

S@M*VFQIGANKDQVM*ELTIAGMGTSALK

### Predicted Fragmentation Pattern

+1

| Seq | # | b: Δ Error | b | y | y: Δ Error | +1 |
|---|---|---|---|---|---|---|
| S@ | 1 | --- | 379.135 | --- | --- | 27 |
| M* | 2 | --- | 526.170 | 2785.393 | --- | 26 |
| V | 3 | --- | 625.239 | 2638.358 | --- | 25 |
| F | 4 | -1005.247 | **772.307** | 2539.289 | --- | 24 |
| Q | 5 | --- | 900.366 | 2392.221 | --- | 23 |
| I | 6 | --- | 1013.450 | 2264.162 | --- | 22 |
| G | 7 | -292.686 | **1070.471** | 2151.078 | --- | 21 |
| A | 8 | -855.585 | **1141.508** | 2094.057 | --- | 20 |
| N | 9 | --- | 1255.551 | 2023.020 | --- | 19 |
| K | 10 | --- | 1383.646 | 1908.977 | --- | 18 |
| D | 11 | --- | 1498.673 | 1780.882 | --- | 17 |
| Q | 12 | --- | 1626.732 | 1665.855 | --- | 16 |
| V | 13 | --- | 1725.800 | 1537.796 | --- | 15 |
| M* | 14 | --- | 1872.835 | 1438.728 | --- | 14 |
| E | 15 | --- | 2001.878 | 1291.693 | --- | 13 |
| L | 16 | --- | 2114.962 | 1162.650 | --- | 12 |
| T | 17 | --- | 2216.010 | **1049.566** | 296.337 | 11 |
| I | 18 | --- | 2329.094 | **948.518** | 116.039 | 10 |
| A | 19 | --- | 2400.131 | **835.434** | 139.727 | 9 |
| G | 20 | --- | 2457.152 | **764.397** | 22.987 | 8 |

| | | | | | | |
|---|---|---|---|---|---|---|
| M | 21 | --- | 2588.193 | **707.376** | **86.497** | 7 |
| G | 22 | --- | 2645.214 | **576.335** | **171.729** | 6 |
| T | 23 | --- | 2746.262 | **519.314** | **418.881** | 5 |
| S | 24 | --- | 2833.294 | **418.266** | **269.403** | 4 |
| A | 25 | --- | 2904.331 | **331.234** | **2838.541** | 3 |
| L | 26 | --- | 3017.415 | 260.197 | --- | 2 |
| K | 27 | --- | --- | 147.113 | --- | 1 |

**+2**

| Seq | # | b: Δ Error | b | y | y: Δ Error | +1 |
|---|---|---|---|---|---|---|
| S@ | 1 | --- | 190.071 | --- | --- | 27 |
| M* | 2 | --- | 263.589 | 1393.200 | --- | 26 |
| V | 3 | --- | 313.123 | 1319.683 | --- | 25 |
| F | 4 | --- | 386.657 | 1270.148 | --- | 24 |
| Q | 5 | --- | 450.686 | 1196.614 | --- | 23 |
| I | 6 | --- | 507.228 | 1132.585 | --- | 22 |
| G | 7 | --- | 535.739 | **1076.043** | **800.272** | 21 |
| A | 8 | **-233.886** | **571.258** | **1047.532** | **-55.598** | 20 |
| N | 9 | --- | 628.279 | **1012.014** | **-396.586** | 19 |
| K | 10 | **-1433.775** | **692.327** | **954.992** | **-295.708** | 18 |
| D | 11 | --- | 749.840 | 890.945 | --- | 17 |
| Q | 12 | **923.541** | **813.869** | 833.431 | --- | 16 |
| V | 13 | --- | 863.404 | 769.402 | --- | 15 |
| M* | 14 | --- | 936.921 | 719.868 | --- | 14 |
| E | 15 | **388.992** | **1001.443** | 646.350 | --- | 13 |
| L | 16 | --- | 1057.985 | **581.829** | **-667.685** | 12 |
| T | 17 | --- | 1108.509 | 525.287 | --- | 11 |
| I | 18 | --- | 1165.051 | 474.763 | --- | 10 |
| A | 19 | --- | 1200.569 | **418.221** | **377.595** | 9 |
| G | 20 | **600.921** | **1229.080** | 382.702 | --- | 8 |
| M | 21 | --- | 1294.600 | 354.191 | --- | 7 |
| G | 22 | --- | 1323.111 | 288.671 | --- | 6 |
| T | 23 | --- | 1373.635 | 260.160 | --- | 5 |
| S | 24 | --- | 1417.151 | 209.637 | --- | 4 |
| A | 25 | --- | 1452.669 | 166.121 | --- | 3 |
| L | 26 | --- | 1509.211 | 130.602 | --- | 2 |
| K | 27 | --- | --- | 74.060 | --- | 1 |

**+3**

| Seq | # | b: Δ Error | b | y | y: Δ Error | +1 |
|---|---|---|---|---|---|---|
| S@ | 1 | --- | 127.050 | --- | --- | 27 |
| M* | 2 | --- | 176.062 | **929.136** | **-251.022** | 26 |
| V | 3 | --- | 209.084 | **880.124** | **-38.143** | 25 |
| F | 4 | --- | 258.107 | 847.101 | --- | 24 |
| Q | 5 | --- | 300.793 | 798.079 | --- | 23 |
| I | 6 | --- | 338.488 | 755.392 | --- | 22 |
| G | 7 | --- | 357.495 | **717.698** | **306.622** | 21 |
| A | 8 | --- | 381.174 | **698.691** | **-322.727** | 20 |
| N | 9 | **-1935.631** | **419.189** | **675.011** | **-543.528** | 19 |
| K | 10 | --- | 461.887 | **636.997** | **-379.711** | 18 |
| D | 11 | --- | 500.229 | **594.299** | **-1427.535** | 17 |
| Q | 12 | --- | 542.915 | 555.957 | --- | 16 |
| V | 13 | **860.394** | **575.938** | 513.270 | --- | 15 |
| M* | 14 | --- | 624.950 | 480.248 | --- | 14 |
| E | 15 | --- | 667.964 | 431.236 | --- | 13 |
| L | 16 | **-233.492** | **705.659** | 388.222 | --- | 12 |
| T | 17 | **300.244** | **739.341** | 350.527 | --- | 11 |
| I | 18 | --- | 777.036 | 316.844 | --- | 10 |

| | | | | | | |
|---|---|---|---|---|---|---|
| A | 19 | 974.886 | **800.715** | 279.150 | --- | 9 |
| G | 20 | 458.059 | **819.722** | 255.471 | --- | 8 |
| M | 21 | --- | 863.402 | 236.463 | --- | 7 |
| G | 22 | 898.297 | **882.410** | 192.783 | --- | 6 |
| T | 23 | 1037.526 | **916.092** | 173.776 | --- | 5 |
| S | 24 | 401.061 | **945.103** | 140.094 | --- | 4 |
| A | 25 | 998.175 | **968.782** | 111.083 | --- | 3 |
| L | 26 | 637.650 | **1006.477** | 87.404 | --- | 2 |
| K | 27 | --- | --- | 49.709 | --- | 1 |

**Soleil Index**

| | | |
|---|---|---|
| 274.1454 | not explained | 16% |
| 714.8871 | not explained | 29% |
| 723.0048 | b3-17 frag loss of ammonia | 19% |
| 733.6037 | b2-11 frag loss of ammonia | 22% |
| 765.4578 | not explained | 57% |
| 766.5876 | not explained | 15% |
| 952.4990 | b3-25 frag loss of ammonia | 40% |
| 953.9440 | y1-18 frag doubly charged | 18% |
| 958.2672 | not explained | 33% |
| 959.1539 | not explained | 65% |
| 963.2248 | not explained | 46% |
| 1010.5789 | y1-19 frag doubly charged | 31% |
| 1084.1610 | not explained | 18% |
| 1099.7013 | not explained | 33% |

| Run: | 76752 | Scans: | 7857 - 7857 | m/z Range: | 171.24835 | to | 1646.0291 |
|---|---|---|---|---|---|---|---|
| Search: | 76752 | Precursor Mass: | 685.99884 | | | | |
| Mods: | C=160.03 *+15.99 #+220.06 @+291.1 | Charge: | +3 | | | | |

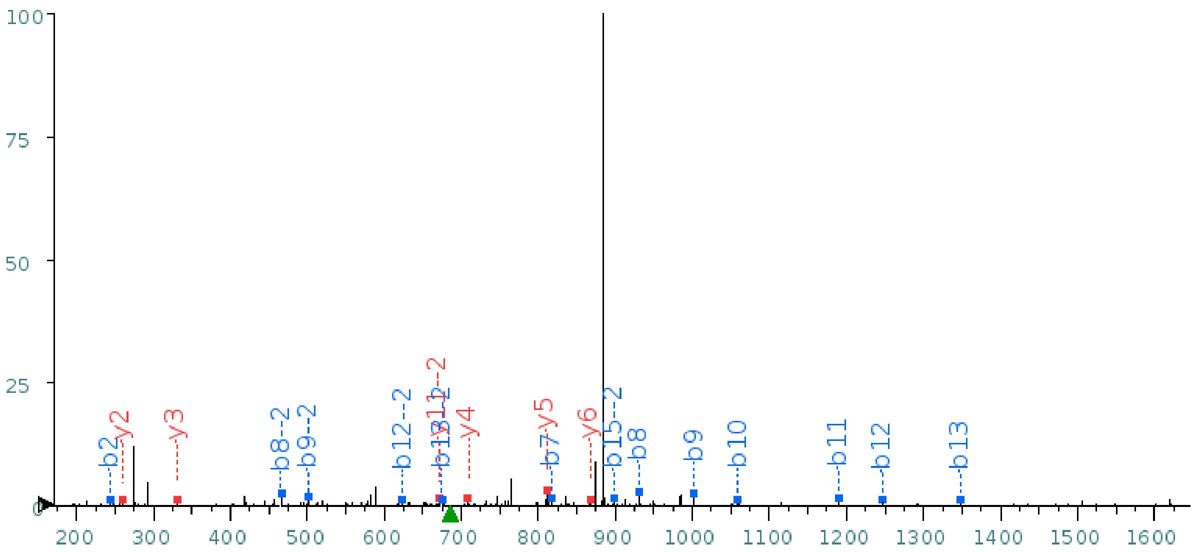

### Sequence

DQVMELTIAGMGTS@ALK

### Predicted Fragmentation Pattern

+1

| Seq | # | b: Δ Error | b | y | y: Δ Error | +1 |
|---|---|---|---|---|---|---|
| D | 1 | --- | 116.034 | --- | --- | 17 |
| Q | 2 | 1051.817 | **244.093** | 1940.956 | --- | 16 |
| V | 3 | --- | 343.161 | 1812.897 | --- | 15 |
| M | 4 | --- | 474.202 | 1713.829 | --- | 14 |
| E | 5 | --- | 603.244 | 1582.788 | --- | 13 |
| L | 6 | --- | 716.328 | 1453.745 | --- | 12 |
| T | 7 | 200.239 | **817.376** | 1340.661 | --- | 11 |
| I | 8 | -108.193 | **930.460** | 1239.614 | --- | 10 |
| A | 9 | 188.143 | **1001.497** | 1126.530 | --- | 9 |
| G | 10 | 150.361 | **1058.519** | 1055.493 | --- | 8 |
| M | 11 | 183.795 | **1189.559** | 998.471 | --- | 7 |
| G | 12 | 276.124 | **1246.581** | **867.431** | 194.621 | 6 |
| T | 13 | 53.438 | **1347.628** | **810.409** | 154.532 | 5 |
| S@ | 14 | --- | 1725.756 | **709.361** | -939.232 | 4 |
| A | 15 | --- | 1796.793 | **331.234** | -2422.607 | 3 |
| L | 16 | --- | 1909.877 | **260.197** | 1119.888 | 2 |
| K | 17 | --- | --- | 147.113 | --- | 1 |

+2

| Seq | # | b: Δ Error | b | y | y: Δ Error | +1 |
|---|---|---|---|---|---|---|

| | | | | | | |
|---|---|---|---|---|---|---|
| D | 1 | --- | 58.521 | --- | --- | 17 |
| Q | 2 | --- | 122.550 | 970.981 | --- | 16 |
| V | 3 | --- | 172.084 | 906.952 | --- | 15 |
| M | 4 | --- | 237.604 | 857.418 | --- | 14 |
| E | 5 | --- | 302.126 | 791.898 | --- | 13 |
| L | 6 | --- | 358.668 | 727.376 | --- | 12 |
| T | 7 | --- | 409.192 | **670.834** | -201.079 | 11 |
| I | 8 | 1576.983 | **465.734** | 620.310 | --- | 10 |
| A | 9 | 1811.778 | **501.252** | 563.768 | --- | 9 |
| G | 10 | --- | 529.763 | 528.250 | --- | 8 |
| M | 11 | --- | 595.283 | 499.739 | --- | 7 |
| G | 12 | 2.148 | **623.794** | 434.219 | --- | 6 |
| T | 13 | 1181.201 | **674.318** | 405.708 | --- | 5 |
| S@ | 14 | --- | 863.382 | 355.184 | --- | 4 |
| A | 15 | -144.289 | **898.900** | 166.121 | --- | 3 |
| L | 16 | --- | 955.442 | 130.602 | --- | 2 |
| K | 17 | --- | --- | 74.060 | --- | 1 |

**Soleil Index**

**883.3753** **not explained** **100%**

| Run: | 76752 | Scans: | 8061 - 8061 | m/z Range: | 197.30284 to 2000 |
|---|---|---|---|---|---|
| Search: | 76752 | Precursor Mass: | 787.63673 | | |
| Mods: | C=160.03 *+15.99 #+220.06 @+291.1 | Charge: | +4 | | |

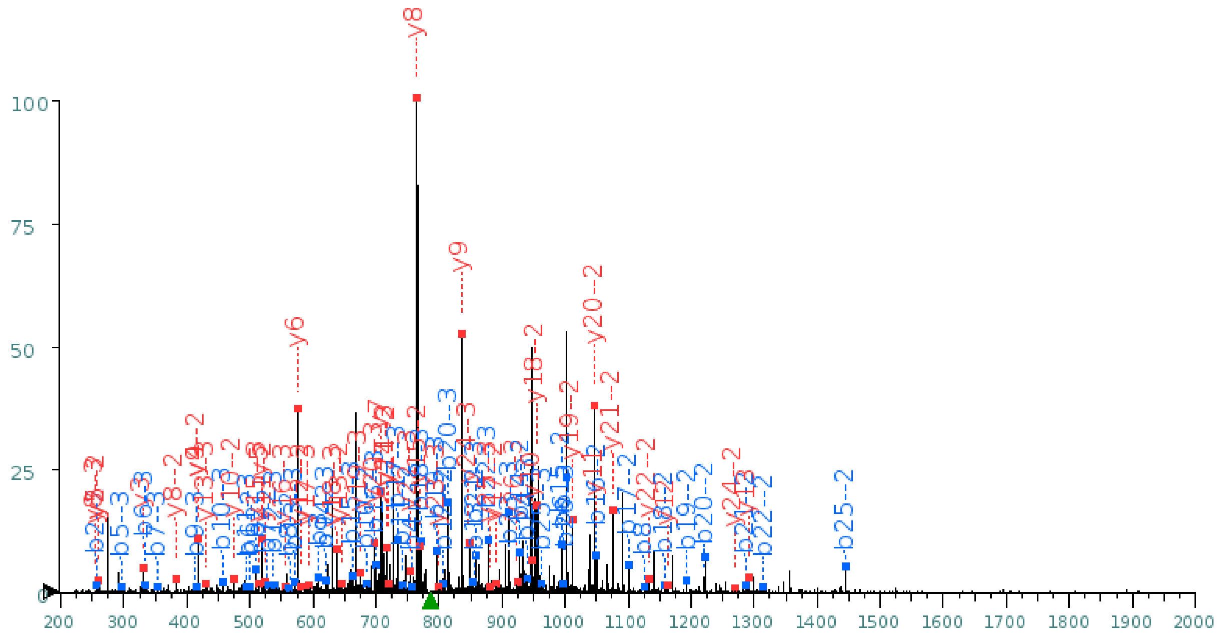

#### Sequence

S@MVFQIGANKDQVM*ELTIAGMGTSALK

#### Predicted Fragmentation Pattern

+1

| Seq | # | b: Δ Error | b | y | y: Δ Error | +1 |
|---|---|---|---|---|---|---|
| S@ | 1 | --- | 379.135 | --- | --- | 27 |
| M | 2 | 463.418 | **510.175** | 2769.398 | --- | 26 |
| V | 3 | 306.010 | **609.244** | 2638.358 | --- | 25 |
| F | 4 | -806.806 | **756.312** | 2539.289 | --- | 24 |
| Q | 5 | --- | 884.371 | 2392.221 | --- | 23 |
| I | 6 | 766.840 | **997.455** | 2264.162 | --- | 22 |
| G | 7 | --- | 1054.476 | 2151.078 | --- | 21 |
| A | 8 | -649.821 | **1125.513** | 2094.057 | --- | 20 |
| N | 9 | --- | 1239.556 | 2023.020 | --- | 19 |
| K | 10 | --- | 1367.651 | 1908.977 | --- | 18 |
| D | 11 | --- | 1482.678 | 1780.882 | --- | 17 |
| Q | 12 | --- | 1610.737 | 1665.855 | --- | 16 |
| V | 13 | --- | 1709.805 | 1537.796 | --- | 15 |
| M* | 14 | --- | 1856.840 | 1438.728 | --- | 14 |
| E | 15 | --- | 1985.883 | **1291.693** | -532.616 | 13 |
| L | 16 | --- | 2098.967 | **1162.650** | -718.667 | 12 |
| T | 17 | --- | 2200.015 | **1049.566** | 35.667 | 11 |
| I | 18 | --- | 2313.099 | **948.518** | -69.017 | 10 |
| A | 19 | --- | 2384.136 | **835.434** | -33.986 | 9 |
| G | 20 | --- | 2441.157 | **764.397** | -64.769 | 8 |

| | | | | | | |
|---|---|---|---|---|---|---|
| M | 21 | --- | 2572.198 | 707.376 | -136.903 | 7 |
| G | 22 | --- | 2629.219 | 576.335 | 10.257 | 6 |
| T | 23 | --- | 2730.267 | 519.314 | -22.858 | 5 |
| S | 24 | --- | 2817.299 | 418.266 | -38.718 | 4 |
| A | 25 | --- | 2888.336 | 331.234 | 463.662 | 3 |
| L | 26 | --- | 3001.420 | 260.197 | 402.367 | 2 |
| K | 27 | --- | --- | 147.113 | --- | 1 |

+2

| Seq | # | b: Δ Error | b | y | y: Δ Error | +1 |
|---|---|---|---|---|---|---|
| S@ | 1 | --- | 190.071 | --- | --- | 27 |
| M | 2 | 2517.779 | 255.591 | 1385.203 | --- | 26 |
| V | 3 | --- | 305.125 | 1319.683 | --- | 25 |
| F | 4 | --- | 378.660 | 1270.148 | -314.030 | 24 |
| Q | 5 | --- | 442.689 | 1196.614 | --- | 23 |
| I | 6 | 657.888 | 499.231 | 1132.585 | -274.734 | 22 |
| G | 7 | 1440.910 | 527.742 | 1076.043 | -441.150 | 21 |
| A | 8 | -1320.293 | 563.260 | 1047.532 | -213.890 | 20 |
| N | 9 | 613.058 | 620.282 | 1012.014 | -62.011 | 19 |
| K | 10 | 440.778 | 684.329 | 954.992 | 21.892 | 18 |
| D | 11 | 1109.767 | 741.843 | 890.945 | 737.436 | 17 |
| Q | 12 | -691.540 | 805.872 | 833.431 | --- | 16 |
| V | 13 | -821.999 | 855.406 | 769.402 | 1008.278 | 15 |
| M* | 14 | -847.975 | 928.924 | 719.868 | -1057.006 | 14 |
| E | 15 | 106.169 | 993.445 | 646.350 | 75.019 | 13 |
| L | 16 | -365.674 | 1049.987 | 581.829 | -517.077 | 12 |
| T | 17 | 724.957 | 1100.511 | 525.287 | 127.108 | 11 |
| I | 18 | -146.332 | 1157.053 | 474.763 | -159.098 | 10 |
| A | 19 | 295.338 | 1192.572 | 418.221 | 69.507 | 9 |
| G | 20 | 525.486 | 1221.082 | 382.702 | -1192.078 | 8 |
| M | 21 | 610.966 | 1286.603 | 354.191 | --- | 7 |
| G | 22 | -62.889 | 1315.113 | 288.671 | --- | 6 |
| T | 23 | --- | 1365.637 | 260.160 | 542.150 | 5 |
| S | 24 | --- | 1409.153 | 209.637 | --- | 4 |
| A | 25 | 452.278 | 1444.672 | 166.121 | --- | 3 |
| L | 26 | --- | 1501.214 | 130.602 | --- | 2 |
| K | 27 | --- | --- | 74.060 | --- | 1 |

+3

| Seq | # | b: Δ Error | b | y | y: Δ Error | +1 |
|---|---|---|---|---|---|---|
| S@ | 1 | --- | 127.050 | --- | --- | 27 |
| M | 2 | --- | 170.730 | 923.804 | 926.703 | 26 |
| V | 3 | --- | 203.753 | 880.124 | 495.941 | 25 |
| F | 4 | --- | 252.776 | 847.101 | 831.695 | 24 |
| Q | 5 | -287.606 | 295.462 | 798.079 | 239.184 | 23 |
| I | 6 | -1796.524 | 333.156 | 755.392 | -496.448 | 22 |
| G | 7 | 378.057 | 352.164 | 717.698 | -125.318 | 21 |
| A | 8 | --- | 375.843 | 698.691 | -446.344 | 20 |
| N | 9 | 1060.921 | 413.857 | 675.011 | 411.172 | 19 |
| K | 10 | -547.469 | 456.555 | 636.997 | 496.720 | 18 |
| D | 11 | -394.365 | 494.898 | 594.299 | -1467.293 | 17 |
| Q | 12 | 1465.644 | 537.584 | 555.957 | -626.118 | 16 |
| V | 13 | -113.998 | 570.607 | 513.270 | 386.437 | 15 |
| M* | 14 | -48.954 | 619.618 | 480.248 | --- | 14 |
| E | 15 | 785.215 | 662.633 | 431.236 | -1711.480 | 13 |
| L | 16 | -1369.059 | 700.327 | 388.222 | --- | 12 |
| T | 17 | 598.063 | 734.010 | 350.527 | --- | 11 |
| I | 18 | -377.183 | 771.704 | 316.844 | --- | 10 |

| | | | | | | |
|---|---|---|---|---|---|---|
| A | 19 | 445.449 | 795.384 | 279.150 | --- | 9 |
| G | 20 | 276.943 | 814.391 | 255.471 | 2988.795 | 8 |
| M | 21 | 547.371 | 858.071 | 236.463 | --- | 7 |
| G | 22 | 258.337 | 877.078 | 192.783 | --- | 6 |
| T | 23 | 545.492 | 910.761 | 173.776 | --- | 5 |
| S | 24 | 998.489 | 939.771 | 140.094 | --- | 4 |
| A | 25 | -671.844 | 963.450 | 111.083 | --- | 3 |
| L | 26 | 379.316 | 1001.145 | 87.404 | --- | 2 |
| K | 27 | --- | --- | 49.709 | --- | 1 |

**Soleil Index**

| | | |
|---|---|---|
| 274.0206 | not explained | 16% |
| 631.2343 | not explained | 25% |
| 668.9927 | not explained | 37% |
| 710.5455 | not explained | 18% |
| 728.2385 | not explained | 24% |
| 765.8452 | not explained | 83% |
| 945.9002 | b3-25 frag loss of water/ b3-25 frag loss of ammonia | 25% |
| 946.5031 | b3-25 frag loss of ammonia | 32% |
| 947.1627 | b3-25 frag loss of ammonia | 50% |
| 953.1608 | not explained | 19% |
| 957.7407 | not explained | 26% |
| 1002.7200 | not explained | 53% |
| 1091.7728 | not explained | 20% |

| | | | |
|---|---|---|---|
| **Run:** | 76752 | **Scans:** | 8462 - 8462 |
| **Search:** | 76752 | **Precursor Mass:** | 783.63893 |
| **Mods:** | C=160.03 *+15.99 #+220.06 @+291.1 | **Charge:** | +4 |

**m/z Range:** 235.60861 **to** 1695.9356

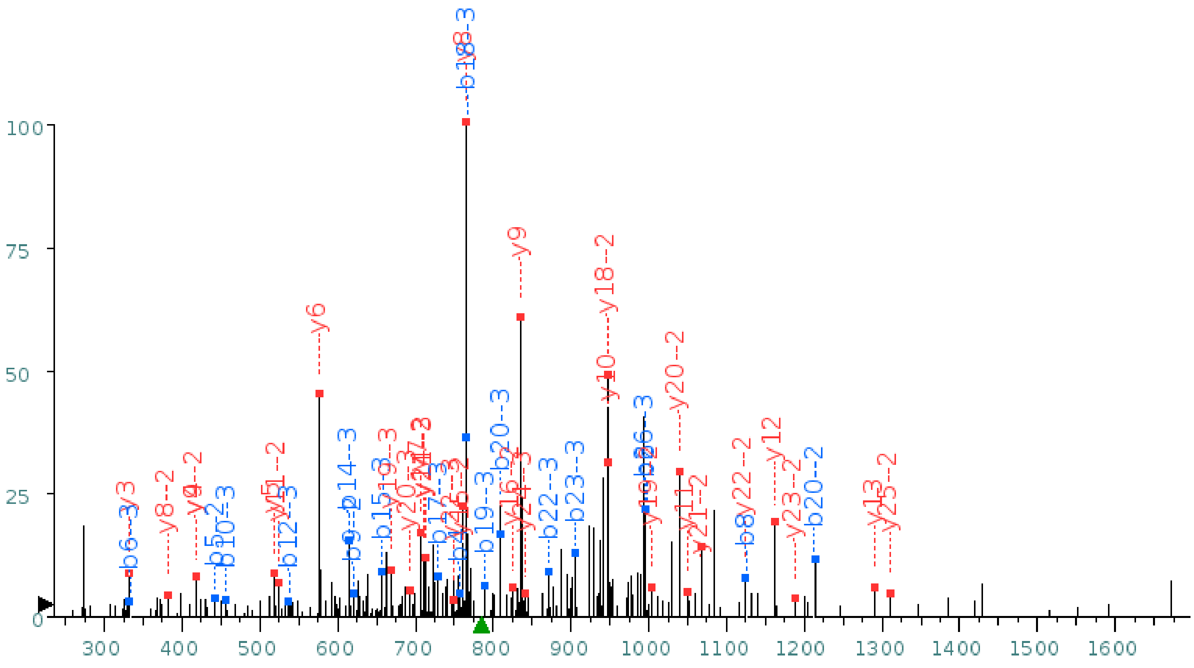

**Sequence**

S@MVFQIGANKDQVMELTIAGMGTSALK

**Predicted Fragmentation Pattern**

**+1**

| Seq | # | b: Δ Error | b | y | y: Δ Error | +1 |
|---|---|---|---|---|---|---|
| S@ | 1 | --- | 379.135 | --- | --- | 27 |
| M | 2 | --- | 510.175 | 2753.403 | --- | 26 |
| V | 3 | --- | 609.244 | 2622.363 | --- | 25 |
| F | 4 | 429.515 | **756.312** | 2523.295 | --- | 24 |
| Q | 5 | --- | 884.371 | 2376.226 | --- | 23 |
| I | 6 | -723.337 | **997.455** | 2248.168 | --- | 22 |
| G | 7 | --- | 1054.476 | 2135.084 | --- | 21 |
| A | 8 | -255.981 | **1125.513** | 2078.062 | --- | 20 |
| N | 9 | --- | 1239.556 | 2007.025 | --- | 19 |
| K | 10 | --- | 1367.651 | 1892.982 | --- | 18 |
| D | 11 | --- | 1482.678 | 1764.887 | --- | 17 |
| Q | 12 | --- | 1610.737 | 1649.860 | --- | 16 |
| V | 13 | --- | 1709.805 | 1521.802 | --- | 15 |
| M | 14 | --- | 1840.846 | 1422.733 | --- | 14 |
| E | 15 | --- | 1969.888 | **1291.693** | 10.594 | 13 |
| L | 16 | --- | 2082.972 | **1162.650** | -228.935 | 12 |
| T | 17 | --- | 2184.020 | **1049.566** | 140.556 | 11 |
| I | 18 | --- | 2297.104 | **948.518** | 29.697 | 10 |
| A | 19 | --- | 2368.141 | **835.434** | 56.531 | 9 |
| G | 20 | --- | 2425.163 | **764.397** | 58.596 | 8 |

| | | | | | | |
|---|---|---|---|---|---|---|
| M | 21 | --- | 2556.203 | **707.376** | **220.541** | 7 |
| G | 22 | --- | 2613.224 | **576.335** | **170.564** | 6 |
| T | 23 | --- | 2714.272 | **519.314** | **-118.071** | 5 |
| S | 24 | --- | 2801.304 | **418.266** | **126.162** | 4 |
| A | 25 | --- | 2872.341 | **331.234** | **277.876** | 3 |
| L | 26 | --- | 2985.425 | 260.197 | --- | 2 |
| K | 27 | --- | --- | 147.113 | --- | 1 |

**+2**

| Seq | # | b: Δ Error | b | y | y: Δ Error | +1 |
|---|---|---|---|---|---|---|
| S@ | 1 | --- | 190.071 | --- | --- | 27 |
| M | 2 | --- | 255.591 | 1377.205 | --- | 26 |
| V | 3 | --- | 305.125 | **1311.685** | **-598.342** | 25 |
| F | 4 | --- | 378.660 | 1262.151 | --- | 24 |
| Q | 5 | **-302.197** | **442.689** | **1188.617** | **507.993** | 23 |
| I | 6 | --- | 499.231 | **1124.587** | **566.819** | 22 |
| G | 7 | --- | 527.742 | **1068.045** | **-425.675** | 21 |
| A | 8 | --- | 563.260 | **1039.535** | **-177.219** | 20 |
| N | 9 | **-398.267** | **620.282** | **1004.016** | **158.409** | 19 |
| K | 10 | --- | 684.329 | **946.995** | **552.358** | 18 |
| D | 11 | --- | 741.843 | 882.947 | --- | 17 |
| Q | 12 | --- | 805.872 | **825.434** | **-1055.947** | 16 |
| V | 13 | --- | 855.406 | **761.404** | **-1235.013** | 15 |
| M | 14 | --- | 920.926 | **711.870** | **1191.797** | 14 |
| E | 15 | --- | 985.448 | 646.350 | --- | 13 |
| L | 16 | --- | 1041.990 | 581.829 | --- | 12 |
| T | 17 | --- | 1092.514 | **525.287** | **300.860** | 11 |
| I | 18 | --- | 1149.056 | 474.763 | --- | 10 |
| A | 19 | --- | 1184.574 | **418.221** | **234.369** | 9 |
| G | 20 | **404.203** | **1213.085** | **382.702** | **-1041.348** | 8 |
| M | 21 | --- | 1278.605 | 354.191 | --- | 7 |
| G | 22 | --- | 1307.116 | 288.671 | --- | 6 |
| T | 23 | --- | 1357.640 | 260.160 | --- | 5 |
| S | 24 | --- | 1401.156 | 209.637 | --- | 4 |
| A | 25 | --- | 1436.674 | 166.121 | --- | 3 |
| L | 26 | --- | 1493.216 | 130.602 | --- | 2 |
| K | 27 | --- | --- | 74.060 | --- | 1 |

**+3**

| Seq | # | b: Δ Error | b | y | y: Δ Error | +1 |
|---|---|---|---|---|---|---|
| S@ | 1 | --- | 127.050 | --- | --- | 27 |
| M | 2 | --- | 170.730 | 918.473 | --- | 26 |
| V | 3 | --- | 203.753 | 874.793 | --- | 25 |
| F | 4 | --- | 252.776 | **841.770** | **-985.908** | 24 |
| Q | 5 | --- | 295.462 | 792.747 | --- | 23 |
| I | 6 | **-2411.182** | **333.156** | **750.061** | **-428.991** | 22 |
| G | 7 | --- | 352.164 | **712.366** | **496.109** | 21 |
| A | 8 | --- | 375.843 | **693.359** | **-439.773** | 20 |
| N | 9 | --- | 413.857 | **669.680** | **-372.564** | 19 |
| K | 10 | **16.920** | **456.555** | 631.666 | --- | 18 |
| D | 11 | --- | 494.898 | 588.967 | --- | 17 |
| Q | 12 | **-722.610** | **537.584** | 550.625 | --- | 16 |
| V | 13 | --- | 570.607 | 507.939 | --- | 15 |
| M | 14 | **981.959** | **614.287** | 474.916 | --- | 14 |
| E | 15 | **-370.080** | **657.301** | 431.236 | --- | 13 |
| L | 16 | --- | 694.996 | 388.222 | --- | 12 |
| T | 17 | **107.637** | **728.678** | 350.527 | --- | 11 |
| I | 18 | **-1061.282** | **766.373** | 316.844 | --- | 10 |

| | | | | | | |
|---|---|---|---|---|---|---|
| A | 19 | -983.951 | **790.052** | 279.150 | --- | 9 |
| G | 20 | -14.498 | **809.059** | 255.471 | --- | 8 |
| M | 21 | --- | 852.739 | 236.463 | --- | 7 |
| G | 22 | 644.620 | **871.746** | 192.783 | --- | 6 |
| T | 23 | 784.968 | **905.429** | 173.776 | --- | 5 |
| S | 24 | --- | 934.440 | 140.094 | --- | 4 |
| A | 25 | --- | 958.119 | 111.083 | --- | 3 |
| L | 26 | 923.411 | **995.813** | 87.404 | --- | 2 |
| K | 27 | --- | --- | 49.709 | --- | 1 |

**Soleil Index**

| | | |
|---|---|---|
| 274.1593 | not explained | 18% |
| 810.1478 | not explained | 23% |
| 836.4810 | b2-13 frag loss of water/ b3-21 frag loss of ammonia | 24% |
| 922.7877 | not explained | 18% |
| 929.1862 | y2-18 frag loss of water/ y2-18 frag loss of ammonia | 18% |
| 938.6080 | not explained | 15% |
| 942.1539 | not explained | 28% |
| 994.5011 | not explained | 41% |
| 1030.1840 | not explained | 15% |
| 1083.7479 | not explained | 17% |
| 1084.5914 | not explained | 22% |

| | | | | | |
|---|---|---|---|---|---|
| **Run:** | 76752 | **Scans:** | 10505 - 10505 | **m/z Range:** | 203.29324 to 1810.0238 |
| **Search:** | 76752 | **Precursor Mass:** | 805.37936 | | |
| **Mods:** | C=160.03 *+15.99 #+220.06 @+291.1 | **Charge:** | +3 | | |

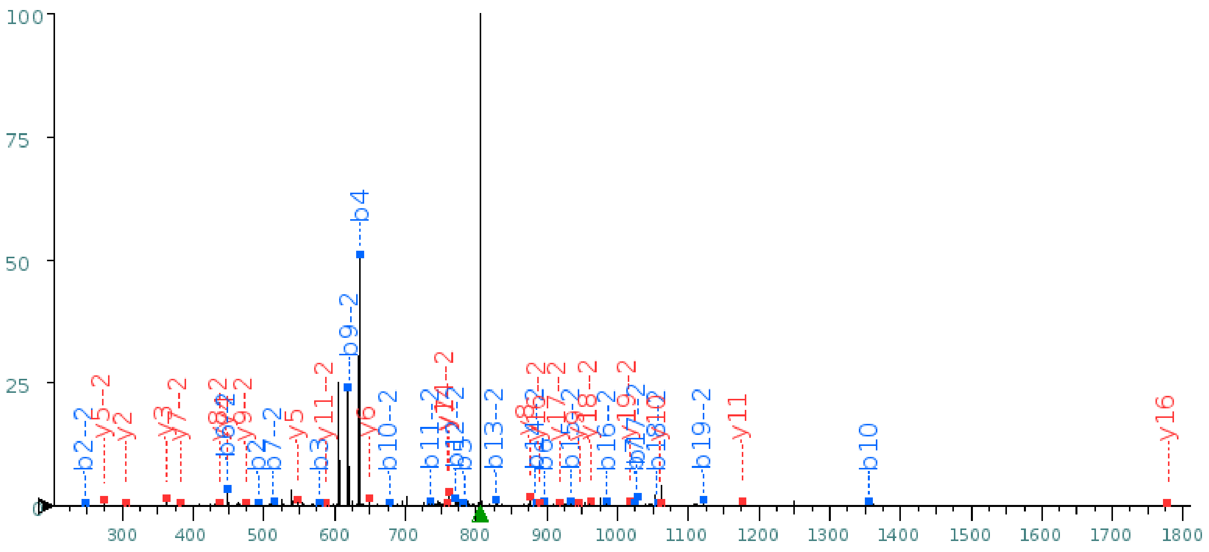

**Sequence**

S@ISGM*IETIDNAINTVSGER

**Predicted Fragmentation Pattern**

| Seq | # | b: Δ Error | b | y | y: Δ Error | +1 |
|---|---|---|---|---|---|---|
| S@ | 1 | --- | 379.135 | --- | --- | 20 |
| I | 2 | 551.666 | **492.219** | 2035.996 | --- | 19 |
| S | 3 | -1310.757 | **579.251** | 1922.912 | --- | 18 |
| G | 4 | -1474.161 | **636.272** | 1835.880 | --- | 17 |
| M* | 5 | -1073.641 | **783.308** | **1778.859** | -493.720 | 16 |
| I | 6 | 238.235 | **896.392** | 1631.824 | --- | 15 |
| E | 7 | -559.347 | **1025.434** | 1518.739 | --- | 14 |
| T | 8 | --- | 1126.482 | 1389.697 | --- | 13 |
| I | 9 | --- | 1239.566 | 1288.649 | --- | 12 |
| D | 10 | 159.927 | **1354.593** | **1175.565** | 118.431 | 11 |
| N | 11 | --- | 1468.636 | **1060.538** | 274.141 | 10 |
| A | 12 | --- | 1539.673 | **946.495** | -929.255 | 9 |
| I | 13 | --- | 1652.757 | **875.458** | 35.495 | 8 |
| N | 14 | --- | 1766.800 | **762.374** | 30.777 | 7 |
| T | 15 | --- | 1867.848 | **648.331** | 27.372 | 6 |
| V | 16 | --- | 1966.916 | **547.283** | 16.386 | 5 |
| S | 17 | --- | 2053.948 | **448.215** | 119.862 | 4 |
| G | 18 | --- | 2110.970 | **361.183** | 78.332 | 3 |
| E | 19 | --- | 2240.012 | **304.162** | 592.867 | 2 |
| R | 20 | --- | --- | 175.119 | --- | 1 |

| | | | | +2 | | |
|---|---|---|---|---|---|---|
| Seq | # | b: Δ Error | b | y | y: Δ Error | +1 |
| S@ | 1 | --- | 190.071 | --- | --- | 20 |
| I | 2 | -845.390 | 246.613 | 1018.502 | 259.318 | 19 |
| S | 3 | --- | 290.129 | 961.960 | 327.545 | 18 |
| G | 4 | --- | 318.640 | 918.444 | 136.973 | 17 |
| M* | 5 | --- | 392.157 | 889.933 | 320.339 | 16 |
| I | 6 | -960.900 | 448.700 | 816.415 | --- | 15 |
| E | 7 | 1174.118 | 513.221 | 759.873 | 298.221 | 14 |
| T | 8 | --- | 563.745 | 695.352 | --- | 13 |
| I | 9 | -1603.105 | 620.287 | 644.828 | --- | 12 |
| D | 10 | -330.196 | 677.800 | 588.286 | -1312.399 | 11 |
| N | 11 | -365.130 | 734.822 | 530.773 | --- | 10 |
| A | 12 | 436.664 | 770.340 | 473.751 | 1037.378 | 9 |
| I | 13 | 402.787 | 826.882 | 438.233 | -20.409 | 8 |
| N | 14 | 447.762 | 883.904 | 381.691 | 457.464 | 7 |
| T | 15 | 348.597 | 934.428 | 324.669 | --- | 6 |
| V | 16 | 154.450 | 983.962 | 274.145 | 375.084 | 5 |
| S | 17 | 258.035 | 1027.478 | 224.611 | --- | 4 |
| G | 18 | 548.046 | 1055.988 | 181.095 | --- | 3 |
| E | 19 | 354.054 | 1120.510 | 152.584 | --- | 2 |
| R | 20 | --- | --- | 88.063 | --- | 1 |

Soleil Index

| | | |
|---|---|---|
| 605.2715 | not explained | 25% |
| 634.3469 | not explained | 30% |
| 805.3434 | precursor | 100% |
| 806.3130 | precursor | 33% |

| Run: | 76752 | Scans: | 11455 - 11455 | m/z Range: | 210.2386 to 1822.2085 |
|---|---|---|---|---|---|
| Search: | 76752 | Precursor Mass: | 795.63546 | | |
| Mods: | C=160.03 *+15.99 #+220.06 @+291.1 | Charge: | +4 | | |

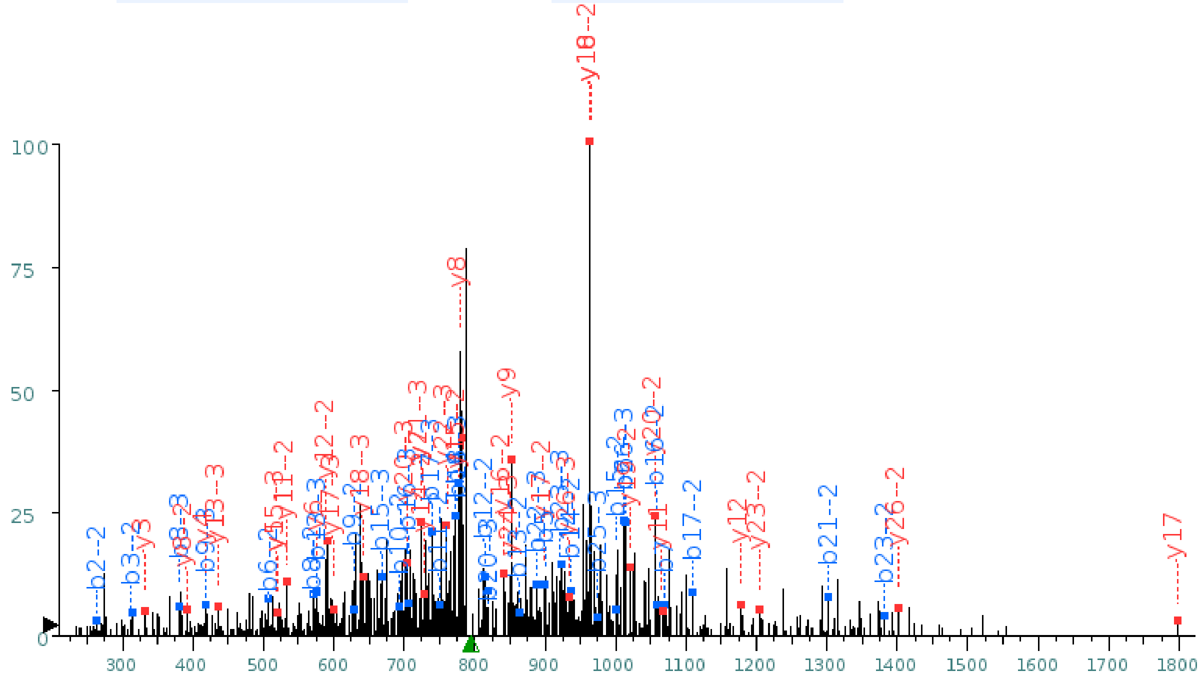

### Sequence

S@M*VFQIGANKDQVM*ELTIAGM*GTSALK

### Predicted Fragmentation Pattern

+1

| Seq | # | b: Δ Error | b | y | y: Δ Error | +1 |
|---|---|---|---|---|---|---|
| S@ | 1 | --- | 379.135 | --- | --- | 27 |
| M* | 2 | --- | 526.170 | 2801.388 | --- | 26 |
| V | 3 | --- | 625.239 | 2654.353 | --- | 25 |
| F | 4 | -878.323 | **772.307** | 2555.284 | --- | 24 |
| Q | 5 | -973.712 | **900.366** | 2408.216 | --- | 23 |
| I | 6 | 23.780 | **1013.450** | 2280.157 | --- | 22 |
| G | 7 | -710.812 | **1070.471** | 2167.073 | --- | 21 |
| A | 8 | --- | 1141.508 | 2110.052 | --- | 20 |
| N | 9 | --- | 1255.551 | 2039.015 | --- | 19 |
| K | 10 | --- | 1383.646 | 1924.972 | --- | 18 |
| D | 11 | --- | 1498.673 | **1796.877** | 184.527 | 17 |
| Q | 12 | --- | 1626.732 | 1681.850 | --- | 16 |
| V | 13 | --- | 1725.800 | 1553.791 | --- | 15 |
| M* | 14 | --- | 1872.835 | 1454.723 | --- | 14 |
| E | 15 | --- | 2001.878 | 1307.688 | --- | 13 |
| L | 16 | --- | 2114.962 | **1178.645** | 28.448 | 12 |
| T | 17 | --- | 2216.010 | **1065.561** | 464.706 | 11 |
| I | 18 | --- | 2329.094 | **964.513** | -582.216 | 10 |

| | | | | | | |
|---|---|---|---|---|---|---|
| A | 19 | --- | 2400.131 | 851.429 | 102.367 | 9 |
| G | 20 | --- | 2457.152 | 780.392 | 419.520 | 8 |
| M* | 21 | --- | 2604.188 | 723.371 | -164.487 | 7 |
| G | 22 | --- | 2661.209 | 576.335 | -1736.432 | 6 |
| T | 23 | --- | 2762.257 | 519.314 | 165.634 | 5 |
| S | 24 | --- | 2849.289 | 418.266 | 223.605 | 4 |
| A | 25 | --- | 2920.326 | 331.234 | 481.795 | 3 |
| L | 26 | --- | 3033.410 | 260.197 | --- | 2 |
| K | 27 | --- | --- | 147.113 | --- | 1 |

+2

| Seq | # | b: Δ Error | b | y | y: Δ Error | +1 |
|---|---|---|---|---|---|---|
| S@ | 1 | --- | 190.071 | --- | --- | 27 |
| M* | 2 | 2379.393 | 263.589 | 1401.198 | -75.700 | 26 |
| V | 3 | 943.184 | 313.123 | 1327.680 | --- | 25 |
| F | 4 | --- | 386.657 | 1278.146 | --- | 24 |
| Q | 5 | --- | 450.686 | 1204.612 | -434.659 | 23 |
| I | 6 | -1951.192 | 507.228 | 1140.582 | --- | 22 |
| G | 7 | --- | 535.739 | 1084.040 | --- | 21 |
| A | 8 | 1213.279 | 571.258 | 1055.530 | -131.538 | 20 |
| N | 9 | -958.608 | 628.279 | 1020.011 | -86.082 | 19 |
| K | 10 | -649.817 | 692.327 | 962.990 | 998.405 | 18 |
| D | 11 | -200.836 | 749.840 | 898.942 | 608.807 | 17 |
| Q | 12 | 810.272 | 813.869 | 841.429 | 389.509 | 16 |
| V | 13 | 129.761 | 863.404 | 777.399 | 420.154 | 15 |
| M* | 14 | -577.327 | 936.921 | 727.865 | 232.155 | 14 |
| E | 15 | 261.451 | 1001.443 | 654.347 | --- | 13 |
| L | 16 | -722.035 | 1057.985 | 589.826 | 439.915 | 12 |
| T | 17 | 601.818 | 1108.509 | 533.284 | 278.238 | 11 |
| I | 18 | --- | 1165.051 | 482.760 | --- | 10 |
| A | 19 | --- | 1200.569 | 426.218 | --- | 9 |
| G | 20 | --- | 1229.080 | 390.700 | 539.684 | 8 |
| M* | 21 | 200.420 | 1302.598 | 362.189 | --- | 7 |
| G | 22 | --- | 1331.108 | 288.671 | --- | 6 |
| T | 23 | -682.732 | 1381.632 | 260.160 | --- | 5 |
| S | 24 | --- | 1425.148 | 209.637 | --- | 4 |
| A | 25 | --- | 1460.667 | 166.121 | --- | 3 |
| L | 26 | --- | 1517.209 | 130.602 | --- | 2 |
| K | 27 | --- | --- | 74.060 | --- | 1 |

+3

| Seq | # | b: Δ Error | b | y | y: Δ Error | +1 |
|---|---|---|---|---|---|---|
| S@ | 1 | --- | 127.050 | --- | --- | 27 |
| M* | 2 | --- | 176.062 | 934.468 | 842.128 | 26 |
| V | 3 | --- | 209.084 | 885.456 | --- | 25 |
| F | 4 | --- | 258.107 | 852.433 | -1076.530 | 24 |
| Q | 5 | --- | 300.793 | 803.410 | --- | 23 |
| I | 6 | --- | 338.488 | 760.724 | -697.930 | 22 |
| G | 7 | --- | 357.495 | 723.029 | 307.352 | 21 |
| A | 8 | -2313.329 | 381.174 | 704.022 | 99.799 | 20 |
| N | 9 | -1981.530 | 419.189 | 680.343 | --- | 19 |
| K | 10 | --- | 461.887 | 642.329 | 50.080 | 18 |
| D | 11 | --- | 500.229 | 599.630 | -647.653 | 17 |
| Q | 12 | --- | 542.915 | 561.288 | --- | 16 |
| V | 13 | -1046.452 | 575.938 | 518.602 | 1535.916 | 15 |
| M* | 14 | --- | 624.950 | 485.579 | --- | 14 |
| E | 15 | 1217.902 | 667.964 | 436.567 | -800.564 | 13 |
| L | 16 | -143.937 | 705.659 | 393.553 | --- | 12 |

| | | | | | | |
|---|---|---|---|---|---|---|
| T | 17 | 361.210 | **739.341** | 355.858 | --- | 11 |
| I | 18 | 887.149 | **777.036** | 322.176 | --- | 10 |
| A | 19 | --- | 800.715 | 284.481 | --- | 9 |
| G | 20 | -228.736 | **819.722** | 260.802 | --- | 8 |
| M* | 21 | --- | 868.734 | 241.795 | --- | 7 |
| G | 22 | 298.586 | **887.741** | 192.783 | --- | 6 |
| T | 23 | 1001.943 | **921.424** | 173.776 | --- | 5 |
| S | 24 | --- | 950.434 | 140.094 | --- | 4 |
| A | 25 | -401.303 | **974.114** | 111.083 | --- | 3 |
| L | 26 | -881.486 | **1011.808** | 87.404 | --- | 2 |
| K | 27 | --- | --- | 49.709 | --- | 1 |

**Soleil Index**

| | | |
|---|---|---|
| 630.2529 | not explained | 21% |
| 636.8936 | y2-13 frag loss of water/ y2-13 frag loss of ammonia | 27% |
| 674.4332 | b2-10 frag loss of water/ b2-10 frag loss of ammonia | 20% |
| 700.7680 | y2-26 frag doubly charged | 17% |
| 709.5375 | y2-14 frag loss of water | 18% |
| 724.6679 | not explained | 19% |
| 730.4218 | b2-25 frag doubly charged | 19% |
| 753.6120 | b1-4 frag loss of water | 24% |
| 765.2208 | not explained | 16% |
| 766.9911 | not explained | 17% |
| 770.3421 | not explained | 20% |
| 778.4191 | not explained | 58% |
| 779.2259 | not explained | 34% |
| 781.8410 | b3-19 frag loss of water | 46% |
| 784.0728 | b3-19 frag loss of ammonia | 23% |
| 787.4933 | precursor loss of two waters | 79% |
| 871.5663 | b3-22 frag loss of ammonia | 18% |
| 954.2322 | not explained | 27% |
| 957.5816 | b3-25 frag loss of ammonia | 19% |
| 968.6898 | not explained | 17% |
| 978.8351 | not explained | 18% |
| 1002.9281 | y2-19 frag loss of water/ y2-19 frag loss of ammonia | 17% |
| 1027.6051 | not explained | 17% |
| 1075.8214 | not explained | 17% |

**MS/MS data of trypsin digested peptides with site score A and site score B above 19 from *Clostridium botulinum CbFla* flagellin protein co-expressed with *Clostridium botulinum* Maf in EV136 cells. Results of dataset 2 (MS/MS results obtained in April 2019).**

| | | | |
|---|---|---|---|
| **Run:** | 77660 | **Scans:** | 9594 - 9594 |
| **Search:** | 77660 | **Precursor Mass:** | 714.84613 |
| **Mods:** | C=160.03 *+15.99 #+220.06 @+291.1 | **Charge:** | +2 |

**m/z Range:** 191.25385 **to** 1183.7534

**Sequence**

IASQTQFNT@K

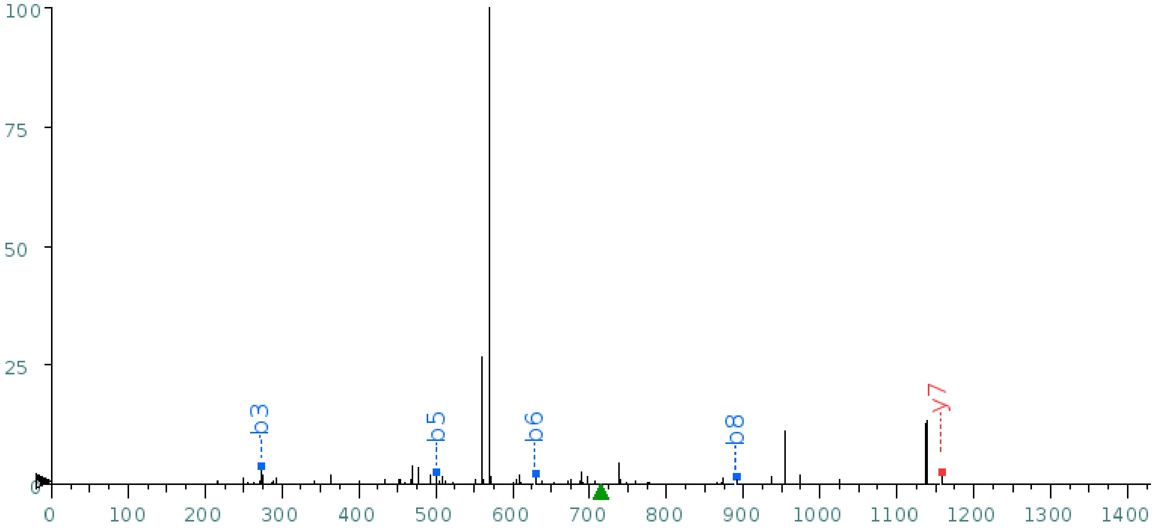

**Predicted Fragmentation Pattern**

| Seq | # | b: Δ Error | b | y | y: Δ Error | +1 |
|---|---|---|---|---|---|---|
| I | 1 | --- | 114.091 | --- | --- | 10 |
| A | 2 | --- | 185.128 | 1315.601 | --- | 9 |
| S | 3 | 730.483 | **272.160** | 1244.564 | --- | 8 |
| Q | 4 | --- | 400.219 | **1157.532** | 163.034 | 7 |
| T | 5 | 436.297 | **501.267** | 1029.474 | --- | 6 |
| Q | 6 | 447.679 | **629.325** | 928.426 | --- | 5 |
| F | 7 | --- | 776.394 | 800.367 | --- | 4 |
| N | 8 | 308.071 | **890.437** | 653.299 | --- | 3 |
| T@ | 9 | --- | 1282.580 | 539.256 | --- | 2 |
| K | 10 | --- | --- | 147.113 | --- | 1 |

| | | | |
|---|---|---|---|
| **Run:** | 77660 | **Scans:** | 9901 - 9901 |
| **Search:** | 77660 | **Precursor Mass:** | 611.81214 |
| **Mods:** | C=160.03 *+15.99 #+220.06 @+291.1 | **Charge:** | +2 |

**m/z Range:** 172.17535 **to** 1132.0225

**Sequence**

VGS@INIGSGK

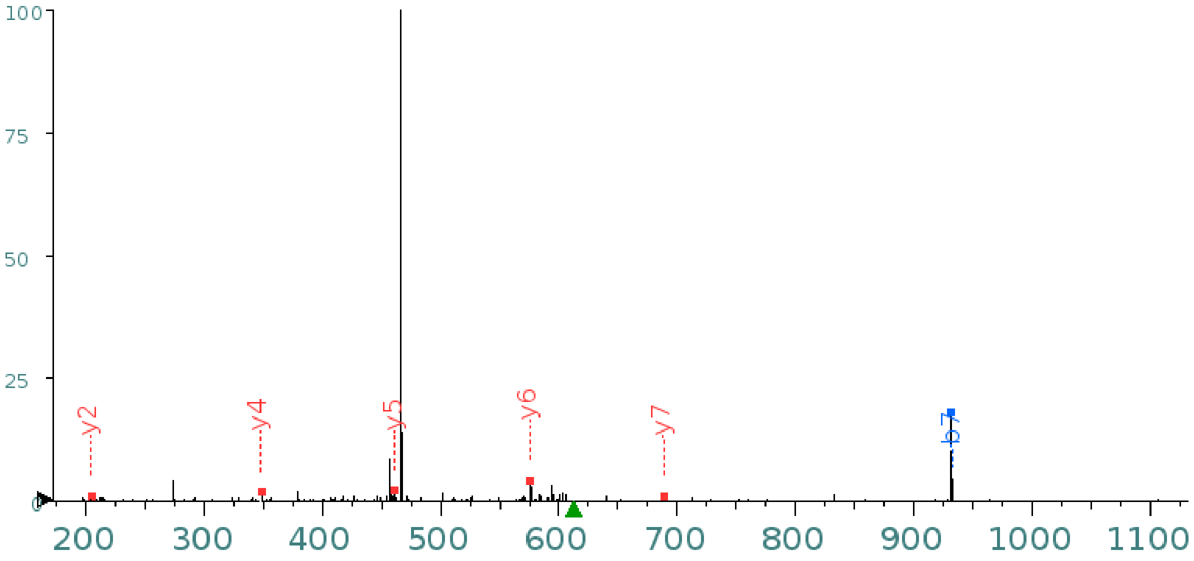

**Predicted Fragmentation Pattern**

| Seq | # | b: Δ Error | b | y | y: Δ Error | +1 |
|---|---|---|---|---|---|---|
| V | 1 | --- | 100.076 | --- | --- | 10 |
| G | 2 | --- | 157.097 | 1123.548 | --- | 9 |
| S@ | 3 | --- | 535.225 | 1066.526 | --- | 8 |
| I | 4 | --- | 648.309 | **688.399** | 202.890 | 7 |
| N | 5 | --- | 762.352 | **575.315** | 13.272 | 6 |
| I | 6 | --- | 875.436 | **461.272** | 202.729 | 5 |
| G | 7 | -935.096 | **932.457** | **348.188** | 477.576 | 4 |
| S | 8 | --- | 1019.489 | 291.166 | --- | 3 |
| G | 9 | --- | 1076.511 | **204.134** | 3511.330 | 2 |
| K | 10 | --- | --- | 147.113 | --- | 1 |

| | | | | | | |
|---|---|---|---|---|---|---|
| **Run:** | 77660 | **Scans:** | 9910 - 9910 | **m/z Range:** | 296.70062 | to 976.02875 |
| **Search:** | 77660 | **Precursor Mass:** | 679.32813 | | | |
| **Mods:** | C=160.03 *+15.99 #+220.06 @+291.1 | **Charge:** | +2 | | | |

**Sequence**

IASQTQFNT#K

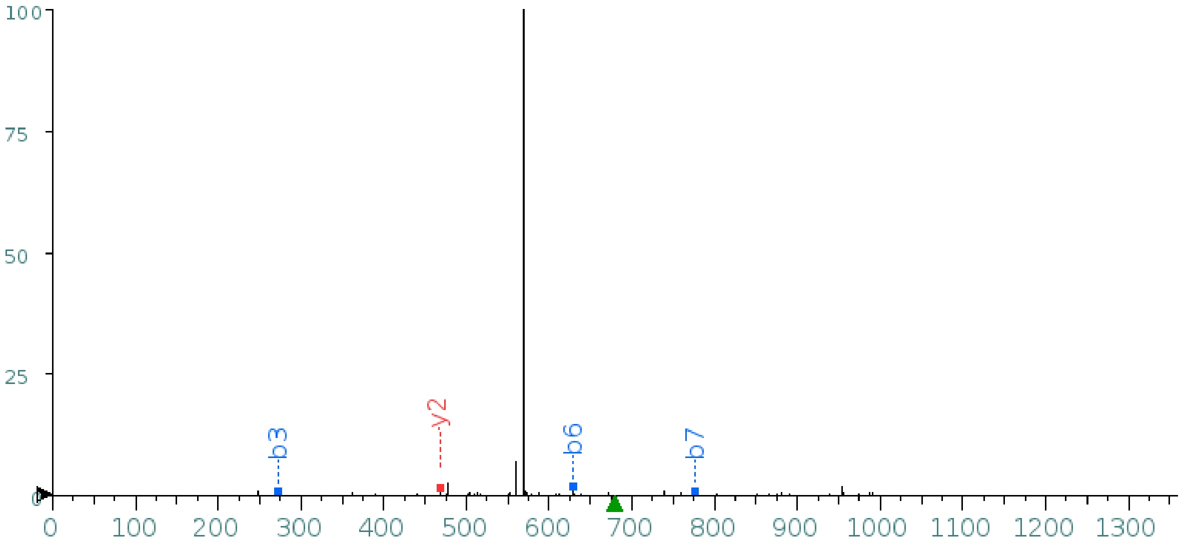

**Predicted Fragmentation Pattern**

| Seq | # | b: Δ Error | b | y | y: Δ Error | +1 |
|---|---|---|---|---|---|---|
| I | 1 | --- | 114.091 | --- | --- | 10 |
| A | 2 | --- | 185.128 | 1244.564 | --- | 9 |
| S | 3 | 385.729 | **272.160** | 1173.527 | --- | 8 |
| Q | 4 | --- | 400.219 | 1086.495 | --- | 7 |
| T | 5 | --- | 501.267 | 958.436 | --- | 6 |
| Q | 6 | -29.773 | **629.325** | 857.389 | --- | 5 |
| F | 7 | 324.815 | **776.394** | 729.330 | --- | 4 |
| N | 8 | --- | 890.437 | 582.262 | --- | 3 |
| T# | 9 | --- | 1211.543 | **468.219** | 200.507 | 2 |
| K | 10 | --- | --- | 147.113 | --- | 1 |

| | | | | | | | |
|---|---|---|---|---|---|---|---|
| **Run:** | 77660 | **Scans:** | 10033 - 10033 | **m/z Range:** | 223.32788 | to | 1033.6852 |
| **Search:** | 77660 | **Precursor Mass:** | 679.32807 | | | | |
| **Mods:** | C=160.03 *+15.99 #+220.06 @+291.1 | **Charge:** | +2 | | | | |

**Sequence**

IAS#QTQFNTK

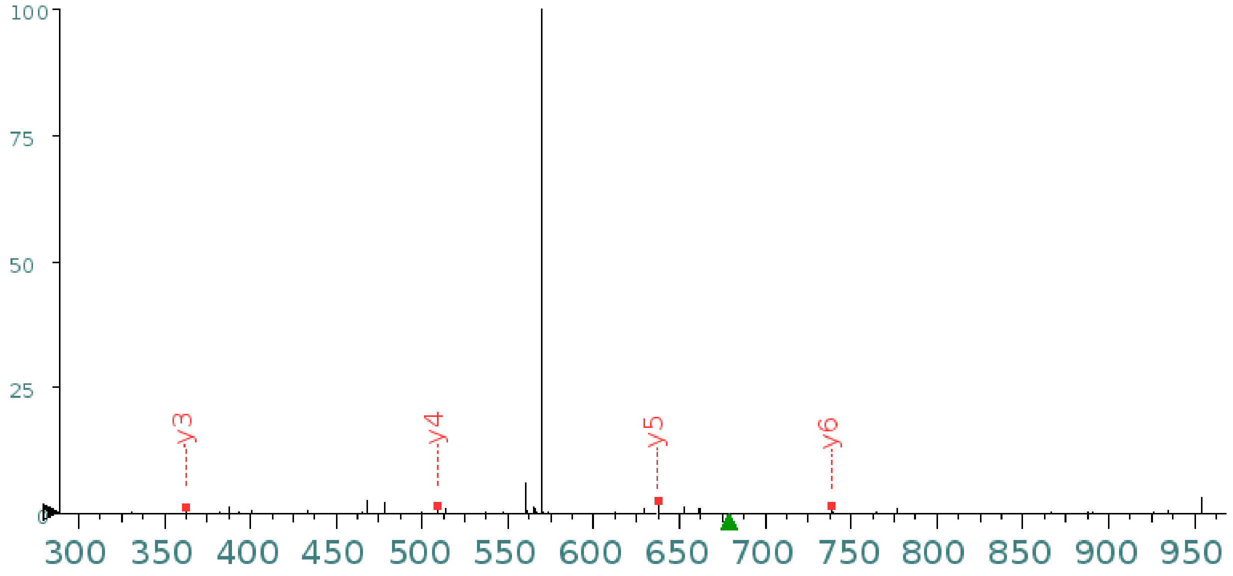

**Predicted Fragmentation Pattern**

| Seq | # | b: Δ Error | b | y | y: Δ Error | +1 |
|---|---|---|---|---|---|---|
| I | 1 | --- | 114.091 | --- | --- | 10 |
| A | 2 | --- | 185.128 | 1244.564 | --- | 9 |
| S# | 3 | --- | 492.219 | 1173.527 | --- | 8 |
| Q | 4 | --- | 620.277 | 866.437 | --- | 7 |
| T | 5 | --- | 721.325 | **738.378** | 386.914 | 6 |
| Q | 6 | --- | 849.384 | **637.330** | 1314.939 | 5 |
| F | 7 | --- | 996.452 | **509.272** | 261.252 | 4 |
| N | 8 | --- | 1110.495 | **362.203** | -100.894 | 3 |
| T | 9 | --- | 1211.543 | 248.160 | --- | 2 |
| K | 10 | --- | --- | 147.113 | --- | 1 |

| | | | |
|---|---|---|---|
| **Run:** | 77660 | **Scans:** | 10358 - 10358 |
| **Search:** | 77660 | **Precursor Mass:** | 714.84631 |
| **Mods:** | C=160.03 *+15.99 #+220.06 @+291.1 | **Charge:** | +2 |

**m/z Range:** 187.1082 to 1224.6843

**Sequence**

IAS@QTQFNTK

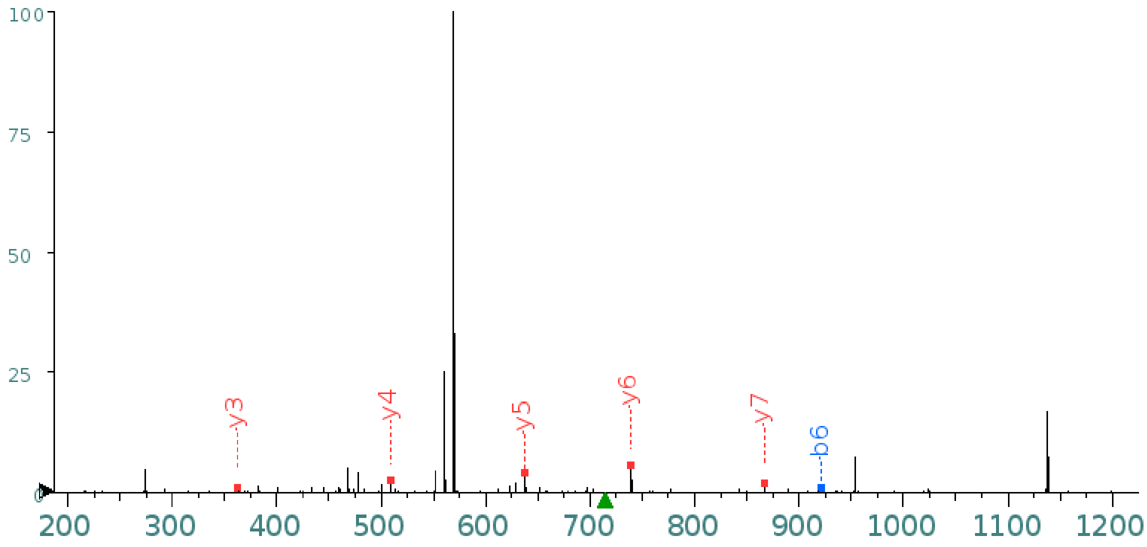

**Predicted Fragmentation Pattern**

| Seq | # | b: Δ Error | b | y | y: Δ Error | +1 |
|---|---|---|---|---|---|---|
| I | 1 | --- | 114.091 | --- | --- | 10 |
| A | 2 | --- | 185.128 | 1315.601 | --- | 9 |
| S@ | 3 | --- | 563.256 | 1244.564 | --- | 8 |
| Q | 4 | --- | 691.314 | **866.437** | 13.159 | 7 |
| T | 5 | --- | 792.362 | **738.378** | -1.196 | 6 |
| Q | 6 | 59.788 | **920.421** | **637.330** | 243.351 | 5 |
| F | 7 | --- | 1067.489 | **509.272** | 157.267 | 4 |
| N | 8 | --- | 1181.532 | **362.203** | 39.653 | 3 |
| T | 9 | --- | 1282.580 | 248.160 | --- | 2 |
| K | 10 | --- | --- | 147.113 | --- | 1 |

| Run: | 77660 | Scans: | 11631 - 11631 | m/z Range: | 195.3013 to 1397.0189[illegible] |
|---|---|---|---|---|---|
| Search: | 77660 | Precursor Mass: | 721.84043 | | |
| Mods: | C=160.03 *+15.99 #+220.06 @+291.1 | Charge: | +2 | | |

### Sequence

VGS@INIGS#GK

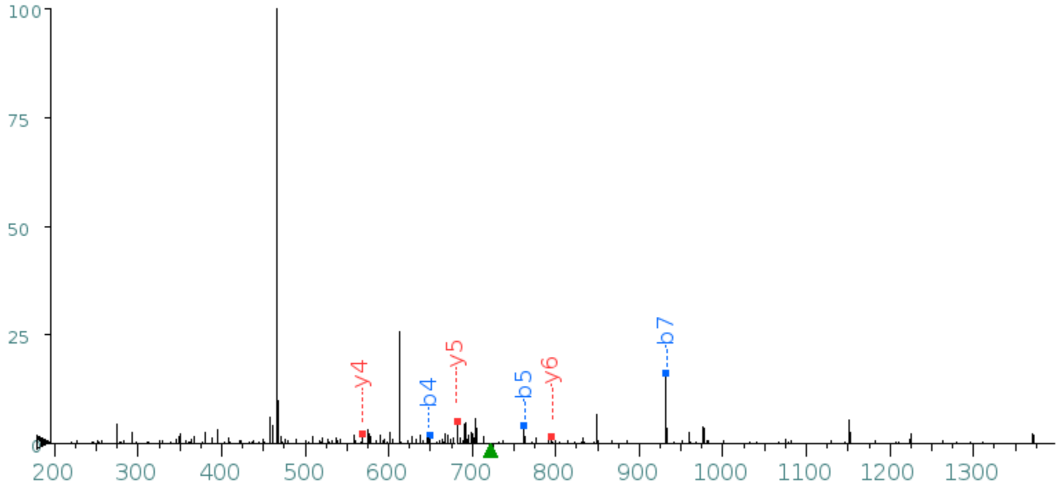

### Predicted Fragmentation Pattern

| Seq | # | b: Δ Error | b | y | y: Δ Error | +1 |
|---|---|---|---|---|---|---|
| V | 1 | --- | 100.076 | --- | --- | 10 |
| G | 2 | --- | 157.097 | 1343.606 | --- | 9 |
| S@ | 3 | --- | 535.225 | 1286.585 | --- | 8 |
| I | 4 | 751.167 | **648.309** | 908.457 | --- | 7 |
| N | 5 | -1272.959 | **762.352** | **795.373** | -1187.243 | 6 |
| I | 6 | --- | 875.436 | **681.330** | 693.356 | 5 |
| G | 7 | -793.139 | **932.457** | **568.246** | -791.102 | 4 |
| S# | 8 | --- | 1239.547 | 511.225 | --- | 3 |
| G | 9 | --- | 1296.569 | 204.134 | --- | 2 |
| K | 10 | --- | --- | 147.113 | --- | 1 |

| Run: | 77660 | Scans: | 11712 - 11712 | m/z Range: | 160.89908 to 1049.5668[illegible] |
|---|---|---|---|---|---|
| Search: | 77660 | Precursor Mass: | 576.29316 | | |
| Mods: | C=160.03 *+15.99 #+220.06 @+291.1 | Charge: | +2 | | |

**Sequence**

VGS#INIGSGK

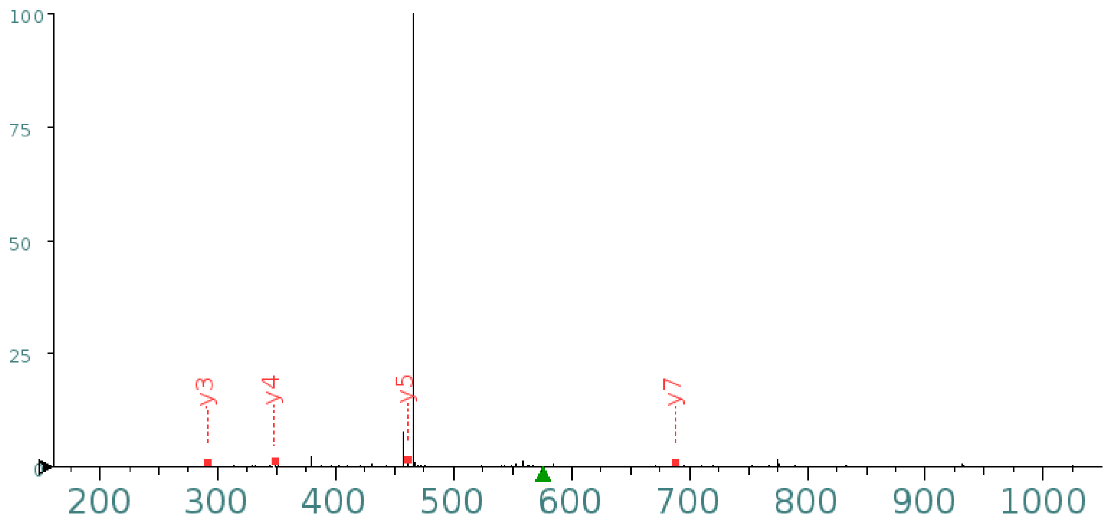

**Predicted Fragmentation Pattern**

| Seq | # | b: Δ Error | b | y | y: Δ Error | +1 |
|---|---|---|---|---|---|---|
| V | 1 | --- | 100.076 | --- | --- | 10 |
| G | 2 | --- | 157.097 | 1052.511 | --- | 9 |
| S# | 3 | --- | 464.187 | 995.489 | --- | 8 |
| I | 4 | --- | 577.272 | **688.399** | -99.241 | 7 |
| N | 5 | --- | 691.314 | 575.315 | --- | 6 |
| I | 6 | --- | 804.399 | **461.272** | 101.271 | 5 |
| G | 7 | --- | 861.420 | **348.188** | 264.752 | 4 |
| S | 8 | --- | 948.452 | **291.166** | 429.625 | 3 |
| G | 9 | --- | 1005.473 | 204.134 | --- | 2 |
| K | 10 | --- | --- | 147.113 | --- | 1 |

| | | | | | |
|---|---|---|---|---|---|
| **Run:** | 77660 | **Scans:** | 11937 - 11937 | **m/z Range:** | 154.14447 to 958.42426 |
| **Search:** | 77660 | **Precursor Mass:** | 611.8118 | | |
| **Mods:** | C=160.03 *+15.99 #+220.06 @+291.1 | **Charge:** | +2 | | |

**Adjusted Sequence**

VGS@INIGSGK

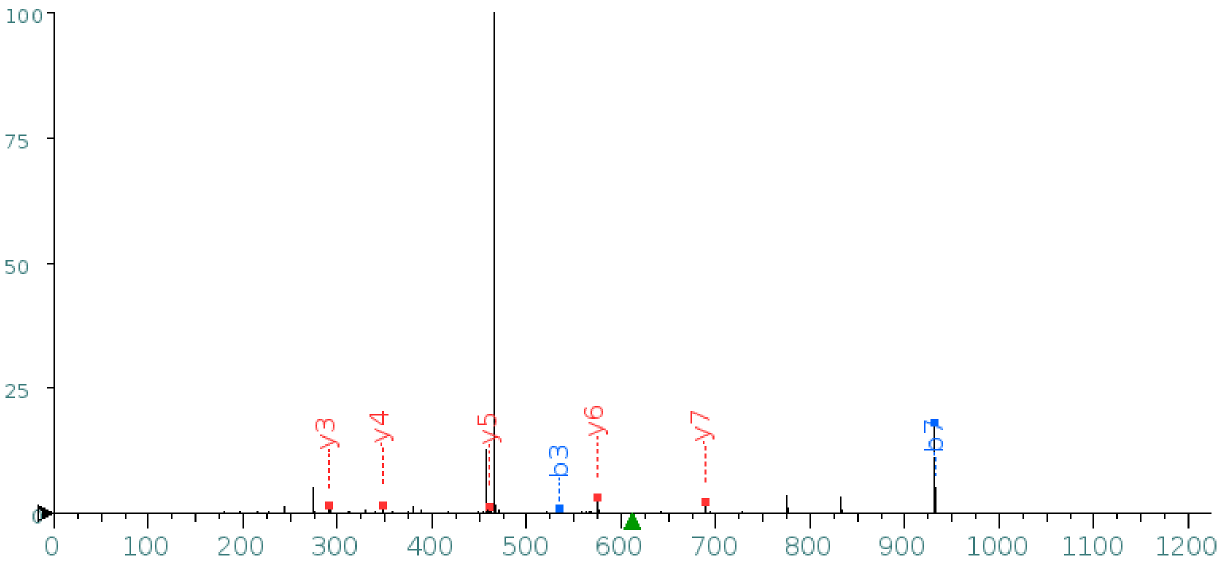

**Predicted Fragmentation Pattern**

| Seq | # | b: Δ Error | b | y | y: Δ Error | +1 |
|---|---|---|---|---|---|---|
| V | 1 | --- | 100.076 | --- | --- | 10 |
| G | 2 | --- | 157.097 | 1123.548 | --- | 9 |
| S@ | 3 | -1607.402 | **535.225** | 1066.526 | --- | 8 |
| I | 4 | --- | 648.309 | **688.399** | 52.734 | 7 |
| N | 5 | --- | 762.352 | **575.315** | 183.938 | 6 |
| I | 6 | --- | 875.436 | **461.272** | 137.121 | 5 |
| G | 7 | -938.506 | **932.457** | **348.188** | 123.170 | 4 |
| S | 8 | --- | 1019.489 | **291.166** | 282.784 | 3 |
| G | 9 | --- | 1076.511 | 204.134 | --- | 2 |
| K | 10 | --- | --- | 147.113 | --- | 1 |

| Run: | 77660 | Scans: | 12042 - 12042 | m/z Range: | 191.06302 to 1416.8439 |
|---|---|---|---|---|---|
| Search: | 77660 | Precursor Mass: | 714.84705 | | |
| Mods: | C=160.03 *+15.99 #+220.06 @+291.1 | Charge: | +2 | | |

**Sequence**

IAS@QTQFNTK

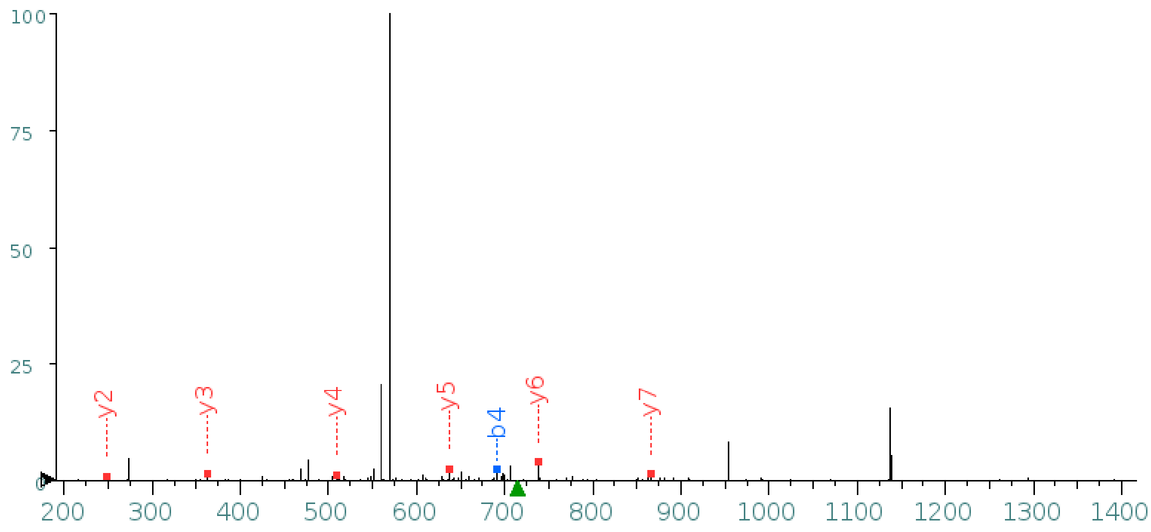

**Predicted Fragmentation Pattern**

| Seq | # | b: Δ Error | b | y | y: Δ Error | +1 |
|---|---|---|---|---|---|---|
| I | 1 | --- | 114.091 | --- | --- | 10 |
| A | 2 | --- | 185.128 | 1315.601 | --- | 9 |
| S@ | 3 | --- | 563.256 | 1244.564 | --- | 8 |
| Q | 4 | 780.265 | **691.314** | **866.437** | 185.571 | 7 |
| T | 5 | --- | 792.362 | **738.378** | 129.888 | 6 |
| Q | 6 | --- | 920.421 | **637.330** | 539.805 | 5 |
| F | 7 | --- | 1067.489 | **509.272** | 104.907 | 4 |
| N | 8 | --- | 1181.532 | **362.203** | 497.924 | 3 |
| T | 9 | --- | 1282.580 | **248.160** | 100.558 | 2 |
| K | 10 | --- | --- | 147.113 | --- | 1 |

| Run: | 77660 | Scans: | 12056 - 12056 | m/z Range: | 159.15001 to 957.60083 |
|---|---|---|---|---|---|
| Search: | 77660 | Precursor Mass: | 611.81176 | | |
| Mods: | C=160.03 *+15.99 #+220.06 @+291.1 | Charge: | +2 | | |

**Sequence**

VGS@INIGSGK

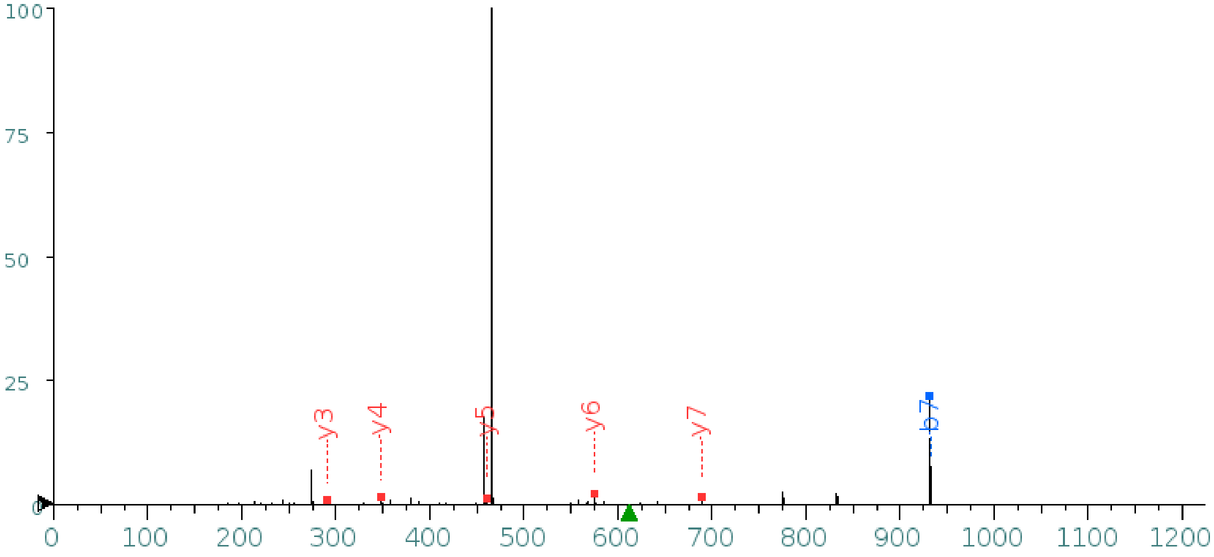

**Predicted Fragmentation Pattern**

| Seq | # | b: Δ Error | b | y | y: Δ Error | +1 |
|---|---|---|---|---|---|---|
| V | 1 | --- | 100.076 | --- | --- | 10 |
| G | 2 | --- | 157.097 | 1123.548 | --- | 9 |
| S@ | 3 | --- | 535.225 | 1066.526 | --- | 8 |
| I | 4 | --- | 648.309 | **688.399** | 150.420 | 7 |
| N | 5 | --- | 762.352 | **575.315** | 158.379 | 6 |
| I | 6 | --- | 875.436 | **461.272** | 232.752 | 5 |
| G | 7 | -955.164 | **932.457** | **348.188** | 350.768 | 4 |
| S | 8 | --- | 1019.489 | **291.166** | 148.474 | 3 |
| G | 9 | --- | 1076.511 | 204.134 | --- | 2 |
| K | 10 | --- | --- | 147.113 | --- | 1 |

| | | | | | |
|---|---|---|---|---|---|
| **Run:** | 77660 | **Scans:** | 12153 - 12153 | **m/z Range:** | 201.16187 to 1248.8261 |
| **Search:** | 77660 | **Precursor Mass:** | 757.35875 | | |
| **Mods:** | C=160.03 *+15.99 #+220.06 @+291.1 | **Charge:** | +2 | | |

#### Adjusted Sequence

VGS@INIGS@GK

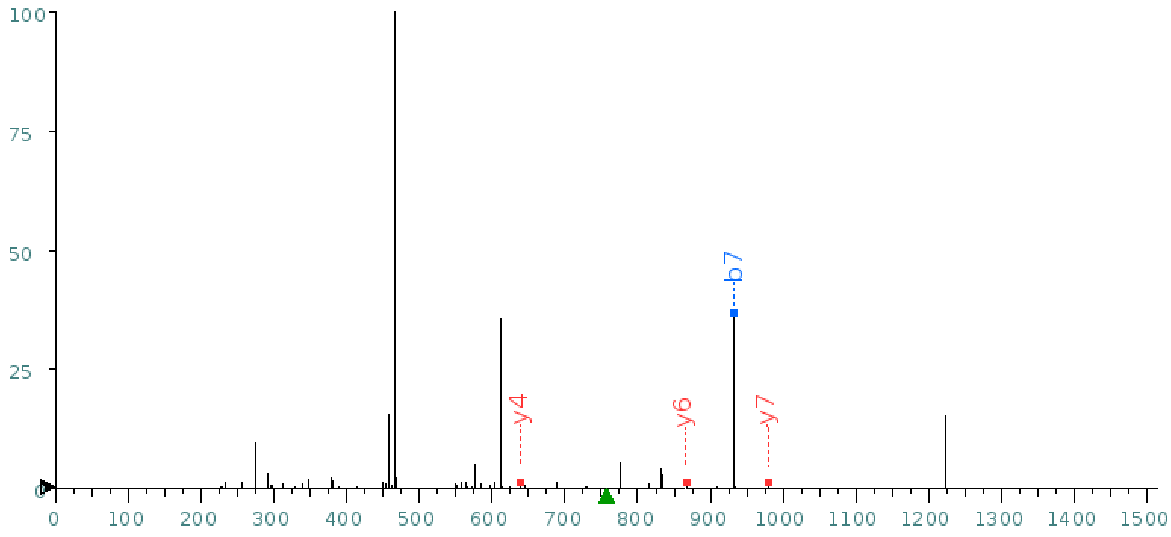

#### Predicted Fragmentation Pattern

| Seq | # | b: Δ Error | b | y | y: Δ Error | +1 |
|---|---|---|---|---|---|---|
| V | 1 | --- | 100.076 | --- | --- | 10 |
| G | 2 | --- | 157.097 | 1414.643 | --- | 9 |
| S@ | 3 | --- | 535.225 | 1357.622 | --- | 8 |
| I | 4 | --- | 648.309 | **979.494** | 304.084 | 7 |
| N | 5 | --- | 762.352 | **866.410** | 330.054 | 6 |
| I | 6 | --- | 875.436 | 752.367 | --- | 5 |
| G | 7 | -807.955 | **932.457** | **639.283** | 375.014 | 4 |
| S@ | 8 | --- | 1310.585 | 582.262 | --- | 3 |
| G | 9 | --- | 1367.606 | 204.134 | --- | 2 |
| K | 10 | --- | --- | 147.113 | --- | 1 |

| | | | | | |
|---|---|---|---|---|---|
| **Run:** | 77660 | **Scans:** | 12176 - 12176 | **m/z Range:** | 153.14464 to 1091.7565 |
| **Search:** | 77660 | **Precursor Mass:** | 611.81169 | | |
| **Mods:** | C=160.03 *+15.99 #+220.06 @+291.1 | **Charge:** | +2 | | |

**Sequence**

VGS@INIGSGK

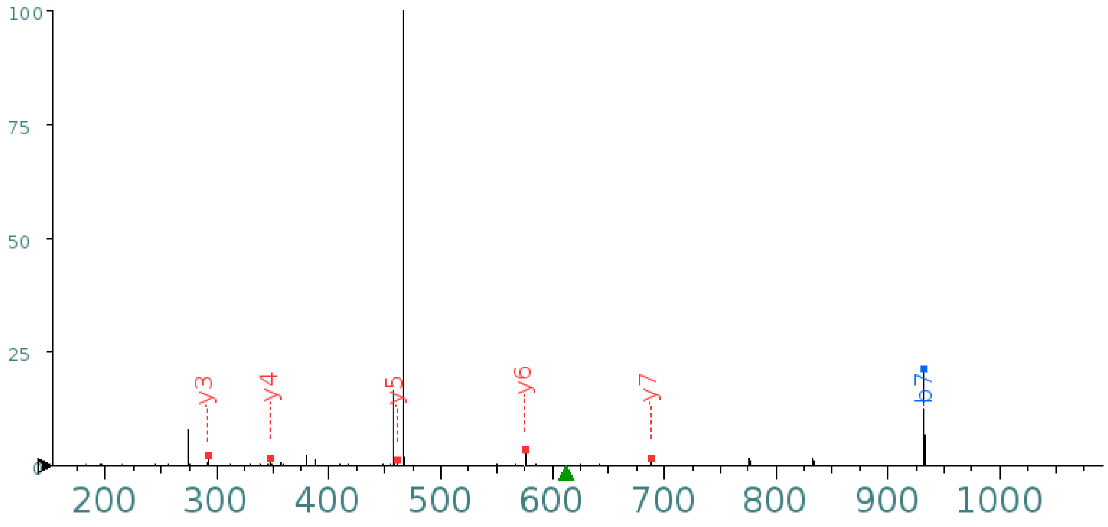

**Predicted Fragmentation Pattern**

| Seq | # | b: Δ Error | b | y | y: Δ Error | +1 |
|---|---|---|---|---|---|---|
| V | 1 | --- | 100.076 | --- | --- | 10 |
| G | 2 | --- | 157.097 | 1123.548 | --- | 9 |
| S@ | 3 | --- | 535.225 | 1066.526 | --- | 8 |
| I | 4 | --- | 648.309 | **688.399** | 183.037 | 7 |
| N | 5 | --- | 762.352 | **575.315** | 174.711 | 6 |
| I | 6 | --- | 875.436 | **461.272** | 370.281 | 5 |
| G | 7 | -937.260 | **932.457** | **348.188** | 269.658 | 4 |
| S | 8 | --- | 1019.489 | **291.166** | 3343.880 | 3 |
| G | 9 | --- | 1076.511 | 204.134 | --- | 2 |
| K | 10 | --- | --- | 147.113 | --- | 1 |

| Run: | 77660 | Scans: | 12289 - 12289 | m/z Range: | 148.19473 to 1091.7951 |
|---|---|---|---|---|---|
| Search: | 77660 | Precursor Mass: | 611.81151 | | |
| Mods: | C=160.03 *+15.99 #+220.06 @+291.1 | Charge: | +2 | | |

**Sequence**

VGS@INIGSGK

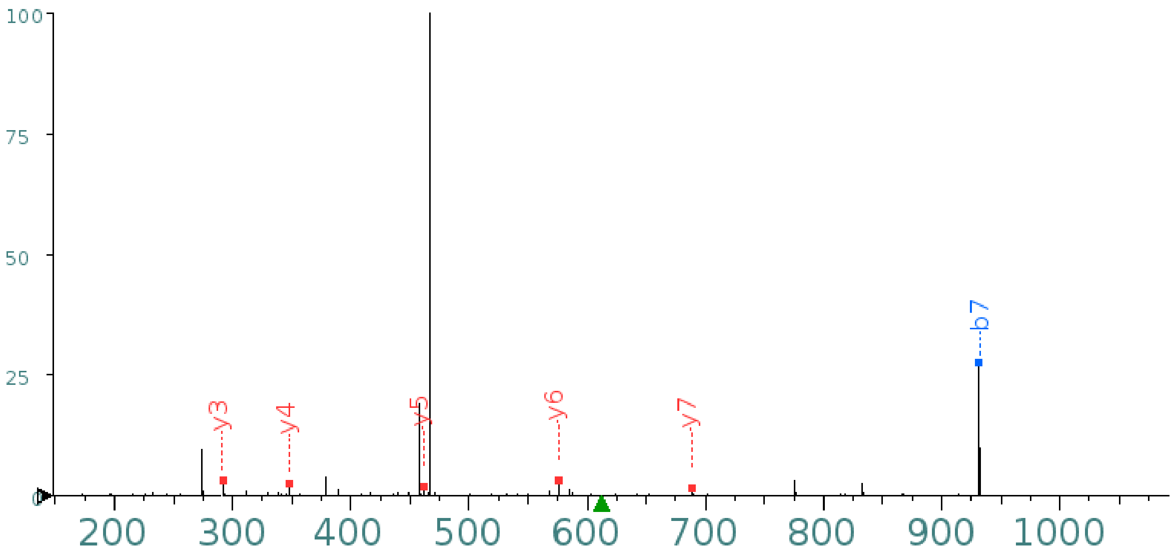

**Predicted Fragmentation Pattern**

| Seq | # | b: Δ Error | b | y | y: Δ Error | +1 |
|---|---|---|---|---|---|---|
| V | 1 | --- | 100.076 | --- | --- | 10 |
| G | 2 | --- | 157.097 | 1123.548 | --- | 9 |
| S@ | 3 | --- | 535.225 | 1066.526 | --- | 8 |
| I | 4 | --- | 648.309 | **688.399** | **179.669** | 7 |
| N | 5 | --- | 762.352 | **575.315** | **136.424** | 6 |
| I | 6 | --- | 875.436 | **461.272** | **215.426** | 5 |
| G | 7 | **-928.407** | **932.457** | **348.188** | **241.450** | 4 |
| S | 8 | --- | 1019.489 | **291.166** | **3416.337** | 3 |
| G | 9 | --- | 1076.511 | 204.134 | --- | 2 |
| K | 10 | --- | --- | 147.113 | --- | 1 |

| Run: | 77660 | Scans: | 13427 - 13427 | m/z Range: | 189.13503 to 1341.7919 |
|---|---|---|---|---|---|
| Search: | 77660 | Precursor Mass: | 757.35983 | | |
| Mods: | C=160.03 *+15.99 #+220.06 @+291.1 | Charge: | +2 | | |

**Sequence**

VGS@INIGS@GK

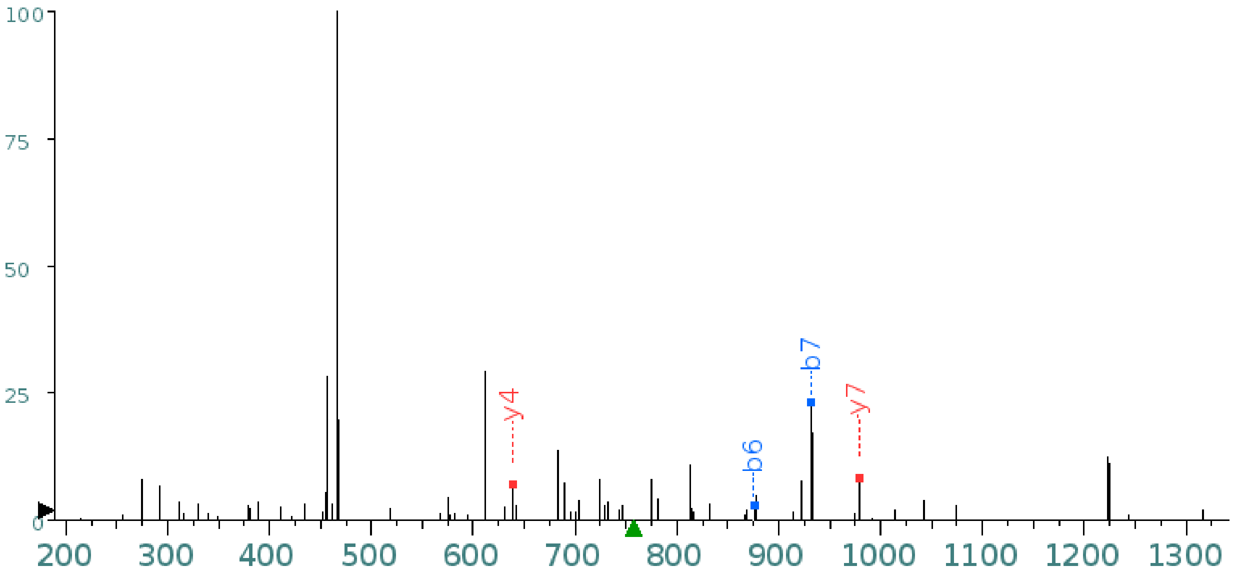

**Predicted Fragmentation Pattern**

| Seq | # | b: Δ Error | b | y | y: Δ Error | +1 |
|---|---|---|---|---|---|---|
| V | 1 | --- | 100.076 | --- | --- | 10 |
| G | 2 | --- | 157.097 | 1414.643 | --- | 9 |
| S@ | 3 | --- | 535.225 | 1357.622 | --- | 8 |
| I | 4 | --- | 648.309 | **979.494** | -970.003 | 7 |
| N | 5 | --- | 762.352 | 866.410 | --- | 6 |
| I | 6 | 155.116 | **875.436** | 752.367 | --- | 5 |
| G | 7 | -871.358 | **932.457** | **639.283** | 159.834 | 4 |
| S@ | 8 | --- | 1310.585 | 582.262 | --- | 3 |
| G | 9 | --- | 1367.606 | 204.134 | --- | 2 |
| K | 10 | --- | --- | 147.113 | --- | 1 |

| | | | | | |
|---|---|---|---|---|---|
| **Run:** | 77660 | **Scans:** | 13451 - 13451 | **m/z Range:** | 148.31651 to 1152.5868 |
| **Search:** | 77660 | **Precursor Mass:** | 576.29343 | | |
| **Mods:** | C=160.03 *+15.99 #+220.06 @+291.1 | **Charge:** | +2 | | |

**Sequence**

VGS#INIGSGK

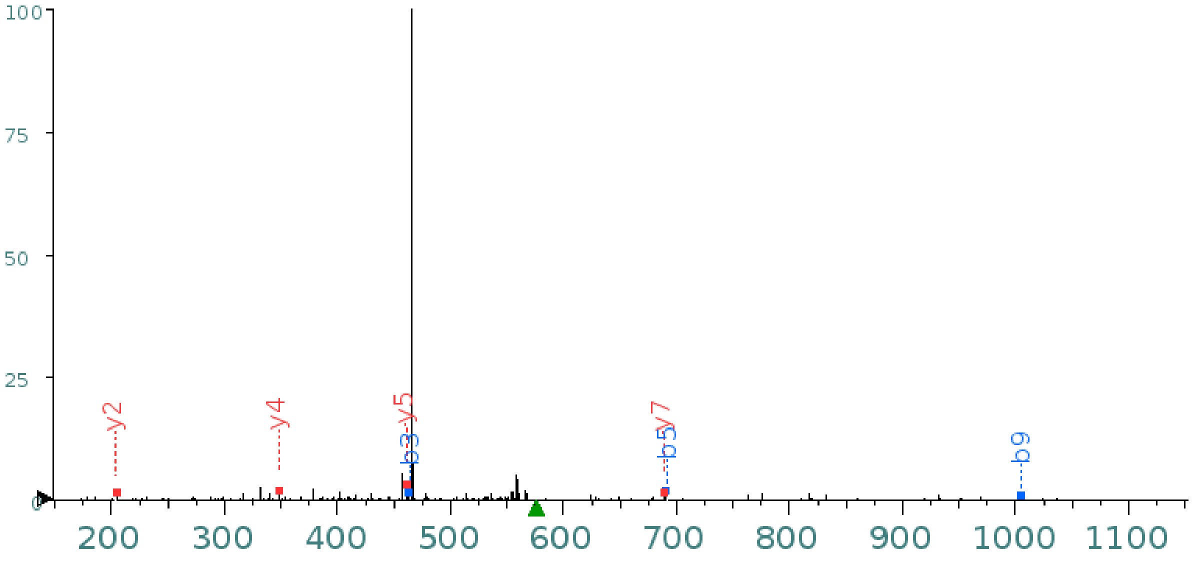

**Predicted Fragmentation Pattern**

| Seq | # | b: Δ Error | b | y | y: Δ Error | +1 |
|---|---|---|---|---|---|---|
| V | 1 | --- | 100.076 | --- | --- | 10 |
| G | 2 | --- | 157.097 | 1052.511 | --- | 9 |
| S# | 3 | -1888.799 | **464.187** | 995.489 | --- | 8 |
| I | 4 | --- | 577.272 | **688.399** | 175.681 | 7 |
| N | 5 | -698.350 | **691.314** | 575.315 | --- | 6 |
| I | 6 | --- | 804.399 | **461.272** | 234.604 | 5 |
| G | 7 | --- | 861.420 | **348.188** | 486.157 | 4 |
| S | 8 | --- | 948.452 | 291.166 | --- | 3 |
| G | 9 | -594.253 | **1005.473** | **204.134** | 806.307 | 2 |
| K | 10 | --- | --- | 147.113 | --- | 1 |

| Run: | 77660 | Scans: | 13595 - 13595 | m/z Range: | 148.08362 to 1100.5515 |
|---|---|---|---|---|---|
| Search: | 77660 | Precursor Mass: | 611.81222 | | |
| Mods: | C=160.03 *+15.99 #+220.06 @+291.1 | Charge: | +2 | | |

**Sequence**

VGSINIGS@GK

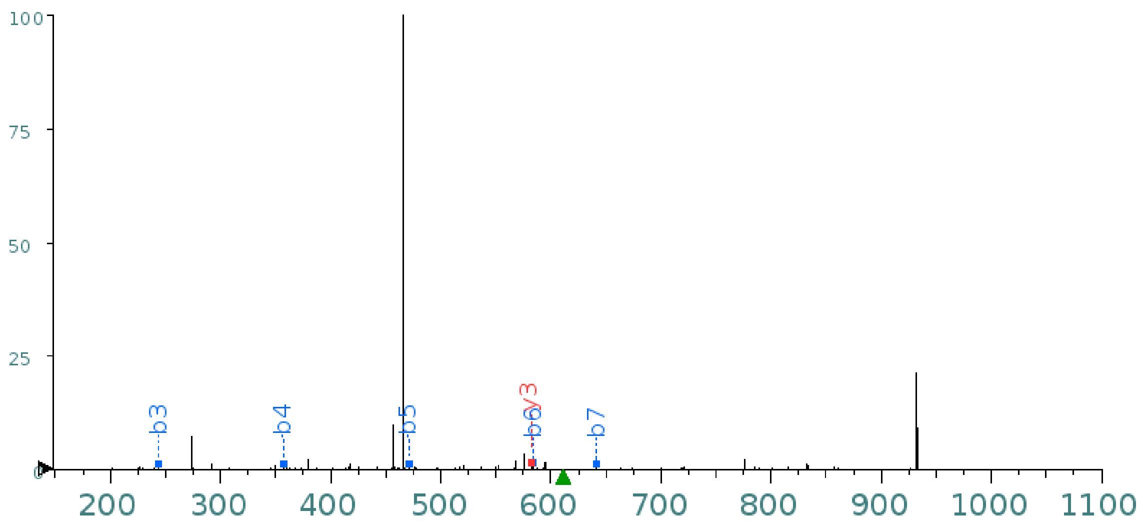

**Predicted Fragmentation Pattern**

| Seq | # | b: Δ Error | b | y | y: Δ Error | +1 |
|---|---|---|---|---|---|---|
| V | 1 | --- | 100.076 | --- | --- | 10 |
| G | 2 | --- | 157.097 | 1123.548 | --- | 9 |
| S | 3 | -39.258 | **244.129** | 1066.526 | --- | 8 |
| I | 4 | 585.412 | **357.213** | 979.494 | --- | 7 |
| N | 5 | 416.460 | **471.256** | 866.410 | --- | 6 |
| I | 6 | -54.887 | **584.340** | 752.367 | --- | 5 |
| G | 7 | 214.160 | **641.362** | 639.283 | --- | 4 |
| S@ | 8 | --- | 1019.489 | **582.262** | 489.189 | 3 |
| G | 9 | --- | 1076.511 | 204.134 | --- | 2 |
| K | 10 | --- | --- | 147.113 | --- | 1 |

| Run: | 77660 | Scans: | 13707 - 13707 | m/z Range: | 153.23021 to 1175.0046 |
|---|---|---|---|---|---|
| Search: | 77660 | Precursor Mass: | 611.81209 | | |
| Mods: | C=160.03 *+15.99 #+220.06 @+291.1 | Charge: | +2 | | |

**Sequence**

VGS@INIGSGK

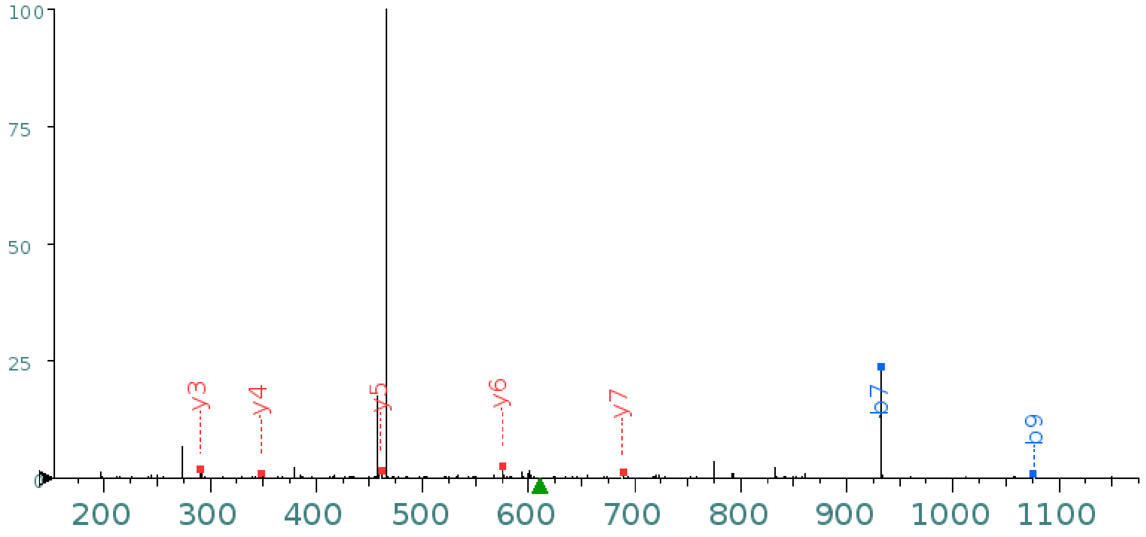

**Predicted Fragmentation Pattern**

| Seq | # | b: Δ Error | b | y | y: Δ Error | +1 |
|---|---|---|---|---|---|---|
| V | 1 | --- | 100.076 | --- | --- | 10 |
| G | 2 | --- | 157.097 | 1123.548 | --- | 9 |
| S@ | 3 | --- | 535.225 | 1066.526 | --- | 8 |
| I | 4 | --- | 648.309 | **688.399** | 454.349 | 7 |
| N | 5 | --- | 762.352 | **575.315** | 168.772 | 6 |
| I | 6 | --- | 875.436 | **461.272** | 383.503 | 5 |
| G | 7 | -864.145 | **932.457** | **348.188** | 1243.692 | 4 |
| S | 8 | --- | 1019.489 | **291.166** | 315.885 | 3 |
| G | 9 | -523.015 | **1076.511** | 204.134 | --- | 2 |
| K | 10 | --- | --- | 147.113 | --- | 1 |

| Run: | 77660 | Scans: | 13775 - 13775 | m/z Range: | 189.28705 to 1392.9510 |
|---|---|---|---|---|---|
| Search: | 77660 | Precursor Mass: | 757.35965 | | |
| Mods: | C=160.03 *+15.99 #+220.06 @+291.1 | Charge: | +2 | | |

**Sequence**

VGS@INIGS@GK

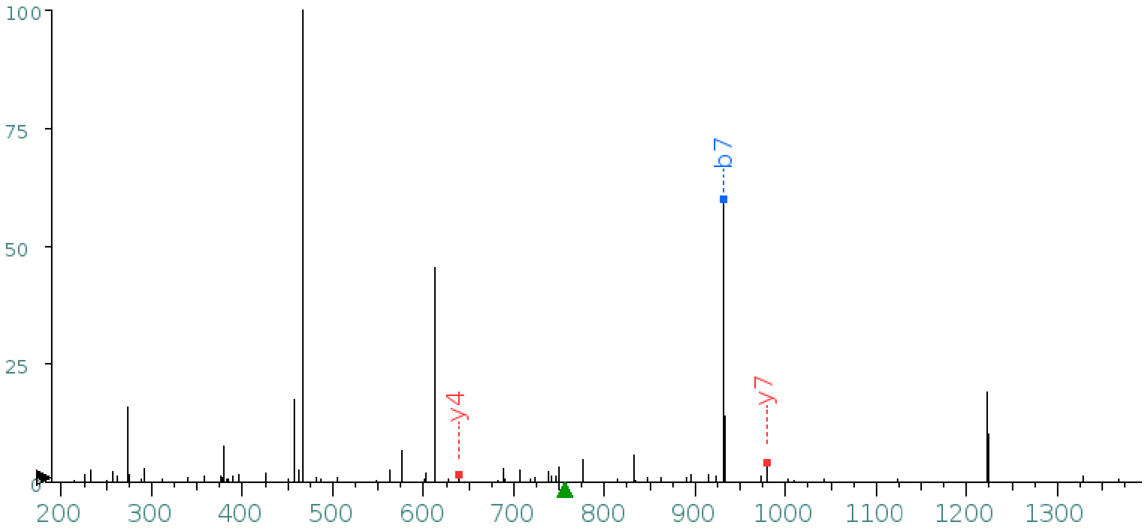

**Predicted Fragmentation Pattern**

| Seq | # | b: Δ Error | b | y | y: Δ Error | +1 |
|---|---|---|---|---|---|---|
| V | 1 | --- | 100.076 | --- | --- | 10 |
| G | 2 | --- | 157.097 | 1414.643 | --- | 9 |
| S@ | 3 | --- | 535.225 | 1357.622 | --- | 8 |
| I | 4 | --- | 648.309 | **979.494** | **137.284** | 7 |
| N | 5 | --- | 762.352 | 866.410 | --- | 6 |
| I | 6 | --- | 875.436 | 752.367 | --- | 5 |
| G | 7 | **-855.228** | **932.457** | **639.283** | **485.001** | 4 |
| S@ | 8 | --- | 1310.585 | 582.262 | --- | 3 |
| G | 9 | --- | 1367.606 | 204.134 | --- | 2 |
| K | 10 | --- | --- | 147.113 | --- | 1 |

| Run: | 77660 | Scans: | 13826 - 13826 | m/z Range: | 154.27045 to 1003.7132 |
|---|---|---|---|---|---|
| Search: | 77660 | Precursor Mass: | 611.81201 | | |
| Mods: | C=160.03 *+15.99 #+220.06 @+291.1 | Charge: | +2 | | |

Options
Show Options

(m/z)

Sequence

VGS@INIGSGK

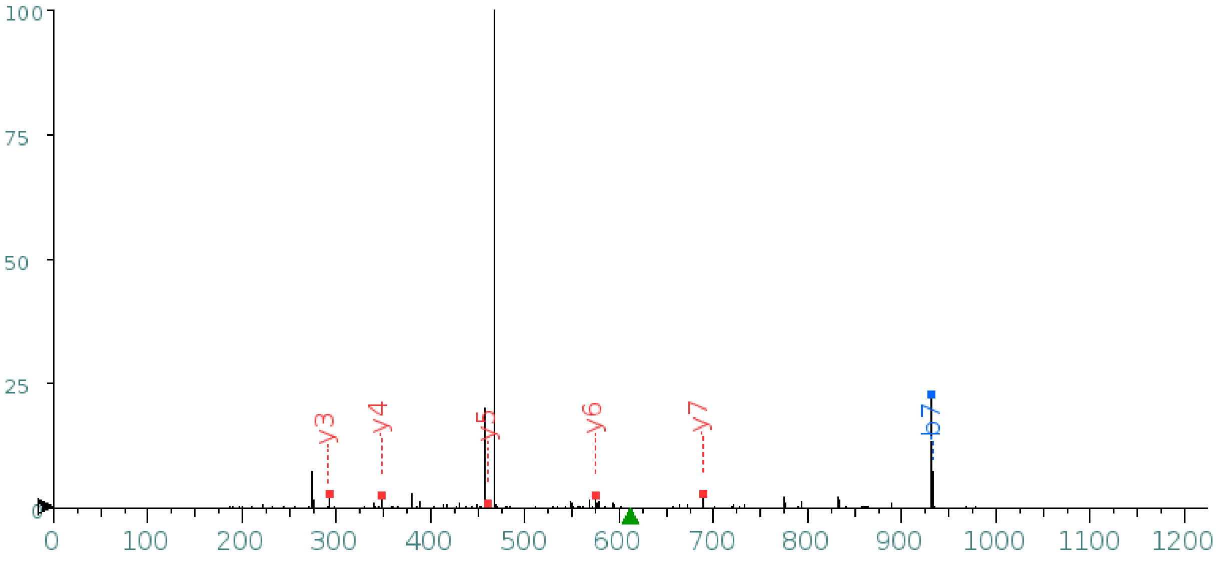

### Predicted Fragmentation Pattern

| Seq | # | b: Δ Error | b | y | y: Δ Error | +1 |
|---|---|---|---|---|---|---|
| V | 1 | --- | 100.076 | --- | --- | 10 |
| G | 2 | --- | 157.097 | 1123.548 | --- | 9 |
| S@ | 3 | --- | 535.225 | 1066.526 | --- | 8 |
| I | 4 | --- | 648.309 | **688.399** | -59.336 | 7 |
| N | 5 | --- | 762.352 | **575.315** | 458.110 | 6 |
| I | 6 | --- | 875.436 | **461.272** | -1994.126 | 5 |
| G | 7 | -947.163 | **932.457** | **348.188** | 497.715 | 4 |
| S | 8 | --- | 1019.489 | **291.166** | 3376.362 | 3 |
| G | 9 | --- | 1076.511 | 204.134 | --- | 2 |
| K | 10 | --- | --- | 147.113 | --- | 1 |

| | | | | | |
|---|---|---|---|---|---|
| **Run:** | 77660 | **Scans:** | 13882 - 13882 | **m/z Range:** | 189.13501 to 1282.9034 |
| **Search:** | 77660 | **Precursor Mass:** | 757.35965 | | |
| **Mods:** | C=160.03 *+15.99 #+220.06 @+291.1 | **Charge:** | +2 | | |

**Sequence**

VGS@INIGS@GK

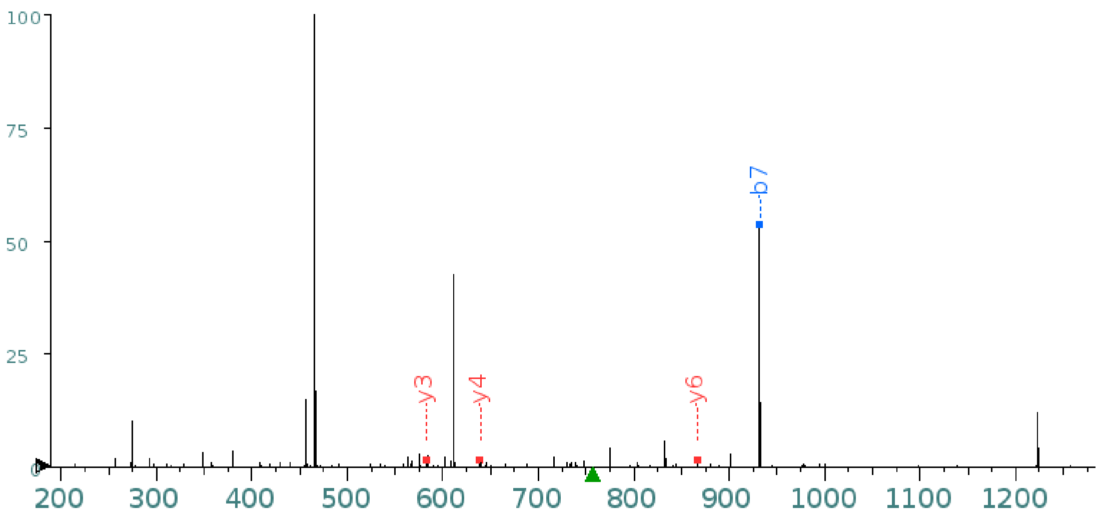

**Predicted Fragmentation Pattern**

| Seq | # | b: Δ Error | b | y | y: Δ Error | +1 |
|---|---|---|---|---|---|---|
| V | 1 | --- | 100.076 | --- | --- | 10 |
| G | 2 | --- | 157.097 | 1414.643 | --- | 9 |
| S@ | 3 | --- | 535.225 | 1357.622 | --- | 8 |
| I | 4 | --- | 648.309 | 979.494 | --- | 7 |
| N | 5 | --- | 762.352 | **866.410** | **77.115** | 6 |
| I | 6 | --- | 875.436 | 752.367 | --- | 5 |
| G | 7 | **-981.987** | **932.457** | **639.283** | **-940.889** | 4 |
| S@ | 8 | --- | 1310.585 | **582.262** | **500.813** | 3 |
| G | 9 | --- | 1367.606 | 204.134 | --- | 2 |
| K | 10 | --- | --- | 147.113 | --- | 1 |

| | | | | | |
|---|---|---|---|---|---|
| **Run:** | 77660 | **Scans:** | 14050 - 14050 | **m/z Range:** | 154.07898 to 1004.7475 |
| **Search:** | 77660 | **Precursor Mass:** | 611.8119 | | |
| **Mods:** | C=160.03 *+15.99 #+220.06 @+291.1 | | | | |

**Sequence**

VGS@INIGSGK

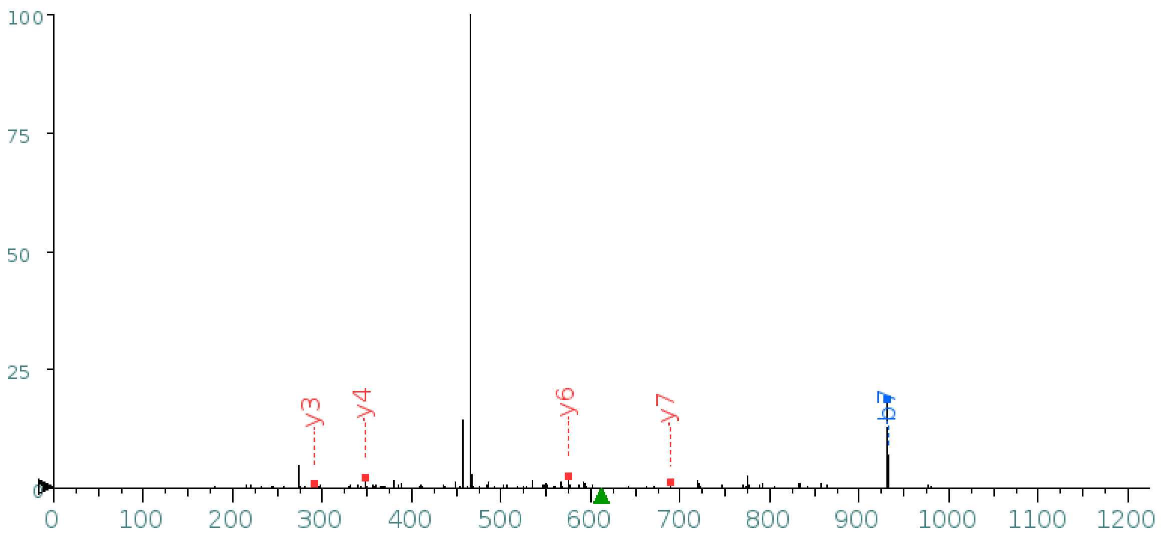

**Predicted Fragmentation Pattern**

| Seq | # | b: Δ Error | b | y | y: Δ Error | +1 |
|---|---|---|---|---|---|---|
| V | 1 | --- | 100.076 | --- | --- | 10 |
| G | 2 | --- | 157.097 | 1123.548 | --- | 9 |
| S@ | 3 | --- | 535.225 | 1066.526 | --- | 8 |
| I | 4 | --- | 648.309 | **688.399** | 283.533 | 7 |
| N | 5 | --- | 762.352 | **575.315** | 303.019 | 6 |
| I | 6 | --- | 875.436 | 461.272 | --- | 5 |
| G | 7 | -836.344 | **932.457** | **348.188** | 388.429 | 4 |
| S | 8 | --- | 1019.489 | **291.166** | 1326.911 | 3 |
| G | 9 | --- | 1076.511 | 204.134 | --- | 2 |
| K | 10 | --- | --- | 147.113 | --- | 1 |

| Run: | 77660 | Scans: | 14160 - 14160 | m/z Range: | 154.0507 to 1082.8726 |
|---|---|---|---|---|---|
| Search: | 77660 | Precursor Mass: | 611.81201 | | |
| Mods: | C=160.03 *+15.99 #+220.06 @+291.1 | Charge: | +2 | | |

**Sequence**

VGS@INIGSGK

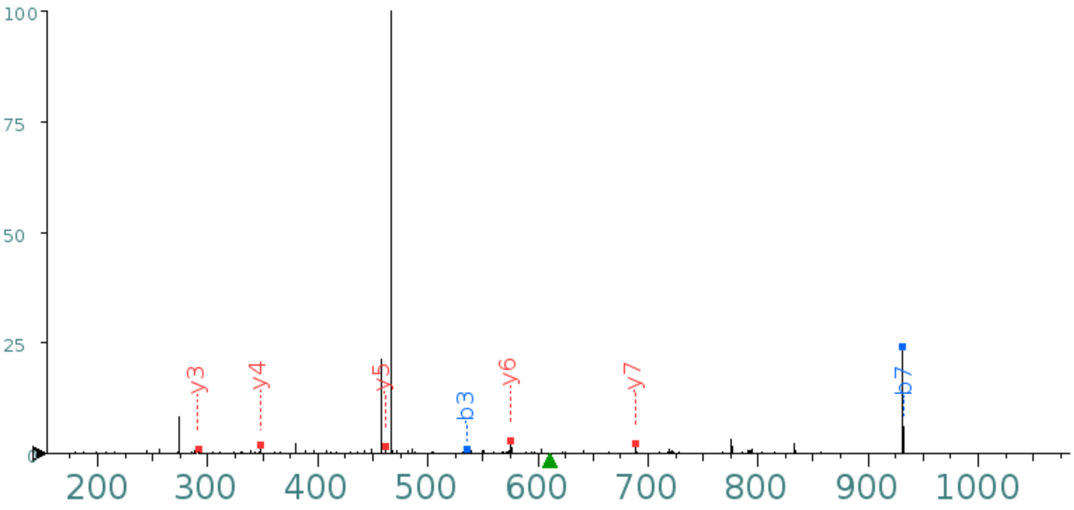

**Predicted Fragmentation Pattern**

| Seq | # | b: Δ Error | b | y | y: Δ Error | +1 |
|---|---|---|---|---|---|---|
| V | 1 | --- | 100.076 | --- | --- | 10 |
| G | 2 | --- | 157.097 | 1123.548 | --- | 9 |
| S@ | 3 | 304.854 | **535.225** | 1066.526 | --- | 8 |
| I | 4 | --- | 648.309 | **688.399** | 242.947 | 7 |
| N | 5 | --- | 762.352 | **575.315** | 264.318 | 6 |
| I | 6 | --- | 875.436 | **461.272** | 273.222 | 5 |
| G | 7 | -869.784 | **932.457** | **348.188** | 677.530 | 4 |
| S | 8 | --- | 1019.489 | **291.166** | 3157.902 | 3 |
| G | 9 | --- | 1076.511 | 204.134 | --- | 2 |
| K | 10 | --- | --- | 147.113 | --- | 1 |

| | | | | | |
|---|---|---|---|---|---|
| **Run:** | 77660 | **Scans:** | 14274 - 14274 | **m/z Range:** | 153.28543 to 1175.9201 |
| **Search:** | 77660 | **Precursor Mass:** | 611.81197 | | |
| **Mods:** | C=160.03 *+15.99 #+220.06 @+291.1 | **Charge:** | +2 | | |

**Sequence**

VGS@INIGSGK

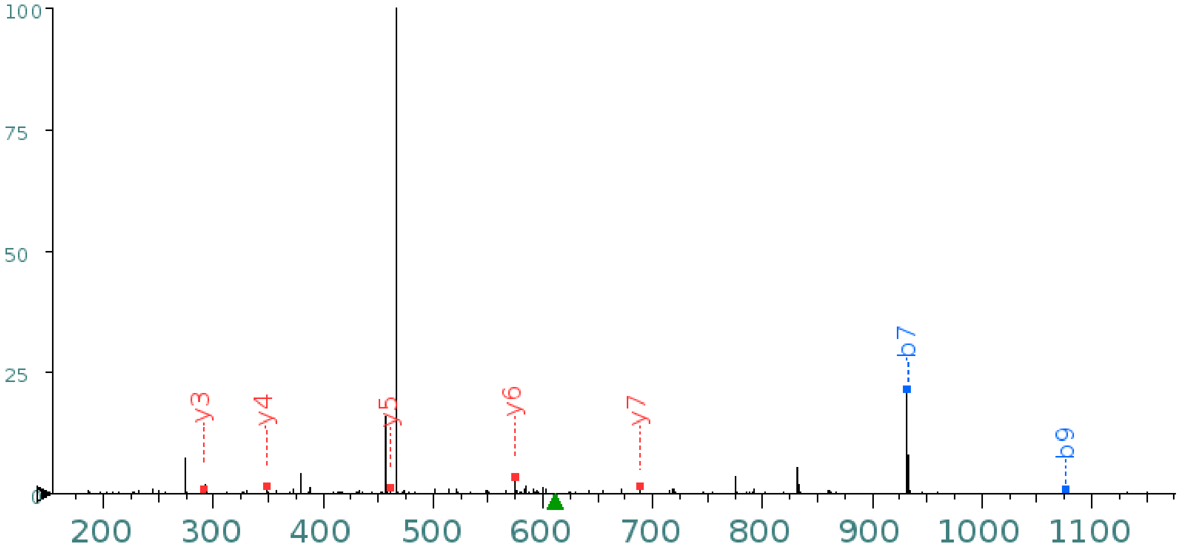

**Predicted Fragmentation Pattern**

| Seq | # | b: Δ Error | b | y | y: Δ Error | +1 |
|---|---|---|---|---|---|---|
| V | 1 | --- | 100.076 | --- | --- | 10 |
| G | 2 | --- | 157.097 | 1123.548 | --- | 9 |
| S@ | 3 | --- | 535.225 | 1066.526 | --- | 8 |
| I | 4 | --- | 648.309 | **688.399** | 406.158 | 7 |
| N | 5 | --- | 762.352 | **575.315** | -140.366 | 6 |
| I | 6 | --- | 875.436 | **461.272** | 84.602 | 5 |
| G | 7 | -944.999 | **932.457** | **348.188** | 129.128 | 4 |
| S | 8 | --- | 1019.489 | **291.166** | 121.230 | 3 |
| G | 9 | -771.784 | **1076.511** | 204.134 | --- | 2 |
| K | 10 | --- | --- | 147.113 | --- | 1 |

| Run: | 77660 | Scans: | 14381 - 14381 | m/z Range: | 154.24881 to 1028.6646 |
|---|---|---|---|---|---|
| Search: | 77660 | Precursor Mass: | 611.81196 | | |
| Mods: | C=160.03 *+15.99 #+220.06 @+291.1 | Charge: | +2 | | |

**Sequence**

VGS@INIGSGK

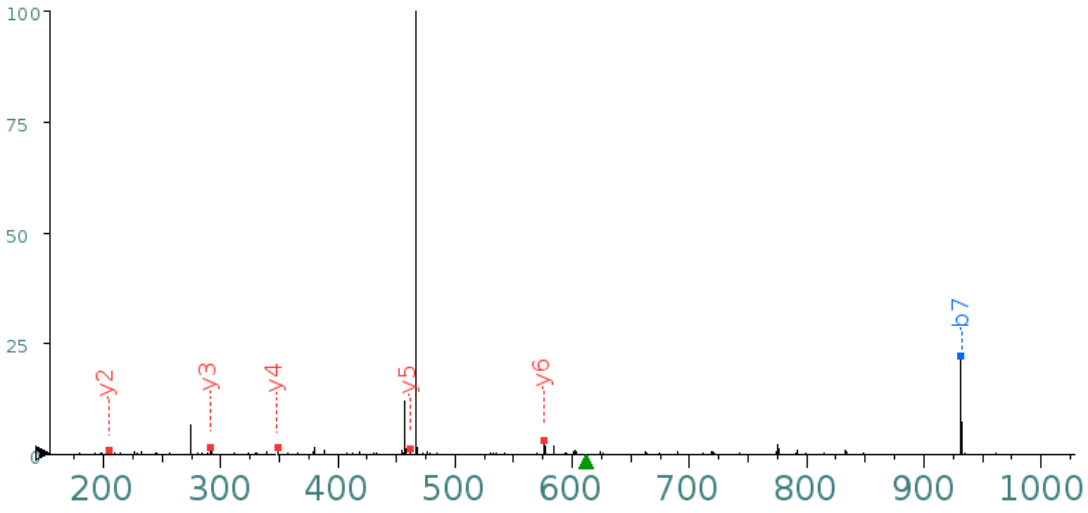

**Predicted Fragmentation Pattern**

| Seq | # | b: Δ Error | b | y | y: Δ Error | +1 |
|---|---|---|---|---|---|---|
| V | 1 | --- | 100.076 | --- | --- | 10 |
| G | 2 | --- | 157.097 | 1123.548 | --- | 9 |
| S@ | 3 | --- | 535.225 | 1066.526 | --- | 8 |
| I | 4 | --- | 648.309 | 688.399 | --- | 7 |
| N | 5 | --- | 762.352 | **575.315** | 52.735 | 6 |
| I | 6 | --- | 875.436 | **461.272** | 610.734 | 5 |
| G | 7 | -865.588 | **932.457** | **348.188** | 441.411 | 4 |
| S | 8 | --- | 1019.489 | **291.166** | 465.439 | 3 |
| G | 9 | --- | 1076.511 | **204.134** | 663.373 | 2 |
| K | 10 | --- | --- | 147.113 | --- | 1 |

| Run: | 77660 | Scans: | 14511 - 14511 | m/z Range: | 154.05838 | to | 1119.0396 |
|---|---|---|---|---|---|---|---|
| Search: | 77660 | Precursor Mass: | 611.8119 | | | | |
| Mods: | C=160.03 *+15.99 #+220.06 @+291.1 | Charge: | +2 | | | | |

### Sequence

VGS@INIGSGK

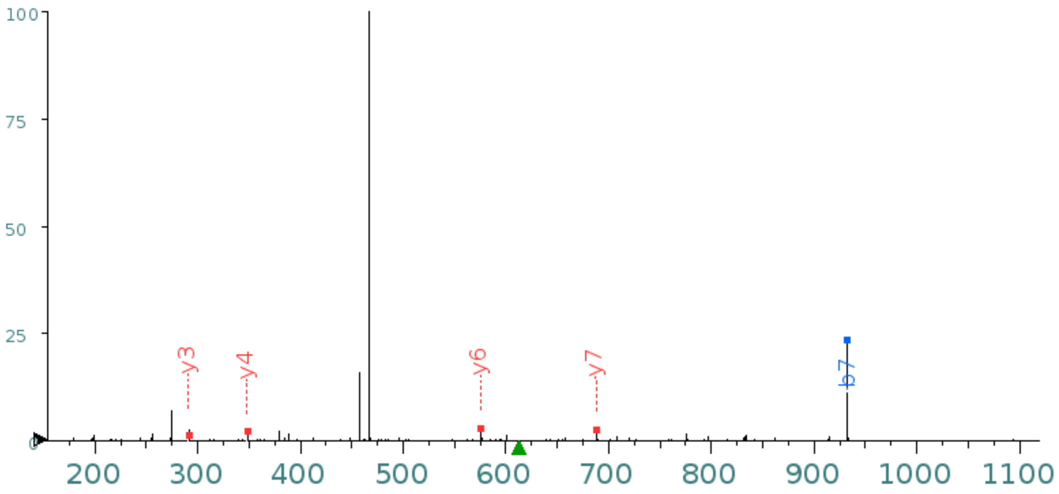

### Predicted Fragmentation Pattern

| Seq | # | b: Δ Error | b | y | y: Δ Error | +1 |
|---|---|---|---|---|---|---|
| V | 1 | --- | 100.076 | --- | --- | 10 |
| G | 2 | --- | 157.097 | 1123.548 | --- | 9 |
| S@ | 3 | --- | 535.225 | 1066.526 | --- | 8 |
| I | 4 | --- | 648.309 | **688.399** | 83.496 | 7 |
| N | 5 | --- | 762.352 | **575.315** | 393.662 | 6 |
| I | 6 | --- | 875.436 | 461.272 | --- | 5 |
| G | 7 | -915.358 | **932.457** | **348.188** | 413.301 | 4 |
| S | 8 | --- | 1019.489 | **291.166** | 403.654 | 3 |
| G | 9 | --- | 1076.511 | 204.134 | --- | 2 |
| K | 10 | --- | --- | 147.113 | --- | 1 |

| | | | |
|---|---|---|---|
| **Run:** | 77660 | **Scans:** | 14777 - 14777 |
| **Search:** | 77660 | **Precursor Mass:** | 611.81185 |
| **Mods:** | C=160.03 *+15.99 #+220.06 @+291.1 | **Charge:** | +2 |

**m/z Range:** 148.22731 **to** 1082.5311

**Sequence**

VGS@INIGSGK

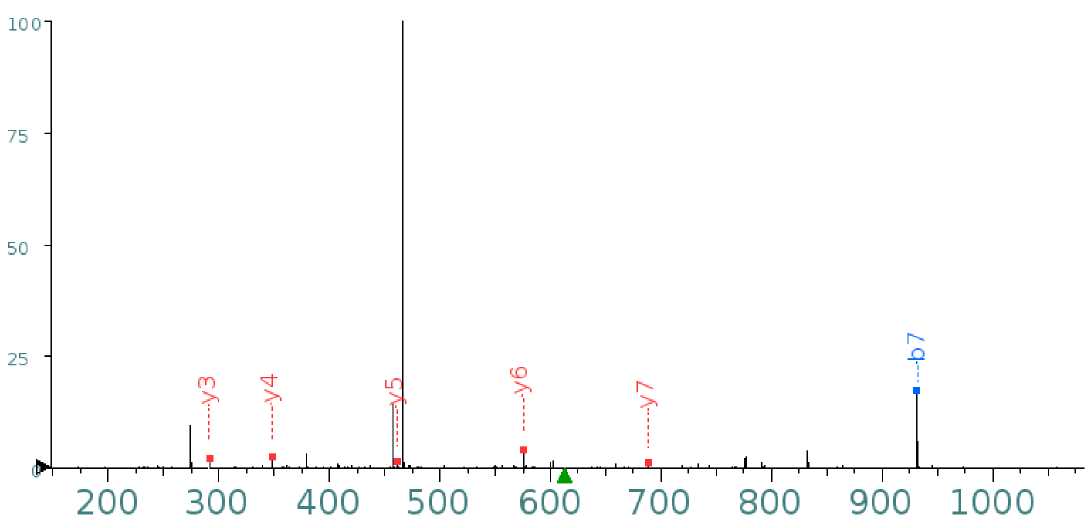

**Predicted Fragmentation Pattern**

| Seq | # | b: Δ Error | b | y | y: Δ Error | +1 |
|---|---|---|---|---|---|---|
| V | 1 | --- | 100.076 | --- | --- | 10 |
| G | 2 | --- | 157.097 | 1123.548 | --- | 9 |
| S@ | 3 | --- | 535.225 | 1066.526 | --- | 8 |
| I | 4 | --- | 648.309 | **688.399** | 370.189 | 7 |
| N | 5 | --- | 762.352 | **575.315** | 52.311 | 6 |
| I | 6 | --- | 875.436 | **461.272** | 629.236 | 5 |
| G | 7 | -875.554 | **932.457** | **348.188** | 305.047 | 4 |
| S | 8 | --- | 1019.489 | **291.166** | 630.441 | 3 |
| G | 9 | --- | 1076.511 | 204.134 | --- | 2 |
| K | 10 | --- | --- | 147.113 | --- | 1 |

| Run: | 77660 | Scans: | 14893 - 14893 | m/z Range: | 153.26353 to 1072.6663 |
|---|---|---|---|---|---|
| Search: | 77660 | Precursor Mass: | 611.81187 | | |
| Mods: | C=160.03 *+15.99 #+220.06 @+291.1 | Charge: | +2 | | |

**Sequence**

VGS@INIGSGK

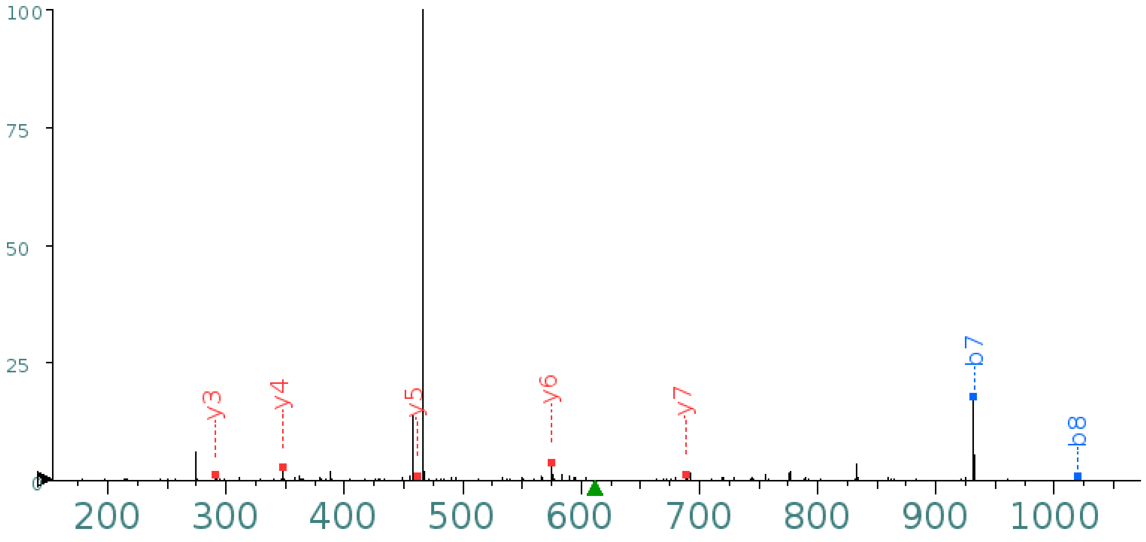

**Predicted Fragmentation Pattern**

| Seq | # | b: Δ Error | b | y | y: Δ Error | +1 |
|---|---|---|---|---|---|---|
| V | 1 | --- | 100.076 | --- | --- | 10 |
| G | 2 | --- | 157.097 | 1123.548 | --- | 9 |
| S@ | 3 | --- | 535.225 | 1066.526 | --- | 8 |
| I | 4 | --- | 648.309 | **688.399** | 176.567 | 7 |
| N | 5 | --- | 762.352 | **575.315** | 467.120 | 6 |
| I | 6 | --- | 875.436 | **461.272** | 210.003 | 5 |
| G | 7 | -774.323 | **932.457** | **348.188** | 397.274 | 4 |
| S | 8 | 79.066 | **1019.489** | **291.166** | 528.682 | 3 |
| G | 9 | --- | 1076.511 | 204.134 | --- | 2 |
| K | 10 | --- | --- | 147.113 | --- | 1 |

| Run: | 77660 | Scans: | 15007 - 15007 | m/z Range: | 148.23814 to 1003.8187[illegible] |
|---|---|---|---|---|---|
| Search: | 77660 | Precursor Mass: | 611.81189 | | |
| Mods: | C=160.03 *+15.99 #+220.06 @+291.1 | Charge: | +2 | | |

### Sequence

VGS@INIGSGK

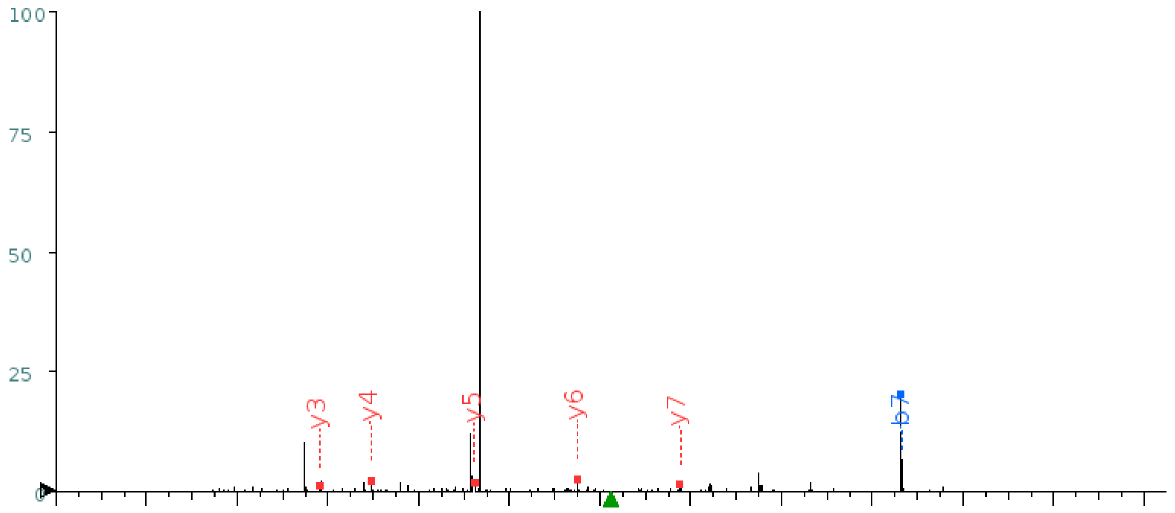

### Predicted Fragmentation Pattern

| Seq | # | b: Δ Error | b | y | y: Δ Error | +1 |
|---|---|---|---|---|---|---|
| V | 1 | --- | 100.076 | --- | --- | 10 |
| G | 2 | --- | 157.097 | 1123.548 | --- | 9 |
| S@ | 3 | --- | 535.225 | 1066.526 | --- | 8 |
| I | 4 | --- | 648.309 | **688.399** | -829.522 | 7 |
| N | 5 | --- | 762.352 | **575.315** | 63.979 | 6 |
| I | 6 | --- | 875.436 | **461.272** | 299.273 | 5 |
| G | 7 | -984.020 | **932.457** | **348.188** | 29.490 | 4 |
| S | 8 | --- | 1019.489 | **291.166** | 435.071 | 3 |
| G | 9 | --- | 1076.511 | 204.134 | --- | 2 |
| K | 10 | --- | --- | 147.113 | --- | 1 |

| Run: | 77660 | Scans: | 15122 - 15122 | m/z Range: | 154.29453 to 1175.8139 |
|---|---|---|---|---|---|
| Search: | 77660 | Precursor Mass: | 611.81193 | | |
| Mods: | C=160.03 *+15.99 #+220.06 @+291.1 | Charge: | +2 | | |

**Sequence**

VGS@INIGSGK

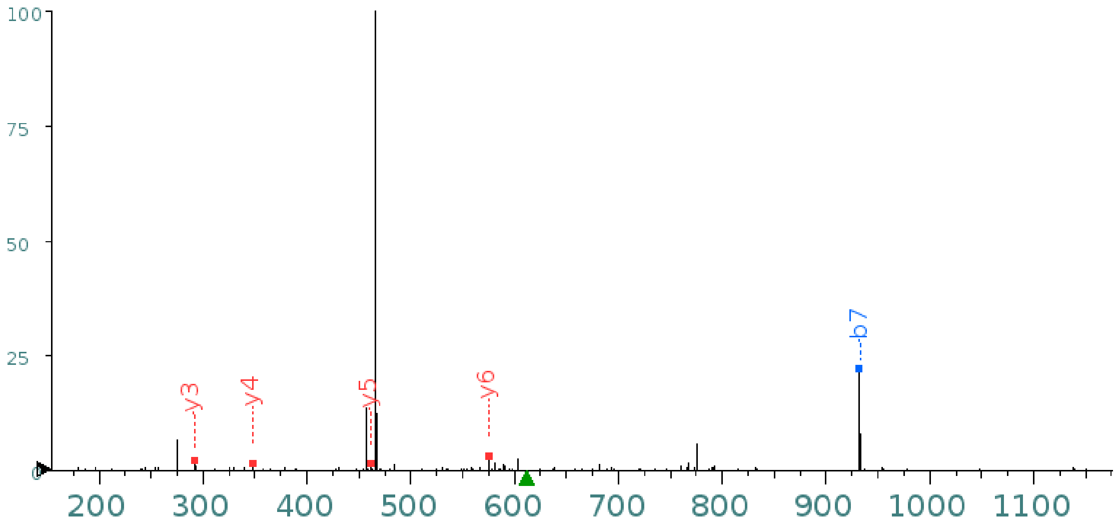

**Predicted Fragmentation Pattern**

| Seq | # | b: Δ Error | b | y | y: Δ Error | +1 |
|---|---|---|---|---|---|---|
| V | 1 | --- | 100.076 | --- | --- | 10 |
| G | 2 | --- | 157.097 | 1123.548 | --- | 9 |
| S@ | 3 | --- | 535.225 | 1066.526 | --- | 8 |
| I | 4 | --- | 648.309 | 688.399 | --- | 7 |
| N | 5 | --- | 762.352 | **575.315** | 326.980 | 6 |
| I | 6 | --- | 875.436 | **461.272** | 170.587 | 5 |
| G | 7 | -927.161 | **932.457** | **348.188** | 502.443 | 4 |
| S | 8 | --- | 1019.489 | **291.166** | 3296.403 | 3 |
| G | 9 | --- | 1076.511 | 204.134 | --- | 2 |
| K | 10 | --- | --- | 147.113 | --- | 1 |

| Run: | 77660 | Scans: | 15513 - 15513 | m/z Range: | 154.18517 to 1210.0252 |
|---|---|---|---|---|---|
| Search: | 77660 | Precursor Mass: | 611.81262 | | |
| Mods: | C=160.03 *+15.99 #+220.06 @+291.1 | Charge: | +2 | | |

#### Sequence

VGS@INIGSGK

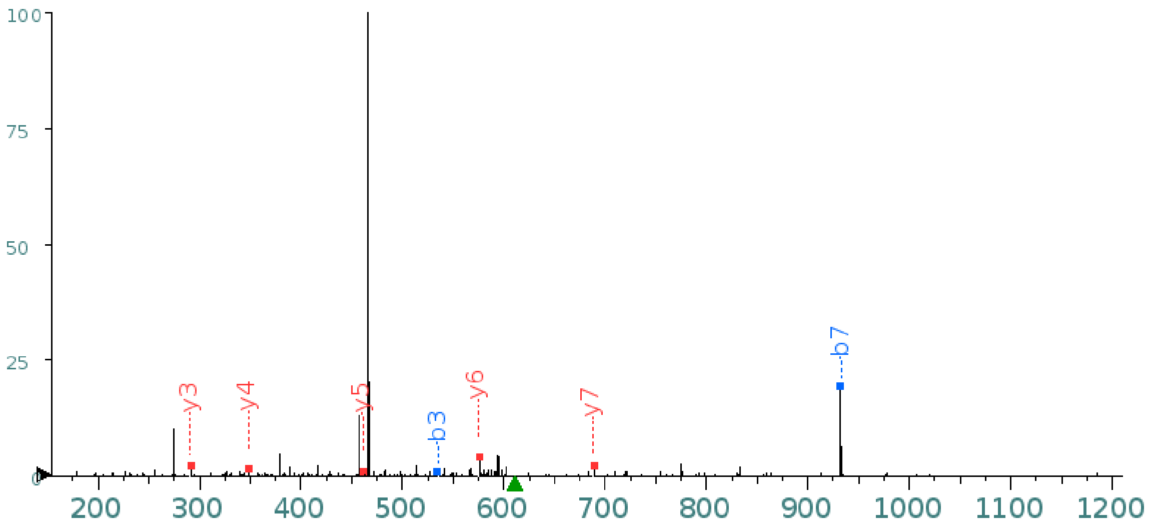

#### Predicted Fragmentation Pattern

| Seq | # | b: Δ Error | b | y | y: Δ Error | +1 |
|---|---|---|---|---|---|---|
| V | 1 | --- | 100.076 | --- | --- | 10 |
| G | 2 | --- | 157.097 | 1123.548 | --- | 9 |
| S@ | 3 | -1416.499 | **535.225** | 1066.526 | --- | 8 |
| I | 4 | --- | 648.309 | **688.399** | 568.960 | 7 |
| N | 5 | --- | 762.352 | **575.315** | 450.585 | 6 |
| I | 6 | --- | 875.436 | **461.272** | 895.057 | 5 |
| G | 7 | -938.638 | **932.457** | **348.188** | 732.845 | 4 |
| S | 8 | --- | 1019.489 | **291.166** | 3368.554 | 3 |
| G | 9 | --- | 1076.511 | 204.134 | --- | 2 |
| K | 10 | --- | --- | 147.113 | --- | 1 |

| | | | | | |
|---|---|---|---|---|---|
| **Run:** | 77660 | **Scans:** | 15596 - 15596 | **m/z Range:** | 180.1864 to 1414.8564 |
| **Search:** | 77660 | **Precursor Mass:** | 714.84725 | | |
| **Mods:** | C=160.03 *+15.99 #+220.06 @+291.1 | **Charge:** | +2 | | |

**Sequence**

IAS@QTQFNTK

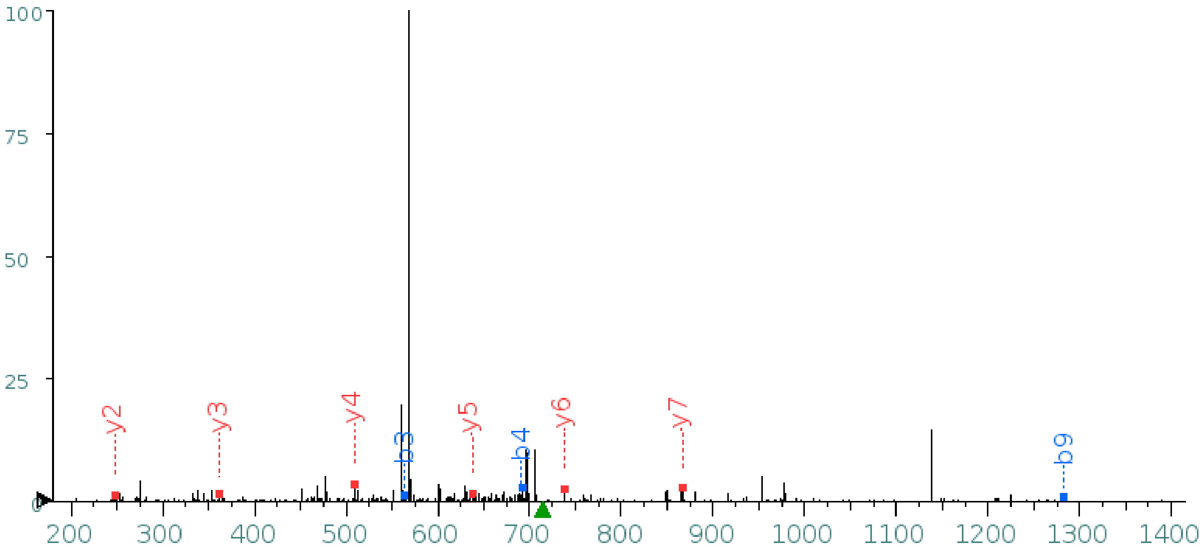

**Predicted Fragmentation Pattern**

| Seq | # | b: Δ Error | b | y | y: Δ Error | +1 |
|---|---|---|---|---|---|---|
| I | 1 | --- | 114.091 | --- | --- | 10 |
| A | 2 | --- | 185.128 | 1315.601 | --- | 9 |
| S@ | 3 | 629.973 | **563.256** | 1244.564 | --- | 8 |
| Q | 4 | 334.816 | **691.314** | **866.437** | 389.037 | 7 |
| T | 5 | --- | 792.362 | **738.378** | 209.876 | 6 |
| Q | 6 | --- | 920.421 | **637.330** | 397.628 | 5 |
| F | 7 | --- | 1067.489 | **509.272** | 385.933 | 4 |
| N | 8 | --- | 1181.532 | **362.203** | 813.888 | 3 |
| T | 9 | -376.214 | **1282.580** | **248.160** | 728.195 | 2 |
| K | 10 | --- | --- | 147.113 | --- | 1 |

| | | | | | |
|---|---|---|---|---|---|
| **Run:** | 77660 | **Scans:** | 15743 - 15743 | **m/z Range:** | 148.51132 to 1117.6464[illegible] |
| **Search:** | 77660 | **Precursor Mass:** | 611.81222 | | |
| **Mods:** | C=160.03 *+15.99 #+220.06 @+291.1 | **Charge:** | +2 | | |

**Sequence**

VGS@INIGSGK

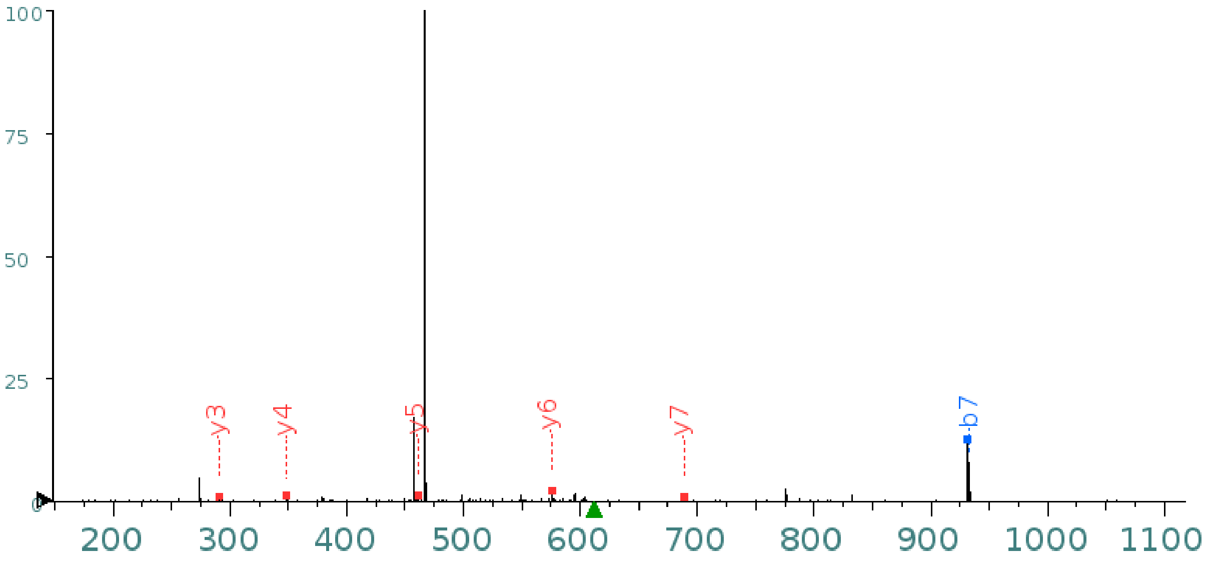

**Predicted Fragmentation Pattern**

| Seq | # | b: Δ Error | b | y | y: Δ Error | +1 |
|---|---|---|---|---|---|---|
| V | 1 | --- | 100.076 | --- | --- | 10 |
| G | 2 | --- | 157.097 | 1123.548 | --- | 9 |
| S@ | 3 | --- | 535.225 | 1066.526 | --- | 8 |
| I | 4 | --- | 648.309 | **688.399** | **505.723** | 7 |
| N | 5 | --- | 762.352 | **575.315** | **227.098** | 6 |
| I | 6 | --- | 875.436 | **461.272** | **158.286** | 5 |
| G | 7 | **-1044.100** | **932.457** | **348.188** | **425.561** | 4 |
| S | 8 | --- | 1019.489 | **291.166** | **436.642** | 3 |
| G | 9 | --- | 1076.511 | 204.134 | --- | 2 |
| K | 10 | --- | --- | 147.113 | --- | 1 |

| | | | | | |
|---|---|---|---|---|---|
| **Run:** | 77660 | **Scans:** | 15959 - 15959 | **m/z Range:** | 148.09207 to 1214.6392 |
| **Search:** | 77660 | **Precursor Mass:** | 611.81213 | | |
| **Mods:** | C=160.03 *+15.99 #+220.06 @+291.1 | **Charge:** | +2 | | |

**Sequence**

VGS@INIGSGK

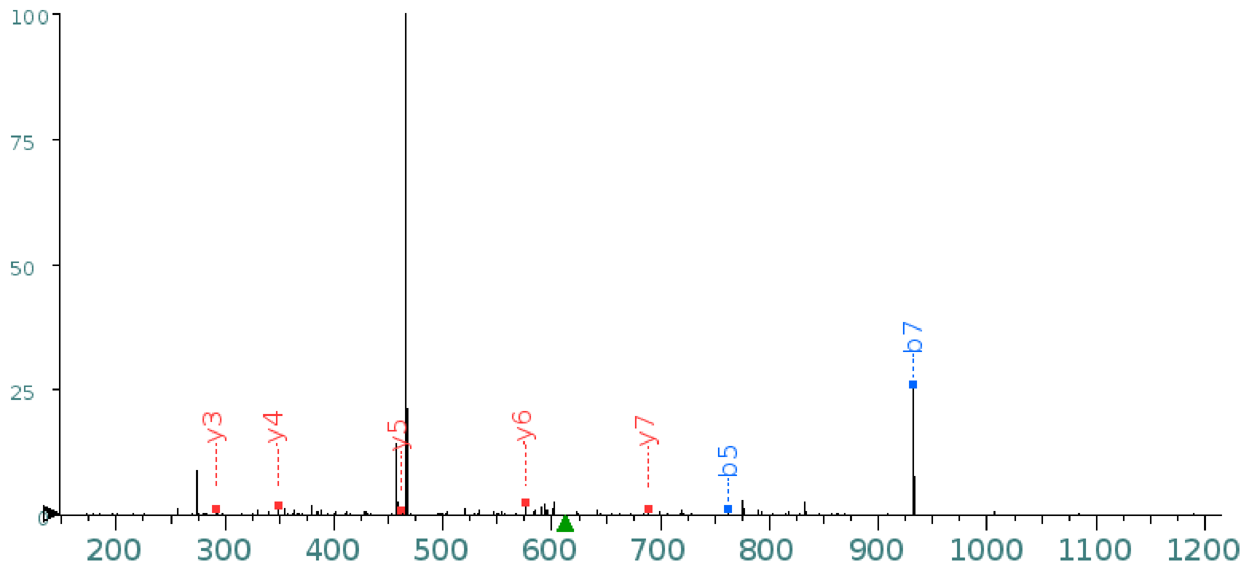

**Predicted Fragmentation Pattern**

| Seq | # | b: Δ Error | b | y | y: Δ Error | +1 |
|---|---|---|---|---|---|---|
| V | 1 | --- | 100.076 | --- | --- | 10 |
| G | 2 | --- | 157.097 | 1123.548 | --- | 9 |
| S@ | 3 | --- | 535.225 | 1066.526 | --- | 8 |
| I | 4 | --- | 648.309 | **688.399** | 305.509 | 7 |
| N | 5 | -958.496 | **762.352** | **575.315** | 346.275 | 6 |
| I | 6 | --- | 875.436 | **461.272** | 837.729 | 5 |
| G | 7 | -959.230 | **932.457** | **348.188** | 452.708 | 4 |
| S | 8 | --- | 1019.489 | **291.166** | 11.193 | 3 |
| G | 9 | --- | 1076.511 | 204.134 | --- | 2 |
| K | 10 | --- | --- | 147.113 | --- | 1 |

| Run: | 77660 | Scans: | 16082 - 16082 | m/z Range: | 154.20549 | to | 1072.7659 |
|---|---|---|---|---|---|---|---|
| Search: | 77660 | Precursor Mass: | 611.81207 | | | | |
| Mods: | C=160.03 *+15.99 #+220.06 @+291.1 | Charge: | +2 | | | | |

**Sequence**

VGS@INIGSGK

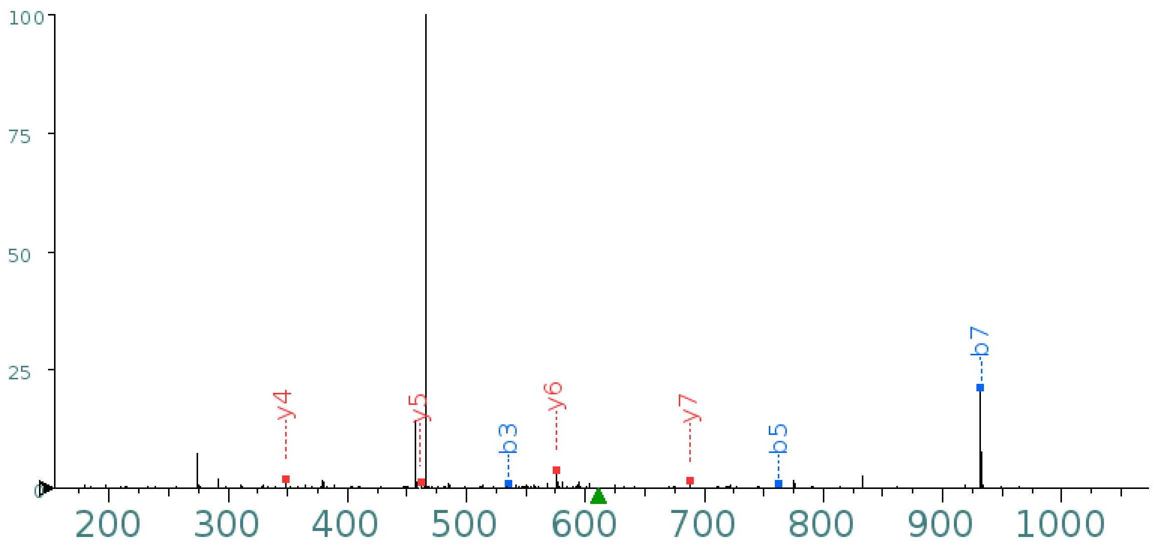

**Predicted Fragmentation Pattern**

| Seq | # | b: Δ Error | b | y | y: Δ Error | +1 |
|---|---|---|---|---|---|---|
| V | 1 | --- | 100.076 | --- | --- | 10 |
| G | 2 | --- | 157.097 | 1123.548 | --- | 9 |
| S@ | 3 | 1111.773 | **535.225** | 1066.526 | --- | 8 |
| I | 4 | --- | 648.309 | **688.399** | 144.747 | 7 |
| N | 5 | -176.047 | **762.352** | **575.315** | 479.626 | 6 |
| I | 6 | --- | 875.436 | **461.272** | 743.535 | 5 |
| G | 7 | -879.161 | **932.457** | **348.188** | 644.444 | 4 |
| S | 8 | --- | 1019.489 | 291.166 | --- | 3 |
| G | 9 | --- | 1076.511 | 204.134 | --- | 2 |
| K | 10 | --- | --- | 147.113 | --- | 1 |

| | | | | | |
|---|---|---|---|---|---|
| **Run:** | 77660 | **Scans:** | 16202 - 16202 | **m/z Range:** | 154.1841 to 1142.6473 |
| **Search:** | 77660 | **Precursor Mass:** | 611.81202 | | |
| **Mods:** | C=160.03 *+15.99 #+220.06 @+291.1 | **Charge:** | +2 | | |

**Sequence**

VGS@INIGSGK

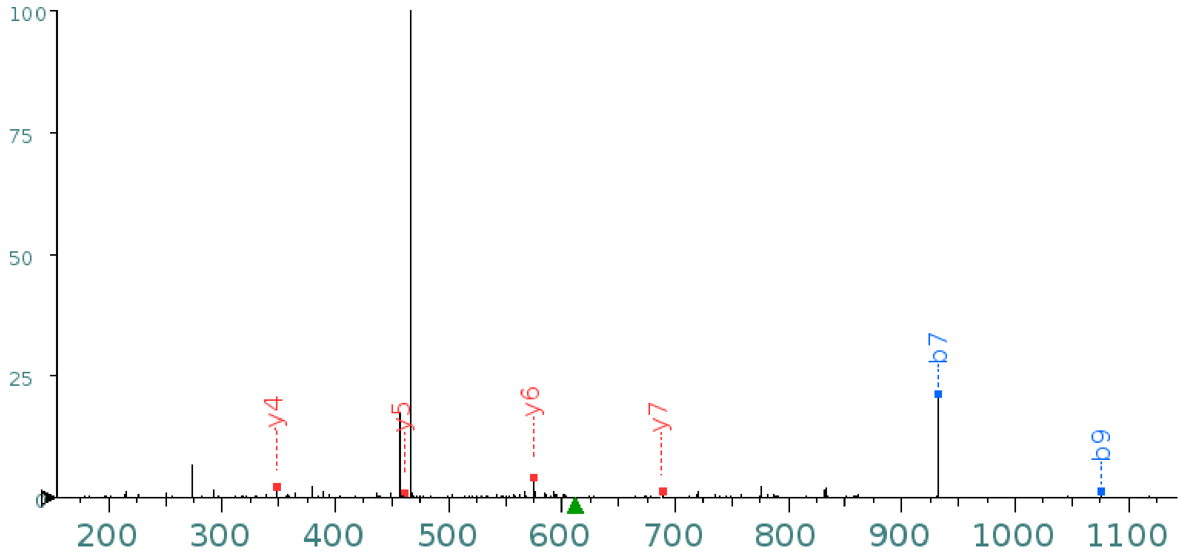

**Predicted Fragmentation Pattern**

| Seq | # | b: Δ Error | b | y | y: Δ Error | +1 |
|---|---|---|---|---|---|---|
| V | 1 | --- | 100.076 | --- | --- | 10 |
| G | 2 | --- | 157.097 | 1123.548 | --- | 9 |
| S@ | 3 | --- | 535.225 | 1066.526 | --- | 8 |
| I | 4 | --- | 648.309 | **688.399** | **315.787** | 7 |
| N | 5 | --- | 762.352 | **575.315** | **159.863** | 6 |
| I | 6 | --- | 875.436 | **461.272** | **-90.193** | 5 |
| G | 7 | **-987.562** | **932.457** | **348.188** | **479.502** | 4 |
| S | 8 | --- | 1019.489 | 291.166 | --- | 3 |
| G | 9 | **-710.234** | **1076.511** | 204.134 | --- | 2 |
| K | 10 | --- | --- | 147.113 | --- | 1 |

| Run: | 77660 | Scans: | 16330 - 16330 | m/z Range: | 154.14664 to 991.63422 |
|---|---|---|---|---|---|
| Search: | 77660 | Precursor Mass: | 611.81203 | | |
| Mods: | C=160.03 *+15.99 #+220.06 @+291.1 | Charge: | +2 | | |

**Sequence**

VGS@INIGSGK

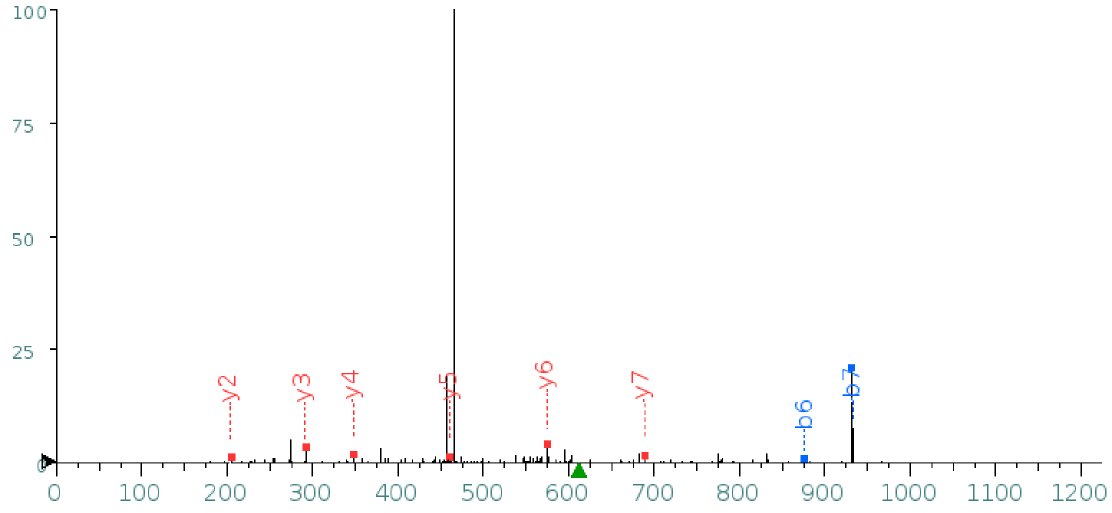

**Predicted Fragmentation Pattern**

| Seq | # | b: Δ Error | b | y | y: Δ Error | +1 |
|---|---|---|---|---|---|---|
| V | 1 | --- | 100.076 | --- | --- | 10 |
| G | 2 | --- | 157.097 | 1123.548 | --- | 9 |
| S@ | 3 | --- | 535.225 | 1066.526 | --- | 8 |
| I | 4 | --- | 648.309 | **688.399** | 191.368 | 7 |
| N | 5 | --- | 762.352 | **575.315** | 199.633 | 6 |
| I | 6 | 294.910 | **875.436** | **461.272** | -105.016 | 5 |
| G | 7 | -936.933 | **932.457** | **348.188** | 230.061 | 4 |
| S | 8 | --- | 1019.489 | **291.166** | 3340.757 | 3 |
| G | 9 | --- | 1076.511 | **204.134** | 2277.822 | 2 |
| K | 10 | --- | --- | 147.113 | --- | 1 |

| Run: | 77660 | Scans: | 16561 - 16561 | m/z Range: | 154.16641 to 1223.6239[illegible] |
|---|---|---|---|---|---|
| Search: | 77660 | Precursor Mass: | 611.81199 | | |
| Mods: | C=160.03 *+15.99 #+220.06 @+291.1 | Charge: | +2 | | |

### Sequence

VGS@INIGSGK

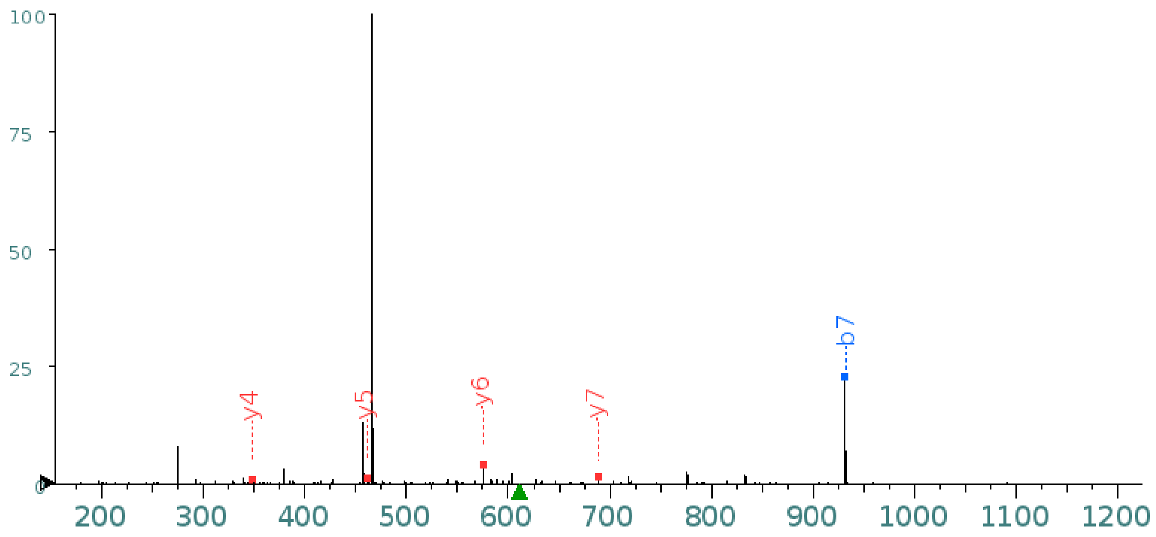

### Predicted Fragmentation Pattern

| Seq | # | b: Δ Error | b | y | y: Δ Error | +1 |
|---|---|---|---|---|---|---|
| V | 1 | --- | 100.076 | --- | --- | 10 |
| G | 2 | --- | 157.097 | 1123.548 | --- | 9 |
| S@ | 3 | --- | 535.225 | 1066.526 | --- | 8 |
| I | 4 | --- | 648.309 | **688.399** | 147.761 | 7 |
| N | 5 | --- | 762.352 | **575.315** | -39.668 | 6 |
| I | 6 | --- | 875.436 | **461.272** | 308.266 | 5 |
| G | 7 | -1009.206 | **932.457** | **348.188** | 172.413 | 4 |
| S | 8 | --- | 1019.489 | 291.166 | --- | 3 |
| G | 9 | --- | 1076.511 | 204.134 | --- | 2 |
| K | 10 | --- | --- | 147.113 | --- | 1 |

| Run: | 77660 | Scans: | 16672 - 16672 | m/z Range: | 147.50533 | to | 1146.6921 |
|---|---|---|---|---|---|---|---|
| Search: | 77660 | Precursor Mass: | 611.81207 | | | | |
| Mods: | C=160.03 *+15.99 #+220.06 @+291.1 | Charge: | +2 | | | | |

**Sequence**

VGS@INIGSGK

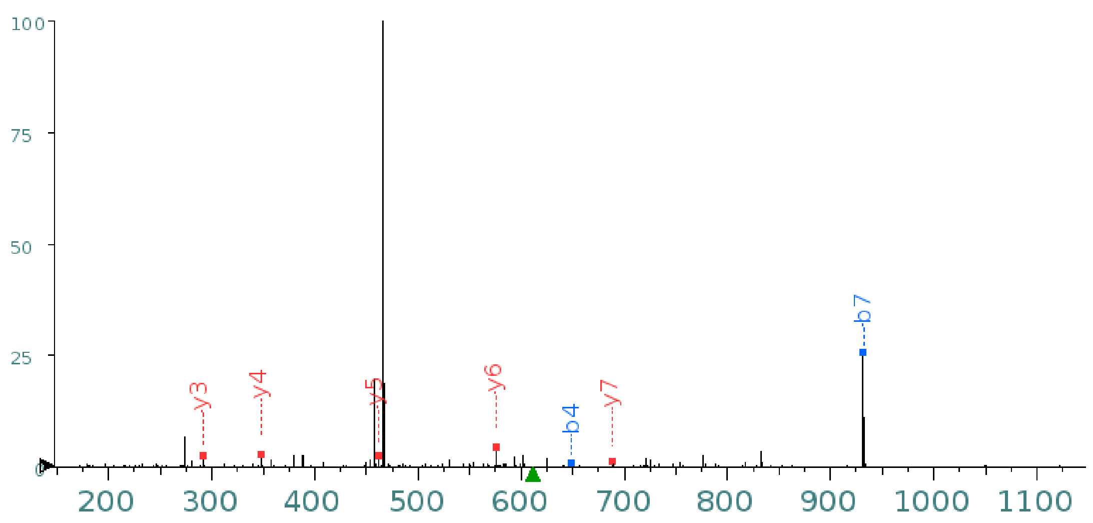

**Predicted Fragmentation Pattern**

| Seq | # | b: Δ Error | b | y | y: Δ Error | +1 |
|---|---|---|---|---|---|---|
| V | 1 | --- | 100.076 | --- | --- | 10 |
| G | 2 | --- | 157.097 | 1123.548 | --- | 9 |
| S@ | 3 | --- | 535.225 | 1066.526 | --- | 8 |
| I | 4 | 405.114 | 648.309 | 688.399 | 127.108 | 7 |
| N | 5 | --- | 762.352 | 575.315 | 131.758 | 6 |
| I | 6 | --- | 875.436 | 461.272 | -78.547 | 5 |
| G | 7 | -907.816 | 932.457 | 348.188 | 513.476 | 4 |
| S | 8 | --- | 1019.489 | 291.166 | 3011.861 | 3 |
| G | 9 | --- | 1076.511 | 204.134 | --- | 2 |
| K | 10 | --- | --- | 147.113 | --- | 1 |

| | | | | | |
|---|---|---|---|---|---|
| **Run:** | 77660 | **Scans:** | 16786 - 16786 | **m/z Range:** | 154.23781 to 987.69952 |
| **Search:** | 77660 | **Precursor Mass:** | 611.81197 | | |
| **Mods:** | C=160.03 *+15.99 #+220.06 @+291.1 | **Charge:** | +2 | | |

**Sequence**

VGS@INIGSGK

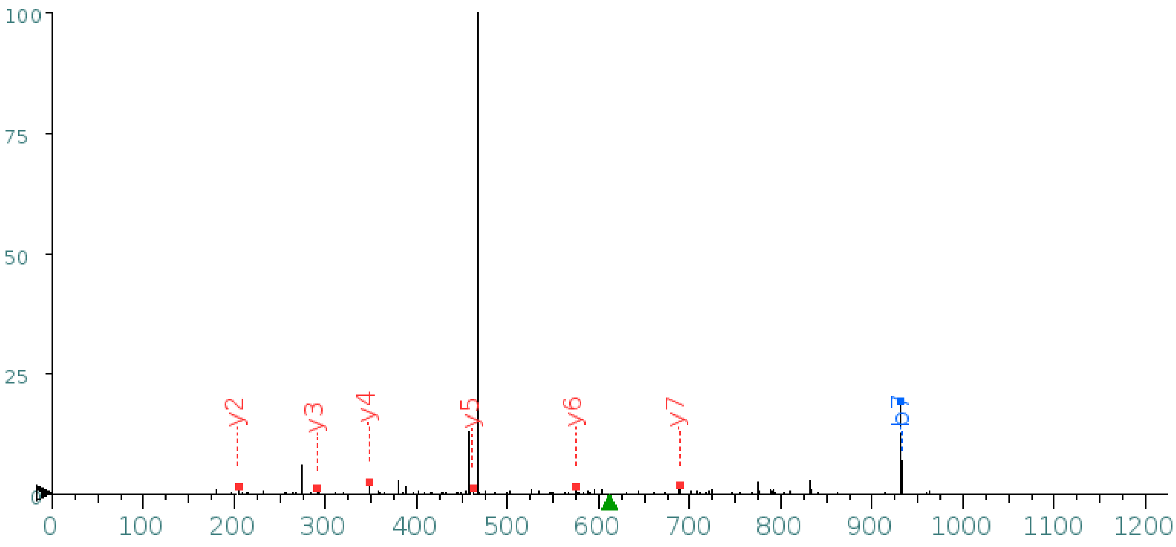

**Predicted Fragmentation Pattern**

| Seq | # | b: Δ Error | b | y | y: Δ Error | +1 |
|---|---|---|---|---|---|---|
| V | 1 | --- | 100.076 | --- | --- | 10 |
| G | 2 | --- | 157.097 | 1123.548 | --- | 9 |
| S@ | 3 | --- | 535.225 | 1066.526 | --- | 8 |
| I | 4 | --- | 648.309 | **688.399** | 411.650 | 7 |
| N | 5 | --- | 762.352 | **575.315** | 207.162 | 6 |
| I | 6 | --- | 875.436 | **461.272** | 294.777 | 5 |
| G | 7 | -963.099 | **932.457** | **348.188** | 90.923 | 4 |
| S | 8 | --- | 1019.489 | **291.166** | 312.952 | 3 |
| G | 9 | --- | 1076.511 | **204.134** | 569.455 | 2 |
| K | 10 | --- | --- | 147.113 | --- | 1 |

| Run: | 77660 | Scans: | 17016 - 17016 | m/z Range: | 148.42015 to 1173.7448 |
|---|---|---|---|---|---|
| Search: | 77660 | Precursor Mass: | 611.81205 | | |
| Mods: | C=160.03 *+15.99 #+220.06 @+291.1 | Charge: | +2 | | |

**Sequence**

VGS@INIGSGK

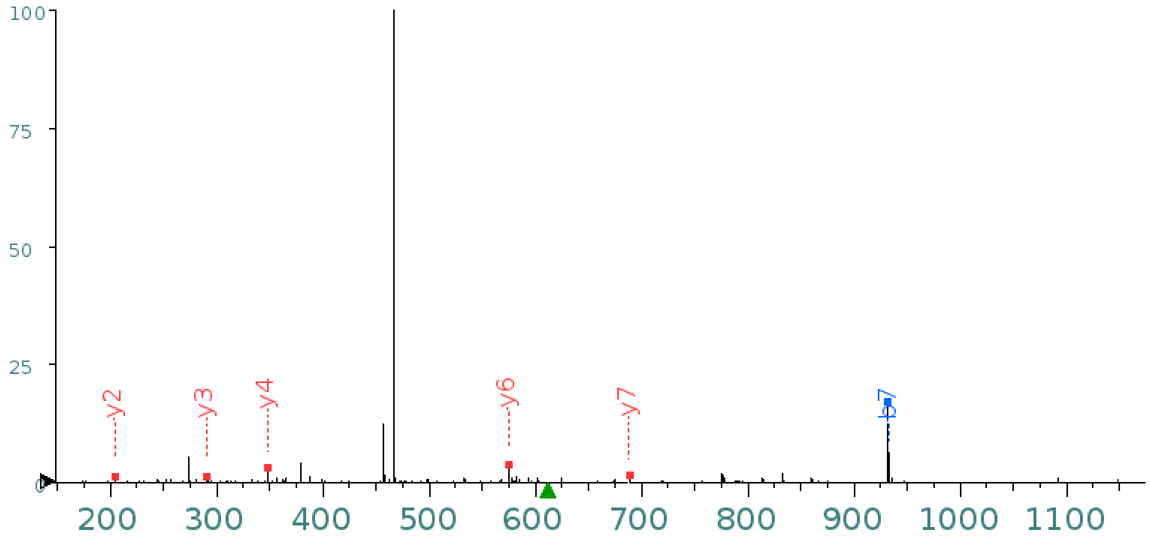

**Predicted Fragmentation Pattern**

| Seq | # | b: Δ Error | b | y | y: Δ Error | +1 |
|---|---|---|---|---|---|---|
| V | 1 | --- | 100.076 | --- | --- | 10 |
| G | 2 | --- | 157.097 | 1123.548 | --- | 9 |
| S@ | 3 | --- | 535.225 | 1066.526 | --- | 8 |
| I | 4 | --- | 648.309 | **688.399** | **318.445** | 7 |
| N | 5 | --- | 762.352 | **575.315** | **172.590** | 6 |
| I | 6 | --- | 875.436 | 461.272 | --- | 5 |
| G | 7 | **-807.759** | **932.457** | **348.188** | **491.761** | 4 |
| S | 8 | --- | 1019.489 | **291.166** | **169.220** | 3 |
| G | 9 | --- | 1076.511 | **204.134** | **-159.345** | 2 |
| K | 10 | --- | --- | 147.113 | --- | 1 |

| Run: | 77660 | Scans: | 17146 - 17146 | m/z Range: | 153.12198 | to | 1135.6097[illegible] |
|---|---|---|---|---|---|---|---|
| Search: | 77660 | Precursor Mass: | 611.81207 | | | | |
| Mods: | C=160.03 *+15.99 #+220.06 @+291.1 | Charge: | +2 | | | | |

**Sequence**

VGS@INIGSGK

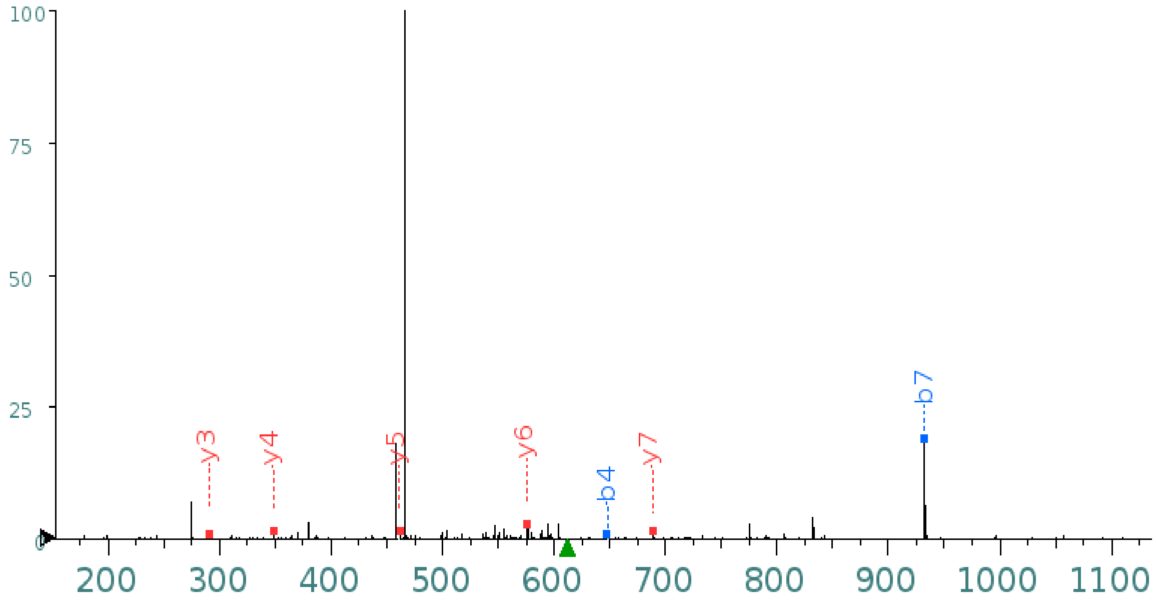

**Predicted Fragmentation Pattern**

| Seq | # | b: Δ Error | b | y | y: Δ Error | +1 |
|---|---|---|---|---|---|---|
| V | 1 | --- | 100.076 | --- | --- | 10 |
| G | 2 | --- | 157.097 | 1123.548 | --- | 9 |
| S@ | 3 | --- | 535.225 | 1066.526 | --- | 8 |
| I | 4 | -1109.430 | **648.309** | **688.399** | 250.303 | 7 |
| N | 5 | --- | 762.352 | **575.315** | 210.025 | 6 |
| I | 6 | --- | 875.436 | **461.272** | 707.068 | 5 |
| G | 7 | -870.571 | **932.457** | **348.188** | 667.377 | 4 |
| S | 8 | --- | 1019.489 | **291.166** | 641.118 | 3 |
| G | 9 | --- | 1076.511 | 204.134 | --- | 2 |
| K | 10 | --- | --- | 147.113 | --- | 1 |

| | | | | | |
|---|---|---|---|---|---|
| **Run:** | 77660 | **Scans:** | 17261 - 17261 | **m/z Range:** | 148.27347 to 1135.9193 |
| **Search:** | 77660 | **Precursor Mass:** | 611.81203 | | |
| **Mods:** | C=160.03 *+15.99 #+220.06 @+291.1 | **Charge:** | +2 | | |

**Sequence**

VGS@INIGSGK

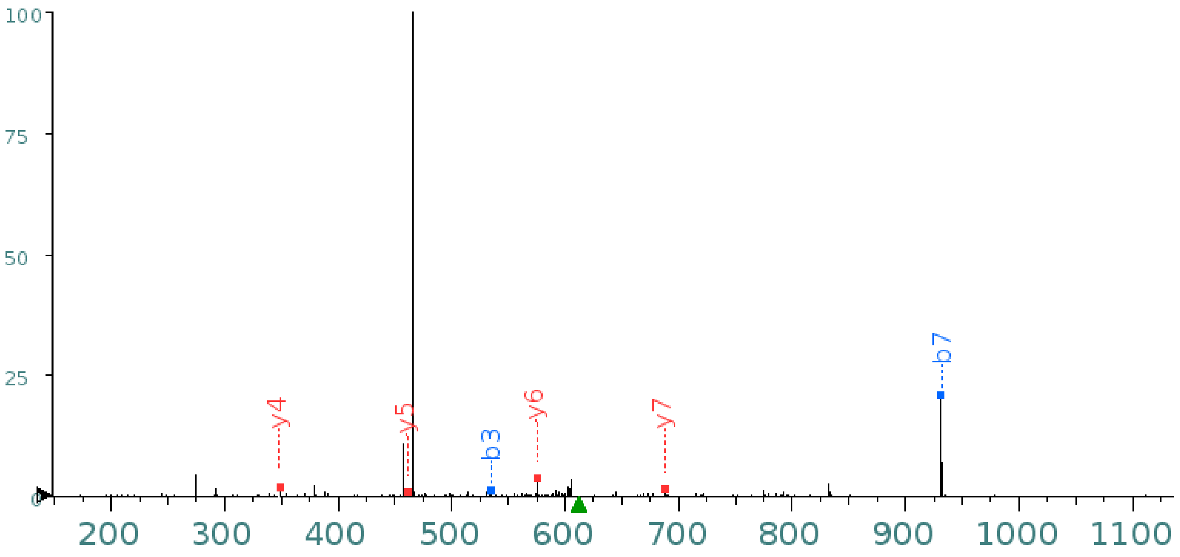

**Predicted Fragmentation Pattern**

| Seq | # | b: Δ Error | b | y | y: Δ Error | +1 |
|---|---|---|---|---|---|---|
| V | 1 | --- | 100.076 | --- | --- | 10 |
| G | 2 | --- | 157.097 | 1123.548 | --- | 9 |
| S@ | 3 | -886.607 | **535.225** | 1066.526 | --- | 8 |
| I | 4 | --- | 648.309 | **688.399** | 260.316 | 7 |
| N | 5 | --- | 762.352 | **575.315** | 382.108 | 6 |
| I | 6 | --- | 875.436 | **461.272** | 140.957 | 5 |
| G | 7 | -958.181 | **932.457** | **348.188** | 744.047 | 4 |
| S | 8 | --- | 1019.489 | 291.166 | --- | 3 |
| G | 9 | --- | 1076.511 | 204.134 | --- | 2 |
| K | 10 | --- | --- | 147.113 | --- | 1 |

| Run: | 77660 | Scans: | 17375 - 17375 | m/z Range: | 154.32268 to 1223.6240[illegible] |
|---|---|---|---|---|---|
| Search: | 77660 | Precursor Mass: | 611.81205 | | |
| Mods: | C=160.03 *+15.99 #+220.06 @+291.1 | Charge: | +2 | | |

**Sequence**

VGS@INIGSGK

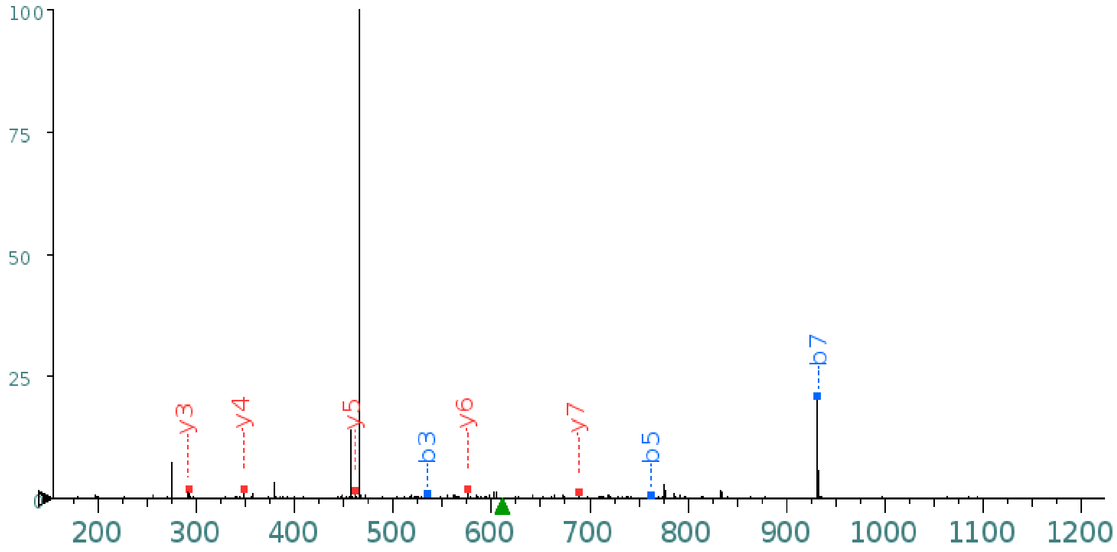

**Predicted Fragmentation Pattern**

| Seq | # | b: Δ Error | b | y | y: Δ Error | +1 |
|---|---|---|---|---|---|---|
| V | 1 | --- | 100.076 | --- | --- | 10 |
| G | 2 | --- | 157.097 | 1123.548 | --- | 9 |
| S@ | 3 | 0.470 | **535.225** | 1066.526 | --- | 8 |
| I | 4 | --- | 648.309 | **688.399** | 53.088 | 7 |
| N | 5 | -708.849 | **762.352** | **575.315** | 160.500 | 6 |
| I | 6 | --- | 875.436 | **461.272** | 557.868 | 5 |
| G | 7 | -984.020 | **932.457** | **348.188** | 424.948 | 4 |
| S | 8 | --- | 1019.489 | **291.166** | 3207.059 | 3 |
| G | 9 | --- | 1076.511 | 204.134 | --- | 2 |
| K | 10 | --- | --- | 147.113 | --- | 1 |

| Run: | 77660 | Scans: | 17490 - 17490 | m/z Range: | 153.30373 to 1065.5953 |
|---|---|---|---|---|---|
| Search: | 77660 | Precursor Mass: | 611.81206 | | |
| Mods: | C=160.03 *+15.99 #+220.06 @+291.1 | Charge: | +2 | | |

**Sequence**

VGS@INIGSGK

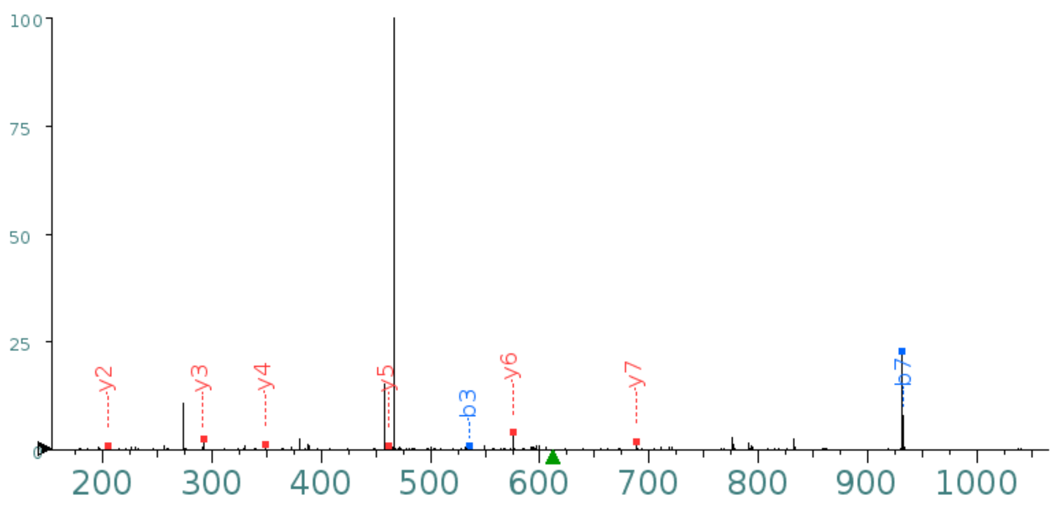

**Predicted Fragmentation Pattern**

| Seq | # | b: Δ Error | b | y | y: Δ Error | +1 |
|---|---|---|---|---|---|---|
| V | 1 | --- | 100.076 | --- | --- | 10 |
| G | 2 | --- | 157.097 | 1123.548 | --- | 9 |
| S@ | 3 | 507.788 | 535.225 | 1066.526 | --- | 8 |
| I | 4 | --- | 648.309 | 688.399 | 345.026 | 7 |
| N | 5 | --- | 762.352 | 575.315 | 359.315 | 6 |
| I | 6 | --- | 875.436 | 461.272 | 733.361 | 5 |
| G | 7 | -947.819 | 932.457 | 348.188 | 356.899 | 4 |
| S | 8 | --- | 1019.489 | 291.166 | 3272.038 | 3 |
| G | 9 | --- | 1076.511 | 204.134 | 971.283 | 2 |
| K | 10 | --- | --- | 147.113 | --- | 1 |

| Run: | 77660 | Scans: | 17721 - 17721 | m/z Range: | 150.40028 to 1212.3941 |
|---|---|---|---|---|---|
| Search: | 77660 | Precursor Mass: | 611.81207 | | |
| Mods: | C=160.03 *+15.99 #+220.06 @+291.1 | Charge: | +2 | | |

**Sequence**

VGS@INIGSGK

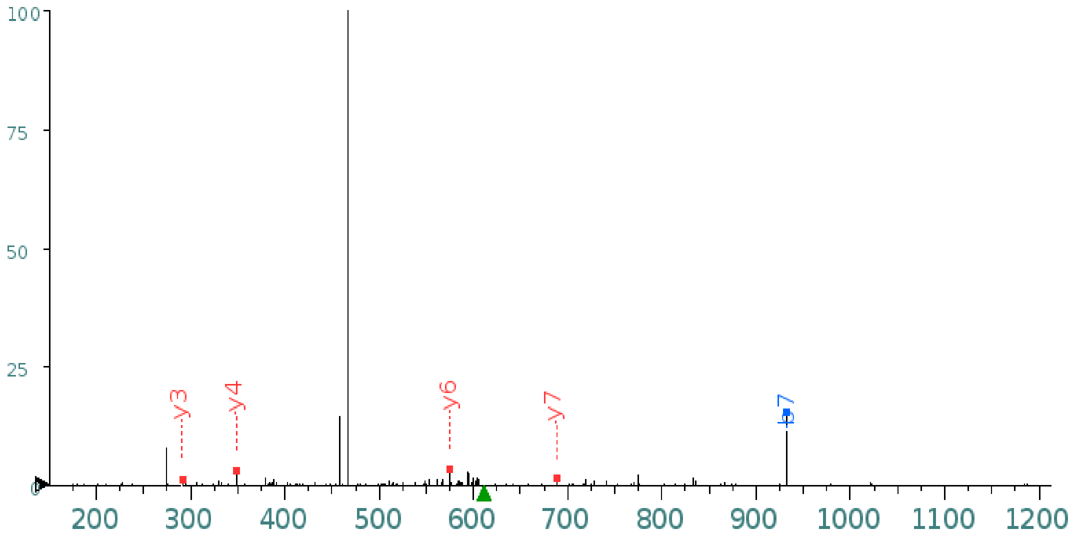

**Predicted Fragmentation Pattern**

| Seq | # | b: Δ Error | b | y | y: Δ Error | +1 |
|---|---|---|---|---|---|---|
| V | 1 | --- | 100.076 | --- | --- | 10 |
| G | 2 | --- | 157.097 | 1123.548 | --- | 9 |
| S@ | 3 | --- | 535.225 | 1066.526 | --- | 8 |
| I | 4 | --- | 648.309 | **688.399** | 154.940 | 7 |
| N | 5 | --- | 762.352 | **575.315** | 428.749 | 6 |
| I | 6 | --- | 875.436 | 461.272 | --- | 5 |
| G | 7 | -899.095 | **932.457** | **348.188** | 191.425 | 4 |
| S | 8 | --- | 1019.489 | **291.166** | 508.579 | 3 |
| G | 9 | --- | 1076.511 | 204.134 | --- | 2 |
| K | 10 | --- | --- | 147.113 | --- | 1 |

| Run: | 77660 | Scans: | 17836 - 17836 | m/z Range: | 154.24097 to 1209.9589[illegible] |
|---|---|---|---|---|---|
| Search: | 77660 | Precursor Mass: | 611.81206 | | |
| Mods: | C=160.03 *+15.99 #+220.06 @+291.1 | Charge: | +2 | | |

### Sequence

VGS@INIGSGK

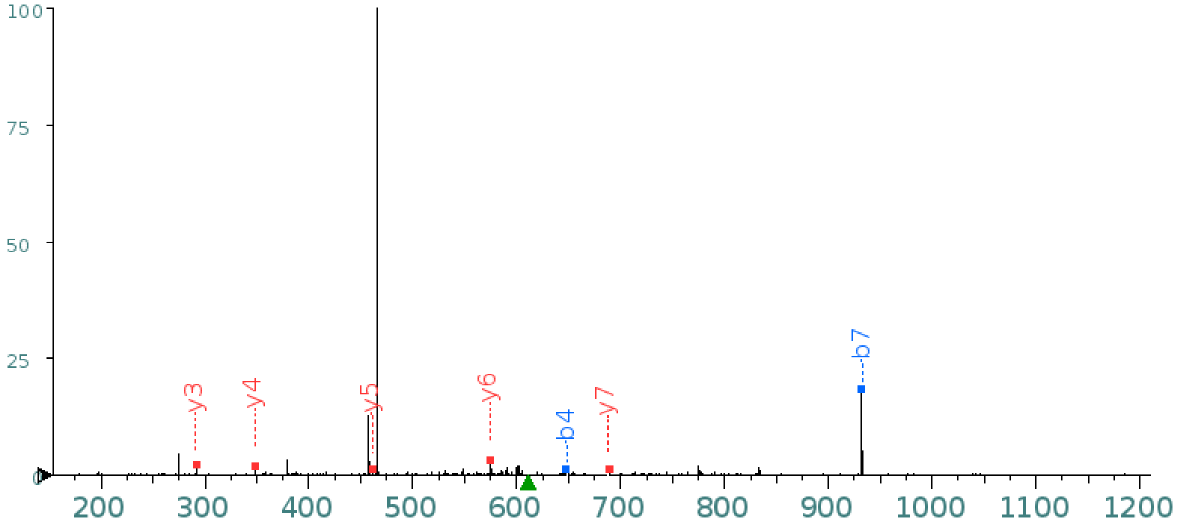

### Predicted Fragmentation Pattern

| Seq | # | b: Δ Error | b | y | y: Δ Error | +1 |
|---|---|---|---|---|---|---|
| V | 1 | --- | 100.076 | --- | --- | 10 |
| G | 2 | --- | 157.097 | 1123.548 | --- | 9 |
| S@ | 3 | --- | 535.225 | 1066.526 | --- | 8 |
| I | 4 | -1067.727 | **648.309** | **688.399** | 339.267 | 7 |
| N | 5 | --- | 762.352 | **575.315** | -49.111 | 6 |
| I | 6 | --- | 875.436 | **461.272** | 473.536 | 5 |
| G | 7 | -961.197 | **932.457** | **348.188** | 607.153 | 4 |
| S | 8 | --- | 1019.489 | **291.166** | 3137.696 | 3 |
| G | 9 | --- | 1076.511 | 204.134 | --- | 2 |
| K | 10 | --- | --- | 147.113 | --- | 1 |

| Run: | 77660 | Scans: | 17905 - 17905 | m/z Range: | 273.28802 to 1808.9906 |
|---|---|---|---|---|---|
| Search: | 77660 | Precursor Mass: | 1008.97078 | | |
| Mods: | C=160.03 *+15.99 #+220.06 @+291.1 | Charge: | +2 | | |

### Sequence

DQVM*ELTIAGM*GT#SALK

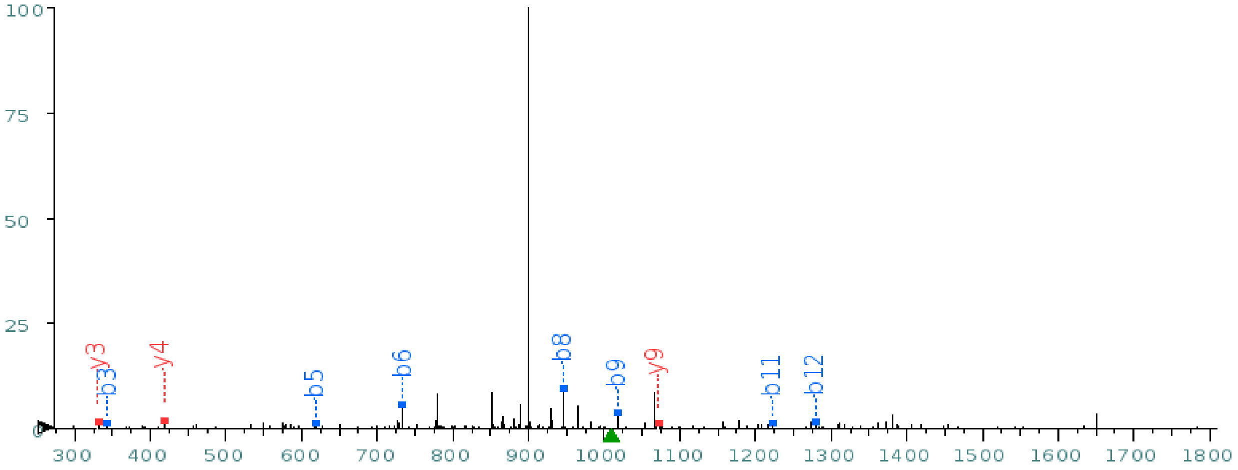

### Predicted Fragmentation Pattern

| Seq | # | b: Δ Error | b | y | y: Δ Error | +1 |
|---|---|---|---|---|---|---|
| D | 1 | --- | 116.034 | --- | --- | 17 |
| Q | 2 | --- | 244.093 | 1901.908 | --- | 16 |
| V | 3 | 841.950 | **343.161** | 1773.850 | --- | 15 |
| M* | 4 | --- | 490.197 | 1674.781 | --- | 14 |
| E | 5 | 226.736 | **619.239** | 1527.746 | --- | 13 |
| L | 6 | 485.130 | **732.323** | 1398.703 | --- | 12 |
| T | 7 | --- | 833.371 | 1285.619 | --- | 11 |
| I | 8 | 148.245 | **946.455** | 1184.571 | --- | 10 |
| A | 9 | 274.004 | **1017.492** | **1071.487** | 394.368 | 9 |
| G | 10 | --- | 1074.514 | 1000.450 | --- | 8 |
| M* | 11 | 276.103 | **1221.549** | 943.429 | --- | 7 |
| G | 12 | 108.817 | **1278.570** | 796.393 | --- | 6 |
| T# | 13 | --- | 1599.676 | 739.372 | --- | 5 |
| S | 14 | --- | 1686.708 | **418.266** | 491.914 | 4 |
| A | 15 | --- | 1757.746 | **331.234** | 984.196 | 3 |
| L | 16 | --- | 1870.830 | 260.197 | --- | 2 |
| K | 17 | --- | --- | 147.113 | --- | 1 |

| Run: | 77660 | Scans: | 18144 - 18144 | m/z Range: | 191.26396 to 1402.5849 |
|---|---|---|---|---|---|
| Search: | 77660 | Precursor Mass: | 714.84664 | | |
| Mods: | C=160.03 *+15.99 #+220.06 @+291.1 | Charge: | +2 | | |

**Sequence**

IAS@QTQFNTK

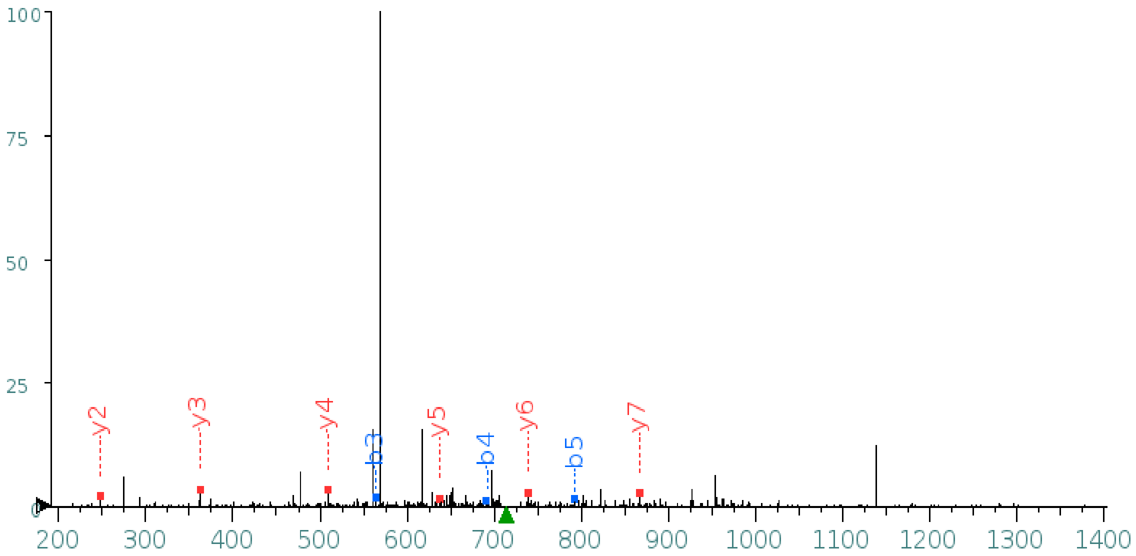

**Predicted Fragmentation Pattern**

| Seq | # | b: Δ Error | b | y | y: Δ Error | +1 |
|---|---|---|---|---|---|---|
| I | 1 | --- | 114.091 | --- | --- | 10 |
| A | 2 | --- | 185.128 | 1315.601 | --- | 9 |
| S@ | 3 | 487.962 | **563.256** | 1244.564 | --- | 8 |
| Q | 4 | -585.027 | **691.314** | **866.437** | 330.751 | 7 |
| T | 5 | -593.447 | **792.362** | **738.378** | 254.575 | 6 |
| Q | 6 | --- | 920.421 | **637.330** | 322.888 | 5 |
| F | 7 | --- | 1067.489 | **509.272** | 219.565 | 4 |
| N | 8 | --- | 1181.532 | **362.203** | 419.302 | 3 |
| T | 9 | --- | 1282.580 | **248.160** | 668.328 | 2 |
| K | 10 | --- | --- | 147.113 | --- | 1 |

| | | | | | |
|---|---|---|---|---|---|
| **Run:** | 77660 | **Scans:** | 18321 - 18321 | **m/z Range:** | 153.00829 to 1170.9732 |
| **Search:** | 77660 | **Precursor Mass:** | 611.81228 | | |
| **Mods:** | C=160.03 *+15.99 #+220.06 @+291.1 | **Charge:** | +2 | | |

**Sequence**

VGS@INIGSGK

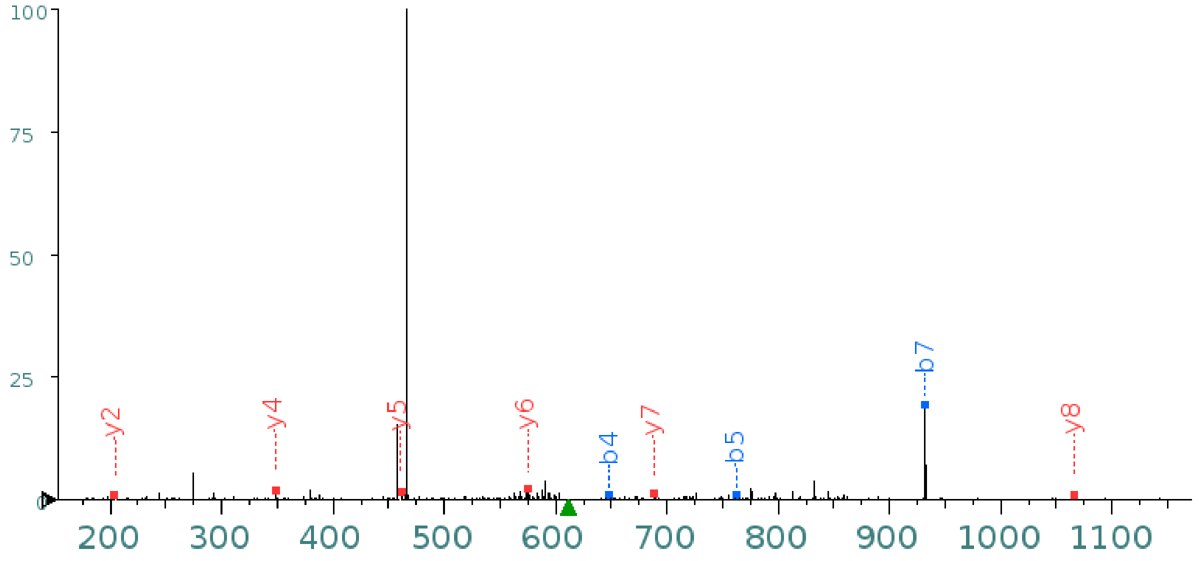

**Predicted Fragmentation Pattern**

| Seq | # | b: Δ Error | b | y | y: Δ Error | +1 |
|---|---|---|---|---|---|---|
| V | 1 | --- | 100.076 | --- | --- | 10 |
| G | 2 | --- | 157.097 | 1123.548 | --- | 9 |
| S@ | 3 | --- | 535.225 | **1066.526** | 221.520 | 8 |
| I | 4 | -1265.611 | **648.309** | **688.399** | 888.123 | 7 |
| N | 5 | -252.619 | **762.352** | **575.315** | 163.681 | 6 |
| I | 6 | --- | 875.436 | **461.272** | 215.691 | 5 |
| G | 7 | -845.065 | **932.457** | **348.188** | 761.725 | 4 |
| S | 8 | --- | 1019.489 | 291.166 | --- | 3 |
| G | 9 | --- | 1076.511 | **204.134** | -4906.653 | 2 |
| K | 10 | --- | --- | 147.113 | --- | 1 |

| Run: | 77660 | Scans: | 18521 - 18521 | m/z Range: | 271.2579 to 1939.3270 |
| --- | --- | --- | --- | --- | --- |
| Search: | 77660 | Precursor Mass: | 1044.48888 | | |
| Mods: | C=160.03 *+15.99 #+220.06 @+291.1 | Charge: | +2 | | |

**Sequence**

DQVM*ELTIAGM*GTS@ALK

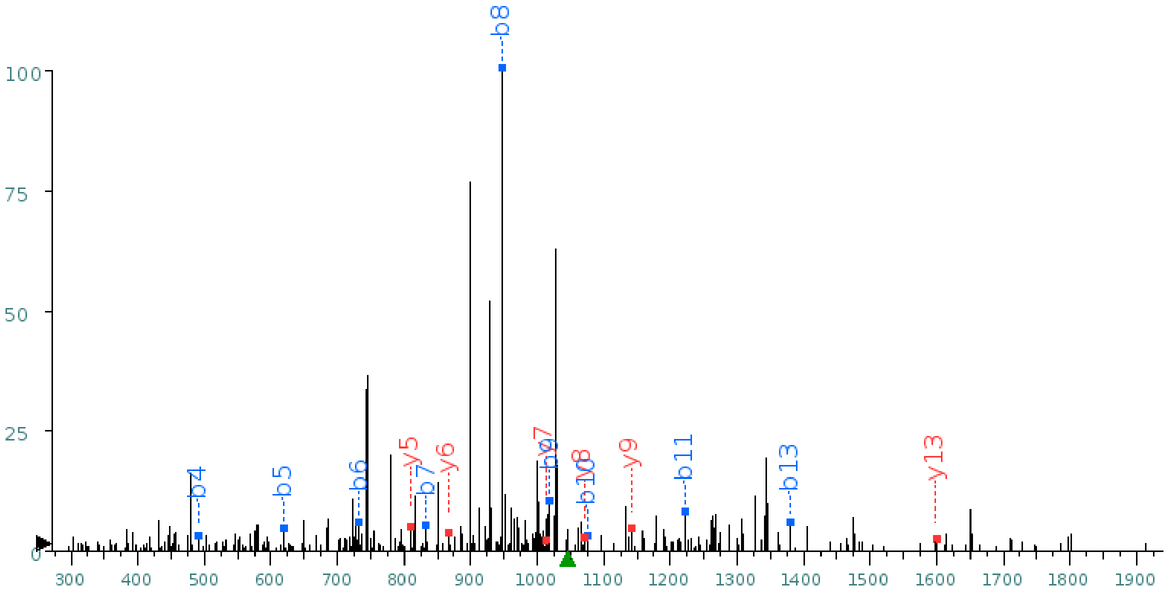

**Predicted Fragmentation Pattern**

| Seq | # | b: Δ Error | b | y | y: Δ Error | +1 |
| --- | --- | --- | --- | --- | --- | --- |
| D | 1 | --- | 116.034 | --- | --- | 17 |
| Q | 2 | --- | 244.093 | 1972.945 | --- | 16 |
| V | 3 | --- | 343.161 | 1844.887 | --- | 15 |
| M* | 4 | 474.444 | **490.197** | 1745.818 | --- | 14 |
| E | 5 | 374.692 | **619.239** | **1598.783** | 297.767 | 13 |
| L | 6 | 284.757 | **732.323** | 1469.740 | --- | 12 |
| T | 7 | -158.336 | **833.371** | 1356.656 | --- | 11 |
| I | 8 | 116.654 | **946.455** | 1255.609 | --- | 10 |
| A | 9 | 76.000 | **1017.492** | **1142.525** | 204.657 | 9 |
| G | 10 | 321.138 | **1074.514** | **1071.487** | -821.292 | 8 |
| M* | 11 | 117.276 | **1221.549** | **1014.466** | 70.118 | 7 |
| G | 12 | --- | 1278.570 | **867.431** | -389.859 | 6 |
| T | 13 | 169.174 | **1379.618** | **810.409** | -1083.800 | 5 |
| S@ | 14 | --- | 1757.746 | 709.361 | --- | 4 |
| A | 15 | --- | 1828.783 | 331.234 | --- | 3 |
| L | 16 | --- | 1941.867 | 260.197 | --- | 2 |
| K | 17 | --- | --- | 147.113 | --- | 1 |

| | | | | | |
|---|---|---|---|---|---|
| **Run:** | 77660 | **Scans:** | 18782 - 18782 | **m/z Range:** | 157.34494 to 1118.5313 |
| **Search:** | 77660 | **Precursor Mass:** | 611.81204 | | |
| **Mods:** | C=160.03 *+15.99 #+220.06 @+291.1 | **Charge:** | +2 | | |

**Sequence**

VGS@INIGSGK

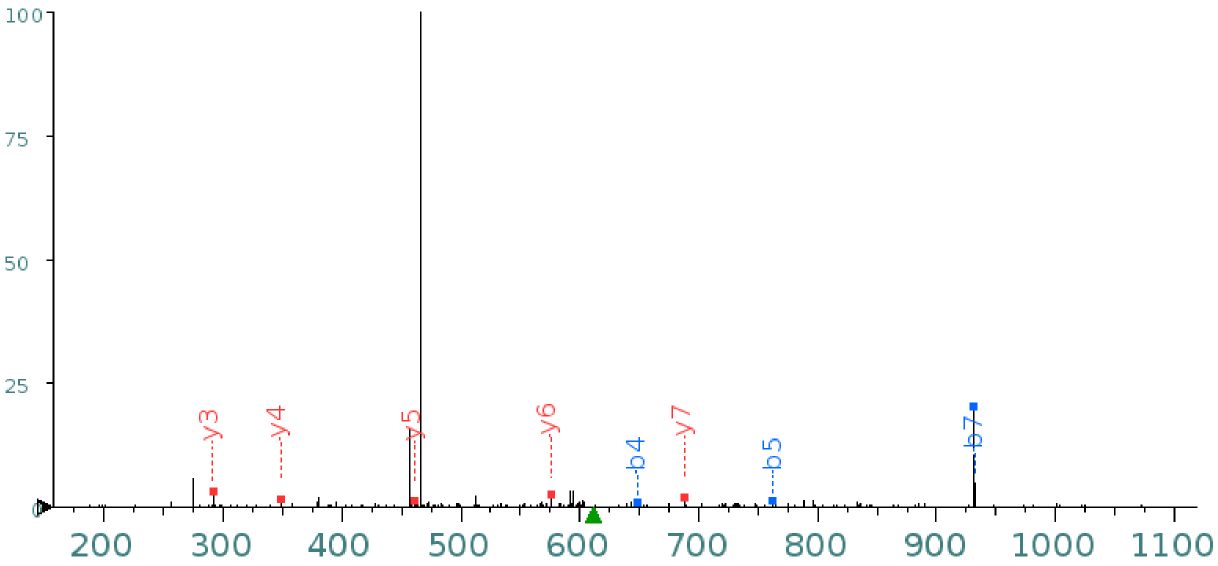

**Predicted Fragmentation Pattern**

| Seq | # | b: Δ Error | b | y | y: Δ Error | +1 |
|---|---|---|---|---|---|---|
| V | 1 | --- | 100.076 | --- | --- | 10 |
| G | 2 | --- | 157.097 | 1123.548 | --- | 9 |
| S@ | 3 | --- | 535.225 | 1066.526 | --- | 8 |
| I | 4 | 315.928 | **648.309** | **688.399** | 92.803 | 7 |
| N | 5 | 300.661 | **762.352** | **575.315** | 8.392 | 6 |
| I | 6 | --- | 875.436 | **461.272** | 279.966 | 5 |
| G | 7 | -981.266 | **932.457** | **348.188** | 378.094 | 4 |
| S | 8 | --- | 1019.489 | **291.166** | 3155.819 | 3 |
| G | 9 | --- | 1076.511 | 204.134 | --- | 2 |
| K | 10 | --- | --- | 147.113 | --- | 1 |

| Run: | 77660 | Scans: | 19225 - 19225 | m/z Range: | 149.15807 to 1223.6244 |
|---|---|---|---|---|---|
| Search: | 77660 | Precursor Mass: | 611.81223 | | |
| Mods: | C=160.03 *+15.99 #+220.06 @+291.1 | Charge: | +2 | | |

**Sequence**

VGS@INIGSGK

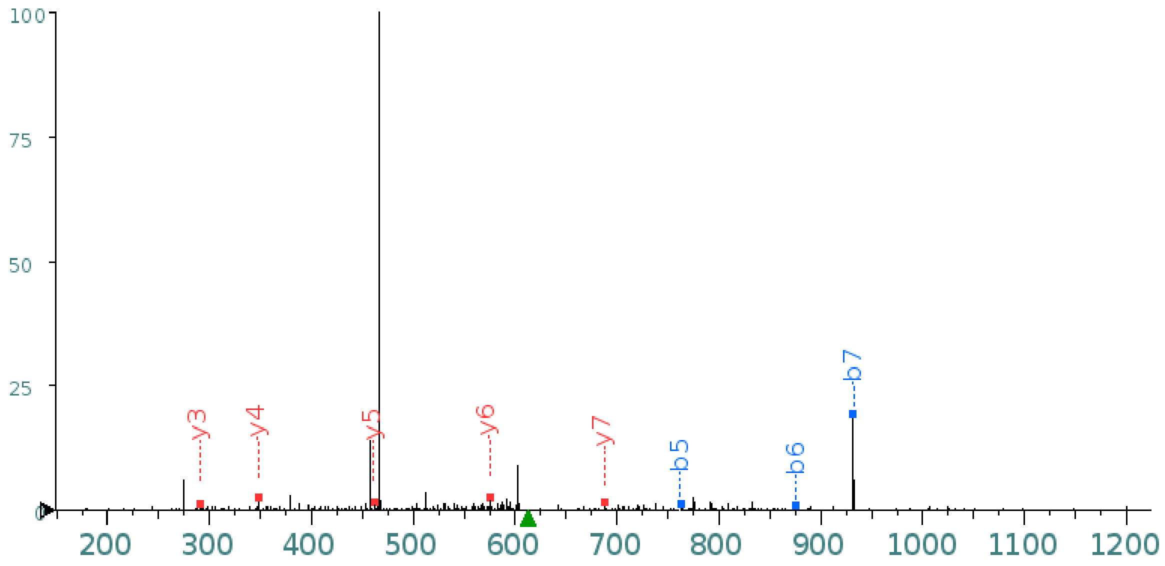

**Predicted Fragmentation Pattern**

| Seq | # | b: Δ Error | b | y | y: Δ Error | +1 |
|---|---|---|---|---|---|---|
| V | 1 | --- | 100.076 | --- | --- | 10 |
| G | 2 | --- | 157.097 | 1123.548 | --- | 9 |
| S@ | 3 | --- | 535.225 | 1066.526 | --- | 8 |
| I | 4 | --- | 648.309 | **688.399** | 261.203 | 7 |
| N | 5 | 773.717 | **762.352** | **575.315** | 399.811 | 6 |
| I | 6 | 362.912 | **875.436** | **461.272** | 491.778 | 5 |
| G | 7 | -838.967 | **932.457** | **348.188** | 547.184 | 4 |
| S | 8 | --- | 1019.489 | **291.166** | 458.318 | 3 |
| G | 9 | --- | 1076.511 | 204.134 | --- | 2 |
| K | 10 | --- | --- | 147.113 | --- | 1 |

| Run: | 77660 | Scans: | 19360 - 19360 | m/z Range: | 148.42674 to 1133.7727 |
|---|---|---|---|---|---|
| Search: | 77660 | Precursor Mass: | 611.81213 | | |
| Mods: | C=160.03 *+15.99 #+220.06 @+291.1 | Charge: | +2 | | |

### Sequence

VGS@INIGSGK

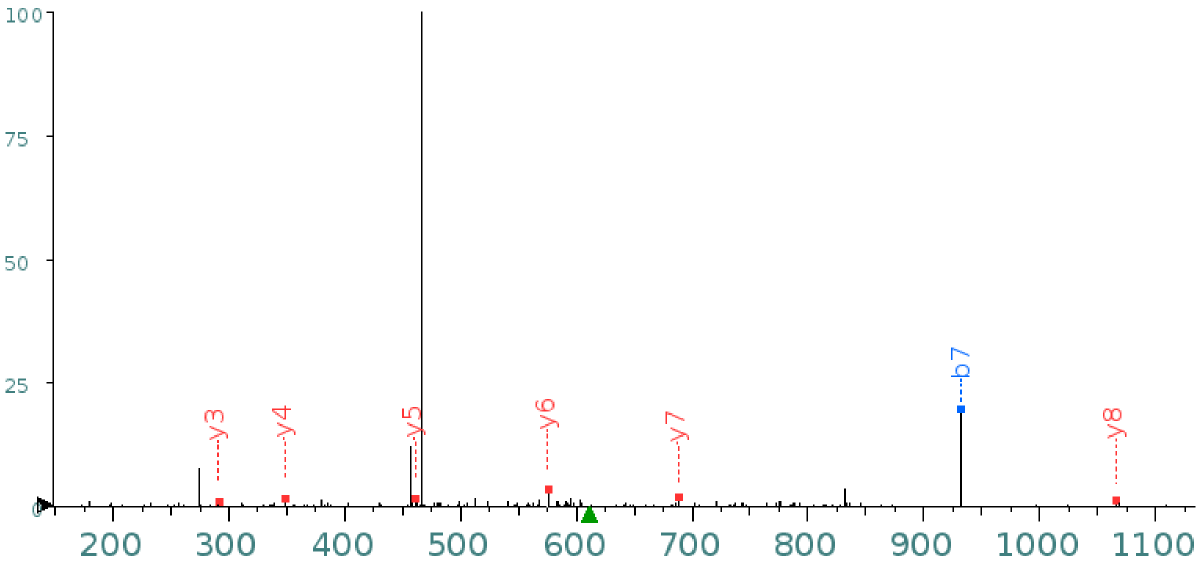

### Predicted Fragmentation Pattern

| Seq | # | b: Δ Error | b | y | y: Δ Error | +1 |
|---|---|---|---|---|---|---|
| V | 1 | --- | 100.076 | --- | --- | 10 |
| G | 2 | --- | 157.097 | 1123.548 | --- | 9 |
| S@ | 3 | --- | 535.225 | **1066.526** | -729.864 | 8 |
| I | 4 | --- | 648.309 | **688.399** | -533.518 | 7 |
| N | 5 | --- | 762.352 | **575.315** | 362.814 | 6 |
| I | 6 | --- | 875.436 | **461.272** | 645.226 | 5 |
| G | 7 | -963.034 | **932.457** | **348.188** | 405.769 | 4 |
| S | 8 | --- | 1019.489 | **291.166** | 449.732 | 3 |
| G | 9 | --- | 1076.511 | 204.134 | --- | 2 |
| K | 10 | --- | --- | 147.113 | --- | 1 |

| Run: | 77660 | Scans: | 19609 - 19609 | m/z Range: | 150.21429 to 1223.6241[illegible] |
|---|---|---|---|---|---|
| Search: | 77660 | Precursor Mass: | 611.81206 | | |
| Mods: | C=160.03 *+15.99 #+220.06 @+291.1 | Charge: | +2 | | |

**Sequence**

VGS@INIGSGK

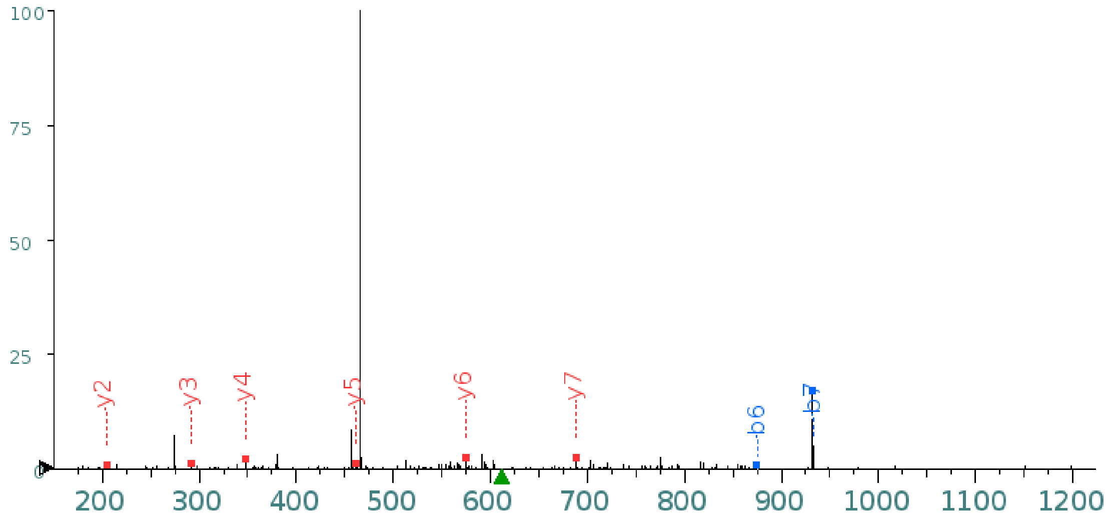

**Predicted Fragmentation Pattern**

| Seq | # | b: Δ Error | b | y | y: Δ Error | +1 |
|---|---|---|---|---|---|---|
| V | 1 | --- | 100.076 | --- | --- | 10 |
| G | 2 | --- | 157.097 | 1123.548 | --- | 9 |
| S@ | 3 | --- | 535.225 | 1066.526 | --- | 8 |
| I | 4 | --- | 648.309 | **688.399** | 282.559 | 7 |
| N | 5 | --- | 762.352 | **575.315** | 373.309 | 6 |
| I | 6 | -823.706 | **875.436** | **461.272** | 255.765 | 5 |
| G | 7 | -952.803 | **932.457** | **348.188** | 556.377 | 4 |
| S | 8 | --- | 1019.489 | **291.166** | 173.830 | 3 |
| G | 9 | --- | 1076.511 | **204.134** | 1687.412 | 2 |
| K | 10 | --- | --- | 147.113 | --- | 1 |

| Run: | 77660 | Scans: | 19723 - 19723 | m/z Range: | 154.17503 to 1223.6240 |
|---|---|---|---|---|---|
| Search: | 77660 | Precursor Mass: | 611.81203 | | |
| Mods: | C=160.03 *+15.99 #+220.06 @+291.1 | Charge: | +2 | | |

**Sequence**

VGS@INIGSGK

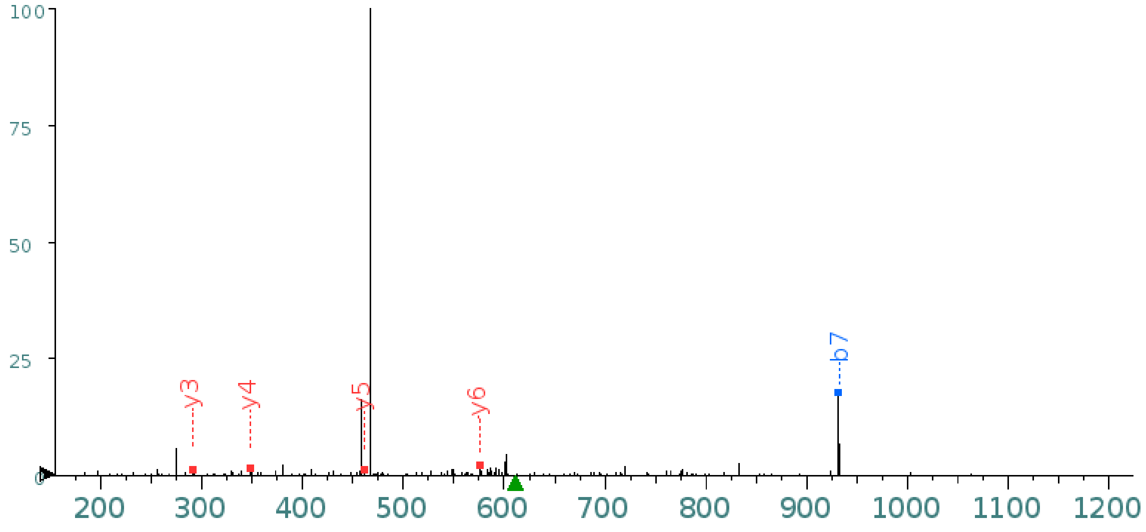

**Predicted Fragmentation Pattern**

| Seq | # | b: Δ Error | b | y | y: Δ Error | +1 |
|---|---|---|---|---|---|---|
| V | 1 | --- | 100.076 | --- | --- | 10 |
| G | 2 | --- | 157.097 | 1123.548 | --- | 9 |
| S@ | 3 | --- | 535.225 | 1066.526 | --- | 8 |
| I | 4 | --- | 648.309 | 688.399 | --- | 7 |
| N | 5 | --- | 762.352 | **575.315** | 512.270 | 6 |
| I | 6 | --- | 875.436 | **461.272** | 398.179 | 5 |
| G | 7 | -867.686 | **932.457** | **348.188** | 683.745 | 4 |
| S | 8 | --- | 1019.489 | **291.166** | 642.270 | 3 |
| G | 9 | --- | 1076.511 | 204.134 | --- | 2 |
| K | 10 | --- | --- | 147.113 | --- | 1 |

| | | | | | |
|---|---|---|---|---|---|
| **Run:** | 77660 | **Scans:** | 19839 - 19839 | **m/z Range:** | 154.16147 to 1088.7208[illegible] |
| **Search:** | 77660 | **Precursor Mass:** | 611.81187 | | |
| **Mods:** | C=160.03 *+15.99 #+220.06 @+291.1 | **Charge:** | +2 | | |

**Sequence**

VGS@INIGSGK

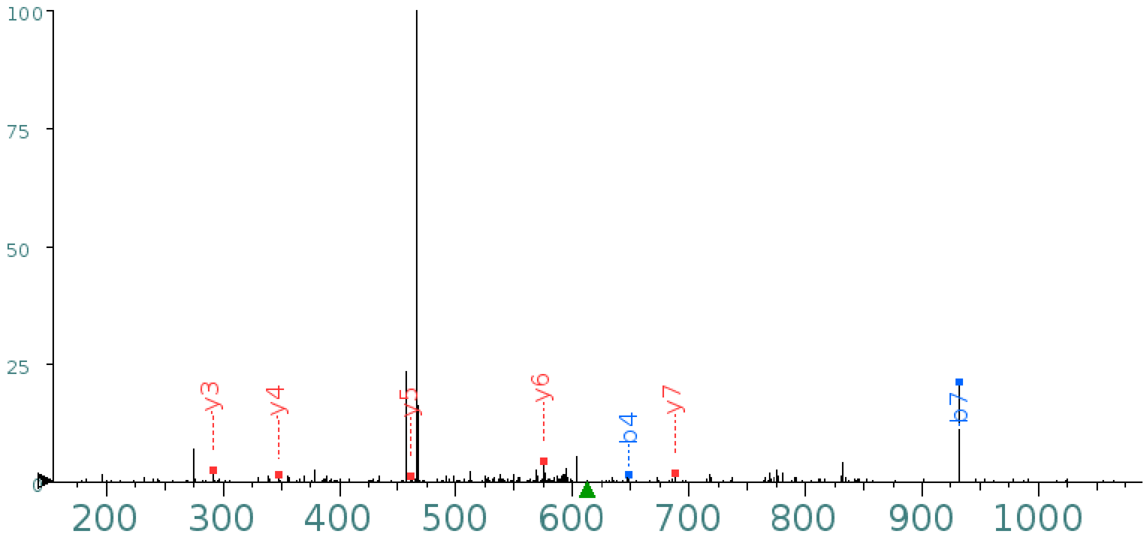

**Predicted Fragmentation Pattern**

| Seq | # | b: Δ Error | b | y | y: Δ Error | +1 |
|---|---|---|---|---|---|---|
| V | 1 | --- | 100.076 | --- | --- | 10 |
| G | 2 | --- | 157.097 | 1123.548 | --- | 9 |
| S@ | 3 | --- | 535.225 | 1066.526 | --- | 8 |
| I | 4 | -428.844 | **648.309** | **688.399** | 211.309 | 7 |
| N | 5 | --- | 762.352 | **575.315** | 155.621 | 6 |
| I | 6 | --- | 875.436 | **461.272** | 283.008 | 5 |
| G | 7 | -942.113 | **932.457** | **348.188** | -69.906 | 4 |
| S | 8 | --- | 1019.489 | **291.166** | 3335.135 | 3 |
| G | 9 | --- | 1076.511 | 204.134 | --- | 2 |
| K | 10 | --- | --- | 147.113 | --- | 1 |

| | | | | | | |
|---|---|---|---|---|---|---|
| **Run:** | 77660 | **Scans:** | 20318 - 20318 | **m/z Range:** | 306.30334 | to 1808.0997 |
| **Search:** | 77660 | **Precursor Mass:** | 1036.49177 | | | |
| **Mods:** | C=160.03 *+15.99 #+220.06 @+291.1 | **Charge:** | +2 | | | |

### Sequence

DQVMELTIAGM*GTS@ALK

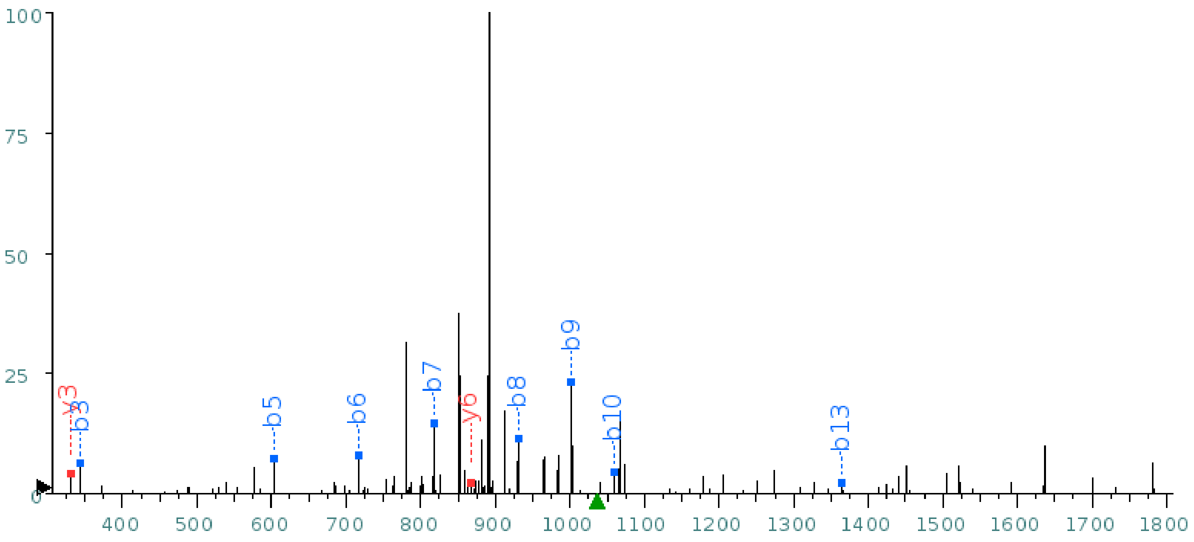

### Predicted Fragmentation Pattern

| Seq | # | b: Δ Error | b | y | y: Δ Error | +1 |
|---|---|---|---|---|---|---|
| D | 1 | --- | 116.034 | --- | --- | 17 |
| Q | 2 | --- | 244.093 | 1956.950 | --- | 16 |
| V | 3 | 880.923 | **343.161** | 1828.892 | --- | 15 |
| M | 4 | --- | 474.202 | 1729.823 | --- | 14 |
| E | 5 | 103.250 | **603.244** | 1598.783 | --- | 13 |
| L | 6 | 356.226 | **716.328** | 1469.740 | --- | 12 |
| T | 7 | 162.394 | **817.376** | 1356.656 | --- | 11 |
| I | 8 | 345.039 | **930.460** | 1255.609 | --- | 10 |
| A | 9 | 117.470 | **1001.497** | 1142.525 | --- | 9 |
| G | 10 | 92.599 | **1058.519** | 1071.487 | --- | 8 |
| M* | 11 | --- | 1205.554 | 1014.466 | --- | 7 |
| G | 12 | --- | 1262.576 | **867.431** | 249.269 | 6 |
| T | 13 | 46.157 | **1363.623** | 810.409 | --- | 5 |
| S@ | 14 | --- | 1741.751 | 709.361 | --- | 4 |
| A | 15 | --- | 1812.788 | **331.234** | 209.362 | 3 |
| L | 16 | --- | 1925.872 | 260.197 | --- | 2 |
| K | 17 | --- | --- | 147.113 | --- | 1 |

| | | | | | |
|---|---|---|---|---|---|
| **Run:** | 77660 | **Scans:** | 20484 - 20484 | **m/z Range:** | 218.42632 to 1691.3475[illegible] |
| **Search:** | 77660 | **Precursor Mass:** | 825.40388 | | |
| **Mods:** | C=160.03 *+15.99 #+220.06 @+291.1 | **Charge:** | +3 | | |

#### Sequence

NLLNGS@AKS@MVFQIGANK

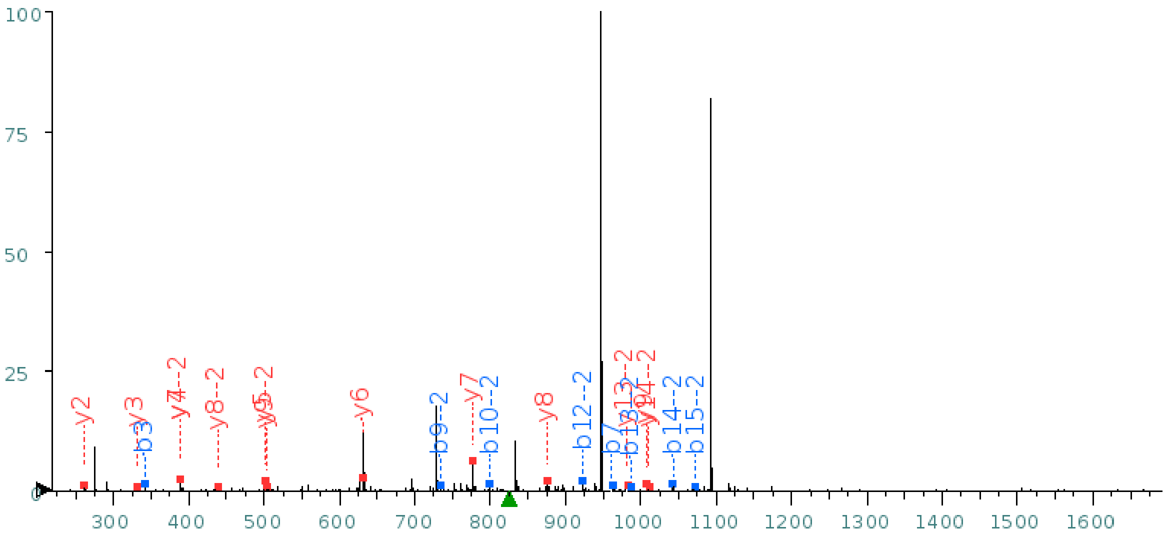

#### Predicted Fragmentation Pattern

+1

| Seq | # | b: Δ Error | b | y | y: Δ Error | +1 |
|---|---|---|---|---|---|---|
| N | 1 | --- | 115.050 | --- | --- | 18 |
| L | 2 | --- | 228.134 | 2360.154 | --- | 17 |
| L | 3 | 641.364 | **341.218** | 2247.070 | --- | 16 |
| N | 4 | --- | 455.261 | 2133.986 | --- | 15 |
| G | 5 | --- | 512.283 | 2019.943 | --- | 14 |
| S@ | 6 | --- | 890.410 | 1962.921 | --- | 13 |
| A | 7 | 690.004 | **961.447** | 1584.794 | --- | 12 |
| K | 8 | --- | 1089.542 | 1513.757 | --- | 11 |
| S@ | 9 | --- | 1467.670 | 1385.662 | --- | 10 |
| M | 10 | --- | 1598.710 | **1007.534** | 257.613 | 9 |
| V | 11 | --- | 1697.779 | **876.494** | 247.975 | 8 |
| F | 12 | --- | 1844.847 | **777.425** | -782.736 | 7 |
| Q | 13 | --- | 1972.906 | **630.357** | 429.310 | 6 |
| I | 14 | --- | 2085.990 | **502.298** | 470.448 | 5 |
| G | 15 | --- | 2143.011 | **389.214** | -105.979 | 4 |
| A | 16 | --- | 2214.048 | **332.193** | 642.365 | 3 |
| N | 17 | --- | 2328.091 | **261.156** | -3516.247 | 2 |
| K | 18 | --- | --- | 147.113 | --- | 1 |

| Run: | 77660 | Scans: | 20492 - 20492 | m/z Range: | 280.18811 to 1821.4140 |
|---|---|---|---|---|---|
| Search: | 77660 | Precursor Mass: | 1000.97352 | | |
| Mods: | C=160.03 *+15.99 #+220.06 @+291.1 | Charge: | +2 | | |

**Sequence**

DQVM*ELTIAGMGT#SALK

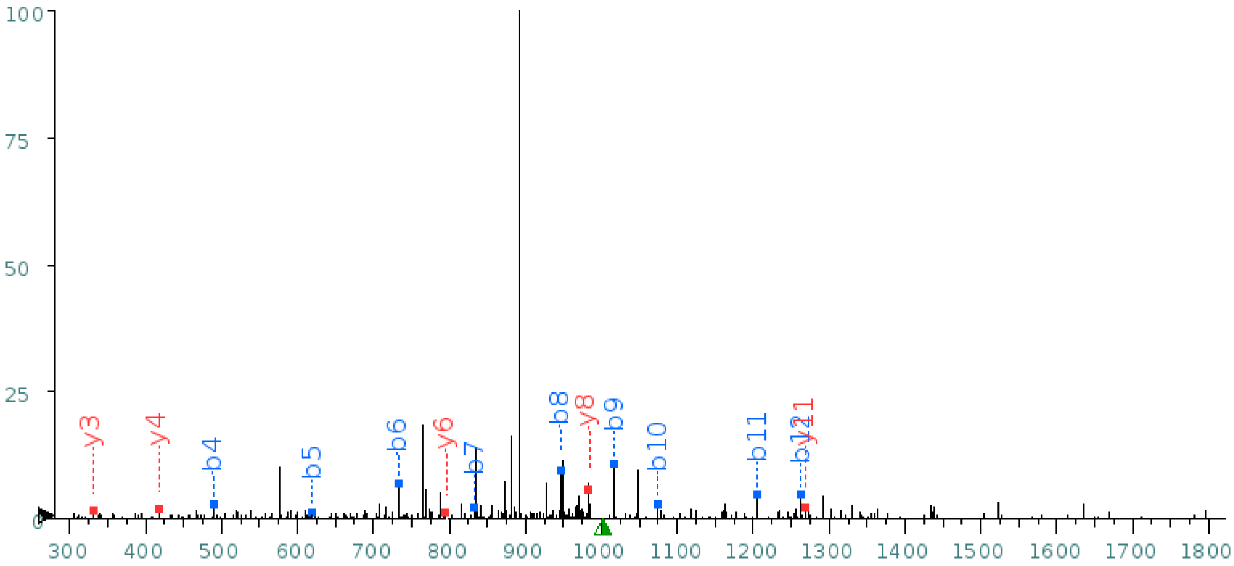

**Predicted Fragmentation Pattern**

| Seq | # | b: Δ Error | b | y | y: Δ Error | +1 |
|---|---|---|---|---|---|---|
| D | 1 | --- | 116.034 | --- | --- | 17 |
| Q | 2 | --- | 244.093 | 1885.913 | --- | 16 |
| V | 3 | --- | 343.161 | 1757.855 | --- | 15 |
| M* | 4 | 432.770 | **490.197** | 1658.786 | --- | 14 |
| E | 5 | -1316.693 | **619.239** | 1511.751 | --- | 13 |
| L | 6 | 37.303 | **732.323** | 1382.708 | --- | 12 |
| T | 7 | 240.198 | **833.371** | **1269.624** | -621.775 | 11 |
| I | 8 | 142.829 | **946.455** | 1168.577 | --- | 10 |
| A | 9 | 134.773 | **1017.492** | 1055.493 | --- | 9 |
| G | 10 | 199.644 | **1074.514** | **984.455** | -680.299 | 8 |
| M | 11 | 186.788 | **1205.554** | 927.434 | --- | 7 |
| G | 12 | 175.278 | **1262.576** | **796.393** | -1109.615 | 6 |
| T# | 13 | --- | 1583.682 | 739.372 | --- | 5 |
| S | 14 | --- | 1670.714 | **418.266** | 676.438 | 4 |
| A | 15 | --- | 1741.751 | **331.234** | 757.484 | 3 |
| L | 16 | --- | 1854.835 | 260.197 | --- | 2 |
| K | 17 | --- | --- | 147.113 | --- | 1 |

| Run: | 77660 | Scans: | 20617 - 20617 | m/z Range: | 309.32706 to 1950.005 |
|---|---|---|---|---|---|
| Search: | 77660 | Precursor Mass: | 1060.50908 | | |
| Mods: | C=160.03 *+15.99 #+220.06 @+291.1 | Charge: | +3 | | |

**Sequence**

S@M*VFQIGANKDQVM*ELTIAGM*GTSALK

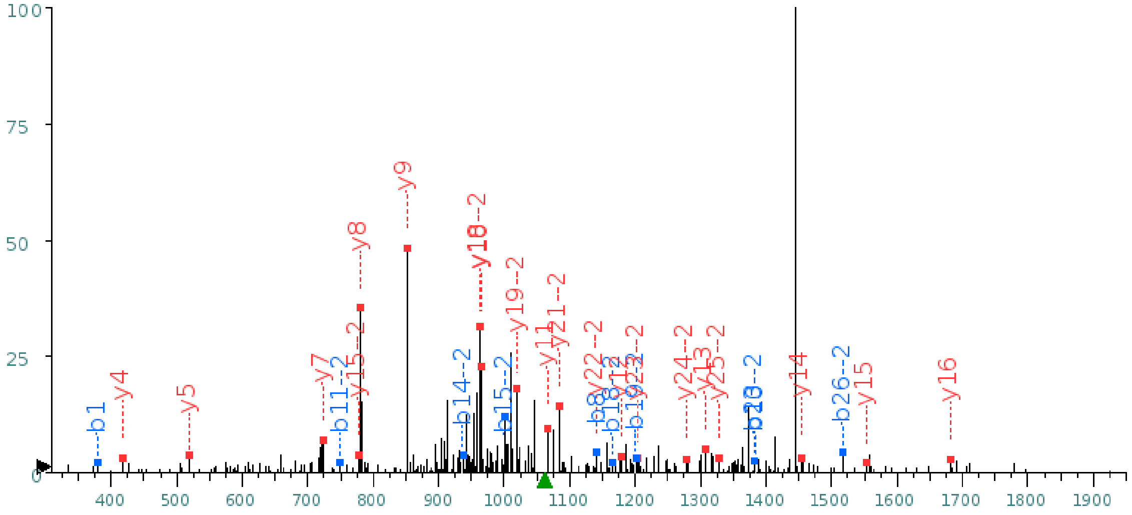

**Predicted Fragmentation Pattern**

+1

| Seq | # | b: Δ Error | b | y | y: Δ Error | +1 |
|---|---|---|---|---|---|---|
| S@ | 1 | 463.532 | **379.135** | --- | --- | 27 |
| M* | 2 | --- | 526.170 | 2801.388 | --- | 26 |
| V | 3 | --- | 625.239 | 2654.353 | --- | 25 |
| F | 4 | --- | 772.307 | 2555.284 | --- | 24 |
| Q | 5 | --- | 900.366 | 2408.216 | --- | 23 |
| I | 6 | --- | 1013.450 | 2280.157 | --- | 22 |
| G | 7 | --- | 1070.471 | 2167.073 | --- | 21 |
| A | 8 | -453.722 | **1141.508** | 2110.052 | --- | 20 |
| N | 9 | --- | 1255.551 | 2039.015 | --- | 19 |
| K | 10 | -493.460 | **1383.646** | 1924.972 | --- | 18 |
| D | 11 | --- | 1498.673 | 1796.877 | --- | 17 |
| Q | 12 | --- | 1626.732 | **1681.850** | 138.136 | 16 |
| V | 13 | --- | 1725.800 | **1553.791** | 18.880 | 15 |
| M* | 14 | --- | 1872.835 | **1454.723** | 115.851 | 14 |
| E | 15 | --- | 2001.878 | **1307.688** | 347.766 | 13 |
| L | 16 | --- | 2114.962 | **1178.645** | 127.237 | 12 |
| T | 17 | --- | 2216.010 | **1065.561** | 190.057 | 11 |
| I | 18 | --- | 2329.094 | **964.513** | -399.472 | 10 |
| A | 19 | --- | 2400.131 | **851.429** | 64.021 | 9 |

**Run:** 77660 **Scans:** 20867 - 20867 **m/z Range:** 306.4469 to 1955.2884

**Search:** 77660 **Precursor Mass:** 1152.20792

**Mods:** C=160.03 *+15.99 #+220.06 @+291.1 **Charge:** +3

### Sequence

S@MVFQIGANKDQVM*ELT@IAGM*GTSALK

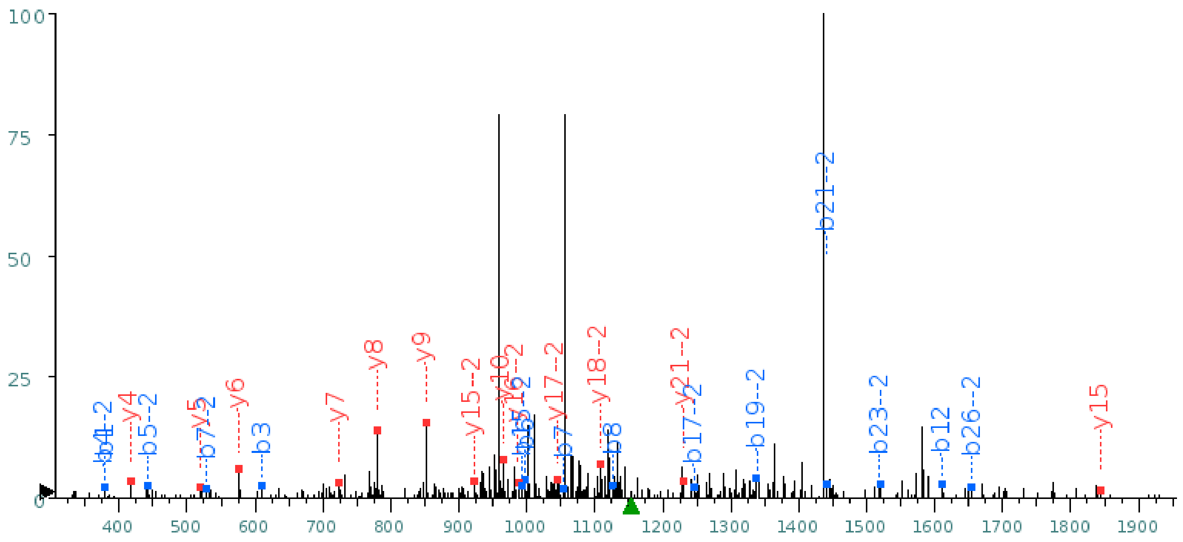

### Predicted Fragmentation Pattern

+1

| Seq | # | b: Δ Error | b | y | y: Δ Error | +1 |
|---|---|---|---|---|---|---|
| S@ | 1 | 93.472 | **379.135** | --- | --- | 27 |
| M | 2 | --- | 510.175 | 3076.489 | --- | 26 |
| V | 3 | 165.722 | **609.244** | 2945.448 | --- | 25 |
| F | 4 | --- | 756.312 | 2846.380 | --- | 24 |
| Q | 5 | --- | 884.371 | 2699.311 | --- | 23 |
| I | 6 | -688.225 | **997.455** | 2571.253 | --- | 22 |
| G | 7 | -172.467 | **1054.476** | 2458.169 | --- | 21 |
| A | 8 | 515.060 | **1125.513** | 2401.147 | --- | 20 |
| N | 9 | --- | 1239.556 | 2330.110 | --- | 19 |
| K | 10 | --- | 1367.651 | 2216.067 | --- | 18 |
| D | 11 | --- | 1482.678 | 2087.972 | --- | 17 |
| Q | 12 | 497.225 | **1610.737** | 1972.945 | --- | 16 |
| V | 13 | --- | 1709.805 | **1844.887** | -428.240 | 15 |
| M* | 14 | --- | 1856.840 | 1745.818 | --- | 14 |
| E | 15 | --- | 1985.883 | 1598.783 | --- | 13 |
| L | 16 | --- | 2098.967 | 1469.740 | --- | 12 |
| T@ | 17 | --- | 2491.110 | 1356.656 | --- | 11 |
| I | 18 | --- | 2604.194 | **964.513** | 234.327 | 10 |
| A | 19 | --- | 2675.231 | **851.429** | 280.437 | 9 |

| | | | | | |
|---|---|---|---|---|---|
| **Run:** | 77660 | **Scans:** | 21014 - 21014 | **m/z Range:** | 267.06055 to 1808.2968 |
| **Search:** | 77660 | **Precursor Mass:** | 1036.49183 | | |
| **Mods:** | C=160.03 *+15.99 #+220.06 @+291.1 | **Charge:** | +2 | | |

**Sequence**

DQVM*ELT@IAGMGTSALK

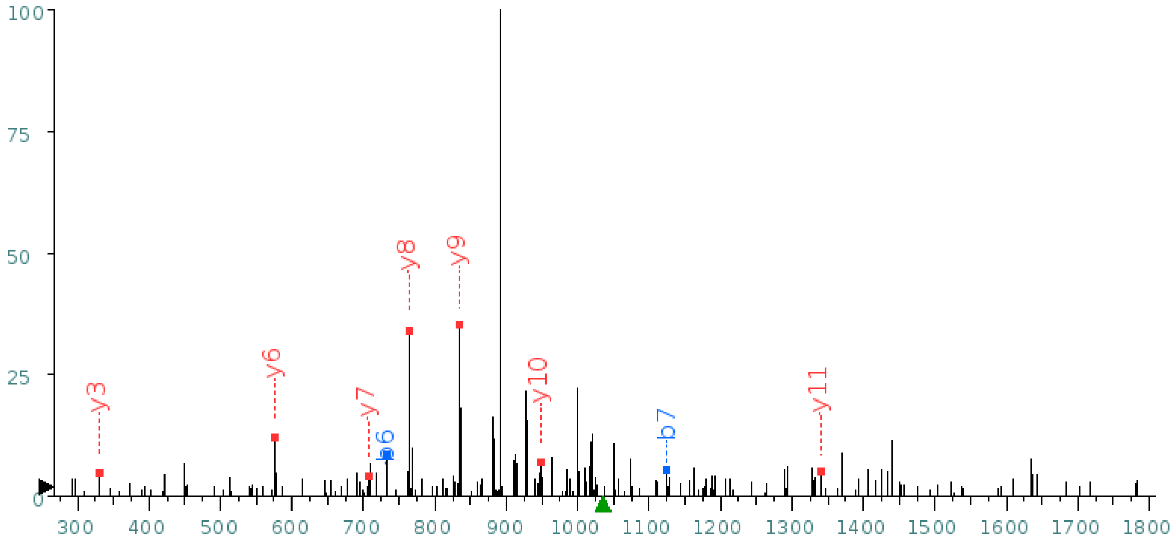

**Predicted Fragmentation Pattern**

| Seq | # | b: Δ Error | b | y | y: Δ Error | +1 |
|---|---|---|---|---|---|---|
| D | 1 | --- | 116.034 | --- | --- | 17 |
| Q | 2 | --- | 244.093 | 1956.950 | --- | 16 |
| V | 3 | --- | 343.161 | 1828.892 | --- | 15 |
| M* | 4 | --- | 490.197 | 1729.823 | --- | 14 |
| E | 5 | --- | 619.239 | 1582.788 | --- | 13 |
| L | 6 | **295.752** | **732.323** | 1453.745 | --- | 12 |
| T@ | 7 | **142.901** | **1124.466** | **1340.661** | **450.415** | 11 |
| I | 8 | --- | 1237.550 | **948.518** | **185.450** | 10 |
| A | 9 | --- | 1308.588 | **835.434** | **42.797** | 9 |
| G | 10 | --- | 1365.609 | **764.397** | **206.594** | 8 |
| M | 11 | --- | 1496.649 | **707.376** | **428.092** | 7 |
| G | 12 | --- | 1553.671 | **576.335** | **458.225** | 6 |
| T | 13 | --- | 1654.719 | 519.314 | --- | 5 |
| S | 14 | --- | 1741.751 | 418.266 | --- | 4 |
| A | 15 | --- | 1812.788 | **331.234** | **429.051** | 3 |
| L | 16 | --- | 1925.872 | 260.197 | --- | 2 |
| K | 17 | --- | --- | 147.113 | --- | 1 |

| Run: | 77660 | Scans: | 21127 - 21127 | m/z Range: | 238.21268 to 1980.3721 |
|---|---|---|---|---|---|
| Search: | 77660 | Precursor Mass: | 890.76361 | | |
| Mods: | C=160.03 *+15.99 #+220.06 @+291.1 | Charge: | +3 | | |

**Sequence**

GAKSIS@GM*IETIDNAINTVSGER

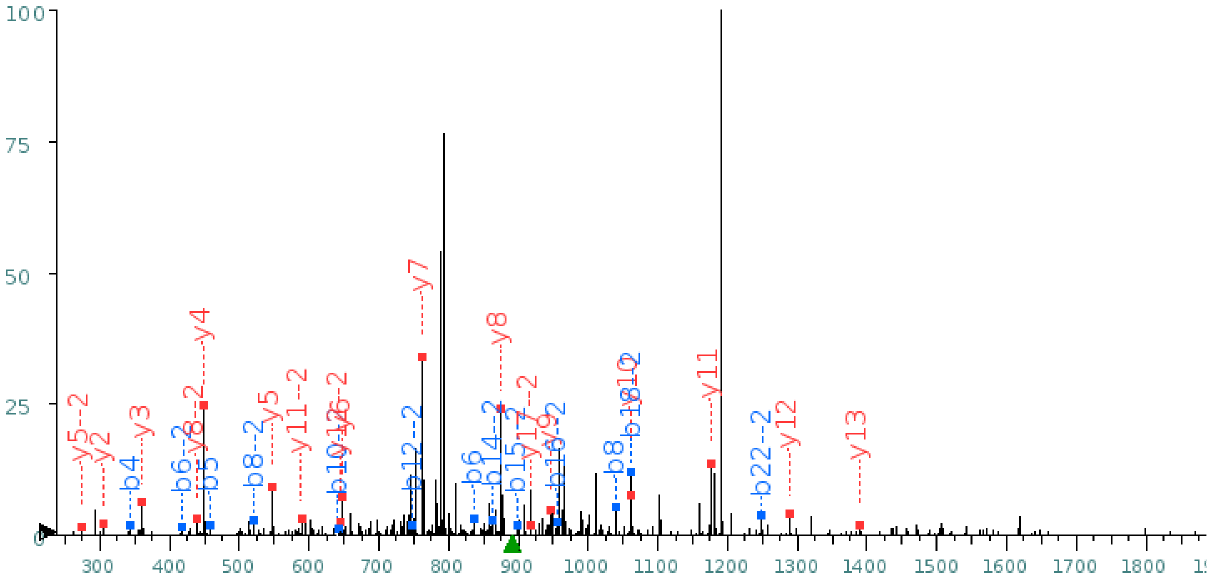

**Predicted Fragmentation Pattern**

+1

| Seq | # | b: Δ Error | b | y | y: Δ Error | +1 |
|---|---|---|---|---|---|---|
| G | 1 | --- | 58.029 | --- | --- | 23 |
| A | 2 | --- | 129.066 | 2613.256 | --- | 22 |
| K | 3 | --- | 257.161 | 2542.219 | --- | 21 |
| S | 4 | -2237.929 | **344.193** | 2414.124 | --- | 20 |
| I | 5 | 221.937 | **457.277** | 2327.092 | --- | 19 |
| S@ | 6 | 261.781 | **835.404** | 2214.008 | --- | 18 |
| G | 7 | --- | 892.426 | 1835.880 | --- | 17 |
| M* | 8 | -612.724 | **1039.461** | 1778.859 | --- | 16 |
| I | 9 | --- | 1152.545 | 1631.824 | --- | 15 |
| E | 10 | --- | 1281.588 | 1518.739 | --- | 14 |
| T | 11 | --- | 1382.636 | **1389.697** | 218.885 | 13 |
| I | 12 | --- | 1495.720 | **1288.649** | 276.054 | 12 |
| D | 13 | --- | 1610.747 | **1175.565** | 96.318 | 11 |
| N | 14 | --- | 1724.789 | **1060.538** | 75.314 | 10 |
| A | 15 | --- | 1795.827 | **946.495** | 245.762 | 9 |
| I | 16 | --- | 1908.911 | **875.458** | 34.519 | 8 |
| N | 17 | --- | 2022.954 | **762.374** | 92.495 | 7 |

| | | | | | |
|---|---|---|---|---|---|
| **Run:** | 77660 | **Scans:** | 21837 - 21837 | **m/z Range:** | 280.39911 to 1823.0761 |
| **Search:** | 77660 | **Precursor Mass:** | 1055.178 | | |
| **Mods:** | C=160.03 *+15.99 #+220.06 @+291.1 | **Charge:** | +3 | | |

**Sequence**

SM*VFQIGANKDQVMELT@IAGM*GTSALK

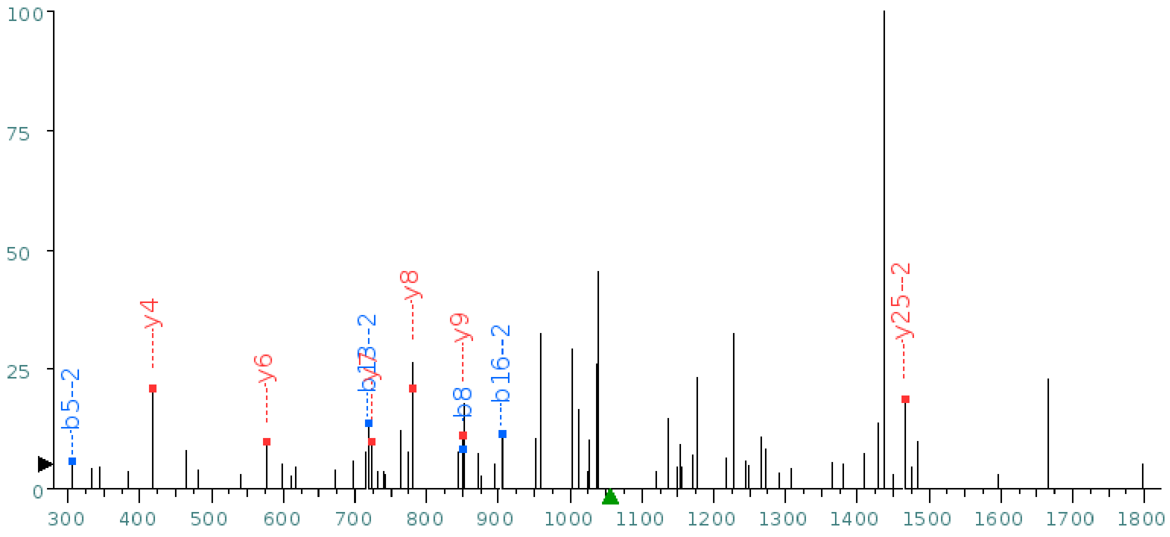

**Predicted Fragmentation Pattern**

+1

| Seq | # | b: Δ Error | b | y | y: Δ Error | +1 |
|---|---|---|---|---|---|---|
| S | 1 | --- | 88.039 | --- | --- | 27 |
| M* | 2 | --- | 235.075 | 3076.489 | --- | 26 |
| V | 3 | --- | 334.143 | 2929.453 | --- | 25 |
| F | 4 | --- | 481.212 | 2830.385 | --- | 24 |
| Q | 5 | --- | 609.270 | 2683.316 | --- | 23 |
| I | 6 | --- | 722.354 | 2555.258 | --- | 22 |
| G | 7 | --- | 779.376 | 2442.174 | --- | 21 |
| A | 8 | 199.735 | **850.413** | 2385.152 | --- | 20 |
| N | 9 | --- | 964.456 | 2314.115 | --- | 19 |
| K | 10 | --- | 1092.551 | 2200.072 | --- | 18 |
| D | 11 | --- | 1207.578 | 2071.977 | --- | 17 |
| Q | 12 | --- | 1335.636 | 1956.950 | --- | 16 |
| V | 13 | --- | 1434.705 | 1828.892 | --- | 15 |
| M | 14 | --- | 1565.745 | 1729.823 | --- | 14 |
| E | 15 | --- | 1694.788 | 1598.783 | --- | 13 |
| L | 16 | --- | 1807.872 | 1469.740 | --- | 12 |
| T@ | 17 | --- | 2200.015 | 1356.656 | --- | 11 |
| I | 18 | --- | 2313.099 | 964.513 | --- | 10 |
| A | 19 | --- | 2384.136 | **851.429** | 10.261 | 9 |
| G | 20 | --- | 2441.157 | **780.392** | 222.164 | 8 |

| Run: | 77660 | Scans: | 22446 - 22446 | m/z Range: | 274.33762 to 1952.1652 |
|---|---|---|---|---|---|
| Search: | 77660 | Precursor Mass: | 1055.17717 | | |
| Mods: | C=160.03 *+15.99 #+220.06 @+291.1 | Charge: | +3 | | |

**Sequence**

S@M*VFQIGANKDQVM*ELTIAGMGTSALK

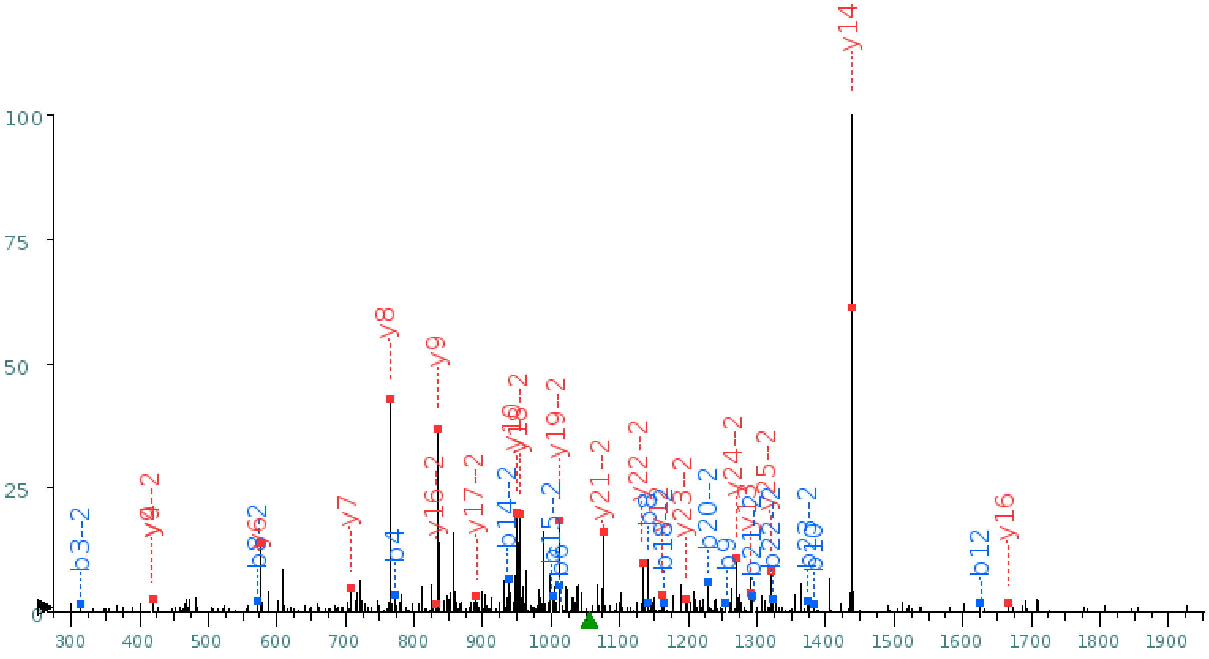

**Predicted Fragmentation Pattern**

| Seq | # | b: Δ Error | b | y | y: Δ Error | +1 |
|---|---|---|---|---|---|---|
| S@ | 1 | --- | 379.135 | --- | --- | 27 |
| M* | 2 | --- | 526.170 | 2785.393 | --- | 26 |
| V | 3 | --- | 625.239 | 2638.358 | --- | 25 |
| F | 4 | **332.992** | **772.307** | 2539.289 | --- | 24 |
| Q | 5 | --- | 900.366 | 2392.221 | --- | 23 |
| I | 6 | **-857.988** | **1013.450** | 2264.162 | --- | 22 |
| G | 7 | --- | 1070.471 | 2151.078 | --- | 21 |
| A | 8 | **-404.167** | **1141.508** | 2094.057 | --- | 20 |
| N | 9 | **-715.071** | **1255.551** | 2023.020 | --- | 19 |
| K | 10 | **-565.439** | **1383.646** | 1908.977 | --- | 18 |
| D | 11 | --- | 1498.673 | 1780.882 | --- | 17 |
| Q | 12 | **-574.697** | **1626.732** | **1665.855** | **66.883** | 16 |
| V | 13 | --- | 1725.800 | 1537.796 | --- | 15 |
| M* | 14 | --- | 1872.835 | **1438.728** | **-509.241** | 14 |
| E | 15 | --- | 2001.878 | **1291.693** | **129.652** | 13 |
| L | 16 | --- | 2114.962 | **1162.650** | **129.757** | 12 |
| T | 17 | --- | 2216.010 | 1049.566 | --- | 11 |
| I | 18 | --- | 2329.094 | **948.518** | **192.911** | 10 |
| A | 19 | --- | 2400.131 | **835.434** | **96.122** | 9 |

| | | | |
|---|---|---|---|
| **Run:** | 77660 | **Scans:** | 22453 - 22453 |
| **Search:** | 77660 | **Precursor Mass:** | 781.70001 |
| **Mods:** | C=160.03 *+15.99 #+220.06 @+291.1 | **Charge:** | +3 |

**m/z Range:** 218.09686 **to** 1921.2325

**Sequence**

S#ISGM*IETIDNAINTVSGER

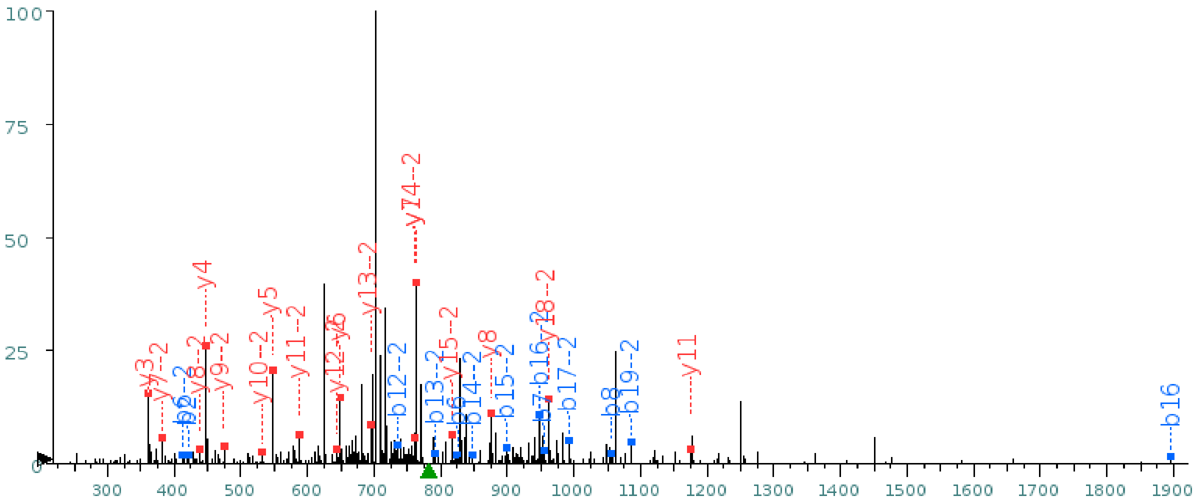

**Predicted Fragmentation Pattern**

| Seq | # | b: Δ Error | b | y | y: Δ Error | +1 |
|---|---|---|---|---|---|---|
| S# | 1 | --- | 308.098 | --- | --- | 20 |
| I | 2 | -106.444 | **421.182** | 2035.996 | --- | 19 |
| S | 3 | --- | 508.214 | 1922.912 | --- | 18 |
| G | 4 | --- | 565.235 | 1835.880 | --- | 17 |
| M* | 5 | --- | 712.271 | 1778.859 | --- | 16 |
| I | 6 | -378.780 | **825.355** | 1631.824 | --- | 15 |
| E | 7 | -323.060 | **954.397** | 1518.739 | --- | 14 |
| T | 8 | -635.779 | **1055.445** | 1389.697 | --- | 13 |
| I | 9 | --- | 1168.529 | 1288.649 | --- | 12 |
| D | 10 | --- | 1283.556 | **1175.565** | 175.422 | 11 |
| N | 11 | --- | 1397.599 | 1060.538 | --- | 10 |
| A | 12 | --- | 1468.636 | 946.495 | --- | 9 |
| I | 13 | --- | 1581.720 | **875.458** | -21.394 | 8 |
| N | 14 | --- | 1695.763 | **762.374** | 12.604 | 7 |
| T | 15 | --- | 1796.811 | **648.331** | 27.184 | 6 |
| V | 16 | 186.431 | **1895.879** | **547.283** | 325.313 | 5 |

| | | | | | |
|---|---|---|---|---|---|
| **Run:** | 77660 | **Scans:** | 22533 - 22533 | **m/z Range:** | 313.26563 to 1684.0655 |
| **Search:** | 77660 | **Precursor Mass:** | 992.97496 | | |
| **Mods:** | C=160.03 *+15.99 #+220.06 @+291.1 | **Charge:** | +2 | | |

**Sequence**

DQVMELT#IAGMGTSALK

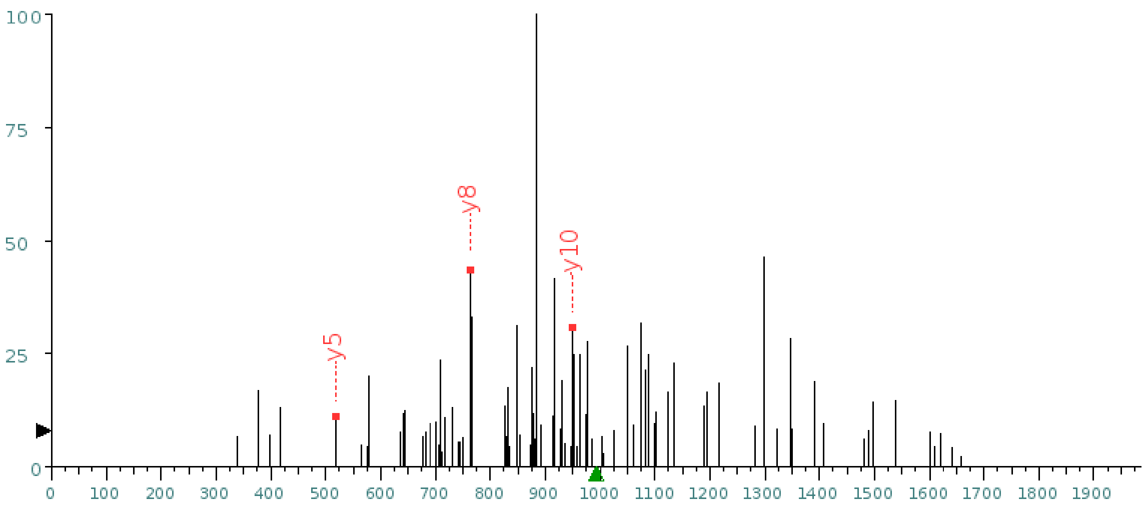

**Predicted Fragmentation Pattern**

| Seq | # | b: Δ Error | b | y | y: Δ Error | +1 |
|---|---|---|---|---|---|---|
| D | 1 | --- | 116.034 | --- | --- | 17 |
| Q | 2 | --- | 244.093 | 1869.918 | --- | 16 |
| V | 3 | --- | 343.161 | 1741.860 | --- | 15 |
| M | 4 | --- | 474.202 | 1642.791 | --- | 14 |
| E | 5 | --- | 603.244 | 1511.751 | --- | 13 |
| L | 6 | --- | 716.328 | 1382.708 | --- | 12 |
| T# | 7 | --- | 1037.434 | 1269.624 | --- | 11 |
| I | 8 | --- | 1150.518 | **948.518** | -80.794 | 10 |
| A | 9 | --- | 1221.556 | 835.434 | --- | 9 |
| G | 10 | --- | 1278.577 | **764.397** | 445.023 | 8 |
| M | 11 | --- | 1409.617 | 707.376 | --- | 7 |
| G | 12 | --- | 1466.639 | 576.335 | --- | 6 |
| T | 13 | --- | 1567.687 | **519.314** | -305.847 | 5 |
| S | 14 | --- | 1654.719 | 418.266 | --- | 4 |
| A | 15 | --- | 1725.756 | 331.234 | --- | 3 |
| L | 16 | --- | 1838.840 | 260.197 | --- | 2 |
| K | 17 | --- | --- | 147.113 | --- | 1 |

| | | | | | |
|---|---|---|---|---|---|
| **Run:** | 77660 | **Scans:** | 22966 - 22966 | **m/z Range:** | 267.4899 to 1823.0941 |
| **Search:** | 77660 | **Precursor Mass:** | 1028.49422 | | |
| **Mods:** | C=160.03 *+15.99 #+220.06 @+291.1 | **Charge:** | +2 | | |

**Sequence**

DQVMELT[@]IAGMGTSALK

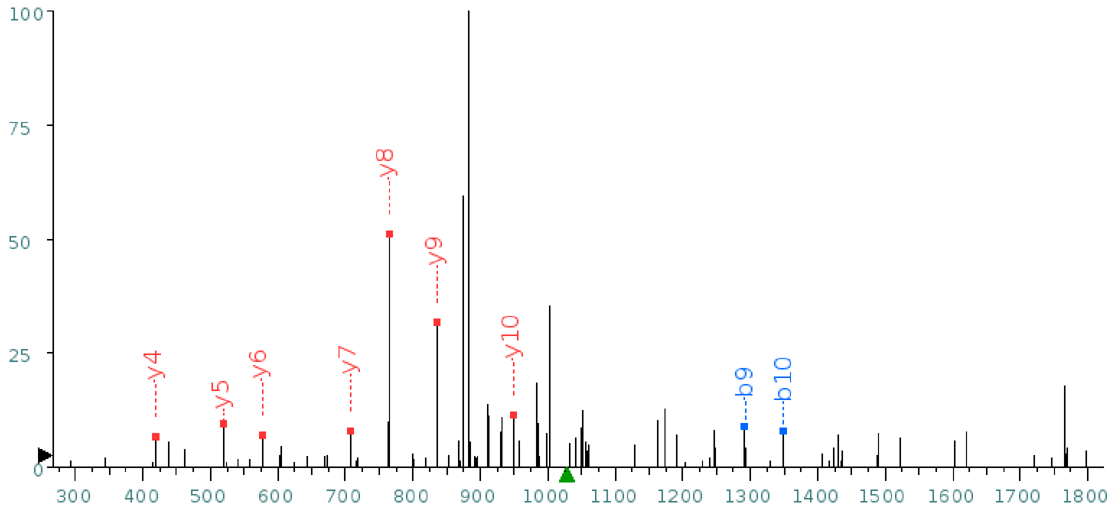

**Predicted Fragmentation Pattern**

| Seq | # | b: Δ Error | b | y | y: Δ Error | +1 |
|---|---|---|---|---|---|---|
| D | 1 | --- | 116.034 | --- | --- | 17 |
| Q | 2 | --- | 244.093 | 1940.956 | --- | 16 |
| V | 3 | --- | 343.161 | 1812.897 | --- | 15 |
| M | 4 | --- | 474.202 | 1713.829 | --- | 14 |
| E | 5 | --- | 603.244 | 1582.788 | --- | 13 |
| L | 6 | --- | 716.328 | 1453.745 | --- | 12 |
| T@ | 7 | --- | 1108.471 | 1340.661 | --- | 11 |
| I | 8 | --- | 1221.556 | **948.518** | 230.474 | 10 |
| A | 9 | -665.270 | **1292.593** | **835.434** | 338.861 | 9 |
| G | 10 | -497.862 | **1349.614** | **764.397** | 121.984 | 8 |
| M | 11 | --- | 1480.655 | **707.376** | 455.851 | 7 |
| G | 12 | --- | 1537.676 | **576.335** | 325.634 | 6 |
| T | 13 | --- | 1638.724 | **519.314** | 594.411 | 5 |
| S | 14 | --- | 1725.756 | **418.266** | 489.363 | 4 |
| A | 15 | --- | 1796.793 | 331.234 | --- | 3 |
| L | 16 | --- | 1909.877 | 260.197 | --- | 2 |
| K | 17 | --- | --- | 147.113 | --- | 1 |

| Run: | 77660 | Scans: | 23200 - 23200 | m/z Range: | 283.34958 to 1931.2939 |
|---|---|---|---|---|---|
| Search: | 77660 | Precursor Mass: | 1026.16762 | | |
| Mods: | C=160.03 *+15.99 #+220.06 @+291.1 | Charge: | +3 | | |

**Sequence**

SMVFQIGANKDQVMELT#IAGM*GTSALK

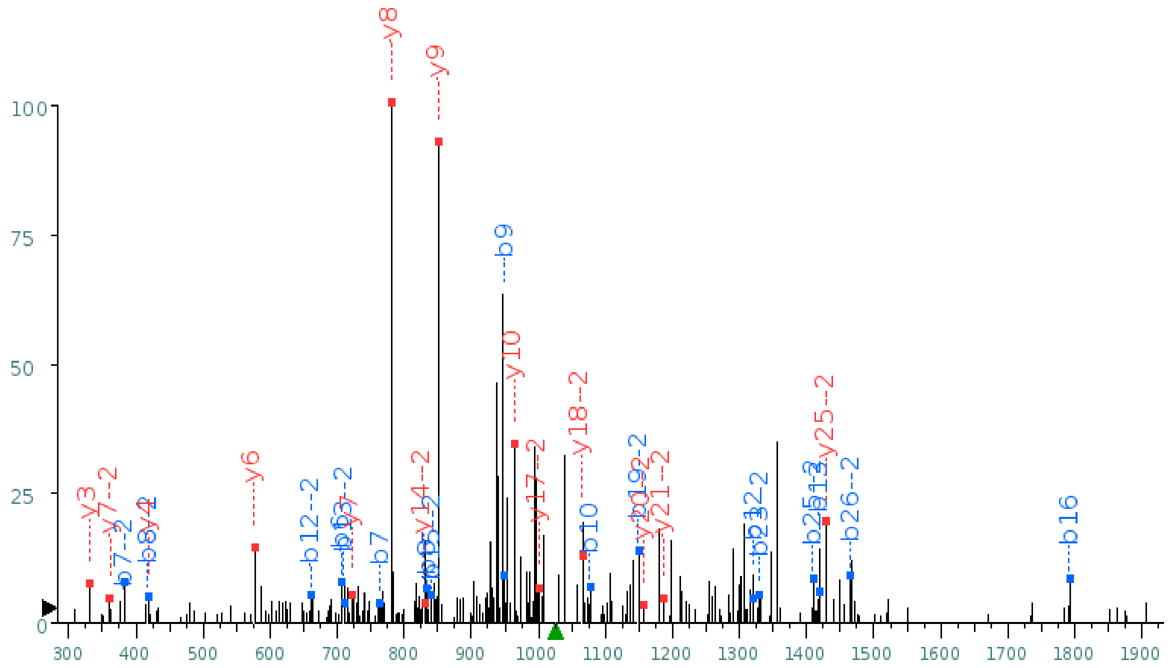

**Predicted Fragmentation Pattern**

| Seq | # | b: Δ Error | b | y | y: Δ Error | +1 |
|---|---|---|---|---|---|---|
| S | 1 | --- | 88.039 | --- | --- | 27 |
| M | 2 | --- | 219.080 | 2989.457 | --- | 26 |
| V | 3 | --- | 318.148 | 2858.416 | --- | 25 |
| F | 4 | --- | 465.217 | 2759.348 | --- | 24 |
| Q | 5 | --- | 593.275 | 2612.279 | --- | 23 |
| I | 6 | 344.469 | **706.359** | 2484.221 | --- | 22 |
| G | 7 | -991.910 | **763.381** | 2371.137 | --- | 21 |
| A | 8 | -860.790 | **834.418** | 2314.115 | --- | 20 |
| N | 9 | -645.037 | **948.461** | 2243.078 | --- | 19 |
| K | 10 | 547.315 | **1076.556** | 2129.035 | --- | 18 |
| D | 11 | --- | 1191.583 | 2000.940 | --- | 17 |
| Q | 12 | 291.752 | **1319.641** | 1885.913 | --- | 16 |
| V | 13 | 157.130 | **1418.710** | 1757.855 | --- | 15 |

| | | | | | |
|---|---|---|---|---|---|
| **Run:** | 77660 | **Scans:** | 23207 - 23207 | **m/z Range:** | 306.34875 to 1840.8714( |
| **Search:** | 77660 | **Precursor Mass:** | 1028.49464 | | |
| **Mods:** | C=160.03 *+15.99 #+220.06 @+291.1 | **Charge:** | +2 | | |

### Sequence

DQVMELT@IAGMGTSALK

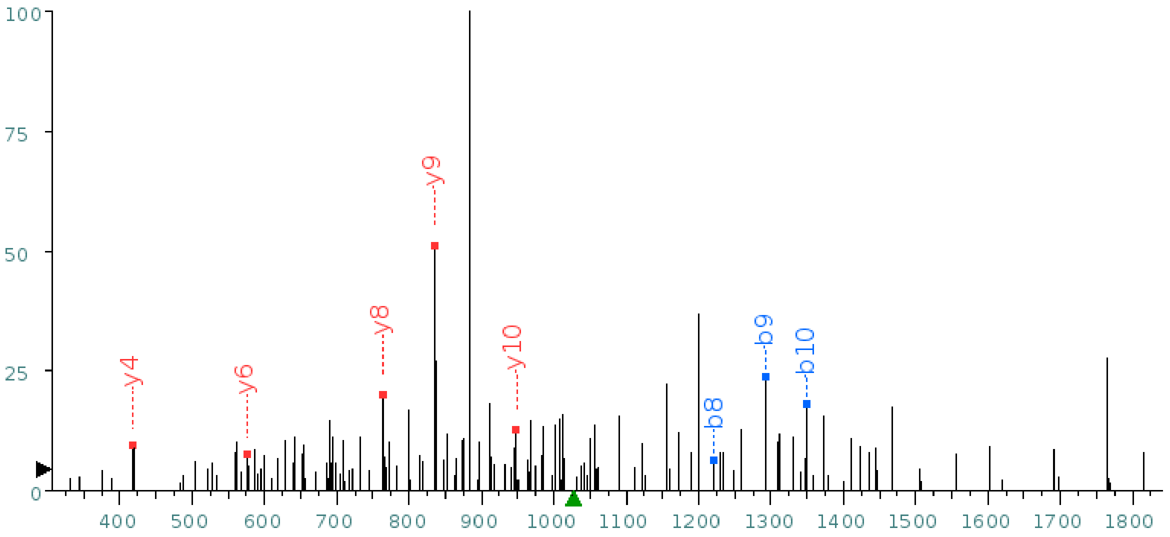

### Predicted Fragmentation Pattern

| Seq | # | b: Δ Error | b | y | y: Δ Error | +1 |
|---|---|---|---|---|---|---|
| D | 1 | --- | 116.034 | --- | --- | 17 |
| Q | 2 | --- | 244.093 | 1940.956 | --- | 16 |
| V | 3 | --- | 343.161 | 1812.897 | --- | 15 |
| M | 4 | --- | 474.202 | 1713.829 | --- | 14 |
| E | 5 | --- | 603.244 | 1582.788 | --- | 13 |
| L | 6 | --- | 716.328 | 1453.745 | --- | 12 |
| T@ | 7 | --- | 1108.471 | 1340.661 | --- | 11 |
| I | 8 | -457.868 | **1221.556** | **948.518** | -481.005 | 10 |
| A | 9 | -512.289 | **1292.593** | **835.434** | 58.138 | 9 |
| G | 10 | 102.048 | **1349.614** | **764.397** | 15.801 | 8 |
| M | 11 | --- | 1480.655 | 707.376 | --- | 7 |
| G | 12 | --- | 1537.676 | **576.335** | -26.809 | 6 |
| T | 13 | --- | 1638.724 | 519.314 | --- | 5 |
| S | 14 | --- | 1725.756 | **418.266** | 98.661 | 4 |
| A | 15 | --- | 1796.793 | 331.234 | --- | 3 |
| L | 16 | --- | 1909.877 | 260.197 | --- | 2 |
| K | 17 | --- | --- | 147.113 | --- | 1 |

| Run: | 77660 | Scans: | 23258 - 23258 | m/z Range: | 307.39368 to 1854.2603 |
|---|---|---|---|---|---|
| Search: | 77660 | Precursor Mass: | 1152.20835 | | |
| Mods: | C=160.03 *+15.99 #+220.06 @+291.1 | Charge: | +3 | | |

### Sequence

S@MVFQIGANKDQVM*ELT@IAGM*GTSALK

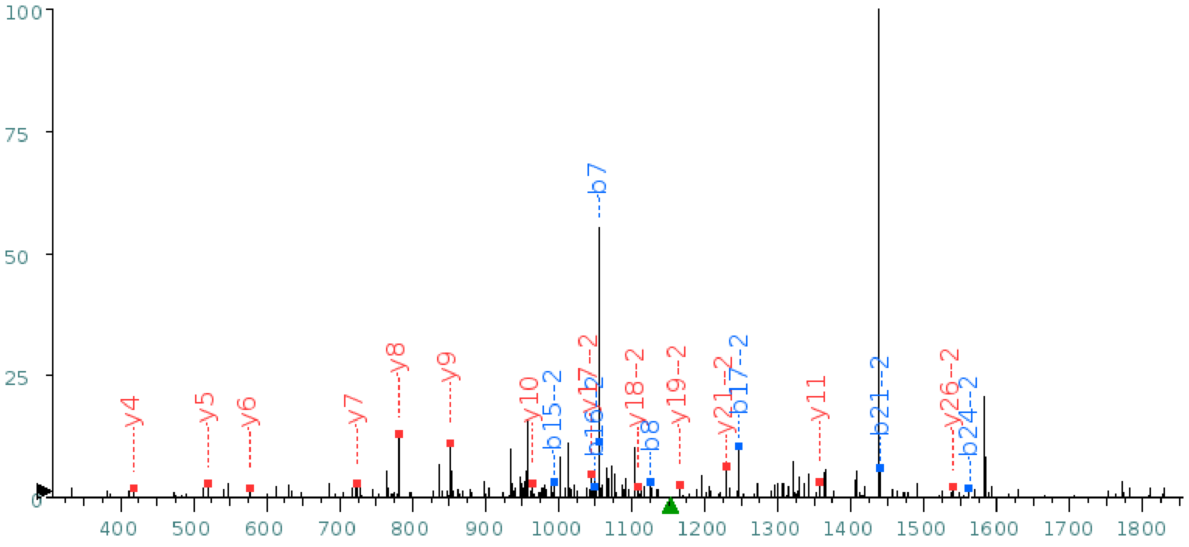

### Predicted Fragmentation Pattern

| | | | +1 | | | |
|---|---|---|---|---|---|---|
| Seq | # | b: Δ Error | b | y | y: Δ Error | +1 |
| S@ | 1 | --- | 379.135 | --- | --- | 27 |
| M | 2 | --- | 510.175 | 3076.489 | --- | 26 |
| V | 3 | --- | 609.244 | 2945.448 | --- | 25 |
| F | 4 | --- | 756.312 | 2846.380 | --- | 24 |
| Q | 5 | --- | 884.371 | 2699.311 | --- | 23 |
| I | 6 | --- | 997.455 | 2571.253 | --- | 22 |
| G | 7 | 563.042 | **1054.476** | 2458.169 | --- | 21 |
| A | 8 | -456.119 | **1125.513** | 2401.147 | --- | 20 |
| N | 9 | --- | 1239.556 | 2330.110 | --- | 19 |
| K | 10 | --- | 1367.651 | 2216.067 | --- | 18 |
| D | 11 | --- | 1482.678 | 2087.972 | --- | 17 |
| Q | 12 | --- | 1610.737 | 1972.945 | --- | 16 |
| V | 13 | --- | 1709.805 | 1844.887 | --- | 15 |
| M* | 14 | --- | 1856.840 | 1745.818 | --- | 14 |
| E | 15 | --- | 1985.883 | 1598.783 | --- | 13 |
| L | 16 | --- | 2098.967 | 1469.740 | --- | 12 |
| T@ | 17 | --- | 2491.110 | **1356.656** | -314.336 | 11 |
| I | 18 | --- | 2604.194 | **964.513** | 346.901 | 10 |
| A | 19 | --- | 2675.231 | **851.429** | 268.257 | 9 |

| Run: | 77660 | Scans: | 23347 - 23347 | m/z Range: | 267.24878 to 1952.2758 |
|---|---|---|---|---|---|
| Search: | 77660 | Precursor Mass: | 1028.49395 | | |
| Mods: | C=160.03 *+15.99 #+220.06 @+291.1 | Charge: | +2 | | |

**Sequence**

DQVMELT@IAGMGTSALK

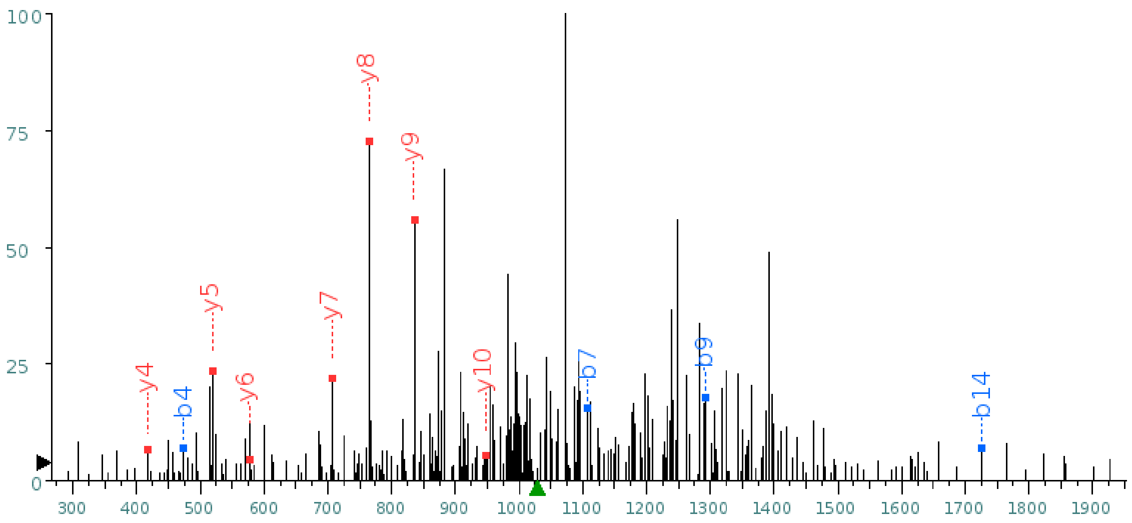

**Predicted Fragmentation Pattern**

| Seq | # | b: Δ Error | b | y | y: Δ Error | +1 |
|---|---|---|---|---|---|---|
| D | 1 | --- | 116.034 | --- | --- | 17 |
| Q | 2 | --- | 244.093 | 1940.956 | --- | 16 |
| V | 3 | --- | 343.161 | 1812.897 | --- | 15 |
| M | 4 | 364.240 | **474.202** | 1713.829 | --- | 14 |
| E | 5 | --- | 603.244 | 1582.788 | --- | 13 |
| L | 6 | --- | 716.328 | 1453.745 | --- | 12 |
| T@ | 7 | 20.139 | **1108.471** | 1340.661 | --- | 11 |
| I | 8 | --- | 1221.556 | **948.518** | 281.540 | 10 |
| A | 9 | 609.340 | **1292.593** | **835.434** | 367.772 | 9 |
| G | 10 | --- | 1349.614 | **764.397** | 390.293 | 8 |
| M | 11 | --- | 1480.655 | **707.376** | 101.594 | 7 |
| G | 12 | --- | 1537.676 | **576.335** | 151.719 | 6 |
| T | 13 | --- | 1638.724 | **519.314** | 492.153 | 5 |
| S | 14 | 210.597 | **1725.756** | **418.266** | 400.355 | 4 |
| A | 15 | --- | 1796.793 | 331.234 | --- | 3 |
| L | 16 | --- | 1909.877 | 260.197 | --- | 2 |
| K | 17 | --- | --- | 147.113 | --- | 1 |

**Run:** 77660

**Scans:** 23373 - 23373

**m/z Range:** 269.20261 to 1989.0351

**Search:** 77660

**Precursor Mass:** 1026.16767

**Mods:** C=160.03 *+15.99 #+220.06 @+291.1

**Charge:** +3

### Sequence

S#MVFQIGANKDQVM*ELTIAGMGTSALK

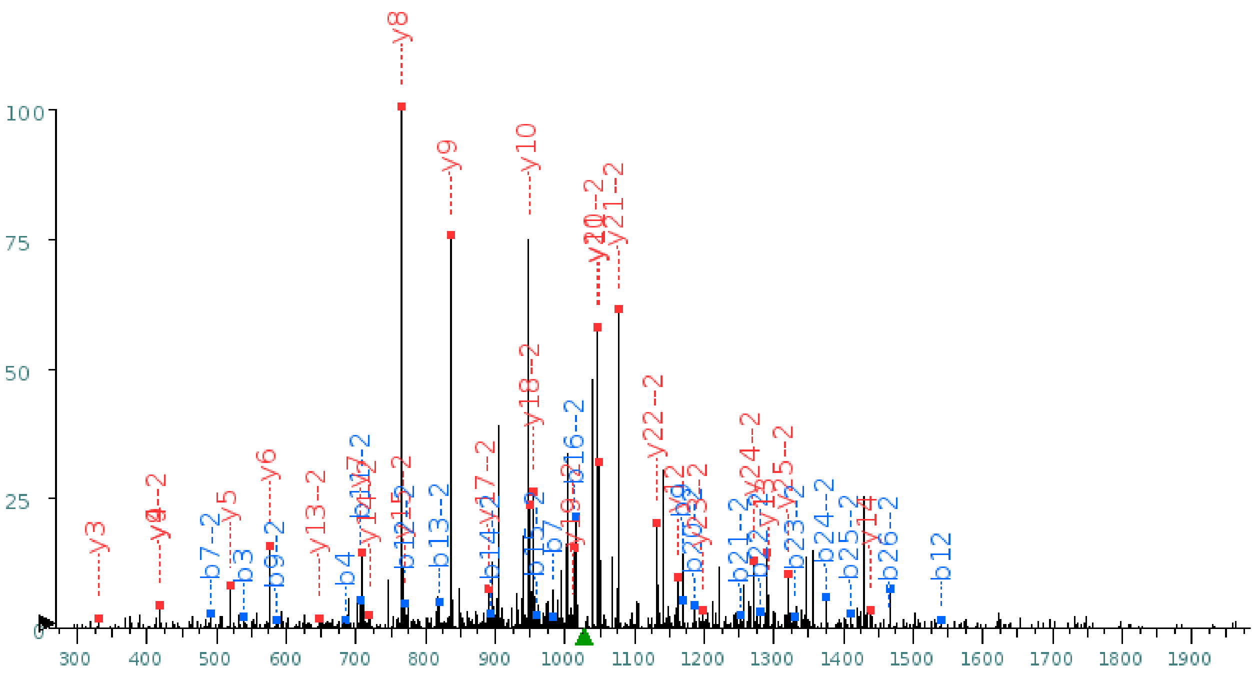

### Predicted Fragmentation Pattern

+1

| Seq | # | b: Δ Error | b | y | y: Δ Error | +1 |
|---|---|---|---|---|---|---|
| S# | 1 | --- | 308.098 | --- | --- | 27 |
| M | 2 | --- | 439.138 | 2769.398 | --- | 26 |
| V | 3 | 159.043 | **538.207** | 2638.358 | --- | 25 |
| F | 4 | 179.327 | **685.275** | 2539.289 | --- | 24 |
| Q | 5 | --- | 813.333 | 2392.221 | --- | 23 |
| I | 6 | --- | 926.418 | 2264.162 | --- | 22 |
| G | 7 | -341.645 | **983.439** | 2151.078 | --- | 21 |
| A | 8 | --- | 1054.476 | 2094.057 | --- | 20 |
| N | 9 | 820.098 | **1168.519** | 2023.020 | --- | 19 |
| K | 10 | --- | 1296.614 | 1908.977 | --- | 18 |
| D | 11 | --- | 1411.641 | 1780.882 | --- | 17 |
| Q | 12 | 126.860 | **1539.700** | 1665.855 | --- | 16 |
| V | 13 | --- | 1638.768 | 1537.796 | --- | 15 |
| M* | 14 | --- | 1785.803 | **1438.728** | 199.019 | 14 |

| Run: | 77660 | Scans: | 23490 - 23490 | m/z Range: | 281.41684 to 2000 |
|---|---|---|---|---|---|
| Search: | 77660 | Precursor Mass: | 1028.49572 | | |
| Mods: | C=160.03 *+15.99 #+220.06 @+291.1 | Charge: | +2 | | |

### Sequence

DQVMELT@IAGMGTSALK

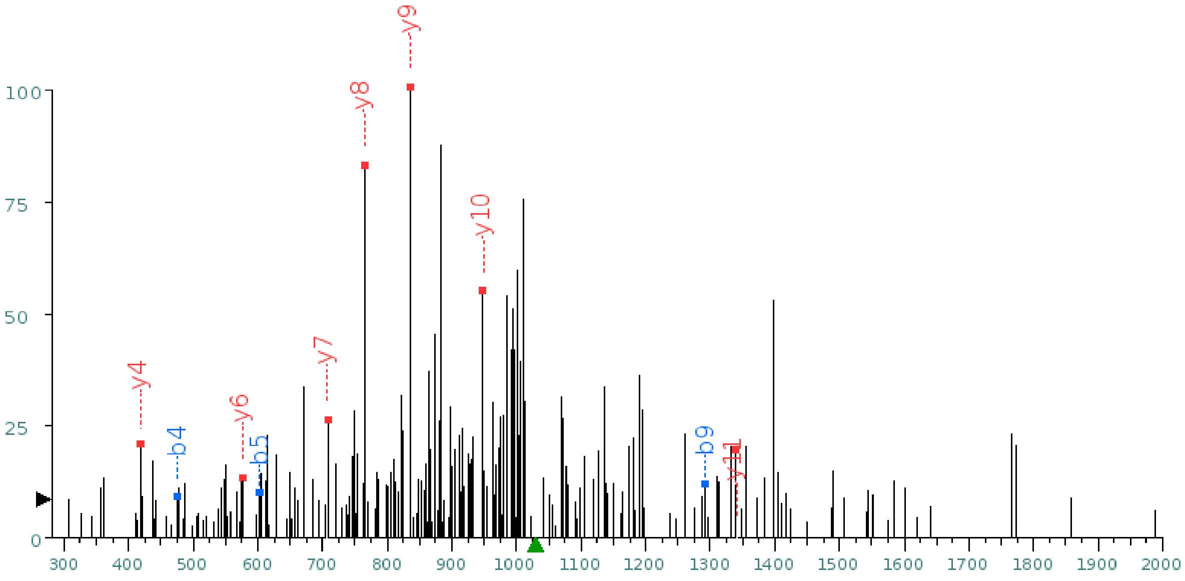

### Predicted Fragmentation Pattern

| Seq | # | b: Δ Error | b | y | y: Δ Error | +1 |
|---|---|---|---|---|---|---|
| D | 1 | --- | 116.034 | --- | --- | 17 |
| Q | 2 | --- | 244.093 | 1940.956 | --- | 16 |
| V | 3 | --- | 343.161 | 1812.897 | --- | 15 |
| M | 4 | 673.405 | **474.202** | 1713.829 | --- | 14 |
| E | 5 | 270.031 | **603.244** | 1582.788 | --- | 13 |
| L | 6 | --- | 716.328 | 1453.745 | --- | 12 |
| T@ | 7 | --- | 1108.471 | **1340.661** | -728.585 | 11 |
| I | 8 | --- | 1221.556 | **948.518** | -252.789 | 10 |
| A | 9 | 134.952 | **1292.593** | **835.434** | -131.681 | 9 |
| G | 10 | --- | 1349.614 | **764.397** | -7.993 | 8 |
| M | 11 | --- | 1480.655 | **707.376** | 336.011 | 7 |
| G | 12 | --- | 1537.676 | **576.335** | 338.016 | 6 |
| T | 13 | --- | 1638.724 | 519.314 | --- | 5 |
| S | 14 | --- | 1725.756 | **418.266** | 418.799 | 4 |
| A | 15 | --- | 1796.793 | 331.234 | --- | 3 |
| L | 16 | --- | 1909.877 | 260.197 | --- | 2 |
| K | 17 | --- | --- | 147.113 | --- | 1 |

| | | | | | |
|---|---|---|---|---|---|
| **Run:** | 77660 | **Scans:** | 23747 - 23747 | **m/z Range:** | 293.2767 to 2000 |
| **Search:** | 77660 | **Precursor Mass:** | 1049.84592 | | |
| **Mods:** | C=160.03 *+15.99 #+220.06 @+291.1 | **Charge:** | +3 | | |

**Sequence**

SMVFQIGANKDQVMELTIAGM*GT@SALK

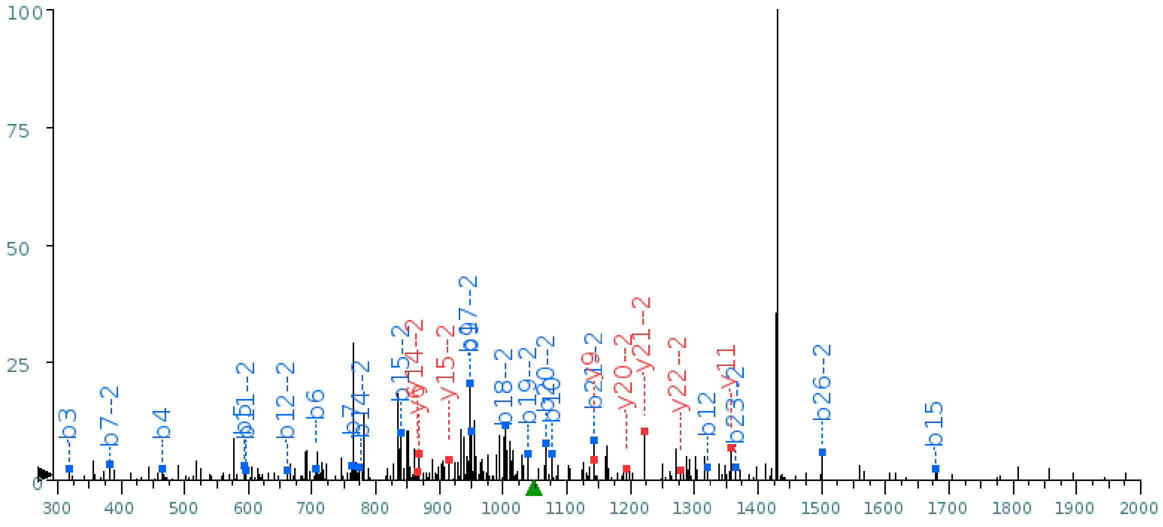

**Predicted Fragmentation Pattern**

| Seq | # | b: Δ Error | b | y | y: Δ Error | +1 |
|---|---|---|---|---|---|---|
| | | | +1 | | | |
| S | 1 | --- | 88.039 | --- | --- | 27 |
| M | 2 | --- | 219.080 | 3060.494 | --- | 26 |
| V | 3 | 403.737 | **318.148** | 2929.453 | --- | 25 |
| F | 4 | 151.438 | **465.217** | 2830.385 | --- | 24 |
| Q | 5 | 57.528 | **593.275** | 2683.316 | --- | 23 |
| I | 6 | 122.504 | **706.359** | 2555.258 | --- | 22 |
| G | 7 | -1167.227 | **763.381** | 2442.174 | --- | 21 |
| A | 8 | --- | 834.418 | 2385.152 | --- | 20 |
| N | 9 | 374.185 | **948.461** | 2314.115 | --- | 19 |
| K | 10 | -910.385 | **1076.556** | 2200.072 | --- | 18 |
| D | 11 | --- | 1191.583 | 2071.977 | --- | 17 |
| Q | 12 | 240.163 | **1319.641** | 1956.950 | --- | 16 |
| V | 13 | --- | 1418.710 | 1828.892 | --- | 15 |
| M | 14 | --- | 1549.750 | 1729.823 | --- | 14 |
| E | 15 | 294.684 | **1678.793** | 1598.783 | --- | 13 |
| L | 16 | --- | 1791.877 | 1469.740 | --- | 12 |
| T | 17 | --- | 1892.924 | **1356.656** | 111.710 | 11 |
| I | 18 | --- | 2006.009 | 1255.609 | --- | 10 |
| A | 19 | --- | 2077.046 | **1142.525** | 221.958 | 9 |
| G | 20 | --- | 2134.067 | 1071.487 | --- | 8 |

| | | | | | |
|---|---|---|---|---|---|
| **Run:** | 77660 | **Scans:** | 23747 - 23747 | **m/z Range:** | 293.2767 to 2000 |
| **Search:** | 77660 | **Precursor Mass:** | 1049.84592 | | |
| **Mods:** | C=160.03 *+15.99 #+220.06 @+291.1 | **Charge:** | +3 | | |

**Sequence**

SMVFQIGANKDQVMELTIAGM*GT@SALK

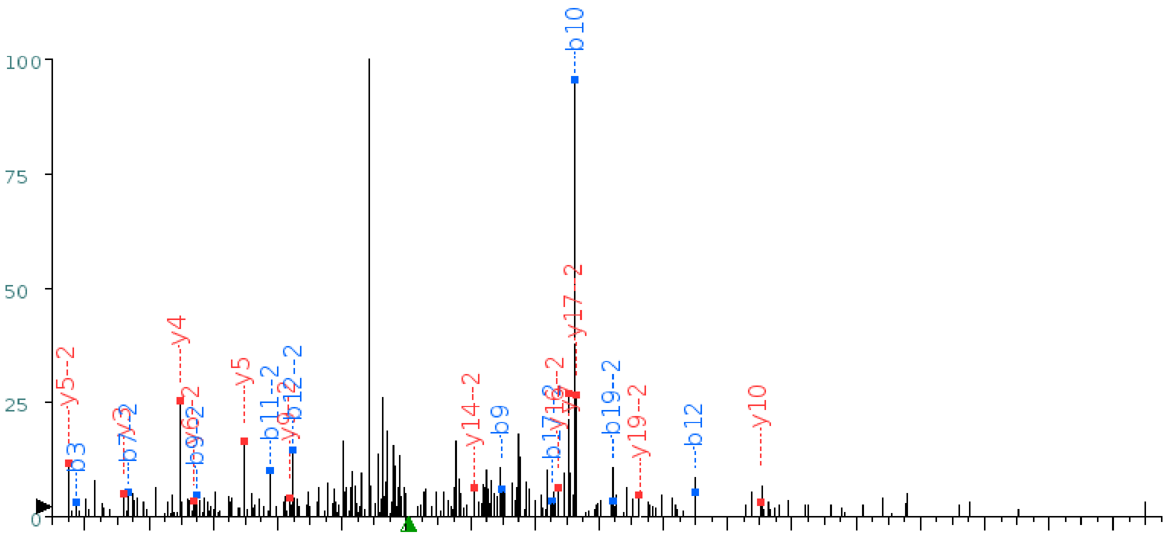

**Predicted Fragmentation Pattern**

| Seq | # | b: Δ Error | b (+1) | y (+1) | y: Δ Error | +1 |
|---|---|---|---|---|---|---|
| S | 1 | --- | 88.039 | --- | --- | 27 |
| M | 2 | --- | 219.080 | 3060.494 | --- | 26 |
| V | 3 | 403.737 | **318.148** | 2929.453 | --- | 25 |
| F | 4 | 151.438 | **465.217** | 2830.385 | --- | 24 |
| Q | 5 | 57.528 | **593.275** | 2683.316 | --- | 23 |
| I | 6 | 122.504 | **706.359** | 2555.258 | --- | 22 |
| G | 7 | -1167.227 | **763.381** | 2442.174 | --- | 21 |
| A | 8 | --- | 834.418 | 2385.152 | --- | 20 |
| N | 9 | 374.185 | **948.461** | 2314.115 | --- | 19 |
| K | 10 | -910.385 | **1076.556** | 2200.072 | --- | 18 |
| D | 11 | --- | 1191.583 | 2071.977 | --- | 17 |
| Q | 12 | 240.163 | **1319.641** | 1956.950 | --- | 16 |
| V | 13 | --- | 1418.710 | 1828.892 | --- | 15 |
| M | 14 | --- | 1549.750 | 1729.823 | --- | 14 |
| E | 15 | 294.684 | **1678.793** | 1598.783 | --- | 13 |
| L | 16 | --- | 1791.877 | 1469.740 | --- | 12 |
| T | 17 | --- | 1892.924 | **1356.656** | 111.710 | 11 |
| I | 18 | --- | 2006.009 | 1255.609 | --- | 10 |
| A | 19 | --- | 2077.046 | **1142.525** | 221.958 | 9 |
| G | 20 | --- | 2134.067 | 1071.487 | --- | 8 |
| M* | 21 | --- | 2281.103 | 1014.466 | --- | 7 |

| | | | | | |
|---|---|---|---|---|---|
| **Run:** | 77660 | **Scans:** | 24786 - 24786 | **m/z Range:** | 202.07262 to 1447.9600[illegible] |
| **Search:** | 77660 | **Precursor Mass:** | 783.64063 | | |
| **Mods:** | C=160.03 *+15.99 #+220.06 @+291.1 | **Charge:** | +4 | | |

### Sequence

SMVFQIGANKDQVMELTIAGMGTS@ALK

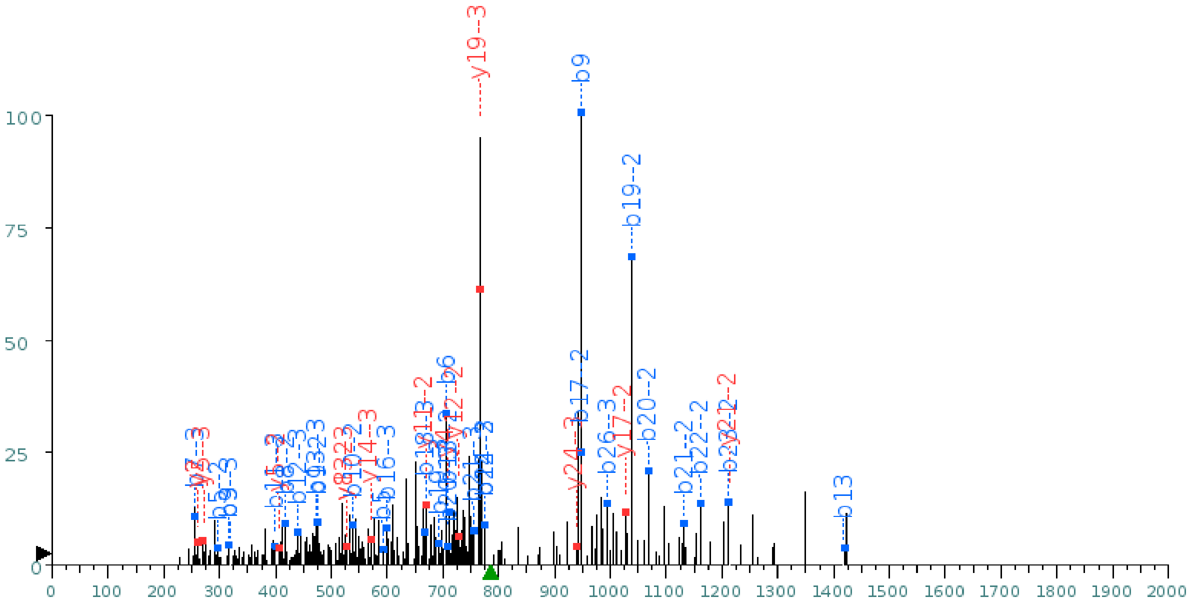

### Predicted Fragmentation Pattern

+1

| Seq | # | b: Δ Error | b | y | y: Δ Error | +1 |
|---|---|---|---|---|---|---|
| S | 1 | --- | 88.039 | --- | --- | 27 |
| M | 2 | --- | 219.080 | 3044.499 | --- | 26 |
| V | 3 | -2727.042 | **318.148** | 2913.458 | --- | 25 |
| F | 4 | --- | 465.217 | 2814.390 | --- | 24 |
| Q | 5 | 234.324 | **593.275** | 2667.322 | --- | 23 |
| I | 6 | 582.734 | **706.359** | 2539.263 | --- | 22 |
| G | 7 | --- | 763.381 | 2426.179 | --- | 21 |
| A | 8 | --- | 834.418 | 2369.157 | --- | 20 |
| N | 9 | -333.205 | **948.461** | 2298.120 | --- | 19 |
| K | 10 | --- | 1076.556 | 2184.077 | --- | 18 |
| D | 11 | --- | 1191.583 | 2055.982 | --- | 17 |
| Q | 12 | --- | 1319.641 | 1940.956 | --- | 16 |
| V | 13 | 246.148 | **1418.710** | 1812.897 | --- | 15 |
| M | 14 | --- | 1549.750 | 1713.829 | --- | 14 |
| E | 15 | --- | 1678.793 | 1582.788 | --- | 13 |
| L | 16 | --- | 1791.877 | 1453.745 | --- | 12 |
| T | 17 | --- | 1892.924 | 1340.661 | --- | 11 |
| I | 18 | --- | 2006.009 | 1239.614 | --- | 10 |
| A | 19 | --- | 2077.046 | 1126.530 | --- | 9 |
| G | 20 | --- | 2134.067 | 1055.493 | --- | 8 |
| M | 21 | --- | 2265.108 | 998.471 | --- | 7 |

| | | | | | |
|---|---|---|---|---|---|
| **Run:** | 77660 | **Scans:** | 25024 - 25024 | **m/z Range:** | 303.31967 to 1931.1270[illegible] |
| **Search:** | 77660 | **Precursor Mass:** | 1146.87738 | | |
| **Mods:** | C=160.03 *+15.99 #+220.06 @+291.1 | **Charge:** | +3 | | |

### Sequence

S@MVFQIGANKDQVM*ELT@IAGMGTSALK

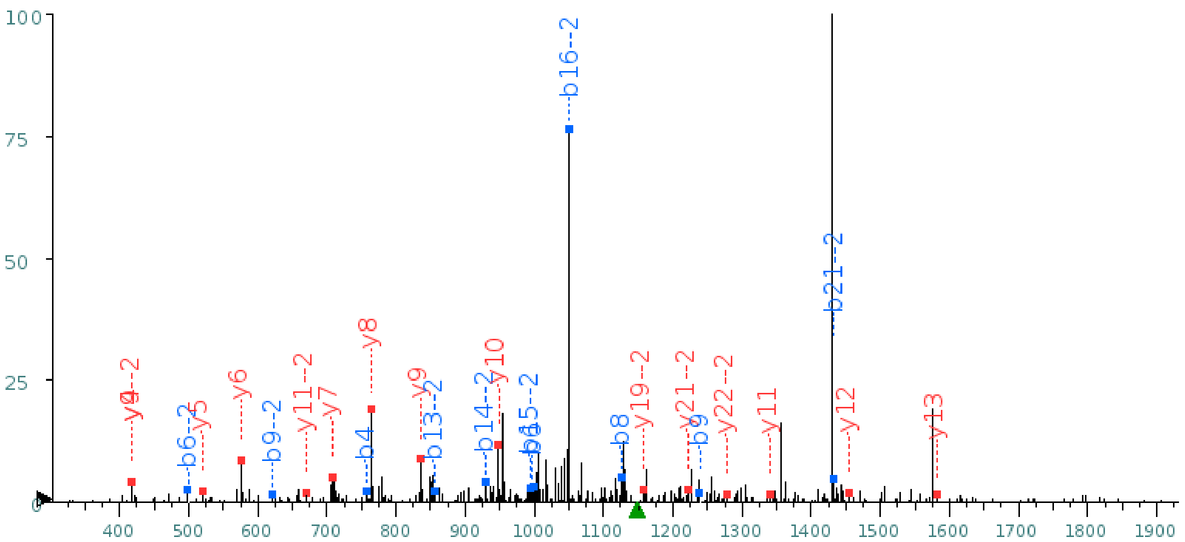

### Predicted Fragmentation Pattern

+1

| Seq | # | b: Δ Error | b | y | y: Δ Error | +1 |
|---|---|---|---|---|---|---|
| S@ | 1 | --- | 379.135 | --- | --- | 27 |
| M | 2 | --- | 510.175 | 3060.494 | --- | 26 |
| V | 3 | --- | 609.244 | 2929.453 | --- | 25 |
| F | 4 | 425.887 | **756.312** | 2830.385 | --- | 24 |
| Q | 5 | --- | 884.371 | 2683.316 | --- | 23 |
| I | 6 | 278.864 | **997.455** | 2555.258 | --- | 22 |
| G | 7 | --- | 1054.476 | 2442.174 | --- | 21 |
| A | 8 | 80.839 | **1125.513** | 2385.152 | --- | 20 |
| N | 9 | -427.213 | **1239.556** | 2314.115 | --- | 19 |
| K | 10 | --- | 1367.651 | 2200.072 | --- | 18 |
| D | 11 | --- | 1482.678 | 2071.977 | --- | 17 |
| Q | 12 | --- | 1610.737 | 1956.950 | --- | 16 |
| V | 13 | --- | 1709.805 | 1828.892 | --- | 15 |
| M* | 14 | --- | 1856.840 | 1729.823 | --- | 14 |
| E | 15 | --- | 1985.883 | **1582.788** | -578.189 | 13 |
| L | 16 | --- | 2098.967 | **1453.745** | 88.861 | 12 |
| T@ | 17 | --- | 2491.110 | **1340.661** | 178.248 | 11 |
| I | 18 | --- | 2604.194 | **948.518** | -561.524 | 10 |
| A | 19 | --- | 2675.231 | **835.434** | 199.542 | 9 |

| | | | | | |
|---|---|---|---|---|---|
| **Run:** | 77660 | **Scans:** | 25108 - 25108 | **m/z Range:** | 276.40558 to 1957.2564 |
| **Search:** | 77660 | **Precursor Mass:** | 1049.84541 | | |
| **Mods:** | C=160.03 *+15.99 #+220.06 @+291.1 | **Charge:** | +3 | | |

**Sequence**

SMVFQIGANKDQVMELTIAGM*GT@SALK

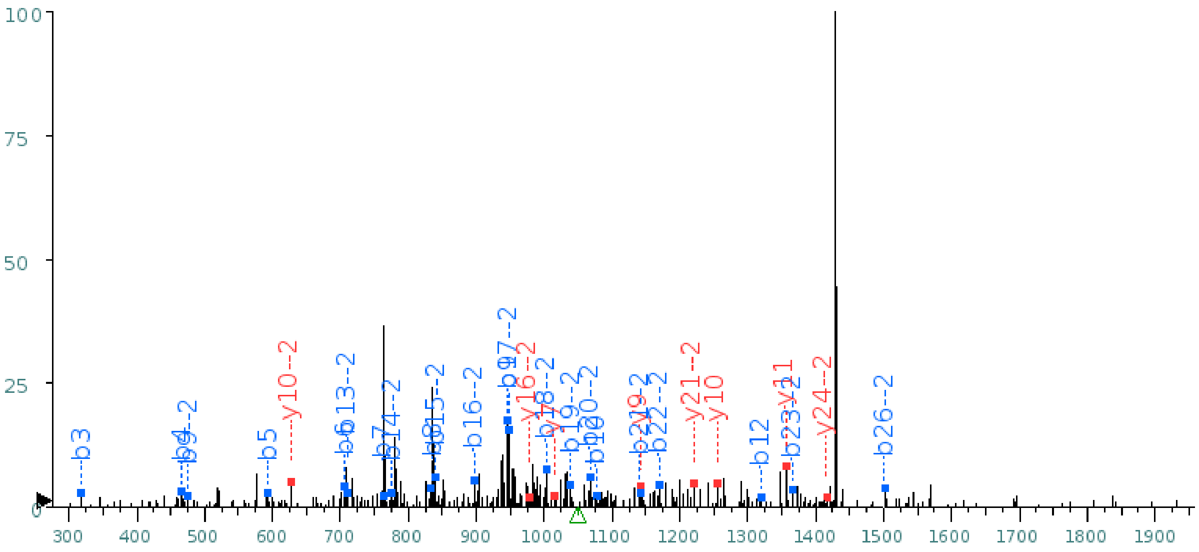

**Predicted Fragmentation Pattern**

+1

| Seq | # | b: Δ Error | b | y | y: Δ Error | +1 |
|---|---|---|---|---|---|---|
| S | 1 | --- | 88.039 | --- | --- | 27 |
| M | 2 | --- | 219.080 | 3060.494 | --- | 26 |
| V | 3 | -97.977 | **318.148** | 2929.453 | --- | 25 |
| F | 4 | 280.415 | **465.217** | 2830.385 | --- | 24 |
| Q | 5 | -1447.407 | **593.275** | 2683.316 | --- | 23 |
| I | 6 | 295.248 | **706.359** | 2555.258 | --- | 22 |
| G | 7 | 308.386 | **763.381** | 2442.174 | --- | 21 |
| A | 8 | -1018.938 | **834.418** | 2385.152 | --- | 20 |
| N | 9 | -812.725 | **948.461** | 2314.115 | --- | 19 |
| K | 10 | 234.377 | **1076.556** | 2200.072 | --- | 18 |
| D | 11 | --- | 1191.583 | 2071.977 | --- | 17 |
| Q | 12 | -54.218 | **1319.641** | 1956.950 | --- | 16 |
| V | 13 | --- | 1418.710 | 1828.892 | --- | 15 |
| M | 14 | --- | 1549.750 | 1729.823 | --- | 14 |
| E | 15 | --- | 1678.793 | 1598.783 | --- | 13 |
| L | 16 | --- | 1791.877 | 1469.740 | --- | 12 |
| T | 17 | --- | 1892.924 | **1356.656** | -363.227 | 11 |

| | | | | | |
|---|---|---|---|---|---|
| **Run:** | 77660 | **Scans:** | 25134 - 25134 | **m/z Range:** | 346.34113 **to** 1942.1866 |
| **Search:** | 77660 | **Precursor Mass:** | 1146.87708 | | |
| **Mods:** | C=160.03 *+15.99 #+220.06 @+291.1 | **Charge:** | +3 | | |

**Sequence**

SMVFQIGANKDQVM*ELTIAGMGT@S@ALK

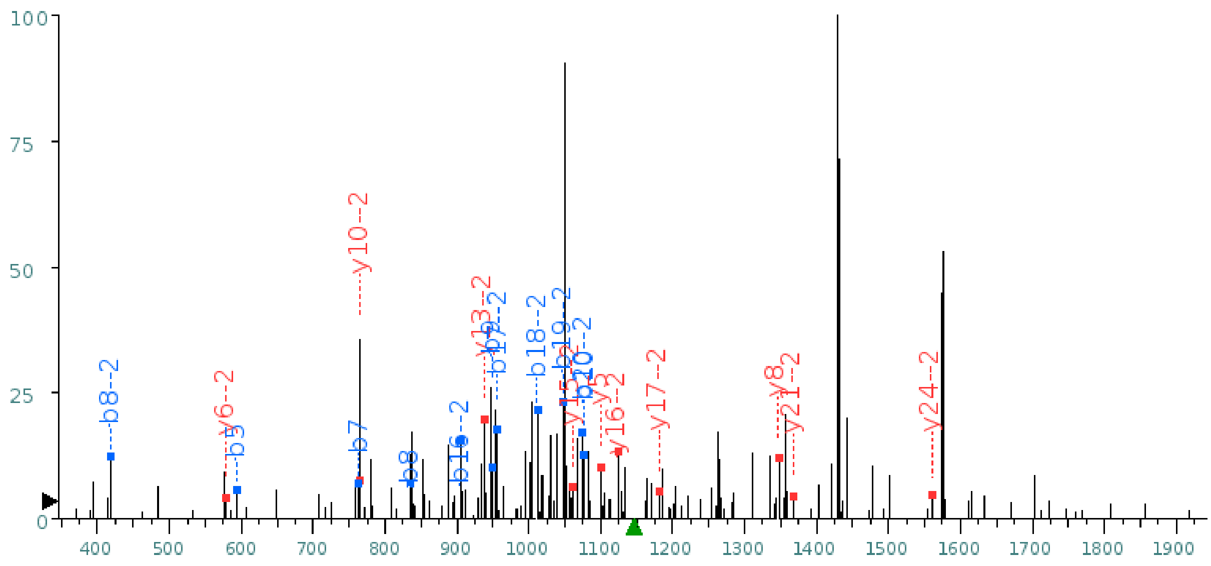

**Predicted Fragmentation Pattern**

| Seq | # | b: Δ Error | b +1 | y +1 | y: Δ Error | +1 |
|---|---|---|---|---|---|---|
| S | 1 | --- | 88.039 | --- | --- | 27 |
| M | 2 | --- | 219.080 | 3351.589 | --- | 26 |
| V | 3 | --- | 318.148 | 3220.549 | --- | 25 |
| F | 4 | --- | 465.217 | 3121.480 | --- | 24 |
| Q | 5 | **202.960** | **593.275** | 2974.412 | --- | 23 |
| I | 6 | --- | 706.359 | 2846.353 | --- | 22 |
| G | 7 | **115.300** | **763.381** | 2733.269 | --- | 21 |
| A | 8 | **342.809** | **834.418** | 2676.248 | --- | 20 |
| N | 9 | **361.324** | **948.461** | 2605.211 | --- | 19 |
| K | 10 | **227.464** | **1076.556** | 2491.168 | --- | 18 |
| D | 11 | --- | 1191.583 | 2363.073 | --- | 17 |
| Q | 12 | --- | 1319.641 | 2248.046 | --- | 16 |
| V | 13 | --- | 1418.710 | 2119.987 | --- | 15 |
| M* | 14 | --- | 1565.745 | 2020.919 | --- | 14 |
| E | 15 | --- | 1694.788 | 1873.883 | --- | 13 |
| L | 16 | --- | 1807.872 | 1744.841 | --- | 12 |
| T | 17 | --- | 1908.919 | 1631.757 | --- | 11 |
| I | 18 | --- | 2022.003 | 1530.709 | --- | 10 |

| | | | | | |
|---|---|---|---|---|---|
| **Run:** | 77660 | **Scans:** | 25308 - 25308 | **m/z Range:** | 301.4064 to 2000 |
| **Search:** | 77660 | **Precursor Mass:** | 1146.87765 | | |
| **Mods:** | C=160.03 *+15.99 #+220.06 @+291.1 | **Charge:** | +3 | | |

**Sequence**

S@MVFQIGANKDQVM*ELT@IAGMGTSALK

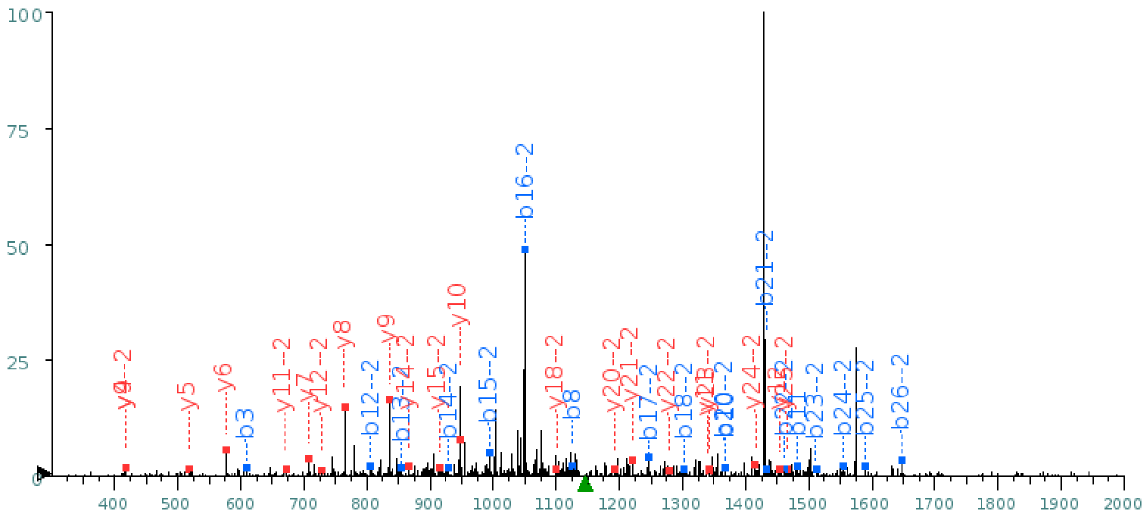

**Predicted Fragmentation Pattern**

| Seq | # | b: Δ Error | b | y | y: Δ Error | +1 |
|---|---|---|---|---|---|---|
| S@ | 1 | --- | 379.135 | --- | --- | 27 |
| M | 2 | --- | 510.175 | 3060.494 | --- | 26 |
| V | 3 | -1546.141 | **609.244** | 2929.453 | --- | 25 |
| F | 4 | --- | 756.312 | 2830.385 | --- | 24 |
| Q | 5 | --- | 884.371 | 2683.316 | --- | 23 |
| I | 6 | --- | 997.455 | 2555.258 | --- | 22 |
| G | 7 | --- | 1054.476 | 2442.174 | --- | 21 |
| A | 8 | -236.883 | **1125.513** | 2385.152 | --- | 20 |
| N | 9 | --- | 1239.556 | 2314.115 | --- | 19 |
| K | 10 | -674.350 | **1367.651** | 2200.072 | --- | 18 |
| D | 11 | 247.433 | **1482.678** | 2071.977 | --- | 17 |
| Q | 12 | --- | 1610.737 | 1956.950 | --- | 16 |
| V | 13 | --- | 1709.805 | 1828.892 | --- | 15 |
| M* | 14 | --- | 1856.840 | 1729.823 | --- | 14 |
| E | 15 | --- | 1985.883 | 1582.788 | --- | 13 |
| L | 16 | --- | 2098.967 | **1453.745** | -374.952 | 12 |
| T@ | 17 | --- | 2491.110 | **1340.661** | 707.067 | 11 |
| I | 18 | --- | 2604.194 | **948.518** | 85.609 | 10 |
| A | 19 | --- | 2675.231 | **835.434** | 22.927 | 9 |
| G | 20 | --- | 2732.253 | **764.397** | 281.454 | 8 |

| Run: | 77660 | Scans: | 25322 - 25322 | m/z Range: | 267.18863 to 2000 |
|---|---|---|---|---|---|
| Search: | 77660 | Precursor Mass: | 1044.51616 | | |
| Mods: | C=160.03 *+15.99 #+220.06 @+291.1 | Charge: | +3 | | |

**Sequence**

SMVFQIGANKDQVMELT@IAGMGTSALK

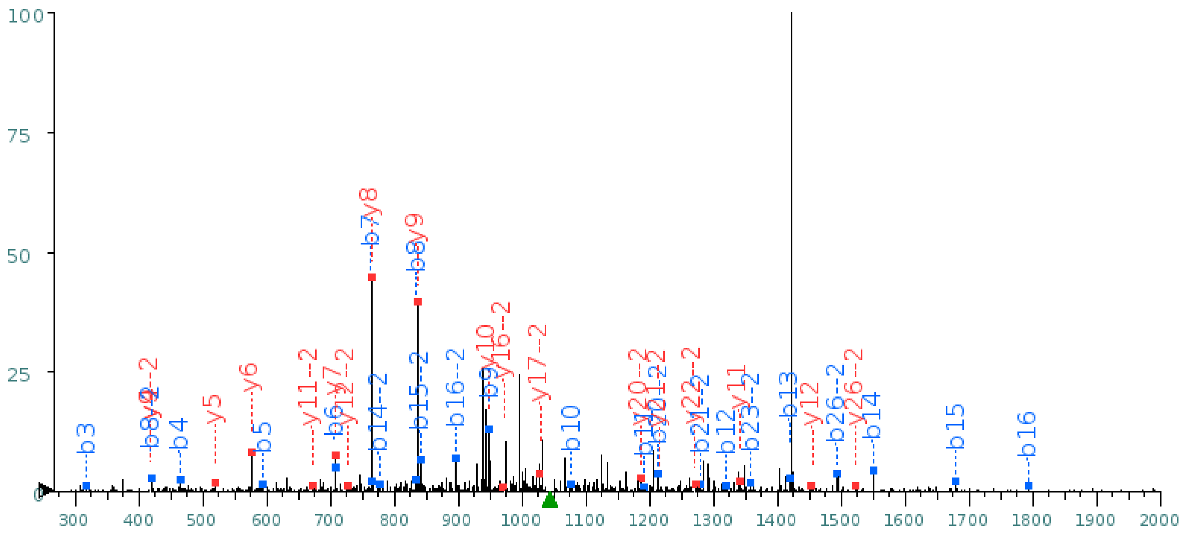

**Predicted Fragmentation Pattern**

| Seq | # | b: Δ Error | b | y | y: Δ Error | +1 |
|---|---|---|---|---|---|---|
| S | 1 | --- | 88.039 | --- | --- | 27 |
| M | 2 | --- | 219.080 | 3044.499 | --- | 26 |
| V | 3 | 423.097 | **318.148** | 2913.458 | --- | 25 |
| F | 4 | 59.160 | **465.217** | 2814.390 | --- | 24 |
| Q | 5 | 11.544 | **593.275** | 2667.322 | --- | 23 |
| I | 6 | 130.106 | **706.359** | 2539.263 | --- | 22 |
| G | 7 | 272.748 | **763.381** | 2426.179 | --- | 21 |
| A | 8 | 363.860 | **834.418** | 2369.157 | --- | 20 |
| N | 9 | 143.282 | **948.461** | 2298.120 | --- | 19 |
| K | 10 | -928.787 | **1076.556** | 2184.077 | --- | 18 |
| D | 11 | 237.385 | **1191.583** | 2055.982 | --- | 17 |
| Q | 12 | 187.921 | **1319.641** | 1940.956 | --- | 16 |
| V | 13 | 226.024 | **1418.710** | 1812.897 | --- | 15 |
| M | 14 | 232.296 | **1549.750** | 1713.829 | --- | 14 |
| E | 15 | 117.700 | **1678.793** | 1582.788 | --- | 13 |
| L | 16 | 208.024 | **1791.877** | **1453.745** | -456.722 | 12 |
| T@ | 17 | --- | 2184.020 | **1340.661** | -468.411 | 11 |
| I | 18 | --- | 2297.104 | **948.518** | 82.649 | 10 |

| | | | | | |
|---|---|---|---|---|---|
| **Run:** | 77660 | **Scans:** | 25351 - 25351 | **m/z Range:** | 267.22607 to 1986.5946 |
| **Search:** | 77660 | **Precursor Mass:** | 1049.8469 | | |
| **Mods:** | C=160.03 *+15.99 #+220.06 @+291.1 | **Charge:** | +3 | | |

**Sequence**

SMVFQIGANKDQVMELT@IAGM*GTSALK

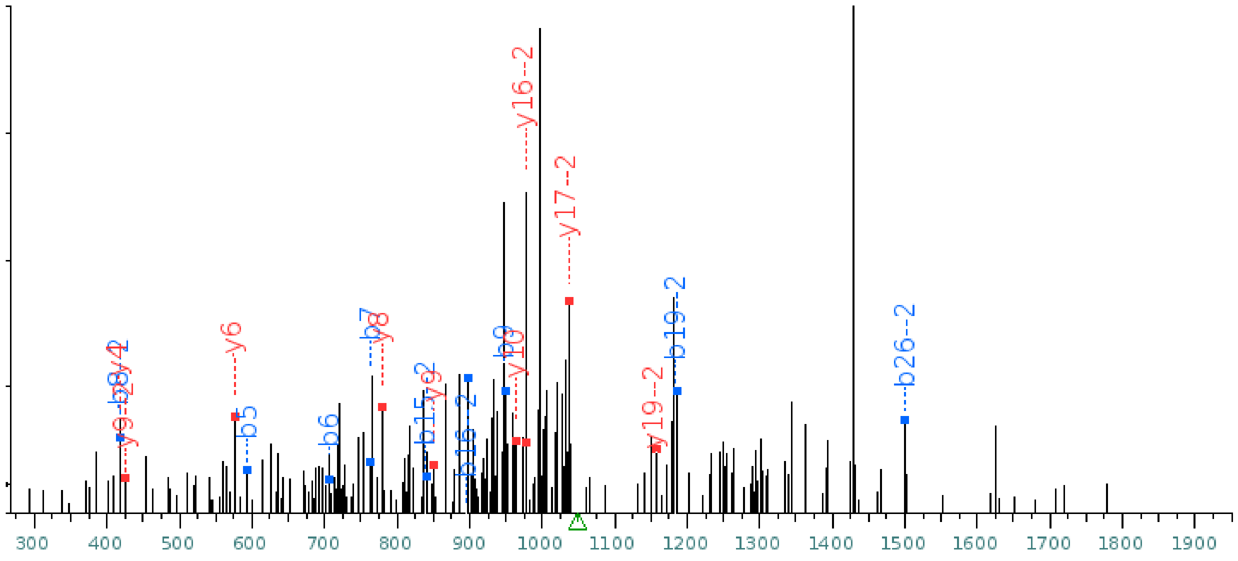

**Predicted Fragmentation Pattern**

| Seq | # | b: Δ Error | b | y | y: Δ Error | +1 |
|---|---|---|---|---|---|---|
| S | 1 | --- | 88.039 | --- | --- | 27 |
| M | 2 | --- | 219.080 | 3060.494 | --- | 26 |
| V | 3 | --- | 318.148 | 2929.453 | --- | 25 |
| F | 4 | --- | 465.217 | 2830.385 | --- | 24 |
| Q | 5 | 28.107 | **593.275** | 2683.316 | --- | 23 |
| I | 6 | 557.187 | **706.359** | 2555.258 | --- | 22 |
| G | 7 | 479.672 | **763.381** | 2442.174 | --- | 21 |
| A | 8 | --- | 834.418 | 2385.152 | --- | 20 |
| N | 9 | 456.873 | **948.461** | 2314.115 | --- | 19 |
| K | 10 | --- | 1076.556 | 2200.072 | --- | 18 |
| D | 11 | --- | 1191.583 | 2071.977 | --- | 17 |
| Q | 12 | --- | 1319.641 | 1956.950 | --- | 16 |
| V | 13 | --- | 1418.710 | 1828.892 | --- | 15 |
| M | 14 | --- | 1549.750 | 1729.823 | --- | 14 |
| E | 15 | --- | 1678.793 | 1598.783 | --- | 13 |
| L | 16 | --- | 1791.877 | 1469.740 | --- | 12 |
| T@ | 17 | --- | 2184.020 | 1356.656 | --- | 11 |
| I | 18 | --- | 2297.104 | **964.513** | 427.458 | 10 |

| Run: | 77660 | Scans: | 25436 - 25436 | m/z Range: | 292.93436 to 1958.2143 |
|---|---|---|---|---|---|
| Search: | 77660 | Precursor Mass: | 1044.5135 | | |
| Mods: | C=160.03 *+15.99 #+220.06 @+291.1 | Charge: | +3 | | |

**Sequence**

S@MVFQIGANKDQVMELTIAGMGTSALK

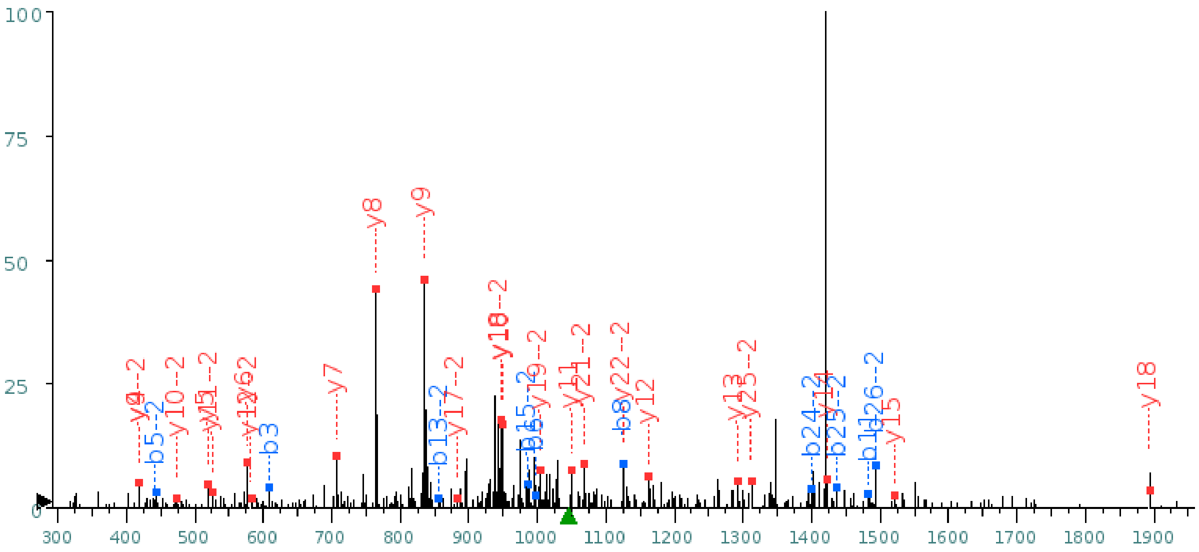

**Predicted Fragmentation Pattern**

**+1**

| Seq | # | b: Δ Error | b | y | y: Δ Error | +1 |
|---|---|---|---|---|---|---|
| S@ | 1 | --- | 379.135 | --- | --- | 27 |
| M | 2 | --- | 510.175 | 2753.403 | --- | 26 |
| V | 3 | -331.263 | **609.244** | 2622.363 | --- | 25 |
| F | 4 | --- | 756.312 | 2523.295 | --- | 24 |
| Q | 5 | --- | 884.371 | 2376.226 | --- | 23 |
| I | 6 | 274.583 | **997.455** | 2248.168 | --- | 22 |
| G | 7 | --- | 1054.476 | 2135.084 | --- | 21 |
| A | 8 | 31.279 | **1125.513** | 2078.062 | --- | 20 |
| N | 9 | --- | 1239.556 | 2007.025 | --- | 19 |
| K | 10 | --- | 1367.651 | **1892.982** | 97.975 | 18 |
| D | 11 | 70.149 | **1482.678** | 1764.887 | --- | 17 |
| Q | 12 | --- | 1610.737 | 1649.860 | --- | 16 |
| V | 13 | --- | 1709.805 | **1521.802** | 103.065 | 15 |
| M | 14 | --- | 1840.846 | **1422.733** | 62.068 | 14 |
| E | 15 | --- | 1969.888 | **1291.693** | -25.602 | 13 |
| L | 16 | --- | 2082.972 | **1162.650** | -554.879 | 12 |
| T | 17 | --- | 2184.020 | **1049.566** | 338.529 | 11 |
| I | 18 | --- | 2297.104 | **948.518** | -55.824 | 10 |
| A | 19 | --- | 2368.141 | **835.434** | 185.812 | 9 |

| | | | | | |
|---|---|---|---|---|---|
| **Run:** | 77660 | **Scans:** | 26087 - 26087 | **m/z Range:** | 271.13864 to 2000 |
| **Search:** | 77660 | **Precursor Mass:** | 1044.51192 | | |
| **Mods:** | C=160.03 *+15.99 #+220.06 @+291.1 | **Charge:** | +3 | | |

### Sequence

SMVFQIGANKDQVMELT@IAGMGTSALK

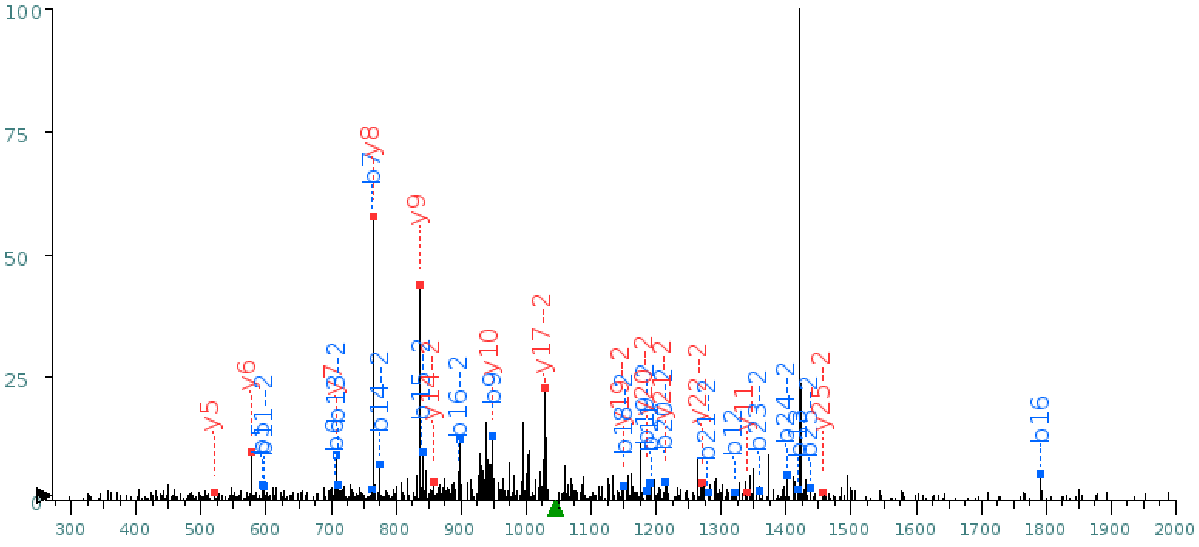

### Predicted Fragmentation Pattern

+1

| Seq | # | b: Δ Error | b | y | y: Δ Error | +1 |
|---|---|---|---|---|---|---|
| S | 1 | --- | 88.039 | --- | --- | 27 |
| M | 2 | --- | 219.080 | 3044.499 | --- | 26 |
| V | 3 | --- | 318.148 | 2913.458 | --- | 25 |
| F | 4 | --- | 465.217 | 2814.390 | --- | 24 |
| Q | 5 | 481.158 | **593.275** | 2667.322 | --- | 23 |
| I | 6 | 1385.439 | **706.359** | 2539.263 | --- | 22 |
| G | 7 | 228.955 | **763.381** | 2426.179 | --- | 21 |
| A | 8 | --- | 834.418 | 2369.157 | --- | 20 |
| N | 9 | 87.180 | **948.461** | 2298.120 | --- | 19 |
| K | 10 | --- | 1076.556 | 2184.077 | --- | 18 |
| D | 11 | -467.796 | **1191.583** | 2055.982 | --- | 17 |
| Q | 12 | 325.032 | **1319.641** | 1940.956 | --- | 16 |
| V | 13 | 108.098 | **1418.710** | 1812.897 | --- | 15 |
| M | 14 | --- | 1549.750 | 1713.829 | --- | 14 |
| E | 15 | --- | 1678.793 | 1582.788 | --- | 13 |
| L | 16 | -44.948 | **1791.877** | 1453.745 | --- | 12 |
| T@ | 17 | --- | 2184.020 | **1340.661** | -604.042 | 11 |
| I | 18 | --- | 2297.104 | **948.518** | 26.544 | 10 |

| Run: | 77660 | Scans: | 26590 - 26590 | m/z Range: | 311.40424 to 2000 |
|---|---|---|---|---|---|
| Search: | 77660 | Precursor Mass: | 1044.51459 | | |
| Mods: | C=160.03 *+15.99 #+220.06 @+291.1 | Charge: | +3 | | |

**Sequence**

S@MVFQIGANKDQVMELTIAGMGTSALK

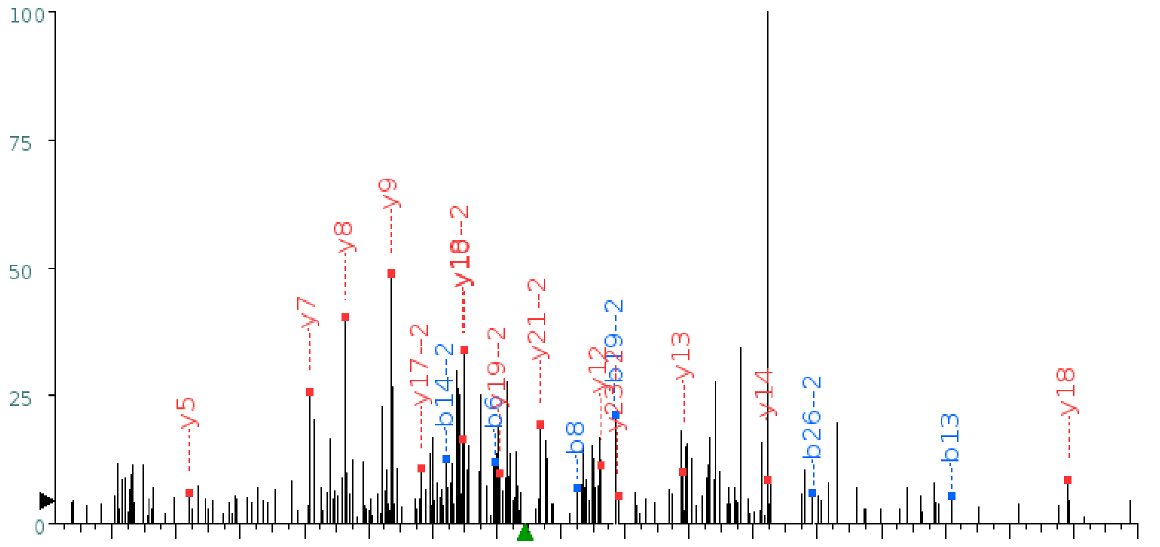

**Predicted Fragmentation Pattern**

| Seq | # | b: Δ Error | b | y | y: Δ Error | +1 |
|---|---|---|---|---|---|---|
| S@ | 1 | --- | 379.135 | --- | --- | 27 |
| M | 2 | --- | 510.175 | 2753.403 | --- | 26 |
| V | 3 | --- | 609.244 | 2622.363 | --- | 25 |
| F | 4 | --- | 756.312 | 2523.295 | --- | 24 |
| Q | 5 | --- | 884.371 | 2376.226 | --- | 23 |
| I | 6 | 369.368 | **997.455** | 2248.168 | --- | 22 |
| G | 7 | --- | 1054.476 | 2135.084 | --- | 21 |
| A | 8 | 184.063 | **1125.513** | 2078.062 | --- | 20 |
| N | 9 | --- | 1239.556 | 2007.025 | --- | 19 |
| K | 10 | --- | 1367.651 | **1892.982** | -501.209 | 18 |
| D | 11 | --- | 1482.678 | 1764.887 | --- | 17 |
| Q | 12 | --- | 1610.737 | 1649.860 | --- | 16 |
| V | 13 | -19.650 | **1709.805** | 1521.802 | --- | 15 |
| M | 14 | --- | 1840.846 | **1422.733** | -133.055 | 14 |
| E | 15 | --- | 1969.888 | **1291.693** | -402.267 | 13 |
| L | 16 | --- | 2082.972 | **1162.650** | -530.599 | 12 |
| T | 17 | --- | 2184.020 | 1049.566 | --- | 11 |
| I | 18 | --- | 2297.104 | **948.518** | 164.994 | 10 |

| Run: | 77660 | Scans: | 28735 - 28735 | m/z Range: | 301.28613 | to | 1996.2379 |
|---|---|---|---|---|---|---|---|
| Search: | 77660 | Precursor Mass: | 1146.87755 | | | | |
| Mods: | C=160.03 *+15.99 #+220.06 @+291.1 | Charge: | +3 | | | | |

**Sequence**

SMVFQIGANKDQVM*ELTIAGMGT@S@ALK

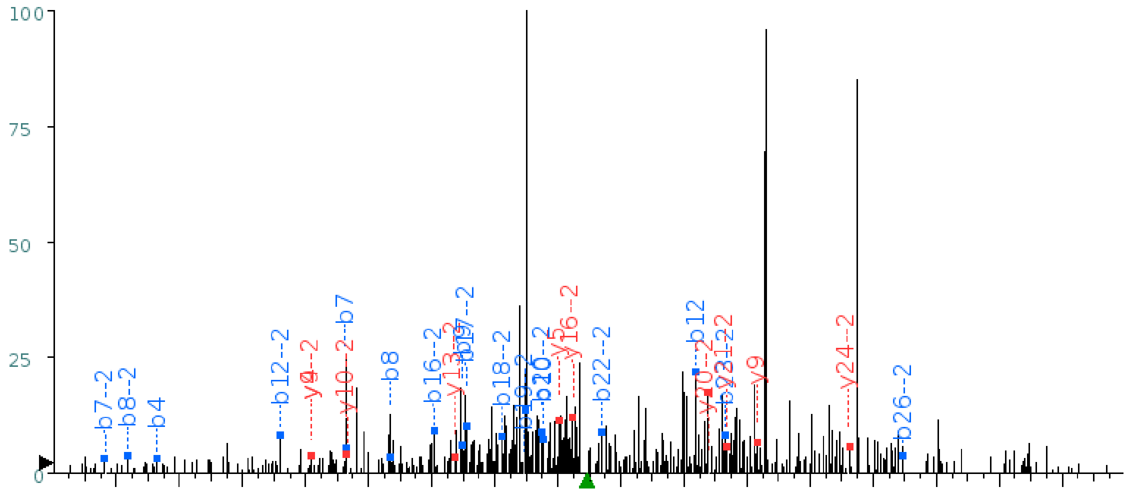

**Predicted Fragmentation Pattern**

| | | | +1 | | | |
|---|---|---|---|---|---|---|
| Seq | # | b: Δ Error | b | y | y: Δ Error | +1 |
| S | 1 | --- | 88.039 | --- | --- | 27 |
| M | 2 | --- | 219.080 | 3351.589 | --- | 26 |
| V | 3 | --- | 318.148 | 3220.549 | --- | 25 |
| F | 4 | 259.238 | **465.217** | 3121.480 | --- | 24 |
| Q | 5 | --- | 593.275 | 2974.412 | --- | 23 |
| I | 6 | --- | 706.359 | 2846.353 | --- | 22 |
| G | 7 | 371.827 | **763.381** | 2733.269 | --- | 21 |
| A | 8 | -660.794 | **834.418** | 2676.248 | --- | 20 |
| N | 9 | -454.346 | **948.461** | 2605.211 | --- | 19 |
| K | 10 | -769.886 | **1076.556** | 2491.168 | --- | 18 |
| D | 11 | --- | 1191.583 | 2363.073 | --- | 17 |
| Q | 12 | 185.610 | **1319.641** | 2248.046 | --- | 16 |
| V | 13 | --- | 1418.710 | 2119.987 | --- | 15 |
| M* | 14 | --- | 1565.745 | 2020.919 | --- | 14 |
| E | 15 | --- | 1694.788 | 1873.883 | --- | 13 |
| L | 16 | --- | 1807.872 | 1744.841 | --- | 12 |
| T | 17 | --- | 1908.919 | 1631.757 | --- | 11 |
| I | 18 | --- | 2022.003 | 1530.709 | --- | 10 |
| A | 19 | --- | 2093.041 | **1417.625** | -152.733 | 9 |
| G | 20 | --- | 2150.062 | 1346.588 | --- | 8 |

| Run: | 77660 | Scans: | 28861 - 28861 | m/z Range: | 303.39774 to 2000 |
|---|---|---|---|---|---|
| Search: | 77660 | Precursor Mass: | 1141.54697 | | |
| Mods: | C=160.03 *+15.99 #+220.06 @+291.1 | Charge: | +3 | | |

**Sequence**

S@MVFQIGANKDQVMELT@IAGMGTSALK

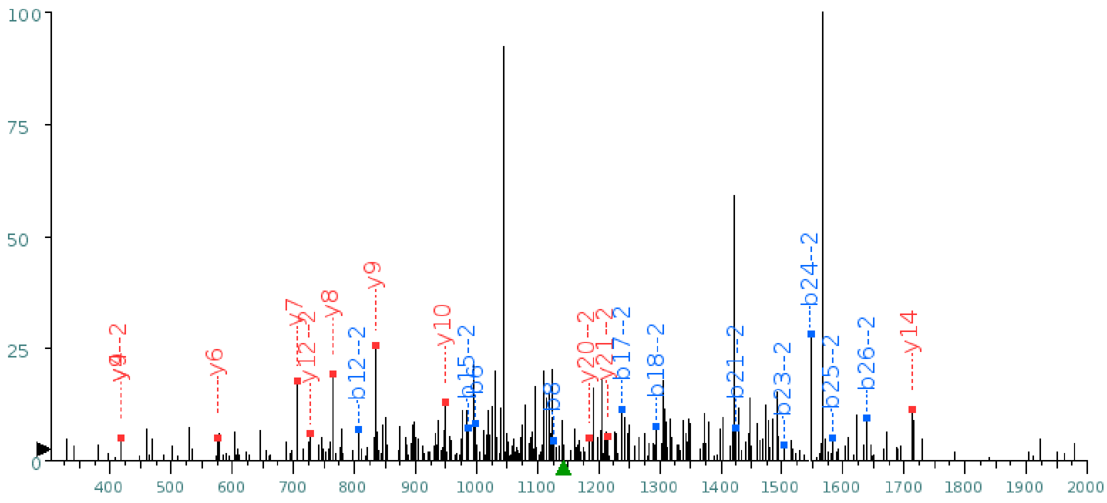

**Predicted Fragmentation Pattern**

+1

| Seq | # | b: Δ Error | b | y | y: Δ Error | +1 |
|---|---|---|---|---|---|---|
| S@ | 1 | --- | 379.135 | --- | --- | 27 |
| M | 2 | --- | 510.175 | 3044.499 | --- | 26 |
| V | 3 | --- | 609.244 | 2913.458 | --- | 25 |
| F | 4 | --- | 756.312 | 2814.390 | --- | 24 |
| Q | 5 | --- | 884.371 | 2667.322 | --- | 23 |
| I | 6 | -565.935 | **997.455** | 2539.263 | --- | 22 |
| G | 7 | --- | 1054.476 | 2426.179 | --- | 21 |
| A | 8 | 486.890 | **1125.513** | 2369.157 | --- | 20 |
| N | 9 | --- | 1239.556 | 2298.120 | --- | 19 |
| K | 10 | --- | 1367.651 | 2184.077 | --- | 18 |
| D | 11 | --- | 1482.678 | 2055.982 | --- | 17 |
| Q | 12 | --- | 1610.737 | 1940.956 | --- | 16 |
| V | 13 | --- | 1709.805 | 1812.897 | --- | 15 |
| M | 14 | --- | 1840.846 | **1713.829** | -230.559 | 14 |
| E | 15 | --- | 1969.888 | 1582.788 | --- | 13 |
| L | 16 | --- | 2082.972 | 1453.745 | --- | 12 |
| T@ | 17 | --- | 2475.115 | 1340.661 | --- | 11 |
| I | 18 | --- | 2588.199 | **948.518** | 283.984 | 10 |
| A | 19 | --- | 2659.236 | **835.434** | 143.452 | 9 |
| G | 20 | --- | 2716.258 | **764.397** | 304.437 | 8 |
| M | 21 | --- | 2847.298 | **707.376** | 244.258 | 7 |

| Run: | 77660 | Scans: | 29105 - 29105 | m/z Range: | 306.10736 | to | 1999.1855 |
|---|---|---|---|---|---|---|---|
| Search: | 77660 | Precursor Mass: | 1141.5463 | | | | |
| Mods: | C=160.03 *+15.99 #+220.06 @+291.1 | Charge: | +3 | | | | |

**Sequence**

S@MVFQIGANKDQVMELT@IAGMGTSALK

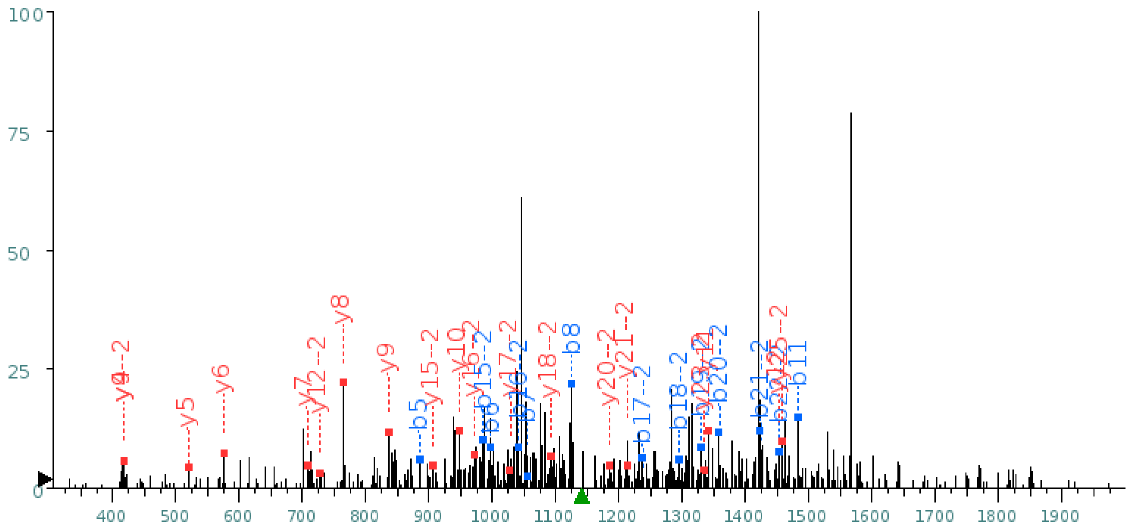

**Predicted Fragmentation Pattern**

| Seq | # | b: Δ Error | b | y | y: Δ Error | +1 |
|---|---|---|---|---|---|---|
| S@ | 1 | --- | 379.135 | --- | --- | 27 |
| M | 2 | --- | 510.175 | 3044.499 | --- | 26 |
| V | 3 | --- | 609.244 | 2913.458 | --- | 25 |
| F | 4 | --- | 756.312 | 2814.390 | --- | 24 |
| Q | 5 | 422.599 | **884.371** | 2667.322 | --- | 23 |
| I | 6 | -650.481 | **997.455** | 2539.263 | --- | 22 |
| G | 7 | 502.677 | **1054.476** | 2426.179 | --- | 21 |
| A | 8 | -564.904 | **1125.513** | 2369.157 | --- | 20 |
| N | 9 | --- | 1239.556 | 2298.120 | --- | 19 |
| K | 10 | --- | 1367.651 | 2184.077 | --- | 18 |
| D | 11 | 47.016 | **1482.678** | 2055.982 | --- | 17 |
| Q | 12 | --- | 1610.737 | 1940.956 | --- | 16 |
| V | 13 | --- | 1709.805 | 1812.897 | --- | 15 |
| M | 14 | --- | 1840.846 | 1713.829 | --- | 14 |
| E | 15 | --- | 1969.888 | 1582.788 | --- | 13 |
| L | 16 | --- | 2082.972 | **1453.745** | -361.170 | 12 |
| T@ | 17 | --- | 2475.115 | **1340.661** | -464.947 | 11 |
| I | 18 | --- | 2588.199 | **948.518** | -228.454 | 10 |
| A | 19 | --- | 2659.236 | **835.434** | 160.689 | 9 |
| G | 20 | --- | 2716.258 | **764.397** | 328.695 | 8 |

| | | | | | |
|---|---|---|---|---|---|
| **Run:** | 77660 | **Scans:** | 29339 - 29339 | **m/z Range:** | 342.44363 **to** 1998.0467! |
| **Search:** | 77660 | **Precursor Mass:** | 1141.54481 | | |
| **Mods:** | C=160.03 *+15.99 #+220.06 @+291.1 | **Charge:** | +3 | | |

**Sequence**

S@MVFQIGANKDQVMELT@IAGMGTSALK

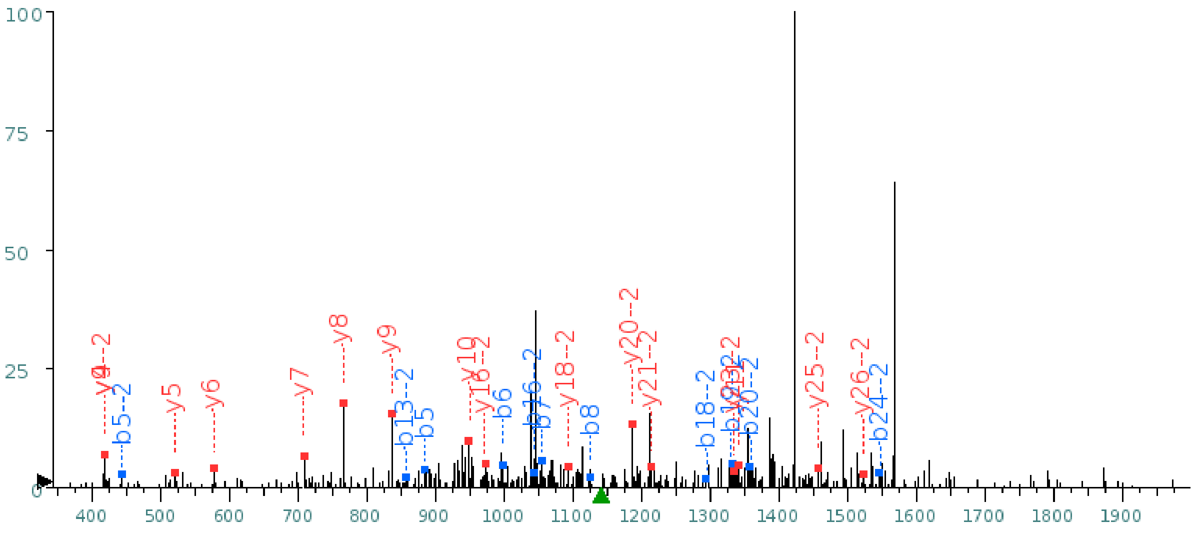

**Predicted Fragmentation Pattern**

**+1**

| Seq | # | b: Δ Error | b | y | y: Δ Error | +1 |
|---|---|---|---|---|---|---|
| S@ | 1 | --- | 379.135 | --- | --- | 27 |
| M | 2 | --- | 510.175 | 3044.499 | --- | 26 |
| V | 3 | --- | 609.244 | 2913.458 | --- | 25 |
| F | 4 | --- | 756.312 | 2814.390 | --- | 24 |
| Q | 5 | -896.080 | **884.371** | 2667.322 | --- | 23 |
| I | 6 | -286.789 | **997.455** | 2539.263 | --- | 22 |
| G | 7 | 639.354 | **1054.476** | 2426.179 | --- | 21 |
| A | 8 | -359.947 | **1125.513** | 2369.157 | --- | 20 |
| N | 9 | --- | 1239.556 | 2298.120 | --- | 19 |
| K | 10 | --- | 1367.651 | 2184.077 | --- | 18 |
| D | 11 | --- | 1482.678 | 2055.982 | --- | 17 |
| Q | 12 | --- | 1610.737 | 1940.956 | --- | 16 |
| V | 13 | --- | 1709.805 | 1812.897 | --- | 15 |
| M | 14 | --- | 1840.846 | 1713.829 | --- | 14 |
| E | 15 | --- | 1969.888 | 1582.788 | --- | 13 |
| L | 16 | --- | 2082.972 | 1453.745 | --- | 12 |
| T@ | 17 | --- | 2475.115 | **1340.661** | -546.795 | 11 |
| I | 18 | --- | 2588.199 | **948.518** | -70.883 | 10 |
| A | 19 | --- | 2659.236 | **835.434** | 205.530 | 9 |

| Run: | 77660 | Scans: | 29383 - 29383 | m/z Range: | 330.38455 to 2000 |
|---|---|---|---|---|---|
| Search: | 77660 | Precursor Mass: | 1199.5661 | | |
| Mods: | C=160.03 *+15.99 #+220.06 @+291.1 | Charge: | +2 | | |

### Sequence

S@ISGMIETIDNAINTVSGER

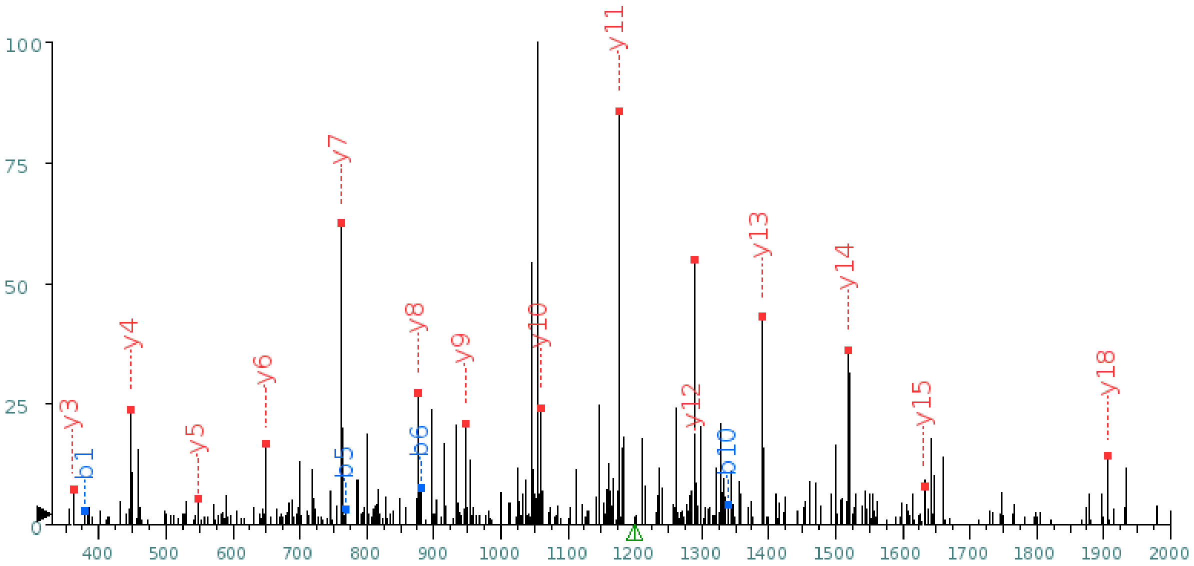

### Predicted Fragmentation Pattern

| Seq | # | b: Δ Error | b | y | y: Δ Error | +1 |
|---|---|---|---|---|---|---|
| S@ | 1 | 577.471 | 379.135 | --- | --- | 20 |
| I | 2 | --- | 492.219 | 2020.002 | --- | 19 |
| S | 3 | --- | 579.251 | 1906.917 | 96.716 | 18 |
| G | 4 | --- | 636.272 | 1819.885 | --- | 17 |
| M | 5 | 257.470 | 767.313 | 1762.864 | --- | 16 |
| I | 6 | 774.360 | 880.397 | 1631.824 | 106.426 | 15 |
| E | 7 | --- | 1009.439 | 1518.739 | 144.535 | 14 |
| T | 8 | --- | 1110.487 | 1389.697 | 160.933 | 13 |
| I | 9 | --- | 1223.571 | 1288.649 | 148.416 | 12 |
| D | 10 | 98.779 | 1338.598 | 1175.565 | 19.278 | 11 |
| N | 11 | --- | 1452.641 | 1060.538 | 194.528 | 10 |
| A | 12 | --- | 1523.678 | 946.495 | 194.068 | 9 |
| I | 13 | --- | 1636.762 | 875.458 | 257.342 | 8 |
| N | 14 | --- | 1750.805 | 762.374 | 272.403 | 7 |
| T | 15 | --- | 1851.853 | 648.331 | 314.595 | 6 |
| V | 16 | --- | 1950.921 | 547.283 | 296.557 | 5 |
| S | 17 | --- | 2037.953 | 448.215 | 273.202 | 4 |
| G | 18 | --- | 2094.975 | 361.183 | 1091.826 | 3 |
| E | 19 | --- | 2224.017 | 304.162 | --- | 2 |
| R | 20 | --- | --- | 175.119 | --- | 1 |

| | | | | | |
|---|---|---|---|---|---|
| **Run:** | 77660 | **Scans:** | 29679 - 29679 | **m/z Range:** | 272.1857 to 2000 |
| **Search:** | 77660 | **Precursor Mass:** | 1049.84614 | | |
| **Mods:** | C=160.03 *+15.99 #+220.06 @+291.1 | **Charge:** | +3 | | |

**Sequence**

SMVFQIGANKDQVMELT@IAGM*GTSALK

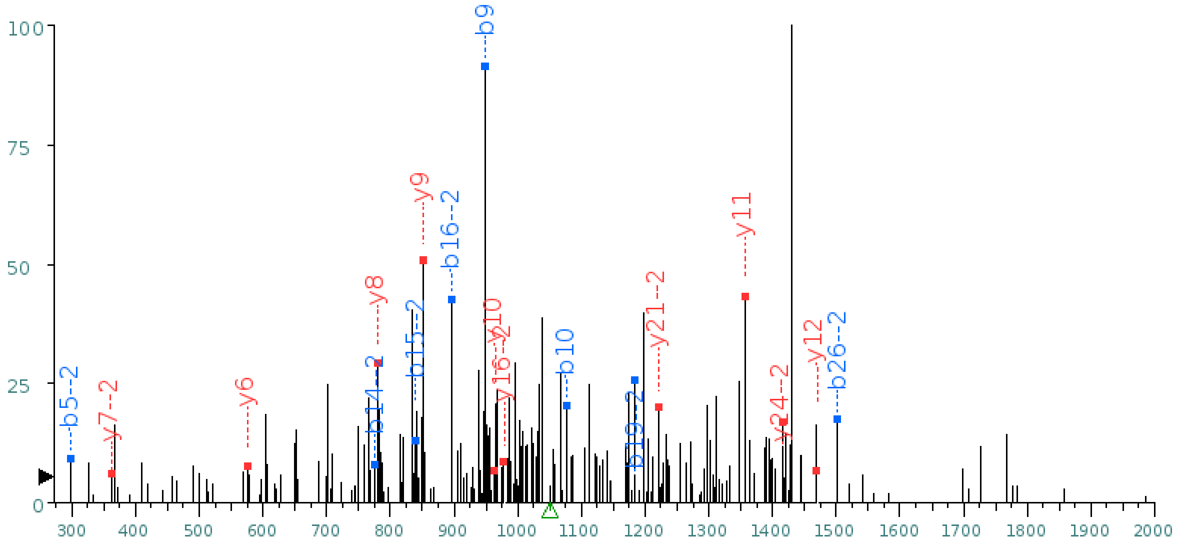

**Predicted Fragmentation Pattern**

| Seq | # | b: Δ Error | b | y | y: Δ Error | +1 |
|---|---|---|---|---|---|---|
| S | 1 | --- | 88.039 | --- | --- | 27 |
| M | 2 | --- | 219.080 | 3060.494 | --- | 26 |
| V | 3 | --- | 318.148 | 2929.453 | --- | 25 |
| F | 4 | --- | 465.217 | 2830.385 | --- | 24 |
| Q | 5 | --- | 593.275 | 2683.316 | --- | 23 |
| I | 6 | --- | 706.359 | 2555.258 | --- | 22 |
| G | 7 | --- | 763.381 | 2442.174 | --- | 21 |
| A | 8 | --- | 834.418 | 2385.152 | --- | 20 |
| N | 9 | -769.154 | **948.461** | 2314.115 | --- | 19 |
| K | 10 | -382.781 | **1076.556** | 2200.072 | --- | 18 |
| D | 11 | --- | 1191.583 | 2071.977 | --- | 17 |
| Q | 12 | --- | 1319.641 | 1956.950 | --- | 16 |
| V | 13 | --- | 1418.710 | 1828.892 | --- | 15 |
| M | 14 | --- | 1549.750 | 1729.823 | --- | 14 |
| E | 15 | --- | 1678.793 | 1598.783 | --- | 13 |
| L | 16 | --- | 1791.877 | **1469.740** | -479.461 | 12 |
| T@ | 17 | --- | 2184.020 | **1356.656** | -212.336 | 11 |
| I | 18 | --- | 2297.104 | **964.513** | -825.493 | 10 |
| A | 19 | --- | 2368.141 | **851.429** | 162.065 | 9 |
| G | 20 | --- | 2425.163 | **780.392** | 67.195 | 8 |

| | | | | | |
|---|---|---|---|---|---|
| **Run:** | 77660 | **Scans:** | 29737 - 29737 | **m/z Range:** | 301.45325 to 2000 |
| **Search:** | 77660 | **Precursor Mass:** | 1141.54604 | | |
| **Mods:** | C=160.03 *+15.99 #+220.06 @+291.1 | **Charge:** | +3 | | |

**Sequence**

S@MVFQIGANKDQVMELT@IAGMGTSALK

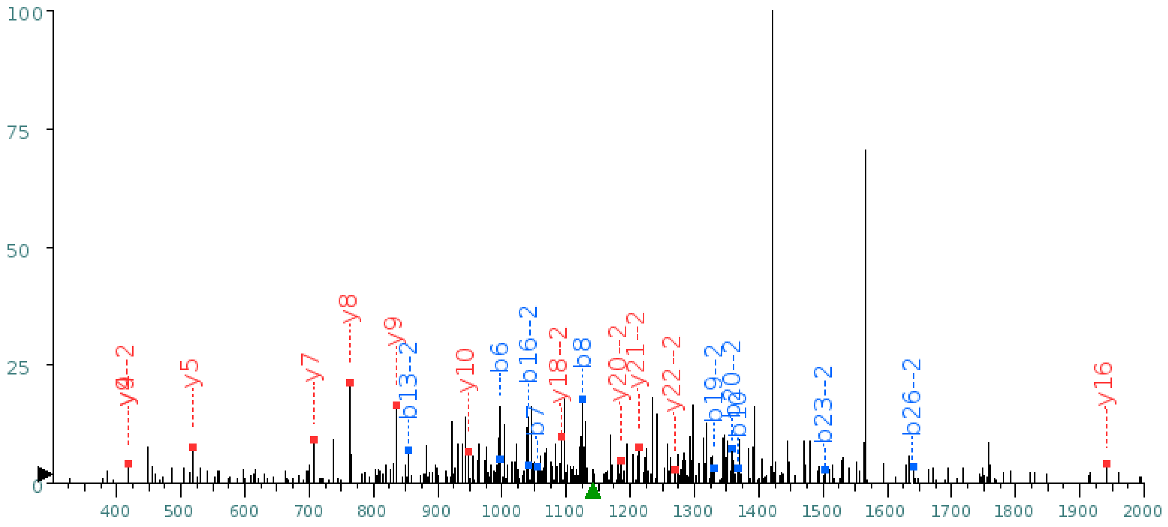

**Predicted Fragmentation Pattern**

| Seq | # | b: Δ Error | b | y | y: Δ Error | +1 |
|---|---|---|---|---|---|---|
| | | | +1 | | | |
| S@ | 1 | --- | 379.135 | --- | --- | 27 |
| M | 2 | --- | 510.175 | 3044.499 | --- | 26 |
| V | 3 | --- | 609.244 | 2913.458 | --- | 25 |
| F | 4 | --- | 756.312 | 2814.390 | --- | 24 |
| Q | 5 | --- | 884.371 | 2667.322 | --- | 23 |
| I | 6 | 174.030 | **997.455** | 2539.263 | --- | 22 |
| G | 7 | 269.477 | **1054.476** | 2426.179 | --- | 21 |
| A | 8 | -839.579 | **1125.513** | 2369.157 | --- | 20 |
| N | 9 | --- | 1239.556 | 2298.120 | --- | 19 |
| K | 10 | -305.453 | **1367.651** | 2184.077 | --- | 18 |
| D | 11 | --- | 1482.678 | 2055.982 | --- | 17 |
| Q | 12 | --- | 1610.737 | **1940.956** | 192.692 | 16 |
| V | 13 | --- | 1709.805 | 1812.897 | --- | 15 |
| M | 14 | --- | 1840.846 | 1713.829 | --- | 14 |
| E | 15 | --- | 1969.888 | 1582.788 | --- | 13 |
| L | 16 | --- | 2082.972 | 1453.745 | --- | 12 |
| T@ | 17 | --- | 2475.115 | 1340.661 | --- | 11 |
| I | 18 | --- | 2588.199 | **948.518** | 337.360 | 10 |
| A | 19 | --- | 2659.236 | **835.434** | 320.463 | 9 |
| G | 20 | --- | 2716.258 | **764.397** | 90.211 | 8 |

| | | | | | |
|---|---|---|---|---|---|
| **Run:** | 77660 | **Scans:** | 31259 - 31259 | **m/z Range:** | 272.28415 to 2000 |
| **Search:** | 77660 | **Precursor Mass:** | 1049.84774 | | |
| **Mods:** | C=160.03 *+15.99 #+220.06 @+291.1 | **Charge:** | +3 | | |

### Sequence

SMVFQIGANKDQVMELT@IAGM*GTSALK

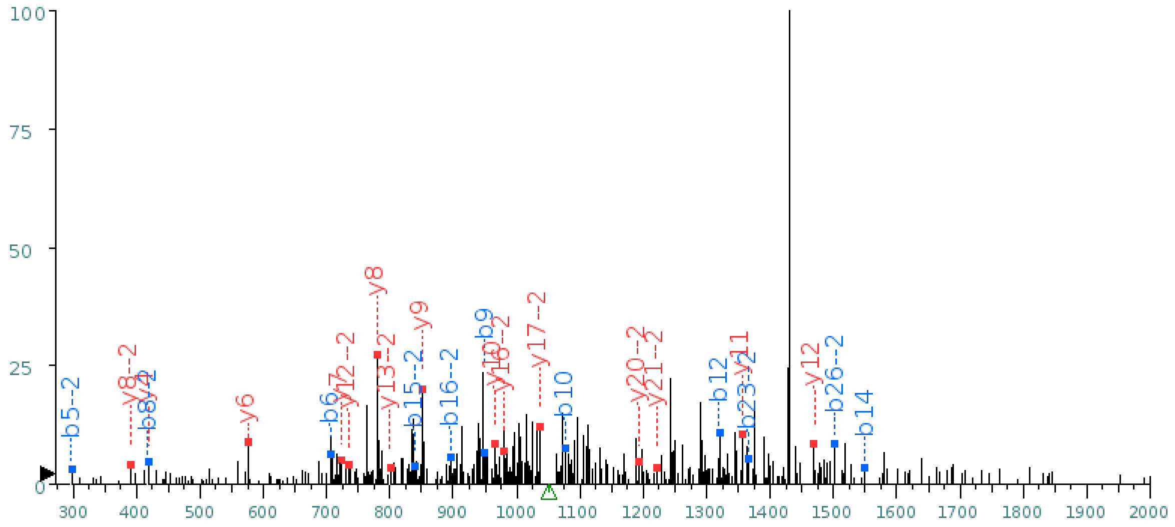

### Predicted Fragmentation Pattern

+1

| Seq | # | b: Δ Error | b | y | y: Δ Error | +1 |
|---|---|---|---|---|---|---|
| S | 1 | --- | 88.039 | --- | --- | 27 |
| M | 2 | --- | 219.080 | 3060.494 | --- | 26 |
| V | 3 | --- | 318.148 | 2929.453 | --- | 25 |
| F | 4 | --- | 465.217 | 2830.385 | --- | 24 |
| Q | 5 | --- | 593.275 | 2683.316 | --- | 23 |
| I | 6 | 2.585 | **706.359** | 2555.258 | --- | 22 |
| G | 7 | --- | 763.381 | 2442.174 | --- | 21 |
| A | 8 | --- | 834.418 | 2385.152 | --- | 20 |
| N | 9 | -456.021 | **948.461** | 2314.115 | --- | 19 |
| K | 10 | 679.478 | **1076.556** | 2200.072 | --- | 18 |
| D | 11 | --- | 1191.583 | 2071.977 | --- | 17 |
| Q | 12 | 587.685 | **1319.641** | 1956.950 | --- | 16 |
| V | 13 | --- | 1418.710 | 1828.892 | --- | 15 |
| M | 14 | -582.448 | **1549.750** | 1729.823 | --- | 14 |
| E | 15 | --- | 1678.793 | 1598.783 | --- | 13 |
| L | 16 | --- | 1791.877 | **1469.740** | -464.497 | 12 |
| T@ | 17 | --- | 2184.020 | **1356.656** | -282.194 | 11 |
| I | 18 | --- | 2297.104 | **964.513** | 105.025 | 10 |
| A | 19 | --- | 2368.141 | **851.429** | 294.479 | 9 |
| G | 20 | --- | 2425.163 | **780.392** | 141.480 | 8 |

| | | | | | |
|---|---|---|---|---|---|
| **Run:** | 77660 | **Scans:** | 31397 - 31397 | **m/z Range:** | 293.11197 to 2000 |
| **Search:** | 77660 | **Precursor Mass:** | 1049.84415 | | |
| **Mods:** | C=160.03 *+15.99 #+220.06 @+291.1 | **Charge:** | +3 | | |

### Sequence

SMVFQIGANKDQVMELT@IAGM*GTSALK

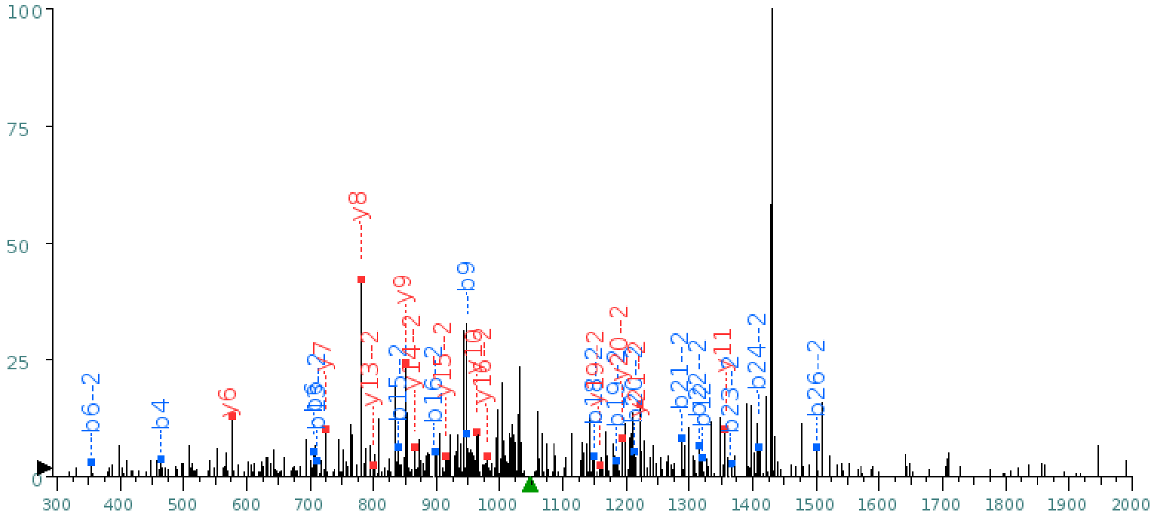

### Predicted Fragmentation Pattern

| | | +1 | | | | |
|---|---|---|---|---|---|---|
| **Seq** | **#** | **b: Δ Error** | **b** | **y** | **y: Δ Error** | **+1** |
| S | 1 | --- | 88.039 | --- | --- | 27 |
| M | 2 | --- | 219.080 | 3060.494 | --- | 26 |
| V | 3 | --- | 318.148 | 2929.453 | --- | 25 |
| F | 4 | -1844.955 | **465.217** | 2830.385 | --- | 24 |
| Q | 5 | --- | 593.275 | 2683.316 | --- | 23 |
| I | 6 | 39.566 | **706.359** | 2555.258 | --- | 22 |
| G | 7 | --- | 763.381 | 2442.174 | --- | 21 |
| A | 8 | --- | 834.418 | 2385.152 | --- | 20 |
| N | 9 | -506.135 | **948.461** | 2314.115 | --- | 19 |
| K | 10 | --- | 1076.556 | 2200.072 | --- | 18 |
| D | 11 | --- | 1191.583 | 2071.977 | --- | 17 |
| Q | 12 | 345.092 | **1319.641** | 1956.950 | --- | 16 |
| V | 13 | --- | 1418.710 | 1828.892 | --- | 15 |
| M | 14 | --- | 1549.750 | 1729.823 | --- | 14 |
| E | 15 | --- | 1678.793 | 1598.783 | --- | 13 |
| L | 16 | --- | 1791.877 | 1469.740 | --- | 12 |
| T@ | 17 | --- | 2184.020 | **1356.656** | -272.831 | 11 |
| I | 18 | --- | 2297.104 | **964.513** | 162.721 | 10 |
| A | 19 | --- | 2368.141 | **851.429** | 80.506 | 9 |
| G | 20 | --- | 2425.163 | **780.392** | 42.796 | 8 |

| | | | | | |
|---|---|---|---|---|---|
| **Run:** | 77660 | **Scans:** | 31680 - 31680 | **m/z Range:** | 311.25717 to 2000 |
| **Search:** | 77660 | **Precursor Mass:** | 1141.54643 | | |
| **Mods:** | C=160.03 *+15.99 #+220.06 @+291.1 | **Charge:** | +3 | | |

#### Sequence

S@MVFQIGANKDQVMELT@IAGMGTSALK

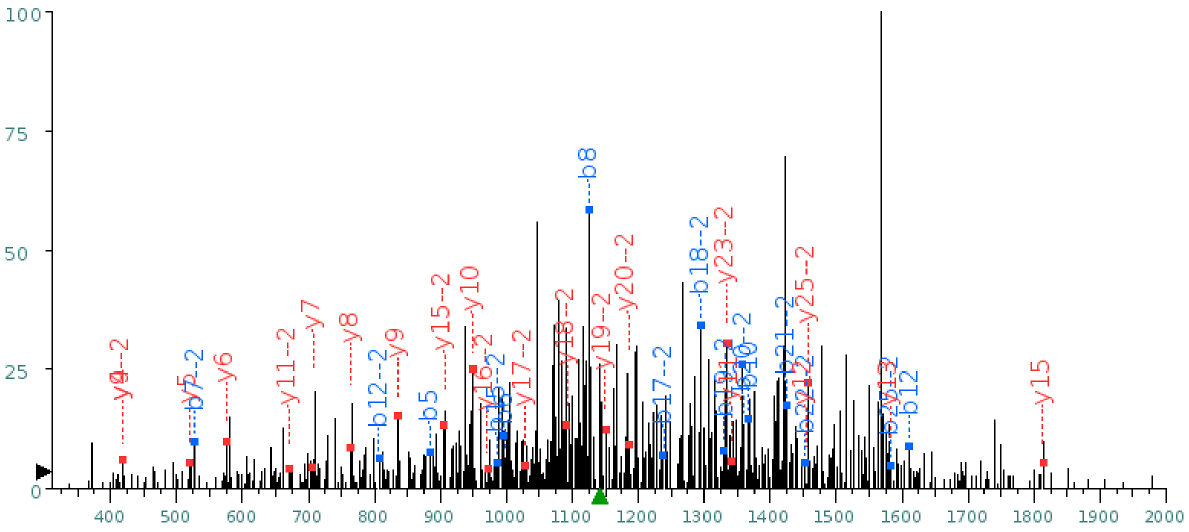

#### Predicted Fragmentation Pattern

+1

| Seq | # | b: Δ Error | b | y | y: Δ Error | +1 |
|---|---|---|---|---|---|---|
| S@ | 1 | --- | 379.135 | --- | --- | 27 |
| M | 2 | --- | 510.175 | 3044.499 | --- | 26 |
| V | 3 | --- | 609.244 | 2913.458 | --- | 25 |
| F | 4 | --- | 756.312 | 2814.390 | --- | 24 |
| Q | 5 | 203.130 | **884.371** | 2667.322 | --- | 23 |
| I | 6 | -831.140 | **997.455** | 2539.263 | --- | 22 |
| G | 7 | --- | 1054.476 | 2426.179 | --- | 21 |
| A | 8 | 199.566 | **1125.513** | 2369.157 | --- | 20 |
| N | 9 | --- | 1239.556 | 2298.120 | --- | 19 |
| K | 10 | -569.166 | **1367.651** | 2184.077 | --- | 18 |
| D | 11 | --- | 1482.678 | 2055.982 | --- | 17 |
| Q | 12 | 21.085 | **1610.737** | 1940.956 | --- | 16 |
| V | 13 | --- | 1709.805 | **1812.897** | 359.526 | 15 |
| M | 14 | --- | 1840.846 | 1713.829 | --- | 14 |
| E | 15 | --- | 1969.888 | **1582.788** | -298.911 | 13 |
| L | 16 | --- | 2082.972 | **1453.745** | -222.036 | 12 |
| T@ | 17 | --- | 2475.115 | **1340.661** | 421.485 | 11 |
| I | 18 | --- | 2588.199 | **948.518** | 213.365 | 10 |
| A | 19 | --- | 2659.236 | **835.434** | 1194.664 | 9 |
| G | 20 | --- | 2716.258 | **764.397** | -982.938 | 8 |

| Run: | 77660 | Scans: | 33267 - 33267 | m/z Range: | 306.34604 to 1993.3598 |
|---|---|---|---|---|---|
| Search: | 77660 | Precursor Mass: | 1146.87909 | | |
| Mods: | C=160.03 *+15.99 #+220.06 @+291.1 | Charge: | +3 | | |

**Sequence**

S@MVFQIGANKDQVM*ELT@IAGMGTSALK

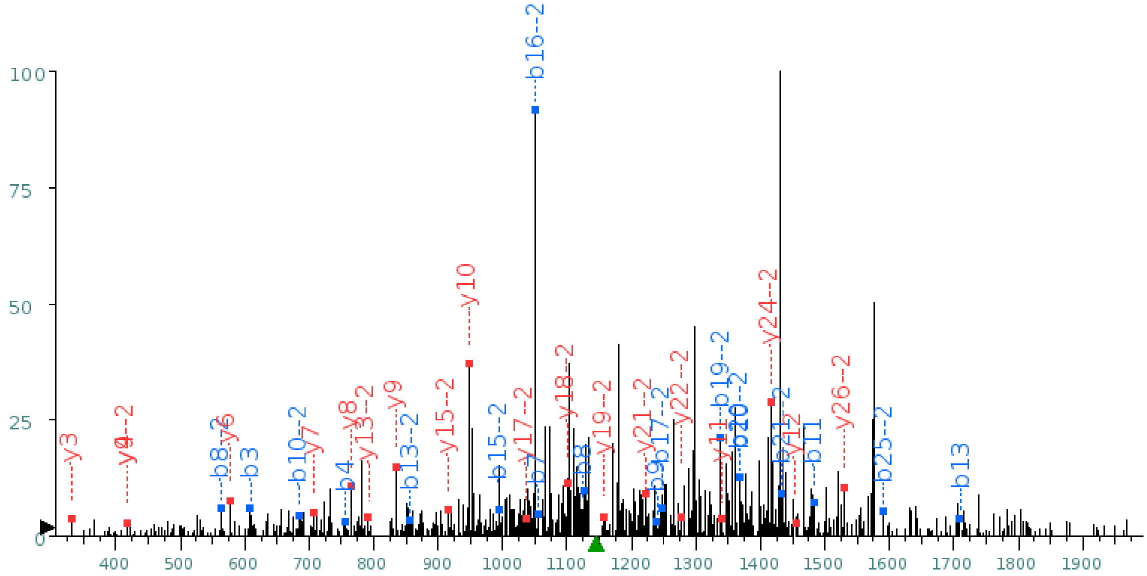

**Predicted Fragmentation Pattern**

+1

| Seq | # | b: Δ Error | b | y | y: Δ Error | +1 |
|---|---|---|---|---|---|---|
| S@ | 1 | --- | 379.135 | --- | --- | 27 |
| M | 2 | --- | 510.175 | 3060.494 | --- | 26 |
| V | 3 | -1373.526 | **609.244** | 2929.453 | --- | 25 |
| F | 4 | -888.857 | **756.312** | 2830.385 | --- | 24 |
| Q | 5 | --- | 884.371 | 2683.316 | --- | 23 |
| I | 6 | --- | 997.455 | 2555.258 | --- | 22 |
| G | 7 | -11.525 | **1054.476** | 2442.174 | --- | 21 |
| A | 8 | 789.425 | **1125.513** | 2385.152 | --- | 20 |
| N | 9 | -423.171 | **1239.556** | 2314.115 | --- | 19 |
| K | 10 | -623.677 | **1367.651** | 2200.072 | --- | 18 |
| D | 11 | 152.461 | **1482.678** | 2071.977 | --- | 17 |
| Q | 12 | --- | 1610.737 | 1956.950 | --- | 16 |
| V | 13 | -397.912 | **1709.805** | 1828.892 | --- | 15 |
| M* | 14 | --- | 1856.840 | 1729.823 | --- | 14 |
| E | 15 | --- | 1985.883 | 1582.788 | --- | 13 |
| L | 16 | --- | 2098.967 | **1453.745** | 677.539 | 12 |
| T@ | 17 | --- | 2491.110 | **1340.661** | -52.540 | 11 |
| I | 18 | --- | 2604.194 | **948.518** | -769.129 | 10 |
| A | 19 | --- | 2675.231 | **835.434** | 252.120 | 9 |
| G | 20 | --- | 2732.253 | **764.397** | -78.425 | 8 |

| | | | |
|---|---|---|---|
| **Run:** | 77660 | **Scans:** | 33278 - 33278 |
| **Search:** | 77660 | **Precursor Mass:** | 1044.5148 |
| **Mods:** | C=160.03 *+15.99 #+220.06 @+291.1 | **Charge:** | +3 |

**m/z Range:** 286.33109 to 1998.4512[illegible]

**Sequence**

SMVFQIGANKDQVMELT@IAGMGTSALK

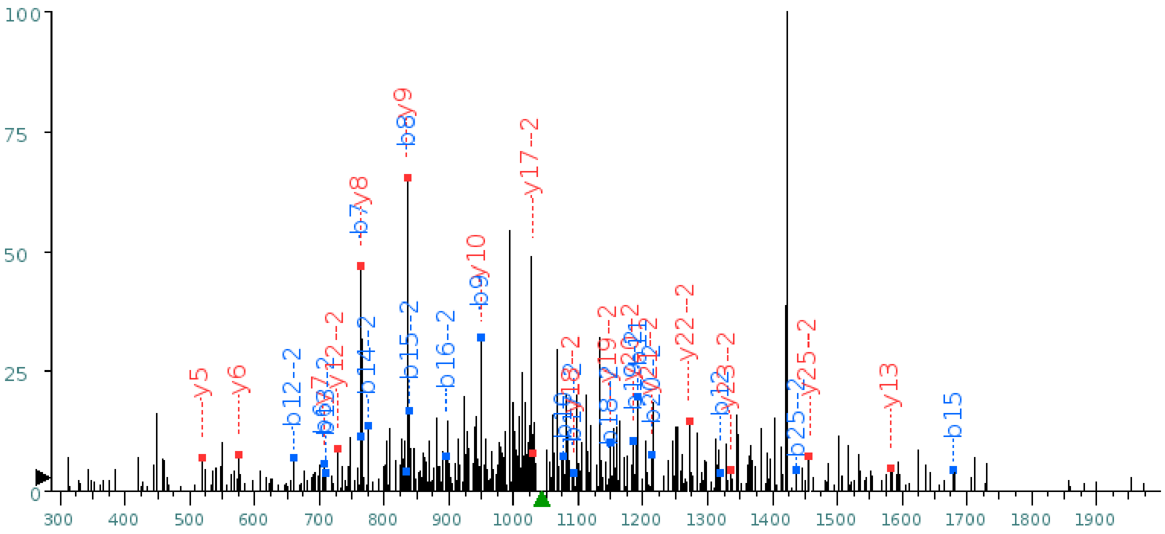

**Predicted Fragmentation Pattern**

| Seq | # | b: Δ Error | +1 b | +1 y | y: Δ Error | +1 |
|---|---|---|---|---|---|---|
| S | 1 | --- | 88.039 | --- | --- | 27 |
| M | 2 | --- | 219.080 | 3044.499 | --- | 26 |
| V | 3 | --- | 318.148 | 2913.458 | --- | 25 |
| F | 4 | --- | 465.217 | 2814.390 | --- | 24 |
| Q | 5 | --- | 593.275 | 2667.322 | --- | 23 |
| I | 6 | 336.180 | **706.359** | 2539.263 | --- | 22 |
| G | 7 | -701.666 | **763.381** | 2426.179 | --- | 21 |
| A | 8 | -784.152 | **834.418** | 2369.157 | --- | 20 |
| N | 9 | 163.996 | **948.461** | 2298.120 | --- | 19 |
| K | 10 | 724.431 | **1076.556** | 2184.077 | --- | 18 |
| D | 11 | 0.796 | **1191.583** | 2055.982 | --- | 17 |
| Q | 12 | 169.705 | **1319.641** | 1940.956 | --- | 16 |
| V | 13 | --- | 1418.710 | 1812.897 | --- | 15 |
| M | 14 | --- | 1549.750 | 1713.829 | --- | 14 |
| E | 15 | 197.587 | **1678.793** | **1582.788** | 99.975 | 13 |
| L | 16 | --- | 1791.877 | 1453.745 | --- | 12 |
| T@ | 17 | --- | 2184.020 | 1340.661 | --- | 11 |
| I | 18 | --- | 2297.104 | **948.518** | 103.365 | 10 |
| A | 19 | --- | 2368.141 | **835.434** | 210.642 | 9 |

**MS/MS data of trypsin digested peptides with site score A and site score B above 19 from *Clostridium botulinum* CbFla flagellin protein co-expressed with *Clostridium botulinum* Maf in EV36 cells**

**Results of dataset 2 (MS/MS results obtained in April 2019).**

| | | | | | |
|---|---|---|---|---|---|
| **Run:** | 77664 | **Scans:** | 19886 - 19886 | **m/z Range:** | 202.38829 to 1462.2663 |
| **Search:** | 77664 | **Precursor Mass:** | 783.3775 | | |
| **Mods:** | C=160.03 *+15.99 #+220.06 @+291.1 | **Charge:** | +3 | | |

**Sequence**

NLLNGS# AKS# M* VFQIGANK

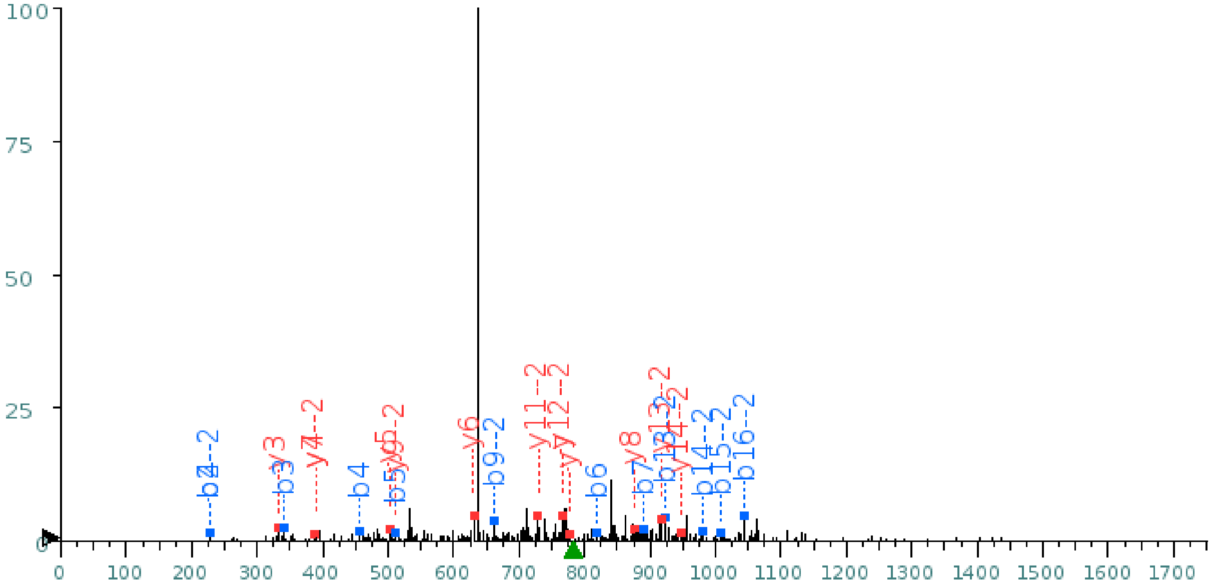

**Predicted Fragmentation Pattern**

| | | | +1 | | | |
|---|---|---|---|---|---|---|
| **Seq** | **#** | **b: Δ Error** | **b** | **y** | **y: Δ Error** | **+1** |
| **N** | **1** | --- | 115.050 | --- | --- | **18** |
| **L** | **2** | **1219.275** | **228.134** | 2234.074 | --- | **17** |
| **L** | **3** | **507.541** | **341.218** | 2120.990 | --- | **16** |
| **N** | **4** | **689.501** | **455.261** | 2007.906 | --- | **15** |
| **G** | **5** | **-1705.838** | **512.283** | 1893.863 | --- | **14** |
| **S#** | **6** | **-401.444** | **819.373** | 1836.842 | --- | **13** |
| **A** | **7** | **29.122** | **890.410** | 1529.752 | --- | **12** |
| **K** | **8** | --- | 1018.505 | 1458.714 | --- | **11** |
| **S#** | **9** | --- | 1325.595 | 1330.620 | --- | **10** |
| **M*** | **10** | --- | 1472.631 | 1023.529 | --- | **9** |
| **V** | **11** | --- | 1571.699 | **876.494** | **-858.024** | **8** |
| **F** | **12** | --- | 1718.768 | **777.425** | **510.429** | **7** |

| Run: | 77664 | Scans: | 21869 - 21869 | m/z Range: | 269.55417 to 2000 |
|---|---|---|---|---|---|
| Search: | 77664 | Precursor Mass: | 1036.82936 | | |
| Mods: | C=160.03 *+15.99 #+220.06 @+291.1 | Charge: | +3 | | |

**Sequence**

S#M*VFQIGANKDQVM*ELTIAGM*GTSALK

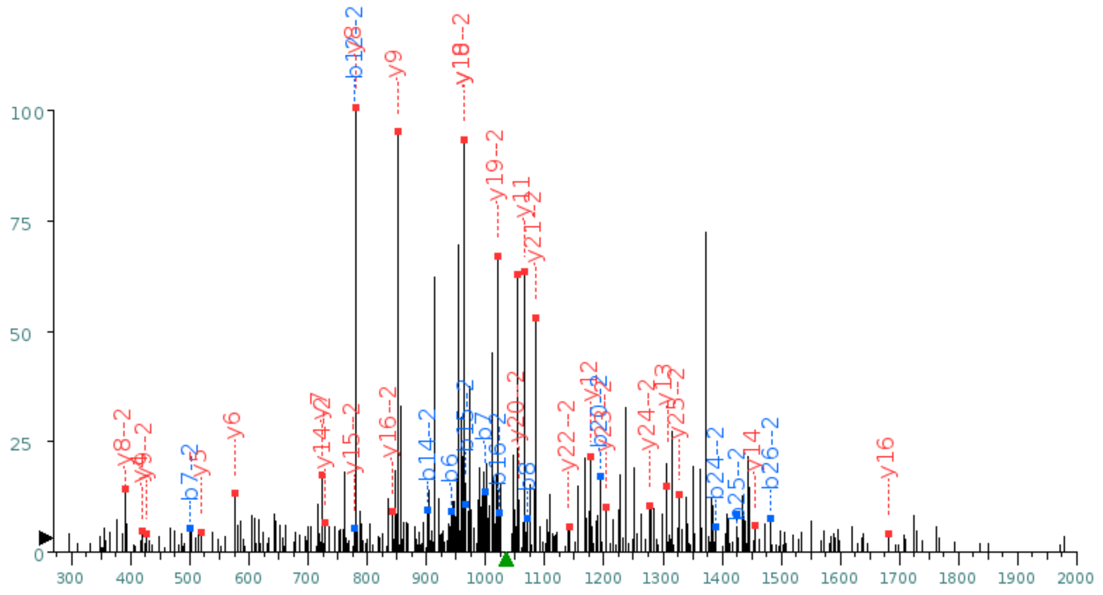

**Predicted Fragmentation Pattern**

| Seq | # | b: Δ Error | b (+1) | y (+1) | y: Δ Error | +1 |
|---|---|---|---|---|---|---|
| S# | 1 | --- | 308.098 | --- | --- | 27 |
| M* | 2 | --- | 455.133 | 2801.388 | --- | 26 |
| V | 3 | --- | 554.201 | 2654.353 | --- | 25 |
| F | 4 | --- | 701.270 | 2555.284 | --- | 24 |
| Q | 5 | --- | 829.328 | 2408.216 | --- | 23 |
| I | 6 | 109.314 | **942.412** | 2280.157 | --- | 22 |
| G | 7 | -86.703 | **999.434** | 2167.073 | --- | 21 |
| A | 8 | -203.233 | **1070.471** | 2110.052 | --- | 20 |
| N | 9 | --- | 1184.514 | 2039.015 | --- | 19 |
| K | 10 | --- | 1312.609 | 1924.972 | --- | 18 |
| D | 11 | --- | 1427.636 | 1796.877 | --- | 17 |
| Q | 12 | --- | 1555.694 | **1681.850** | 150.181 | 16 |
| V | 13 | --- | 1654.763 | 1553.791 | --- | 15 |
| M* | 14 | --- | 1801.798 | **1454.723** | 82.964 | 14 |
| E | 15 | --- | 1930.841 | **1307.688** | -635.661 | 13 |
| L | 16 | --- | 2043.925 | **1178.645** | 83.126 | 12 |
| T | 17 | --- | 2144.973 | **1065.561** | 61.095 | 11 |
| I | 18 | --- | 2258.057 | **964.513** | -809.267 | 10 |
| A | 19 | --- | 2329.094 | **851.429** | 281.297 | 9 |
| G | 20 | --- | 2386.115 | **780.392** | 54.370 | 8 |

| Run: | 77664 | Scans: | 22083 - 22083 | m/z Range: | 306.52396 | to | 1696.7757 |
|---|---|---|---|---|---|---|---|
| Search: | 77664 | Precursor Mass: | 1000.97198 | | | | |
| Mods: | C=160.03 *+15.99 #+220.06 @+291.1 | Charge: | +2 | | | | |

**Sequence**

DQVM*ELT#IAGMGTSALK

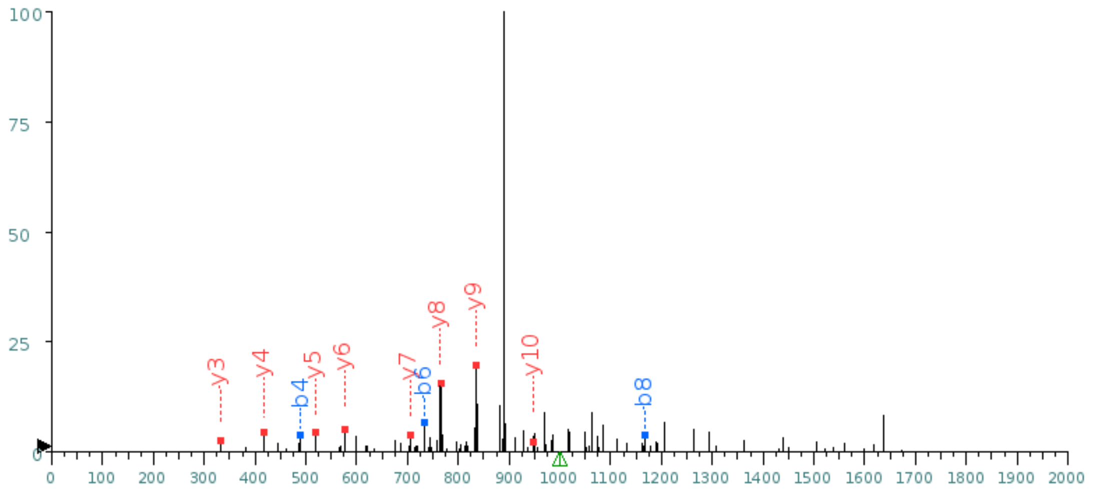

**Predicted Fragmentation Pattern**

| Seq | # | b: Δ Error | b | y | y: Δ Error | +1 |
|---|---|---|---|---|---|---|
| D | 1 | --- | 116.034 | --- | --- | 17 |
| Q | 2 | --- | 244.093 | 1885.913 | --- | 16 |
| V | 3 | --- | 343.161 | 1757.855 | --- | 15 |
| M* | 4 | -1885.198 | **490.197** | 1658.786 | --- | 14 |
| E | 5 | --- | 619.239 | 1511.751 | --- | 13 |
| L | 6 | 171.960 | **732.323** | 1382.708 | --- | 12 |
| T# | 7 | --- | 1053.429 | 1269.624 | --- | 11 |
| I | 8 | 29.401 | **1166.513** | **948.518** | 59.745 | 10 |
| A | 9 | --- | 1237.550 | **835.434** | 132.643 | 9 |
| G | 10 | --- | 1294.572 | **764.397** | 1292.805 | 8 |
| M | 11 | --- | 1425.612 | **707.376** | 45.863 | 7 |
| G | 12 | --- | 1482.634 | **576.335** | 407.330 | 6 |
| T | 13 | --- | 1583.682 | **519.314** | 272.774 | 5 |
| S | 14 | --- | 1670.714 | **418.266** | -66.447 | 4 |
| A | 15 | --- | 1741.751 | **331.234** | 874.670 | 3 |
| L | 16 | --- | 1854.835 | 260.197 | --- | 2 |
| K | 17 | --- | --- | 147.113 | --- | 1 |

**Soleil Index**

| | | |
|---|---|---|
| 381.2327 | y1-8 frag doubly charged | 1% |
| 444.4595 | not explained | 2% |
| 462.3806 | not explained | 1% |

| Run: | 77664 | Scans: | 22504 - 22504 | m/z Range: | 270.43472 to 1952.3678 |
|---|---|---|---|---|---|
| Search: | 77664 | Precursor Mass: | 1036.49137 | | |
| Mods: | C=160.03 *+15.99 #+220.06 @+291.1 | Charge: | +2 | | |

#### Sequence

DQVM*ELTIAGMGTS@ALK

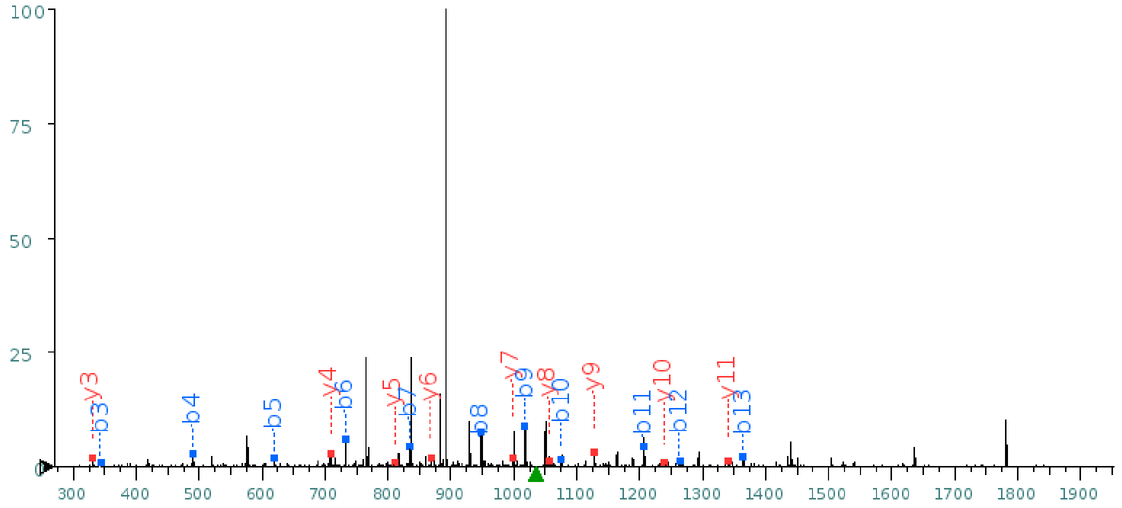

#### Predicted Fragmentation Pattern

| Seq | # | b: Δ Error | b | y | y: Δ Error | +1 |
|---|---|---|---|---|---|---|
| D | 1 | --- | 116.034 | --- | --- | 17 |
| Q | 2 | --- | 244.093 | 1956.950 | --- | 16 |
| V | 3 | 231.385 | 343.161 | 1828.892 | --- | 15 |
| M* | 4 | 644.772 | 490.197 | 1729.823 | --- | 14 |
| E | 5 | 579.117 | 619.239 | 1582.788 | --- | 13 |
| L | 6 | 201.953 | 732.323 | 1453.745 | --- | 12 |
| T | 7 | 214.576 | 833.371 | 1340.661 | 284.275 | 11 |
| I | 8 | 235.915 | 946.455 | 1239.614 | -642.113 | 10 |
| A | 9 | 160.200 | 1017.492 | 1126.530 | 200.450 | 9 |
| G | 10 | 142.633 | 1074.514 | 1055.493 | 204.925 | 8 |
| M | 11 | 164.216 | 1205.554 | 998.471 | 66.760 | 7 |
| G | 12 | 351.246 | 1262.576 | 867.431 | 285.698 | 6 |
| T | 13 | 157.228 | 1363.623 | 810.409 | 456.583 | 5 |
| S@ | 14 | --- | 1741.751 | 709.361 | -1160.481 | 4 |
| A | 15 | --- | 1812.788 | 331.234 | 927.551 | 3 |
| L | 16 | --- | 1925.872 | 260.197 | --- | 2 |
| K | 17 | --- | --- | 147.113 | --- | 1 |

| Run: | 77664 | Scans: | 22759 - 22759 | m/z Range: | 294.55688 to 1965.2176 |
|---|---|---|---|---|---|
| Search: | 77664 | Precursor Mass: | 1031.49769 | | |
| Mods: | C=160.03 *+15.99 #+220.06 @+291.1 | Charge: | +3 | | |

### Sequence

S#M*VFQIGANKDQVMELTIAGM*GTSALK

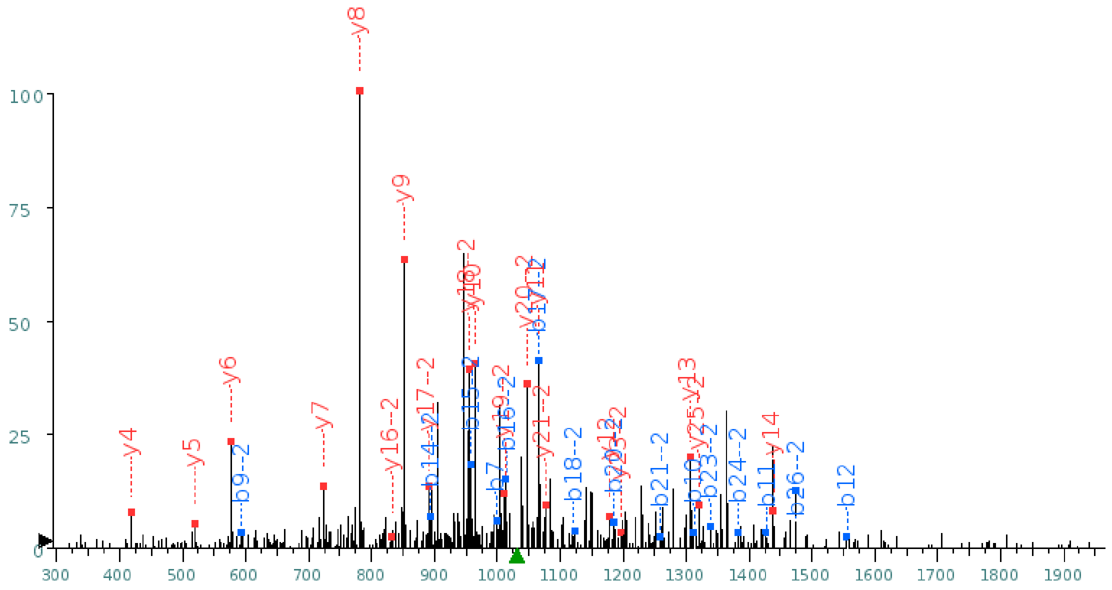

### Predicted Fragmentation Pattern

+1

| Seq | # | b: Δ Error | b | y | y: Δ Error | +1 |
|---|---|---|---|---|---|---|
| S# | 1 | --- | 308.098 | --- | --- | 27 |
| M* | 2 | --- | 455.133 | 2785.393 | --- | 26 |
| V | 3 | --- | 554.201 | 2638.358 | --- | 25 |
| F | 4 | --- | 701.270 | 2539.289 | --- | 24 |
| Q | 5 | --- | 829.328 | 2392.221 | --- | 23 |
| I | 6 | --- | 942.412 | 2264.162 | --- | 22 |
| G | 7 | -680.512 | **999.434** | 2151.078 | --- | 21 |
| A | 8 | --- | 1070.471 | 2094.057 | --- | 20 |
| N | 9 | --- | 1184.514 | 2023.020 | --- | 19 |
| K | 10 | -553.315 | **1312.609** | 1908.977 | --- | 18 |
| D | 11 | -500.127 | **1427.636** | 1780.882 | --- | 17 |
| Q | 12 | -308.786 | **1555.694** | 1665.855 | --- | 16 |
| V | 13 | --- | 1654.763 | 1537.796 | --- | 15 |
| M | 14 | --- | 1785.803 | **1438.728** | -359.273 | 14 |
| E | 15 | --- | 1914.846 | **1307.688** | -204.777 | 13 |
| L | 16 | --- | 2027.930 | **1178.645** | 257.165 | 12 |
| T | 17 | --- | 2128.978 | **1065.561** | 78.162 | 11 |
| I | 18 | --- | 2242.062 | **964.513** | 57.698 | 10 |

| | | | | | |
|---|---|---|---|---|---|
| **Run:** | 77664 | **Scans:** | 24233 - 24233 | **m/z Range:** | 306.23224 to 2000 |
| **Search:** | 77664 | **Precursor Mass:** | 1026.16643 | | |
| **Mods:** | C=160.03 *+15.99 #+220.06 @+291.1 | **Charge:** | +3 | | |

**Sequence**

SMVFQIGANKDQVMELT#IAGM*GTSALK

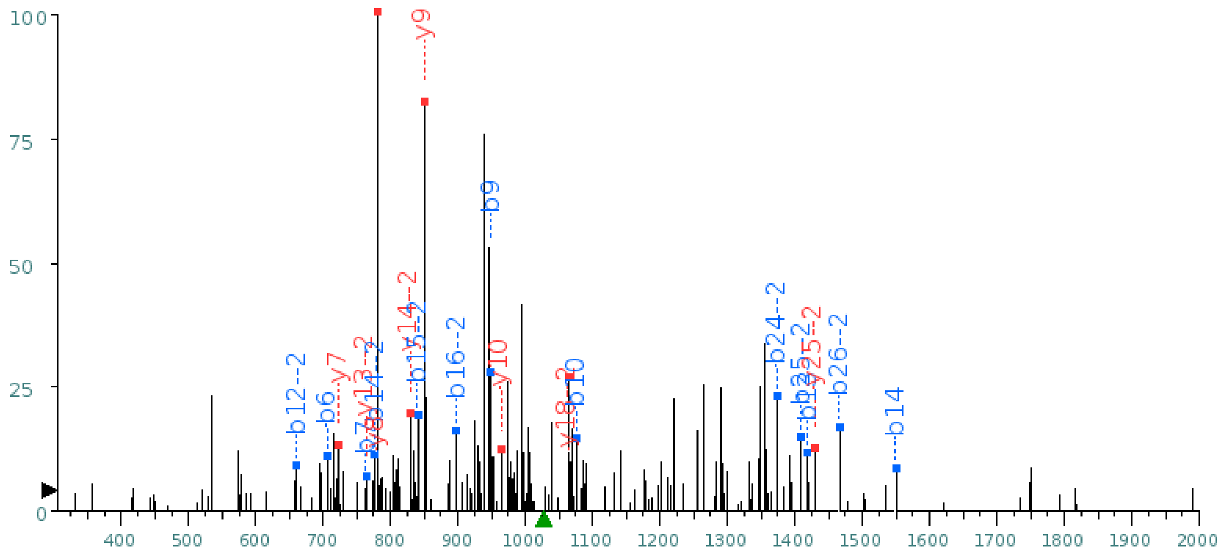

**Predicted Fragmentation Pattern**

**+1**

| Seq | # | b: Δ Error | b | y | y: Δ Error | +1 |
|---|---|---|---|---|---|---|
| **S** | **1** | --- | 88.039 | --- | --- | **27** |
| **M** | **2** | --- | 219.080 | 2989.457 | --- | **26** |
| **V** | **3** | --- | 318.148 | 2858.416 | --- | **25** |
| **F** | **4** | --- | 465.217 | 2759.348 | --- | **24** |
| **Q** | **5** | --- | 593.275 | 2612.279 | --- | **23** |
| **I** | **6** | **-33.017** | **706.359** | 2484.221 | --- | **22** |
| **G** | **7** | **318.614** | **763.381** | 2371.137 | --- | **21** |
| **A** | **8** | --- | 834.418 | 2314.115 | --- | **20** |
| **N** | **9** | **-744.534** | **948.461** | 2243.078 | --- | **19** |
| **K** | **10** | **-38.498** | **1076.556** | 2129.035 | --- | **18** |
| **D** | **11** | --- | 1191.583 | 2000.940 | --- | **17** |
| **Q** | **12** | --- | 1319.641 | 1885.913 | --- | **16** |
| **V** | **13** | **106.808** | **1418.710** | 1757.855 | --- | **15** |
| **M** | **14** | **175.134** | **1549.750** | 1658.786 | --- | **14** |
| **E** | **15** | --- | 1678.793 | 1527.746 | --- | **13** |
| **L** | **16** | --- | 1791.877 | 1398.703 | --- | **12** |
| **T#** | **17** | --- | 2112.983 | 1285.619 | --- | **11** |

| Run: | 77664 | Scans: | 24371 - 24371 | m/z Range: | 288.26691 to 2000 |
|---|---|---|---|---|---|
| Search: | 77664 | Precursor Mass: | 1049.84544 | | |
| Mods: | C=160.03 *+15.99 #+220.06 @+291.1 | Charge: | +3 | | |

### Sequence

S@MVFQIGANKDQVM*ELTIAGMGTSALK

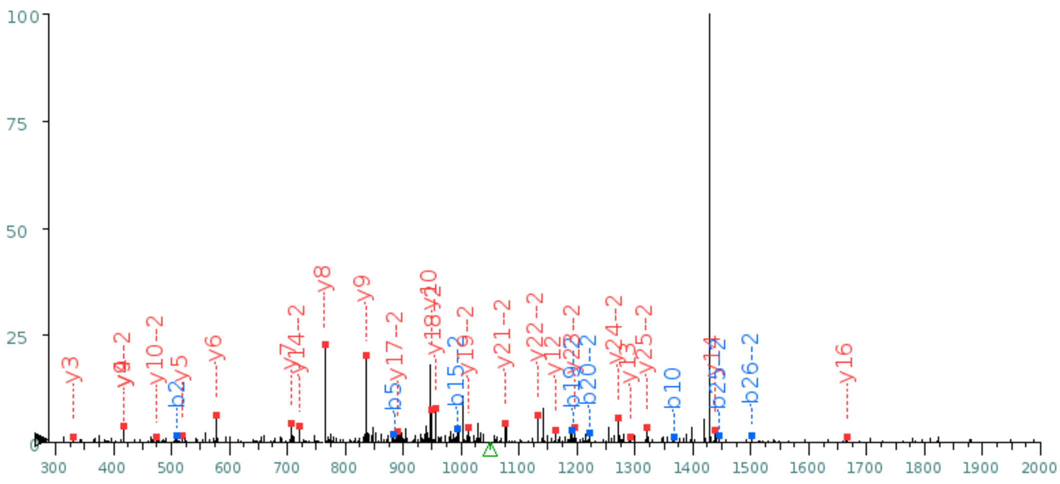

### Predicted Fragmentation Pattern

| Seq | # | b: Δ Error | +1 b | +1 y | y: Δ Error | +1 |
|---|---|---|---|---|---|---|
| S@ | 1 | --- | 379.135 | --- | --- | 27 |
| M | 2 | 30.431 | **510.175** | 2769.398 | --- | 26 |
| V | 3 | --- | 609.244 | 2638.358 | --- | 25 |
| F | 4 | --- | 756.312 | 2539.289 | --- | 24 |
| Q | 5 | -720.082 | **884.371** | 2392.221 | --- | 23 |
| I | 6 | --- | 997.455 | 2264.162 | --- | 22 |
| G | 7 | --- | 1054.476 | 2151.078 | --- | 21 |
| A | 8 | --- | 1125.513 | 2094.057 | --- | 20 |
| N | 9 | --- | 1239.556 | 2023.020 | --- | 19 |
| K | 10 | -291.878 | **1367.651** | 1908.977 | --- | 18 |
| D | 11 | --- | 1482.678 | 1780.882 | --- | 17 |
| Q | 12 | --- | 1610.737 | **1665.855** | 222.920 | 16 |
| V | 13 | --- | 1709.805 | 1537.796 | --- | 15 |
| M* | 14 | --- | 1856.840 | **1438.728** | 130.061 | 14 |
| E | 15 | --- | 1985.883 | **1291.693** | -145.075 | 13 |
| L | 16 | --- | 2098.967 | **1162.650** | 208.790 | 12 |
| T | 17 | --- | 2200.015 | 1049.566 | --- | 11 |
| I | 18 | --- | 2313.099 | **948.518** | 72.226 | 10 |
| A | 19 | --- | 2384.136 | **835.434** | 48.495 | 9 |
| G | 20 | --- | 2441.157 | **764.397** | 81.749 | 8 |
| M | 21 | --- | 2572.198 | **707.376** | 127.646 | 7 |
| G | 22 | --- | 2629.219 | **576.335** | 174.799 | 6 |

| Run: | 77664 | Scans: | 24419 - 24419 | m/z Range: | 318.12466 to 1935.9945 |
|---|---|---|---|---|---|
| Search: | 77664 | Precursor Mass: | 1028.49325 | | |
| Mods: | C=160.03 *+15.99 #+220.06 @+291.1 | Charge: | +2 | | |

### Sequence

DQVMELT@IAGMGTSALK

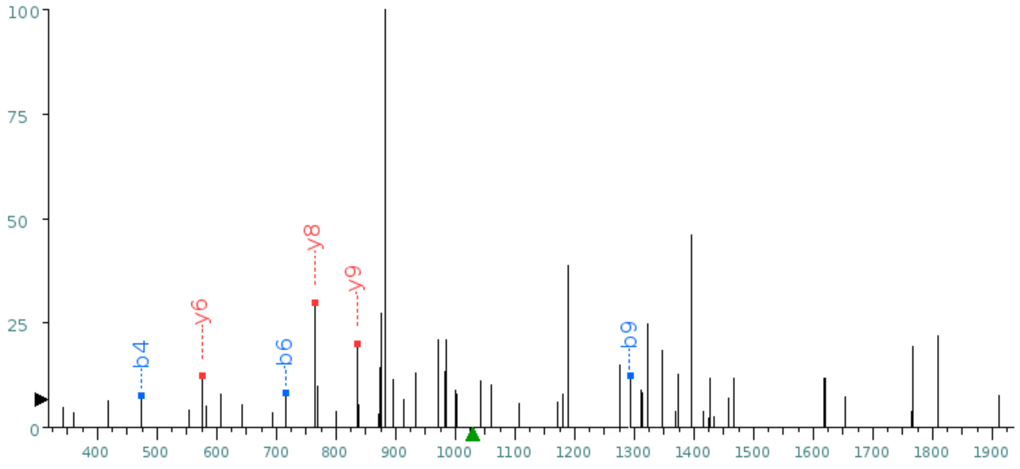

### Predicted Fragmentation Pattern

| Seq | # | b: Δ Error | b | y | y: Δ Error | +1 |
|---|---|---|---|---|---|---|
| D | 1 | --- | 116.034 | --- | --- | 17 |
| Q | 2 | --- | 244.093 | 1940.956 | --- | 16 |
| V | 3 | --- | 343.161 | 1812.897 | --- | 15 |
| M | 4 | 343.532 | **474.202** | 1713.829 | --- | 14 |
| E | 5 | --- | 603.244 | 1582.788 | --- | 13 |
| L | 6 | 402.884 | **716.328** | 1453.745 | --- | 12 |
| T@ | 7 | --- | 1108.471 | 1340.661 | --- | 11 |
| I | 8 | --- | 1221.556 | 948.518 | --- | 10 |
| A | 9 | 215.952 | **1292.593** | **835.434** | 265.847 | 9 |
| G | 10 | --- | 1349.614 | **764.397** | 285.604 | 8 |
| M | 11 | --- | 1480.655 | 707.376 | --- | 7 |
| G | 12 | --- | 1537.676 | **576.335** | 360.875 | 6 |
| T | 13 | --- | 1638.724 | 519.314 | --- | 5 |
| S | 14 | --- | 1725.756 | 418.266 | --- | 4 |
| A | 15 | --- | 1796.793 | 331.234 | --- | 3 |
| L | 16 | --- | 1909.877 | 260.197 | --- | 2 |
| K | 17 | --- | --- | 147.113 | --- | 1 |

| | | | | | | |
|---|---|---|---|---|---|---|
| **Run:** | 77664 | **Scans:** | 24535 - 24535 | **m/z Range:** | 318.34988 | to 1910.4957: |
| **Search:** | 77664 | **Precursor Mass:** | 1028.49333 | | | |
| **Mods:** | C=160.03 *+15.99 #+220.06 @+291.1 | **Charge:** | +2 | | | |

**Sequence**

DQVMELT@IAGMGTSALK

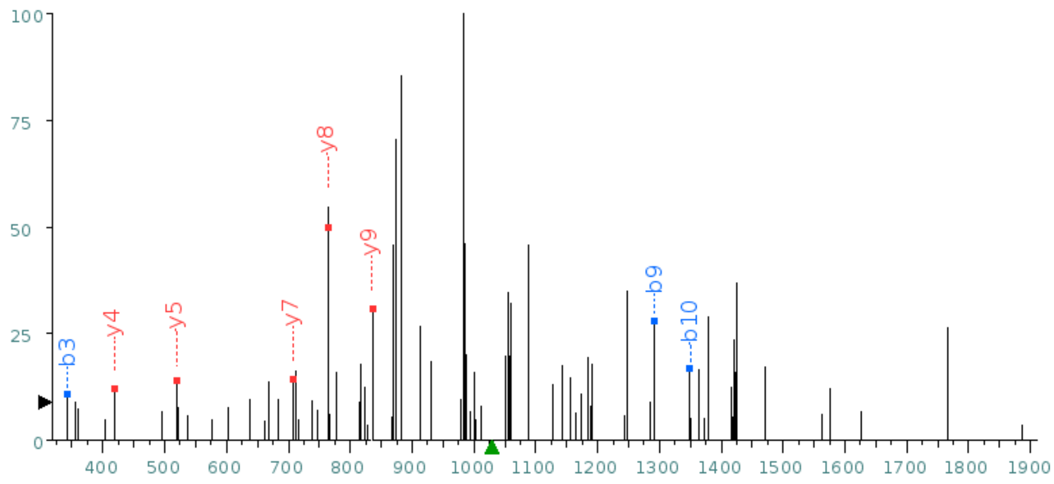

**Predicted Fragmentation Pattern**

| Seq | # | b: Δ Error | b | y | y: Δ Error | +1 |
|---|---|---|---|---|---|---|
| D | 1 | --- | 116.034 | --- | --- | 17 |
| Q | 2 | --- | 244.093 | 1940.956 | --- | 16 |
| V | 3 | 549.508 | **343.161** | 1812.897 | --- | 15 |
| M | 4 | --- | 474.202 | 1713.829 | --- | 14 |
| E | 5 | --- | 603.244 | 1582.788 | --- | 13 |
| L | 6 | --- | 716.328 | 1453.745 | --- | 12 |
| T@ | 7 | --- | 1108.471 | 1340.661 | --- | 11 |
| I | 8 | --- | 1221.556 | 948.518 | --- | 10 |
| A | 9 | -661.866 | **1292.593** | **835.434** | 215.170 | 9 |
| G | 10 | -699.803 | **1349.614** | **764.397** | 333.004 | 8 |
| M | 11 | --- | 1480.655 | **707.376** | 55.525 | 7 |
| G | 12 | --- | 1537.676 | 576.335 | --- | 6 |
| T | 13 | --- | 1638.724 | **519.314** | 330.682 | 5 |
| S | 14 | --- | 1725.756 | **418.266** | 619.820 | 4 |
| A | 15 | --- | 1796.793 | 331.234 | --- | 3 |
| L | 16 | --- | 1909.877 | 260.197 | --- | 2 |
| K | 17 | --- | --- | 147.113 | --- | 1 |

| | | | | | | | |
|---|---|---|---|---|---|---|---|
| **Run:** | 77664 | **Scans:** | 24797 - 24797 | **m/z Range:** | 306.44461 | to | 1953.3535: |
| **Search:** | 77664 | **Precursor Mass:** | 1028.49364 | | | | |
| **Mods:** | C=160.03 *+15.99 #+220.06 @+291.1 | **Charge:** | +2 | | | | |

**Sequence**

DQVMELT@IAGMGTSALK

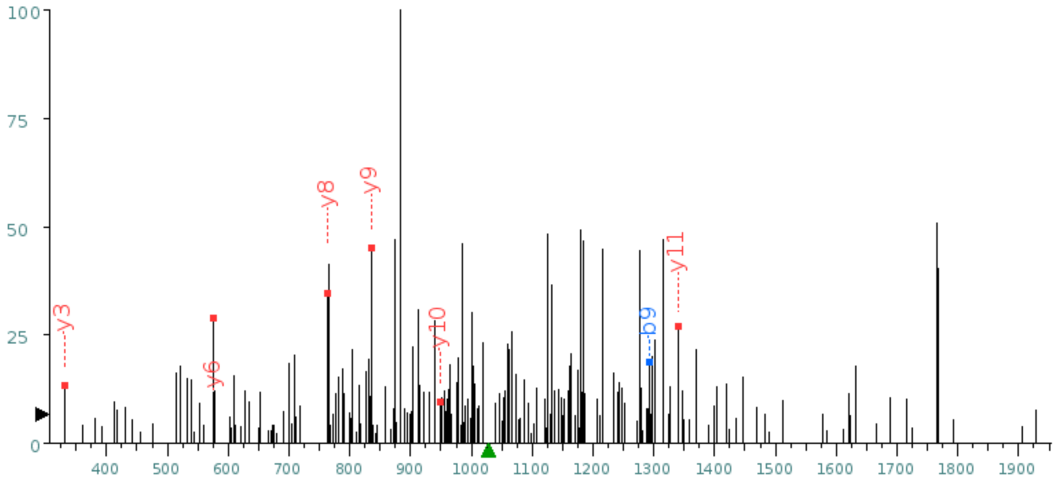

**Predicted Fragmentation Pattern**

| Seq | # | b: Δ Error | b | y | y: Δ Error | +1 |
|---|---|---|---|---|---|---|
| D | 1 | --- | 116.034 | --- | --- | 17 |
| Q | 2 | --- | 244.093 | 1940.956 | --- | 16 |
| V | 3 | --- | 343.161 | 1812.897 | --- | 15 |
| M | 4 | --- | 474.202 | 1713.829 | --- | 14 |
| E | 5 | --- | 603.244 | 1582.788 | --- | 13 |
| L | 6 | --- | 716.328 | 1453.745 | --- | 12 |
| T@ | 7 | --- | 1108.471 | **1340.661** | -558.189 | 11 |
| I | 8 | --- | 1221.556 | **948.518** | 59.230 | 10 |
| A | 9 | 225.297 | **1292.593** | **835.434** | 168.722 | 9 |
| G | 10 | --- | 1349.614 | **764.397** | 66.740 | 8 |
| M | 11 | --- | 1480.655 | 707.376 | --- | 7 |
| G | 12 | --- | 1537.676 | **576.335** | 589.406 | 6 |
| T | 13 | --- | 1638.724 | 519.314 | --- | 5 |
| S | 14 | --- | 1725.756 | 418.266 | --- | 4 |
| A | 15 | --- | 1796.793 | **331.234** | 635.486 | 3 |
| L | 16 | --- | 1909.877 | 260.197 | --- | 2 |
| K | 17 | --- | --- | 147.113 | --- | 1 |

| Run: | 77664 | Scans: | 25297 - 25297 | m/z Range: | 272.27286 | to | 1951.3121 |
|---|---|---|---|---|---|---|---|
| Search: | 77664 | Precursor Mass: | 1049.84654 | | | | |
| Mods: | C=160.03 *+15.99 #+220.06 @+291.1 | Charge: | +3 | | | | |

### Sequence

SMVFQIGANKDQVM*ELT@IAGMGTSALK

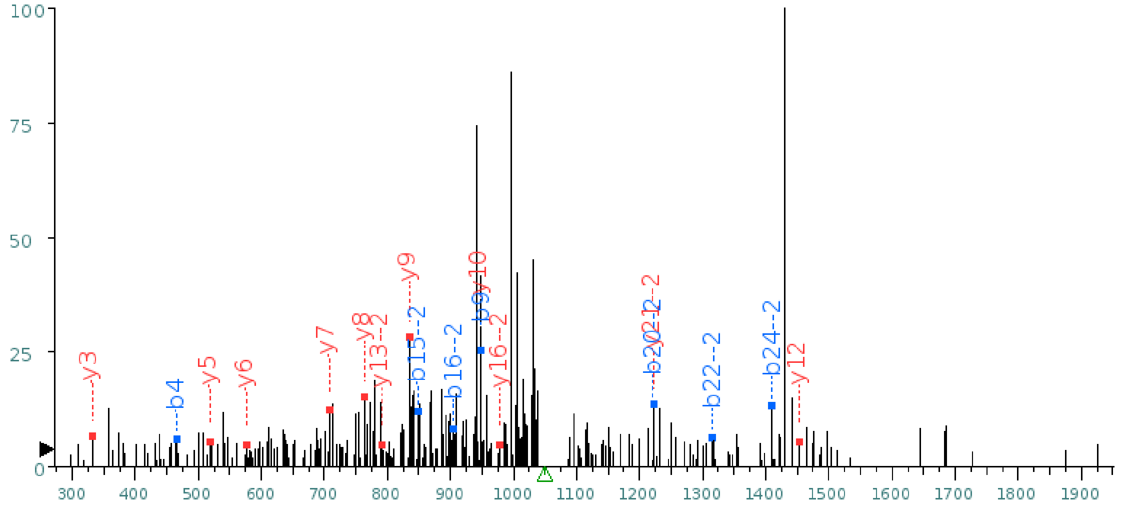

### Predicted Fragmentation Pattern

| | | | +1 | | | |
|---|---|---|---|---|---|---|
| Seq | # | b: Δ Error | b | y | y: Δ Error | +1 |
| S | 1 | --- | 88.039 | --- | --- | 27 |
| M | 2 | --- | 219.080 | 3060.494 | --- | 26 |
| V | 3 | --- | 318.148 | 2929.453 | --- | 25 |
| F | 4 | 422.008 | **465.217** | 2830.385 | --- | 24 |
| Q | 5 | --- | 593.275 | 2683.316 | --- | 23 |
| I | 6 | --- | 706.359 | 2555.258 | --- | 22 |
| G | 7 | --- | 763.381 | 2442.174 | --- | 21 |
| A | 8 | --- | 834.418 | 2385.152 | --- | 20 |
| N | 9 | 445.814 | **948.461** | 2314.115 | --- | 19 |
| K | 10 | --- | 1076.556 | 2200.072 | --- | 18 |
| D | 11 | --- | 1191.583 | 2071.977 | --- | 17 |
| Q | 12 | --- | 1319.641 | 1956.950 | --- | 16 |
| V | 13 | --- | 1418.710 | 1828.892 | --- | 15 |
| M* | 14 | --- | 1565.745 | 1729.823 | --- | 14 |
| E | 15 | --- | 1694.788 | 1582.788 | --- | 13 |
| L | 16 | --- | 1807.872 | **1453.745** | 188.085 | 12 |
| T@ | 17 | --- | 2200.015 | 1340.661 | --- | 11 |
| I | 18 | --- | 2313.099 | **948.518** | 385.200 | 10 |

| | | | | | | |
|---|---|---|---|---|---|---|
| **Run:** | 77664 | **Scans:** | 26816 - 26816 | **m/z Range:** | 360.099 | to 2000 |
| **Search:** | 77664 | **Precursor Mass:** | 1141.54522 | | | |
| **Mods:** | C=160.03 *+15.99 #+220.06 @+291.1 | **Charge:** | +3 | | | |

**Sequence**

S@MVFQIGANKDQVMELT@IAGMGTSALK

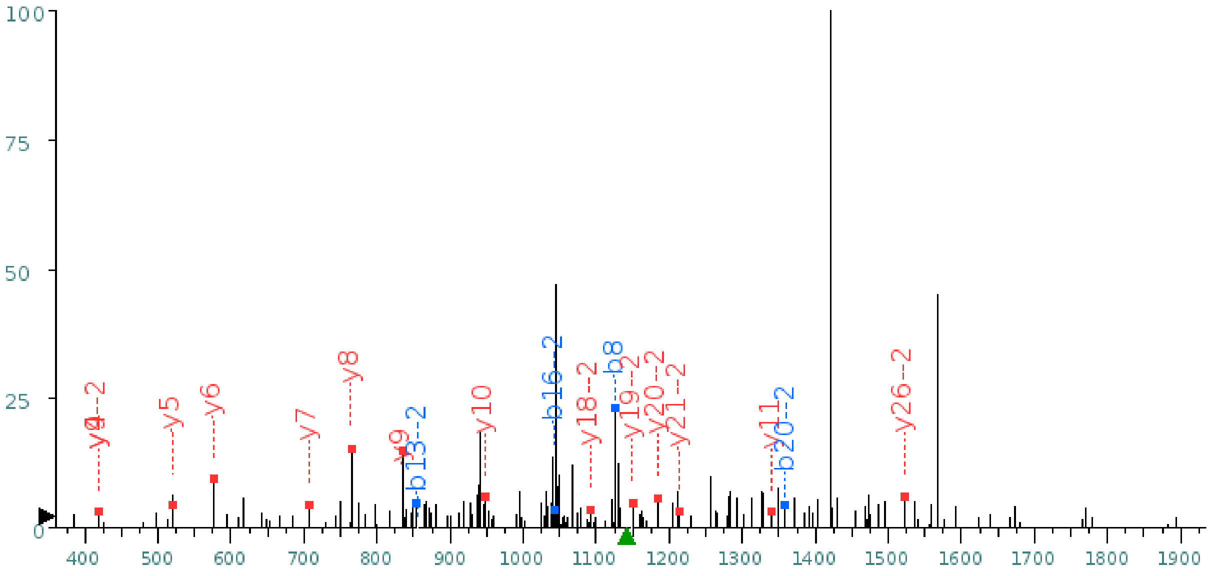

**Predicted Fragmentation Pattern**

+1

| Seq | # | b: Δ Error | b | y | y: Δ Error | +1 |
|---|---|---|---|---|---|---|
| S@ | 1 | --- | 379.135 | --- | --- | 27 |
| M | 2 | --- | 510.175 | 3044.499 | --- | 26 |
| V | 3 | --- | 609.244 | 2913.458 | --- | 25 |
| F | 4 | --- | 756.312 | 2814.390 | --- | 24 |
| Q | 5 | --- | 884.371 | 2667.322 | --- | 23 |
| I | 6 | --- | 997.455 | 2539.263 | --- | 22 |
| G | 7 | --- | 1054.476 | 2426.179 | --- | 21 |
| A | 8 | -526.903 | **1125.513** | 2369.157 | --- | 20 |
| N | 9 | --- | 1239.556 | 2298.120 | --- | 19 |
| K | 10 | --- | 1367.651 | 2184.077 | --- | 18 |
| D | 11 | --- | 1482.678 | 2055.982 | --- | 17 |
| Q | 12 | --- | 1610.737 | 1940.956 | --- | 16 |
| V | 13 | --- | 1709.805 | 1812.897 | --- | 15 |
| M | 14 | --- | 1840.846 | 1713.829 | --- | 14 |
| E | 15 | --- | 1969.888 | 1582.788 | --- | 13 |
| L | 16 | --- | 2082.972 | 1453.745 | --- | 12 |
| T@ | 17 | --- | 2475.115 | **1340.661** | 285.276 | 11 |
| I | 18 | --- | 2588.199 | **948.518** | 83.936 | 10 |

| Run: | 77664 | Scans: | 26833 - 26833 | m/z Range: | 251.16833 to 2000 |
|---|---|---|---|---|---|
| Search: | 77664 | Precursor Mass: | 861.75266 | | |
| Mods: | C=160.03 *+15.99 #+220.06 @+291.1 | Charge: | +3 | | |

**Sequence**

GAKSISGMIETIDNAINT#VSGER

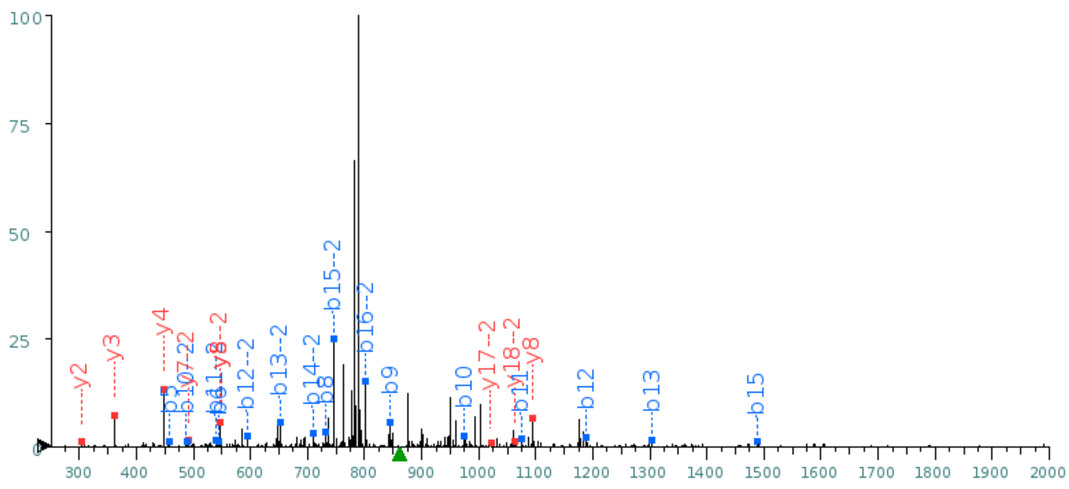

**Predicted Fragmentation Pattern**

| Seq | # | b: Δ Error | b +1 | y +1 | y: Δ Error | +1 |
|---|---|---|---|---|---|---|
| G | 1 | --- | 58.029 | --- | --- | 23 |
| A | 2 | --- | 129.066 | 2526.224 | --- | 22 |
| K | 3 | --- | 257.161 | 2455.187 | --- | 21 |
| S | 4 | --- | 344.193 | 2327.092 | --- | 20 |
| I | 5 | 80.896 | **457.277** | 2240.060 | --- | 19 |
| S | 6 | 170.679 | **544.309** | 2126.976 | --- | 18 |
| G | 7 | --- | 601.330 | 2039.944 | --- | 17 |
| M | 8 | 249.573 | **732.371** | 1982.922 | --- | 16 |
| I | 9 | -73.417 | **845.455** | 1851.882 | --- | 15 |
| E | 10 | 208.727 | **974.498** | 1738.798 | --- | 14 |
| T | 11 | 141.799 | **1075.545** | 1609.755 | --- | 13 |
| I | 12 | 193.949 | **1188.629** | 1508.707 | --- | 12 |
| D | 13 | 164.697 | **1303.656** | 1395.623 | --- | 11 |
| N | 14 | --- | 1417.699 | 1280.596 | --- | 10 |
| A | 15 | 46.693 | **1488.736** | 1166.554 | --- | 9 |
| I | 16 | --- | 1601.820 | **1095.516** | -143.932 | 8 |
| N | 17 | --- | 1715.863 | 982.432 | --- | 7 |
| T# | 18 | --- | 2036.969 | 868.389 | --- | 6 |
| V | 19 | --- | 2136.038 | **547.283** | 252.195 | 5 |
| S | 20 | --- | 2223.070 | **448.215** | 115.165 | 4 |
| G | 21 | --- | 2280.091 | **361.183** | 289.319 | 3 |
| E | 22 | --- | 2409.134 | **304.162** | 370.642 | 2 |

| Run: | 77664 | Scans: | 27133 - 27133 | m/z Range: | to |
|---|---|---|---|---|---|
| Search: | 77664 | Precursor Mass: | 885.43233 | | |
| Mods: | C=160.03 *+15.99 #+220.06 @+291.1 | Charge: | +3 | | |

**Sequence**

GAKSISGMIETIDNAINT@VSGER

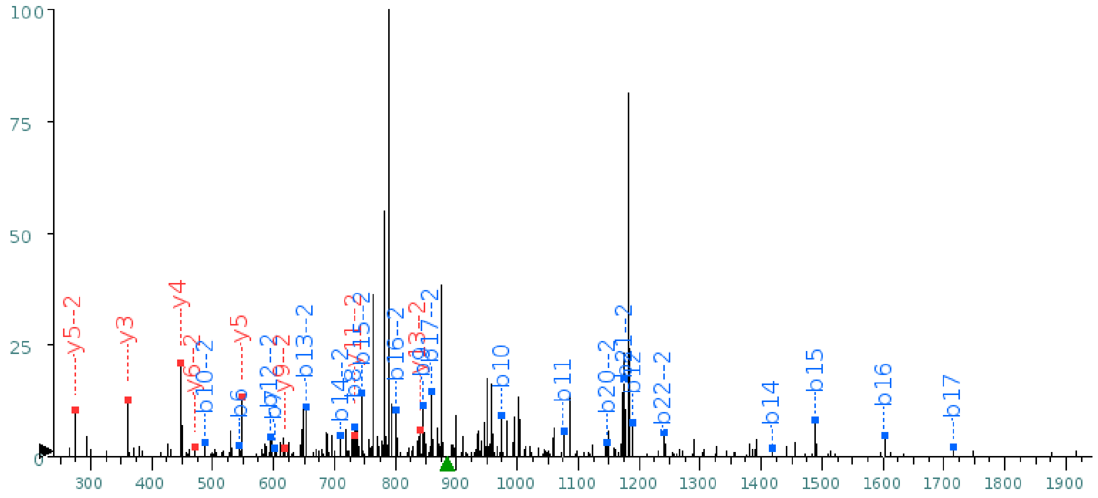

**Predicted Fragmentation Pattern**

+1

| Seq | # | b: Δ Error | b | y | y: Δ Error | +1 |
|---|---|---|---|---|---|---|
| G | 1 | --- | 58.029 | --- | --- | 23 |
| A | 2 | --- | 129.066 | 2597.261 | --- | 22 |
| K | 3 | --- | 257.161 | 2526.224 | --- | 21 |
| S | 4 | --- | 344.193 | 2398.129 | --- | 20 |
| I | 5 | --- | 457.277 | 2311.097 | --- | 19 |
| S | 6 | 390.001 | **544.309** | 2198.013 | --- | 18 |
| G | 7 | 359.355 | **601.330** | 2110.981 | --- | 17 |
| M | 8 | -18.134 | **732.371** | 2053.959 | --- | 16 |
| I | 9 | -83.959 | **845.455** | 1922.919 | --- | 15 |
| E | 10 | 200.463 | **974.498** | 1809.835 | --- | 14 |
| T | 11 | 164.378 | **1075.545** | 1680.792 | --- | 13 |
| I | 12 | 110.173 | **1188.629** | 1579.745 | --- | 12 |
| D | 13 | --- | 1303.656 | 1466.661 | --- | 11 |
| N | 14 | 257.086 | **1417.699** | 1351.634 | --- | 10 |
| A | 15 | 180.562 | **1488.736** | 1237.591 | --- | 9 |
| I | 16 | 144.150 | **1601.820** | 1166.554 | --- | 8 |
| N | 17 | 178.974 | **1715.863** | 1053.469 | --- | 7 |
| T@ | 18 | --- | 2108.006 | 939.427 | --- | 6 |
| V | 19 | --- | 2207.075 | **547.283** | 104.590 | 5 |
| S | 20 | --- | 2294.107 | **448.215** | -57.765 | 4 |
| G | 21 | --- | 2351.128 | **361.183** | 442.984 | 3 |

**MS/MS data of trypsin digested peptides with site score A and site score B above 19 from *Clostridium botulinum Cb*Fla flagellin protein co-expressed with *Clostridium botulinum* Maf in BL21(DE3) cells. Results of dataset 2 (MS/MS results obtained in April 2019).**

| | | | | | |
|---|---|---|---|---|---|
| **Run:** | 77662 | **Scans:** | 9318 - 9318 | **m/z Range:** | 193.2561 to 1136.6317 |
| **Search:** | 77662 | **Precursor Mass:** | 679.3286 | | |
| **Mods:** | C=160.03 *+15.99 #+220.06 @+291.1 | **Charge:** | +2 | | |

**Sequence**

IAS#QTQFNTK

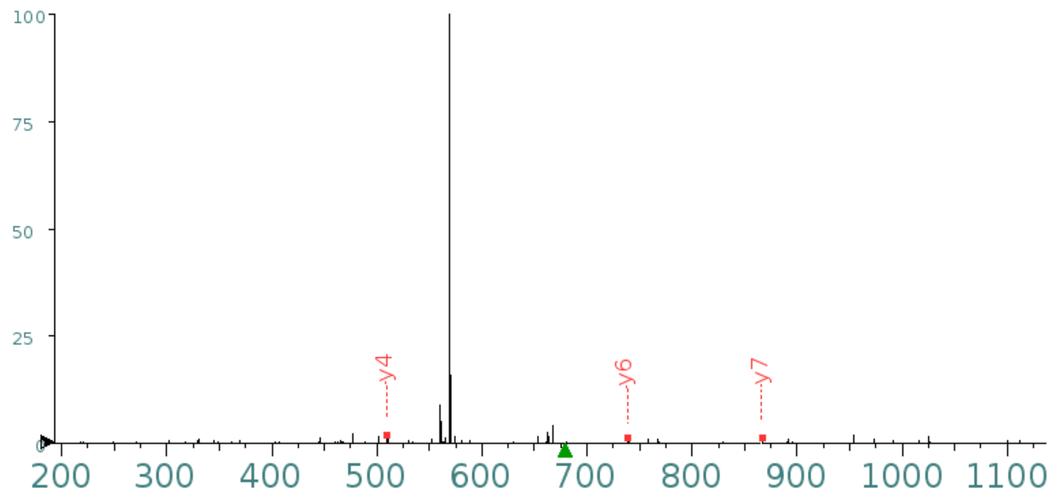

**Predicted Fragmentation Pattern**

| Seq | # | b: Δ Error | b | y | y: Δ Error | +1 |
|---|---|---|---|---|---|---|
| I | 1 | --- | 114.091 | --- | --- | 10 |
| A | 2 | --- | 185.128 | 1244.564 | --- | 9 |
| S# | 3 | --- | 492.219 | 1173.527 | --- | 8 |
| Q | 4 | --- | 620.277 | 866.437 | 113.881 | 7 |
| T | 5 | --- | 721.325 | 738.378 | 54.763 | 6 |
| Q | 6 | --- | 849.384 | 637.330 | --- | 5 |
| F | 7 | --- | 996.452 | 509.272 | 320.662 | 4 |
| N | 8 | --- | 1110.495 | 362.203 | --- | 3 |
| T | 9 | --- | 1211.543 | 248.160 | --- | 2 |
| K | 10 | --- | --- | 147.113 | --- | 1 |

**Soleil Index**

| | | |
|---|---|---|
| 569.4458 | not explained | 100% |
| 570.2626 | not explained | 16% |

| | | | | | |
|---|---|---|---|---|---|
| **Run:** | 77662 | **Scans:** | 9639 - 9639 | **m/z Range:** | 223.2569 to 1162.8313 |
| **Search:** | 77662 | **Precursor Mass:** | 679.32712 | | |
| **Mods:** | C=160.03 *+15.99 #+220.06 @+291.1 | **Charge:** | +2 | | |

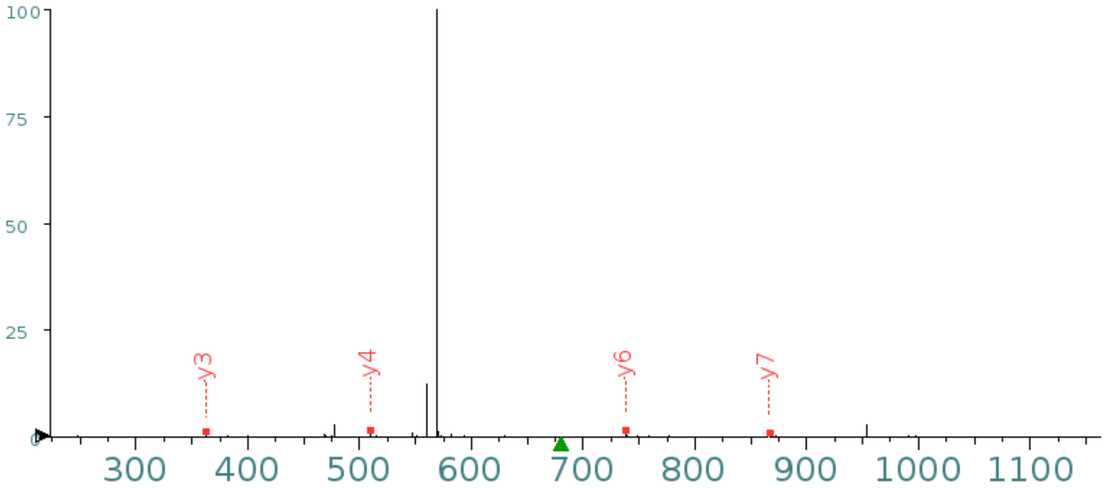

**Sequence**

IAS#QTQFNTK

**Predicted Fragmentation Pattern**

| Seq | # | b: Δ Error | b | y | y: Δ Error | +1 |
|---|---|---|---|---|---|---|
| I | 1 | --- | 114.091 | --- | --- | 10 |
| A | 2 | --- | 185.128 | 1244.564 | --- | 9 |
| S# | 3 | --- | 492.219 | 1173.527 | --- | 8 |
| Q | 4 | --- | 620.277 | **866.437** | 285.977 | 7 |
| T | 5 | --- | 721.325 | **738.378** | 239.704 | 6 |
| Q | 6 | --- | 849.384 | 637.330 | --- | 5 |
| F | 7 | --- | 996.452 | **509.272** | 306.409 | 4 |
| N | 8 | --- | 1110.495 | **362.203** | 149.501 | 3 |
| T | 9 | --- | 1211.543 | 248.160 | --- | 2 |
| K | 10 | --- | --- | 147.113 | --- | 1 |

**Soleil Index**

| | | |
|---|---|---|
| **569.4003** | **not explained** | **100%** |

| | | | | | |
|---|---|---|---|---|---|
| **Run:** | 77662 | **Scans:** | 9803 - 9803 | **m/z Range:** | 180.29677 to 1249.6579 |
| **Search:** | 77662 | **Precursor Mass:** | 686.3214 | | |
| **Mods:** | C=160.03 *+15.99 #+220.06 @+291.1 | **Charge:** | +2 | | |

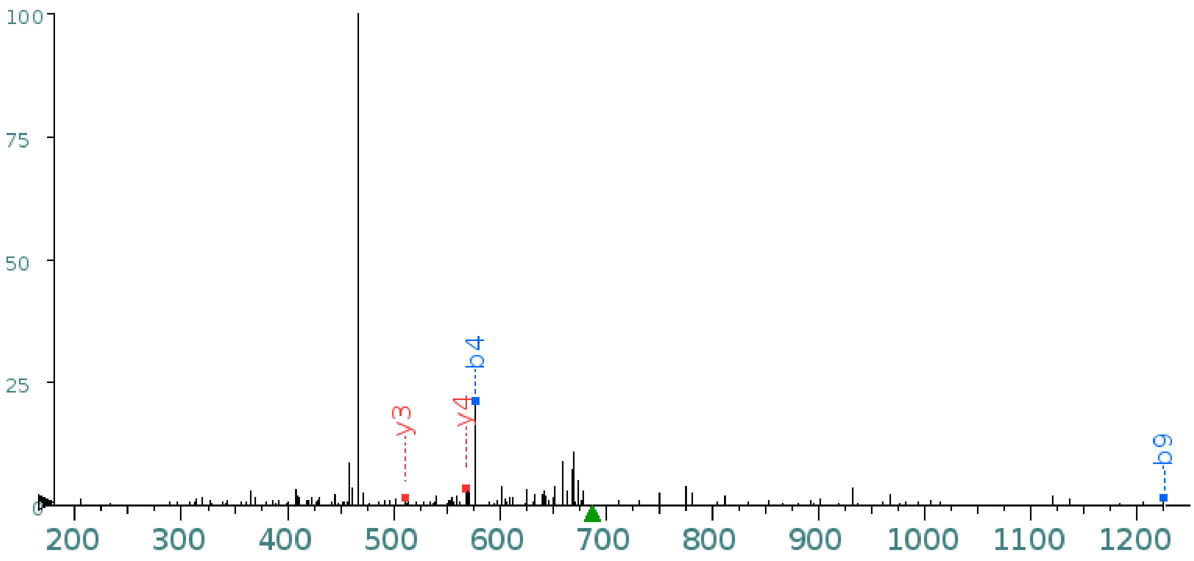

**Sequence**

VGS#INIGS#GK

**Predicted Fragmentation Pattern**

| Seq | # | b: Δ Error | b | y | y: Δ Error | +1 |
|---|---|---|---|---|---|---|
| V | 1 | --- | 100.076 | --- | --- | 10 |
| G | 2 | --- | 157.097 | 1272.569 | --- | 9 |
| S# | 3 | --- | 464.187 | 1215.547 | --- | 8 |
| I | 4 | -1721.183 | **577.272** | 908.457 | --- | 7 |
| N | 5 | --- | 691.314 | 795.373 | --- | 6 |
| I | 6 | --- | 804.399 | 681.330 | --- | 5 |
| G | 7 | --- | 861.420 | **568.246** | -360.538 | 4 |
| S# | 8 | --- | 1168.510 | **511.225** | 31.901 | 3 |
| G | 9 | -713.533 | **1225.532** | 204.134 | --- | 2 |
| K | 10 | --- | --- | 147.113 | --- | 1 |

**Soleil Index**

| | | |
|---|---|---|
| 466.3061 | not explained | 100% |

| | | | | | |
|---|---|---|---|---|---|
| **Run:** | 77662 | **Scans:** | 11353 - 11353 | **m/z Range:** | 191.23517 to 1234.7959 |
| **Search:** | 77662 | **Precursor Mass:** | 679.32812 | | |
| **Mods:** | C=160.03 *+15.99 #+220.06 @+291.1 | **Charge:** | +2 | | |

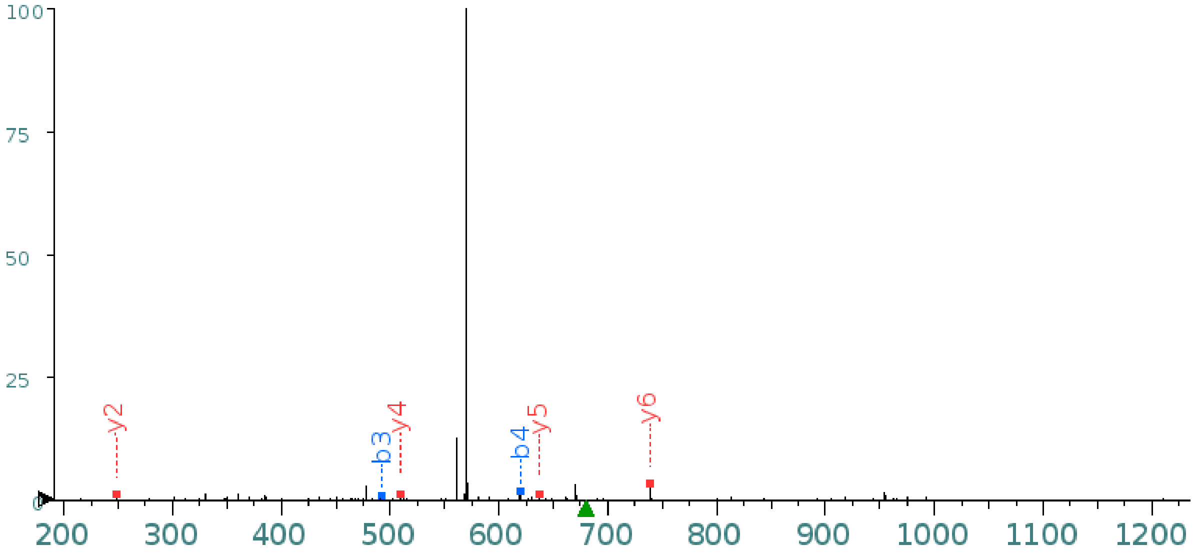

**Sequence**

IAS#QTQFNTK

**Predicted Fragmentation Pattern**

| Seq | # | b: Δ Error | b | y | y: Δ Error | +1 |
|---|---|---|---|---|---|---|
| I | 1 | --- | 114.091 | --- | --- | 10 |
| A | 2 | --- | 185.128 | 1244.564 | --- | 9 |
| S# | 3 | 345.527 | **492.219** | 1173.527 | --- | 8 |
| Q | 4 | -680.924 | **620.277** | 866.437 | --- | 7 |
| T | 5 | --- | 721.325 | **738.378** | 338.674 | 6 |
| Q | 6 | --- | 849.384 | **637.330** | 382.222 | 5 |
| F | 7 | --- | 996.452 | **509.272** | 457.900 | 4 |
| N | 8 | --- | 1110.495 | 362.203 | --- | 3 |
| T | 9 | --- | 1211.543 | **248.160** | 618.710 | 2 |
| K | 10 | --- | --- | 147.113 | --- | 1 |

**Soleil Index**

| | | |
|---|---|---|
| 569.4179 | not explained | 100% |

| Run: | 77662 | Scans: | 11443 - 11443 | m/z Range: | 203.18138 to 1020.8418 |
|---|---|---|---|---|---|
| Search: | 77662 | Precursor Mass: | 576.29298 | | |
| Mods: | C=160.03 *+15.99 #+220.06 @+291.1 | Charge: | +2 | | |

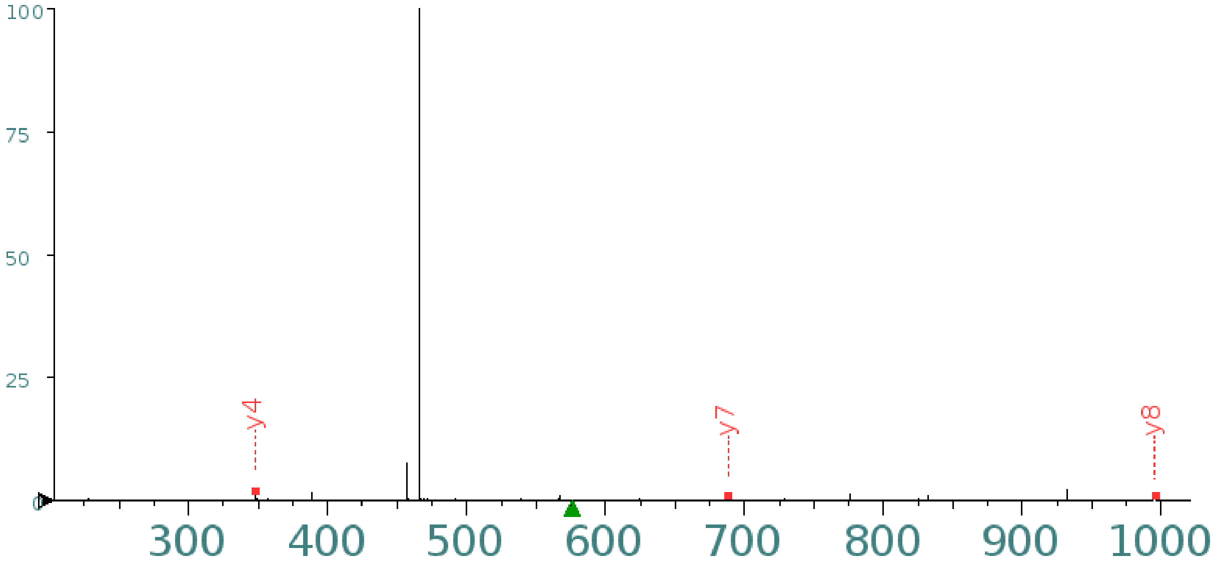

**Sequence**

VGS#INIGSGK

**Predicted Fragmentation Pattern**

| Seq | # | b: Δ Error | b | y | y: Δ Error | +1 |
|---|---|---|---|---|---|---|
| V | 1 | --- | 100.076 | --- | --- | 10 |
| G | 2 | --- | 157.097 | 1052.511 | --- | 9 |
| S# | 3 | --- | 464.187 | **995.489** | **354.125** | 8 |
| I | 4 | --- | 577.272 | **688.399** | **296.648** | 7 |
| N | 5 | --- | 691.314 | 575.315 | --- | 6 |
| I | 6 | --- | 804.399 | 461.272 | --- | 5 |
| G | 7 | --- | 861.420 | **348.188** | **384.050** | 4 |
| S | 8 | --- | 948.452 | 291.166 | --- | 3 |
| G | 9 | --- | 1005.473 | 204.134 | --- | 2 |
| K | 10 | --- | --- | 147.113 | --- | 1 |

**Soleil Index**

| | | |
|---|---|---|
| 466.3631 | not explained | 100% |

| | | | |
|---|---|---|---|
| **Run:** | 77662 | **Scans:** | 11560 - 11560 |
| **Search:** | 77662 | **Precursor Mass:** | 576.29311 |
| **Mods:** | C=160.03 *+15.99 #+220.06 @+291.1 | **Charge:** | +2 |

**m/z Range:** 160.26895 **to** 957.87189

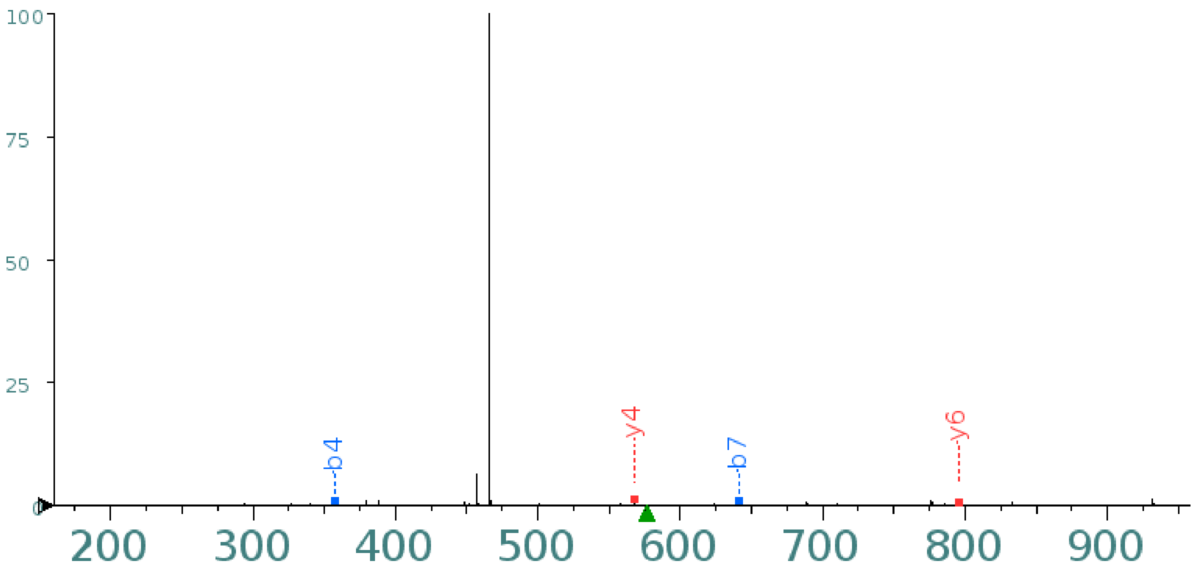

**Sequence**

VGSINIGS#GK

**Predicted Fragmentation Pattern**

| Seq | # | b: Δ Error | b | y | y: Δ Error | +1 |
|---|---|---|---|---|---|---|
| V | 1 | --- | 100.076 | --- | --- | 10 |
| G | 2 | --- | 157.097 | 1052.511 | --- | 9 |
| S | 3 | --- | 244.129 | 995.489 | --- | 8 |
| I | 4 | 385.865 | 357.213 | 908.457 | --- | 7 |
| N | 5 | --- | 471.256 | 795.373 | 236.674 | 6 |
| I | 6 | --- | 584.340 | 681.330 | --- | 5 |
| G | 7 | 231.757 | 641.362 | 568.246 | -1372.693 | 4 |
| S# | 8 | --- | 948.452 | 511.225 | --- | 3 |
| G | 9 | --- | 1005.473 | 204.134 | --- | 2 |
| K | 10 | --- | --- | 147.113 | --- | 1 |

**Soleil Index**

466.4103 **not explained** **100%**

| | | | | | | |
|---|---|---|---|---|---|---|
| **Run:** | 77662 | **Scans:** | 11775 - 11775 | **m/z Range:** | 196.94934 | to 956.69086 |
| **Search:** | 77662 | **Precursor Mass:** | 576.29323 | | | |
| **Mods:** | C=160.03 *+15.99 #+220.06 @+291.1 | **Charge:** | +2 | | | |

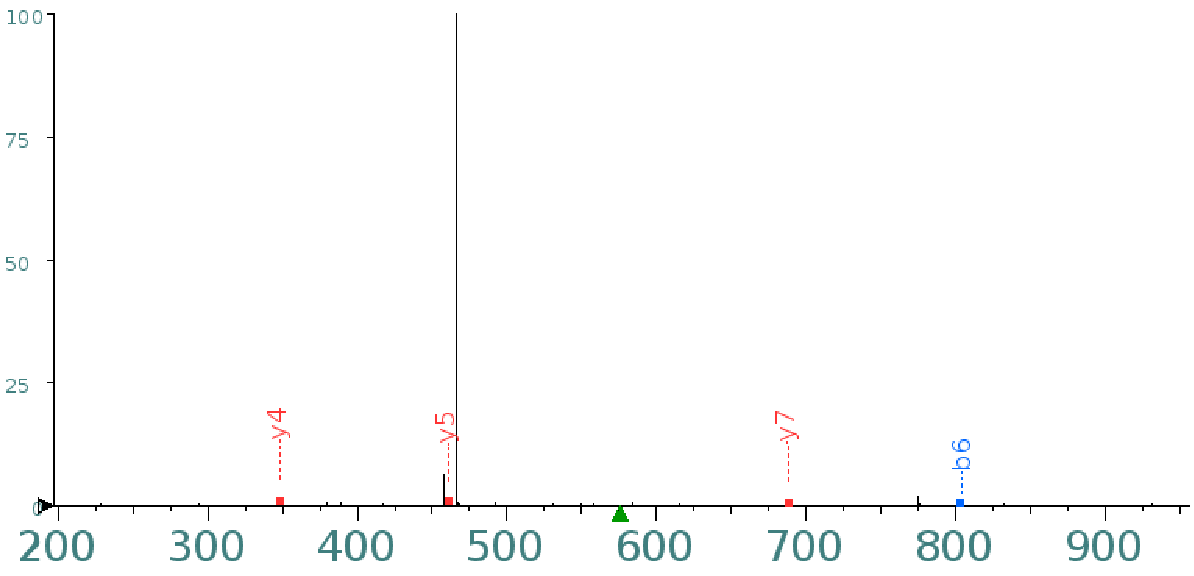

**Sequence**

VGS#INIGSGK

**Predicted Fragmentation Pattern**

| Seq | # | b: Δ Error | b | y | y: Δ Error | +1 |
|---|---|---|---|---|---|---|
| V | 1 | --- | 100.076 | --- | --- | 10 |
| G | 2 | --- | 157.097 | 1052.511 | --- | 9 |
| S# | 3 | --- | 464.187 | 995.489 | --- | 8 |
| I | 4 | --- | 577.272 | **688.399** | 477.734 | 7 |
| N | 5 | --- | 691.314 | 575.315 | --- | 6 |
| I | 6 | -1022.850 | **804.399** | **461.272** | 316.200 | 5 |
| G | 7 | --- | 861.420 | **348.188** | 454.984 | 4 |
| S | 8 | --- | 948.452 | 291.166 | --- | 3 |
| G | 9 | --- | 1005.473 | 204.134 | --- | 2 |
| K | 10 | --- | --- | 147.113 | --- | 1 |

**Soleil Index**

| | | |
|---|---|---|
| 466.3775 | not explained | 100% |

| Run: | 77662 | Scans: | 11892 - 11892 | m/z Range: | 178.1163 | to | 957.3631 |
|---|---|---|---|---|---|---|---|
| Search: | 77662 | Precursor Mass: | 576.29313 | | | | |
| Mods: | C=160.03 *+15.99 #+220.06 @+291.1 | Charge: | +2 | | | | |

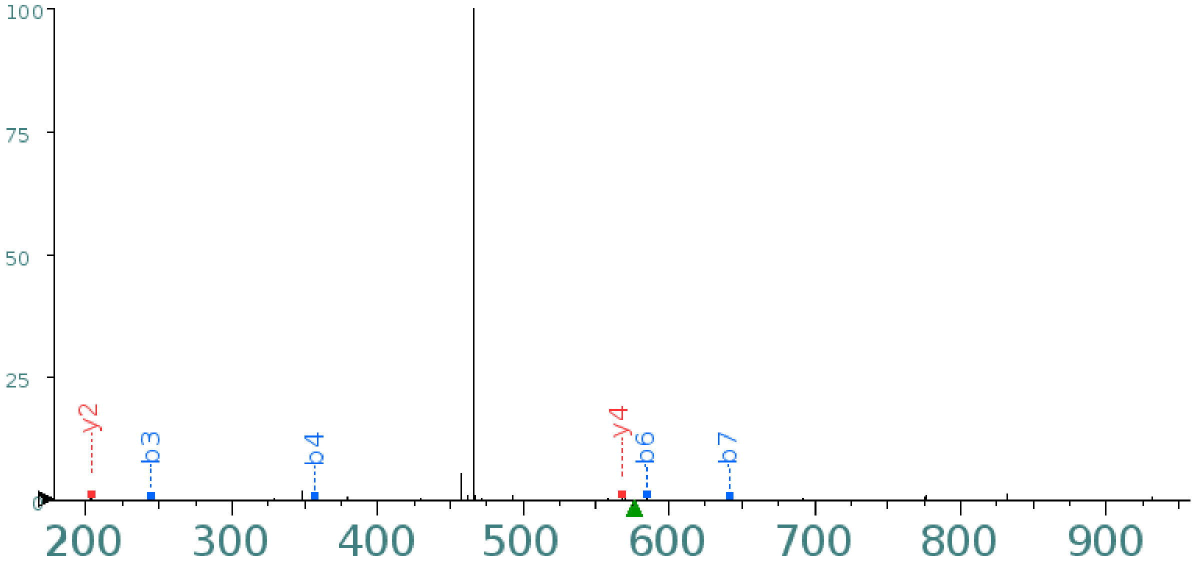

**Sequence**

VGSINIGS#GK

**Predicted Fragmentation Pattern**

| Seq | # | b: Δ Error | b | y | y: Δ Error | +1 |
|---|---|---|---|---|---|---|
| V | 1 | --- | 100.076 | --- | --- | 10 |
| G | 2 | --- | 157.097 | 1052.511 | --- | 9 |
| S | 3 | 277.306 | **244.129** | 995.489 | --- | 8 |
| I | 4 | 398.499 | **357.213** | 908.457 | --- | 7 |
| N | 5 | --- | 471.256 | 795.373 | --- | 6 |
| I | 6 | 342.958 | **584.340** | 681.330 | --- | 5 |
| G | 7 | 436.130 | **641.362** | **568.246** | -1527.597 | 4 |
| S# | 8 | --- | 948.452 | 511.225 | --- | 3 |
| G | 9 | --- | 1005.473 | **204.134** | 12.453 | 2 |
| K | 10 | --- | --- | 147.113 | --- | 1 |

**Soleil Index**

| | | |
|---|---|---|
| 466.3326 | not explained | 100% |

| | | | | | |
|---|---|---|---|---|---|
| **Run:** | 77662 | **Scans:** | 12655 - 12655 | **m/z Range:** | 179.0148 to 956.91644 |
| **Search:** | 77662 | **Precursor Mass:** | 576.29344 | | |
| **Mods:** | C=160.03 *+15.99 #+220.06 @+291.1 | **Charge:** | +2 | | |

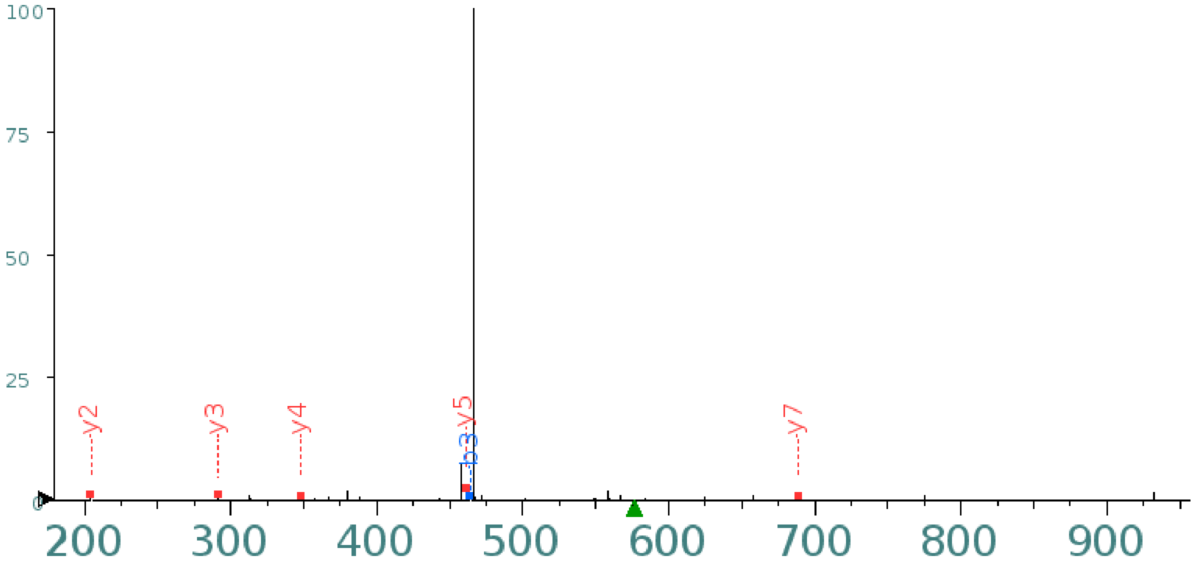

**Sequence**

VGS#INIGSGK

**Predicted Fragmentation Pattern**

| Seq | # | b: Δ Error | b | y | y: Δ Error | +1 |
|---|---|---|---|---|---|---|
| V | 1 | --- | 100.076 | --- | --- | 10 |
| G | 2 | --- | 157.097 | 1052.511 | --- | 9 |
| S# | 3 | -1539.952 | **464.187** | 995.489 | --- | 8 |
| I | 4 | --- | 577.272 | **688.399** | 252.695 | 7 |
| N | 5 | --- | 691.314 | 575.315 | --- | 6 |
| I | 6 | --- | 804.399 | **461.272** | 307.605 | 5 |
| G | 7 | --- | 861.420 | **348.188** | 473.198 | 4 |
| S | 8 | --- | 948.452 | **291.166** | 115.991 | 3 |
| G | 9 | --- | 1005.473 | **204.134** | -585.581 | 2 |
| K | 10 | --- | --- | 147.113 | --- | 1 |

**Soleil Index**

| | | |
|---|---|---|
| 466.4405 | not explained | 100% |

| | | | | | |
|---|---|---|---|---|---|
| **Run:** | 77662 | **Scans:** | 13458 - 13458 | **m/z Range:** | 219.17911 to 979.64697 |
| **Search:** | 77662 | **Precursor Mass:** | 576.29355 | | |
| **Mods:** | C=160.03 *+15.99 #+220.06 @+291.1 | **Charge:** | +2 | | |

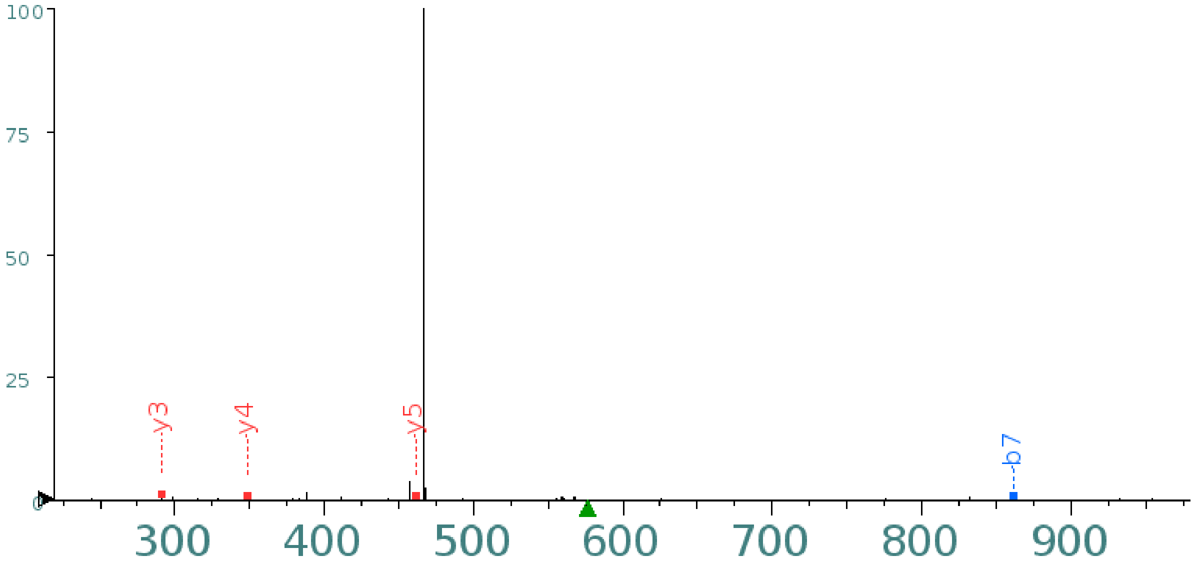

**Sequence**

VGS#INIGSGK

**Predicted Fragmentation Pattern**

| Seq | # | b: Δ Error | b | y | y: Δ Error | +1 |
|---|---|---|---|---|---|---|
| V | 1 | --- | 100.076 | --- | --- | 10 |
| G | 2 | --- | 157.097 | 1052.511 | --- | 9 |
| S# | 3 | --- | 464.187 | 995.489 | --- | 8 |
| I | 4 | --- | 577.272 | 688.399 | --- | 7 |
| N | 5 | --- | 691.314 | 575.315 | --- | 6 |
| I | 6 | --- | 804.399 | **461.272** | 44.911 | 5 |
| G | 7 | 68.207 | **861.420** | **348.188** | 435.369 | 4 |
| S | 8 | --- | 948.452 | **291.166** | 151.617 | 3 |
| G | 9 | --- | 1005.473 | 204.134 | --- | 2 |
| K | 10 | --- | --- | 147.113 | --- | 1 |

**Soleil Index**

| | | |
|---|---|---|
| 466.3629 | not explained | 100% |

| | | | | | |
|---|---|---|---|---|---|
| **Run:** | 77662 | **Scans:** | 13675 - 13675 | **m/z Range:** | 177.02266 to 1344.0434 |
| **Search:** | 77662 | **Precursor Mass:** | 679.32803 | | |
| **Mods:** | C=160.03 *+15.99 #+220.06 @+291.1 | **Charge:** | +2 | | |

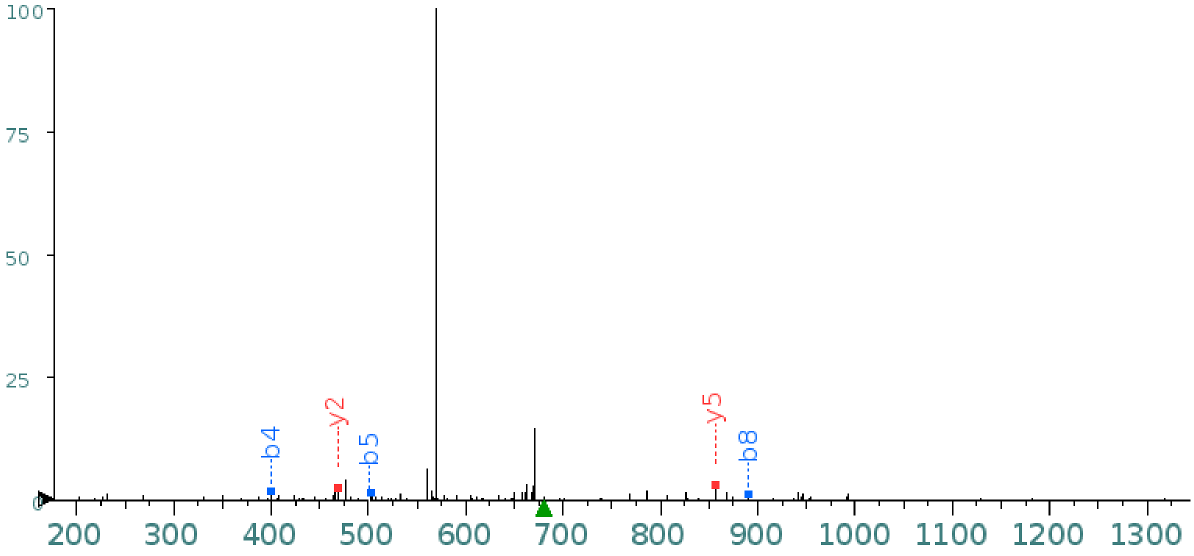

**Sequence**

IASQTQFNT#K

**Predicted Fragmentation Pattern**

| Seq | # | b: Δ Error | b | y | y: Δ Error | +1 |
|---|---|---|---|---|---|---|
| I | 1 | --- | 114.091 | --- | --- | 10 |
| A | 2 | --- | 185.128 | 1244.564 | --- | 9 |
| S | 3 | --- | 272.160 | 1173.527 | --- | 8 |
| Q | 4 | -3.290 | **400.219** | 1086.495 | --- | 7 |
| T | 5 | 938.117 | **501.267** | 958.436 | --- | 6 |
| Q | 6 | --- | 629.325 | **857.389** | -58.412 | 5 |
| F | 7 | --- | 776.394 | 729.330 | --- | 4 |
| N | 8 | -114.154 | **890.437** | 582.262 | --- | 3 |
| T# | 9 | --- | 1211.543 | **468.219** | 156.593 | 2 |
| K | 10 | --- | --- | 147.113 | --- | 1 |

**Soleil Index**

| | | |
|---|---|---|
| 569.3335 | not explained | 100% |

| | | | | | |
|---|---|---|---|---|---|
| **Run:** | 77662 | **Scans:** | 13805 - 13805 | **m/z Range:** | 212.13155 to 1047.6303 |
| **Search:** | 77662 | **Precursor Mass:** | 576.2935 | | |
| **Mods:** | C=160.03 *+15.99 #+220.06 @+291.1 | **Charge:** | +2 | | |

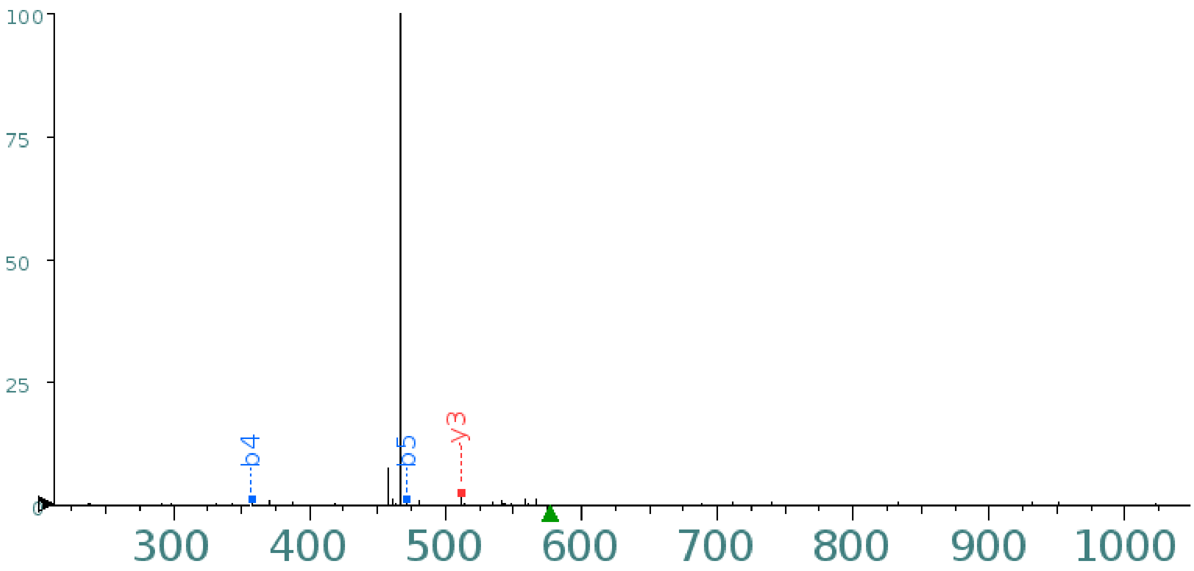

**Sequence**

VGSINIGS#GK

**Predicted Fragmentation Pattern**

| Seq | # | b: Δ Error | b | y | y: Δ Error | +1 |
|---|---|---|---|---|---|---|
| V | 1 | --- | 100.076 | --- | --- | 10 |
| G | 2 | --- | 157.097 | 1052.511 | --- | 9 |
| S | 3 | --- | 244.129 | 995.489 | --- | 8 |
| I | 4 | 343.350 | **357.213** | 908.457 | --- | 7 |
| N | 5 | 28.862 | **471.256** | 795.373 | --- | 6 |
| I | 6 | --- | 584.340 | 681.330 | --- | 5 |
| G | 7 | --- | 641.362 | 568.246 | --- | 4 |
| S# | 8 | --- | 948.452 | **511.225** | 565.733 | 3 |
| G | 9 | --- | 1005.473 | 204.134 | --- | 2 |
| K | 10 | --- | --- | 147.113 | --- | 1 |

**Soleil Index**

| | | |
|---|---|---|
| 466.3886 | not explained | 100% |

| | | | | | |
|---|---|---|---|---|---|
| **Run:** | 77662 | **Scans:** | 13906 - 13906 | **m/z Range:** | 191.18785 to 1251.7965 |
| **Search:** | 77662 | **Precursor Mass:** | 686.32264 | | |
| **Mods:** | C=160.03 *+15.99 #+220.06 @+291.1 | **Charge:** | +2 | | |

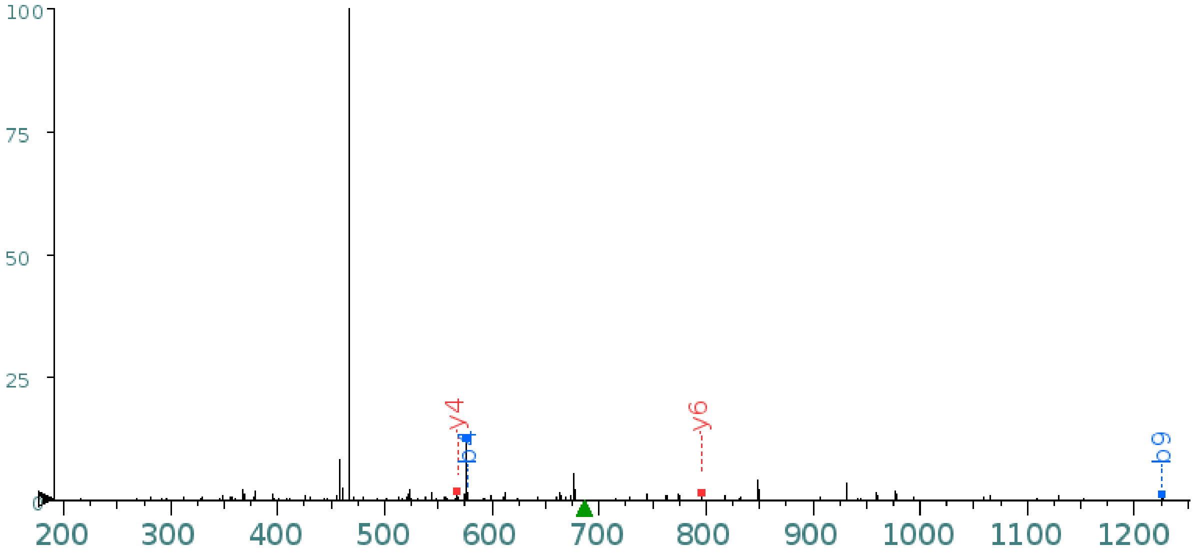

**Sequence**

VGS#INIGS#GK

**Predicted Fragmentation Pattern**

| Seq | # | b: Δ Error | b | y | y: Δ Error | +1 |
|---|---|---|---|---|---|---|
| V | 1 | --- | 100.076 | --- | --- | 10 |
| G | 2 | --- | 157.097 | 1272.569 | --- | 9 |
| S# | 3 | --- | 464.187 | 1215.547 | --- | 8 |
| I | 4 | -1686.385 | **577.272** | 908.457 | --- | 7 |
| N | 5 | --- | 691.314 | **795.373** | 333.308 | 6 |
| I | 6 | --- | 804.399 | 681.330 | --- | 5 |
| G | 7 | --- | 861.420 | **568.246** | -1459.619 | 4 |
| S# | 8 | --- | 1168.510 | 511.225 | --- | 3 |
| G | 9 | 46.369 | **1225.532** | 204.134 | --- | 2 |
| K | 10 | --- | --- | 147.113 | --- | 1 |

**Soleil Index**

| | | |
|---|---|---|
| 466.3044 | not explained | 100% |

| | | | | | |
|---|---|---|---|---|---|
| **Run:** | 77662 | **Scans:** | 14512 - 14512 | **m/z Range:** | 161.11572 to 1000.6856 |
| **Search:** | 77662 | **Precursor Mass:** | 576.29338 | | |
| **Mods:** | C=160.03 *+15.99 #+220.06 @+291.1 | **Charge:** | +2 | | |

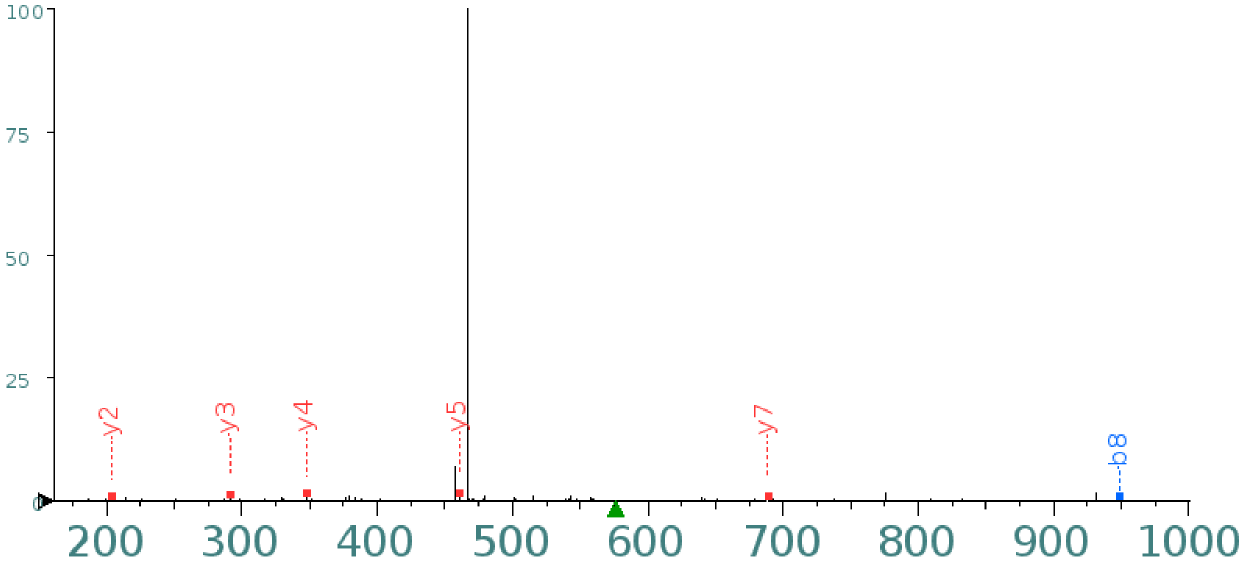

**Sequence**

VGS#INIGSGK

**Predicted Fragmentation Pattern**

| Seq | # | b: Δ Error | b | y | y: Δ Error | +1 |
|---|---|---|---|---|---|---|
| V | 1 | --- | 100.076 | --- | --- | 10 |
| G | 2 | --- | 157.097 | 1052.511 | --- | 9 |
| S# | 3 | --- | 464.187 | 995.489 | --- | 8 |
| I | 4 | --- | 577.272 | **688.399** | 616.250 | 7 |
| N | 5 | --- | 691.314 | 575.315 | --- | 6 |
| I | 6 | --- | 804.399 | **461.272** | 282.346 | 5 |
| G | 7 | --- | 861.420 | **348.188** | 659.849 | 4 |
| S | 8 | 400.301 | **948.452** | **291.166** | 417.478 | 3 |
| G | 9 | --- | 1005.473 | **204.134** | -178.936 | 2 |
| K | 10 | --- | --- | 147.113 | --- | 1 |

**Soleil Index**

| | | |
|---|---|---|
| 466.2990 | not explained | 100% |

| | | | | | |
|---|---|---|---|---|---|
| **Run:** | 77662 | **Scans:** | 14857 - 14857 | **m/z Range:** | 148.36975 to 978.37634 |
| **Search:** | 77662 | **Precursor Mass:** | 576.29343 | | |
| **Mods:** | C=160.03 *+15.99 #+220.06 @+291.1 | **Charge:** | +2 | | |

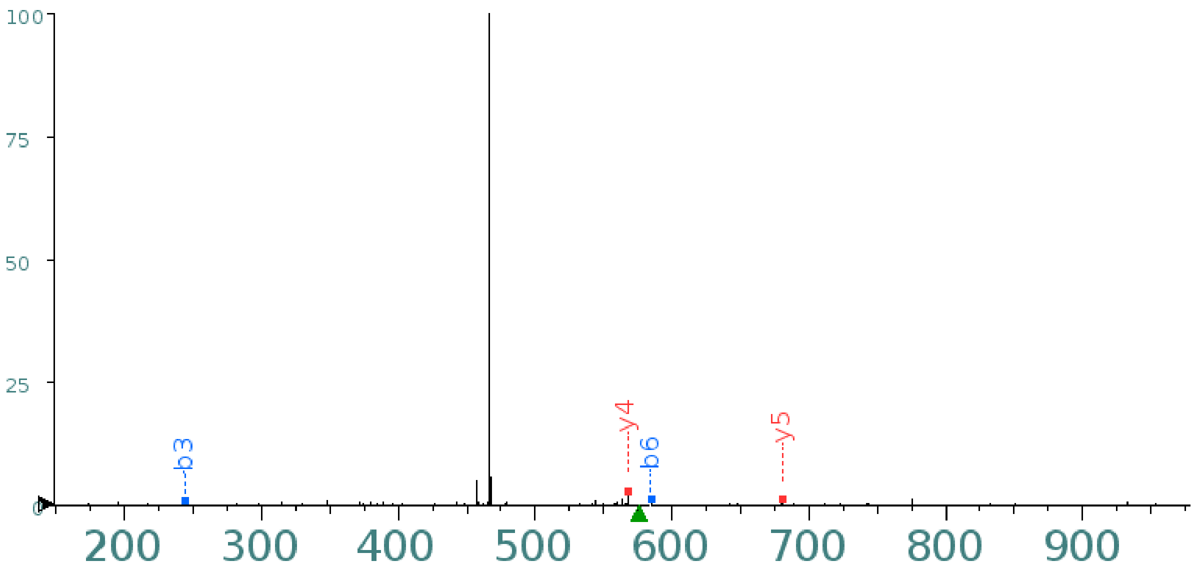

**Sequence**

VGSINIGS#GK

**Predicted Fragmentation Pattern**

| Seq | # | b: Δ Error | b | y | y: Δ Error | +1 |
|---|---|---|---|---|---|---|
| V | 1 | --- | 100.076 | --- | --- | 10 |
| G | 2 | --- | 157.097 | 1052.511 | --- | 9 |
| S | 3 | 231.702 | 244.129 | 995.489 | --- | 8 |
| I | 4 | --- | 357.213 | 908.457 | --- | 7 |
| N | 5 | --- | 471.256 | 795.373 | --- | 6 |
| I | 6 | 491.260 | 584.340 | 681.330 | -196.740 | 5 |
| G | 7 | --- | 641.362 | 568.246 | -1469.098 | 4 |
| S# | 8 | --- | 948.452 | 511.225 | --- | 3 |
| G | 9 | --- | 1005.473 | 204.134 | --- | 2 |
| K | 10 | --- | --- | 147.113 | --- | 1 |

**Soleil Index**

| | | |
|---|---|---|
| 466.3721 | not explained | 100% |

| | | | | | |
|---|---|---|---|---|---|
| **Run:** | 77662 | **Scans:** | 16526 - 16526 | **m/z Range:** | 184.06534 to 1280.8560[illegible] |
| **Search:** | 77662 | **Precursor Mass:** | 686.32263 | | |
| **Mods:** | C=160.03 *+15.99 #+220.06 @+291.1 | **Charge:** | +2 | | |

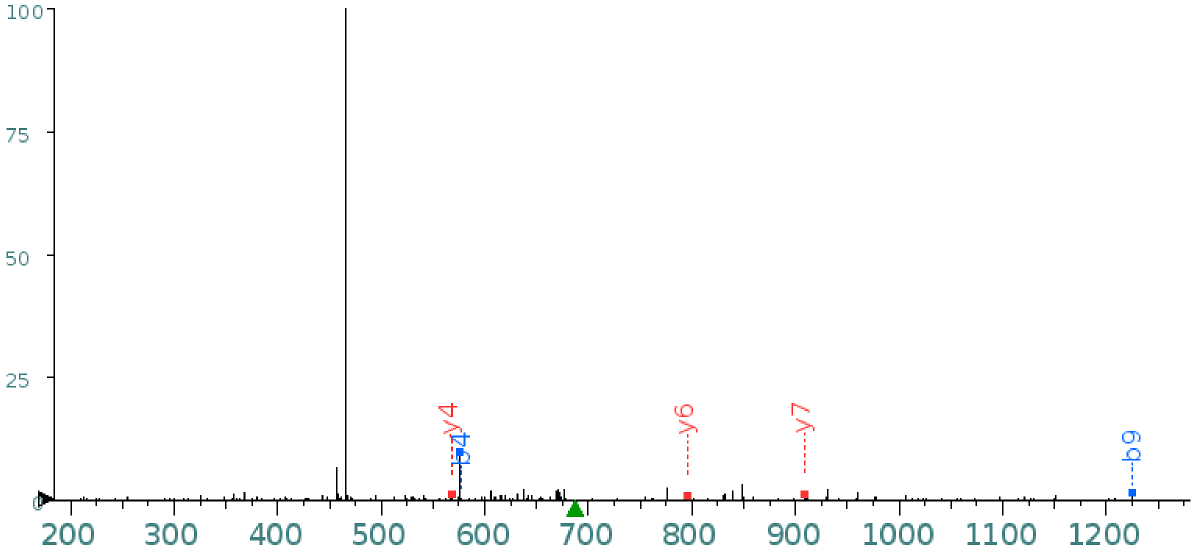

**Sequence**

VGS#INIGS#GK

**Predicted Fragmentation Pattern**

| Seq | # | b: Δ Error | b | y | y: Δ Error | +1 |
|---|---|---|---|---|---|---|
| V | 1 | --- | 100.076 | --- | --- | 10 |
| G | 2 | --- | 157.097 | 1272.569 | --- | 9 |
| S# | 3 | --- | 464.187 | 1215.547 | --- | 8 |
| I | 4 | -1569.279 | **577.272** | **908.457** | 412.055 | 7 |
| N | 5 | --- | 691.314 | **795.373** | 354.627 | 6 |
| I | 6 | --- | 804.399 | 681.330 | --- | 5 |
| G | 7 | --- | 861.420 | **568.246** | 427.367 | 4 |
| S# | 8 | --- | 1168.510 | 511.225 | --- | 3 |
| G | 9 | -473.896 | **1225.532** | 204.134 | --- | 2 |
| K | 10 | --- | --- | 147.113 | --- | 1 |

**Soleil Index**

| | | |
|---|---|---|
| 466.3449 | not explained | 100% |

**Run:** 77662

**Scans:** 16636 - 16636

**m/z Range:** 176.03006 to 957.88385

**Search:** 77662

**Precursor Mass:** 576.29353

**Mods:** C=160.03 *+15.99 #+220.06 @+291.1

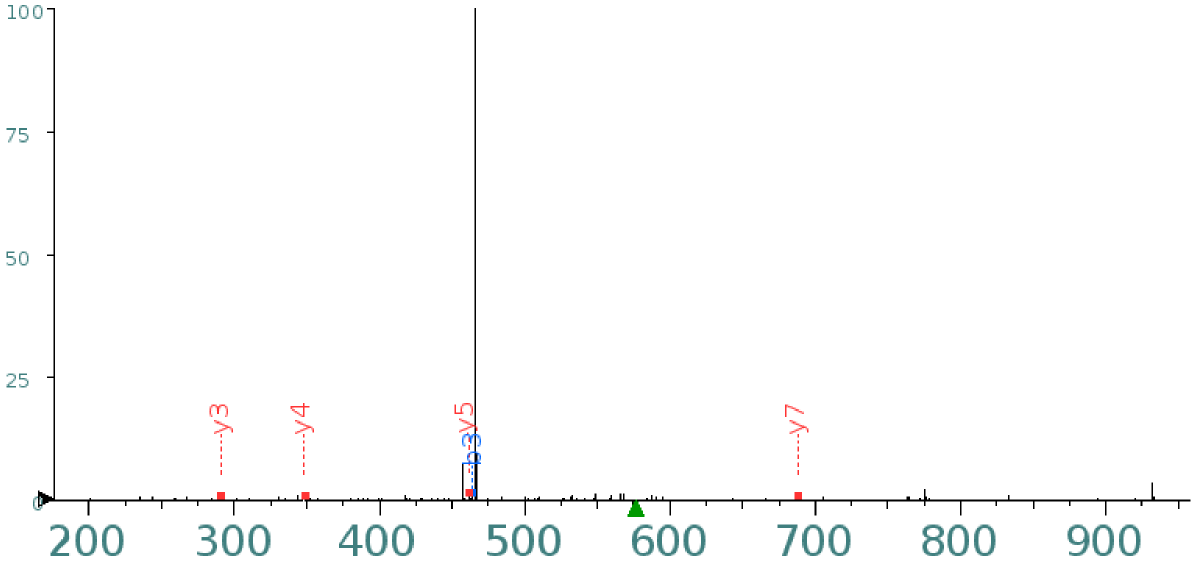

**Sequence**

VGS#INIGSGK

**Predicted Fragmentation Pattern**

| Seq | # | b: Δ Error | b | y | y: Δ Error | +1 |
|---|---|---|---|---|---|---|
| V | 1 | --- | 100.076 | --- | --- | 10 |
| G | 2 | --- | 157.097 | 1052.511 | --- | 9 |
| S# | 3 | -1748.519 | **464.187** | 995.489 | --- | 8 |
| I | 4 | --- | 577.272 | **688.399** | 369.568 | 7 |
| N | 5 | --- | 691.314 | 575.315 | --- | 6 |
| I | 6 | --- | 804.399 | **461.272** | 459.126 | 5 |
| G | 7 | --- | 861.420 | **348.188** | 664.489 | 4 |
| S | 8 | --- | 948.452 | **291.166** | 373.701 | 3 |
| G | 9 | --- | 1005.473 | 204.134 | --- | 2 |
| K | 10 | --- | --- | 147.113 | --- | 1 |

**Soleil Index**

466.2802 **not explained** **100%**

| Run: | 77662 | Scans: | 16886 - 16886 | m/z Range: | 176.29419 to 1351.7557[illegible] |
|---|---|---|---|---|---|
| Search: | 77662 | Precursor Mass: | 686.32243 | | |
| Mods: | C=160.03 *+15.99 #+220.06 @+291.1 | Charge: | +2 | | |

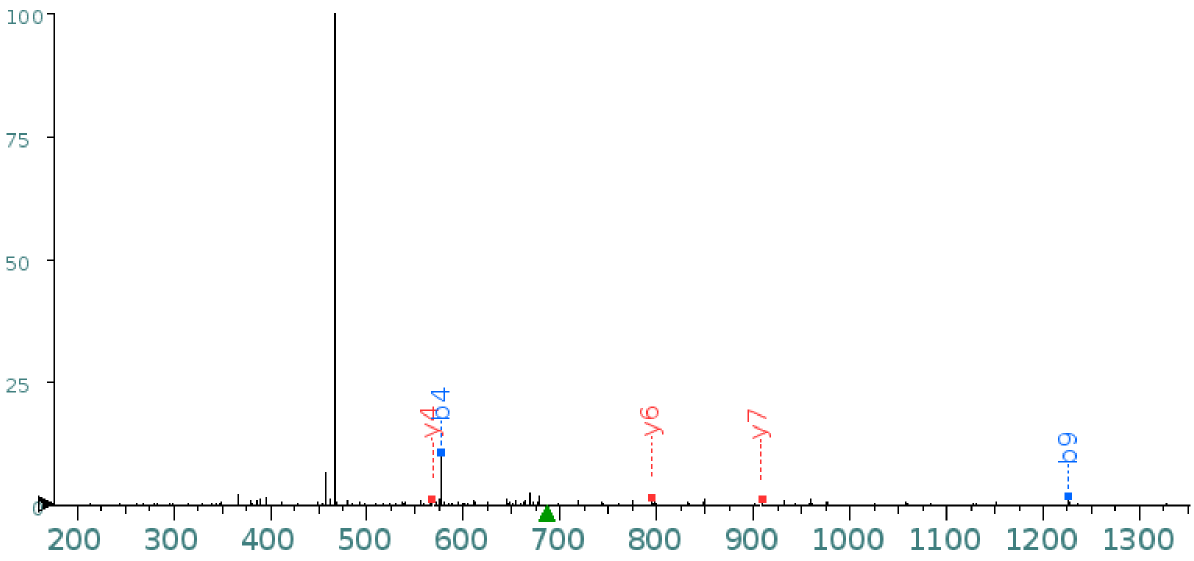

**Sequence**

VGS#INIGS#GK

**Predicted Fragmentation Pattern**

| Seq | # | b: Δ Error | b | y | y: Δ Error | +1 |
|---|---|---|---|---|---|---|
| V | 1 | --- | 100.076 | --- | --- | 10 |
| G | 2 | --- | 157.097 | 1272.569 | --- | 9 |
| S# | 3 | --- | 464.187 | 1215.547 | --- | 8 |
| I | 4 | -1348.399 | 577.272 | 908.457 | 259.377 | 7 |
| N | 5 | --- | 691.314 | 795.373 | 139.023 | 6 |
| I | 6 | --- | 804.399 | 681.330 | --- | 5 |
| G | 7 | --- | 861.420 | 568.246 | -1464.035 | 4 |
| S# | 8 | --- | 1168.510 | 511.225 | --- | 3 |
| G | 9 | 219.737 | 1225.532 | 204.134 | --- | 2 |
| K | 10 | --- | --- | 147.113 | --- | 1 |

**Soleil Index**

| | | |
|---|---|---|
| 466.2746 | not explained | 100% |

| Run: | 77662 | Scans: | 18050 - 18050 | m/z Range: | 372.47162 to 1675.8526 |
|---|---|---|---|---|---|
| Search: | 77662 | Precursor Mass: | 1008.97025 | | |
| Mods: | C=160.03 *+15.99 #+220.06 @+291.1 | Charge: | +2 | | |

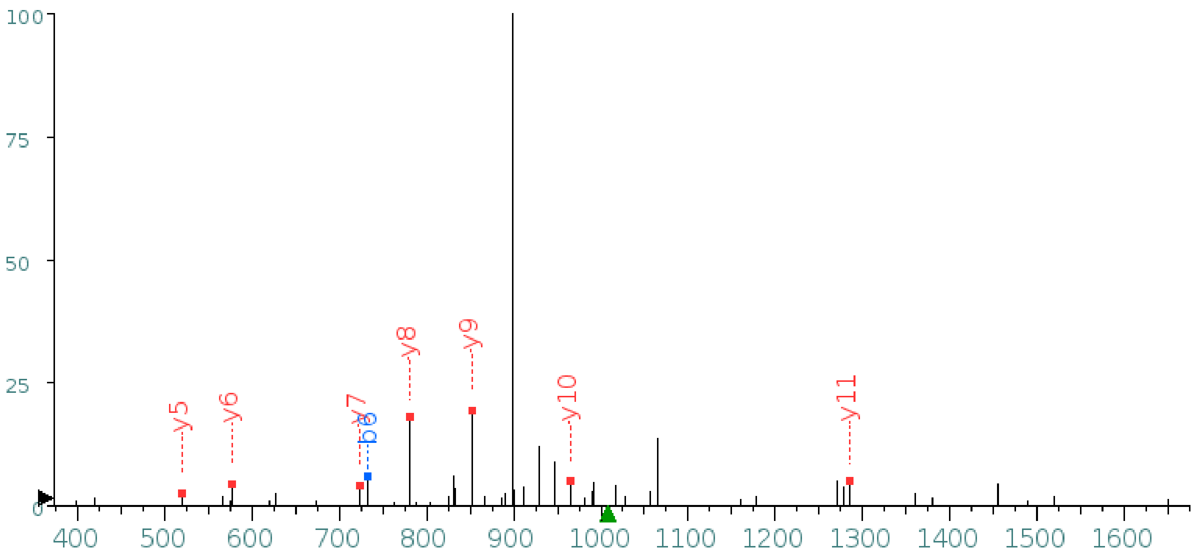

**Sequence**

DQVM*ELT#IAGM*GTSALK

**Predicted Fragmentation Pattern**

| Seq | # | b: Δ Error | b | y | y: Δ Error | +1 |
|---|---|---|---|---|---|---|
| D | 1 | --- | 116.034 | --- | --- | 17 |
| Q | 2 | --- | 244.093 | 1901.908 | --- | 16 |
| V | 3 | --- | 343.161 | 1773.850 | --- | 15 |
| M* | 4 | --- | 490.197 | 1674.781 | --- | 14 |
| E | 5 | --- | 619.239 | 1527.746 | --- | 13 |
| L | 6 | 256.685 | **732.323** | 1398.703 | --- | 12 |
| T# | 7 | --- | 1053.429 | **1285.619** | 128.332 | 11 |
| I | 8 | --- | 1166.513 | **964.513** | 207.508 | 10 |
| A | 9 | --- | 1237.550 | **851.429** | 124.942 | 9 |
| G | 10 | --- | 1294.572 | **780.392** | 231.858 | 8 |
| M* | 11 | --- | 1441.607 | **723.371** | 481.713 | 7 |
| G | 12 | --- | 1498.629 | **576.335** | 282.769 | 6 |
| T | 13 | --- | 1599.676 | **519.314** | 376.839 | 5 |
| S | 14 | --- | 1686.708 | 418.266 | --- | 4 |
| A | 15 | --- | 1757.746 | 331.234 | --- | 3 |
| L | 16 | --- | 1870.830 | 260.197 | --- | 2 |
| K | 17 | --- | --- | 147.113 | --- | 1 |

**Soleil Index**

| | | |
|---|---|---|
| 899.0646 | not explained | 100% |

| | | | | | |
|---|---|---|---|---|---|
| **Run:** | 77662 | **Scans:** | 18078 - 18078 | **m/z Range:** | 347.45917 to 1675.922 |
| **Search:** | 77662 | **Precursor Mass:** | 1008.97025 | | |
| **Mods:** | C=160.03 *+15.99 #+220.06 @+291.1 | **Charge:** | +2 | | |

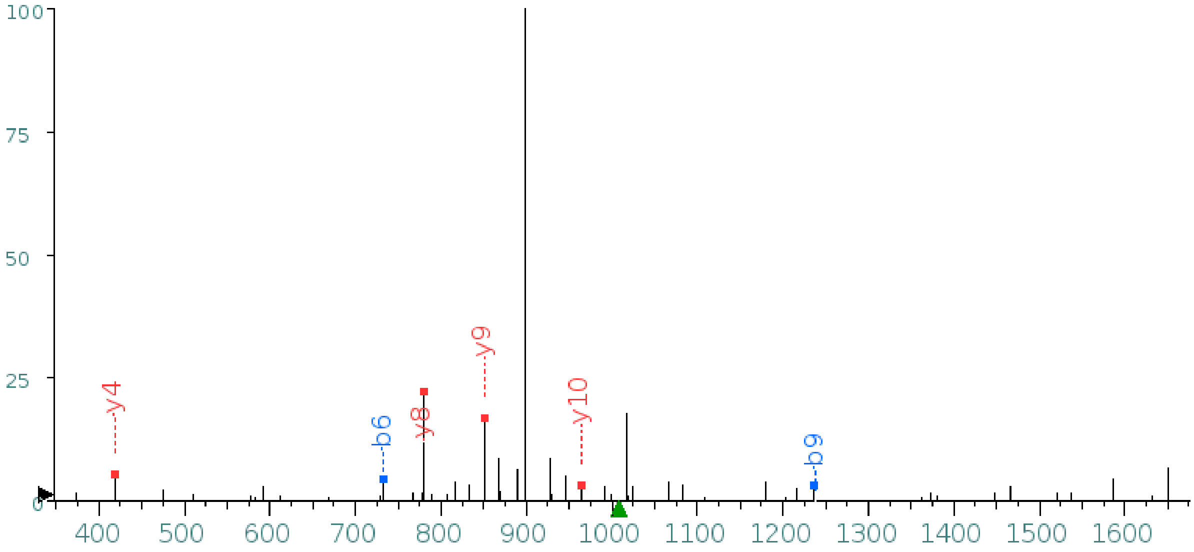

**Sequence**

DQVM*ELT#IAGM*GTSALK

**Predicted Fragmentation Pattern**

| Seq | # | b: Δ Error | b | y | y: Δ Error | +1 |
|---|---|---|---|---|---|---|
| D | 1 | --- | 116.034 | --- | --- | 17 |
| Q | 2 | --- | 244.093 | 1901.908 | --- | 16 |
| V | 3 | --- | 343.161 | 1773.850 | --- | 15 |
| M* | 4 | --- | 490.197 | 1674.781 | --- | 14 |
| E | 5 | --- | 619.239 | 1527.746 | --- | 13 |
| L | 6 | 441.164 | **732.323** | 1398.703 | --- | 12 |
| T# | 7 | --- | 1053.429 | 1285.619 | --- | 11 |
| I | 8 | --- | 1166.513 | **964.513** | 306.554 | 10 |
| A | 9 | -424.530 | **1237.550** | **851.429** | 314.969 | 9 |
| G | 10 | --- | 1294.572 | **780.392** | 265.316 | 8 |
| M* | 11 | --- | 1441.607 | 723.371 | --- | 7 |
| G | 12 | --- | 1498.629 | 576.335 | --- | 6 |
| T | 13 | --- | 1599.676 | 519.314 | --- | 5 |
| S | 14 | --- | 1686.708 | **418.266** | 519.757 | 4 |
| A | 15 | --- | 1757.746 | 331.234 | --- | 3 |
| L | 16 | --- | 1870.830 | 260.197 | --- | 2 |
| K | 17 | --- | --- | 147.113 | --- | 1 |

**Soleil Index**

| | | |
|---|---|---|
| 899.1660 | not explained | 100% |
| 1017.7391 | not explained | 18% |

| | | | |
|---|---|---|---|
| **Run:** | 77662 | **Scans:** | 18340 - 18340 |
| **Search:** | 77662 | **Precursor Mass:** | 783.37749 |
| **Mods:** | C=160.03 *+15.99 #+220.06 @+291.1 | **Charge:** | +3 |

**m/z Range:** 198.36079 to 1972.3951

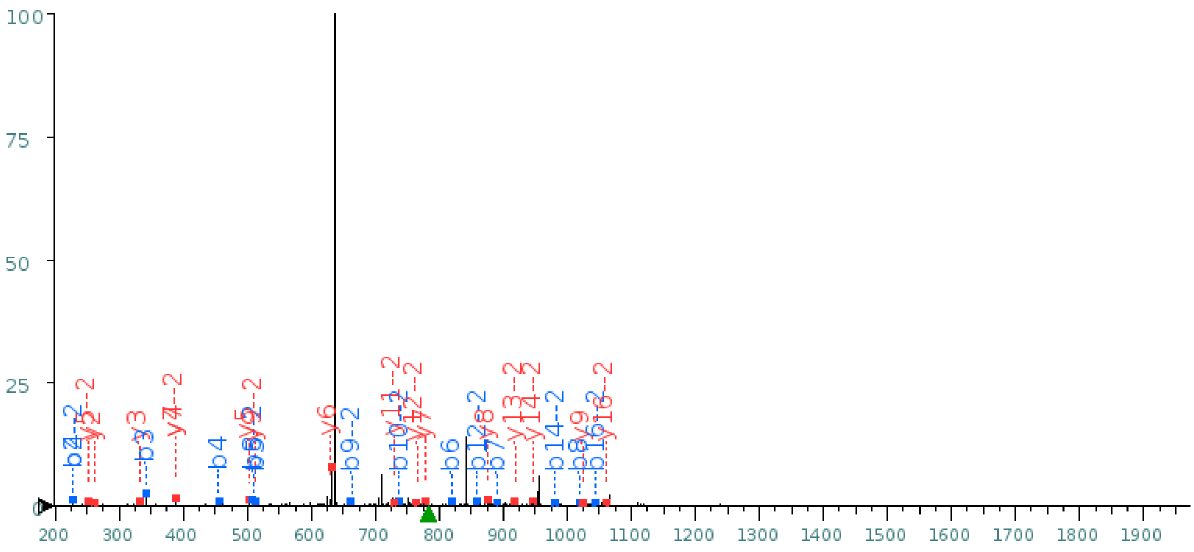

**Sequence**

NLLNGS#AKS#M*VFQIGANK

**Predicted Fragmentation Pattern**

+1

| Seq | # | b: Δ Error | b | y | y: Δ Error | +1 |
|---|---|---|---|---|---|---|
| N | 1 | --- | 115.050 | --- | --- | 18 |
| L | 2 | 409.547 | **228.134** | 2234.074 | --- | 17 |
| L | 3 | 146.272 | **341.218** | 2120.990 | --- | 16 |
| N | 4 | 461.583 | **455.261** | 2007.906 | --- | 15 |
| G | 5 | 341.453 | **512.283** | 1893.863 | --- | 14 |
| S# | 6 | -33.452 | **819.373** | 1836.842 | --- | 13 |
| A | 7 | -563.923 | **890.410** | 1529.752 | --- | 12 |
| K | 8 | 342.001 | **1018.505** | 1458.714 | --- | 11 |
| S# | 9 | --- | 1325.595 | 1330.620 | --- | 10 |
| M* | 10 | --- | 1472.631 | **1023.529** | 260.879 | 9 |
| V | 11 | --- | 1571.699 | **876.494** | 386.254 | 8 |
| F | 12 | --- | 1718.768 | **777.425** | 481.174 | 7 |
| Q | 13 | --- | 1846.826 | **630.357** | 1532.608 | 6 |
| I | 14 | --- | 1959.910 | **502.298** | 408.532 | 5 |
| G | 15 | --- | 2016.932 | **389.214** | 284.972 | 4 |
| A | 16 | --- | 2087.969 | **332.193** | 340.879 | 3 |
| N | 17 | --- | 2202.012 | **261.156** | 407.775 | 2 |
| K | 18 | --- | --- | 147.113 | --- | 1 |

| | | | +2 | | | |
|---|---|---|---|---|---|---|
| Seq | # | b: Δ Error | b | y | y: Δ Error | +1 |
| N | 1 | --- | 58.029 | --- | --- | 18 |
| L | 2 | --- | 114.571 | 1117.541 | --- | 17 |
| L | 3 | --- | 171.113 | **1060.999** | -362.931 | 16 |
| N | 4 | 409.547 | **228.134** | 1004.457 | --- | 15 |
| G | 5 | --- | 256.645 | **947.435** | -557.774 | 14 |
| S# | 6 | --- | 410.190 | **918.925** | -1070.879 | 13 |
| A | 7 | --- | 445.709 | **765.379** | -659.413 | 12 |
| K | 8 | -1802.440 | **509.756** | **729.861** | 341.784 | 11 |
| S# | 9 | -1350.572 | **663.301** | 665.813 | --- | 10 |
| M* | 10 | 1093.516 | **736.819** | **512.268** | 369.739 | 9 |
| V | 11 | --- | 786.353 | 438.751 | --- | 8 |
| F | 12 | -736.002 | **859.887** | **389.216** | 279.807 | 7 |
| Q | 13 | --- | 923.917 | 315.682 | --- | 6 |
| I | 14 | 338.597 | **980.459** | **251.653** | 2343.738 | 5 |
| G | 15 | --- | 1008.970 | 195.111 | --- | 4 |
| A | 16 | -626.169 | **1044.488** | 166.600 | --- | 3 |
| N | 17 | --- | 1101.510 | 131.082 | --- | 2 |
| K | 18 | --- | --- | 74.060 | --- | 1 |

**Soleil Index**

637.0648 **not explained** **100%**

| | | | |
|---|---|---|---|
| **Run:** | 77662 | **Scans:** | 18432 - 18432 |
| **Search:** | 77662 | **Precursor Mass:** | 1008.97003 |
| **Mods:** | C=160.03 *+15.99 #+220.06 @+291.1 | **Charge:** | +2 |

**m/z Range:** 331.55884 **to** 1869.1358

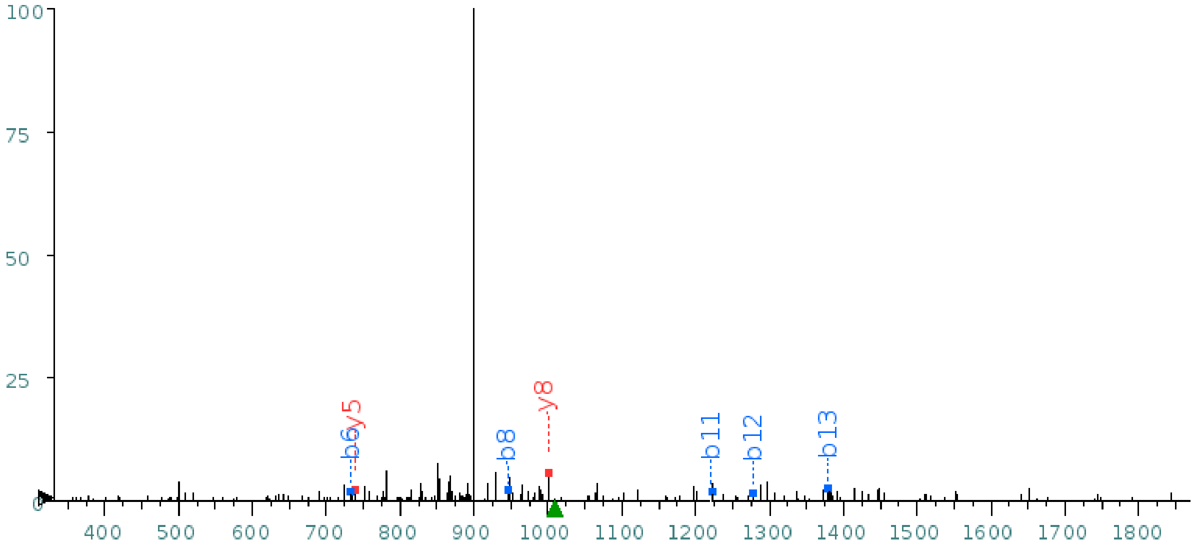

**Sequence**

DQVM*ELTIAGM*GTS#ALK

**Predicted Fragmentation Pattern**

| Seq | # | b: Δ Error | b | y | y: Δ Error | +1 |
|---|---|---|---|---|---|---|
| D | 1 | --- | 116.034 | --- | --- | 17 |
| Q | 2 | --- | 244.093 | 1901.908 | --- | 16 |
| V | 3 | --- | 343.161 | 1773.850 | --- | 15 |
| M* | 4 | --- | 490.197 | 1674.781 | --- | 14 |
| E | 5 | --- | 619.239 | 1527.746 | --- | 13 |
| L | 6 | 253.103 | **732.323** | 1398.703 | --- | 12 |
| T | 7 | --- | 833.371 | 1285.619 | --- | 11 |
| I | 8 | 276.908 | **946.455** | 1184.571 | --- | 10 |
| A | 9 | --- | 1017.492 | 1071.487 | --- | 9 |
| G | 10 | --- | 1074.514 | **1000.450** | -350.672 | 8 |
| M* | 11 | 245.840 | **1221.549** | 943.429 | --- | 7 |
| G | 12 | -533.041 | **1278.570** | 796.393 | --- | 6 |
| T | 13 | 315.981 | **1379.618** | **739.372** | 283.506 | 5 |
| S# | 14 | --- | 1686.708 | 638.324 | --- | 4 |
| A | 15 | --- | 1757.746 | 331.234 | --- | 3 |
| L | 16 | --- | 1870.830 | 260.197 | --- | 2 |
| K | 17 | --- | --- | 147.113 | --- | 1 |

**Soleil Index**

| | | |
|---|---|---|
| 899.4590 | not explained | 100% |

| Run: | 77662 | Scans: | 18572 - 18572 | m/z Range: | 274.21793 | to | 1914.1964 |
|---|---|---|---|---|---|---|---|
| Search: | 77662 | Precursor Mass: | 1008.96979 | | | | |
| Mods: | C=160.03 *+15.99 #+220.06 @+291.1 | Charge: | +2 | | | | |

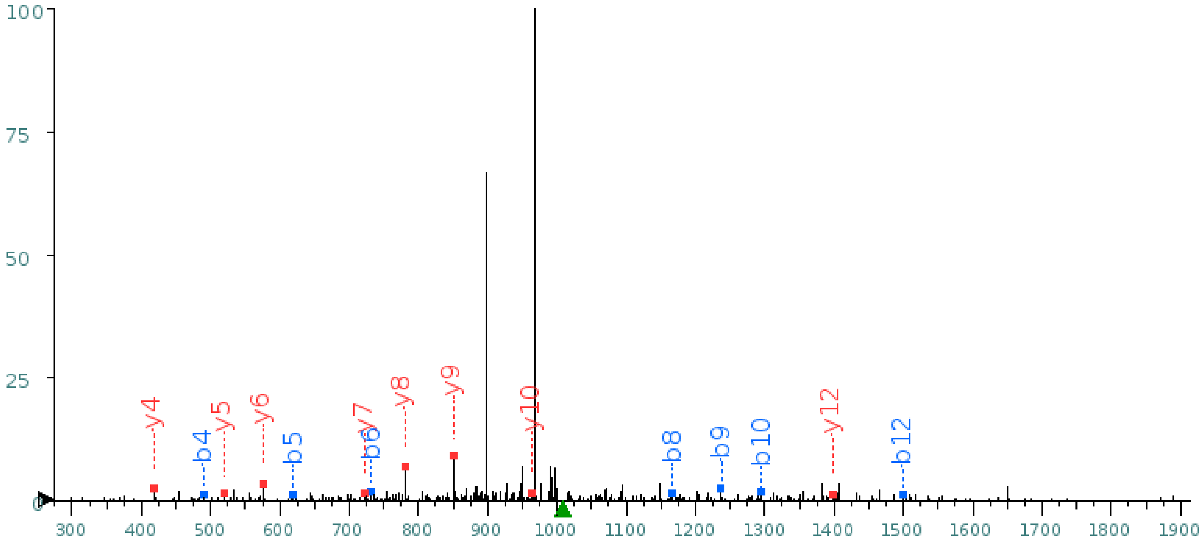

### Sequence

DQVM*ELT#IAGM*GTSALK

### Predicted Fragmentation Pattern

| Seq | # | b: Δ Error | b | y | y: Δ Error | +1 |
|---|---|---|---|---|---|---|
| D | 1 | --- | 116.034 | --- | --- | 17 |
| Q | 2 | --- | 244.093 | 1901.908 | --- | 16 |
| V | 3 | --- | 343.161 | 1773.850 | --- | 15 |
| M* | 4 | 145.003 | **490.197** | 1674.781 | --- | 14 |
| E | 5 | 555.685 | **619.239** | 1527.746 | --- | 13 |
| L | 6 | 87.637 | **732.323** | **1398.703** | 349.941 | 12 |
| T# | 7 | --- | 1053.429 | 1285.619 | --- | 11 |
| I | 8 | 230.582 | **1166.513** | **964.513** | -56.587 | 10 |
| A | 9 | -269.066 | **1237.550** | **851.429** | -4.650 | 9 |
| G | 10 | -589.392 | **1294.572** | **780.392** | 244.053 | 8 |
| M* | 11 | --- | 1441.607 | **723.371** | -1124.911 | 7 |
| G | 12 | 284.300 | **1498.629** | **576.335** | 128.110 | 6 |
| T | 13 | --- | 1599.676 | **519.314** | 596.172 | 5 |
| S | 14 | --- | 1686.708 | **418.266** | 297.550 | 4 |
| A | 15 | --- | 1757.746 | 331.234 | --- | 3 |
| L | 16 | --- | 1870.830 | 260.197 | --- | 2 |
| K | 17 | --- | --- | 147.113 | --- | 1 |

**Soleil Index**

| | | |
|---|---|---|
| 898.8270 | not explained | 67% |
| 968.8347 | not explained | 100% |

| | | | |
|---|---|---|---|
| **Run:** | 77662 | **Scans:** | 18830 - 18830 |
| **Search:** | 77662 | **Precursor Mass:** | 1008.97031 |
| **Mods:** | C=160.03 *+15.99 #+220.06 @+291.1 | | |

**m/z Range:** 271.3671 **to** 2000

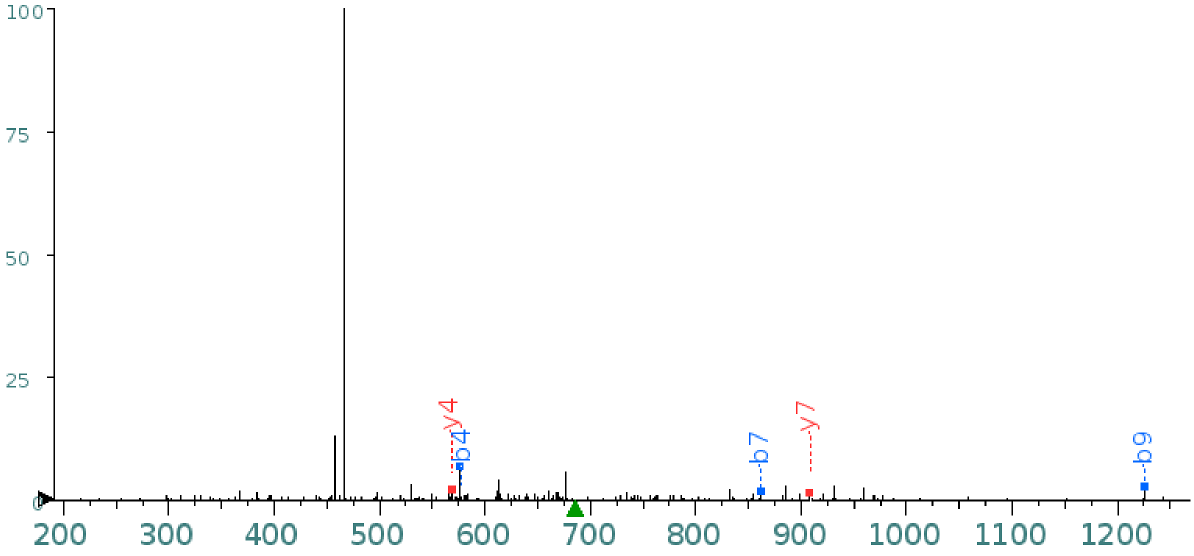

**Sequence**

VGS#INIGS#GK

**Predicted Fragmentation Pattern**

| Seq | # | b: Δ Error | b | y | y: Δ Error | +1 |
|---|---|---|---|---|---|---|
| V | 1 | --- | 100.076 | --- | --- | 10 |
| G | 2 | --- | 157.097 | 1272.569 | --- | 9 |
| S# | 3 | --- | 464.187 | 1215.547 | --- | 8 |
| I | 4 | -1587.416 | **577.272** | **908.457** | -428.315 | 7 |
| N | 5 | --- | 691.314 | 795.373 | --- | 6 |
| I | 6 | --- | 804.399 | 681.330 | --- | 5 |
| G | 7 | 503.498 | **861.420** | **568.246** | -70.622 | 4 |
| S# | 8 | --- | 1168.510 | 511.225 | --- | 3 |
| G | 9 | 267.624 | **1225.532** | 204.134 | --- | 2 |
| K | 10 | --- | --- | 147.113 | --- | 1 |

**Soleil Index**

| | | |
|---|---|---|
| 466.2672 | not explained | 100% |

| Run: | 77662 | Scans: | 19052 - 19052 | m/z Range: | 261.3186 to 1896.2403 |
|---|---|---|---|---|---|
| Search: | 77662 | Precursor Mass: | 1008.97105 | | |
| Mods: | C=160.03 *+15.99 #+220.06 @+291.1 | Charge: | +2 | | |

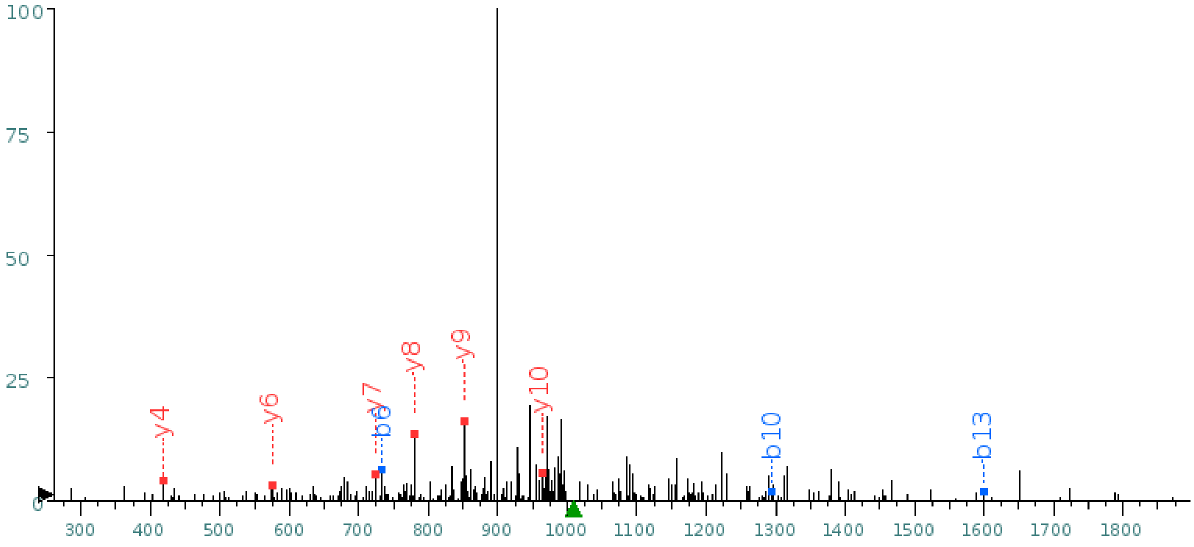

**Sequence**

DQVM*ELT#IAGM*GTSALK

**Predicted Fragmentation Pattern**

| Seq | # | b: Δ Error | b | y | y: Δ Error | +1 |
|---|---|---|---|---|---|---|
| D | 1 | --- | 116.034 | --- | --- | 17 |
| Q | 2 | --- | 244.093 | 1901.908 | --- | 16 |
| V | 3 | --- | 343.161 | 1773.850 | --- | 15 |
| M* | 4 | --- | 490.197 | 1674.781 | --- | 14 |
| E | 5 | --- | 619.239 | 1527.746 | --- | 13 |
| L | 6 | 283.424 | **732.323** | 1398.703 | --- | 12 |
| T# | 7 | --- | 1053.429 | 1285.619 | --- | 11 |
| I | 8 | --- | 1166.513 | **964.513** | 364.987 | 10 |
| A | 9 | --- | 1237.550 | **851.429** | 342.908 | 9 |
| G | 10 | -482.065 | **1294.572** | **780.392** | 262.189 | 8 |
| M* | 11 | --- | 1441.607 | **723.371** | 307.108 | 7 |
| G | 12 | --- | 1498.629 | **576.335** | 193.219 | 6 |
| T | 13 | 53.543 | **1599.676** | 519.314 | --- | 5 |
| S | 14 | --- | 1686.708 | **418.266** | 757.164 | 4 |
| A | 15 | --- | 1757.746 | 331.234 | --- | 3 |
| L | 16 | --- | 1870.830 | 260.197 | --- | 2 |
| K | 17 | --- | --- | 147.113 | --- | 1 |

**Soleil Index**

| | | |
|---|---|---|
| 899.0729 | not explained | 100% |
| 946.7196 | y1-10 frag loss of water/ y1-10 frag loss of ammonia | 19% |
| 970.0048 | not explained | 17% |
| 991.5348 | precursor loss of two waters | 16% |

| | | | | | | | |
|---|---|---|---|---|---|---|---|
| **Run:** | 77662 | **Scans:** | 19627 - 19627 | **m/z Range:** | 296.245 | to | 1973.1881 |
| **Search:** | 77662 | **Precursor Mass:** | 1008.97333 | | | | |
| **Mods:** | C=160.03 *+15.99 #+220.06 @+291.1 | **Charge:** | +2 | | | | |

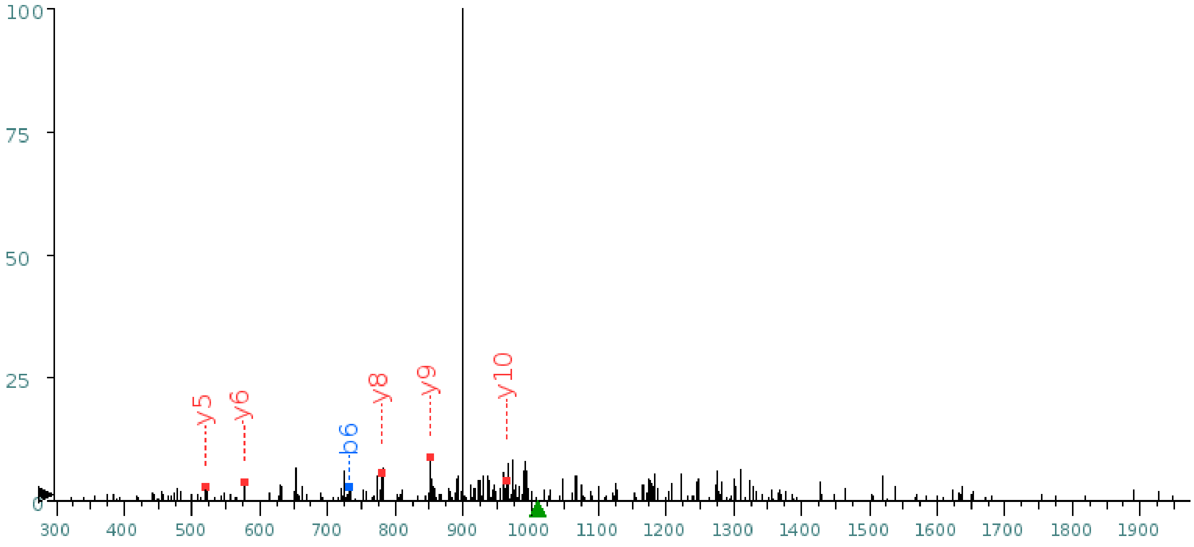

**Sequence**

DQVM*ELT#IAGM*GTSALK

**Predicted Fragmentation Pattern**

| Seq | # | b: Δ Error | b | y | y: Δ Error | +1 |
|---|---|---|---|---|---|---|
| D | 1 | --- | 116.034 | --- | --- | 17 |
| Q | 2 | --- | 244.093 | 1901.908 | --- | 16 |
| V | 3 | --- | 343.161 | 1773.850 | --- | 15 |
| M* | 4 | --- | 490.197 | 1674.781 | --- | 14 |
| E | 5 | --- | 619.239 | 1527.746 | --- | 13 |
| L | 6 | 475.638 | **732.323** | 1398.703 | --- | 12 |
| T# | 7 | --- | 1053.429 | 1285.619 | --- | 11 |
| I | 8 | --- | 1166.513 | **964.513** | 63.140 | 10 |
| A | 9 | --- | 1237.550 | **851.429** | 121.932 | 9 |
| G | 10 | --- | 1294.572 | **780.392** | 44.047 | 8 |
| M* | 11 | --- | 1441.607 | 723.371 | --- | 7 |
| G | 12 | --- | 1498.629 | **576.335** | -87.816 | 6 |
| T | 13 | --- | 1599.676 | **519.314** | 53.417 | 5 |
| S | 14 | --- | 1686.708 | 418.266 | --- | 4 |
| A | 15 | --- | 1757.746 | 331.234 | --- | 3 |
| L | 16 | --- | 1870.830 | 260.197 | --- | 2 |
| K | 17 | --- | --- | 147.113 | --- | 1 |

**Soleil Index**

| | | |
|---|---|---|
| **899.2987** | **not explained** | **100%** |

| | | | |
|---|---|---|---|
| **Run:** | 77662 | **Scans:** | 20462 - 20462 |
| **Search:** | 77662 | **Precursor Mass:** | 1036.83028 |
| **Mods:** | C=160.03 *+15.99 #+220.06 @+291.1 | **Charge:** | +3 |

**m/z Range:** 283.42279 to 1849.9783

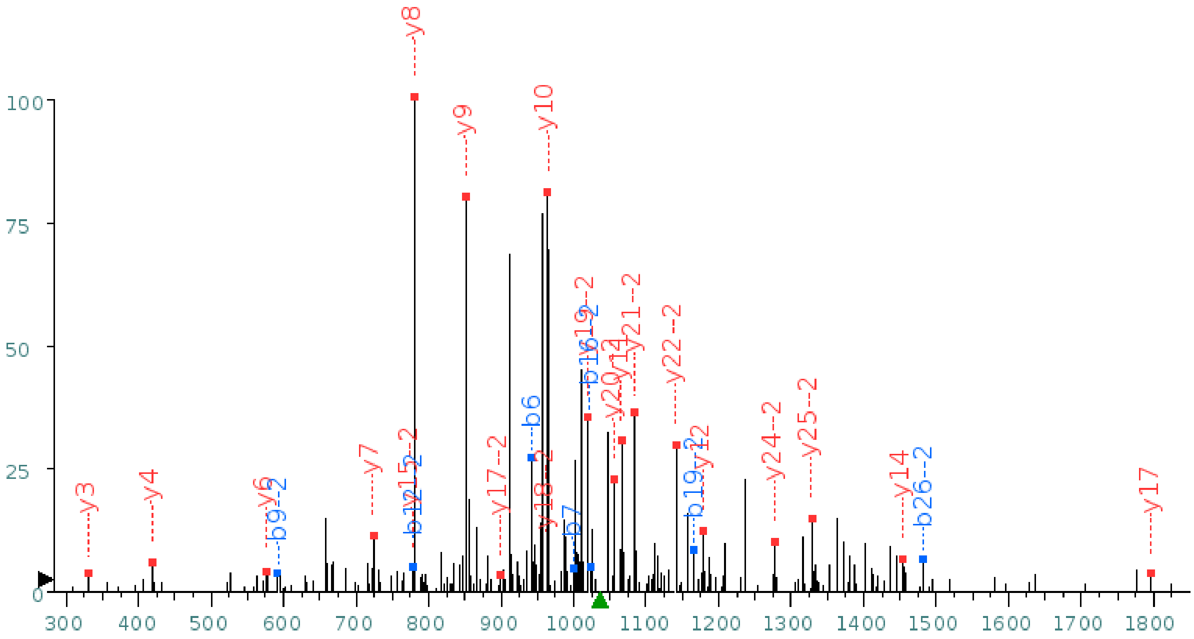

**Sequence**

S#M*VFQIGANKDQVM*ELTIAGM*GTSALK

**Predicted Fragmentation Pattern**

+1

| Seq | # | b: Δ Error | b | y | y: Δ Error | +1 |
|---|---|---|---|---|---|---|
| S# | 1 | --- | 308.098 | --- | --- | 27 |
| M* | 2 | --- | 455.133 | 2801.388 | --- | 26 |
| V | 3 | --- | 554.201 | 2654.353 | --- | 25 |
| F | 4 | --- | 701.270 | 2555.284 | --- | 24 |
| Q | 5 | --- | 829.328 | 2408.216 | --- | 23 |
| I | 6 | 256.535 | **942.412** | 2280.157 | --- | 22 |
| G | 7 | -272.789 | **999.434** | 2167.073 | --- | 21 |
| A | 8 | --- | 1070.471 | 2110.052 | --- | 20 |
| N | 9 | --- | 1184.514 | 2039.015 | --- | 19 |
| K | 10 | --- | 1312.609 | 1924.972 | --- | 18 |
| D | 11 | --- | 1427.636 | **1796.877** | 116.409 | 17 |
| Q | 12 | --- | 1555.694 | 1681.850 | --- | 16 |
| V | 13 | --- | 1654.763 | 1553.791 | --- | 15 |
| M* | 14 | --- | 1801.798 | **1454.723** | -30.732 | 14 |
| E | 15 | --- | 1930.841 | 1307.688 | --- | 13 |
| L | 16 | --- | 2043.925 | **1178.645** | 191.843 | 12 |
| T | 17 | --- | 2144.973 | **1065.561** | 112.180 | 11 |
| I | 18 | --- | 2258.057 | **964.513** | -995.775 | 10 |

| | | | | | | |
|---|---|---|---|---|---|---|
| A | 19 | --- | 2329.094 | **851.429** | **194.742** | **9** |
| G | 20 | --- | 2386.115 | **780.392** | **165.562** | **8** |
| M* | 21 | --- | 2533.151 | **723.371** | **188.028** | **7** |
| G | 22 | --- | 2590.172 | **576.335** | **312.087** | **6** |
| T | 23 | --- | 2691.220 | 519.314 | --- | **5** |
| S | 24 | --- | 2778.252 | **418.266** | **177.803** | **4** |
| A | 25 | --- | 2849.289 | **331.234** | **501.216** | **3** |
| L | 26 | --- | 2962.373 | 260.197 | --- | **2** |
| K | 27 | --- | --- | 147.113 | --- | **1** |

**+2**

| Seq | # | b: Δ Error | b | y | y: Δ Error | +1 |
|---|---|---|---|---|---|---|
| S# | 1 | --- | 154.552 | --- | --- | 27 |
| M* | 2 | --- | 228.070 | 1401.198 | --- | 26 |
| V | 3 | --- | 277.604 | **1327.680** | 415.879 | 25 |
| F | 4 | --- | 351.139 | **1278.146** | 340.396 | 24 |
| Q | 5 | --- | 415.168 | 1204.612 | --- | 23 |
| I | 6 | --- | 471.710 | **1140.582** | 357.816 | 22 |
| G | 7 | --- | 500.221 | **1084.040** | 175.167 | 21 |
| A | 8 | --- | 535.739 | **1055.530** | -32.065 | 20 |
| N | 9 | -1066.451 | **592.761** | **1020.011** | -150.004 | 19 |
| K | 10 | --- | 656.808 | **962.990** | 585.499 | 18 |
| D | 11 | --- | 714.322 | **898.942** | 135.445 | 17 |
| Q | 12 | -208.837 | **778.351** | 841.429 | --- | 16 |
| V | 13 | --- | 827.885 | **777.399** | 1013.952 | 15 |
| M* | 14 | --- | 901.403 | 727.865 | --- | 14 |
| E | 15 | --- | 965.924 | 654.347 | --- | 13 |
| L | 16 | 669.996 | **1022.466** | 589.826 | --- | 12 |
| T | 17 | --- | 1072.990 | 533.284 | --- | 11 |
| I | 18 | --- | 1129.532 | 482.760 | --- | 10 |
| A | 19 | 81.724 | **1165.051** | 426.218 | --- | 9 |
| G | 20 | --- | 1193.561 | 390.700 | --- | 8 |
| M* | 21 | --- | 1267.079 | 362.189 | --- | 7 |
| G | 22 | --- | 1295.590 | 288.671 | --- | 6 |
| T | 23 | --- | 1346.114 | 260.160 | --- | 5 |
| S | 24 | --- | 1389.630 | 209.637 | --- | 4 |
| A | 25 | --- | 1425.148 | 166.121 | --- | 3 |
| L | 26 | 243.423 | **1481.690** | 130.602 | --- | 2 |
| K | 27 | --- | --- | 74.060 | --- | 1 |

---

**Soleil Index**

| | | |
|---|---|---|
| 781.5055 | not explained | 32% |
| 852.5532 | not explained | 23% |
| 856.2280 | not explained | 19% |
| 912.7147 | not explained | 68% |
| 957.8252 | not explained | 77% |
| 1001.8660 | y2-19 frag loss of water | 27% |

| | | |
|---|---|---|
| 1010.9290 | not explained | 45% |
| 1046.3788 | not explained | 31% |
| 1047.3591 | y1-11 frag loss of water | 32% |
| 1156.9332 | not explained | 16% |
| 1236.5889 | not explained | 23% |
| 1363.0342 | not explained | 15% |

| Run: | 77662 | Scans: | 21050 - 21050 | m/z Range: | 261.50168 to 2000 |
|---|---|---|---|---|---|
| Search: | 77662 | Precursor Mass: | 1000.97545 | | |
| Mods: | C=160.03 *+15.99 #+220.06 @+291.1 | Charge: | +2 | | |

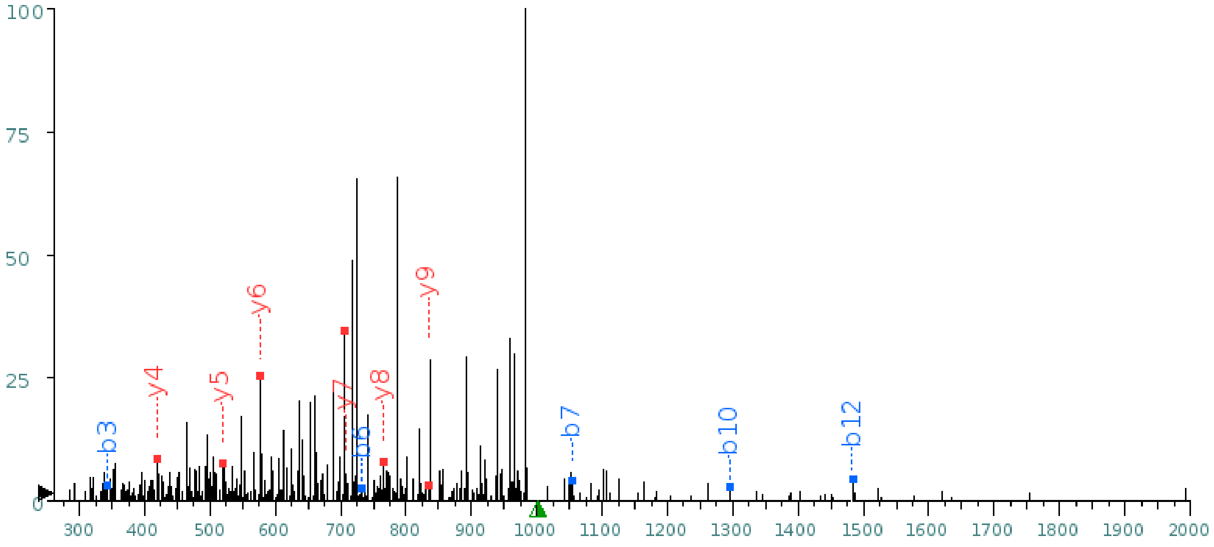

### Sequence

DQVM*ELT#IAGMGTSALK

### Predicted Fragmentation Pattern

| Seq | # | b: Δ Error | b | y | y: Δ Error | +1 |
|---|---|---|---|---|---|---|
| D | 1 | --- | 116.034 | --- | --- | 17 |
| Q | 2 | --- | 244.093 | 1885.913 | --- | 16 |
| V | 3 | 634.603 | **343.161** | 1757.855 | --- | 15 |
| M* | 4 | --- | 490.197 | 1658.786 | --- | 14 |
| E | 5 | --- | 619.239 | 1511.751 | --- | 13 |
| L | 6 | 251.604 | **732.323** | 1382.708 | --- | 12 |
| T# | 7 | -92.755 | **1053.429** | 1269.624 | --- | 11 |
| I | 8 | --- | 1166.513 | 948.518 | --- | 10 |
| A | 9 | --- | 1237.550 | **835.434** | 177.925 | 9 |
| G | 10 | 64.126 | **1294.572** | **764.397** | -0.967 | 8 |
| M | 11 | --- | 1425.612 | **707.376** | -1248.052 | 7 |
| G | 12 | 300.725 | **1482.634** | **576.335** | 38.637 | 6 |
| T | 13 | --- | 1583.682 | **519.314** | 274.301 | 5 |
| S | 14 | --- | 1670.714 | **418.266** | 31.618 | 4 |
| A | 15 | --- | 1741.751 | 331.234 | --- | 3 |
| L | 16 | --- | 1854.835 | 260.197 | --- | 2 |
| K | 17 | --- | --- | 147.113 | --- | 1 |

Soleil Index

| | | |
|---|---|---|
| 465.4417 | not explained | 16% |
| 548.3816 | not explained | 17% |
| 635.4576 | y1-11 frag doubly charged | 20% |
| 653.4886 | not explained | 20% |
| 659.4662 | not explained | 21% |
| 688.6488 | y1-7 frag loss of water | 24% |
| 716.5791 | not explained | 49% |
| 724.3345 | not explained | 65% |
| 742.5707 | not explained | 17% |
| 787.4387 | not explained | 66% |
| 837.6167 | not explained | 28% |
| 891.0585 | not explained | 29% |
| 939.5891 | not explained | 27% |
| 957.6686 | not explained | 33% |
| 965.5592 | not explained | 30% |
| 983.6110 | precursor loss of two waters | 100% |

| | | | | | |
|---|---|---|---|---|---|
| **Run:** | 77662 | **Scans:** | 23175 - 23175 | **m/z Range:** | 301.61353 to 1795.9007( |
| **Search:** | 77662 | **Precursor Mass:** | 1099.52062 | | |
| **Mods:** | C=160.03 *+15.99 #+220.06 @+291.1 | | | | |

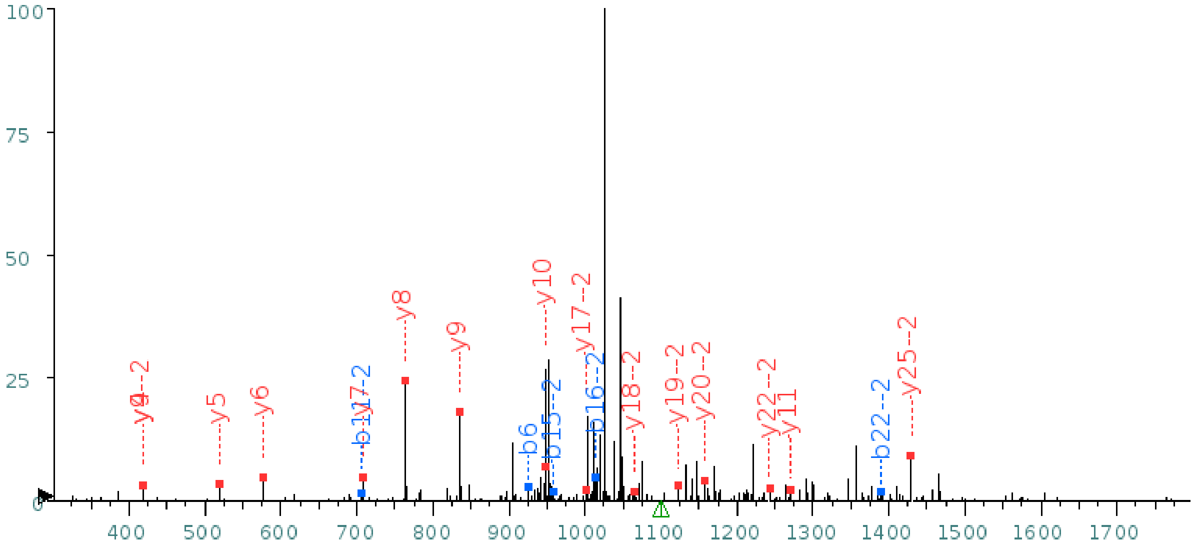

**Sequence**

S#MVFQIGANKDQVM*ELT#IAGMGTSALK

**Predicted Fragmentation Pattern**

+1

| Seq | # | b: Δ Error | b | y | y: Δ Error | +1 |
|---|---|---|---|---|---|---|
| S# | 1 | --- | 308.098 | --- | --- | 27 |
| M | 2 | --- | 439.138 | 2989.457 | --- | 26 |
| V | 3 | --- | 538.207 | 2858.416 | --- | 25 |
| F | 4 | --- | 685.275 | 2759.348 | --- | 24 |
| Q | 5 | --- | 813.333 | 2612.279 | --- | 23 |
| I | 6 | -104.717 | **926.418** | 2484.221 | --- | 22 |
| G | 7 | --- | 983.439 | 2371.137 | --- | 21 |
| A | 8 | --- | 1054.476 | 2314.115 | --- | 20 |
| N | 9 | --- | 1168.519 | 2243.078 | --- | 19 |
| K | 10 | --- | 1296.614 | 2129.035 | --- | 18 |
| D | 11 | --- | 1411.641 | 2000.940 | --- | 17 |
| Q | 12 | --- | 1539.700 | 1885.913 | --- | 16 |
| V | 13 | --- | 1638.768 | 1757.855 | --- | 15 |
| M* | 14 | --- | 1785.803 | 1658.786 | --- | 14 |
| E | 15 | --- | 1914.846 | 1511.751 | --- | 13 |
| L | 16 | --- | 2027.930 | 1382.708 | --- | 12 |
| T# | 17 | --- | 2349.036 | **1269.624** | 621.979 | 11 |
| I | 18 | --- | 2462.120 | **948.518** | 41.086 | 10 |
| A | 19 | --- | 2533.157 | **835.434** | 136.441 | 9 |
| G | 20 | --- | 2590.179 | **764.397** | 84.782 | 8 |

| | | | | | | |
|---|---|---|---|---|---|---|
| M | 21 | --- | 2721.219 | **707.376** | **286.687** | 7 |
| G | 22 | --- | 2778.241 | **576.335** | **270.915** | 6 |
| T | 23 | --- | 2879.288 | **519.314** | **311.889** | 5 |
| S | 24 | --- | 2966.320 | **418.266** | **603.861** | 4 |
| A | 25 | --- | 3037.357 | 331.234 | --- | 3 |
| L | 26 | --- | 3150.441 | 260.197 | --- | 2 |
| K | 27 | --- | --- | 147.113 | --- | 1 |

**+2**

| Seq | # | b: Δ Error | b | y | y: Δ Error | +1 |
|---|---|---|---|---|---|---|
| S# | 1 | --- | 154.552 | --- | --- | 27 |
| M | 2 | --- | 220.073 | 1495.232 | --- | 26 |
| V | 3 | --- | 269.607 | **1429.712** | **-179.119** | 25 |
| F | 4 | --- | 343.141 | 1380.178 | --- | 24 |
| Q | 5 | --- | 407.170 | 1306.643 | --- | 23 |
| I | 6 | --- | 463.712 | **1242.614** | **335.943** | 22 |
| G | 7 | --- | 492.223 | 1186.072 | --- | 21 |
| A | 8 | --- | 527.742 | **1157.561** | **-250.294** | 20 |
| N | 9 | --- | 584.763 | **1122.043** | **192.977** | 19 |
| K | 10 | --- | 648.811 | **1065.021** | **795.155** | 18 |
| D | 11 | **372.262** | **706.324** | **1000.974** | **957.492** | 17 |
| Q | 12 | --- | 770.353 | 943.460 | --- | 16 |
| V | 13 | --- | 819.888 | 879.431 | --- | 15 |
| M* | 14 | --- | 893.405 | 829.897 | --- | 14 |
| E | 15 | **-439.428** | **957.927** | 756.379 | --- | 13 |
| L | 16 | **12.496** | **1014.469** | 691.858 | --- | 12 |
| T# | 17 | --- | 1175.022 | 635.316 | --- | 11 |
| I | 18 | --- | 1231.564 | 474.763 | --- | 10 |
| A | 19 | --- | 1267.082 | **418.221** | **712.017** | 9 |
| G | 20 | --- | 1295.593 | 382.702 | --- | 8 |
| M | 21 | --- | 1361.113 | 354.191 | --- | 7 |
| G | 22 | **149.643** | **1389.624** | 288.671 | --- | 6 |
| T | 23 | --- | 1440.148 | 260.160 | --- | 5 |
| S | 24 | --- | 1483.664 | 209.637 | --- | 4 |
| A | 25 | --- | 1519.182 | 166.121 | --- | 3 |
| L | 26 | --- | 1575.724 | 130.602 | --- | 2 |
| K | 27 | --- | --- | 74.060 | --- | 1 |

**Soleil Index**

| | | |
|---|---|---|
| 947.2838 | not explained | 27% |
| 953.3064 | not explained | 29% |
| 1002.5751 | not explained | 17% |
| 1011.7205 | not explained | 16% |
| 1026.4431 | not explained | 100% |
| 1047.3733 | y2-18 frag loss of water/ y2-18 frag loss of ammonia | 41% |

| Run: | 77662 | Scans: | 23214 - 23214 | m/z Range: | 274.35818 | to | 1937.1983 |
|---|---|---|---|---|---|---|---|
| Search: | 77662 | Precursor Mass: | 1026.16773 | | | | |
| Mods: | C=160.03 *+15.99 #+220.06 @+291.1 | Charge: | +3 | | | | |

#### Sequence

S#MVFQIGANKDQVM*ELTIAGMGTSALK

#### Predicted Fragmentation Pattern

+1

| Seq | # | b: Δ Error | b | y | y: Δ Error | +1 |
|---|---|---|---|---|---|---|
| S# | 1 | -90.472 | **308.098** | --- | --- | 27 |
| M | 2 | --- | 439.138 | 2769.398 | --- | 26 |
| V | 3 | --- | 538.207 | 2638.358 | --- | 25 |
| F | 4 | -1258.244 | **685.275** | 2539.289 | --- | 24 |
| Q | 5 | --- | 813.333 | 2392.221 | --- | 23 |
| I | 6 | --- | 926.418 | 2264.162 | --- | 22 |
| G | 7 | --- | 983.439 | 2151.078 | --- | 21 |
| A | 8 | --- | 1054.476 | 2094.057 | --- | 20 |
| N | 9 | --- | 1168.519 | 2023.020 | --- | 19 |
| K | 10 | --- | 1296.614 | 1908.977 | --- | 18 |
| D | 11 | --- | 1411.641 | 1780.882 | --- | 17 |
| Q | 12 | --- | 1539.700 | 1665.855 | --- | 16 |
| V | 13 | --- | 1638.768 | 1537.796 | --- | 15 |
| M* | 14 | --- | 1785.803 | 1438.728 | --- | 14 |
| E | 15 | --- | 1914.846 | 1291.693 | --- | 13 |
| L | 16 | --- | 2027.930 | 1162.650 | --- | 12 |
| T | 17 | --- | 2128.978 | **1049.566** | 113.696 | 11 |
| I | 18 | --- | 2242.062 | **948.518** | 72.484 | 10 |
| A | 19 | --- | 2313.099 | **835.434** | 58.722 | 9 |
| G | 20 | --- | 2370.120 | **764.397** | 63.546 | 8 |
| M | 21 | --- | 2501.161 | **707.376** | 184.403 | 7 |
| G | 22 | --- | 2558.182 | **576.335** | 160.824 | 6 |
| T | 23 | --- | 2659.230 | 519.314 | --- | 5 |
| S | 24 | --- | 2746.262 | **418.266** | 39.716 | 4 |
| A | 25 | --- | 2817.299 | 331.234 | --- | 3 |
| L | 26 | --- | 2930.383 | 260.197 | --- | 2 |
| K | 27 | --- | --- | 147.113 | --- | 1 |

+2

| Seq | # | b: Δ Error | b | y | y: Δ Error | +1 |
|---|---|---|---|---|---|---|
| S# | 1 | --- | 154.552 | --- | --- | 27 |
| M | 2 | --- | 220.073 | 1385.203 | --- | 26 |
| V | 3 | --- | 269.607 | **1319.683** | 559.985 | 25 |
| F | 4 | --- | 343.141 | **1270.148** | 230.654 | 24 |
| Q | 5 | --- | 407.170 | **1196.614** | 24.457 | 23 |
| I | 6 | --- | 463.712 | **1132.585** | -100.496 | 22 |
| G | 7 | --- | 492.223 | **1076.043** | -126.959 | 21 |
| A | 8 | --- | 527.742 | **1047.532** | -177.168 | 20 |
| N | 9 | --- | 584.763 | **1012.014** | -63.820 | 19 |
| K | 10 | --- | 648.811 | **954.992** | 454.815 | 18 |
| D | 11 | --- | 706.324 | **890.945** | -456.221 | 17 |
| Q | 12 | --- | 770.353 | **833.431** | 294.473 | 16 |

| | | | | | | |
|---|---|---|---|---|---|---|
| V | 13 | --- | 819.888 | 769.402 | --- | 15 |
| M* | 14 | 893.563 | **893.405** | 719.868 | --- | 14 |
| E | 15 | 20.793 | **957.927** | 646.350 | --- | 13 |
| L | 16 | --- | 1014.469 | 581.829 | --- | 12 |
| T | 17 | --- | 1064.992 | 525.287 | --- | 11 |
| I | 18 | --- | 1121.535 | **474.763** | 1655.368 | 10 |
| A | 19 | --- | 1157.053 | **418.221** | 147.933 | 9 |
| G | 20 | 653.630 | **1185.564** | 382.702 | --- | 8 |
| M | 21 | --- | 1251.084 | 354.191 | --- | 7 |
| G | 22 | --- | 1279.595 | 288.671 | --- | 6 |
| T | 23 | --- | 1330.119 | 260.160 | --- | 5 |
| S | 24 | --- | 1373.635 | 209.637 | --- | 4 |
| A | 25 | --- | 1409.153 | 166.121 | --- | 3 |
| L | 26 | --- | 1465.695 | 130.602 | --- | 2 |
| K | 27 | --- | --- | 74.060 | --- | 1 |

**Soleil Index**

| | | |
|---|---|---|
| 765.5930 | not explained | 20% |
| 836.5511 | not explained | 15% |
| 848.1848 | not explained | 20% |
| 904.3293 | not explained | 29% |
| 923.3197 | not explained | 18% |
| 946.1885 | not explained | 65% |
| 947.1870 | not explained | 77% |
| 993.9470 | y2-19 frag loss of water | 18% |
| 1002.9022 | not explained | 77% |
| 1038.8167 | not explained | 21% |
| 1067.2983 | not explained | 23% |
| 1102.7030 | b2-18 frag loss of water | 17% |
| 1104.1760 | b2-18 frag loss of water/ b2-18 frag loss of ammonia | 36% |
| 1133.6095 | not explained | 16% |
| 1355.9689 | b2-24 frag loss of water/ b2-24 frag loss of ammonia | 35% |
| 1429.3108 | not explained | 28% |

| | | | | | |
|---|---|---|---|---|---|
| **Run:** | 77662 | **Scans:** | 29538 - 29538 | **m/z Range:** | 271.34949 to 2000 |
| **Search:** | 77662 | **Precursor Mass:** | 1020.83445 | | |
| **Mods:** | C=160.03 *+15.99 #+220.06 @+291.1 | **Charge:** | +3 | | |

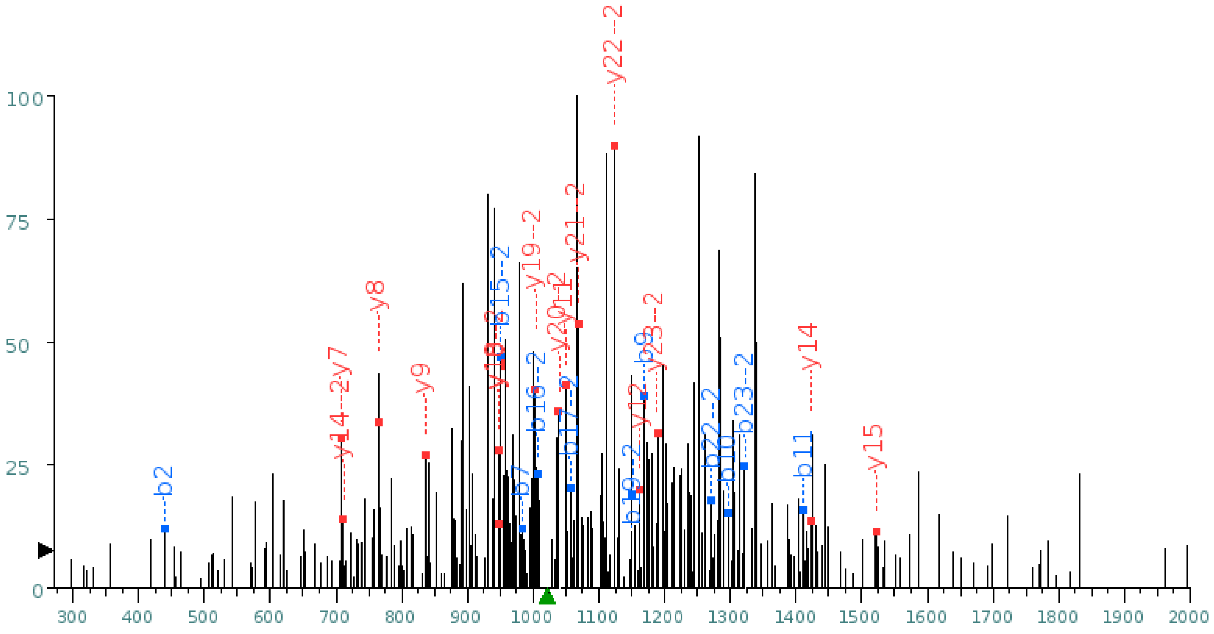

**Sequence**

S#MVFQIGANKDQVMELTIAGMGTSALK

**Predicted Fragmentation Pattern**

| Seq | # | b: Δ Error | b | y | y: Δ Error | +1 |
|---|---|---|---|---|---|---|
| S# | 1 | --- | 308.098 | --- | --- | 27 |
| M | 2 | 512.681 | **439.138** | 2753.403 | --- | 26 |
| V | 3 | --- | 538.207 | 2622.363 | --- | 25 |
| F | 4 | --- | 685.275 | 2523.295 | --- | 24 |
| Q | 5 | --- | 813.333 | 2376.226 | --- | 23 |
| I | 6 | --- | 926.418 | 2248.168 | --- | 22 |
| G | 7 | -381.519 | **983.439** | 2135.084 | --- | 21 |
| A | 8 | --- | 1054.476 | 2078.062 | --- | 20 |
| N | 9 | 166.369 | **1168.519** | 2007.025 | --- | 19 |
| K | 10 | -504.978 | **1296.614** | 1892.982 | --- | 18 |
| D | 11 | -570.693 | **1411.641** | 1764.887 | --- | 17 |
| Q | 12 | --- | 1539.700 | 1649.860 | --- | 16 |
| V | 13 | --- | 1638.768 | **1521.802** | 129.129 | 15 |
| M | 14 | --- | 1769.808 | **1422.733** | 621.615 | 14 |
| E | 15 | --- | 1898.851 | 1291.693 | --- | 13 |
| L | 16 | --- | 2011.935 | **1162.650** | 165.444 | 12 |
| T | 17 | --- | 2112.983 | **1049.566** | -80.179 | 11 |
| I | 18 | --- | 2226.067 | **948.518** | -535.628 | 10 |

| | | | | | | |
|---|---|---|---|---|---|---|
| A | 19 | --- | 2297.104 | **835.434** | **241.604** | **9** |
| G | 20 | --- | 2354.125 | **764.397** | **302.362** | **8** |
| M | 21 | --- | 2485.166 | **707.376** | **464.299** | **7** |
| G | 22 | --- | 2542.187 | 576.335 | --- | 6 |
| T | 23 | --- | 2643.235 | 519.314 | --- | 5 |
| S | 24 | --- | 2730.267 | 418.266 | --- | 4 |
| A | 25 | --- | 2801.304 | 331.234 | --- | 3 |
| L | 26 | --- | 2914.388 | 260.197 | --- | 2 |
| K | 27 | --- | --- | 147.113 | --- | 1 |

**+2**

| Seq | # | b: Δ Error | b | y | y: Δ Error | +1 |
|---|---|---|---|---|---|---|
| S# | 1 | --- | 154.552 | --- | --- | 27 |
| M | 2 | --- | 220.073 | 1377.205 | --- | 26 |
| V | 3 | --- | 269.607 | 1311.685 | --- | 25 |
| F | 4 | --- | 343.141 | 1262.151 | --- | 24 |
| Q | 5 | --- | 407.170 | **1188.617** | **746.982** | 23 |
| I | 6 | --- | 463.712 | **1124.587** | **-567.604** | 22 |
| G | 7 | --- | 492.223 | **1068.045** | **213.703** | 21 |
| A | 8 | --- | 527.742 | **1039.535** | **-293.763** | 20 |
| N | 9 | --- | 584.763 | **1004.016** | **-201.246** | 19 |
| K | 10 | --- | 648.811 | **946.995** | **46.395** | 18 |
| D | 11 | --- | 706.324 | 882.947 | --- | 17 |
| Q | 12 | --- | 770.353 | 825.434 | --- | 16 |
| V | 13 | --- | 819.888 | 761.404 | --- | 15 |
| M | 14 | --- | 885.408 | **711.870** | **-678.584** | 14 |
| E | 15 | **-176.295** | **949.929** | 646.350 | --- | 13 |
| L | 16 | **-43.420** | **1006.471** | 581.829 | --- | 12 |
| T | 17 | **-102.364** | **1056.995** | 525.287 | --- | 11 |
| I | 18 | --- | 1113.537 | 474.763 | --- | 10 |
| A | 19 | **766.790** | **1149.056** | 418.221 | --- | 9 |
| G | 20 | --- | 1177.566 | 382.702 | --- | 8 |
| M | 21 | --- | 1243.087 | 354.191 | --- | 7 |
| G | 22 | **-626.130** | **1271.597** | 288.671 | --- | 6 |
| T | 23 | **-123.881** | **1322.121** | 260.160 | --- | 5 |
| S | 24 | --- | 1365.637 | 209.637 | --- | 4 |
| A | 25 | --- | 1401.156 | 166.121 | --- | 3 |
| L | 26 | --- | 1457.698 | 130.602 | --- | 2 |
| K | 27 | --- | --- | 74.060 | --- | 1 |

Soleil Index

| | | |
|---|---|---|
| **541.4449** | **not explained** | **19%** |
| **577.5176** | **not explained** | **18%** |
| **603.5410** | **not explained** | **23%** |
| **619.5950** | **not explained** | **18%** |
| **743.9552** | **y2-15 frag loss of water/ y2-15 frag loss of ammonia** | **18%** |
| **757.5702** | **not explained** | **16%** |

| | | |
|---|---|---|
| 765.4822 | not explained | 43% |
| 766.4877 | not explained | 16% |
| 784.3429 | not explained | 22% |
| 840.7681 | not explained | 26% |
| 853.5100 | not explained | 19% |
| 875.6691 | not explained | 18% |
| 877.2913 | not explained | 32% |
| 891.8563 | not explained | 30% |
| 892.6193 | not explained | 62% |
| 897.7035 | not explained | 16% |
| 902.6185 | not explained | 41% |
| 906.8203 | not explained | 23% |
| 930.6331 | y1-10 frag loss of water/ y1-10 frag loss of ammonia/ y2-18 frag loss of ammonia | 80% |
| 938.1492 | not explained | 18% |
| 942.0612 | not explained | 77% |
| 955.7772 | not explained | 23% |
| 957.7711 | not explained | 51% |
| 960.6813 | not explained | 24% |
| 961.6669 | not explained | 23% |
| 963.0978 | not explained | 22% |
| 968.7513 | not explained | 31% |
| 971.8722 | not explained | 22% |
| 979.2302 | not explained | 66% |

| | | |
|---|---|---|
| 996.1238 | not explained | 16% |
| 997.7936 | not explained | 22% |
| 1001.2252 | not explained | 48% |
| 1002.9781 | y1-19 frag doubly charged | 15% |
| 1009.3223 | precursor loss of two waters | 18% |
| 1034.8101 | not explained | 30% |
| 1066.4205 | not explained | 100% |
| 1070.2280 | not explained | 30% |
| 1088.5334 | not explained | 16% |
| 1101.7435 | not explained | 19% |
| 1105.9025 | y2-22 frag loss of water | 27% |
| 1110.8718 | not explained | 88% |
| 1112.0732 | b1-18 frag doubly charged | 52% |
| 1130.0574 | b2-19 frag loss of water | 24% |
| 1132.0450 | b2-19 frag loss of water/ b2-19 frag loss of ammonia | 23% |
| 1151.0533 | b1-9 frag loss of water/ b1-9 frag loss of ammonia | 43% |
| 1161.2745 | b2-20 frag loss of ammonia | 19% |
| 1173.0707 | not explained | 29% |
| 1175.7629 | not explained | 26% |
| 1181.5756 | not explained | 27% |
| 1198.2476 | not explained | 45% |
| 1201.0100 | not explained | 29% |
| 1204.0667 | not explained | 17% |

| | | |
|---|---|---|
| 1212.8402 | not explained | 21% |
| 1213.4778 | not explained | 24% |
| 1222.3573 | not explained | 23% |
| 1225.7008 | b2-21 frag loss of water/ b2-21 frag loss of ammonia | 24% |
| 1236.5712 | not explained | 29% |
| 1237.7166 | not explained | 18% |
| 1238.6295 | not explained | 20% |
| 1239.2704 | not explained | 17% |
| 1239.9962 | not explained | 19% |
| 1244.8525 | y2-24 frag loss of water/ y2-24 frag loss of ammonia | 42% |
| 1251.0990 | not explained | 34% |
| 1252.5974 | b2-22 frag loss of water | 92% |
| 1260.5792 | not explained | 31% |
| 1283.2111 | not explained | 68% |
| 1284.3500 | not explained | 51% |
| 1289.5111 | not explained | 20% |
| 1304.1971 | b2-23 frag loss of water/ b2-23 frag loss of ammonia | 34% |
| 1307.7067 | not explained | 19% |
| 1312.9132 | not explained | 31% |
| 1337.9382 | not explained | 84% |
| 1339.0255 | not explained | 50% |
| 1363.3114 | not explained | 17% |
| 1388.5201 | not explained | 17% |

| | | |
|---|---|---|
| 1402.9209 | not explained | 18% |
| 1424.2264 | not explained | 31% |
| 1445.4080 | not explained | 25% |
| 1587.0411 | not explained | 24% |
| 1831.0717 | not explained | 23% |

| | | | |
|---|---|---|---|
| Run: | 77662 | Scans: | 33441 - 33441 |
| Search: | 77662 | Precursor Mass: | 1026.16812 |
| Mods: | C=160.03 *+15.99 #+220.06 @+291.1 | Charge: | +3 |

m/z Range: 270.17297 to 2000

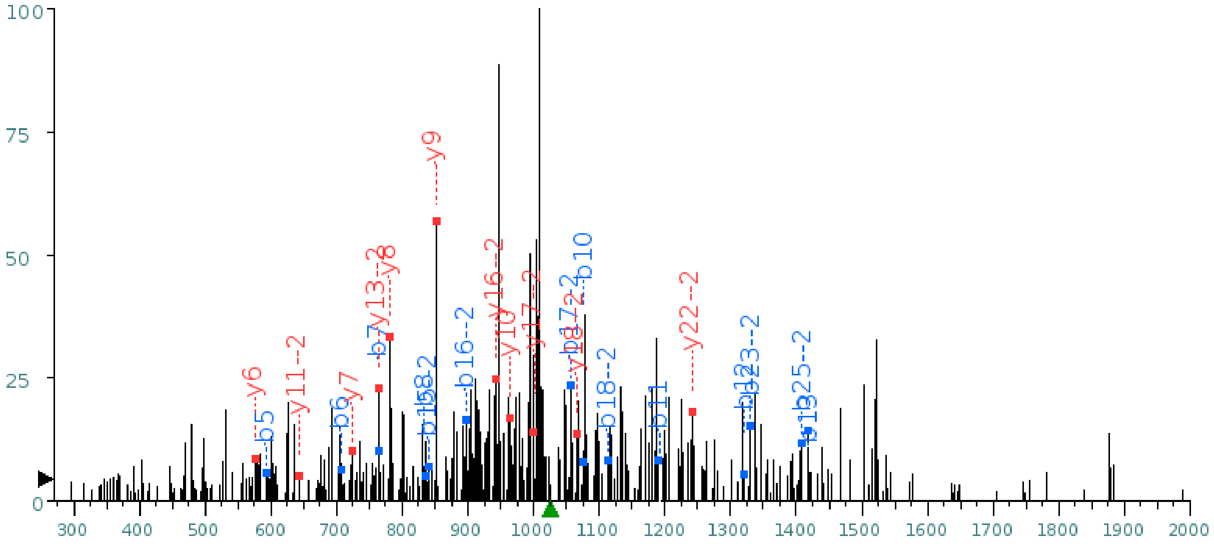

### Sequence

SMVFQIGANKDQVMELT#IAGM*GTSALK

### Predicted Fragmentation Pattern

+1

| Seq | # | b: Δ Error | b | y | y: Δ Error | +1 |
|---|---|---|---|---|---|---|
| S | 1 | --- | 88.039 | --- | --- | 27 |
| M | 2 | --- | 219.080 | 2989.457 | --- | 26 |
| V | 3 | --- | 318.148 | 2858.416 | --- | 25 |
| F | 4 | --- | 465.217 | 2759.348 | --- | 24 |
| Q | 5 | 407.768 | **593.275** | 2612.279 | --- | 23 |
| I | 6 | 1401.293 | **706.359** | 2484.221 | --- | 22 |
| G | 7 | 448.359 | **763.381** | 2371.137 | --- | 21 |
| A | 8 | 312.107 | **834.418** | 2314.115 | --- | 20 |
| N | 9 | --- | 948.461 | 2243.078 | --- | 19 |
| K | 10 | 92.573 | **1076.556** | 2129.035 | --- | 18 |
| D | 11 | -808.446 | **1191.583** | 2000.940 | --- | 17 |
| Q | 12 | -91.132 | **1319.641** | 1885.913 | --- | 16 |
| V | 13 | 86.161 | **1418.710** | 1757.855 | --- | 15 |
| M | 14 | --- | 1549.750 | 1658.786 | --- | 14 |
| E | 15 | --- | 1678.793 | 1527.746 | --- | 13 |
| L | 16 | --- | 1791.877 | 1398.703 | --- | 12 |
| T# | 17 | --- | 2112.983 | 1285.619 | --- | 11 |
| I | 18 | --- | 2226.067 | **964.513** | -560.993 | 10 |

| | | | | | | |
|---|---|---|---|---|---|---|
| A | 19 | --- | 2297.104 | **851.429** | **285.165** | 9 |
| G | 20 | --- | 2354.125 | **780.392** | **155.398** | 8 |
| M* | 21 | --- | 2501.161 | **723.371** | **138.769** | 7 |
| G | 22 | --- | 2558.182 | **576.335** | **314.098** | 6 |
| T | 23 | --- | 2659.230 | 519.314 | --- | 5 |
| S | 24 | --- | 2746.262 | 418.266 | --- | 4 |
| A | 25 | --- | 2817.299 | 331.234 | --- | 3 |
| L | 26 | --- | 2930.383 | 260.197 | --- | 2 |
| K | 27 | --- | --- | 147.113 | --- | 1 |

**+2**

| Seq | # | b: Δ Error | b | y | y: Δ Error | +1 |
|---|---|---|---|---|---|---|
| S | 1 | --- | 44.523 | --- | --- | 27 |
| M | 2 | --- | 110.044 | 1495.232 | --- | 26 |
| V | 3 | --- | 159.578 | 1429.712 | --- | 25 |
| F | 4 | --- | 233.112 | 1380.178 | --- | 24 |
| Q | 5 | --- | 297.141 | 1306.643 | --- | 23 |
| I | 6 | --- | 353.683 | **1242.614** | **76.312** | 22 |
| G | 7 | --- | 382.194 | 1186.072 | --- | 21 |
| A | 8 | --- | 417.713 | 1157.561 | --- | 20 |
| N | 9 | --- | 474.734 | 1122.043 | --- | 19 |
| K | 10 | --- | 538.782 | **1065.021** | **651.861** | 18 |
| D | 11 | --- | 596.295 | **1000.974** | **-459.244** | 17 |
| Q | 12 | --- | 660.324 | **943.460** | **339.820** | 16 |
| V | 13 | --- | 709.858 | 879.431 | --- | 15 |
| M | 14 | --- | 775.379 | 829.897 | --- | 14 |
| E | 15 | **-232.722** | **839.900** | **764.377** | **100.007** | 13 |
| L | 16 | **678.876** | **896.442** | 699.855 | --- | 12 |
| T# | 17 | **483.053** | **1056.995** | **643.313** | **156.155** | 11 |
| I | 18 | **-540.475** | **1113.537** | 482.760 | --- | 10 |
| A | 19 | --- | 1149.056 | 426.218 | --- | 9 |
| G | 20 | --- | 1177.566 | 390.700 | --- | 8 |
| M* | 21 | --- | 1251.084 | 362.189 | --- | 7 |
| G | 22 | --- | 1279.595 | 288.671 | --- | 6 |
| T | 23 | **154.089** | **1330.119** | 260.160 | --- | 5 |
| S | 24 | --- | 1373.635 | 209.637 | --- | 4 |
| A | 25 | **-100.665** | **1409.153** | 166.121 | --- | 3 |
| L | 26 | --- | 1465.695 | 130.602 | --- | 2 |
| K | 27 | --- | --- | 74.060 | --- | 1 |

**Soleil Index**

| | | |
|---|---|---|
| 479.3036 | not explained | 16% |
| 530.6574 | not explained | 19% |
| 625.5980 | b2-21 frag doubly charged/ y2-11 frag loss of water/ y2-11 frag loss of ammonia | 20% |
| 635.0447 | not explained | 16% |
| 693.3398 | b2-13 frag loss of ammonia | 19% |
| 704.4109 | b2-25 frag doubly charged/ y1-7 frag loss of water | 15% |

| | | |
|---|---|---|
| 782.2419 | not explained | 19% |
| 799.7325 | not explained | 18% |
| 803.4866 | not explained | 17% |
| 831.6086 | not explained | 16% |
| 878.3478 | b2-16 frag loss of water/ y1-15 frag loss of water | 18% |
| 891.9592 | not explained | 15% |
| 905.1285 | not explained | 22% |
| 911.6952 | not explained | 25% |
| 915.0287 | not explained | 20% |
| 917.1128 | not explained | 18% |
| 932.5902 | not explained | 23% |
| 939.6682 | not explained | 21% |
| 947.2678 | y1-10 frag loss of water/ y1-10 frag loss of ammonia | 89% |
| 961.6427 | not explained | 21% |
| 974.2059 | not explained | 21% |
| 978.3494 | not explained | 22% |
| 993.2720 | not explained | 20% |
| 993.9496 | not explained | 18% |
| 994.7120 | not explained | 50% |
| 998.7812 | not explained | 30% |
| 999.8209 | y1-17 frag doubly charged | 17% |
| 1003.3032 | not explained | 18% |
| 1004.6951 | not explained | 53% |

| | | |
|---|---|---|
| 1007.6864 | not explained | 37% |
| 1008.5409 | not explained | 100% |
| 1009.8464 | not explained | 57% |
| 1011.7126 | not explained | 23% |
| 1014.7483 | precursor loss of two waters | 23% |
| 1047.5306 | y2-18 frag loss of water/ y2-18 frag loss of ammonia | 23% |
| 1049.8518 | not explained | 19% |
| 1067.7123 | not explained | 20% |
| 1078.6337 | not explained | 38% |
| 1096.8021 | b2-18 frag loss of ammonia | 18% |
| 1117.1666 | not explained | 15% |
| 1132.4513 | b2-19 frag loss of ammonia | 23% |
| 1134.6515 | not explained | 18% |
| 1170.3947 | not explained | 21% |
| 1179.3827 | not explained | 23% |
| 1187.8203 | not explained | 33% |
| 1205.7860 | not explained | 21% |
| 1224.7427 | y2-22 frag loss of water/ y2-22 frag loss of ammonia | 21% |
| 1316.9618 | not explained | 20% |
| 1337.8351 | not explained | 22% |
| 1345.4562 | not explained | 15% |
| 1466.7950 | not explained | 19% |
| 1503.4904 | not explained | 24% |

| | | |
|---|---|---|
| 1519.3165 | not explained | 20% |
| 1522.7472 | not explained | 33% |

**MS/MS data of trypsin digested peptides with site score A and site score B above 19 from *Clostridium botulinum Cb*Fla flagellin protein co-expressed with *Clostridium botulinum* Maf in BL21(DE3) cells. Results of dataset 3 (MS/MS results obtained in July 2019).**

| | | | | | |
|---|---|---|---|---|---|
| **Run:** | 79495 | **Scans:** | 9956 - 9956 | **m/z Range:** | 205.19757 to 1016.7053 |
| **Search:** | 79495 | **Precursor Mass:** | 679.3319 | | |
| **Mods:** | C=160.03 *+15.99 #+220.06 @+291.1 | **Charge:** | +2 | | |

**Sequence**

IASQT#QFNTK

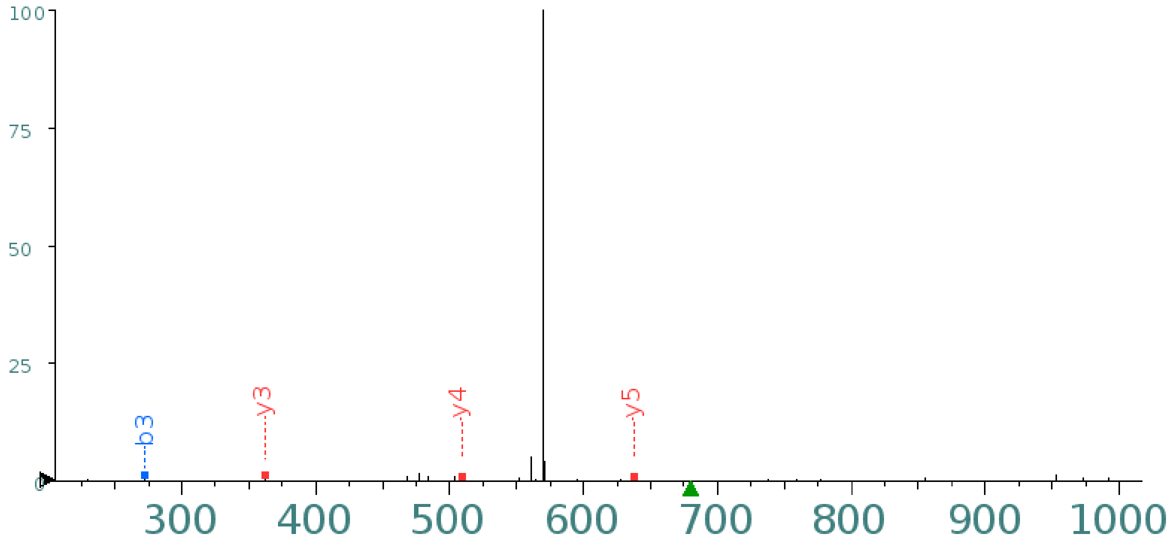

**Predicted Fragmentation Pattern**

| Seq | # | b: Δ Error | b | y | y: Δ Error | +1 |
|---|---|---|---|---|---|---|
| I | 1 | --- | 114.091 | --- | --- | 10 |
| A | 2 | --- | 185.128 | 1244.564 | --- | 9 |
| S | 3 | -380.793 | **272.160** | 1173.527 | --- | 8 |
| Q | 4 | --- | 400.219 | 1086.495 | --- | 7 |
| T# | 5 | --- | 721.325 | 958.436 | --- | 6 |
| Q | 6 | --- | 849.384 | **637.330** | -34.123 | 5 |
| F | 7 | --- | 996.452 | **509.272** | 302.097 | 4 |
| N | 8 | --- | 1110.495 | **362.203** | 424.521 | 3 |
| T | 9 | --- | 1211.543 | 248.160 | --- | 2 |
| K | 10 | --- | --- | 147.113 | --- | 1 |

**Soleil Index**

| | | |
|---|---|---|
| **569.3867** | **not explained** | **100%** |

| | | | | | |
|---|---|---|---|---|---|
| **Run:** | 79495 | **Scans:** | 11842 - 11842 | **m/z Range:** | 160.00352 to 957.53973 |
| **Search:** | 79495 | **Precursor Mass:** | 576.29651 | | |
| **Mods:** | C=160.03 *+15.99 #+220.06 @+291.1 | **Charge:** | +2 | | |

**Sequence**

VGS#INIGSGK

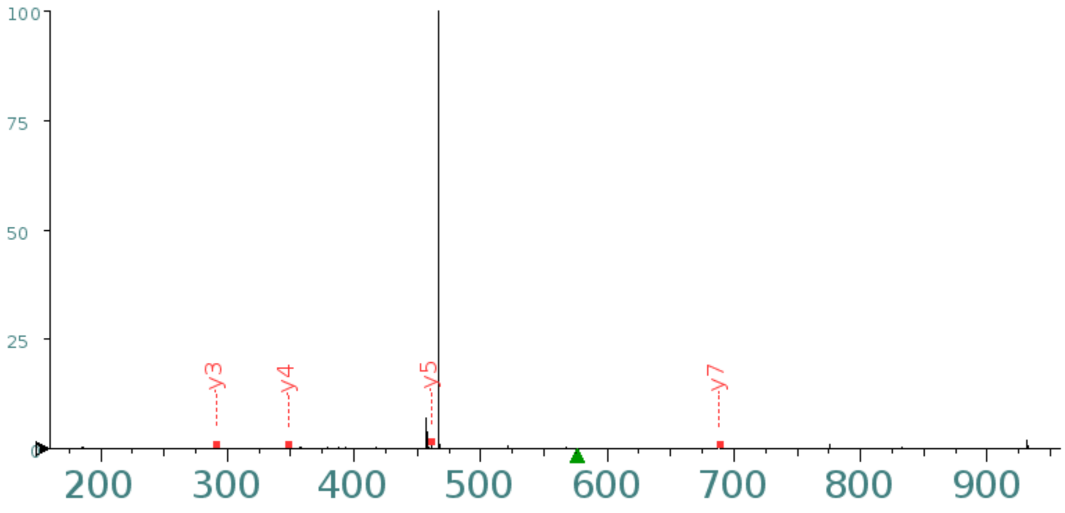

**Predicted Fragmentation Pattern**

| Seq | # | b: Δ Error | b | y | y: Δ Error | +1 |
|---|---|---|---|---|---|---|
| V | 1 | --- | 100.076 | --- | --- | 10 |
| G | 2 | --- | 157.097 | 1052.511 | --- | 9 |
| S# | 3 | --- | 464.187 | 995.489 | --- | 8 |
| I | 4 | --- | 577.272 | **688.399** | 373.555 | 7 |
| N | 5 | --- | 691.314 | 575.315 | --- | 6 |
| I | 6 | --- | 804.399 | **461.272** | 304.563 | 5 |
| G | 7 | --- | 861.420 | **348.188** | 148.230 | 4 |
| S | 8 | --- | 948.452 | **291.166** | 357.991 | 3 |
| G | 9 | --- | 1005.473 | 204.134 | --- | 2 |
| K | 10 | --- | --- | 147.113 | --- | 1 |

**Soleil Index**

| 466.3667 | not explained | 100% |
|---|---|---|

| | | | | | |
|---|---|---|---|---|---|
| **Run:** | 79495 | **Scans:** | 18885 - 18885 | **m/z Range:** | 306.36993 to 1766.0141 |
| **Search:** | 79495 | **Precursor Mass:** | 1008.97655 | | |
| **Mods:** | C=160.03 *+15.99 #+220.06 @+291.1 | **Charge:** | +2 | | |

**Sequence**

DQVM*ELT#IAGM*GTSALK

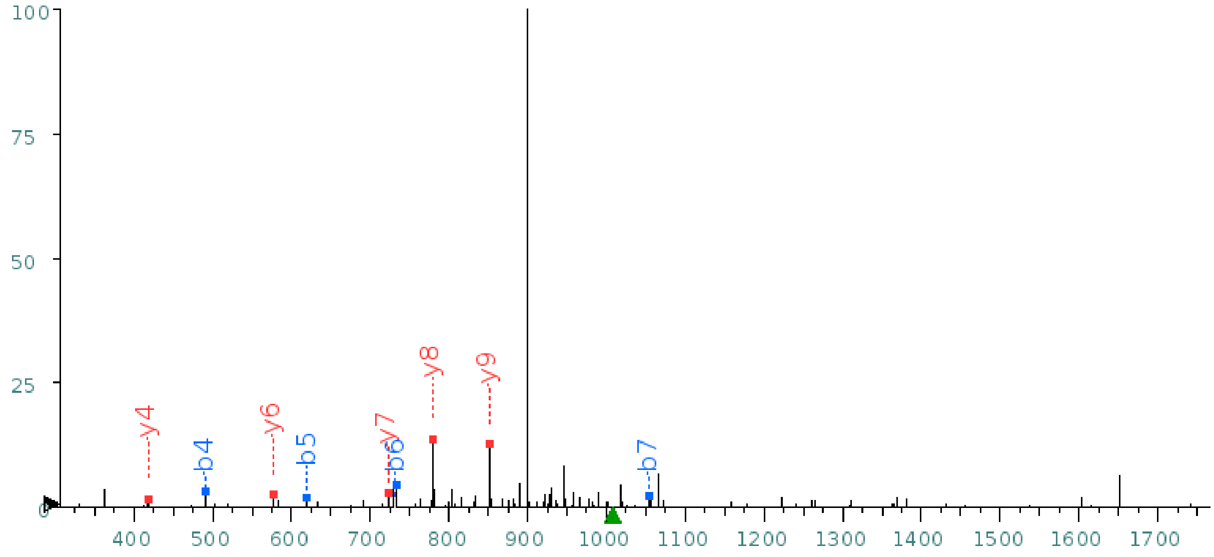

#### Predicted Fragmentation Pattern

| Seq | # | b: Δ Error | b | y | y: Δ Error | +1 |
|---|---|---|---|---|---|---|
| D | 1 | --- | 116.034 | --- | --- | 17 |
| Q | 2 | --- | 244.093 | 1901.908 | --- | 16 |
| V | 3 | --- | 343.161 | 1773.850 | --- | 15 |
| M* | 4 | 506.785 | **490.197** | 1674.781 | --- | 14 |
| E | 5 | 586.993 | **619.239** | 1527.746 | --- | 13 |
| L | 6 | 376.709 | **732.323** | 1398.703 | --- | 12 |
| T# | 7 | 329.060 | **1053.429** | 1285.619 | --- | 11 |
| I | 8 | --- | 1166.513 | 964.513 | --- | 10 |
| A | 9 | --- | 1237.550 | **851.429** | 165.218 | 9 |
| G | 10 | --- | 1294.572 | **780.392** | 126.780 | 8 |
| M* | 11 | --- | 1441.607 | **723.371** | 1346.756 | 7 |
| G | 12 | --- | 1498.629 | **576.335** | 125.675 | 6 |
| T | 13 | --- | 1599.676 | 519.314 | --- | 5 |
| S | 14 | --- | 1686.708 | **418.266** | 296.602 | 4 |
| A | 15 | --- | 1757.746 | 331.234 | --- | 3 |
| L | 16 | --- | 1870.830 | 260.197 | --- | 2 |
| K | 17 | --- | --- | 147.113 | --- | 1 |

#### Soleil Index

| | | |
|---|---|---|
| 898.9457 | not explained | 30% |
| 899.6405 | not explained | 100% |

| | | | | | |
|---|---|---|---|---|---|
| **Run:** | 79495 | **Scans:** | 19253 - 19253 | **m/z Range:** | 203.29776 to 1736.0164 |
| **Search:** | 79495 | **Precursor Mass:** | 783.38282 | | |
| **Mods:** | C=160.03 *+15.99 #+220.06 @+291.1 | **Charge:** | +3 | | |

#### Sequence

NLLNGS#AKS#M*VFQIGANK

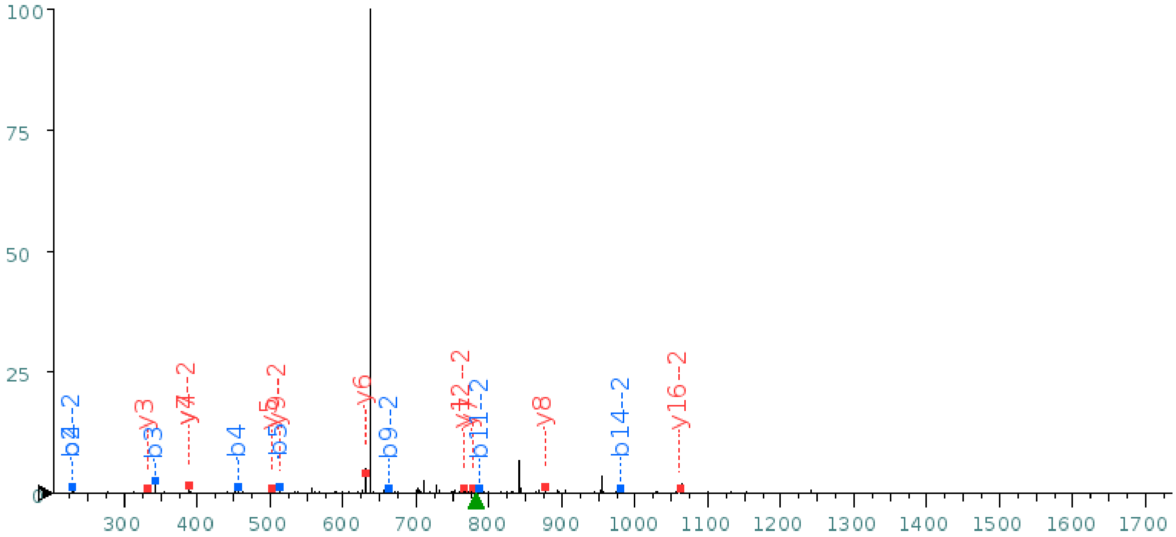

#### Predicted Fragmentation Pattern

+1

| Seq | # | b: Δ Error | b | y | y: Δ Error | +1 |
|---|---|---|---|---|---|---|
| N | 1 | --- | 115.050 | --- | --- | 18 |
| L | 2 | 716.137 | **228.134** | 2234.074 | --- | 17 |
| L | 3 | 722.373 | **341.218** | 2120.990 | --- | 16 |
| N | 4 | 224.448 | **455.261** | 2007.906 | --- | 15 |
| G | 5 | 412.766 | **512.283** | 1893.863 | --- | 14 |
| S# | 6 | --- | 819.373 | 1836.842 | --- | 13 |
| A | 7 | --- | 890.410 | 1529.752 | --- | 12 |
| K | 8 | --- | 1018.505 | 1458.714 | --- | 11 |
| S# | 9 | --- | 1325.595 | 1330.620 | --- | 10 |
| M* | 10 | --- | 1472.631 | 1023.529 | --- | 9 |
| V | 11 | --- | 1571.699 | **876.494** | 325.435 | 8 |
| F | 12 | --- | 1718.768 | **777.425** | 362.881 | 7 |
| Q | 13 | --- | 1846.826 | **630.357** | 855.862 | 6 |
| I | 14 | --- | 1959.910 | **502.298** | 426.865 | 5 |
| G | 15 | --- | 2016.932 | **389.214** | -38.617 | 4 |

| | | | | | | |
|---|---|---|---|---|---|---|
| A | 16 | --- | 2087.969 | **332.193** | 455.070 | 3 |
| N | 17 | --- | 2202.012 | 261.156 | --- | 2 |
| K | 18 | --- | --- | 147.113 | --- | 1 |

+2

| Seq | # | b: Δ Error | b | y | y: Δ Error | +1 |
|---|---|---|---|---|---|---|
| N | 1 | --- | 58.029 | --- | --- | 18 |
| L | 2 | --- | 114.571 | 1117.541 | --- | 17 |
| L | 3 | --- | 171.113 | **1060.999** | 343.503 | 16 |
| N | 4 | 716.137 | **228.134** | 1004.457 | --- | 15 |
| G | 5 | --- | 256.645 | 947.435 | --- | 14 |
| S# | 6 | --- | 410.190 | 918.925 | --- | 13 |
| A | 7 | --- | 445.709 | **765.379** | 796.689 | 12 |
| K | 8 | --- | 509.756 | 729.861 | --- | 11 |
| S# | 9 | -953.617 | **663.301** | 665.813 | --- | 10 |
| M* | 10 | --- | 736.819 | **512.268** | 441.050 | 9 |
| V | 11 | -1016.043 | **786.353** | 438.751 | --- | 8 |
| F | 12 | --- | 859.887 | **389.216** | -43.784 | 7 |
| Q | 13 | --- | 923.917 | 315.682 | --- | 6 |
| I | 14 | -165.989 | **980.459** | 251.653 | --- | 5 |
| G | 15 | --- | 1008.970 | 195.111 | --- | 4 |
| A | 16 | --- | 1044.488 | 166.600 | --- | 3 |
| N | 17 | --- | 1101.510 | 131.082 | --- | 2 |
| K | 18 | --- | --- | 74.060 | --- | 1 |

**Soleil Index**

**637.1848** **not explained** **100%**

| | | | | | |
|---|---|---|---|---|---|
| **Run:** | 79495 | **Scans:** | 19373 - 19373 | **m/z Range:** | 355.00281 to 1675.9608 |
| **Search:** | 79495 | **Precursor Mass:** | 1008.97842 | | |
| **Mods:** | C=160.03 *+15.99 #+220.06 @+291.1 | **Charge:** | +2 | | |

**Sequence**

DQVM*ELT#IAGM*GTSALK

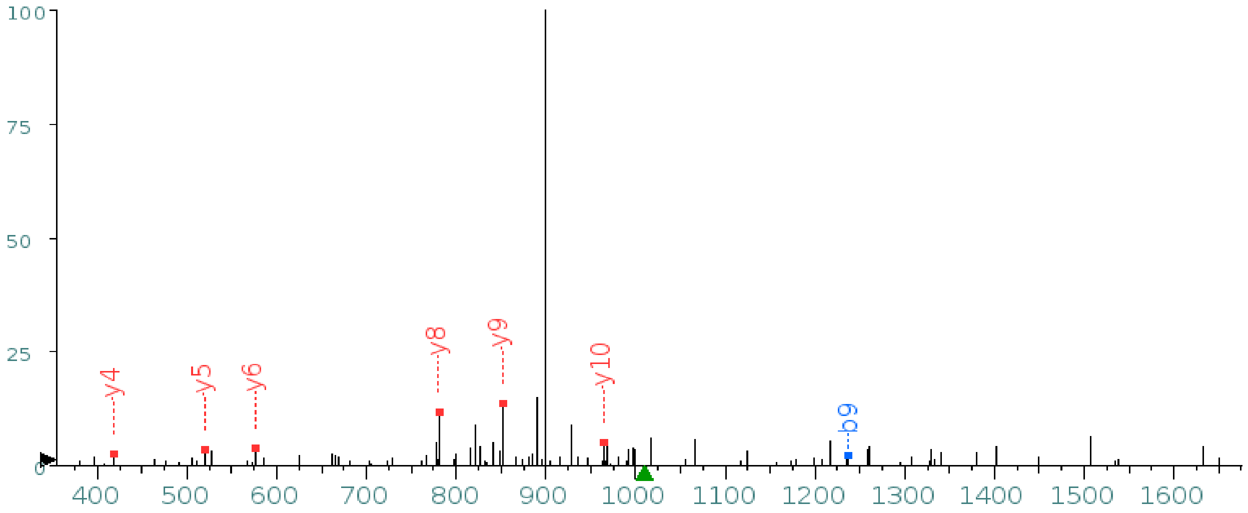

**Predicted Fragmentation Pattern**

| Seq | # | b: Δ Error | b | y | y: Δ Error | +1 |
|---|---|---|---|---|---|---|
| D | 1 | --- | 116.034 | --- | --- | 17 |
| Q | 2 | --- | 244.093 | 1901.908 | --- | 16 |
| V | 3 | --- | 343.161 | 1773.850 | --- | 15 |
| M* | 4 | --- | 490.197 | 1674.781 | --- | 14 |
| E | 5 | --- | 619.239 | 1527.746 | --- | 13 |
| L | 6 | --- | 732.323 | 1398.703 | --- | 12 |
| T# | 7 | --- | 1053.429 | 1285.619 | --- | 11 |
| I | 8 | --- | 1166.513 | **964.513** | **653.253** | 10 |
| A | 9 | **-556.441** | **1237.550** | **851.429** | **297.058** | 9 |
| G | 10 | --- | 1294.572 | **780.392** | **347.700** | 8 |
| M* | 11 | --- | 1441.607 | 723.371 | --- | 7 |
| G | 12 | --- | 1498.629 | **576.335** | **430.927** | 6 |
| T | 13 | --- | 1599.676 | **519.314** | **700.286** | 5 |
| S | 14 | --- | 1686.708 | **418.266** | **-119.062** | 4 |
| A | 15 | --- | 1757.746 | 331.234 | --- | 3 |
| L | 16 | --- | 1870.830 | 260.197 | --- | 2 |
| K | 17 | --- | --- | 147.113 | --- | 1 |

**Soleil Index**

| | | |
|---|---|---|
| **890.1161** | **not explained** | **15%** |
| **899.2133** | **not explained** | **100%** |

| | | | | | |
|---|---|---|---|---|---|
| **Run:** | 79495 | **Scans:** | 21368 - 21368 | **m/z Range:** | 306.31903 to 1983.5196 |
| **Search:** | 79495 | **Precursor Mass:** | 1036.83797 | | |
| **Mods:** | C=160.03 *+15.99 #+220.06 @+291.1 | **Charge:** | +3 | | |

**Sequence**

SM*VFQIGANKDQVM*ELT#IAGM*GTSALK

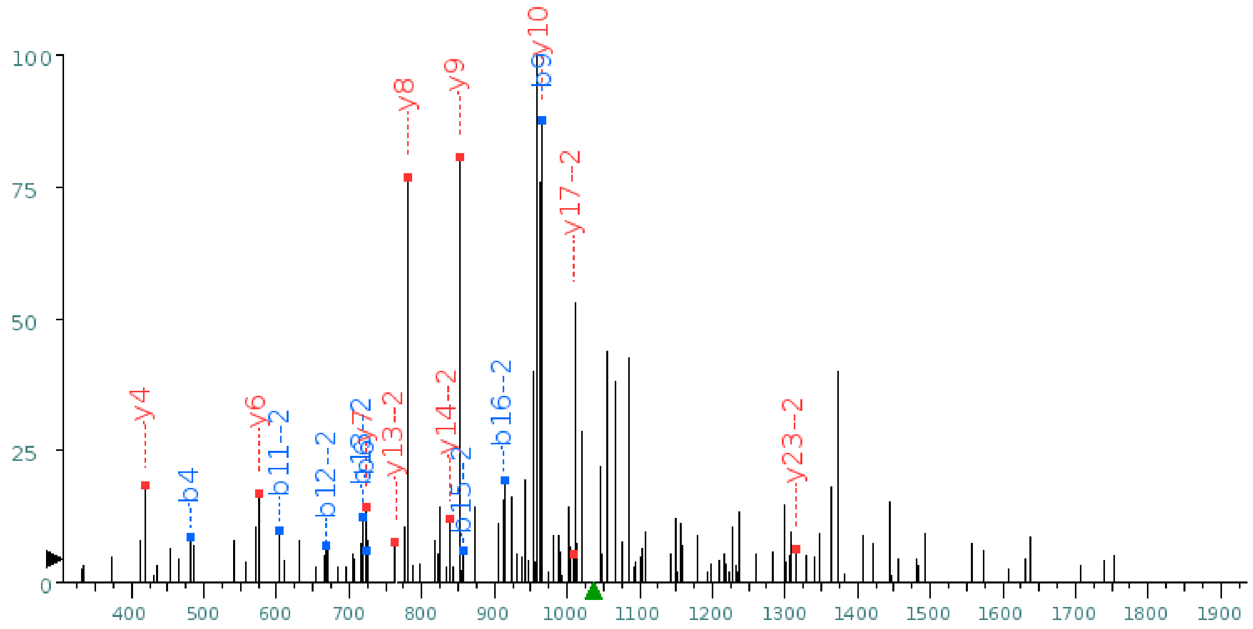

**Predicted Fragmentation Pattern**

| Seq | # | b: Δ Error | b +1 | y +1 | y: Δ Error | +1 |
|---|---|---|---|---|---|---|
| S | 1 | --- | 88.039 | --- | --- | 27 |
| M* | 2 | --- | 235.075 | 3021.446 | --- | 26 |
| V | 3 | --- | 334.143 | 2874.411 | --- | 25 |
| F | 4 | **472.530** | **481.212** | 2775.343 | --- | 24 |
| Q | 5 | --- | 609.270 | 2628.274 | --- | 23 |
| I | 6 | **115.848** | **722.354** | 2500.216 | --- | 22 |
| G | 7 | --- | 779.376 | 2387.132 | --- | 21 |
| A | 8 | --- | 850.413 | 2330.110 | --- | 20 |
| N | 9 | **-297.388** | **964.456** | 2259.073 | --- | 19 |
| K | 10 | --- | 1092.551 | 2145.030 | --- | 18 |
| D | 11 | --- | 1207.578 | 2016.935 | --- | 17 |
| Q | 12 | --- | 1335.636 | 1901.908 | --- | 16 |
| V | 13 | --- | 1434.705 | 1773.850 | --- | 15 |

| | | | | | | |
|---|---|---|---|---|---|---|
| M* | 14 | --- | 1581.740 | 1674.781 | --- | 14 |
| E | 15 | --- | 1710.783 | 1527.746 | --- | 13 |
| L | 16 | --- | 1823.867 | 1398.703 | --- | 12 |
| T# | 17 | --- | 2144.973 | 1285.619 | --- | 11 |
| I | 18 | --- | 2258.057 | 964.513 | -357.041 | 10 |
| A | 19 | --- | 2329.094 | 851.429 | 250.345 | 9 |
| G | 20 | --- | 2386.115 | 780.392 | 66.726 | 8 |
| M* | 21 | --- | 2533.151 | 723.371 | 418.572 | 7 |
| G | 22 | --- | 2590.172 | 576.335 | 252.074 | 6 |
| T | 23 | --- | 2691.220 | 519.314 | --- | 5 |
| S | 24 | --- | 2778.252 | 418.266 | 668.350 | 4 |
| A | 25 | --- | 2849.289 | 331.234 | --- | 3 |
| L | 26 | --- | 2962.373 | 260.197 | --- | 2 |
| K | 27 | --- | --- | 147.113 | --- | 1 |

+2

| Seq | # | b: Δ Error | b | y | y: Δ Error | +1 |
|---|---|---|---|---|---|---|
| S | 1 | --- | 44.523 | --- | --- | 27 |
| M* | 2 | --- | 118.041 | 1511.227 | --- | 26 |
| V | 3 | --- | 167.575 | 1437.709 | --- | 25 |
| F | 4 | --- | 241.109 | 1388.175 | --- | 24 |
| Q | 5 | --- | 305.139 | 1314.641 | 523.034 | 23 |
| I | 6 | --- | 361.681 | 1250.611 | --- | 22 |
| G | 7 | --- | 390.191 | 1194.069 | --- | 21 |
| A | 8 | --- | 425.710 | 1165.559 | --- | 20 |
| N | 9 | --- | 482.731 | 1130.040 | --- | 19 |
| K | 10 | --- | 546.779 | 1073.019 | --- | 18 |
| D | 11 | 340.750 | 604.292 | 1008.971 | -909.815 | 17 |
| Q | 12 | 539.716 | 668.322 | 951.458 | --- | 16 |
| V | 13 | 948.354 | 717.856 | 887.428 | --- | 15 |
| M* | 14 | --- | 791.374 | 837.894 | 791.951 | 14 |
| E | 15 | 659.168 | 855.895 | 764.377 | -1067.245 | 13 |
| L | 16 | 1035.956 | 912.437 | 699.855 | --- | 12 |
| T# | 17 | --- | 1072.990 | 643.313 | --- | 11 |
| I | 18 | --- | 1129.532 | 482.760 | --- | 10 |
| A | 19 | --- | 1165.051 | 426.218 | --- | 9 |
| G | 20 | --- | 1193.561 | 390.700 | --- | 8 |
| M* | 21 | --- | 1267.079 | 362.189 | --- | 7 |
| G | 22 | --- | 1295.590 | 288.671 | --- | 6 |
| T | 23 | --- | 1346.114 | 260.160 | --- | 5 |
| S | 24 | --- | 1389.630 | 209.637 | --- | 4 |
| A | 25 | --- | 1425.148 | 166.121 | --- | 3 |
| L | 26 | --- | 1481.690 | 130.602 | --- | 2 |
| K | 27 | --- | --- | 74.060 | --- | 1 |

**Soleil Index**

| | | |
|---|---|---|
| 781.4493 | not explained | 32% |
| 922.7178 | not explained | 16% |
| 943.2250 | not explained | 19% |
| 954.3511 | not explained | 40% |
| 957.6356 | not explained | 100% |
| 965.7803 | not explained | 23% |
| 1010.5305 | not explained | 53% |

| | | |
|---|---|---|
| 1011.2291 | not explained | 29% |
| 1019.7665 | not explained | 28% |
| 1020.3754 | not explained | 21% |
| 1046.7172 | not explained | 22% |
| 1055.7921 | b2-17 frag loss of water/ b2-17 frag loss of ammonia/ y2-18 frag loss of water/ y2-18 frag loss of ammonia | 44% |
| 1065.8207 | not explained | 38% |
| 1066.8579 | not explained | 26% |
| 1084.4264 | not explained | 42% |
| 1363.7609 | not explained | 18% |
| 1372.7654 | b2-24 frag loss of ammonia | 40% |
| 1445.2024 | not explained | 15% |

| | | | | | | |
|---|---|---|---|---|---|---|
| **Run:** | 79495 | **Scans:** | 21413 - 21413 | **m/z Range:** | 306.38196 | to 1660.9334 |
| **Search:** | 79495 | **Precursor Mass:** | 1000.98026 | | | |
| **Mods:** | C=160.03 *+15.99 #+220.06 @+291.1 | **Charge:** | +2 | | | |

**Sequence**

DQVM*ELTIAGMGT#SALK

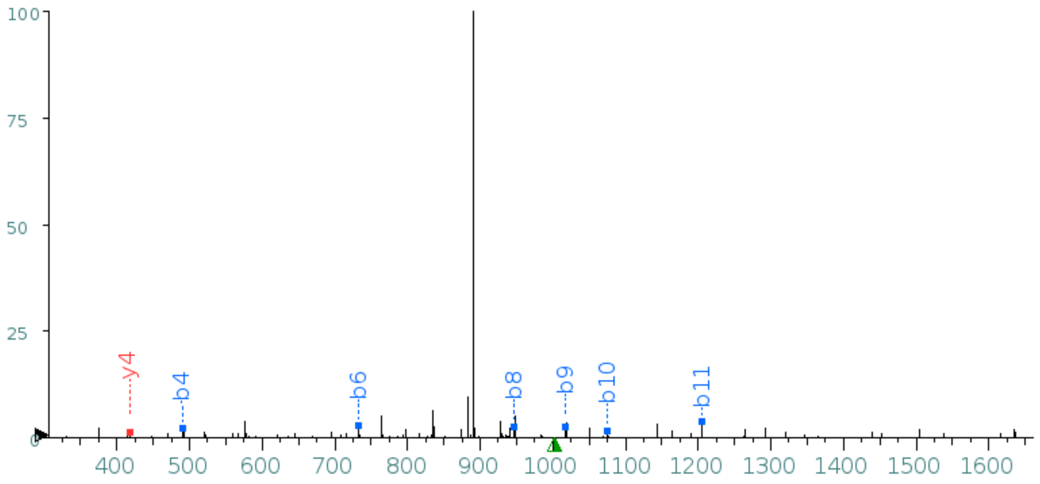

**Predicted Fragmentation Pattern**

| Seq | # | b: Δ Error | b | y | y: Δ Error | +1 |
|---|---|---|---|---|---|---|
| D | 1 | --- | 116.034 | --- | --- | 17 |
| Q | 2 | --- | 244.093 | 1885.913 | --- | 16 |
| V | 3 | --- | 343.161 | 1757.855 | --- | 15 |
| M* | 4 | 375.292 | **490.197** | 1658.786 | --- | 14 |
| E | 5 | --- | 619.239 | 1511.751 | --- | 13 |
| L | 6 | 193.455 | **732.323** | 1382.708 | --- | 12 |
| T | 7 | --- | 833.371 | 1269.624 | --- | 11 |
| I | 8 | 181.510 | **946.455** | 1168.577 | --- | 10 |
| A | 9 | 373.276 | **1017.492** | 1055.493 | --- | 9 |
| G | 10 | 253.922 | **1074.514** | 984.455 | --- | 8 |
| M | 11 | 254.803 | **1205.554** | 927.434 | --- | 7 |
| G | 12 | --- | 1262.576 | 796.393 | --- | 6 |
| T# | 13 | --- | 1583.682 | 739.372 | --- | 5 |
| S | 14 | --- | 1670.714 | **418.266** | 71.159 | 4 |
| A | 15 | --- | 1741.751 | 331.234 | --- | 3 |
| L | 16 | --- | 1854.835 | 260.197 | --- | 2 |
| K | 17 | --- | --- | 147.113 | --- | 1 |

**Soleil Index**

| | | |
|---|---|---|
| 891.3772 | not explained | 100% |

**MS/MS data of GluC digested peptides with site score A and site score B above 19 from *Clostridium botulinum Cb*Fla flagellin protein co-expressed with *Clostridium botulinum* Maf in BL21(DE3) cells. Results of dataset 3 (MS/MS results obtained in July 2019).**

| | | | |
|---|---|---|---|
| **Run:** | 79496 | **Scans:** | 38326 - 38326 |
| **Search:** | 79496 | **Precursor Mass:** | 1129.57212 |
| **Mods:** | C=160.03 *+15.99 #+220.06 @+291.1 | **Charge:** | +4 |

**m/z Range:** 317.57074 to 1992.1889

**Sequence**

LTIAGM*GTSALKVGSINIGSGKGAKSIS#GM*IETIDNAINTVSGE

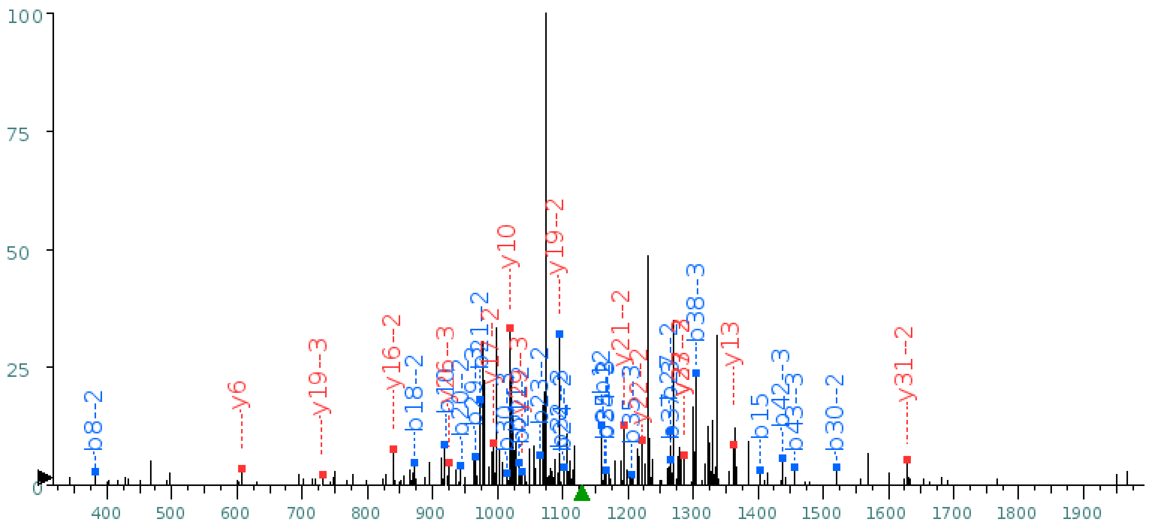

**Predicted Fragmentation Pattern**

+1

| Seq | # | b: Δ Error | b | y | y: Δ Error | +1 |
|---|---|---|---|---|---|---|
| L | 1 | --- | 114.091 | --- | --- | 44 |
| T | 2 | --- | 215.139 | 4402.185 | --- | 43 |
| I | 3 | --- | 328.223 | 4301.137 | --- | 42 |
| A | 4 | --- | 399.260 | 4188.053 | --- | 41 |
| G | 5 | --- | 456.282 | 4117.016 | --- | 40 |
| M* | 6 | --- | 603.317 | 4059.995 | --- | 39 |
| G | 7 | --- | 660.339 | 3912.959 | --- | 38 |
| T | 8 | --- | 761.386 | 3855.938 | --- | 37 |
| S | 9 | --- | 848.418 | 3754.890 | --- | 36 |
| A | 10 | -753.361 | **919.455** | 3667.858 | --- | 35 |
| L | 11 | -568.011 | **1032.539** | 3596.821 | --- | 34 |
| K | 12 | -631.216 | **1160.634** | 3483.737 | --- | 33 |
| V | 13 | --- | 1259.703 | 3355.642 | --- | 32 |

| | | | | | | |
|---|---|---|---|---|---|---|
| G | 14 | --- | 1316.724 | 3256.574 | --- | 31 |
| S | 15 | 43.442 | **1403.756** | 3199.552 | --- | 30 |
| I | 16 | --- | 1516.840 | 3112.520 | --- | 29 |
| N | 17 | --- | 1630.883 | 2999.436 | --- | 28 |
| I | 18 | --- | 1743.967 | 2885.393 | --- | 27 |
| G | 19 | --- | 1800.989 | 2772.309 | --- | 26 |
| S | 20 | --- | 1888.021 | 2715.288 | --- | 25 |
| G | 21 | --- | 1945.042 | 2628.256 | --- | 24 |
| K | 22 | --- | 2073.137 | 2571.234 | --- | 23 |
| G | 23 | --- | 2130.159 | 2443.139 | --- | 22 |
| A | 24 | --- | 2201.196 | 2386.118 | --- | 21 |
| K | 25 | --- | 2329.291 | 2315.081 | --- | 20 |
| S | 26 | --- | 2416.323 | 2186.986 | --- | 19 |
| I | 27 | --- | 2529.407 | 2099.954 | --- | 18 |
| S# | 28 | --- | 2836.497 | 1986.870 | --- | 17 |
| G | 29 | --- | 2893.519 | 1679.779 | --- | 16 |
| M* | 30 | --- | 3040.554 | 1622.758 | --- | 15 |
| I | 31 | --- | 3153.638 | 1475.722 | --- | 14 |
| E | 32 | --- | 3282.681 | **1362.638** | -409.584 | 13 |
| T | 33 | --- | 3383.728 | 1233.596 | --- | 12 |
| I | 34 | --- | 3496.812 | 1132.548 | --- | 11 |
| D | 35 | --- | 3611.839 | **1019.464** | 210.213 | 10 |
| N | 36 | --- | 3725.882 | 904.437 | --- | 9 |
| A | 37 | --- | 3796.919 | 790.394 | --- | 8 |
| I | 38 | --- | 3910.004 | 719.357 | --- | 7 |
| N | 39 | --- | 4024.046 | **606.273** | 475.356 | 6 |
| T | 40 | --- | 4125.094 | 492.230 | --- | 5 |
| V | 41 | --- | 4224.163 | 391.182 | --- | 4 |
| S | 42 | --- | 4311.195 | 292.114 | --- | 3 |
| G | 43 | --- | 4368.216 | 205.082 | --- | 2 |
| E | 44 | --- | --- | 148.060 | --- | 1 |

+2

| Seq | # | b: Δ Error | b | y | y: Δ Error | +1 |
|---|---|---|---|---|---|---|
| L | 1 | --- | 57.549 | --- | --- | 44 |
| T | 2 | --- | 108.073 | 2201.596 | --- | 43 |
| I | 3 | --- | 164.615 | 2151.072 | --- | 42 |
| A | 4 | --- | 200.134 | 2094.530 | --- | 41 |
| G | 5 | --- | 228.644 | 2059.012 | --- | 40 |
| M* | 6 | --- | 302.162 | 2030.501 | --- | 39 |
| G | 7 | --- | 330.673 | 1956.983 | --- | 38 |
| T | 8 | 799.394 | **381.197** | 1928.473 | --- | 37 |
| S | 9 | --- | 424.713 | 1877.949 | --- | 36 |
| A | 10 | --- | 460.231 | 1834.433 | --- | 35 |
| L | 11 | --- | 516.773 | 1798.914 | --- | 34 |
| K | 12 | --- | 580.821 | 1742.372 | --- | 33 |
| V | 13 | --- | 630.355 | 1678.325 | --- | 32 |
| G | 14 | --- | 658.866 | **1628.790** | 613.230 | 31 |
| S | 15 | --- | 702.382 | 1600.280 | --- | 30 |
| I | 16 | --- | 758.924 | 1556.764 | --- | 29 |
| N | 17 | --- | 815.945 | 1500.222 | --- | 28 |
| I | 18 | 370.347 | **872.487** | 1443.200 | --- | 27 |
| G | 19 | --- | 900.998 | 1386.658 | --- | 26 |
| S | 20 | -453.727 | **944.514** | 1358.147 | --- | 25 |
| G | 21 | 548.936 | **973.025** | 1314.631 | --- | 24 |
| K | 22 | 711.856 | **1037.072** | **1286.121** | -341.237 | 23 |
| G | 23 | 506.900 | **1065.583** | **1222.073** | -750.525 | 22 |
| A | 24 | 756.735 | **1101.102** | **1193.563** | -1.138 | 21 |

| | | | | | | |
|---|---|---|---|---|---|---|
| K | 25 | 458.999 | 1165.149 | 1158.044 | --- | 20 |
| S | 26 | --- | 1208.665 | 1093.996 | 63.136 | 19 |
| I | 27 | -661.187 | 1265.207 | 1050.480 | --- | 18 |
| S# | 28 | --- | 1418.752 | 993.938 | -264.444 | 17 |
| G | 29 | --- | 1447.263 | 840.393 | 123.212 | 16 |
| M* | 30 | -630.928 | 1520.781 | 811.883 | --- | 15 |
| I | 31 | --- | 1577.323 | 738.365 | --- | 14 |
| E | 32 | --- | 1641.844 | 681.823 | --- | 13 |
| T | 33 | --- | 1692.368 | 617.302 | --- | 12 |
| I | 34 | --- | 1748.910 | 566.778 | --- | 11 |
| D | 35 | --- | 1806.423 | 510.236 | --- | 10 |
| N | 36 | --- | 1863.445 | 452.722 | --- | 9 |
| A | 37 | --- | 1898.963 | 395.701 | --- | 8 |
| I | 38 | --- | 1955.505 | 360.182 | --- | 7 |
| N | 39 | --- | 2012.527 | 303.640 | --- | 6 |
| T | 40 | --- | 2063.051 | 246.619 | --- | 5 |
| V | 41 | --- | 2112.585 | 196.095 | --- | 4 |
| S | 42 | --- | 2156.101 | 146.561 | --- | 3 |
| G | 43 | --- | 2184.612 | 103.045 | --- | 2 |
| E | 44 | --- | --- | 74.534 | --- | 1 |

**+3**

| Seq | # | b: Δ Error | b | y | y: Δ Error | +1 |
|---|---|---|---|---|---|---|
| L | 1 | --- | 38.702 | --- | --- | 44 |
| T | 2 | --- | 72.385 | 1468.067 | --- | 43 |
| I | 3 | --- | 110.079 | 1434.384 | --- | 42 |
| A | 4 | --- | 133.758 | 1396.689 | --- | 41 |
| G | 5 | --- | 152.765 | 1373.010 | --- | 40 |
| M* | 6 | --- | 201.777 | 1354.003 | --- | 39 |
| G | 7 | --- | 220.784 | 1304.991 | --- | 38 |
| T | 8 | --- | 254.467 | 1285.984 | -235.019 | 37 |
| S | 9 | --- | 283.478 | 1252.302 | --- | 36 |
| A | 10 | --- | 307.157 | 1223.291 | --- | 35 |
| L | 11 | --- | 344.851 | 1199.612 | --- | 34 |
| K | 12 | --- | 387.550 | 1161.917 | --- | 33 |
| V | 13 | --- | 420.572 | 1119.219 | --- | 32 |
| G | 14 | --- | 439.580 | 1086.196 | --- | 31 |
| S | 15 | --- | 468.590 | 1067.189 | --- | 30 |
| I | 16 | --- | 506.285 | 1038.178 | -353.834 | 29 |
| N | 17 | --- | 544.299 | 1000.484 | --- | 28 |
| I | 18 | --- | 581.994 | 962.469 | --- | 27 |
| G | 19 | --- | 601.001 | 924.775 | -751.145 | 26 |
| S | 20 | --- | 630.012 | 905.767 | --- | 25 |
| G | 21 | --- | 649.019 | 876.757 | --- | 24 |
| K | 22 | --- | 691.717 | 857.750 | --- | 23 |
| G | 23 | --- | 710.724 | 815.051 | --- | 22 |
| A | 24 | --- | 734.403 | 796.044 | --- | 21 |
| K | 25 | --- | 777.102 | 772.365 | --- | 20 |
| S | 26 | --- | 806.112 | 729.667 | 1313.393 | 19 |
| I | 27 | --- | 843.807 | 700.656 | --- | 18 |
| S# | 28 | --- | 946.171 | 662.961 | --- | 17 |
| G | 29 | 823.041 | 965.178 | 560.598 | --- | 16 |
| M* | 30 | 838.856 | 1014.190 | 541.591 | --- | 15 |
| I | 31 | --- | 1051.884 | 492.579 | --- | 14 |
| E | 32 | -761.267 | 1094.898 | 454.884 | --- | 13 |
| T | 33 | --- | 1128.581 | 411.870 | --- | 12 |
| I | 34 | -507.508 | 1166.276 | 378.188 | --- | 11 |
| D | 35 | 423.445 | 1204.618 | 340.493 | --- | 10 |

| | | | | | | |
|---|---|---|---|---|---|---|
| N | 36 | --- | 1242.632 | 302.151 | --- | 9 |
| A | 37 | -782.544 | **1266.311** | 264.136 | --- | 8 |
| I | 38 | -410.034 | **1304.006** | 240.457 | --- | 7 |
| N | 39 | --- | 1342.020 | 202.762 | --- | 6 |
| T | 40 | --- | 1375.703 | 164.748 | --- | 5 |
| V | 41 | --- | 1408.726 | 131.066 | --- | 4 |
| S | 42 | -638.497 | **1437.736** | 98.043 | --- | 3 |
| G | 43 | -661.257 | **1456.744** | 69.032 | --- | 2 |
| E | 44 | --- | --- | 50.025 | --- | 1 |

**Soleil Index**

| | | |
|---|---|---|
| 977.6372 | b2-38 frag doubly charged/ y2-17 frag loss of ammonia/ y2-38 frag loss of water | 30% |
| 978.9908 | y2-38 frag doubly charged | 22% |
| 997.8849 | b3-30 frag loss of ammonia | 33% |
| 1021.8340 | y3-29 frag loss of ammonia | 24% |
| 1070.6366 | not explained | 17% |
| 1071.4857 | not explained | 20% |
| 1075.2932 | y2-19 frag loss of water/ y2-42 frag loss of water | 100% |
| 1231.4331 | not explained | 49% |
| 1269.4232 | y2-23 frag loss of ammonia/ y3-37 frag loss of ammonia | 35% |
| 1300.4755 | b1-14 frag loss of ammonia | 17% |
| 1335.9961 | y3-39 frag loss of water/ y3-39 frag loss of ammonia | 32% |
| 1336.8206 | y3-39 frag loss of water/ y3-39 frag loss of ammonia | 17% |
